# *simona:* a comprehensive R package for semantic similarity analysis on bio-ontologies

**DOI:** 10.1101/2023.12.03.569758

**Authors:** Zuguang Gu

## Abstract

**Background:** Bio-ontologies are keys in structuring complex biological information for effective data integration and knowledge representation. Semantic similarity analysis on bio-ontologies quantitatively assesses the degree of similarity between biological concepts based on the semantics encoded in ontologies. It plays an important role in structured and meaningful interpretations and integration of complex data from multiple biological domains.

**Results:** We present *simona*, a novel R package for semantic similarity analysis on general bioontologies. *Simona* implements infrastructures for ontology analysis by offering efficient data structures, fast ontology traversal methods, and elegant visualizations. Moreover, it provides a robust toolbox supporting over 70 methods for semantic similarity analysis. With *simona*, we conducted a benchmark against current semantic similarity methods. The results demonstrate methods are clustered based on their mathematical methodologies, thus guiding researchers in the selection of appropriate methods. Additionally, we explored annotation-based versus topology-based methods, revealing that semantic similarities solely based on ontology topology can efficiently reveal semantic similarity structures, facilitating analysis on less-studied organisms and other ontologies.

**Conclusions:** *Simona* offers a versatile interface and efficient implementation for processing, visualization, and semantic similarity analysis on bio-ontologies. We believe that *simona* will serve as a robust tool for uncovering relationships and enhancing the interoperability of biological knowledge systems.

## Background

Bio-ontologies play a crucial role in organizing and standardizing biological information to facilitate effective data integration and knowledge representation. They are typically constructed as directed acyclic graphs (DAGs) where biological concepts are modelled as terms connected by hierarchical relations. The most widely used bio-ontology, Gene Ontology (GO), provides a controlled vocabulary to describe functions, processes and cellular components of genes, playing a central role in functional interpretation in biology studies. Besides GO, there are a huge amount of bio-ontology resources developed in various biological domains, such as Medical Subject Headings (MeSH), Cell Ontology [1] and the Human Disease Ontology [2]. Additionally, there are bio-ontology databases that allow querying hierarchical relations of biology terms, such as the OBO Foundry [3], BioPortal [4] and Ontobee [5]. These resources are essential for data integration, enabling researchers to combine information from diverse sources and analyze it in a coherent and standardized way.

Semantic similarity analysis on bio-ontologies is a computational approach that quantitatively assesses the degree of similarity between biological concepts based on the semantics encoded in ontologies. It has wide applications in the biology field, such as gene function prediction [6], clustering and summarization of biological entities [7], interpretation of proteinprotein interactions [8], cross-species comparisons [9], and biomedical text mining [10]. As biomedical data continues to increase dramatically in complexity and size, semantic similarity analysis is becoming an important tool for structured and meaningful interpretations and integration of complex data from multiple biological domains.

Semantic similarity analysis is a well-established topic in general ontology studies. Over the last three decades, numerous semantic similarity methods have been proposed. The development of these methods began with the analysis of WordNet [11], a collection of English words organized in a hierarchical structure. As the GO project developed and matured, methods were adapted to apply to GO, and new methods specifically for GO were further developed. A comprehensive overview of state-of-the-art methods for semantic similarity analysis was provided by Mazandu *et al*. [12]. However, to date, only a few methods have been implemented as tools in the biology field, and most of them are only designed for GO. In the R programming ecosystem, there are currently only three packages for semantic similarity analysis: *ontologySimilarity* [13], *GOSemSim* [14] and *GOSim* [15]. These packages are either restricted to GO or have very limited functionalities. Therefore, the current R package development for semantic similarity analysis does not fully leverage the theoretical efforts in this field, limiting practical applications in general bio-ontology analysis and integration to the R/Bioconductor ecosystem. There are also tools implemented in other programming languages such as *SML-Toolkit* [16] in Java and *FastSemSim* [17] in Python, but we focus on the R computing environment in this work.

In this paper, we introduced a new R package named *simona*, designed to be a general-purpose package for semantic similarity analysis on bio-ontologies. *Simona* makes two major contributions: 1) the establishment of an infrastructure for ontology analysis, including an efficient data structure for storing ontology data, fast methods to traverse ontologies, and elegant visualizations; 2) providing a comprehensive toolbox for semantic similarity analysis with more than 70 different methods. *Simona* is implemented by efficient algorithms. Using GO as the test ontology, *simona* has runtime improvement of approximately 2.6x, 31x, and >3000x compared to *ontologySimilarity, GOSemSim*, and *GOSim*, respectively.

Despite the comprehensive set of semantic similarity methods proposed in the field, there is a lack of quantitative comparisons of methods based on real-world datasets in current studies. We performed comparisons of semantic similarity methods, leveraging the comprehensive toolbox provided by *simona*. We grouped methods into clusters, which are reflected by the commonness of their mathematical methodologies, providing users guidance to select a proper one for their studies. Most existing tools for semantic similarity analysis rely on external gene annotations and are majorly restricted to GO, limiting their applicability to organisms with incomplete gene annotation or other ontologies. We compared the annotation-based method and the method solely based on the topology of the ontology without relying on external data. The results demonstrate that the two types of methods generate very similar similarity patterns. This analysis provides evidence for extending the similarity analysis to broader ontologies where gene annotations may not be available or may be incomplete.

## Methods

Mazandu *et al*. [12] provide an excellent and comprehensive overview of current methods for semantic similarity analysis on GO. Nevertheless, these methods can be applied to general ontologies without modification. In *simona*, we have implemented the methods reviewed by Mazandu *et al*. [12] and one additional semantic similarity method proposed by Zhao and Wang [18]. Furthermore, we introduced several new methods in this section with simplified forms, primarily for comparing with other complex methods to reveal their mathematical attributes. In addition to the standard settings introduced in Mazandu *et al*. [12] and original papers, *simona* supports more flexible configurations. One typical scenario is for all distance-based methods, *simona* supports not only the shortest distance-based methods as introduced in the original papers but also the longest distance-based methods. This is because distance is typically a measure of the semantic specificity that a term has or the distinctiveness that two terms exhibit in the ontology.

The semantic similarity analysis can be applied on the following three levels:

- *Single-term level*. At the level of individual terms, a numeric score is assigned to each term in the ontology, measuring its semantic specificity. This measure is often referred to information content (IC). Generally, IC is higher for terms deeper in the DAG, indicating a greater level of semantic information. Many similarity methods are established based on various IC definitions.
- *Term-to-term level*. A similarity is calculated for a pair of terms. This score reflects how closely related two terms are in the ontology, determined by the specificity of their common ancestors or their proximity in the DAG.
- *Group-to-group level*. Semantic similarity is also calculated for two groups of terms. This calculation aggregates similarities from individual terms within the two groups. The resulting score provides a measure of overall similarity between two groups.

In the next three subsections, we only provide brief introductions to the typical methods on the three levels. Readers are encouraged to refer to Mazandu *et al*. [12] or the vignettes of *simona* for the comprehensive definitions of these methods. We adopt similar mathematical notations as presented in Mazandu *et al*. [12], available in its supplementary file.

In total, *simona* provides 11 methods for IC, 34 methods for term-to-term similarity, and 28 methods for group-to-group similarity. Note that for many similarity methods, users have the flexibility to choose a specific IC method. Additionally, for a group similarity method, users can select both a specific IC method and a term-to-term similarity method. Thus, *simona* provides extensive support for a wide range of methods in semantic similarity analysis.

### Information content

IC is a general concept that quantifies the amount of information a term can provide. In information theory, IC is defined as the negative logarithm of the probability of a specific occurring event [19]. Formally, for term *x*,

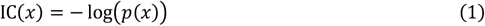

where *p*(*x*) is the probability of observing term *x* if taken as a random variable. In this way, when a term is associated with a small probability, indicating that the corresponding event is rare to occur, it is called informative if it is observed in a dataset.

There are a large number of methods that define *p*(*x*), categorized as either *extrinsic*, relying on data from external sources, or *intrinsic*, which is solely dependent on the ontology itself. In many applications of semantic analysis on GO, *p*(*x*) is often derived from external gene annotations to GO terms [20], which is defined as

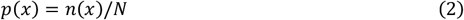

where *n*(*x*) is the number of unique genes annotated to GO term *x*, and *N* is the total number of genes annotated to the whole GO (or one of its namespaces). Due to the hierarchical structure of GO, when a gene is annotated to a specific term, it is also implicitly annotated to all its parent and ancestor terms. Thus, *n*(*x*) also includes genes that are indirectly annotated to the offspring terms of *x*. Then, *p*(*x*) is the probability of a term being annotated if randomly selecting genes from *N*. It is easy to see, if a term is annotated to fewer genes, indicating a more specific biological concept, it becomes more informative for downstream biological meaning interpretation.

Majority uses of ICs in current biological data analysis rely on the external gene annotation, however, IC can also be defined in an intrinsic way, which only relies on the ontology’s topology. Here as examples, we proposed two new and simple forms of the intrinsic definition of *p*(*x*). The first one is defined as the fraction of the number of *x*’s offspring terms to the total number of terms in the ontology

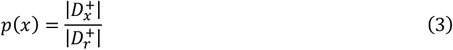

Where 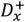 is the set of offspring terms of *x*, including *x* itself, 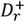 is the set of offspring terms of the root term *r*, including *r* itself (assuming there is only one root term for simplicity), and the notation | · | is the number of elements in a set. This simple definition of *p*(*x*) measures the broadness of a term connects to the rest of the terms in the ontology (we name this method as *IC_offspring* in *simona*).

The second intrinsic *p*(*x*) measures the relative distance to leaf terms in the ontology, defined as:

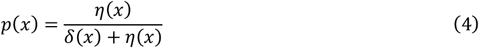

where *η*(*x*) is the height of *x* defined *as* the longest distance from *x* to its connectable leaves, and *δ*(*x*) is the depth of *x* defined as the longest distance from the root. Then in the longest path from the root to the leaf passing through *x, p*(*x*) is the proportion of the length from *x* to the leaf (we name this method *IC_height* in *simona*).

Last, not necessarily restricted to probability-based forms, IC can be generalized as a numeric score, satisfying the only requirement:

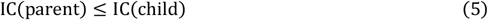

which means a parent term should not be more specific than its child term.

### Term-to-term similarity

The calculation of semantic similarity between two terms relies on their topological relations within the ontology. Simply speaking, if the common ancestors of two terms represent highly specific biological concepts, or if the two terms are closely located in the DAG, it can be concluded that the two terms are similar in contributing similar biological meanings. The evaluation of semantic similarity always involves common ancestors (CAs) shared by the two terms, denoted as:

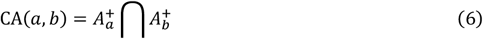

Where 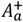 is the set of term *a*’s ancestors, including *a* itself. The notation is the same for term *b*.

A simple definition of the semantic similarity proposed and implemented in *simona* is:

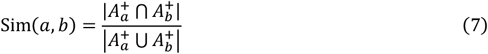

which is a Jaccard-like measurement. Terms *a* and *b* have high similarity if they have a similar inheritance from the upstream of the ontology. Despite its simplicity, we demonstrated in the Results section that this method performs similarly to more sophisticated methods (we name this method as *Sim_Ancestor* in *simona*).

Many similarity methods proposed in current studies make use of a specific type of common ancestors. There are the following three different types of common ancestors.

- *Most informative common ancestor (MICA)*. The common ancestor term with the highest IC is named MICA. IC can be calculated extrinsically or intrinsically.

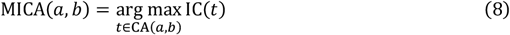
- *Lowest common ancestor (LCA)*. The common ancestor term with the largest depth (*i*.*e*., the lowest in the DAG) is named LCA.

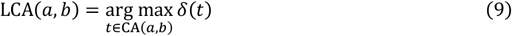
*Nearest common ancestor (NCA)*. Some methods depend on the shortest distance between terms *a* and *b*. The common ancestor of *a* and *b* on the shortest path is named NCA. In Equation 10, *D*_sp_(*t, a*) is the shortest distance from term *t* to *a*, and the same for *D*_sp_(*t, b*).

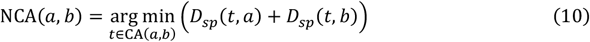

LCA and NCA are defined based on the topology of the DAG. In the DAG, a child term can have more than one parent, thus LCA is possibly a different term from NCA for two terms *a* and *b*.

The next two semantic similarity methods are widely used in ontology analysis. Lin [21] defined the similarity as the IC of the MICA term *c*, normalized by the average ICs of terms *a* and *b*.

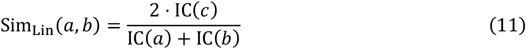

Although originally proposed for general information theory, it has been adopted and widely used for semantic similarities analysis on GO terms, taking ICs calculated from gene annotations.

As a second typical example, Wu and Palmer [22] proposed a similarity method that is purely based on the ontology structure, defined as:

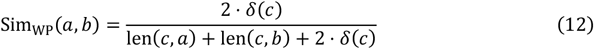

where term *c* is the LCA of terms *a* and *b*, len(*c, a*) is the longest distance from *c* to *a*, and *δ*(*c*) is the depth of *c* which is the longest distance from root *r* to *c*. Note *δ*(*c*) can be written as len(*r,c*) by definition, then the WP method can be rewritten in the Lin-like form:

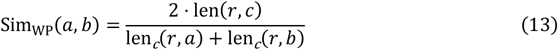

Where len_c_(*r, a*) is the longest distance from *r* to *a*, passing through term *c*.

In the Results section, we used the Lin and the WP methods as two representative methods for comparisons.

### Group-to-group similarity

It calculates a single similarity value between two groups of terms, by aggregating similarities from individual term pairs within the two groups. One typical application is in calculating the semantic or functional similarity of two genes or proteins, where the similarity is calculated based on the two sets of GO terms to which genes are annotated.

In general, there are two types of methods for calculating groupwise similarities. The first type makes use of the pairwise similarities of terms in the two groups. Denote the two groups of terms as *T*_*p*_ and *T*_*q*_, in the “best-match average” (BMA) method [23], the semantic similarity of a single term *x* to a group of terms *T* is first calculated as:

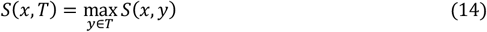

which is the similarity between *x* and its “best matched”, *i*.*e*., the most similar, term in *T*. Based on it, the semantic similarity between group *p* and *q* can be calculated, for example, as the average of similarities between terms in one group and their best-matched terms in the other group.

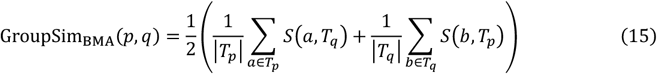

The second type of method treats the two groups of terms as two sub-graphs induced from the complete ontology DAG. For example, the following method named ALN calculates the average distance between the two sub-graphs as the similarity measure [24].

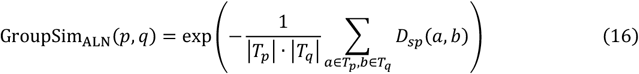

*Simona* supports a large number of groupwise similarity methods. However, due to the heterogeneity of similarity patterns within the groups of terms, the similarity value on the group level may vary significantly among different methods. Thus, in the Results section, we will not perform benchmarking on groupwise similarity methods.

### Implementation

#### The package

*Simona* is a general-purpose package for semantic similarity analysis on bio-ontologies. It employs a standardized data structure in the *ontology_DAG* class and implements efficient functions for traversing DAGs using depth-first search (DFS) or breadth-first search (BFS) using low-level C++ code. *Simona* can efficiently handle large ontologies containing millions of terms in a reasonable time.

It is also useful to examine local structures of DAGs, only focusing on a subset of terms of interest. For example, it can be useful to study how information is transmitted from ancestors to a specific set of offspring terms. *Simona* provides a flexible way to filter the complete DAG. Specifically, to obtain a sub-DAG induced from a single term as the root or a set of terms as leaves, the subsetting method on the complete `dag` object is especially convenient, as demonstrated in the following code.

`dag[term] # a sub-DAG that only contains ‘term’ and its offspring`

`dag[, terms] # a sub-DAG that only contains ‘terms’ and their ancestors`

In the semantic similarity analysis, for two terms, `term1` and `term2`, the following code can be used to extract a sub-DAG containing their common ancestors.

`ancestors1 = dag_ancestors(dag, term1, include_self = TRUE)`

`ancestors2 = dag_ancestors(dag, term2, include_self = TRUE)`

`sub_dag = dag_filter(dag, terms = intersect(ancestors2, ancestors2))`

There is a helper function `CA_terms()` which directly extracts CAs of two terms. The previous code can be simplified to:

`sub_dag = dag_filter(dag, terms = CA_terms(term1, term2))`

To take the union of ancestors of both terms, we can directly use the subsetting method.

`dag[, c(term1, term2)]`

*Simona* offers a comprehensive set of low-level functions designed for term pairs. These functions include calculations for distances considering DAGs in both directed and undirected modes, supporting both shortest-distance and longest-distance measurements. Additionally, there are functions for identifying MICA/LCA/NCA terms. These low-level functions serve as the foundation for implementing more specialized methods for semantic similarity analysis in *simona*.

#### Ontology formats

*Simona* supports the common formats of ontologies, simplifying the analysis of biological ontologies from public databases. For the *obo* format, *simona* provides a parser function, `import_obo()`, which imports and parses hierarchical relations of terms along with their relation types. Other formats, such as *owl* (the RDF/XML format) and *ttl* (the Turtle format), are prioritized to be parsed by an external parser, *ROBOT* [25] where internally they are converted to the *obo* format by *ROBOT* and then read by `import_obo()`. While *ROBOT* serves as a comprehensive and elegant tool for converting between various ontology formats, it may not be memory-efficient for larger ontologies (*e*.*g*., > 200K terms). To address this concern, optionally, *simona* also implements native parsers for the *owl* and *ttl* formats with the `import_owl()` and `import_ttl()` functions. These functions can recognize commonly used tags in the files. However, they may not be universally applicable to all aspects of the two formats.

Since there is no standard data format for external annotations, *simona* accepts annotations as a simple R list object that contains vectors of items annotated to terms. It is not necessary whether items are only annotated to the lowest terms in the ontology or they have already been aggregated. *Simona* processes such aggregation automatically.

#### Treelization

DAG is a generalized form of a tree, allowing a child term to have more than one parent. This may cause difficulties when partitioning a DAG into sub-DAGs where a term is possibly associated to multiple sub-DAGs. *Simona* provides a function `dag_treelize()`, which simplifies a DAG to a tree to let each term strictly have one parent. The reduction is executed in a breadth-first manner. Beginning from the root and to a term *a*, a unique link connects *a* and one of its child term *c* only when *δ*_*c*_ = *δ*_*a*_ + 1, *i*.*e*., the depth of *c* equals the depth of *a* plus one. Once the child *c* is selected, the links connecting *c* and all its other parents are removed. The treelization process can also be described in another way. For a term *c* and the set of its parents {*a*_1_, …, *a*_*k*_}, only the link from the parent with the largest depth in the DAG is kept and all the links from other parents are removed. In this way, the depths of all terms in the reduced tree remain the same as in the original DAG. The resulting tree is also saved as an *ontology_DAG* object but explicitly marked as a tree. Treelization is mainly used for partitioning and visualization on DAG, which will be introduced in the next two subsections.

#### Partitioning

In an ontology, a biological concept may have multiple child concepts that describe it more specifically from distinct aspects. Partitioning the ontology helps achieve a more concise view of the system. In *simona*, ontology partitioning is applied to the treelized DAG, where the tree is cut at specific levels. There are two approaches: cutting the tree at a certain level (defined by the depth) using the `partition_by_level()` function, or dynamically cutting the tree by controlling the number of terms in sub-trees with the `partition_by_size()` function.

#### Visualization

Hierarchical layouts, such as radial layouts, are widely used for ontology visualization [26]. While these layouts are efficient for visualizing small ontologies, challenges arise when dealing with larger ontologies. *Simona* provides functions for visualizing both small and large ontologies.

The `dag_graphviz()` function is designed to visualize small DAGs using the Graphviz hierarchical layout internally through the *DiagrammeR* package [27]. Together with the subsetting functionality on the *ontology_DAG* object, it is a useful tool for studying local structures in DAG. For example, to visualize ancestors of two terms with their CAs highlighted in red:

`sub_dag = dag[, c(term1, term2)]`

`ca = CA_terms(dag, term1, term2)`

`color = rep(“red”, length(ca))`

`names(color) = ca`

`dag_graphviz(sub_dag, node_param = list(color = color))`

*Simona* also implements a radial layout for visualizing complete ontologies with the function `dag_circular_viz()`. The layout is first applied to the treelized DAG, later additional links that are removed from the treelized DAG are added back subsequently. In this layout, terms are positioned on circles with different radii. For a circle where terms with a depth of *d* are located, the radius is calculated as

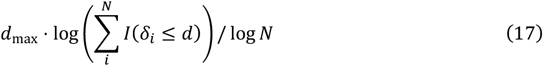

where *d*_max_ is the maximal depth, *N* is the total number of terms in the DAG, *δ*_*i*_ is the depth of term *i* and *I*() denotes the identity function. The radius is set to zero for the root term. This adjustment on the radius helps get rid of the scenario where a single long branch affects the global visualization.

Each term is associated with a sector on the circle. The width of the sector is proportional to the number of its connectable leaf terms in the treelized DAG. Importantly, since the layout is from the tree representation of the DAG, the sectors of child terms are mutually exclusive.

The sum of their widths equals the width of the sector of their parent term. In other words, the sector of a term entirely includes its offspring in the treelized DAG.

The `dag_circular_viz()` function provides a useful tool for visualizing the global structures of ontologies (more examples are shown in Fig. 5 and Supplementary File 6), thereby revealing specific attributes that may be of interest. Another practical use case is to highlight a list of terms on the DAG to observe their proximity and relationships. For example, the following code highlights a list of significant GO terms from a GO enrichment analysis (Fig. 1), where `sig_go_ids` is a vector containing significant GO terms.

`dag_circular_viz(dag, highlight = sig_go_ids, …)`

**Fig. 1.**
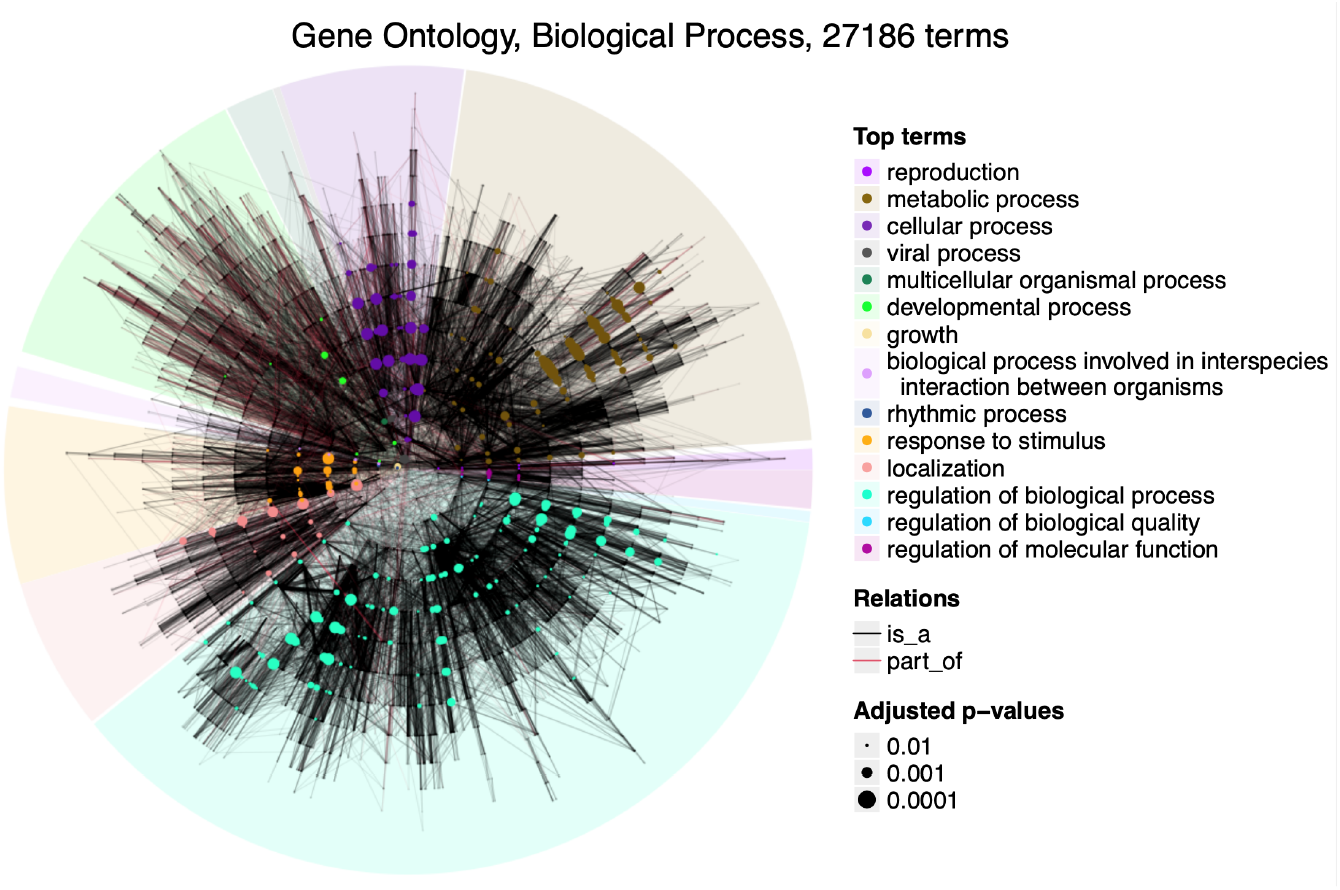
Circular visualization on GO, the Biological Process namespace. A list of significant GO terms from a GO enrichment analysis are highlighted on the plot. Dot sizes are proportional to - log10(adjusted *p*-values) from the enrichment analysis.

#### Semantic similarity analysis

*Simona* provides an extensive set of methods for semantic similarity analysis. The `term_IC()` function supports 11 methods for calculating ICs, `term_sim()` supports 34 methods for calculating term-to-term similarities, and `group_sim()`supports 28 methods for calculating group-to-group similarities. The core calculations are implemented through efficient algorithms in C++ code, and it has significant performance improvement over existing tools. A detailed description of the algorithm for traversing DAGs can be found in Supplementary File 2.

#### Interactive ontology browser

*Simona* implements a `dag_shiny()` function, which generates a web-based Shiny application. Users can visualize and compute semantic similarities for a provided list of terms. Additionally, the application facilitates interactive exploration of parent, sibling, and child terms, and allows users to query common ancestors in the ontology.

## Results and discussion

### Compared to other tools

We compared *simona* (version 1.13.2) to the following R packages supporting semantic similarity analysis: *ontologyIndex* (version 2.12)/*ontologySimilarity* (version 2.7) [13], *GOSemSim* (version 2.30.2) [14] and *GOSim* (1.42.0) [15]. *ontologyIndex* and *ontologySimilarity* are part of a suite of packages called *ontologyX*, where *ontologyIndex* offers methods for IC analysis and *ontologySimilarity* offers methods for semantic similarity analysis. *GOSemSim* and *GOSim* only support GO, so for this comparison, we took the Biological Process (BP) namespace in GO as the test ontology. *GOSemSim* and *GOSim* only support IC calculated based on gene annotation. While *ontologyIndex* supports more general annotation-based IC methods, and when working with GO, it mostly uses genes as the annotation items. Therefore, we only compared ICs and semantic similarities based on gene annotations with human as the selected organism. In the comparison, the GO data is from the *GO*.*db* package (version 3.19.1) or the *obo* file from GO website with the same version (version 2024-01-17, https://release.geneontology.org/2024-01-17/) as in *GO*.*db*. The relation types of “is a”, “part of”, “regulates”, “positively regulates” and “negatively regulates” are used. The package *org*.*Hs*.*eg*.*db* (version 3.19.1) is used as the source of gene annotation to GO terms. R 4.4.1 and Bioconductor 3.19 are used as the working environment. A detailed analysis report can be found in Supplementary File 1.

We first compared ICs calculated by the four tools. As shown in Figs 2A-C, *simona* and *ontologyIndex* generate identical IC values, while another two tools of *GOSemSim* and *GOSim* also generate identical IC values. Although there are systematic shifts in ICs between *simona*/*ontologyIndex* and *GOSemSim*/*GOSim*, they show strong linear relationships.

**Fig. 2.**
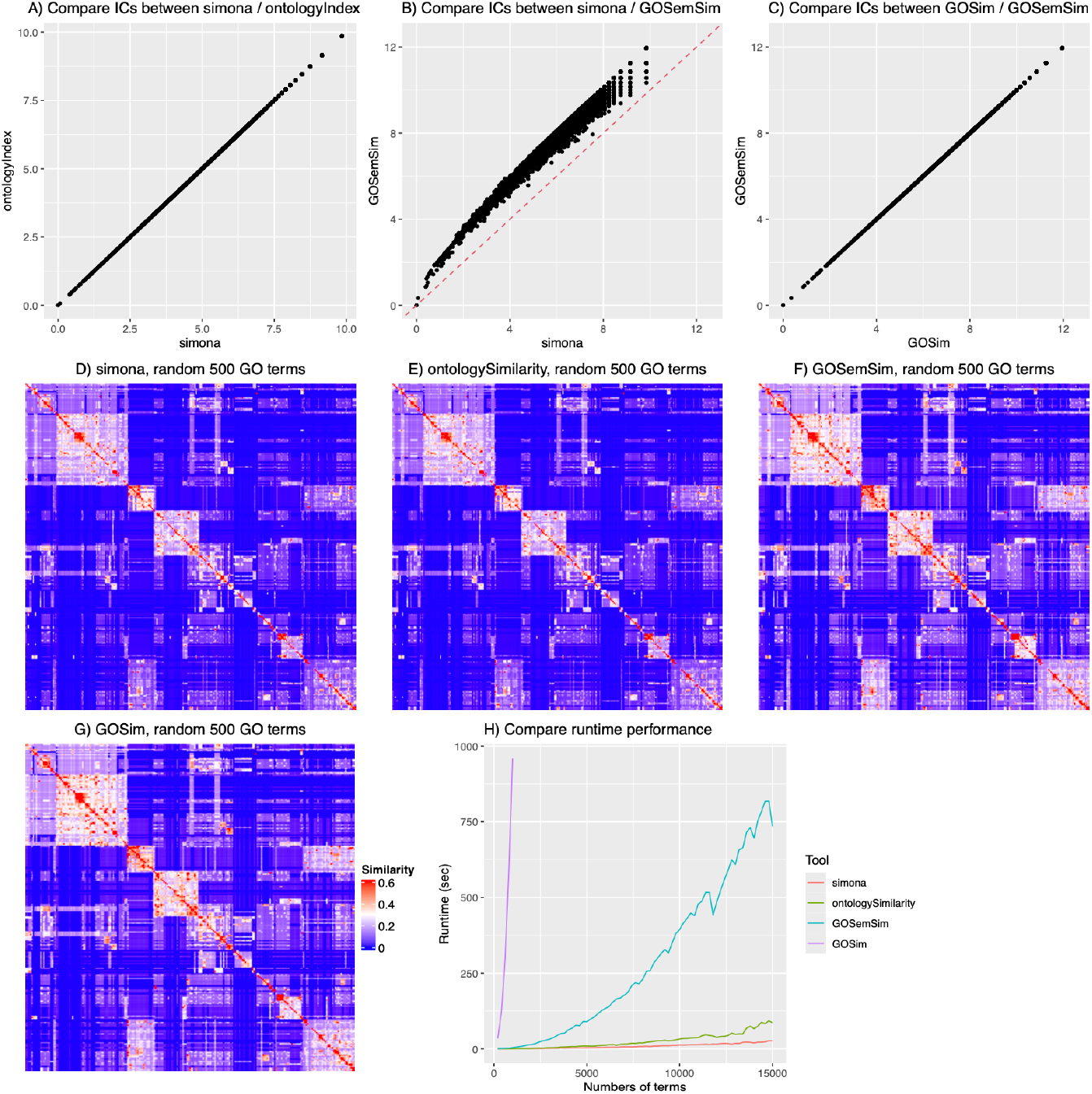
Compare *simona* to other tools. A-C) Pairwise comparisons of ICs of GO terms, calculated by *simona, ontologyIndex, GOSemSim* and *GOSim*. D-G) Heatmaps of semantic similarities of 500 random GO terms, using the Lin method implemented in the four tools. H) Runtime performance on the number of query GO terms.

The difference in ICs between the two groups of methods mainly comes from how they calculate the number of genes annotated to each GO term. Given the DAG structure of GO, when a GO term is annotated with a gene, all its ancestor terms are associated with that gene as well. In this way, to reduce the data size, a gene is normally only annotated to the lowest terms in the GO annotation database, and the aggregation of annotated genes on GO terms is recursively applied within individual software. It is worth noting, however, that there are also cases where a gene is annotated to both ancestor and offspring terms at the same time in the database file, possibly due to incomplete annotations.

*Simona* and *ontologyIndex* use the count of unique genes that are directly annotated to the GO term, as well as to its offspring terms. Let’s denote the set of annotated genes for term *x* as *G*_*x*_ and it is calculated as

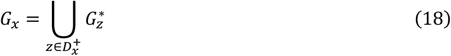

where 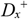 is the set of *x*’s offspring terms, including *x* itself, and 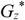 is the set of genes *directly* annotated to term *z*. Then the information content is

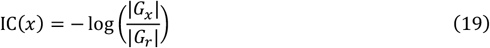

where *G*_*r*_ is the set of total genes annotated to the root term *r, i*.*e*., to the complete ontology.

*GOSemsim* and *GOSim* have a different way of calculating the numbers of annotated genes. They simply add the numbers of genes directly annotated to term *x* and its offspring terms without removing duplicated genes. In the following equation,

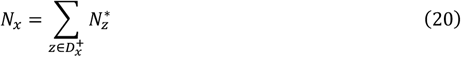

*N*_*x*_ is the number of genes for *x* and 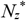 is the number of genes directly annotated to term *z*. The information content is then

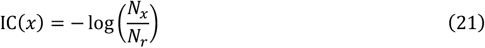

where *N*_*r*_ is the sum of the numbers of genes annotated to the whole ontology.

If a gene is only uniquely annotated to a distinct GO term in the ontology, Equation 19 and 21 give the same values. However, it is common for a gene to be annotated to multiple GO terms (on average, a gene is annotated to 7.3 GO BP terms), so on the common ancestors of these GO terms, the same gene will be counted multiple times. Additionally, even for genes directly annotated to the same GO term, the gene may be duplicated but in different evidence codes (37.6% of GO BP terms are directly annotated to duplicated genes). These factors lead to an overestimation of the number of annotated genes on a GO term when simply adding the numbers from its offspring terms, and in turn, as shown in Fig. 2B, it may overestimate ICs. Given that *p*(*x*) in the IC definition is the probability of observing a GO term when a gene is randomly picked, the method by *simona* and *ontologyIndex* is more proper under this context.

The method of annotation-based IC [19] was originally proposed and applied to WordNet [11] which is a taxonomy of concepts represented as nouns in English words, *e*.*g*., a coin is a subclass of cash. The probability of observing a concept is based on the frequency of that concept, as well as its offspring classes in a corpus gathered from a large collection of text. In that dataset, the frequency of observing the concept can be simply added from its offspring concepts because a concept is unique in the taxonomy and all concepts are represented as words. However, when applying this method to GO, adjustments need to be made because now the corpus consists of genes while concepts are GO terms. The mapping between the two is not one-to-one. Many studies adopt the default implementation without distinguishing duplicated gene annotations [28, 29], while only a few studies explicitly take unique genes [30, 31]. In Supplementary Fig. S1.3, we demonstrated that *GOSemSim* relatively overestimates ICs more for terms with smaller depths, *i*.*e*., the terms closer to the root.

We next compared the semantic similarity calculated by the four tools, using the method proposed by Lin [21]. Since not all GO terms have gene annotation (56.7% of GO BP terms have genes annotated), we randomly sampled 500 GO BP terms that have gene annotations. Figs 2D-G illustrate the similarity heatmaps of the 500 random GO terms calculated by the four tools. Once again, *simona* and *ontologySimilarity* produce identical similarity values, and *GOSemSim* and *GOSim* produce almost identical values. *GOSemSim* estimates systematically higher similarities than *simona*, due to its overestimation of ICs. *GOSemSim* relatively overestimates similarities for the term pairs whose MICA terms have small depths (Supplementary Fig. S1.6). Nevertheless, in general, the similarities calculated by the four tools are highly similar (Supplementary Fig. S1.5).

Last, we benchmarked the runtime performance of the four tools. The key step in Lin’s method is to traverse the DAG to look for the MICA of every term pair. It is also the dominant and most time-consuming part of the computation. We tested the runtime performance on calculating Lin’s similarity with varying numbers of random terms, ranging from 200 to 15,000 in steps of 200, on the GO BP ontology. Due to *GOSim*’s bad runtime performance, we only ran *GOSim* on no more than 1000 terms. Fig. 2H clearly illustrates that *simona* and *ontologySimilarity* show significantly better performance than *GOSemSim* and *GOSim*. Among the four tools, *simona* has the best performance. For example, with 10,000 query terms, *simona* achieves a 2.6x speedup compared to *ontologySimilarity* and a 31x speedup compared to *GOSemSim. Simona* has an even better runtime performance when the number of terms increase. The detailed benchmarking report is in Supplementary File 1.

### Compare IC and semantic similarity methods

Mazandu *et al*. [12] have provided a comprehensive, structured and theoretical overview of the state-of-the-art methods for semantic similarity analysis. However, a quantitative benchmark of IC and semantic similarity methods based on real-world data is still lacking in current studies. The comprehensive toolkit of *simona* makes it possible to perform comparisons of different methods on specific ontologies. In this section, we compared both IC and semantic similarity methods implemented in *simona*. For the benchmark, we took the BP namespace in GO as the test ontology, using relation types of “is a” and “part of”. The gene annotations to GO terms are based on human as the organism. Readers are encouraged to refer to Mazandu *et al*. [12] or the vignettes of *simona* for the comprehensive definitions of the methods mentioned in this section.

ICs for all GO terms are calculated using 11 IC methods implemented in *simona*. Fig. 3A illustrates the heatmap of the Pearson correlation of ICs from different IC methods.

**Fig. 3.**
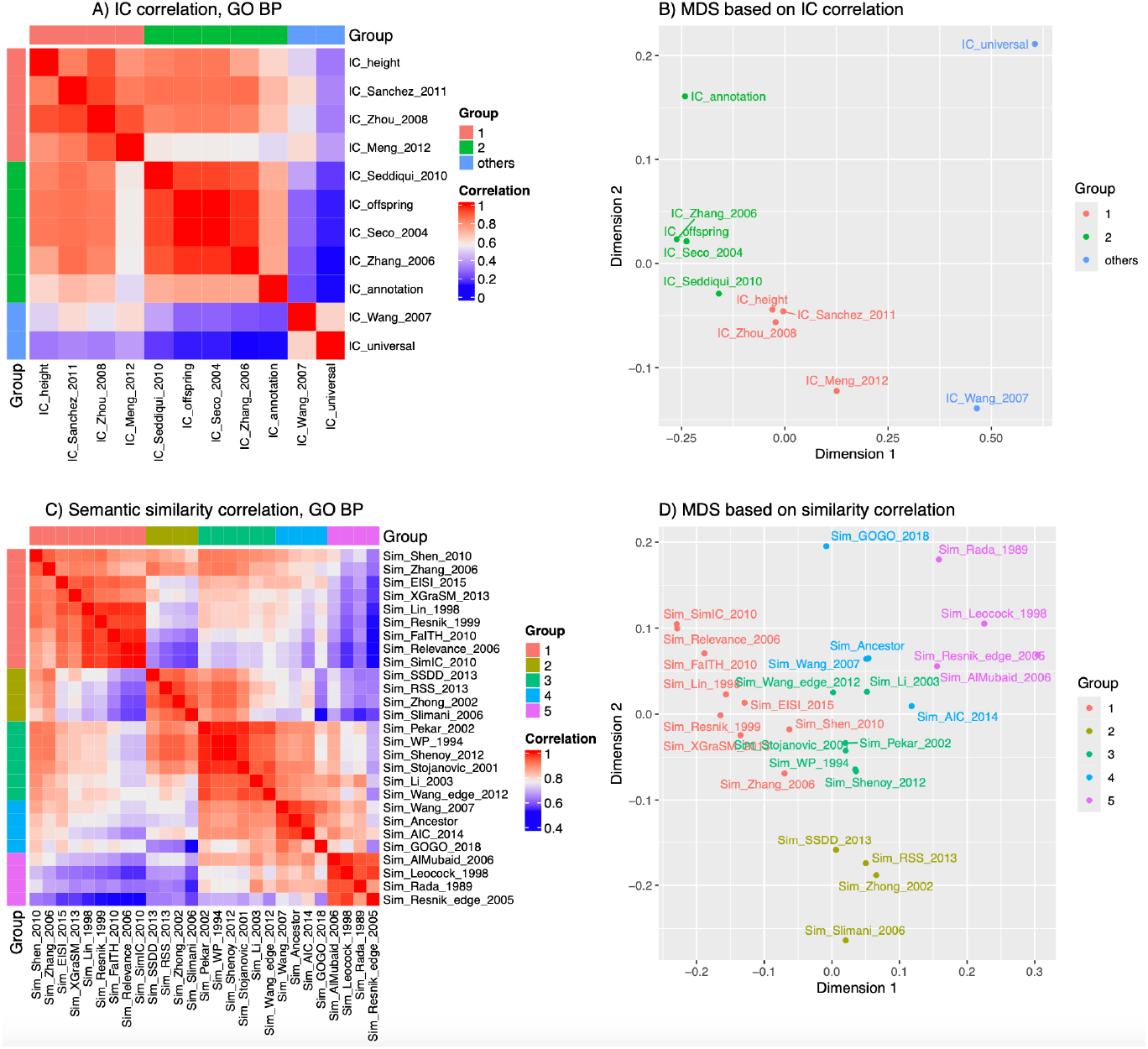
Compare IC and term-to-term similarity methods. A) Heatmap of Pearson correlations of IC values from different methods. B) The MDS on the correlation matrix in Fig. A. C) Heatmap of Pearson correlations of semantic similarity values from different methods. D) The MDS plot on the correlation matrix in Fig. C. In Figs A and C, partitionings on rows and columns are based on hierarchical clustering on the corresponding matrices.

Partitioning on rows and columns of heatmap is based on hierarchical clustering on the dissimilarity matrix of 1-cor where cor is the correlation matrix of ICs (Supplementary Fig. S3.1). In Fig. 3A, the two methods of *IC_Wang* and *IC_universal* are far away from other methods, which is also confirmed in the multidimensional scaling (MDS) analysis in Fig. 3B. These two methods are defined very differently from other IC methods. The remaining IC methods can be classified into two groups. It is interesting to note that, for methods in group 1, the heights or the depths of terms in the DAG play a major role in calculating IC values (*e*.*g*., the *IC_height* method), while for methods in group 2, the aggregation of all offspring terms plays a major role (*e*.*g*., the *IC_offspring* method). Supplementary Fig. S3.3 illustrates pairwise scatterplots of ICs from every two IC methods, which helps to inspect the trend of the correlation and how term depths are weighted in different IC methods.

We next compared term similarity methods. Semantic similarities of 124,750 term pairs, derived from 500 random GO BP terms with gene annotations, are calculated using 27 similarity methods implemented in *simona*. Seven out of a total of 34 similarity methods are removed from the comparison due to large differences from all other methods (the complete analysis can be found in Supplementary Fig S4.1). In this comparison, default settings are used for all similarity methods. Fig. 3C illustrates the heatmap of the Pearson correlation of semantic similarities from 27 methods. Methods are grouped into 5 clusters based on hierarchical clustering on 1-cor where cor is the correlation matrix of semantic similarities (Supplementary Fig S4.2). The grouping of methods can be confirmed on the MDS plot in Fig. 3D.

It is also interesting to note that the grouping of similarity methods reflects their similar methodologies. Group 1 mainly contains methods utilizing ICs from gene annotations. Methods in group 2 are heterogeneous and their mathematical forms are quite different. Methods in group 3 mainly relate to depths of LCA terms. Methods in group 4 relate to aggregation from all common ancestors, specifically the fraction of aggregation from the intersection of common ancestors to the aggregation from the union of common ancestors. Group 5 contains methods more related to the distances of two terms in the DAG. Supplementary Fig. S4.4 illustrates pairwise scatterplots of similarity values from every two term similarity methods, helping to inspect the trend of the correlation and how the depths of MICA/LCA terms are weighted in different similarity methods. Supplementary Fig. S4.5 illustrates similarity heatmaps of the 500 random GO terms from every similarity method for direct comparisons.

### Compare annotation-based and topology-based methods

Many existing R packages as well as other implementations for semantic similarity analysis [32, 33] are only based on ICs calculated from gene annotations. Therefore, they are primarily applied to GO only with well-annotated organisms such as human and mouse. In addition to annotation-based methods, *simona* supports a diverse range of methods based on the topology of the GO DAG, with no need for external annotation data. Here, we compared semantic similarity calculated based on gene annotations with that based on the topology of the GO DAG. For the benchmark, we took the BP namespace in GO as the test ontology, using relation types of “is a” and “part of”. The gene annotations to GO terms are based on human as the organism.

For the IC-based method (referred to here as annotation-based, denoted as “*IC_annotation*” for clarity), the semantic similarity is determined by the IC of the MICA of the term pairs using the Lin method; while for the topology-based method, the semantic similarity is determined by the depth of the LCA using the WP method. As introduced in the Methods section, the two methods have very similar forms. We compared, for a given set of 500 random GO terms resulting in 124,750 term pairs, how different their MICA terms and LCA terms are.

Fig. 4A indicates that for more than 98% of term pairs, their MICA and LCA terms are the same. Among the remaining 1.6% (1,976 term pairs), we observed that none of their MICA and LCA terms has an ancestor or offspring relation (Supplementary File 5). However, Fig. 4B reveals that for 73.0% of these 1,976 term pairs, their MICA and LCA terms are siblings, meaning they share the same parents, with an undirected distance of 2 (An example can be found in Supplementary Fig S5.1).

**Fig. 4.**
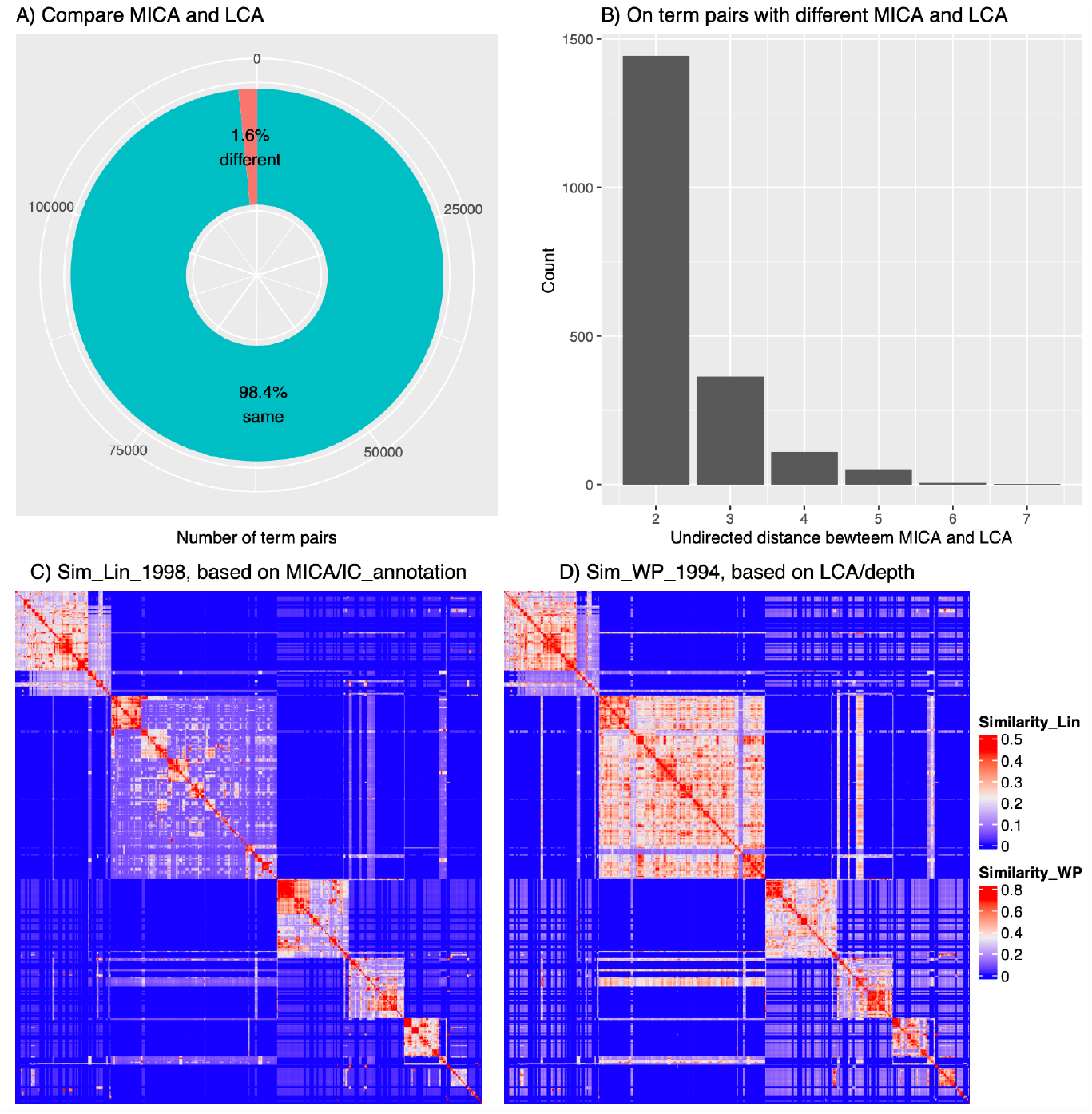
Compare the MICA and LCA of 500 random GO terms. A) Numbers and proportions of term pairs with identical or different terms as their MICA and LCA. B) Distribution of the undirected distance between MICA and LCA for 1,976 term pairs with differing MICA and LCA. C) Heatmap of semantic similarity using the Lin method. D) Heatmap of semantic similarity using the WP method. Row and column orders are the same in Figs C and D.

Note that the depth of a parent is always smaller than its child terms because the depth is defined as the longest distance from the root. This satisfies the requirement of the general IC definition in Equation 5. Under this context, depth can be considered as a special type of IC. On the other hand, Equation 18 and 19 show that *IC_annotation* is calculated based on the aggregation of gene annotations from offspring terms, which takes into account the DAG topology. By taking these two aspects together, we could say that depth and *IC_annotation* have similar forms and we would expect that generally, depth and *IC_annotaiton* have a strong positive correlation (Supplementary Fig. S5.3). In an extreme scenario where the ontology has a strictly binary tree structure and each term has a unique gene annotated, *IC_annotation* has an approximate linear relation to the depth (the last Equation in Supplementary File 5).

As depth can be thought of as an IC measurement, the LCA can be considered as a special type of MICA. According to the high correlation between *IC_annotation* and depth, as well as the large overlap between MICA and LCA terms, we would expect that the semantic similarities based on MICA and LCA should also be highly similar. In Figs 4C and 4D, the two heatmaps illustrate that the similarity patterns from the Lin method and the WP method are highly similar, where the WP method generates a higher level of similarity but also increases the inter-block signals off diagonals. Supplementary Fig. S5.5 illustrates that the two types of similarities exhibit strong positive correlations.

Most current analyses of semantic similarities rely on gene annotations, which require high-quality and complete annotation data. Unfortunately, only a very small number of ontologies, such as GO, have well-annotated data, and they are also limited to well-studied organisms. This significantly restricts the use of semantic similarity analysis on other less-studied organisms and makes it difficult to extend to other ontologies. The results presented in this section deliver a clear message that methods solely based on the topology of the ontology can also efficiently reveal the structure of semantic similarity. Together with the wide range of topology-based methods implemented in *simona*, it will greatly extend the semantic analysis of other organisms and ontologies.

### A gallery of OBO Foundry ontologies

The OBO Foundry [3] serves as a repository for public biological ontologies. We imported 206 ontologies from the OBO Foundry (data retrieved on 2023-08-07) and generated global circular visualization, similarity heatmaps and runtime performance analysis for each ontology. The complete gallery is available in Supplementary File 6.

One use of the OBO Foundry gallery is to explore ontology-specific global structures. Fig. 5 illustrates examples of four ontologies with different structures where each ontology is partitioned by automatically choosing a proper number of clusters.

In Fig. 5A, we visualized the NCBI Taxonomy [34] which contains over 2.5 million terms. NCBI taxonomy has a strict tree structure, with leaf terms as organisms and non-leaf terms indicating taxon classifications on specific levels. The widths of sectors correspond to the numbers of organisms in sub-classifications. For example, the largest cluster of organisms is under the taxon Endopterygota (a subset of insects, the brown cluster on the bottom left) and the term Endopterygota is comparably low in the complete taxonomy. If the depth of the taxonomy represents the extent of classification, we can find taxa of Viruses (the orange cluster on the right), Bacteria (the pink cluster on the top) and Fungi (the cyan cluster on the right) have small depths, thus comparably less well-studied, while the taxon led by Deuterostomia (a subset of animals, wine-red cluster on the bottom right), although only includes a small fraction of organisms in the taxonomy, extends very deep, indicating a well-studied group.

Fig. 5B illustrates the Chemical Entities of Biological Interest [35], a hierarchical organization of approximately 165,000 small chemical compounds. Being different from other ontologies in Fig. 5, it is more densely connected. The average number of parents is 1.45 which is also among the top ontologies in the OBO Foundry. Chemical compounds may be involved in multiple types of chemical reactions, thus it is very likely that they are associated with a large number of parent chemical functions.

**Fig. 5.**
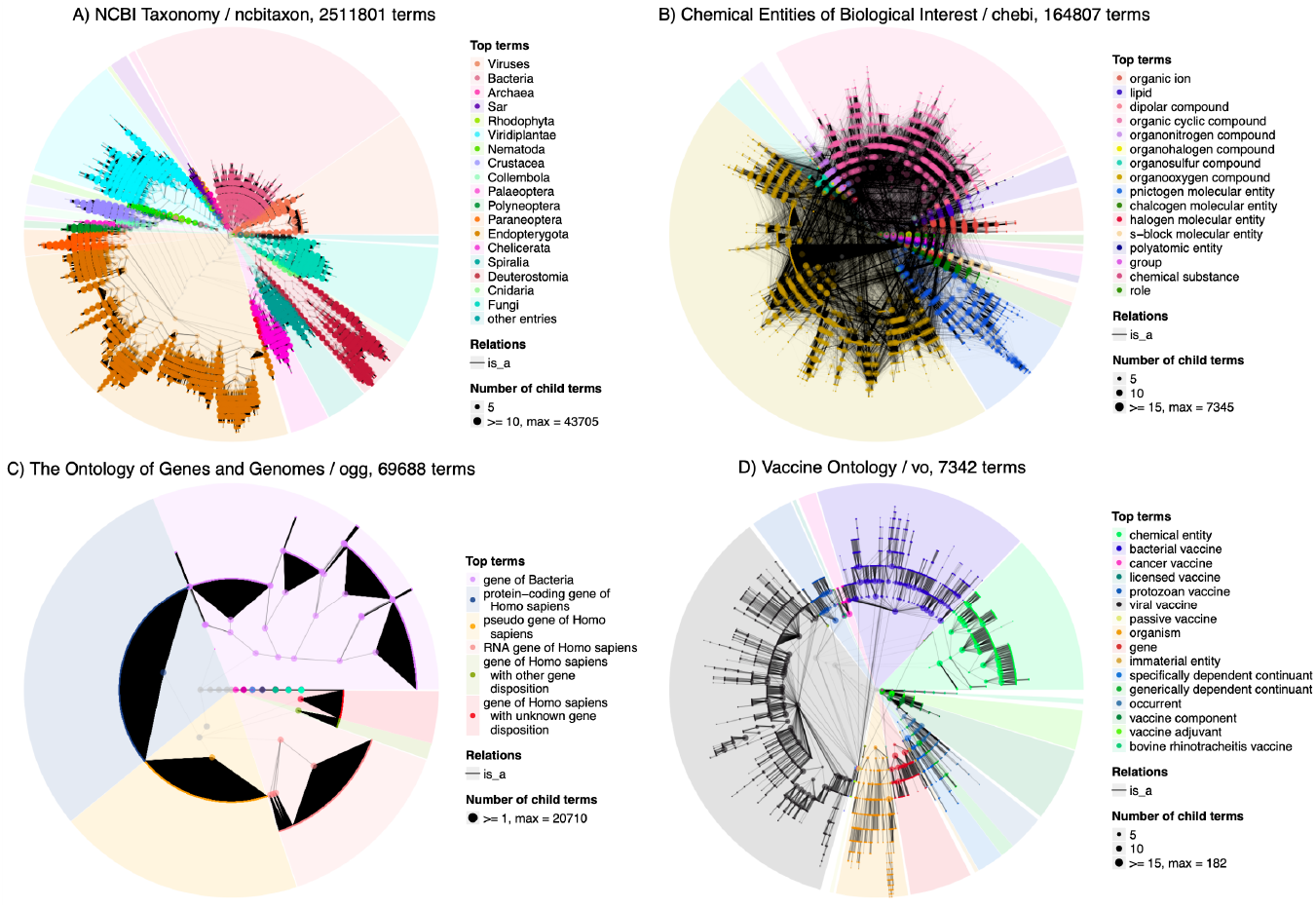
Circular visualization of four bio-ontologies. A) NCBI Taxonomy. B) Chemical Entities of Biological Interest. C) The Ontology of Genes and Genomes, D) Vaccine Ontology.

Fig. 5C illustrates the Ontology of Genes and Genomes (OGG) [36]. It has a very different structure characterized by huge numbers of leaf terms connecting to parents. This structure reflects the construction of OGG. The upper levels contain hierarchical relationships of gene types, *e*.*g*., “protein-coding gene of Homo sapiens” being a subclass of “gene of Homo sapiens”, while at the lowest level, specific genes are assigned to the most specific gene types within corresponding branches, such as all individual protein-coding genes, pseudogenes, RNA genes, *etc*.

Finally, Fig. 5D visualizes the Vaccine Ontology [37]. Despite its relatively small size, it exhibits a well-defined hierarchical structure, closely resembling a tree, with only 1.7% additional links from the treelized DAG.

## Conclusions

*Simona* offers a versatile interface and efficient implementation for processing, visualization, and semantic similarity analysis on bio-ontologies. We believe that *simona* will serve as a robust tool for uncovering relationships and enhancing the interoperability of biological knowledge systems.

## Availability and requirements

**Project name**: simona.

**Project home page**: https://bioconductor.org/packages/simona/.

**Operating system(s)**: Platform independent.

**Programming language**: R.

**License**: MIT.

## List of abbreviations

DAG: directed acyclic graph
GO: Gene ontology
IC: Information content
CA: Common ancestor
MICA: Most informative common ancestor
LCA: Lowest common ancestor
NCA: Nearest common ancestor
BP: Biological process.

## Declarations

### Ethics approval and consent to participate

Not applicable.

### Consent for publication

Not applicable.

## Availability of data and materials

The *simona* package as well as its documentation are freely available from https://bioconductor.org/packages/simona/. Supplementary files are also available from https://jokergoo.github.io/simona_supplementary/.

## Authors’ contributions

Z.G. designed the work, implemented the software, performed the analysis and wrote the manuscript. All authors reviewed the manuscript.

## Supporting information

Supplementary file 1: Compare to other tools. Supplementary file 2: The algorithm.

Supplementary file 3: Compare IC methods.

Supplementary file 4: Compare semantic similarity methods.

Supplementary file 5: Compare topology-based and annotation-based semantic similarity methods.

Supplementary file 6: OBO Foundry gallery.

