## Supplementary File 1 for "*simona:* a comprehensive R package for semantic similarity analysis on bio-ontologies": compare_tools.html

Supplementary File 1: Compare to other tools


### Supplementary File 1: Compare to other tools

###### Zuguang Gu

#### 2024-08-29

- Description of tools
  - simona
  - ontologyIndex and ontologySimilarity
  - GOSemSim
  - GOSim
  - Compare supported GO terms
- Compare Information contents
- Compare semantic similarities
- Runtime
- Session info

In this document, we compare **simona** to the following three R packages: **ontologyIndex**/**ontologySimilarity**, **GOSemSim** and **GOSim**. Because **GOSemSim** and **GOSim** only support Gene Ontology (GO), for this comparison we take GO (the Biological Process namespace) as the test ontology. **ontologyIndex** and **ontologySimilarity** belong to a suite of packages which are called **ontologyX** where **ontologyIndex** provides data structures and **ontologySimilarity** provides methods for semantic similarity analysis.

Different tools may have different GO versions and different default parameters. In the next section, we describe the differences of the four tools and the parameters we selected to make the differences of parameters between tools as small as possible.

**ontologyIndex**/**ontologySimilarity**, **GOSemSim** and **GOSim** mainly support information contents (IC) calculated from gene annotation, so in the comparison we will apply later, we only look at that IC method based on gene annotations, taking human as the organism.

There might be functions having similar names in different packages. To reduce the ambiguity, we add the `package::` prefix when calling functions from respective packages.

#### Description of tools

##### simona

**simona** uses the **GO.db** package as the GO source and the **org.Hs.eg.db** package as the gene annotation source. We take `"is_a"`, `"part_of"` and `"regulates"` as the GO relation types. Note that because `"regulates"` is a parent relation of `"negatively_regulates"` and `"positively_regulates"`, the latter two are automatically included.

In the next code, we calculate the information content based on the gene annotation on each GO term. On each GO term, **simona** takes the number of **unique** genes annotated to the term and to all its offspring terms. If a GO term has no gene annotated, its IC value is `NA`.

```
library(simona)
dag = simona::create_ontology_DAG_from_GO_db(namespace = "BP", org_db = "org.Hs.eg.db", 
    relations = c("part_of", "regulates"))
ic1 = simona::term_IC(dag, method = "IC_annotation")

go_id1 = names(ic1[!is.na(ic1)])
```

##### ontologyIndex and ontologySimilarity

**ontologySimilarity** provides a pre-computed object of ICs for all GO terms (`ontologySimilarity::GO_IC`), but the GO data is very old (generated on 2016). In this comparison, to be consistent to **GO.db**, we regenerate the ontology object with the `go-basic.obo` file downloaded from GO website (https://release.geneontology.org/2024-01-17/) ) with the same version as in **GO.db** (2024-01-17). The gene annotation is from the **org.Hs.eg.db** package.

On each term, **ontologyIndex** takes the number of **unique** genes annotated to the term and to all its offspring terms. See in the source code: https://github.com/cran/ontologyIndex/blob/1f83db5178e1b2501fa681e0b4ca4c117f4c25ef/R/sets.R#L94.

```
library(ontologyIndex)
onto = ontologyIndex::get_ontology("go-basic.obo", 
    propagate_relationships = c("is_a", "part_of", "regulates", "negatively_regulates", "positively_regulates"))

library(org.Hs.eg.db)
tb = toTable(org.Hs.egGO)
tb = tb[tb$Ontology == "BP", ]
# this ensures each GO term has genes annotated
lt = split(tb$go_id, tb$gene_id)

ic2 = ontologyIndex::get_term_info_content(onto, lt)
go_id2 = names(ic2)
```

##### GOSemSim

**GOSemSim** uses **GO.db** package as the GO source and **org.Hs.eg.db** as the gene annotation source. All relation types of `"is_a"`, `"part_of"`, `"regulates"`, `"negatively_regulates"` and `"positively_regulates"` are included.

On each GO term, **GOSemSim** takes the number of **all** genes annotated to the term and to all its offspring terms **without making genes unique**. Also note, a gene can be annotated to a GO term duplicatedly with multiple evidence codes (e.g. the gene with Entrez ID “994” is annotated to a GO term “GO:0000086” twice with evidence code “IBA” and “TAS”, check `org.Hs.egGO2EG[["GO:0000086"]]`). If a GO term has no gene annotated, its IC value is `Inf`.

```
library(GOSemSim)
data = GOSemSim::godata("org.Hs.eg.db", ont = "BP")
ic3 = data@IC[is.finite(data@IC)]
go_id3 = names(ic3)
```

##### GOSim

The settings is the same as **GOSemSim**. `GOSim::calcICs()` calculates the information contents and saves them to the object `IC` in a local file `"ICsBPhumanall.rda"`.

```
library(GOSim)
GOSim::setOntology("BP")
GOSim::calcICs()
load("ICsBPhumanall.rda") # it loads the object `IC`
ic4 = IC[is.finite(IC)]
go_id4 = names(ic4)
```

Note, `GOSim::setOntology()` not only sets the GO namespace, but also loads a pre-computed IC vector to the R session which will be used for calculating similarity. By default, it only loads the IC vector based on an old version of GO shipped with the package. To update the IC vector to the one we just generated, we need to run `GOSim::setOntology()` again by explicitly specifying the directory that contains `"ICsBPhumanall.rda"`.

ICs we just generated in `IC`:

```
head(IC)
```

```
## GO:0000001 GO:0000002 GO:0000003 GO:0000011 GO:0000012 GO:0000017 
##        Inf   8.018341   3.954860        Inf   9.116954  10.563873
```

ICs internally used for calculating semantic similarity. You can see they are different from values in `IC`. The object `GOSimEnv` is generated by **GOSim** and inserted to the global environment of the R session, i.e., `.GlobalEnv`.

```
head(GOSimEnv$IC)
```

```
## GO:0000001 GO:0000002 GO:0000003 GO:0042254 GO:0006467 GO:0000011 
##        Inf   8.666766   3.805017   6.161240   9.493444        Inf
```

We have to manually update the IC values that will be internally used. Now the internal ICs are the same as in `IC`:

```
GOSim::setOntology("BP", DIR = ".")
head(GOSimEnv$IC)
```

```
## GO:0000001 GO:0000002 GO:0000003 GO:0000011 GO:0000012 GO:0000017 
##        Inf   8.018341   3.954860        Inf   9.116954  10.563873
```

##### Compare supported GO terms

We compare the supported GO terms in the four tools. The UpSet plot shows all the four tools share almost the same set of GO terms, except **GOSemSim** loses a tiny amount of them. This is expected because all the tools are based on the same version of GO data.

```
lt = list(simona = go_id1, ontologyIndex = go_id2, GOSemSim = go_id3, GOSim = go_id4)
library(ComplexHeatmap)
UpSet(make_comb_mat(lt), column_title = "Numbers of supported GO BP terms")
```

Figure S1.1. Supported GO BP terms in simona, ontologyIndex, GOSim and GOSemSim.

We take the common GO terms in the four lists for later analyses.

```
go_id_common = intersect(intersect(go_id1, go_id2), intersect(go_id3, go_id4))
length(go_id_common)
```

```
## [1] 15405
```

#### Compare Information contents

We simply make scatterplots for every pair of tools.

```
df = data.frame(
    simona        = ic1[go_id_common],
    ontologyIndex = ic2[go_id_common],
    GOSemSim      = ic3[go_id_common],
    GOSim         = ic4[go_id_common]
)
pairs(df, pch = ".", col = "#00000040", xlim = c(0, 12), ylim = c(0, 12),
    main = "Pairwise comparions of ICs")
```

Figure S1.2. Pairwise comparison of ICs of GO BP, calculated by simona, ontologyIndex, GOSemSim and GOSim.

**simona** and **ontologyIndex** calculate identical IC values, **GOSemSim** and **GOSim** calculate identical IC values. This can be validated by the following code:

```
sum(abs(df$simona - df$ontologyIndex) < 1e-10)/nrow(df)
```

```
## [1] 1
```

```
sum(abs(df$GOSemSim - df$GOSim) < 1e-10)/nrow(df)
```

```
## [1] 1
```

This is expected because **simona**/**ontologyIndex** use the same method for calculating ICs, and **GOSemSim**/**GOSim** use another same method. **simona** and **GOSemSim**, although not identical, still show very strong linear relations, but with a systematic shift.

The difference between **simona** and **GOSemSim** is mainly due to how they calculate the number of genes annotated to each GO term.

Because of the nature of the DAG structure of GO, if a GO term is annotated to a gene, all its ancestor terms are also associated with that gene. The calculation of annotated genes is applied in a recursive way.

**Method used by simona/ontologyIndex**: Denote \(G^\*\_x\) as the set of genes *directly* annotated to the GO term \(x\), and \(G\_x\) as the set of genes annotated to \(x\) after merging from its child terms, \(G\_x\) is the union of all genes annotated to its child terms and the genes directly annotated to \(x\).

\[ G\_x = \left( \bigcup\_{z \in \mathrm{C}\_x} G\_z \right) \bigcup G^\*\_x \]

where \(\mathrm{C}\_x\) is the set of \(x\)’s child terms.

The information content is:

\[ \mathrm{IC}(x) = -\log \left( \frac{|G\_x|}{|G\_r|} \right) \]

where \(G\_r\) is the set of all genes annotated to all GO terms (i.e., the set of genes annotated to the root term \(r\) of the GO DAG).

\(G\_x\) can also be written as:

\[ G\_x = \bigcup\_{z \in \mathrm{D}^+\_x} G\_z^\* \]

where \(\mathrm{D}^+\_x\) is the set of \(x\)’s offspring terms plus \(x\) itself. In this way, \(G\_x\) is the union of all genes annotated to \(x\) and \(x\)’s offspring terms.

**Method used by GOSemsim/GOSim**: Since genes are not uniquified, we cannot use the same notations here. Let’s denote \(N^\*\_x\) as the number of genes *directly* annotated to term \(x\), and \(N\_x\) is the number of genes annotated to \(x\) after merging from its child terms. Note that \(N^\*\_x \ne |G^\*\_x|\) unless all genes directly annotated to \(x\) are unique. \(N\_x\) is calculated as:

\[ N\_x = N^\*\_x + \sum\_{z \in \mathrm{C}\_x} N\_z \]

It can also be written as:

\[ N\_x = \sum\_{z \in \mathrm{D}^+\_x} N^\*\_z \]

The information content is

\[ \mathrm{IC}(x) = -\log \left( \frac{N\_x}{N\_r} \right) \]

where \(N\_r\) is the value for the root term.

According to the two different ways of calculating gene annotation-based IC, the differences come from 1) a gene can be annotated to multiple terms; 2) even for the same term, a gene can be duplicatedly annotated with different evidence codes. As already shown in the scatter plot of pairwise comparison in Figure S1.2, **GOSemsim/GOSim** systematically over-estimates IC values. The two types of methods are the same when a gene is only uniquely annotated to a single GO term.

The next plots show that **GOSemsim** over-estimates the IC values more for the terms with smaller depth (i.e. closer to the root and have more accumulation from offsprings).

```
depth = dag_depth(dag)
df$depth = depth[rownames(df)]
par(mfrow = c(1, 2))
boxplot(df$GOSemSim/df$simona ~ df$depth, xlab = "Depth", ylab = "IC_GOSemSim/IC_simona")
barplot(table(df$depth), xlab = "Depth", ylab = "Counts")
```

Figure S1.3. Systematic shift of IC values from simona to GOSemSim. Left: over-estimation on the IC values on each depth. Right: number of terms on each depth.

#### Compare semantic similarities

We randomly pick 500 GO terms and calculate the semantic similarities among them. For all the four tools, we use the “Lin 1998” method.

```
set.seed(123)
go_id = sample(go_id_common, 500)

m1 = simona::term_sim(dag, go_id, method = "Sim_Lin_1998")
m2 = ontologySimilarity::get_term_sim_mat(ontology = onto, information_content = ic2, 
    method = "lin", row_terms = go_id, col_terms = go_id)
m3 = GOSemSim::termSim(go_id, go_id, semData = data, method = "Lin")
m4 = GOSim::getTermSim(go_id, method = "Lin")
```

We visualize the four matrices as heatmaps. Since they are symmetric and correspond to the same set of GO terms, we align the four heatmaps to let them have the same row orders and column orders.

```
library(circlize)
# set the same color mapping to the four heatmaps
col_fun = colorRamp2(c(0, 0.3, 0.6), c("blue", "white", "red"))

od = hclust(dist(m1))$order

Heatmap(m1, name = "similarity", col = col_fun,
        show_row_names = FALSE, show_column_names = FALSE,
        row_order = od, column_order = od,
        column_title = "simona, random 500 GO terms") +
Heatmap(m2, name = "similarity", col = col_fun,
        show_row_names = FALSE, show_column_names = FALSE,
        row_order = od, column_order = od,
        column_title = "ontologySimilarity, random 500 GO terms") +
Heatmap(m3, name = "similarity", col = col_fun, 
        show_row_names = FALSE, show_column_names = FALSE,
        row_order = od, column_order = od,
        column_title = "GOSemSim, random 500 GO terms") +
Heatmap(m4, name = "similarity", col = col_fun, 
        show_row_names = FALSE, show_column_names = FALSE,
        row_order = od, column_order = od,
        column_title = "GOSim, random 500 GO terms")
```

Figure S1.4. Heatmaps of semantic similarities of 500 random GO BP terms calculated by Lin 1998 method.

**simona** and **ontologySimilarity** generate identical results. **GOSemSim** and **GOSim** generates almost identical results. It can be validated by the following code:

```
sum(abs(m1 - m2) < 1e-10)/250000  # 500*500
```

```
## [1] 1
```

```
sum(abs(m3 - m4) < 1e-10)/250000
```

```
## [1] 0.99996
```

**GOSemSim** and **GOSim** estimate systematically higher similarities than **simona** and **ontologySimilarity** (the heatmaps are more redish), due to that they two over-estimate the ICs.

We can also perform a pairwise comparison on semantic similarities between tools. We can draw the same conclusions as from the heatmaps.

```
df = data.frame(
    simona             = m1[lower.tri(m1)],
    ontologySimilarity = m2[lower.tri(m2)],
    GOSemSim           = m3[lower.tri(m3)],
    GOSim              = m4[lower.tri(m4)]
)
pairs(df, pch = ".", col = "#00000040", xlim = c(0, 1), ylim = c(0, 1),
    main = "Pairwise comparison of semantic similarities")
```

Figure S1.5. Pairwise comparison of semantic similarities from the 500 random GO BP terms.

The next plots show that **GOSemsim** over-estimates the similarity values more for the MICA terms with smaller depth (i.e. closer to the root).

```
mica = MICA_term(dag, go_id)
mica = mica[lower.tri(mica)]
df$depth = depth[mica]
par(mfrow = c(1, 2))
boxplot(df$GOSemSim/df$simona ~ df$depth, xlab = "Depth of MICA", ylab = "sim_GOSemSim/sim_simona")
barplot(table(df$depth), xlab = "Depth of MICA", ylab = "Counts")
```

Figure S1.6. Systematic shift of semantic similarity values from simona to GOSemSim. Left: over-estimation on the similarity values on each depth of MICA terms. Right: number of MICA terms on each depth. A depth of zero means the MICA term is the root term.

#### Runtime

The key step in Lin’s method is to look for MICA of every two terms. It is also the dominant and most time-consuming part in the computation. In the code below, we benchmark the runtime of the four tools on looking for MICA on different numbers of terms.

Because **GOSim** has a bad runtime performance, we only run `GOSim::getTermSim()` on more than 1000 terms.

```
set.seed(666)
k = seq(200, 15000, by = 200)
t1 = t2 = t3 = t4 = rep(NA_real_, length(k))
for(i in seq_along(k)) {
    go_id = sample(go_id_common, k[i])
    t1[i] = system.time(simona::term_sim(dag, go_id, method = "Sim_Lin_1998"))[3]
    t2[i] = system.time(ontologySimilarity::get_term_sim_mat(ontology = onto, information_content = ic2, 
        method = "lin", row_terms = go_id, col_terms = go_id))[3]
    t3[i] = system.time(GOSemSim::termSim(go_id, go_id, semData= data, method = "Lin"))[3]
    if(k[i] <= 1000) t4[i] = system.time(GOSim::getTermSim(go_id, method = "Lin"))[3]
}
```

The following plot shows **simona** and **ontologySimilarity** show far better performance than **GOSemSim** and **GOSim**. **simona** has the best performance among the four tools.

```
par(mfrow = c(1, 2))
plot(NULL, xlim = c(0, 15000), ylim = c(0, max(t3, t4, na.rm = TRUE)), 
    xlab = "Number of terms", ylab = "runtime (sec)",
    main = "Compare runtime of the four tools")
lines(k, t1, col = 2)
lines(k, t2, col = 3)
lines(k, t3, col = 4)
lines(k, t4, col = 7)

legend("top", lty = 1, col = c(2, 3, 4, 7), 
    legend = c("simona", "ontologySimilarity", "GOSemSim", "GOSim"))

# the second plot
plot(NULL, xlim = c(0, 15000), ylim = c(0, max(t2)), 
    xlab = "Number of terms", ylab = "runtime (sec)",
    main = "Compare runtime")
lines(k, t1, col = 2)
lines(k, t2, col = 3)

legend("top", lty = 1, col = c(2, 3), 
    legend = c("simona", "ontologySimilarity"))
```

Figure S1.7. Runtime of simona, ontologySimilarity, GOSemSim and GOSim on calculating semantic similarities of the random 500 GO BP terms. Left: compare all four tools. Right: only compare simona and ontologySimilarity.

We can also directly compare the fold improvement of runtime of **simona** to other tools. It shows that approximately, **simona** has 2.6x speedup than **ontologySimilarity** and 31x speedup than **GOSemSim**.

```
par(mfrow = c(1, 3))
plot(k, t2/t1, xlab = "Number of terms", ylab = "ontologySimilarity / simona", main = "Runtime fold improvement")
plot(k, t3/t1, xlab = "Number of terms", ylab = "GOSemSim / simona", main = "Runtime fold improvement")
plot(k, t3/t2, xlab = "Number of terms", ylab = "GOSemSim / ontologySimilarity", main = "Runtime fold improvement")
```

Figure S1.8. Runtime speed improvement of simone to other tools.

#### Session info

```
sessionInfo()
```

```
## R version 4.4.1 (2024-06-14)
## Platform: x86_64-apple-darwin20
## Running under: macOS Sonoma 14.5
## 
## Matrix products: default
## BLAS:   /Library/Frameworks/R.framework/Versions/4.4-x86_64/Resources/lib/libRblas.0.dylib 
## LAPACK: /Library/Frameworks/R.framework/Versions/4.4-x86_64/Resources/lib/libRlapack.dylib;  LAPACK version 3.12.0
## 
## locale:
## [1] en_GB.UTF-8/en_GB.UTF-8/en_GB.UTF-8/C/en_GB.UTF-8/en_GB.UTF-8
## 
## time zone: Europe/Berlin
## tzcode source: internal
## 
## attached base packages:
## [1] grid      stats4    stats     graphics  grDevices utils     datasets 
## [8] methods   base     
## 
## other attached packages:
##  [1] circlize_0.4.16       ComplexHeatmap_2.20.0 GOSim_1.42.0         
##  [4] annotate_1.82.0       XML_3.99-0.17         GO.db_3.19.1         
##  [7] GOSemSim_2.30.2       org.Hs.eg.db_3.19.1   AnnotationDbi_1.66.0 
## [10] IRanges_2.38.1        S4Vectors_0.42.1      Biobase_2.64.0       
## [13] BiocGenerics_0.50.0   ontologyIndex_2.12    simona_1.3.12        
## [16] knitr_1.48            rmarkdown_2.28       
## 
## loaded via a namespace (and not attached):
##  [1] blob_1.2.4              R.utils_2.12.3          Biostrings_2.72.1      
##  [4] fastmap_1.2.0           promises_1.3.0          digest_0.6.37          
##  [7] mime_0.12               lifecycle_1.0.4         cluster_2.1.6          
## [10] Cairo_1.6-2             topGO_2.56.0            KEGGREST_1.44.1        
## [13] RSQLite_2.3.7           magrittr_2.0.3          compiler_4.4.1         
## [16] rlang_1.1.4             sass_0.4.9              tools_4.4.1            
## [19] igraph_2.0.3            yaml_2.3.10             bit_4.0.5              
## [22] scatterplot3d_0.3-44    xml2_1.3.6              RColorBrewer_1.1-3     
## [25] R.oo_1.26.0             nnet_7.3-19             xtable_1.8-4           
## [28] colorspace_2.1-1        iterators_1.0.14        cli_3.6.3              
## [31] crayon_1.5.3            httr_1.4.7              rjson_0.2.22           
## [34] DBI_1.2.3               cachem_1.1.0            modeltools_0.2-23      
## [37] zlibbioc_1.50.0         parallel_4.4.1          XVector_0.44.0         
## [40] matrixStats_1.3.0       vctrs_0.6.5             yulab.utils_0.1.7      
## [43] Matrix_1.7-0            SparseM_1.84-2          jsonlite_1.8.8         
## [46] GetoptLong_1.0.5        RBGL_1.80.0             bit64_4.0.5            
## [49] clue_0.3-65             foreach_1.5.2           jquerylib_0.1.4        
## [52] glue_1.7.0              codetools_0.2-20        Polychrome_1.5.1       
## [55] shape_1.4.6.1           later_1.3.2             GenomeInfoDb_1.40.1    
## [58] UCSC.utils_1.0.0        rappdirs_0.3.3          htmltools_0.5.8.1      
## [61] graph_1.82.0            GenomeInfoDbData_1.2.12 R6_2.5.1               
## [64] httr2_1.0.3             doParallel_1.0.17       ontologySimilarity_2.7 
## [67] lattice_0.22-6          evaluate_0.24.0         shiny_1.9.1            
## [70] highr_0.11              png_0.1-8               R.methodsS3_1.8.2      
## [73] memoise_2.0.1           corpcor_1.6.10          httpuv_1.6.15          
## [76] bslib_0.8.0             flexmix_2.3-19          Rcpp_1.0.13            
## [79] xfun_0.47               fs_1.6.4                pkgconfig_2.0.3        
## [82] GlobalOptions_0.1.2
```
