## Supplementary File 1 for "*simona:* a comprehensive R package for semantic similarity analysis on bio-ontologies": jquery.sticky.js

```
// Sticky Plugin v1.0.4 for jQuery
// =============
// Author: Anthony Garand
// Improvements by German M. Bravo (Kronuz) and Ruud Kamphuis (ruudk)
// Improvements by Leonardo C. Daronco (daronco)
// Created: 02/14/2011
// Date: 07/20/2015
// Website: http://stickyjs.com/
// Description: Makes an element on the page stick on the screen as you scroll
//              It will only set the 'top' and 'position' of your element, you
//              might need to adjust the width in some cases.

(function (factory) {
    if (typeof define === 'function' && define.amd) {
        // AMD. Register as an anonymous module.
        define(['jquery'], factory);
    } else if (typeof module === 'object' && module.exports) {
        // Node/CommonJS
        module.exports = factory(require('jquery'));
    } else {
        // Browser globals
        factory(jQuery);
    }
}(function ($) {
    var slice = Array.prototype.slice; // save ref to original slice()
    var splice = Array.prototype.splice; // save ref to original slice()

  var defaults = {
      topSpacing: 0,
      bottomSpacing: 0,
      className: 'is-sticky',
      wrapperClassName: 'sticky-wrapper',
      center: false,
      getWidthFrom: '',
      widthFromWrapper: true, // works only when .getWidthFrom is empty
      responsiveWidth: false,
      zIndex: 'inherit'
    },
    $window = $(window),
    $document = $(document),
    sticked = [],
    windowHeight = $window.height(),
    scroller = function() {
      var scrollTop = $window.scrollTop(),
        documentHeight = $document.height(),
        dwh = documentHeight - windowHeight,
        extra = (scrollTop > dwh) ? dwh - scrollTop : 0;

      for (var i = 0, l = sticked.length; i < l; i++) {
        var s = sticked[i],
          elementTop = s.stickyWrapper.offset().top,
          etse = elementTop - s.topSpacing - extra;

        //update height in case of dynamic content
        s.stickyWrapper.css('height', s.stickyElement.outerHeight());

        if (scrollTop <= etse) {
          if (s.currentTop !== null) {
            s.stickyElement
              .css({
                'width': '',
                'position': '',
                'top': '',
                'z-index': ''
              });
            s.stickyElement.parent().removeClass(s.className);
            s.stickyElement.trigger('sticky-end', [s]);
            s.currentTop = null;
          }
        }
        else {
          var newTop = documentHeight - s.stickyElement.outerHeight()
            - s.topSpacing - s.bottomSpacing - scrollTop - extra;
          if (newTop < 0) {
            newTop = newTop + s.topSpacing;
          } else {
            newTop = s.topSpacing;
          }
          if (s.currentTop !== newTop) {
            var newWidth;
            if (s.getWidthFrom) {
                padding =  s.stickyElement.innerWidth() - s.stickyElement.width();
                newWidth = $(s.getWidthFrom).width() - padding || null;
            } else if (s.widthFromWrapper) {
                newWidth = s.stickyWrapper.width();
            }
            if (newWidth == null) {
                newWidth = s.stickyElement.width();
            }
            s.stickyElement
              .css('width', newWidth)
              .css('position', 'fixed')
              .css('top', newTop)
              .css('z-index', s.zIndex);

            s.stickyElement.parent().addClass(s.className);

            if (s.currentTop === null) {
              s.stickyElement.trigger('sticky-start', [s]);
            } else {
              // sticky is started but it have to be repositioned
              s.stickyElement.trigger('sticky-update', [s]);
            }

            if (s.currentTop === s.topSpacing && s.currentTop > newTop || s.currentTop === null && newTop < s.topSpacing) {
              // just reached bottom || just started to stick but bottom is already reached
              s.stickyElement.trigger('sticky-bottom-reached', [s]);
            } else if(s.currentTop !== null && newTop === s.topSpacing && s.currentTop < newTop) {
              // sticky is started && sticked at topSpacing && overflowing from top just finished
              s.stickyElement.trigger('sticky-bottom-unreached', [s]);
            }

            s.currentTop = newTop;
          }

          // Check if sticky has reached end of container and stop sticking
          var stickyWrapperContainer = s.stickyWrapper.parent();
          var unstick = (s.stickyElement.offset().top + s.stickyElement.outerHeight() >= stickyWrapperContainer.offset().top + stickyWrapperContainer.outerHeight()) && (s.stickyElement.offset().top <= s.topSpacing);

          if( unstick ) {
            s.stickyElement
              .css('position', 'absolute')
              .css('top', '')
              .css('bottom', 0)
              .css('z-index', '');
          } else {
            s.stickyElement
              .css('position', 'fixed')
              .css('top', newTop)
              .css('bottom', '')
              .css('z-index', s.zIndex);
          }
        }
      }
    },
    resizer = function() {
      windowHeight = $window.height();

      for (var i = 0, l = sticked.length; i < l; i++) {
        var s = sticked[i];
        var newWidth = null;
        if (s.getWidthFrom) {
            if (s.responsiveWidth) {
                newWidth = $(s.getWidthFrom).width();
            }
        } else if(s.widthFromWrapper) {
            newWidth = s.stickyWrapper.width();
        }
        if (newWidth != null) {
            s.stickyElement.css('width', newWidth);
        }
      }
    },
    methods = {
      init: function(options) {
        return this.each(function() {
          var o = $.extend({}, defaults, options);
          var stickyElement = $(this);

          var stickyId = stickyElement.attr('id');
          var wrapperId = stickyId ? stickyId + '-' + defaults.wrapperClassName : defaults.wrapperClassName;
          var wrapper = $('<div></div>')
            .attr('id', wrapperId)
            .addClass(o.wrapperClassName);

          stickyElement.wrapAll(function() {
            if ($(this).parent("#" + wrapperId).length == 0) {
                    return wrapper;
            }
});

          var stickyWrapper = stickyElement.parent();

          if (o.center) {
            stickyWrapper.css({width:stickyElement.outerWidth(),marginLeft:"auto",marginRight:"auto"});
          }

          if (stickyElement.css("float") === "right") {
            stickyElement.css({"float":"none"}).parent().css({"float":"right"});
          }

          o.stickyElement = stickyElement;
          o.stickyWrapper = stickyWrapper;
          o.currentTop    = null;

          sticked.push(o);

          methods.setWrapperHeight(this);
          methods.setupChangeListeners(this);
        });
      },

      setWrapperHeight: function(stickyElement) {
        var element = $(stickyElement);
        var stickyWrapper = element.parent();
        if (stickyWrapper) {
          stickyWrapper.css('height', element.outerHeight());
        }
      },

      setupChangeListeners: function(stickyElement) {
        if (window.MutationObserver) {
          var mutationObserver = new window.MutationObserver(function(mutations) {
            if (mutations[0].addedNodes.length || mutations[0].removedNodes.length) {
              methods.setWrapperHeight(stickyElement);
            }
          });
          mutationObserver.observe(stickyElement, {subtree: true, childList: true});
        } else {
          if (window.addEventListener) {
            stickyElement.addEventListener('DOMNodeInserted', function() {
              methods.setWrapperHeight(stickyElement);
            }, false);
            stickyElement.addEventListener('DOMNodeRemoved', function() {
              methods.setWrapperHeight(stickyElement);
            }, false);
          } else if (window.attachEvent) {
            stickyElement.attachEvent('onDOMNodeInserted', function() {
              methods.setWrapperHeight(stickyElement);
            });
            stickyElement.attachEvent('onDOMNodeRemoved', function() {
              methods.setWrapperHeight(stickyElement);
            });
          }
        }
      },
      update: scroller,
      unstick: function(options) {
        return this.each(function() {
          var that = this;
          var unstickyElement = $(that);

          var removeIdx = -1;
          var i = sticked.length;
          while (i-- > 0) {
            if (sticked[i].stickyElement.get(0) === that) {
                splice.call(sticked,i,1);
                removeIdx = i;
            }
          }
          if(removeIdx !== -1) {
            unstickyElement.unwrap();
            unstickyElement
              .css({
                'width': '',
                'position': '',
                'top': '',
                'float': '',
                'z-index': ''
              })
            ;
          }
        });
      }
    };

  // should be more efficient than using $window.scroll(scroller) and $window.resize(resizer):
  if (window.addEventListener) {
    window.addEventListener('scroll', scroller, false);
    window.addEventListener('resize', resizer, false);
  } else if (window.attachEvent) {
    window.attachEvent('onscroll', scroller);
    window.attachEvent('onresize', resizer);
  }

  $.fn.sticky = function(method) {
    if (methods[method]) {
      return methods[method].apply(this, slice.call(arguments, 1));
    } else if (typeof method === 'object' || !method ) {
      return methods.init.apply( this, arguments );
    } else {
      $.error('Method ' + method + ' does not exist on jQuery.sticky');
    }
  };

  $.fn.unstick = function(method) {
    if (methods[method]) {
      return methods[method].apply(this, slice.call(arguments, 1));
    } else if (typeof method === 'object' || !method ) {
      return methods.unstick.apply( this, arguments );
    } else {
      $.error('Method ' + method + ' does not exist on jQuery.sticky');
    }
  };
  $(function() {
    setTimeout(scroller, 0);
  });
}));
```
