## Supplementary File 2 for "*simona:* a comprehensive R package for semantic similarity analysis on bio-ontologies": algorithm.html

Supplementary File 2: The Algorithm


### Supplementary File 2: The Algorithm

###### Zuguang Gu

#### 2024-08-29

- The algorithm
- Time complexity
- Session info

#### The algorithm

**The task**: for a given set of query terms (\(n \ge 2\)), calculate their semantic similarities.

The key step is to look for the tuple of three items \((a, x\_1, x\_2)\) where \(a\) is a common ancestor of the two terms \(x\_1\) and \(x\_2\). **simona** implements a “top-down” algorithm to traverse the DAG to obtain such tuples where \(a\) is always a common ancestor of \(x\_1\) and \(x\_2\).

Figure S2.1. Algorithm for looking for LCA terms implemented in simona.

We use a simplified DAG in Figure S2.1 as an example to explain the algorithm used in **simona**. In this example, we want to calculate depths of LCA (the lowest common ancestors) terms of four query terms of \(x\_1\), \(x\_2\), \(x\_3\) and \(x\_4\) (where semantic similarities can be further calculated based on the LCA depth, and the process is the same as looking for MICA terms). The depths of LCA terms of the four query terms are saved in a symmetric matrix denoted as \(M\). The algorithm includes two steps:

1. Find all the ancestors of \(x\_1\), \(x\_2\), \(x\_3\) and \(x\_4\), also including the four query terms. This can be done by the breadth-first search (BFS) or depth-first search (DFS). We denote this set of ancestor terms as \(A\). In Figure S2.1A, \(A\) includes terms in blue and red.
2. For every term \(a\) in \(A\), there are two sub-steps:

   1. Look for the subset of query terms that are offsprings of \(a\). This can be done by BFS or DFS. Let’s denote this subset of query terms as \(D\).
   2. If \(D\) has at least two terms, then for every pair of terms in \(D\), \(a\) is always their common ancestor. Thus, we only need to update values in the sub-matrix of \(M\) that corresponds to terms in \(D\), by comparing to the depth of \(a\).

   Figure S2.1 B-F as well as matrices in the bottom right of Figure S2.1 illustrate this process. Note in the figures, we start from term \(a\_1\) which is the root in \(A\), but the order of B-F does not matter and it can be randomly permutated.

The advantage of this algorithm is, when traversing from each ancestor to query terms, we can always ensure in the tuple \((a, x\_i, x\_j)\), \(a\) is a common ancestor of \(x\_i\) and \(x\_j\).

Current tools, e.g. **GOSemSim** and **ontologySimilarity**, use similar implementations, which is a “bottom-up” method, going through every pair of query terms. The algorithm can be summarized in the following four steps:

1. For query term \(x\_1\), get the set of its ancestors, denoted as \(A\_1\).
2. For query term \(x\_2\), get the set of its ancestors, denoted as \(A\_2\).
3. Get the intersection set of \(A\_1\) and \(A\_2\), denoted as \(A\_\mathrm{intersect}\).
4. Obtain the LCA term from \(A\_\mathrm{intersect}\).

This algorithm is not effcient, because many of the ancestor-offspring relations of \((a, x)\) are either repeatedly visited, or unnecessarily visited.

Let’s take the DAG in Figure S2.2 as an example to illustrate the difference between the two algorithms. In Figure S2.2, the DAG has a strict tree structure of 7 terms, and we want to find the LCA terms of \(x\_1\), \(x\_2\), \(x\_3\) and \(x\_4\).

Figure S2.2. A tiny DAG example.

Let’s count how many times term \(a\_2\) is visited by the two algorithms.

**The “bottom-up” methods used in existing tools**: The steps where \(a\_2\) is visited and how it is visited are listed as follows:

```
# the pair of x1 and x4
x1 -> a2  # obtain x1's ancestors
a2 -> a1  # obtain x1's ancestors.
a2 -> a1  # compare a2 to x4's ancestors. Here a1 is also an ancestor of x4
a2 -> a3  # compare a2 to x4's ancestors
a2 -> x4  # compare a2 to x4's ancestors

# the pair of x1 and x2
# since x1's ancestore are already obtained, we do not need to do it again
a2 -> a1  # compare a2 to x2's ancestors
a2 -> a2  # compare a2 to x2's ancestors
a2 -> x2  # compare a2 to x2's ancestors

# the pair of x1 and x3
# since x1's ancestore are already obtained, we do not need to do it again
a2 -> a1  # compare a2 to x3's ancestors
a2 -> a3  # compare a2 to x3's ancestors
a2 -> x3  # compare a2 to x3's ancestors

# the pair of x2 and x3
x2 -> a2  # obtain x2's ancestors
a2 -> a1  # obtain x2's ancestors
a2 -> a1  # compare a2 to x3's ancestors. Here a1 is also an ancestor of x3
a2 -> a3  # compare a2 to x3's ancestors
a2 -> x3  # compare a2 to x3's ancestors

# the pair of x2 and x4
# since x2's ancestore are already obtained, we do not need to do it again
a2 -> a1  # compare a2 to x4's ancestors
a2 -> a3  # compare a2 to x4's ancestors
a2 -> x3  # compare a2 to x4's ancestors
```

The total number when \(a\_2\) is visited is 19.

**The “top-down” methods by simona**:

```
# get all ancestors of x1, x2, x3 and x4
x1 -> a2
a2 -> a1
x2 -> a2  # because a2 is already visited from x1, the traversal from x2 stops here

# traverse down from a1
a1 -> a2
a2 -> x1
a2 -> x2

# traverse down from a2
a2 -> x1
a2 -> x2

# obtain the tuple (a2, x_i, x_j)
a2 -> x1
a2 -> x2
```

Now \(a\_2\) is only visited 10 times.

Let’s make a comparison of how \(a\_2\) is visited by the two algorithms. In the following table, numbers represent the times of each traversal step.

| traversal | bottom-up | top-down (simona) |
| --- | --- | --- |
| a2 -> a1 | 7 | 1 |
| a1 -> a2 | 0 | 1 |
| a2 -> a2 | 1 | 0 |
| a2 -> a3 | 4 | 0 |
| a2 -> x1 | 0 | 3 |
| a2 -> x2 | 1 | 3 |
| a2 -> x3 | 3 | 0 |
| a2 -> x4 | 1 | 0 |
| x1 -> a2 | 1 | 1 |
| x2 -> a2 | 1 | 1 |

From the table above, we can see in the “bottom-up” method, the traversal `a2 -> a1` is performed 7 times. It is visited at least once when obtaining ancestor terms of every query term. This traversal is repeated unnecessarily, because \(a\_1\) is the root and it is a common ancestor of every pair of the query terms, with no need to test to other terms in the DAG again and again. Another type of unnecessary comparison is between \(a\_2\) and \(a\_3\) which is visited 4 times by the bottom-up method. \(a\_2\) and \(a\_3\) are not connected and there is no need to compare the two terms.

There are also traversals in the “top-down” methods that are more visited than the “bottom-up” method, such as `a2 -> x1`. This link locates quite low in the DAG, and it is visited from every of \(a\_2\)’s ancestors in the DAG. Nevertheless, the total run time is still effcient for the “top-down” method.

#### Time complexity

The top-down algorithm by **simona** has a time complexity approximate to \(O(n^2)\). We can emperically validated it by two DAGs. The first one `binary_tree` is a tree where each term has two children and all leaf terms have the same depth of 11.

```
library(simona)
binary_tree = dag_random_tree(n_children = 2, max = 2^12 - 1)
binary_tree
```

```
## An ontology_DAG object:
##   Source: dag_random_tree 
##   4095 terms / 4094 relations / a tree
##   Root: 1 
##   Terms: 1, 10, 100, 1000, ...
##   Max depth: 11 
##   Aspect ratio: 186.18:1
```

The second DAG is based on `binary_tree` where each term has a probability of 0.6 to add other 2 ~ 10 terms as its new child terms that are lower than the term.

```
set.seed(123)
random_dag = dag_add_random_children(binary_tree, p_add = 0.6, new_children = c(2, 10))
random_dag
```

```
## An ontology_DAG object:
##   Source: dag_add_random_children 
##   4095 terms / 11618 relations
##   Root: 1 
##   Terms: 1, 10, 100, 1000, ...
##   Max depth: 11 
##   Avg number of parents: 2.84
##   Avg number of children: 1.73
##   Aspect ratio: 186.18:1 (based on the longest distance from root)
##                 79.64:1 (based on the shortest distance from root)
```

In `binary_tree`, average number of parents is 1 (exclude the root), and in `random_dag`, the average number of parents is higher, which means it is more densely connected in `random_dag`:

```
mean(n_parents(random_dag))
```

```
## [1] 2.837118
```

Let’s visualize these two DAGs:

```
library(grid)
pushViewport(viewport(x = 0.25, width = 0.5))
dag_circular_viz(binary_tree, newpage = FALSE)
popViewport()

pushViewport(viewport(x = 0.75, width = 0.5))
dag_circular_viz(random_dag, edge_transparency = 0.96, newpage = FALSE)
popViewport()
```

Figure S2.3. Visualization of the binary tree and a random dense DAG.

And we compare the runtime performance on the two DAGs.

```
benchmark_runtime = function(dag, by = 200, max = 10000) {
    invisible(dag_depth(dag))  # depth will be cached

    n_terms = dag_n_terms(dag)
    k = seq(by, min(max, floor(n_terms/by)*by), by = by)
    t = rep(NA_real_, length(k))
    for(i in seq_along(k)) {
        terms = sample(n_terms, k[i]) # numeric indicies are also allowed
        t[i] = system.time(term_sim(dag, terms, method = "Sim_WP_1994"))[3]
    }
    data.frame(k = k, t = t)
}

lt = list()
lt[[1]] = benchmark_runtime(binary_tree, by = 100)
lt[[2]] = benchmark_runtime(random_dag, by = 100)
```

```
par(mfrow = c(1, 2))
plot(NULL, xlim = c(0, 4100), ylim = c(0, max(lt[[1]]$t, lt[[2]]$t)),
    xlab = "Numbers of random terms", ylab = "runtime (sec)")
for(i in seq_along(lt)) {
    x = lt[[i]]$k
    y = lt[[i]]$t
    lines(x, y, col = i + 1)
}
legend("topleft", lty = 1, col = 2:3, legend = c("binary_tree", "random_dag"))

plot(NULL, xlim = c(0, 1), ylim = c(0, 1),
    xlab = "Numbers of random terms, scaled", ylab = "runtime, scaled")
for(i in seq_along(lt)) {
    x = lt[[i]]$k
    y = lt[[i]]$t
    x = x/max(x)
    y = y/max(y)
    lines(x, y, col = i + 1)
}
curve(x^1, from = 0, to = 1, lty = 2, add = TRUE)
curve(x^2, from = 0, to = 1, lty = 2, add = TRUE)
legend("topleft", lty = 1, col = 2:3, legend = c("binary_tree", "random_dag"))
```

Figure S2.4. Runtime performance on the binary tree and the random dense DAG. Left: absolute time complexity. Right: relative time complexity.

Figure S2.4 shows that when the DAG has a more complex structure, i.e. a term has more than one parent, the scaling factor of the time complexity increases.

#### Session info

```
sessionInfo()
```

```
## R version 4.4.1 (2024-06-14)
## Platform: x86_64-apple-darwin20
## Running under: macOS Sonoma 14.5
## 
## Matrix products: default
## BLAS:   /Library/Frameworks/R.framework/Versions/4.4-x86_64/Resources/lib/libRblas.0.dylib 
## LAPACK: /Library/Frameworks/R.framework/Versions/4.4-x86_64/Resources/lib/libRlapack.dylib;  LAPACK version 3.12.0
## 
## locale:
## [1] en_GB.UTF-8/en_GB.UTF-8/en_GB.UTF-8/C/en_GB.UTF-8/en_GB.UTF-8
## 
## time zone: Europe/Berlin
## tzcode source: internal
## 
## attached base packages:
## [1] grid      stats     graphics  grDevices utils     datasets  methods  
## [8] base     
## 
## other attached packages:
## [1] simona_1.3.12  knitr_1.48     rmarkdown_2.28
## 
## loaded via a namespace (and not attached):
##  [1] sass_0.4.9            xml2_1.3.6            shape_1.4.6.1        
##  [4] digest_0.6.37         magrittr_2.0.3        evaluate_0.24.0      
##  [7] RColorBrewer_1.1-3    iterators_1.0.14      circlize_0.4.16      
## [10] fastmap_1.2.0         foreach_1.5.2         doParallel_1.0.17    
## [13] jsonlite_1.8.8        GlobalOptions_0.1.2   promises_1.3.0       
## [16] ComplexHeatmap_2.20.0 codetools_0.2-20      jquerylib_0.1.4      
## [19] cli_3.6.3             shiny_1.9.1           rlang_1.1.4          
## [22] crayon_1.5.3          scatterplot3d_0.3-44  cachem_1.1.0         
## [25] yaml_2.3.10           tools_4.4.1           parallel_4.4.1       
## [28] colorspace_2.1-1      httpuv_1.6.15         GetoptLong_1.0.5     
## [31] BiocGenerics_0.50.0   R6_2.5.1              mime_0.12            
## [34] png_0.1-8             matrixStats_1.3.0     stats4_4.4.1         
## [37] lifecycle_1.0.4       S4Vectors_0.42.1      IRanges_2.38.1       
## [40] clue_0.3-65           cluster_2.1.6         pkgconfig_2.0.3      
## [43] bslib_0.8.0           later_1.3.2           Rcpp_1.0.13          
## [46] xfun_0.47             highr_0.11            xtable_1.8-4         
## [49] rjson_0.2.22          htmltools_0.5.8.1     igraph_2.0.3         
## [52] compiler_4.4.1        Polychrome_1.5.1
```
