## Supplementary figures and images for "*simona:* a comprehensive R package for semantic similarity analysis on bio-ontologies"

### f1.png

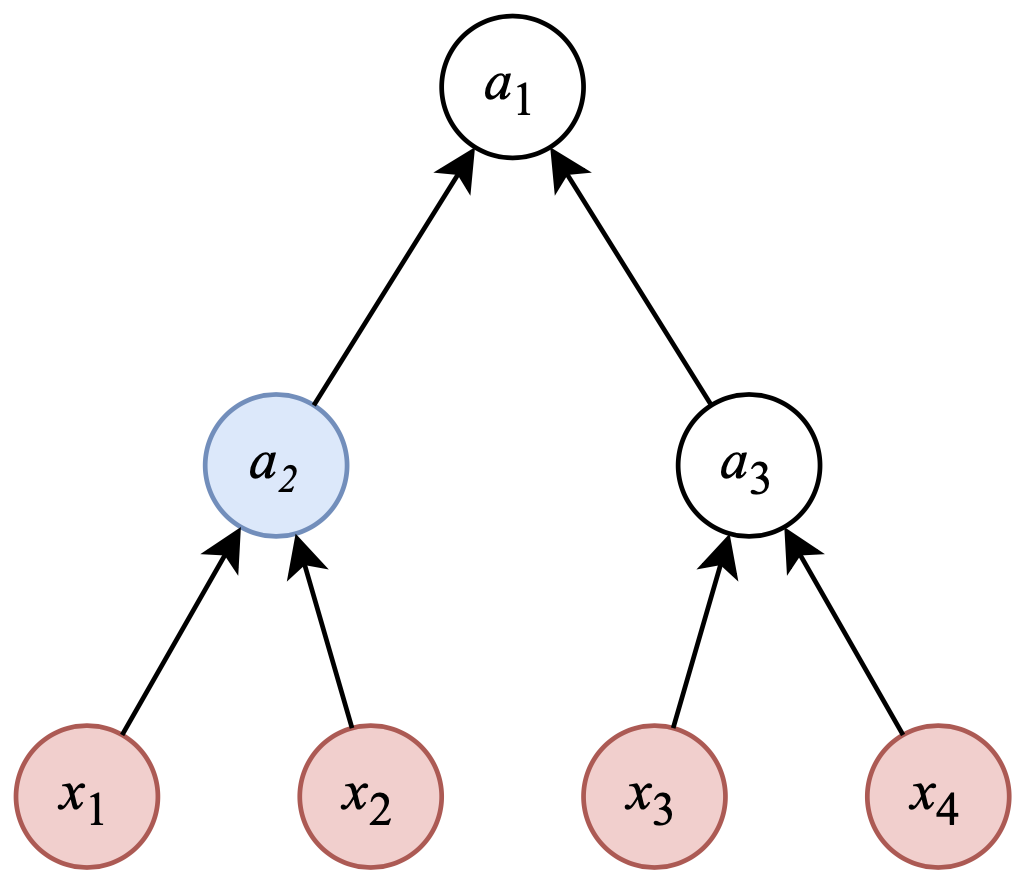

### f2.png

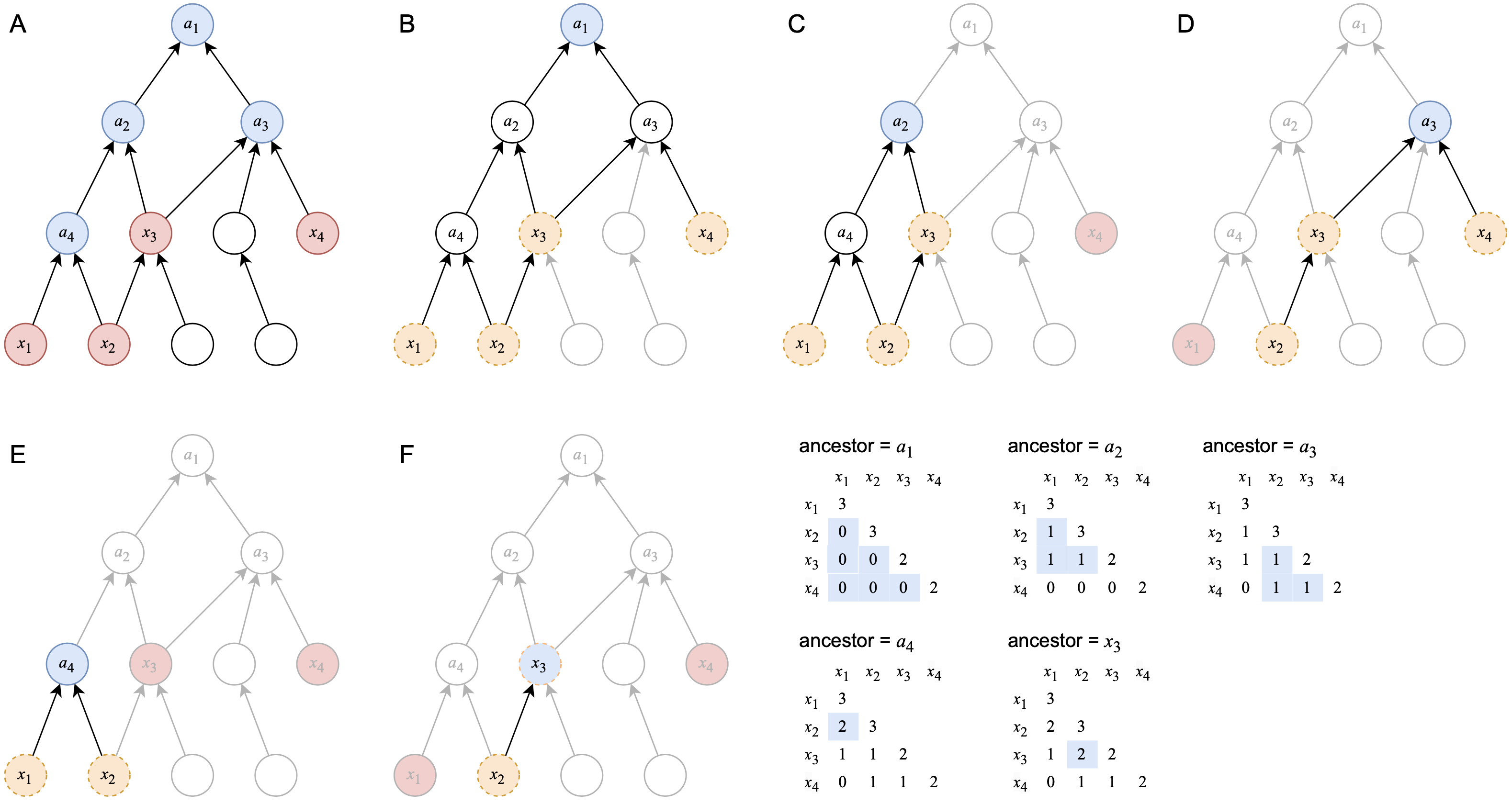

### go_bp_random_500_sim_Sim_AIC_2014.png

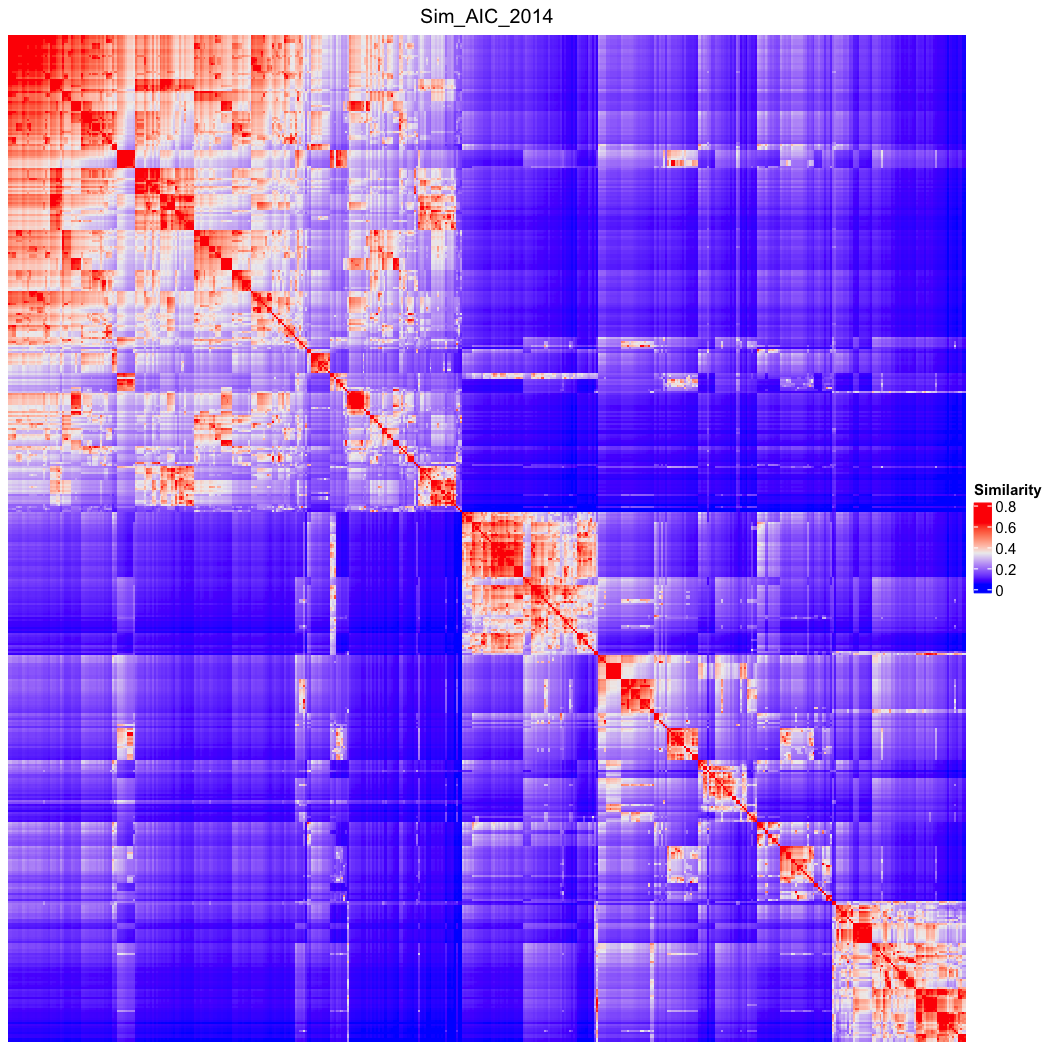

### go_bp_random_500_sim_Sim_AIC_2014_Lin_order.png

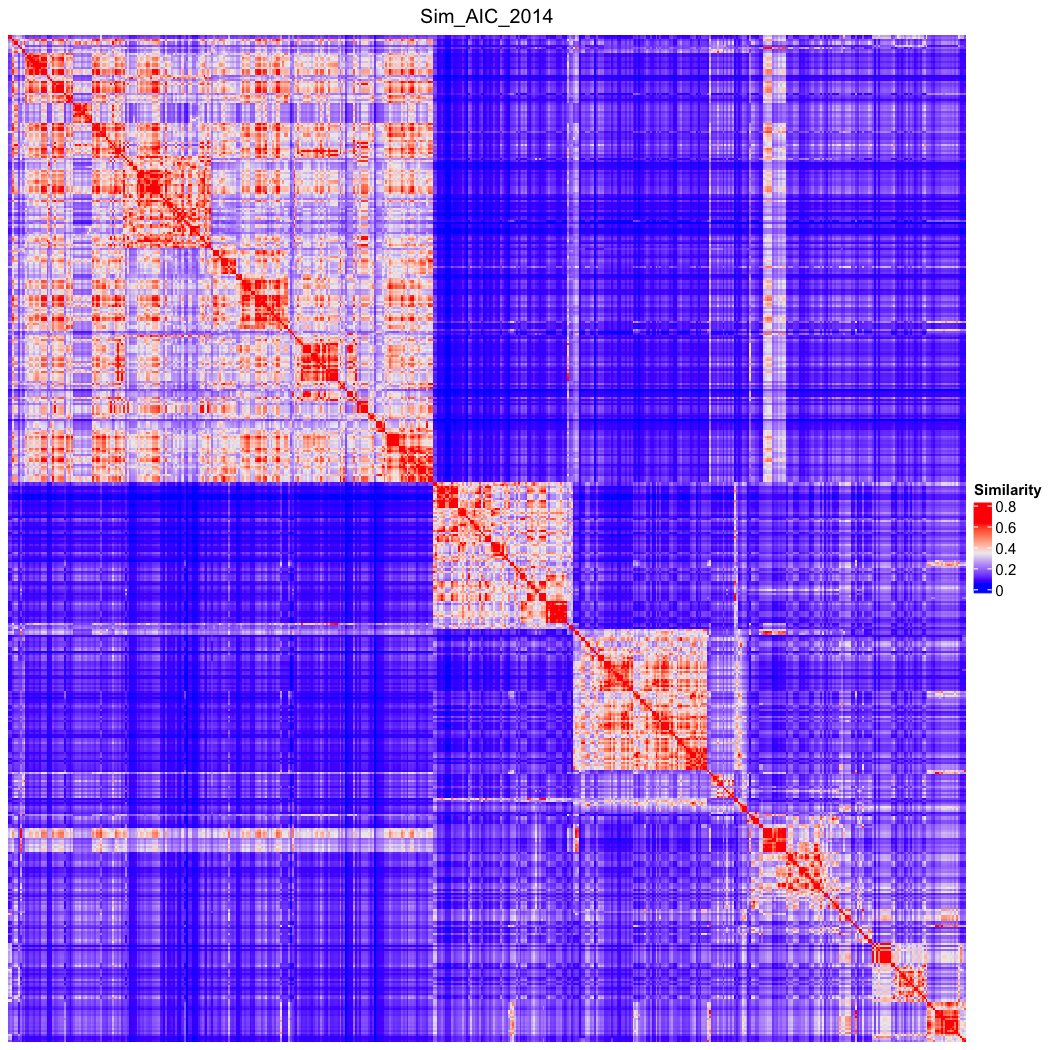

### go_bp_random_500_sim_Sim_AlMubaid_2006.png

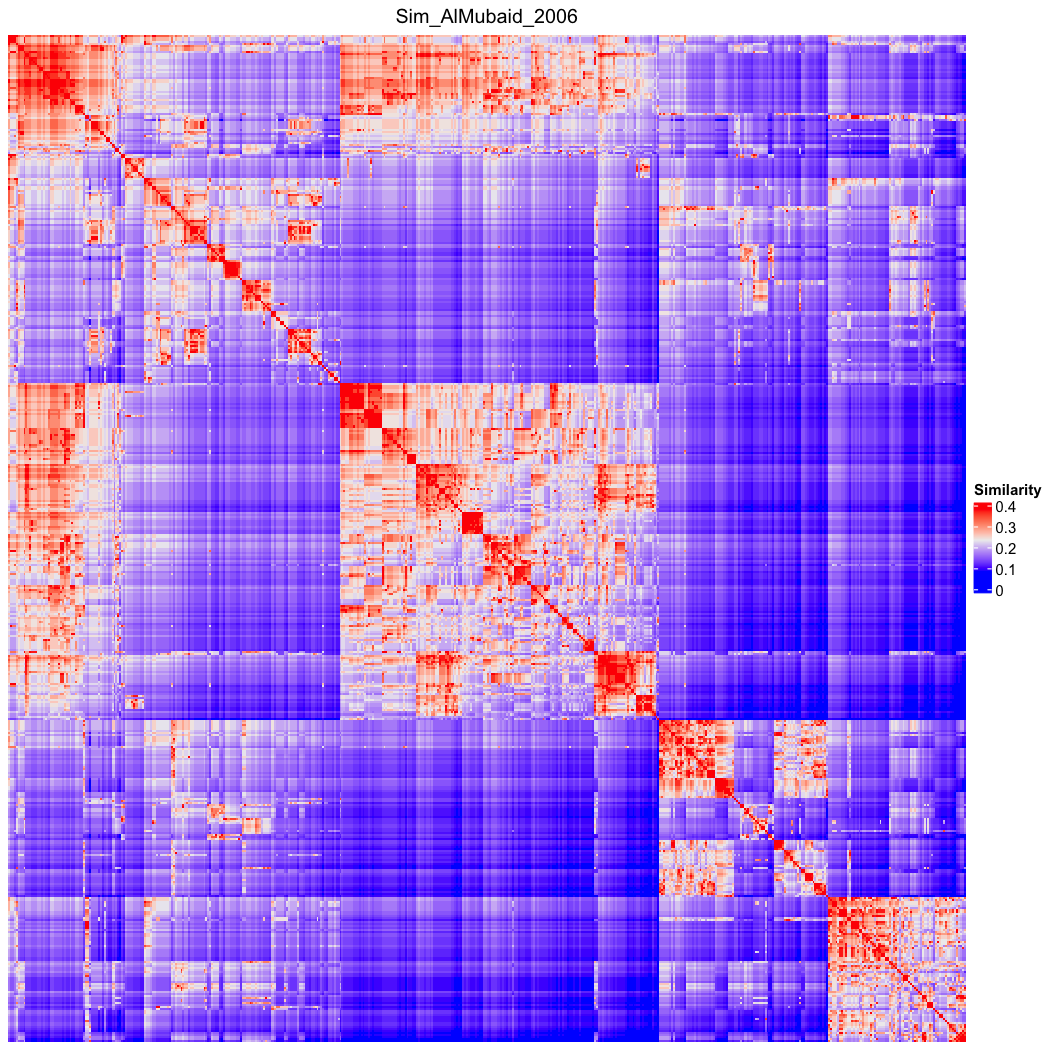

### go_bp_random_500_sim_Sim_AlMubaid_2006_Lin_order.png

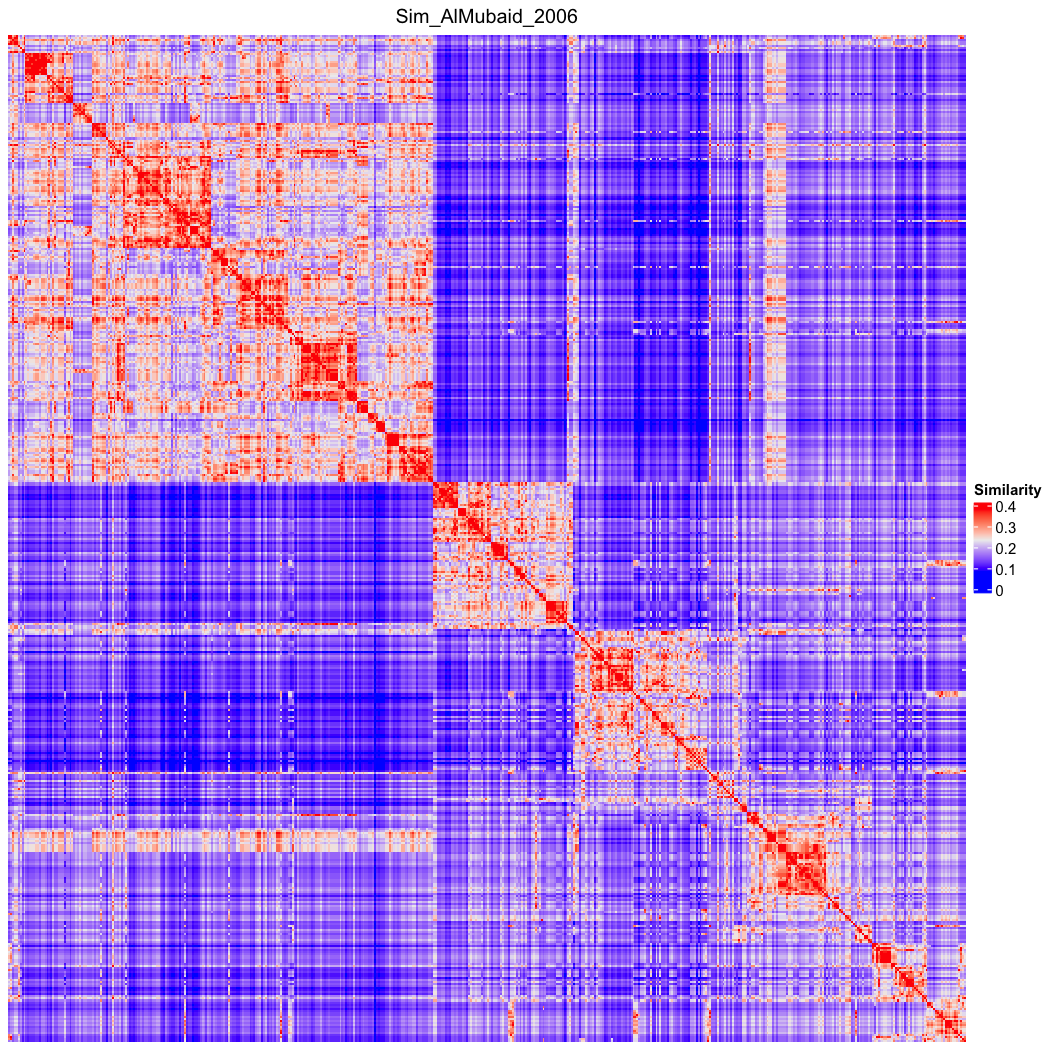

### go_bp_random_500_sim_Sim_Ancestor.png

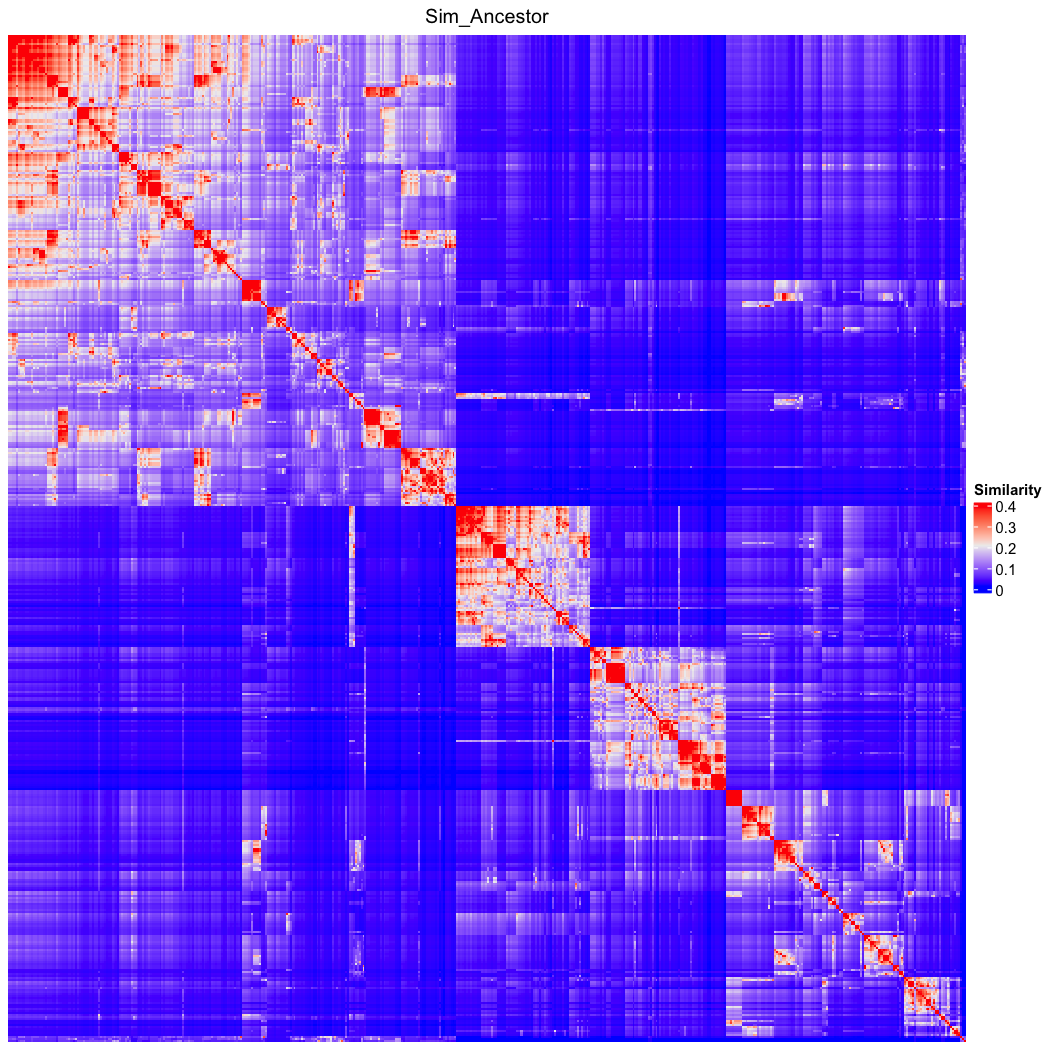

### go_bp_random_500_sim_Sim_Ancestor_Lin_order.png

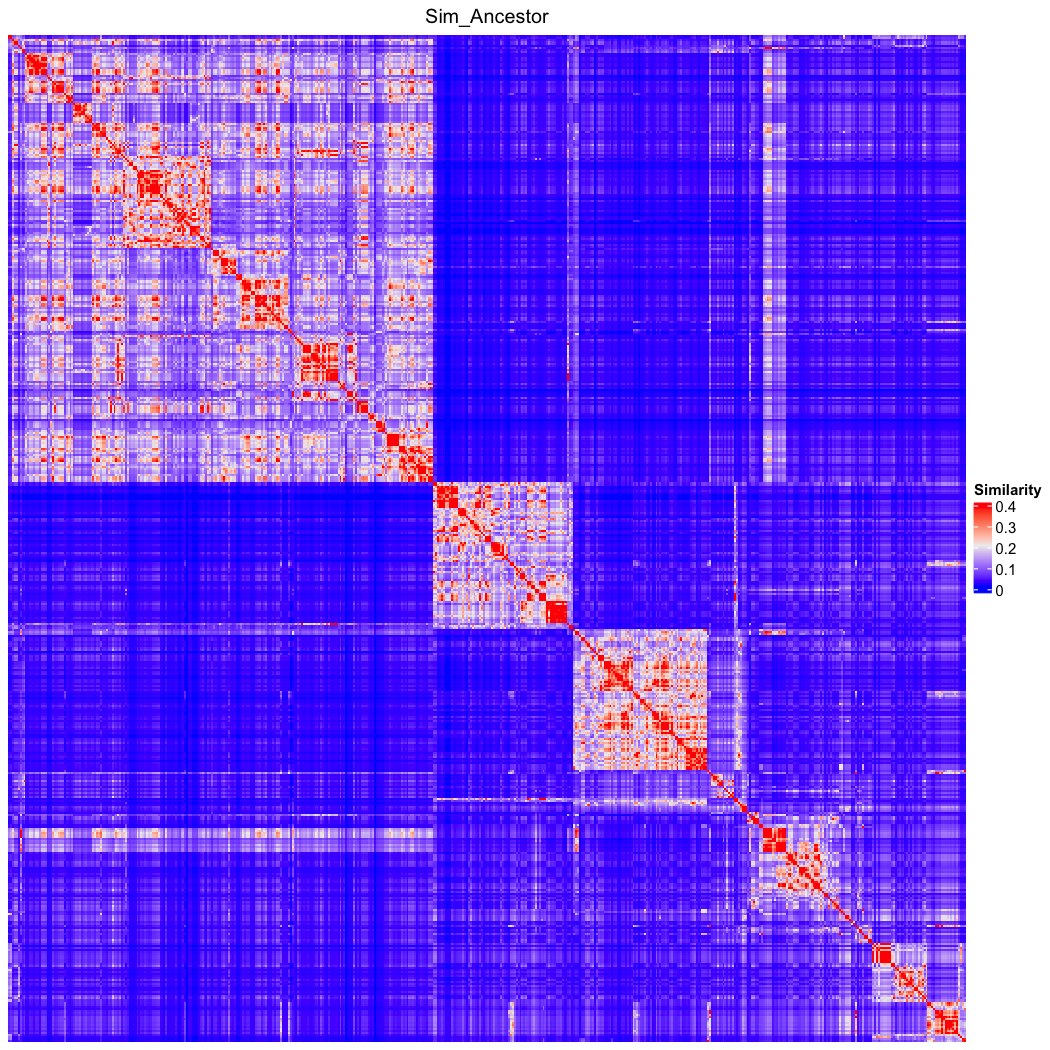

### go_bp_random_500_sim_Sim_Dice.png

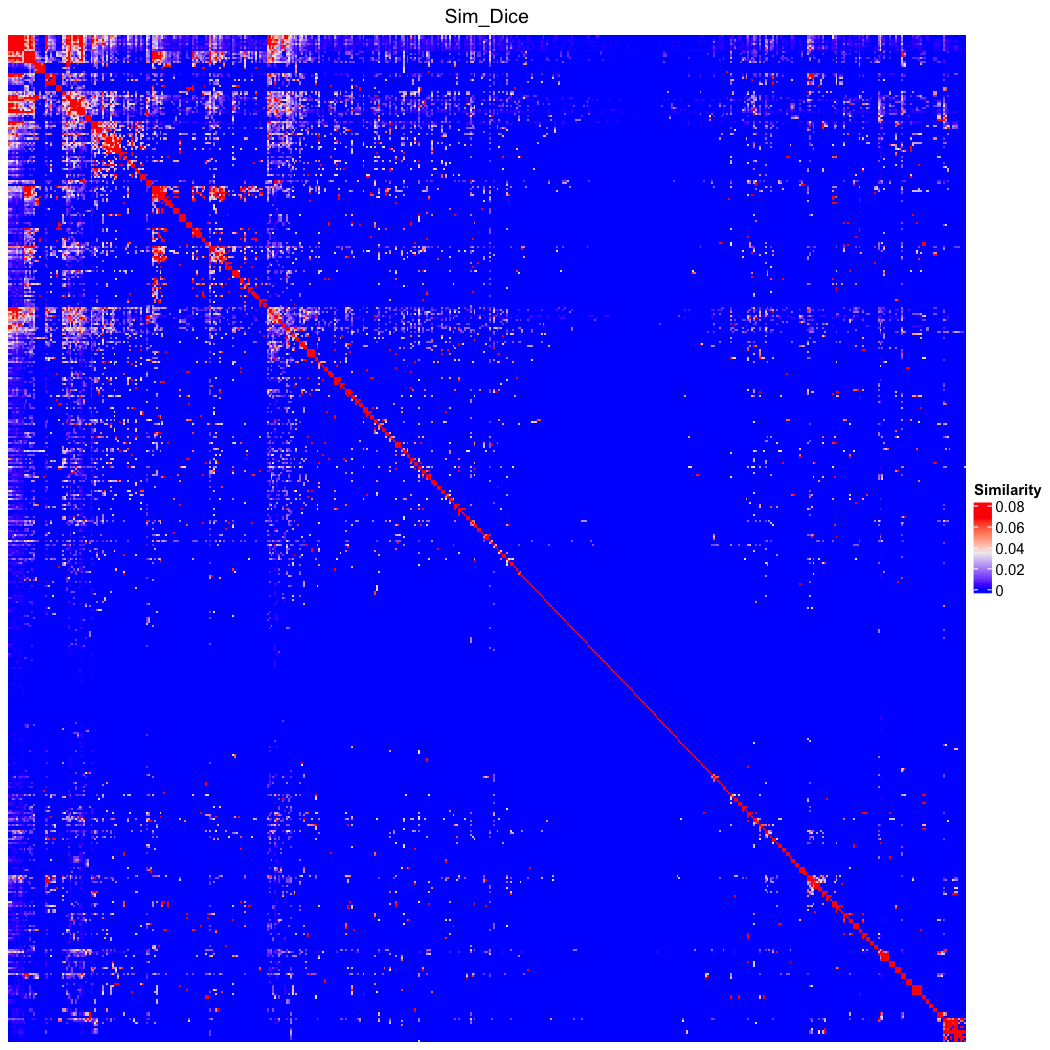

### go_bp_random_500_sim_Sim_Dice_Lin_order.png

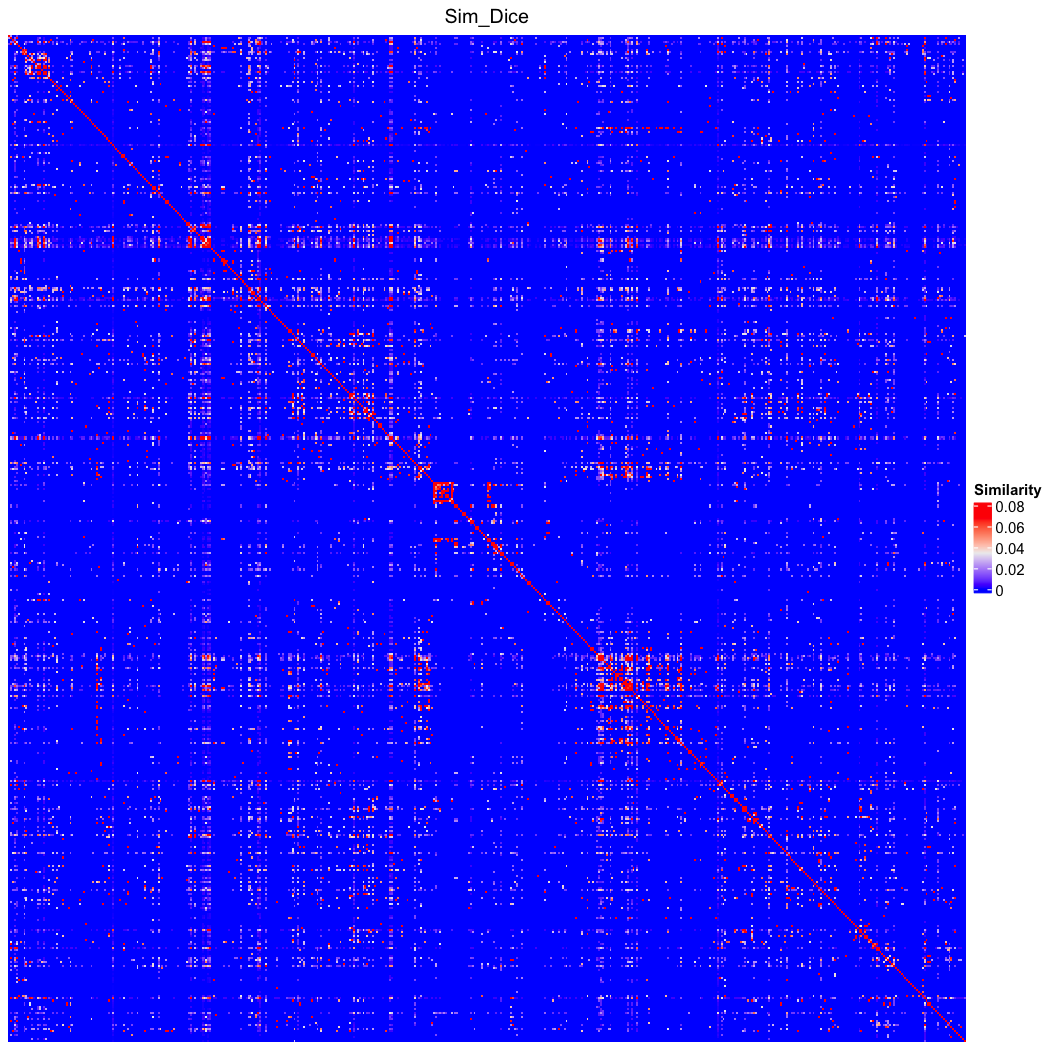

### go_bp_random_500_sim_Sim_EISI_2015.png

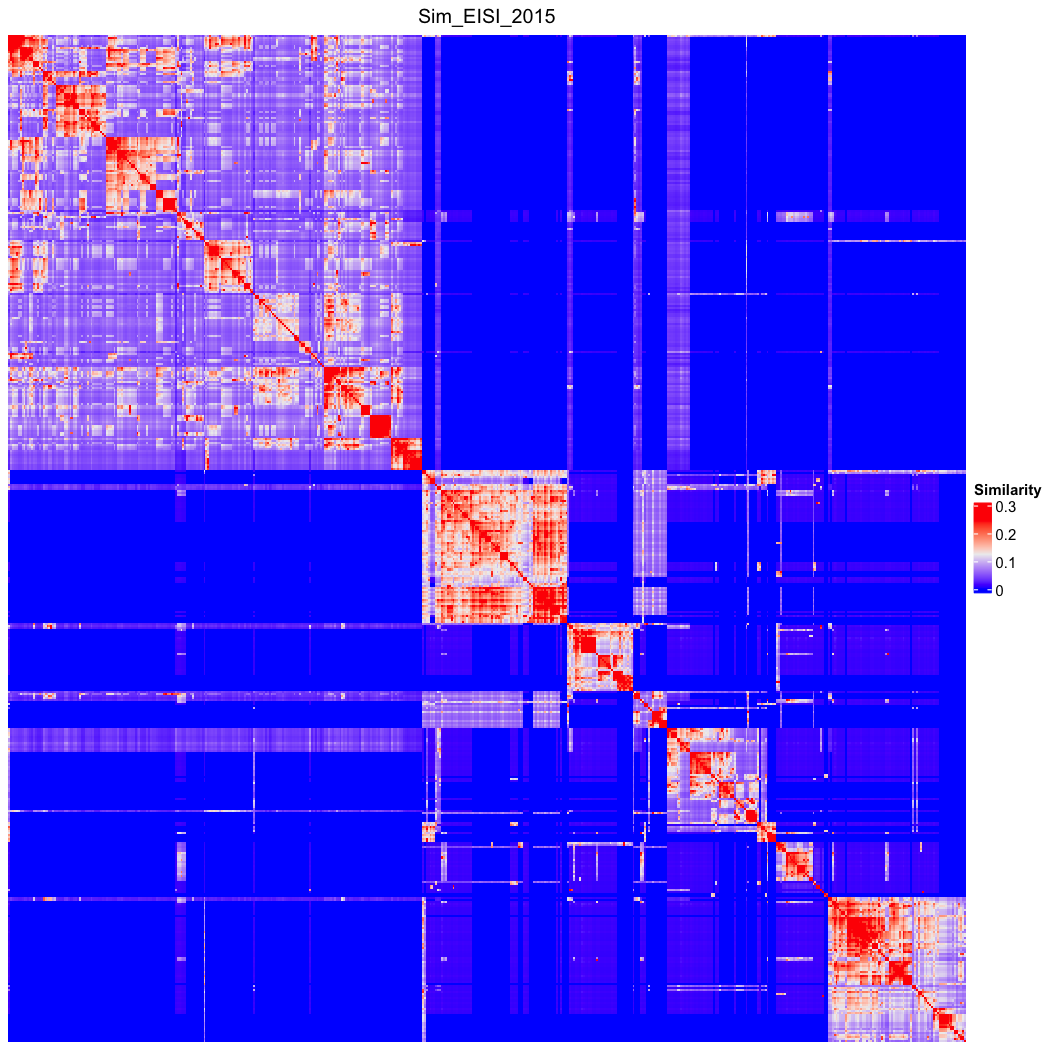

### go_bp_random_500_sim_Sim_EISI_2015_Lin_order.png

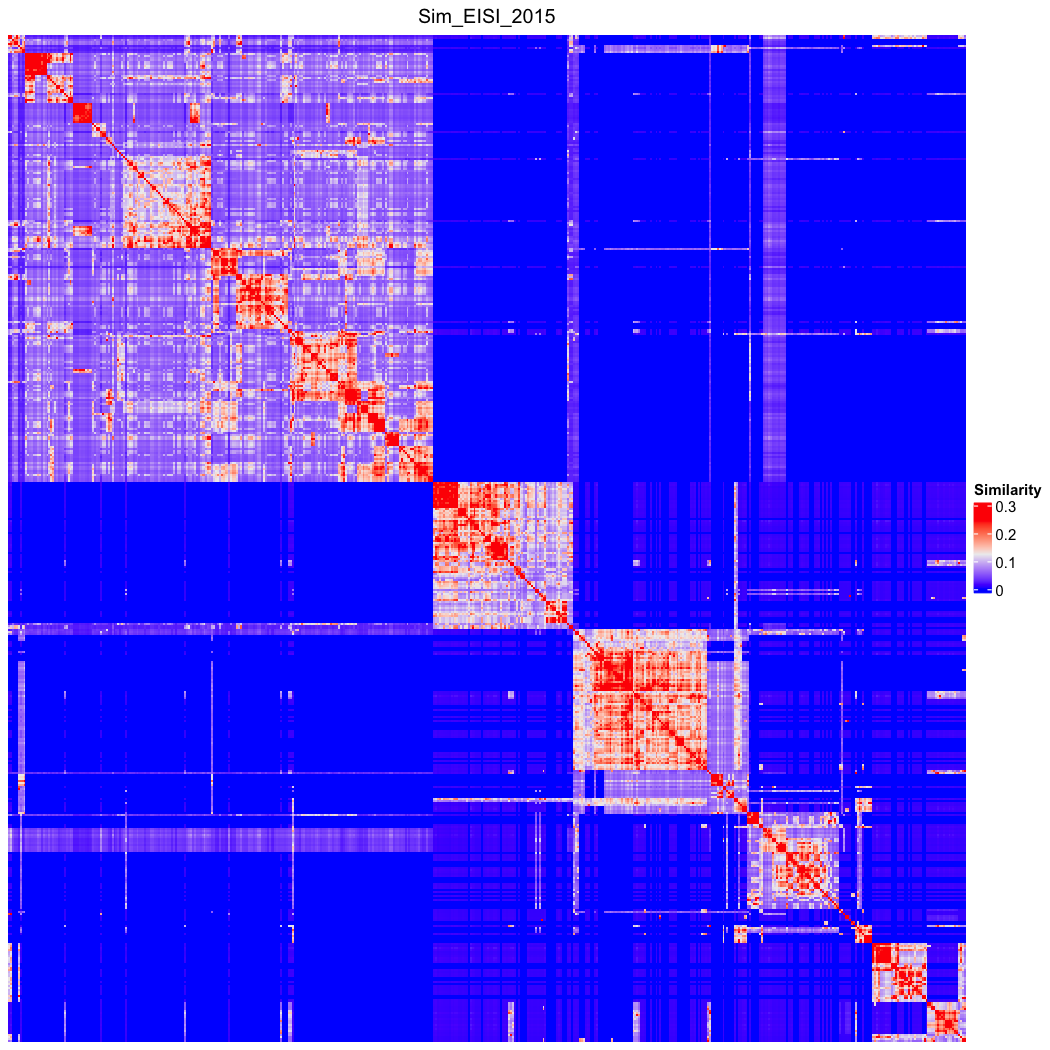

### go_bp_random_500_sim_Sim_FaITH_2010.png

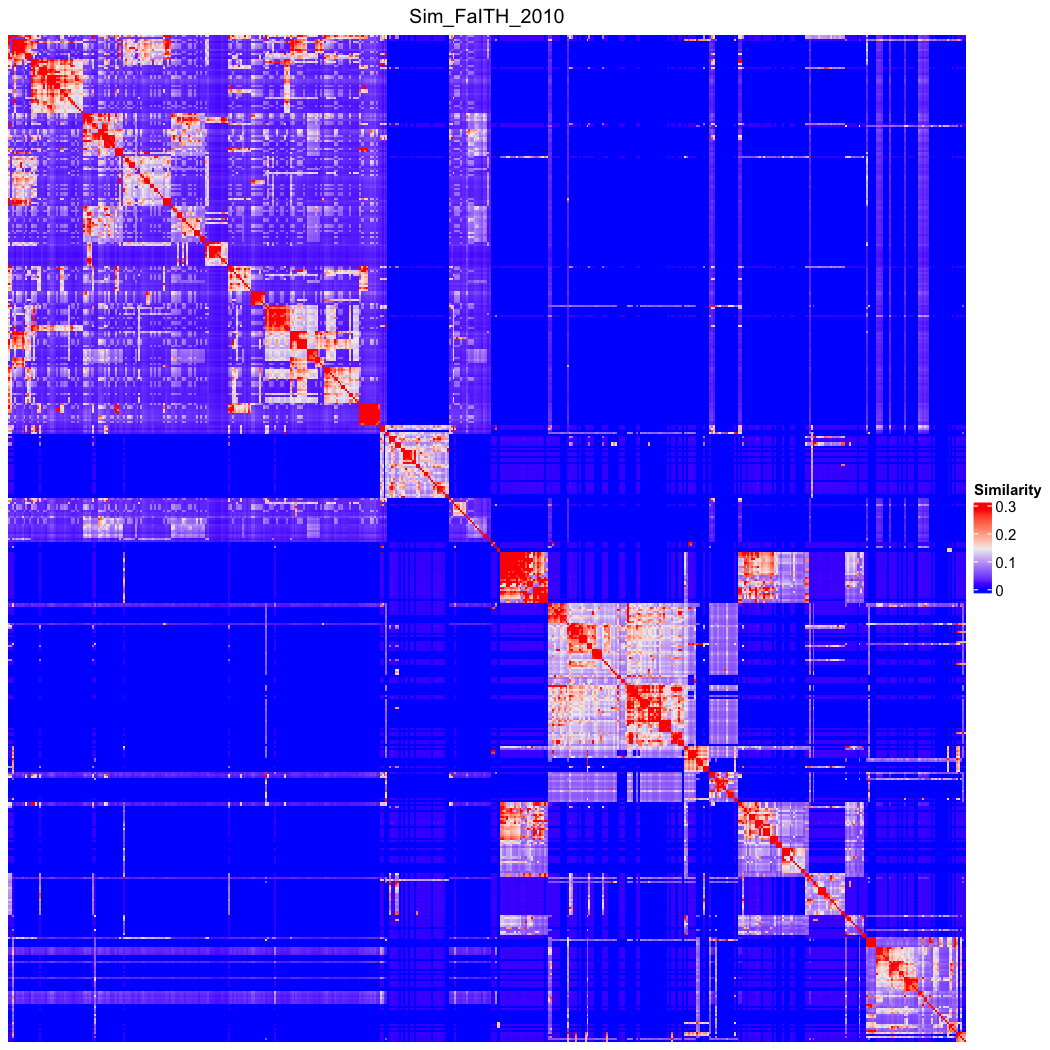

### go_bp_random_500_sim_Sim_FaITH_2010_Lin_order.png

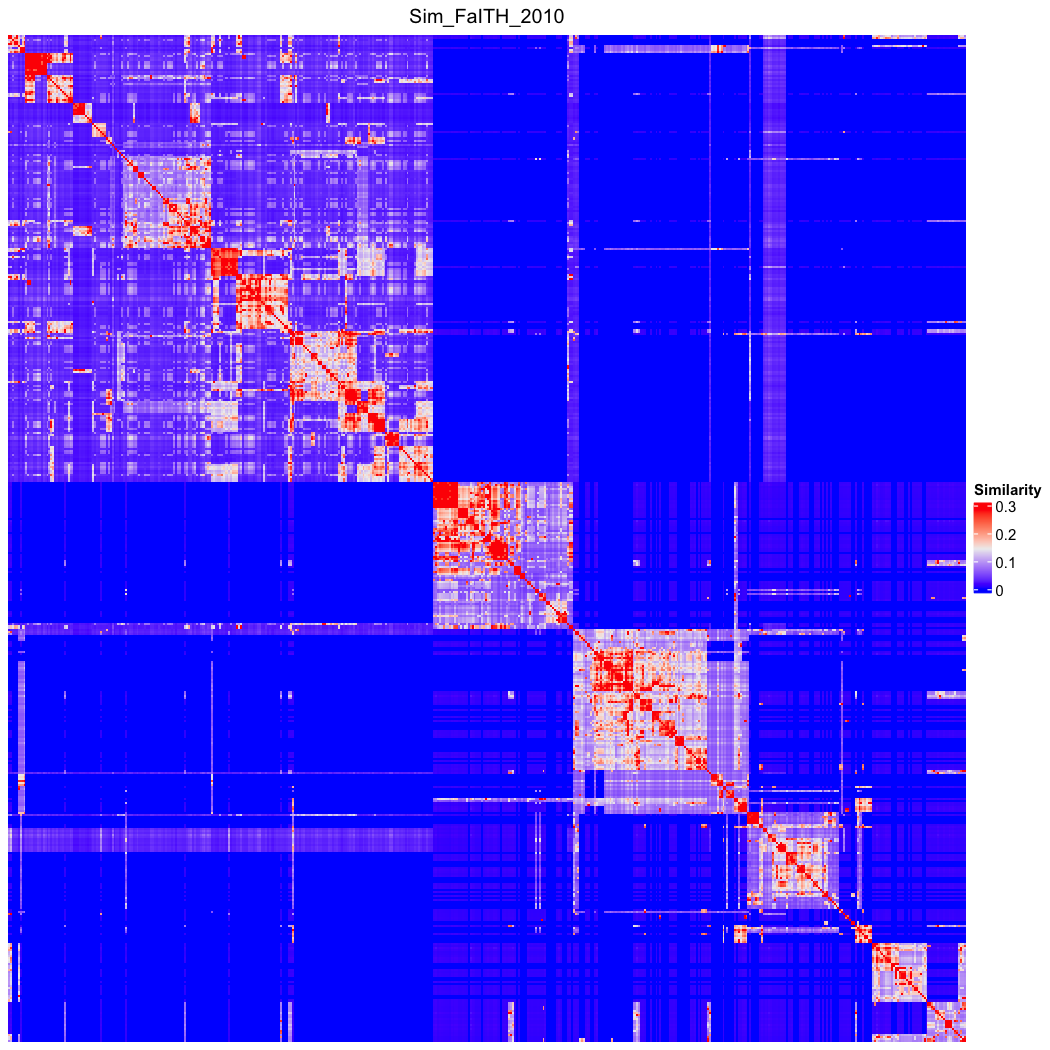

### go_bp_random_500_sim_Sim_GOGO_2018.png

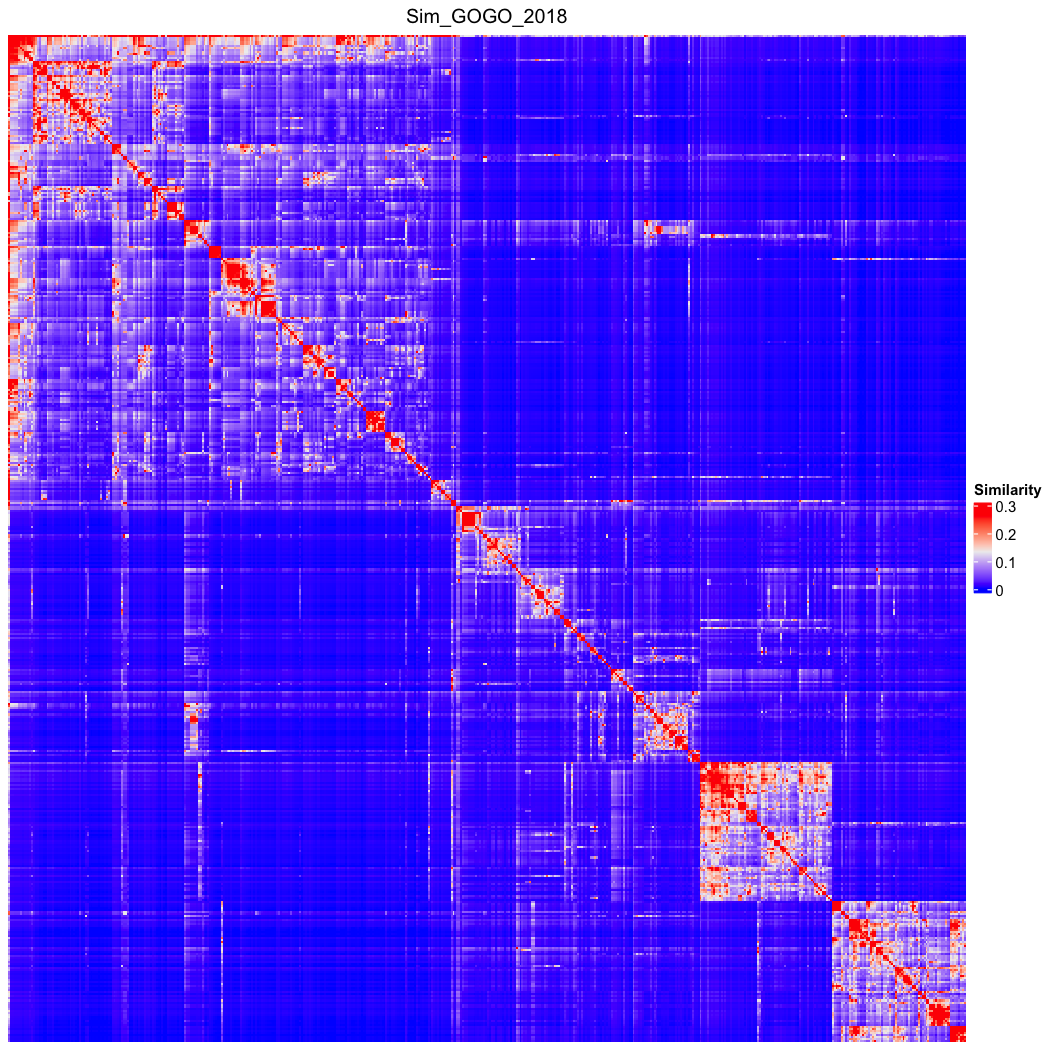

### go_bp_random_500_sim_Sim_GOGO_2018_Lin_order.png

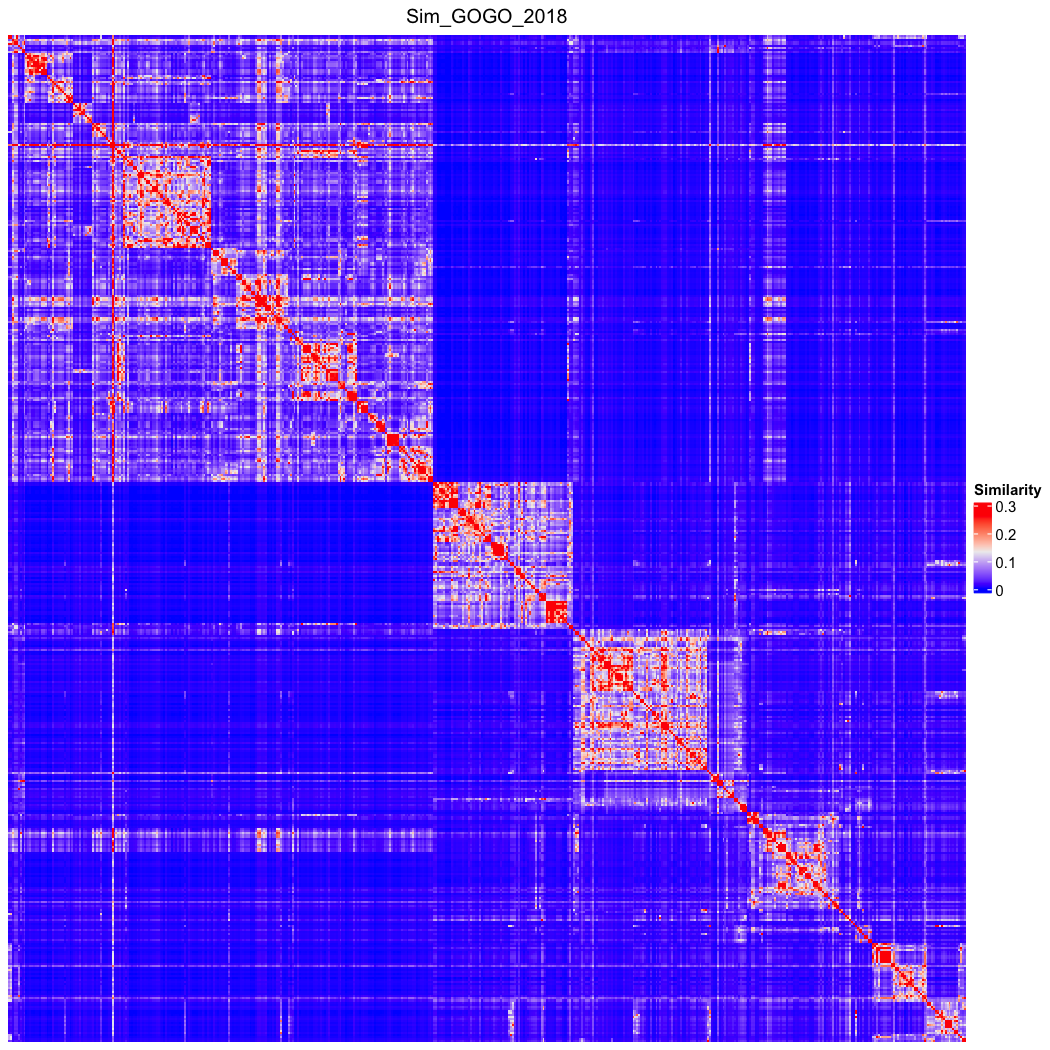

### go_bp_random_500_sim_Sim_HRSS_2013.png

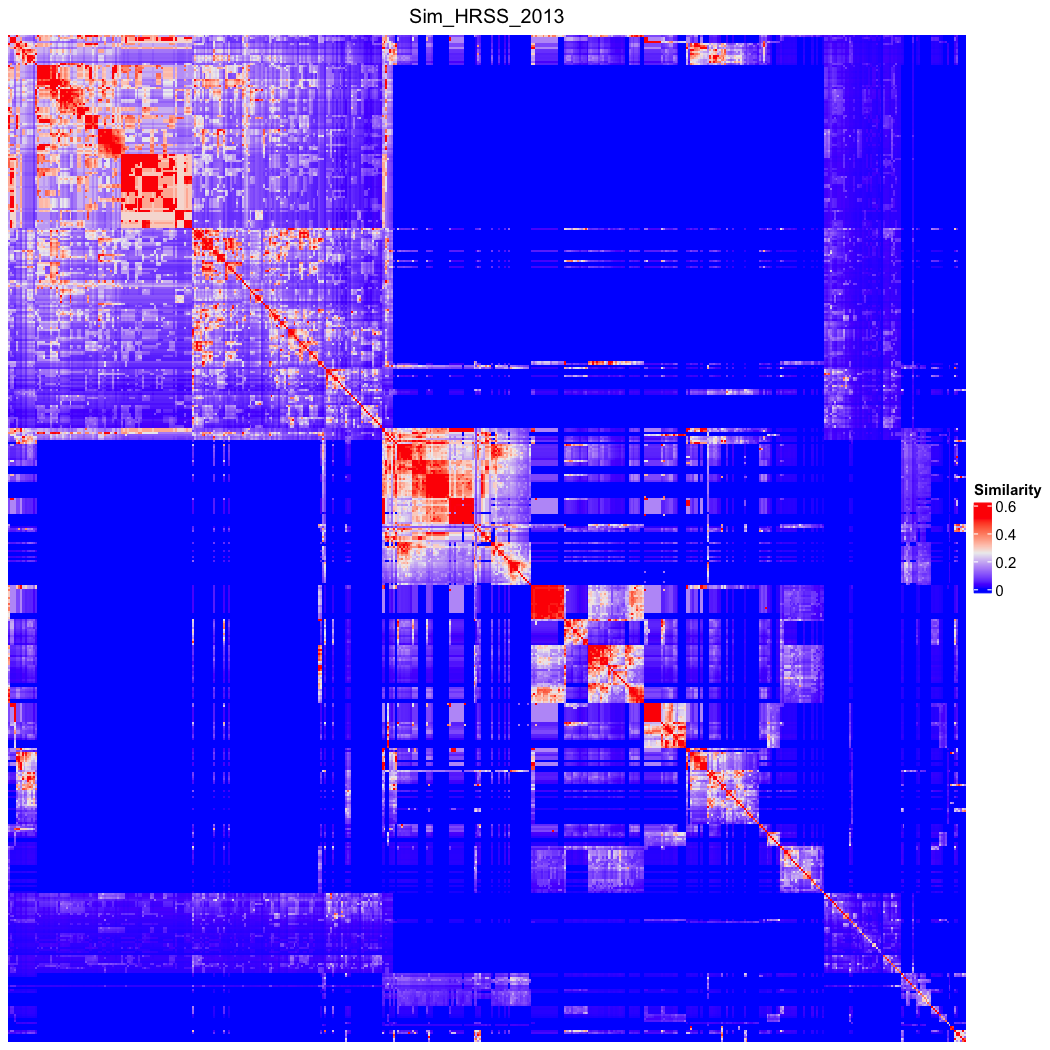

### go_bp_random_500_sim_Sim_HRSS_2013_Lin_order.png

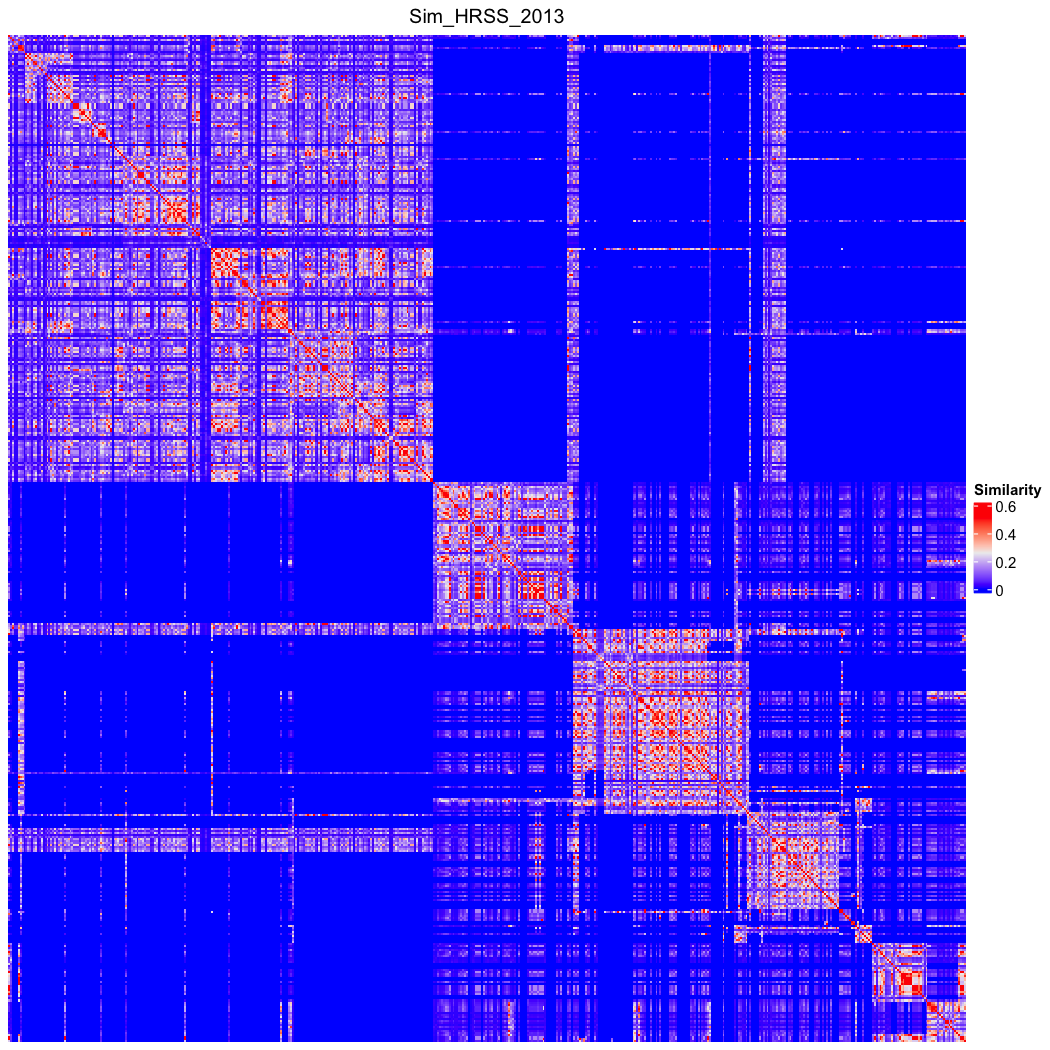

### go_bp_random_500_sim_Sim_Jaccard.png

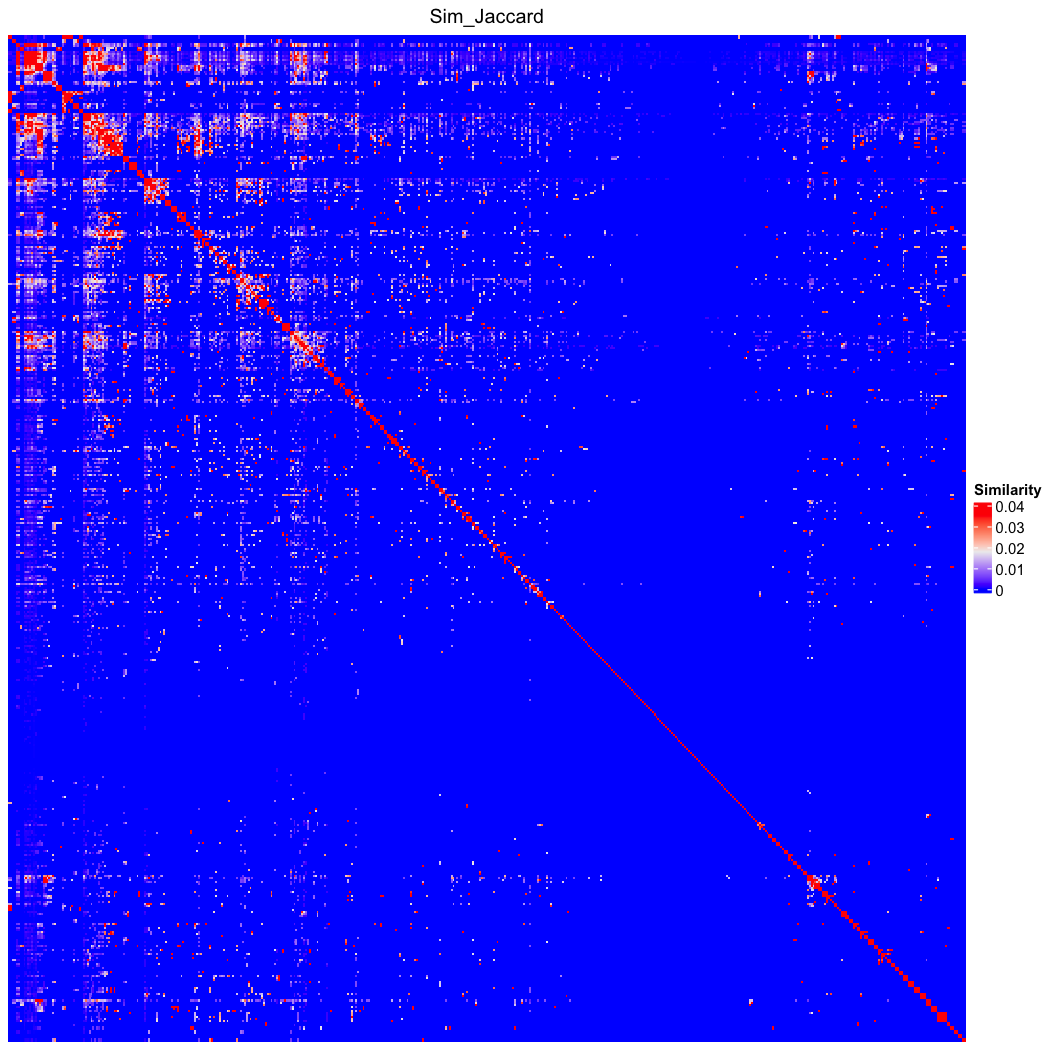

### go_bp_random_500_sim_Sim_Jaccard_Lin_order.png

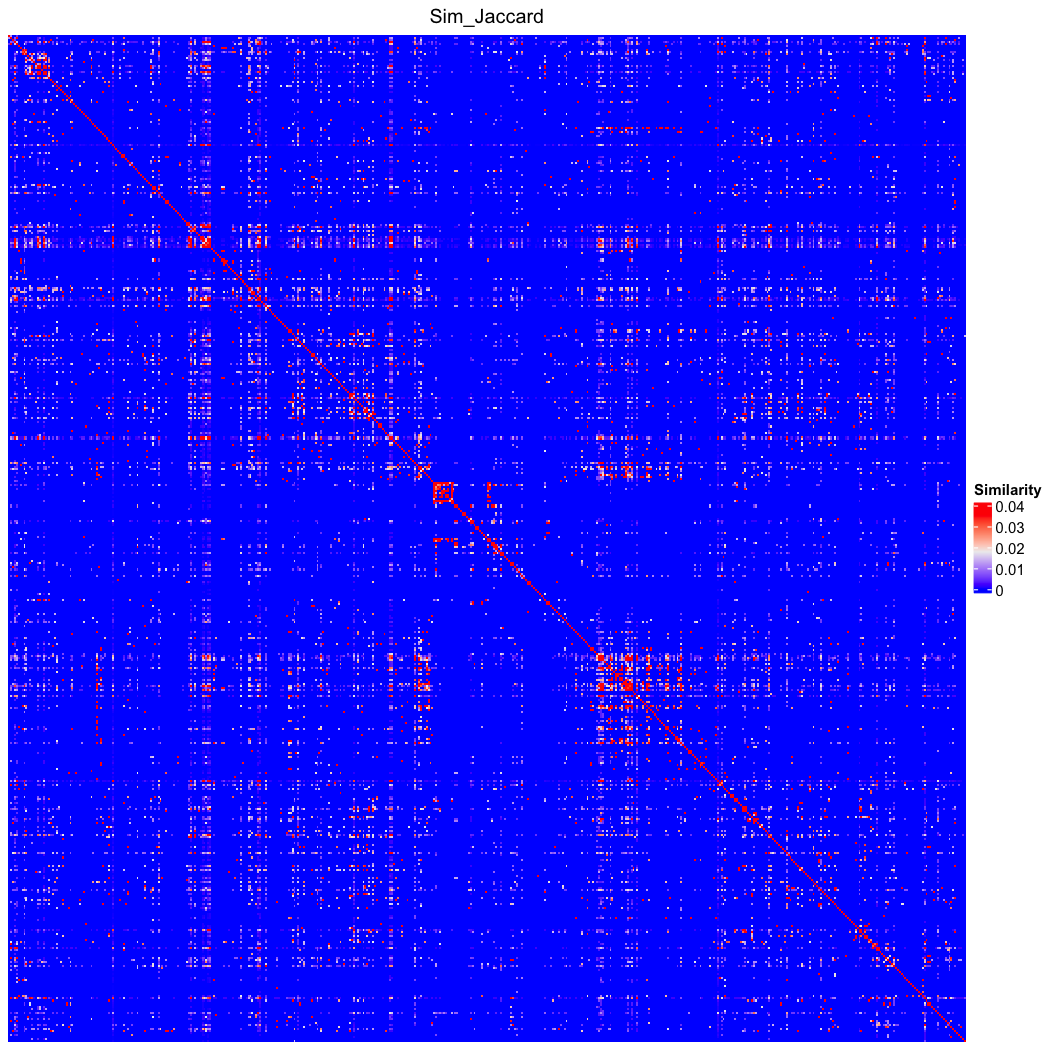

### go_bp_random_500_sim_Sim_Jiang_1997.png

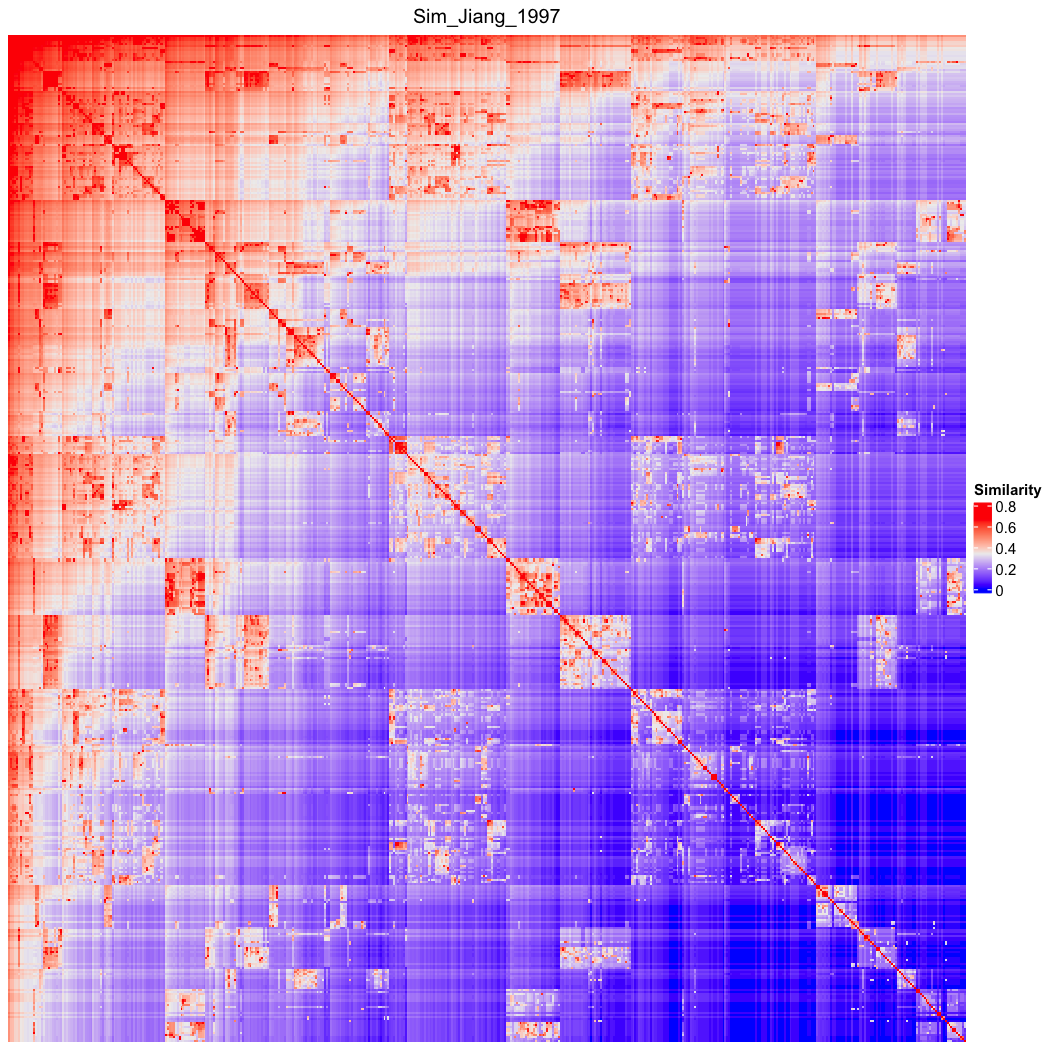

### go_bp_random_500_sim_Sim_Jiang_1997_Lin_order.png

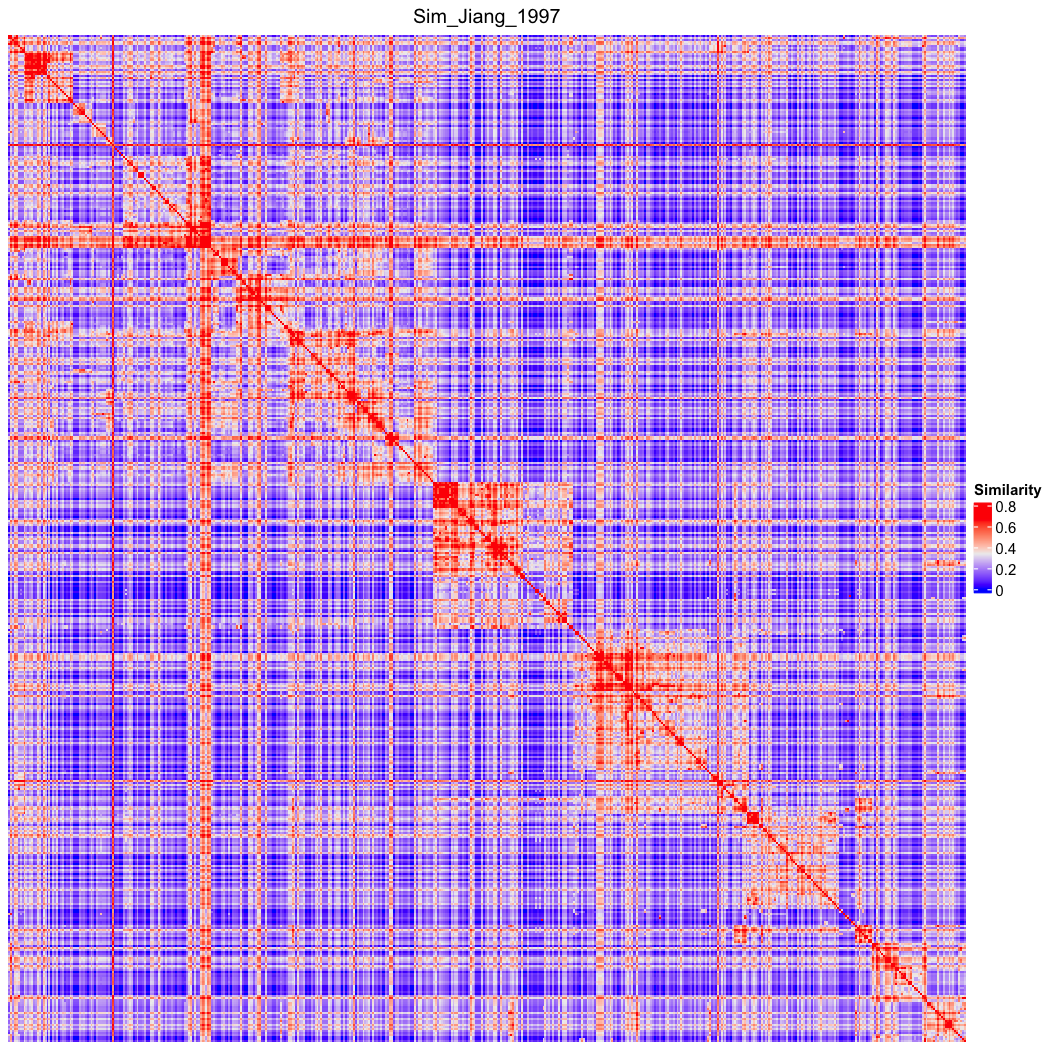

### go_bp_random_500_sim_Sim_Kappa.png

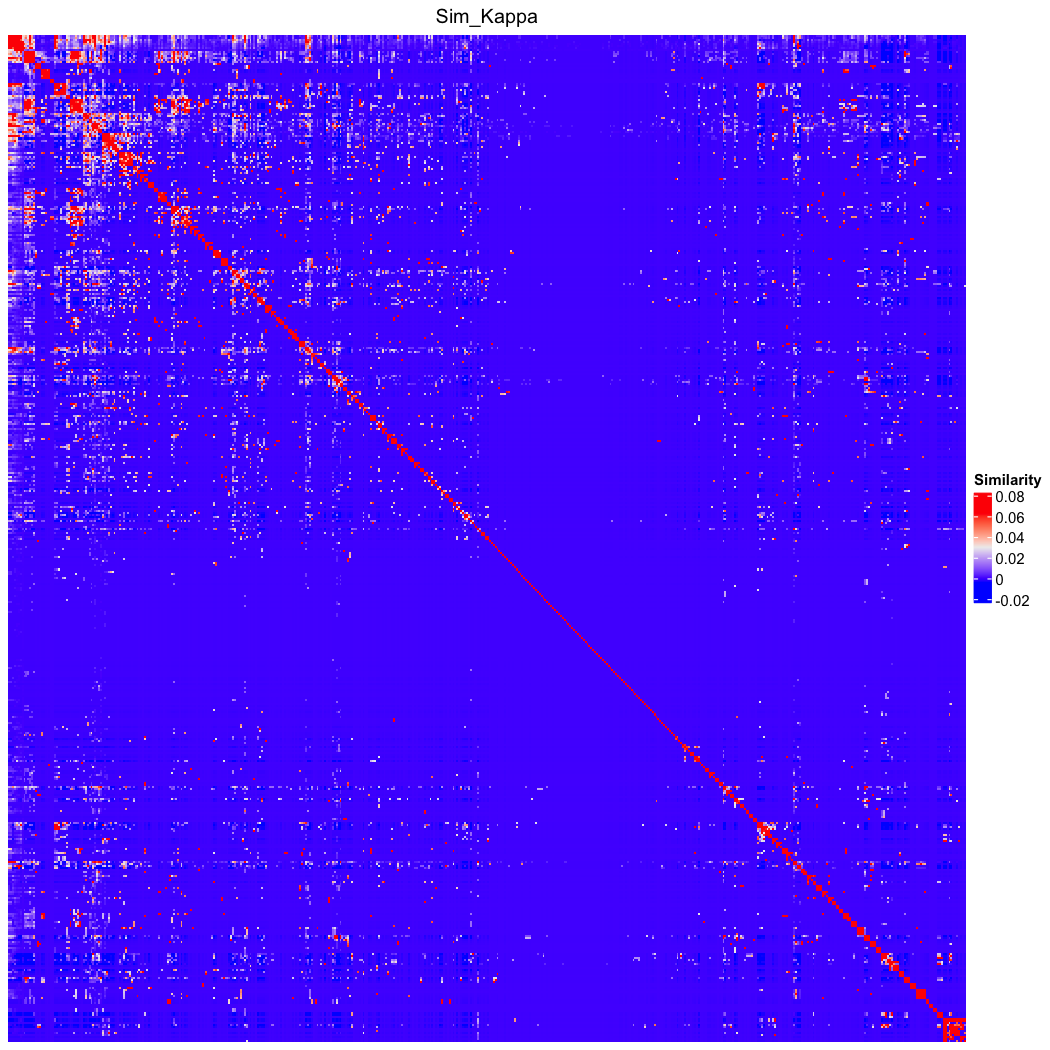

### go_bp_random_500_sim_Sim_Kappa_Lin_order.png

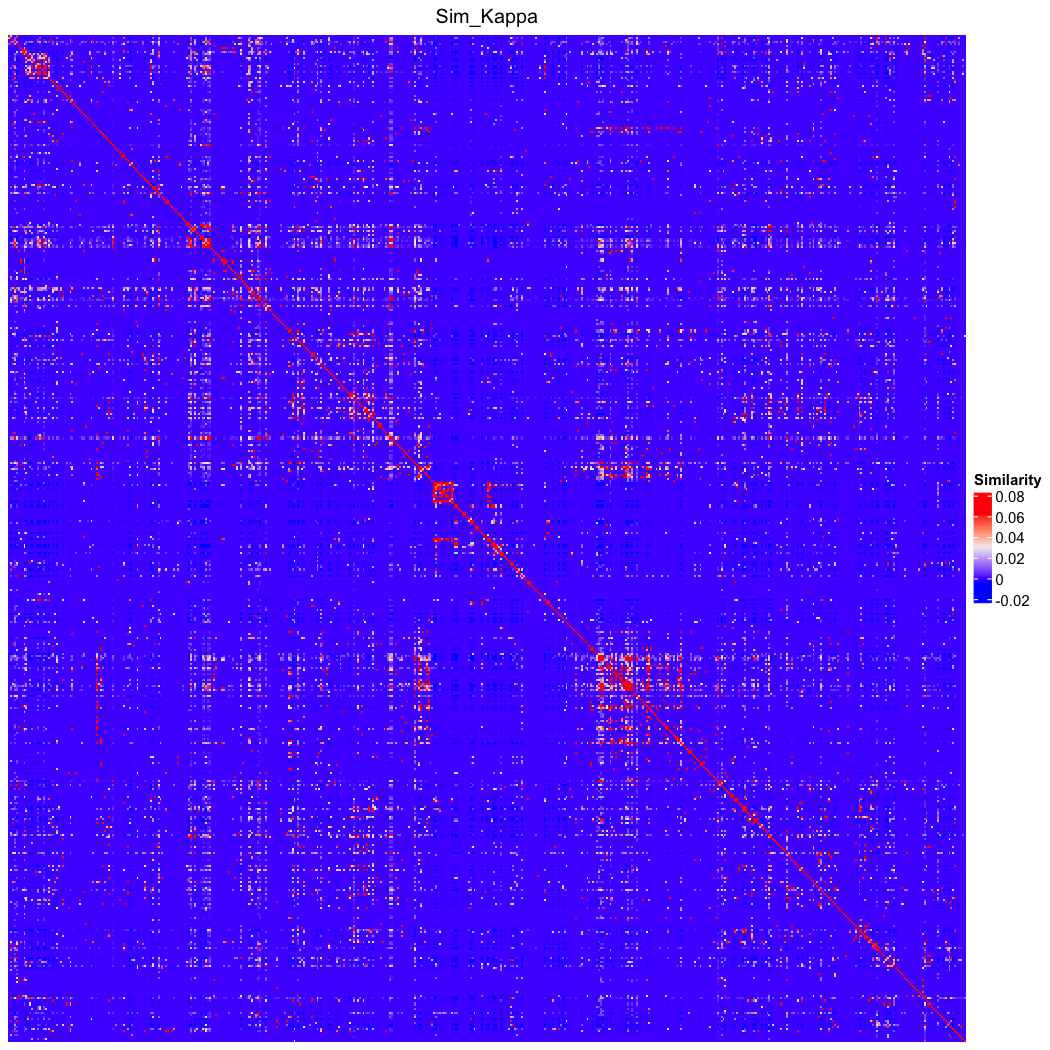

### go_bp_random_500_sim_Sim_Leocock_1998.png

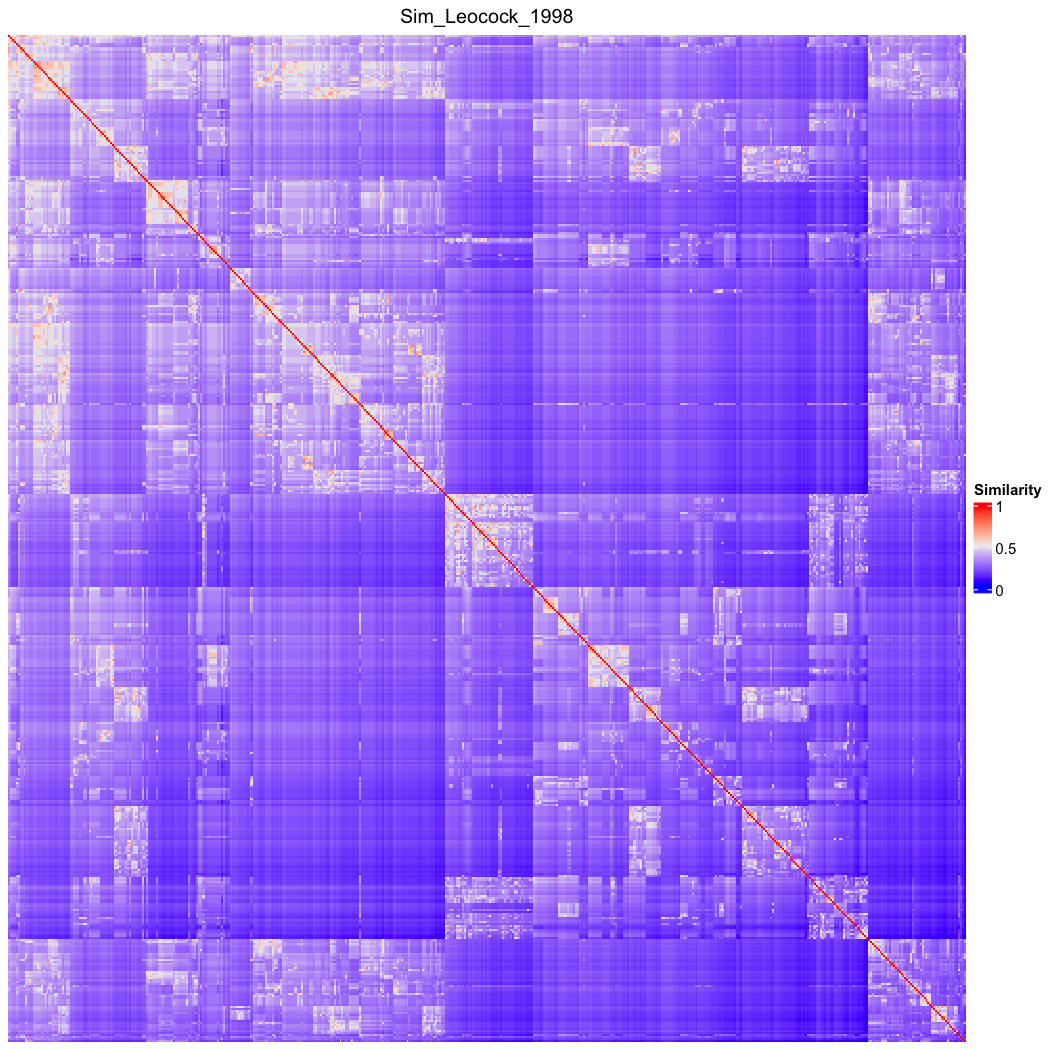

### go_bp_random_500_sim_Sim_Leocock_1998_Lin_order.png

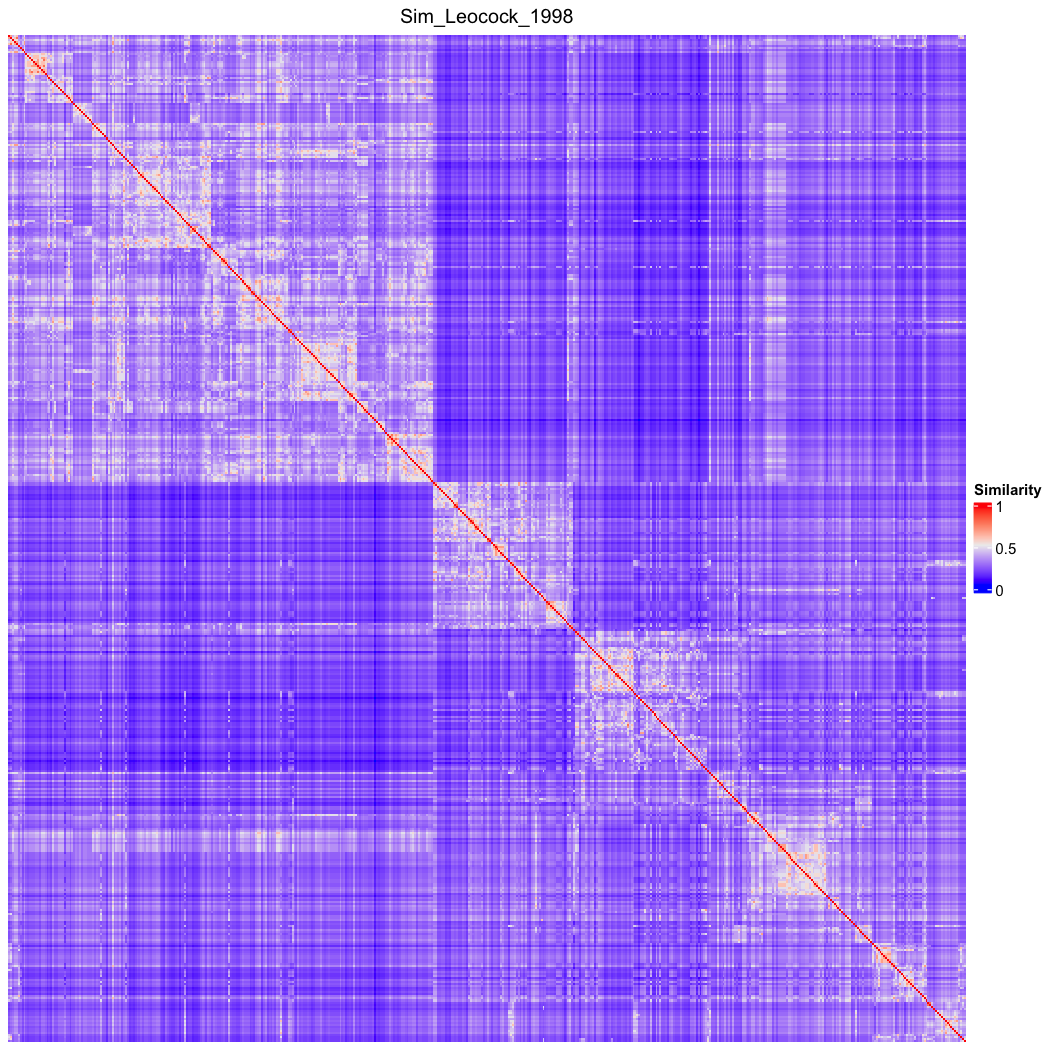

### go_bp_random_500_sim_Sim_Li_2003.png

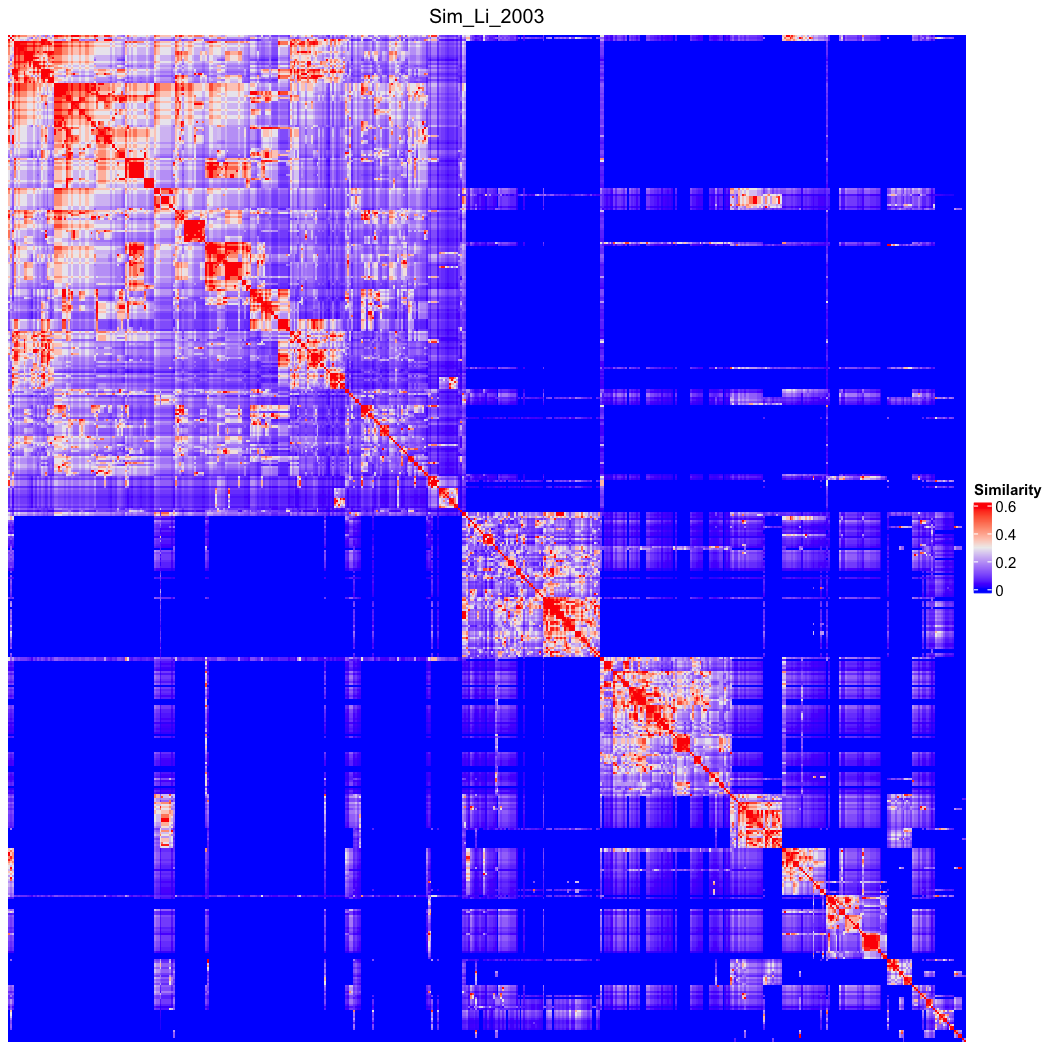

### go_bp_random_500_sim_Sim_Li_2003_Lin_order.png

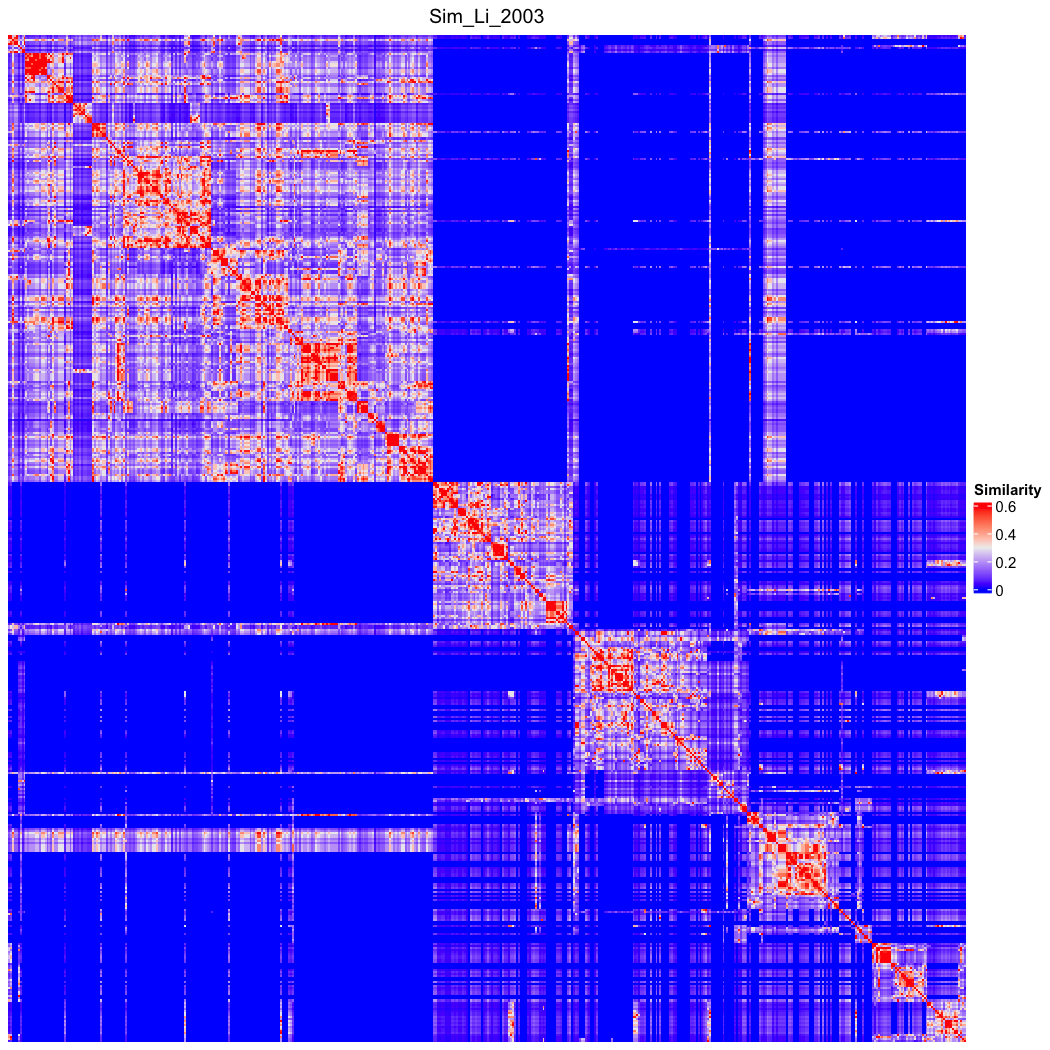

### go_bp_random_500_sim_Sim_Lin_1998.png

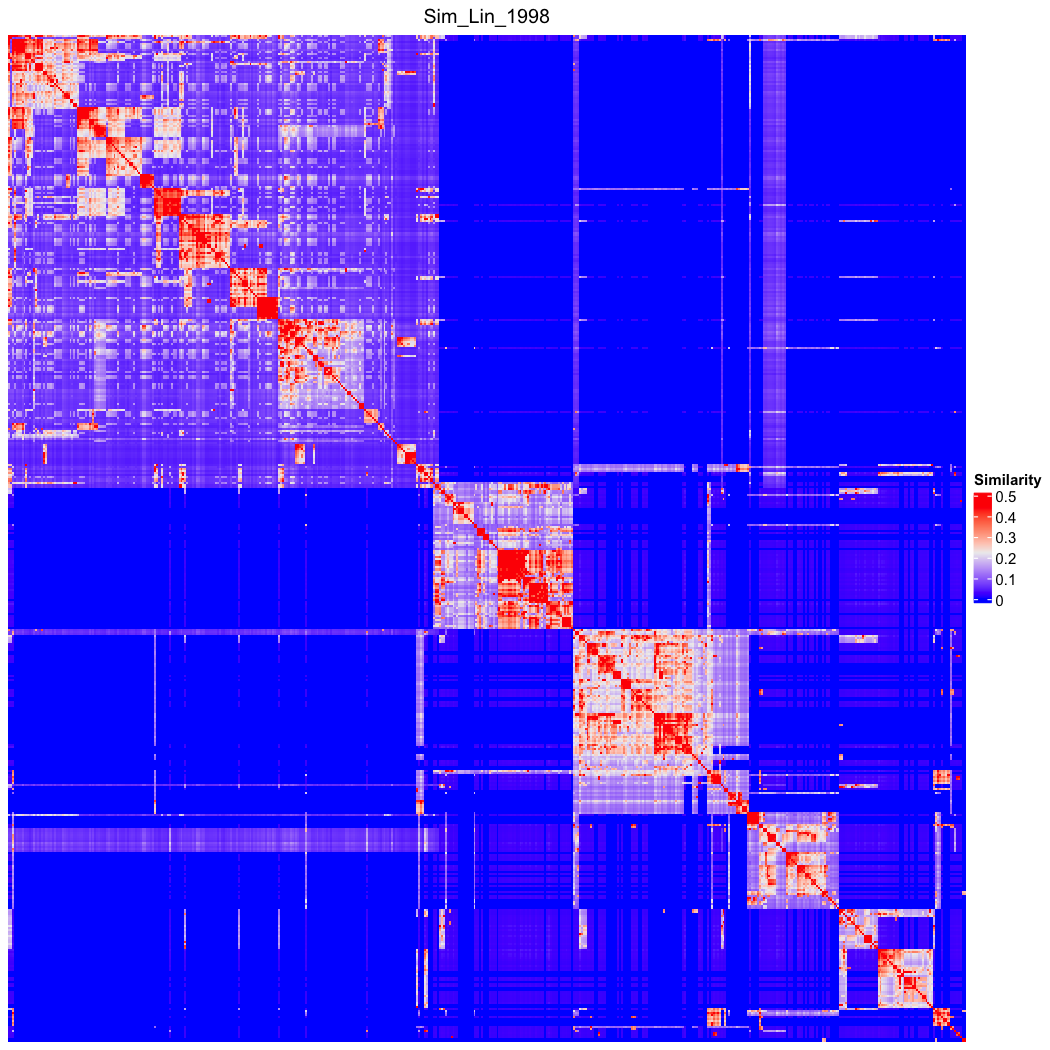

### go_bp_random_500_sim_Sim_Lin_1998_Lin_order.png

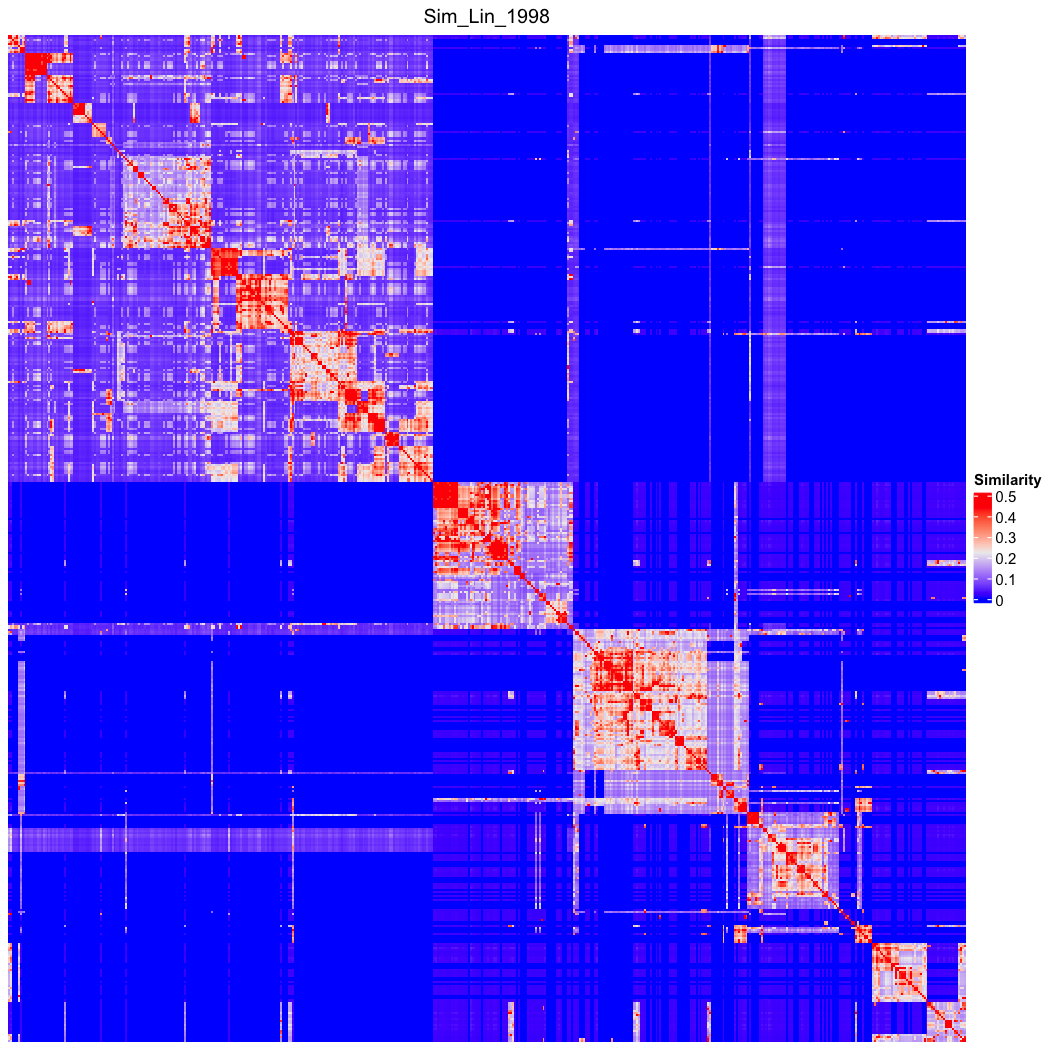

### go_bp_random_500_sim_Sim_Overlap.png

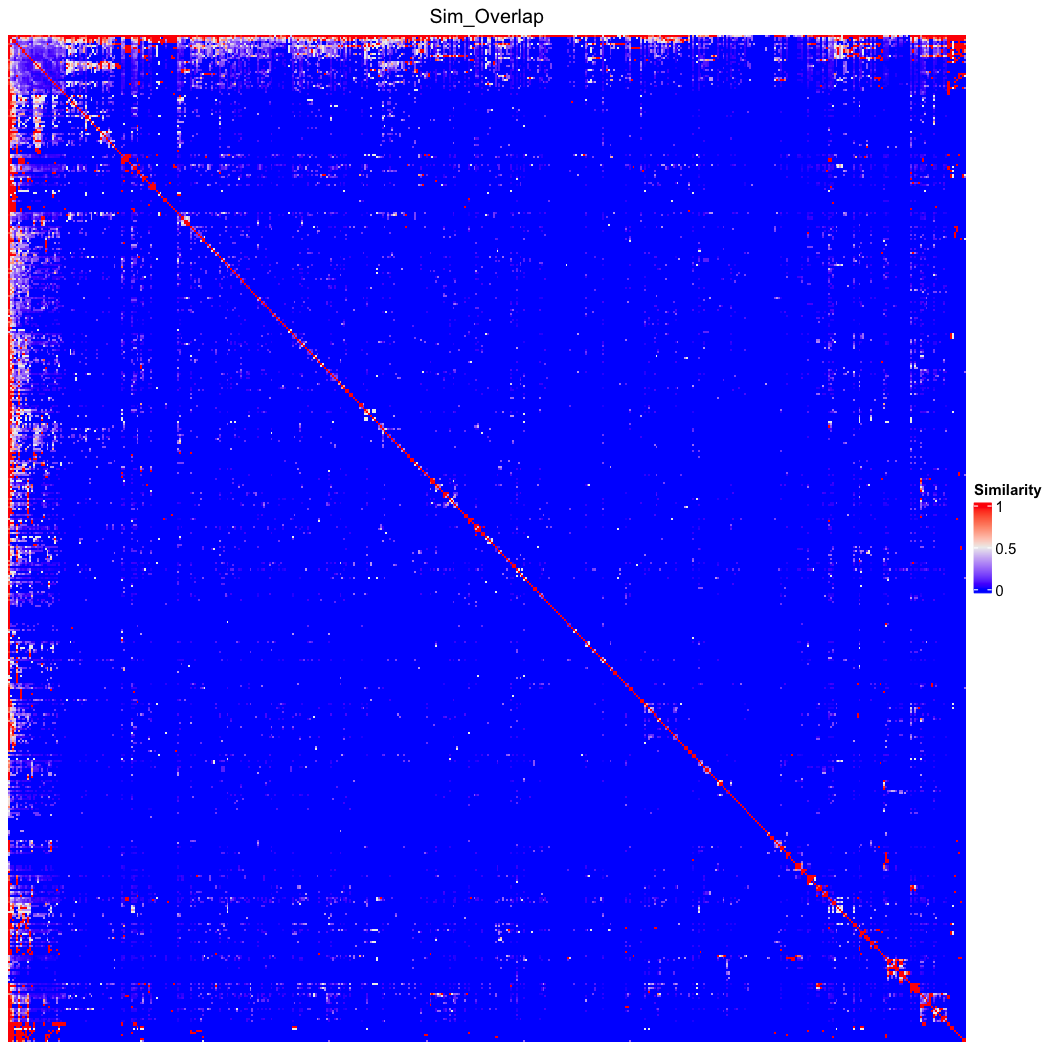

### go_bp_random_500_sim_Sim_Overlap_Lin_order.png

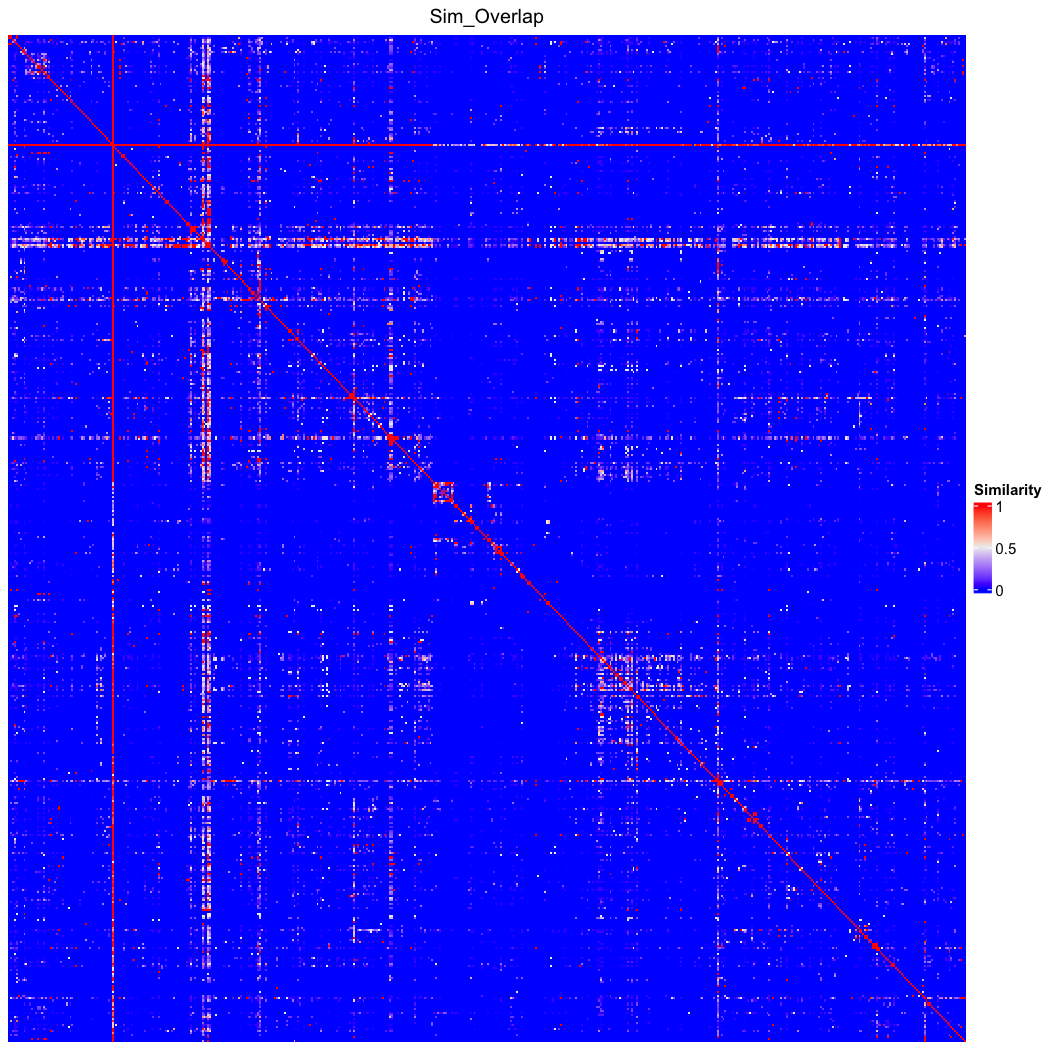

### go_bp_random_500_sim_Sim_Pekar_2002.png

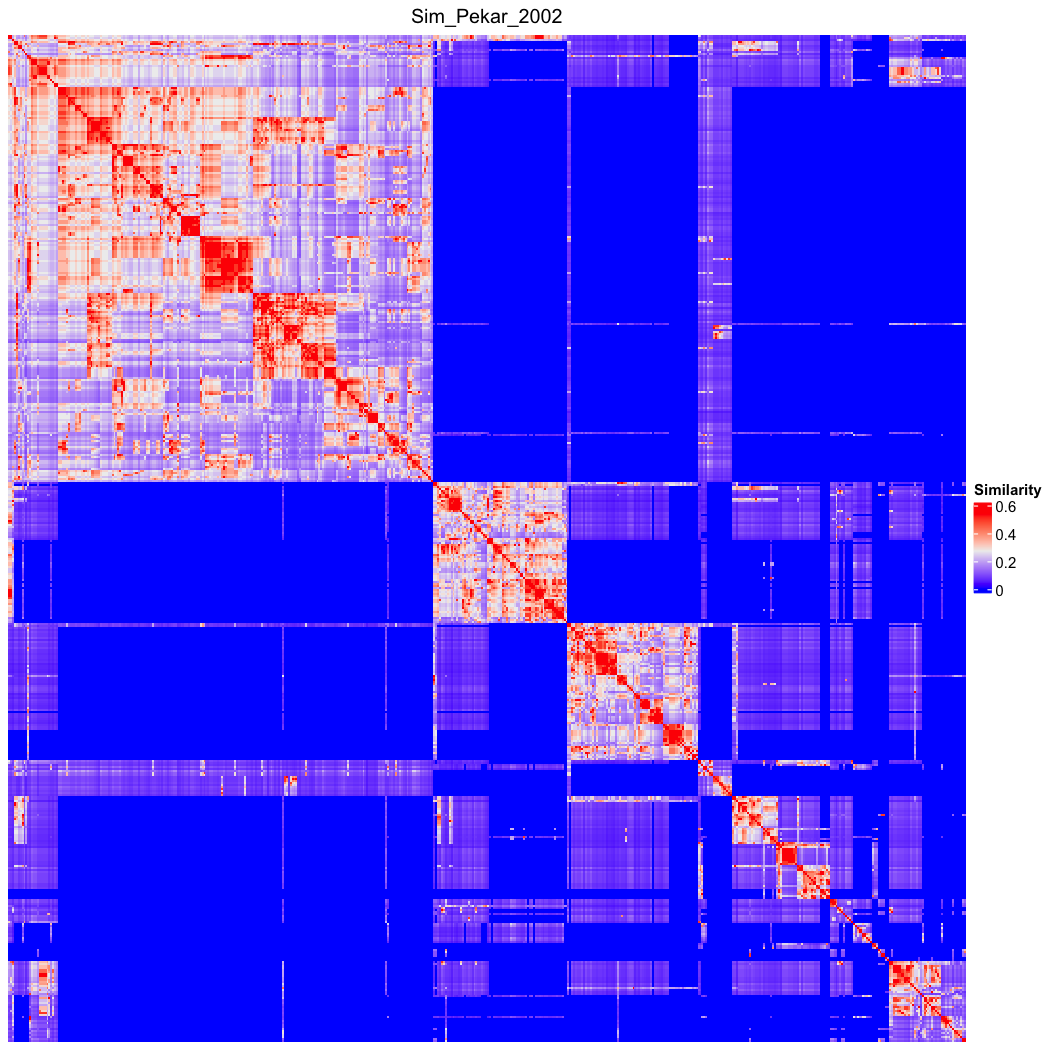

### go_bp_random_500_sim_Sim_Pekar_2002_Lin_order.png

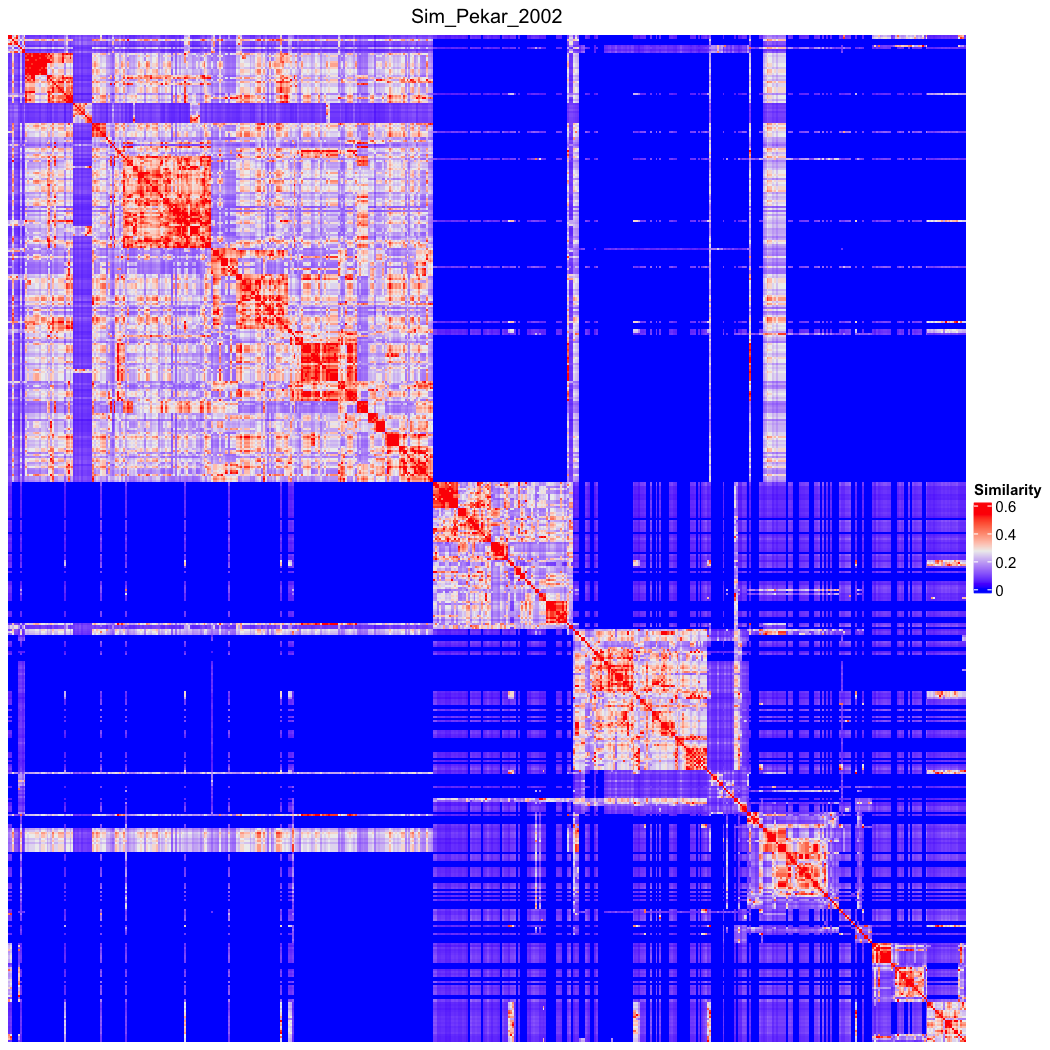

### go_bp_random_500_sim_Sim_Rada_1989.png

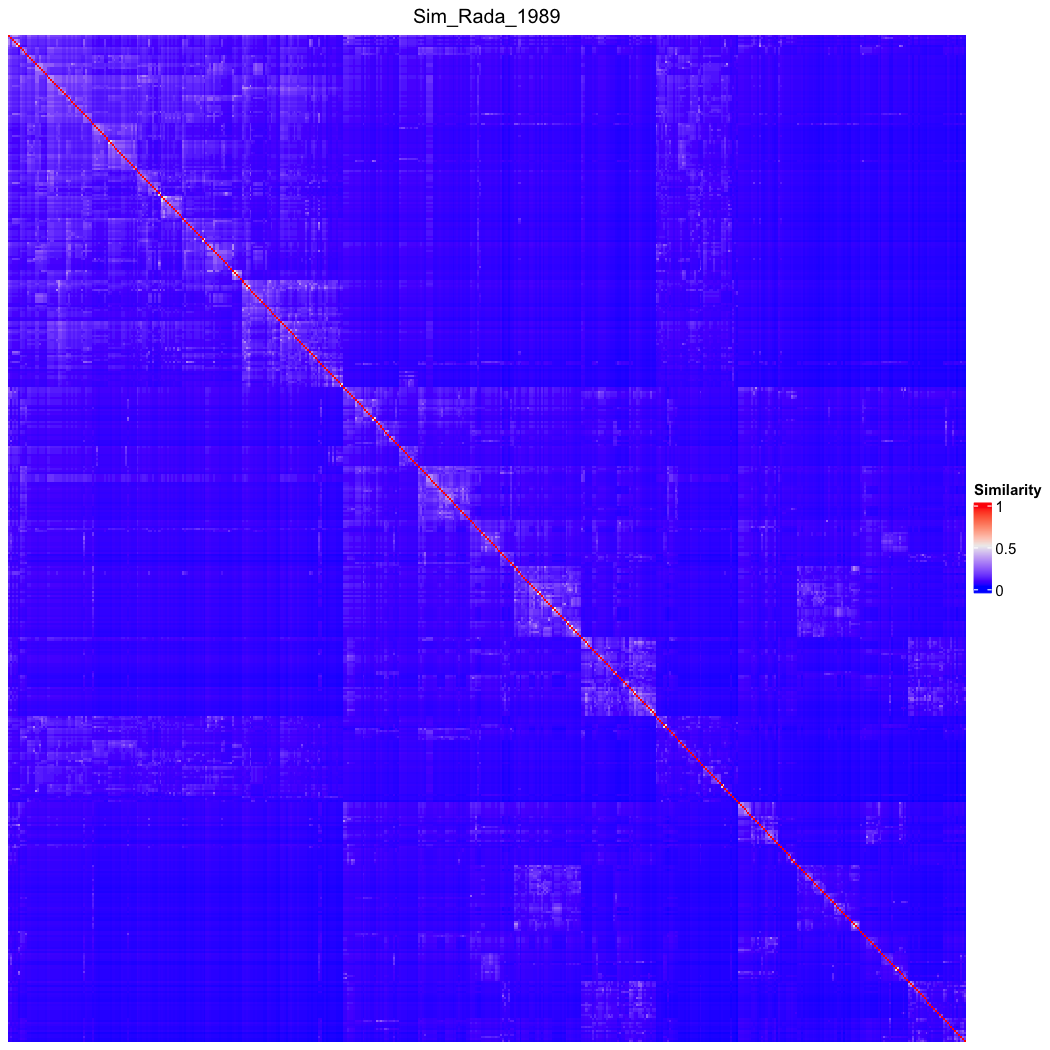

### go_bp_random_500_sim_Sim_Rada_1989_Lin_order.png

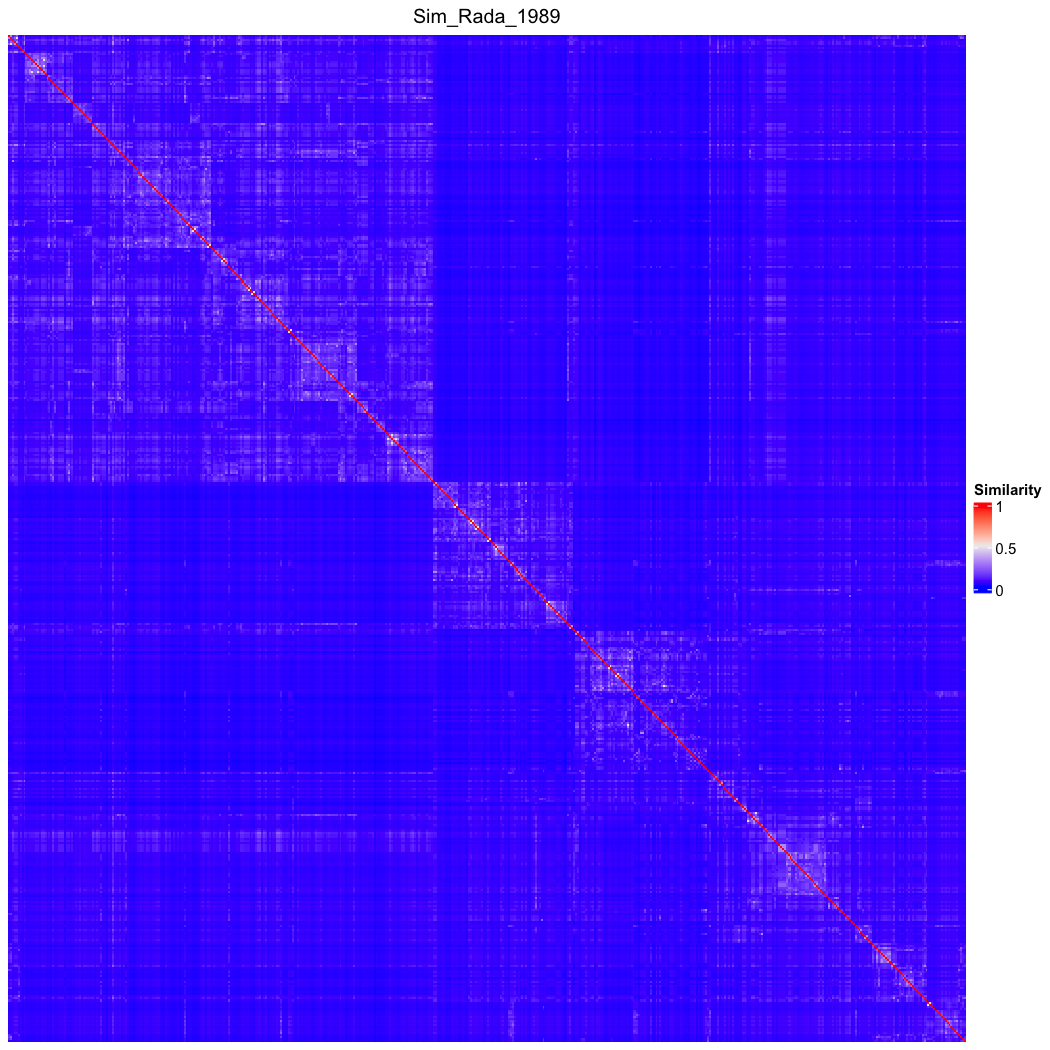

### go_bp_random_500_sim_Sim_Relevance_2006.png

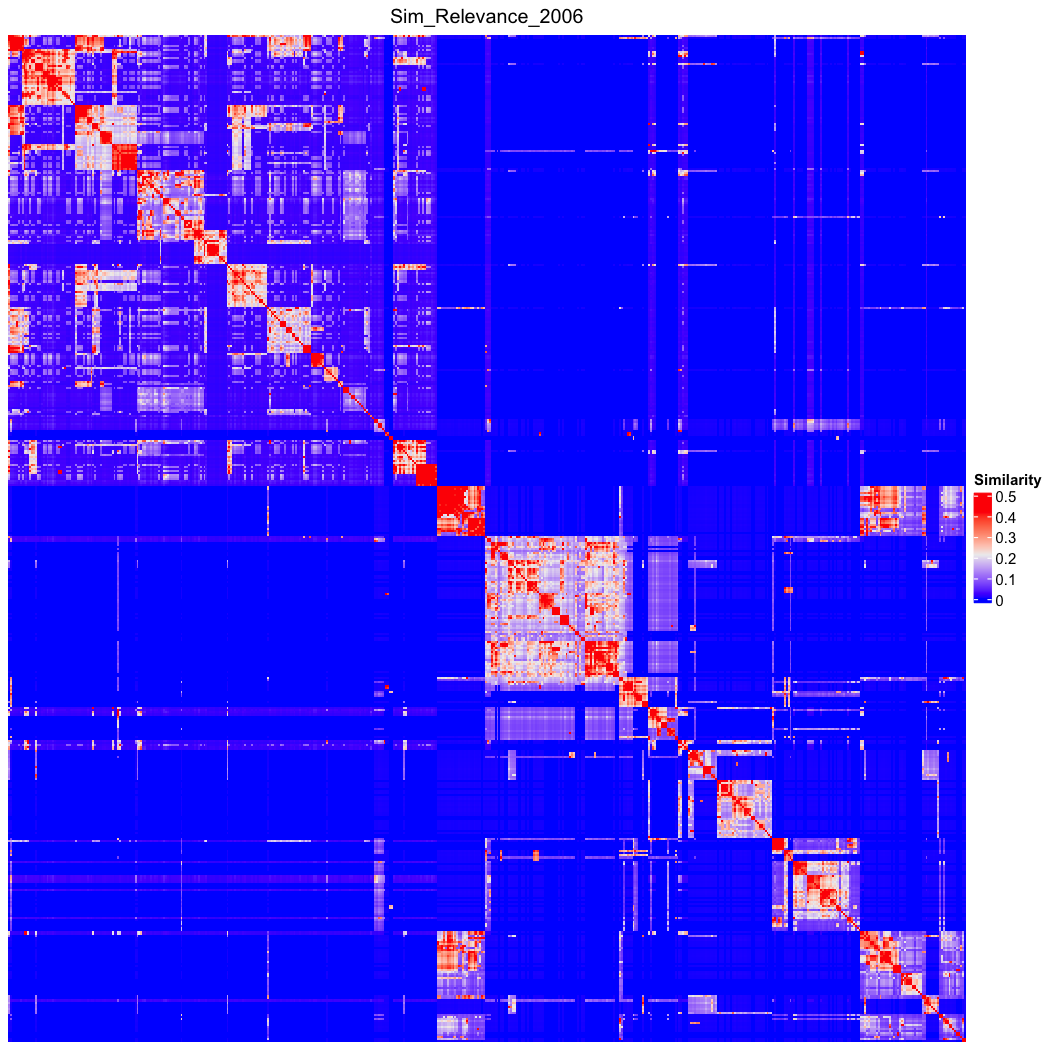

### go_bp_random_500_sim_Sim_Relevance_2006_Lin_order.png

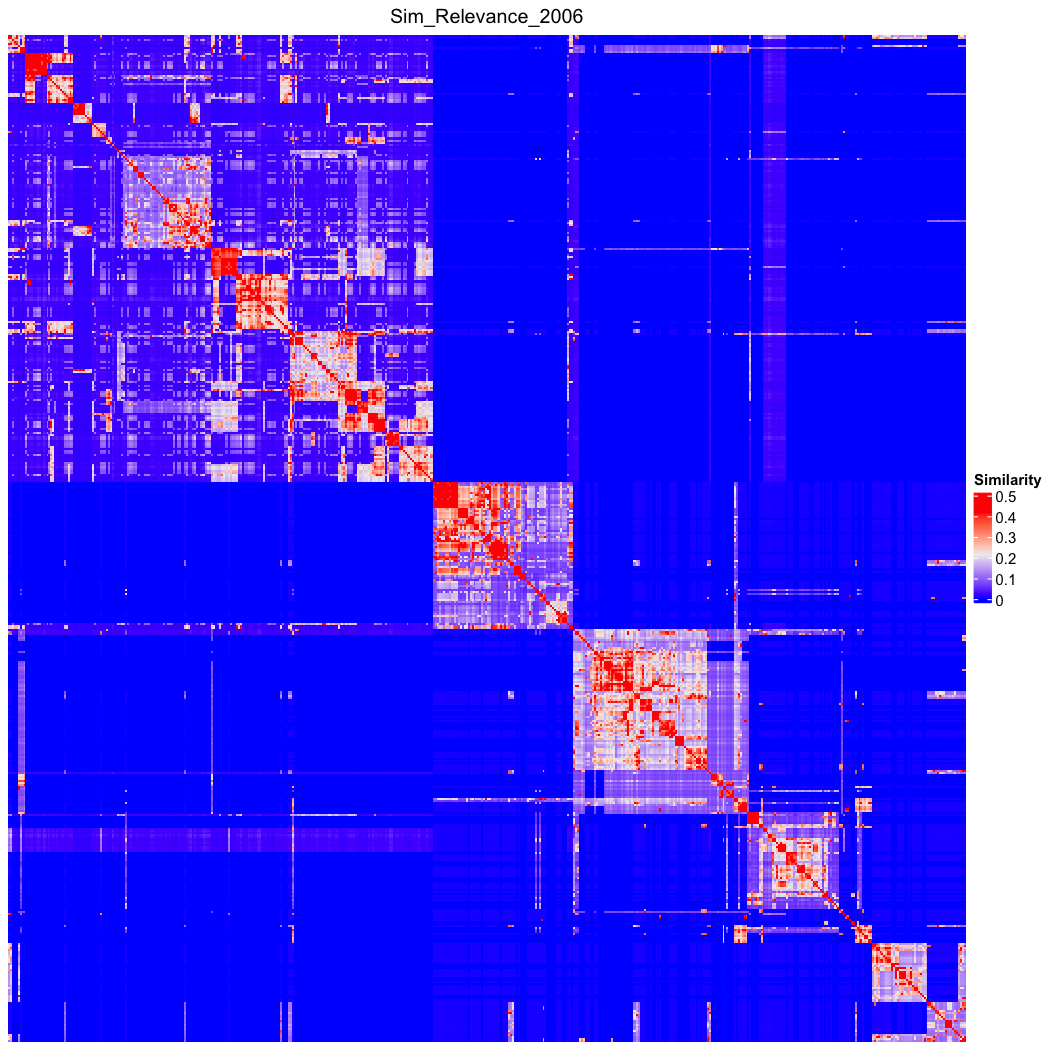

### go_bp_random_500_sim_Sim_Resnik_1999.png

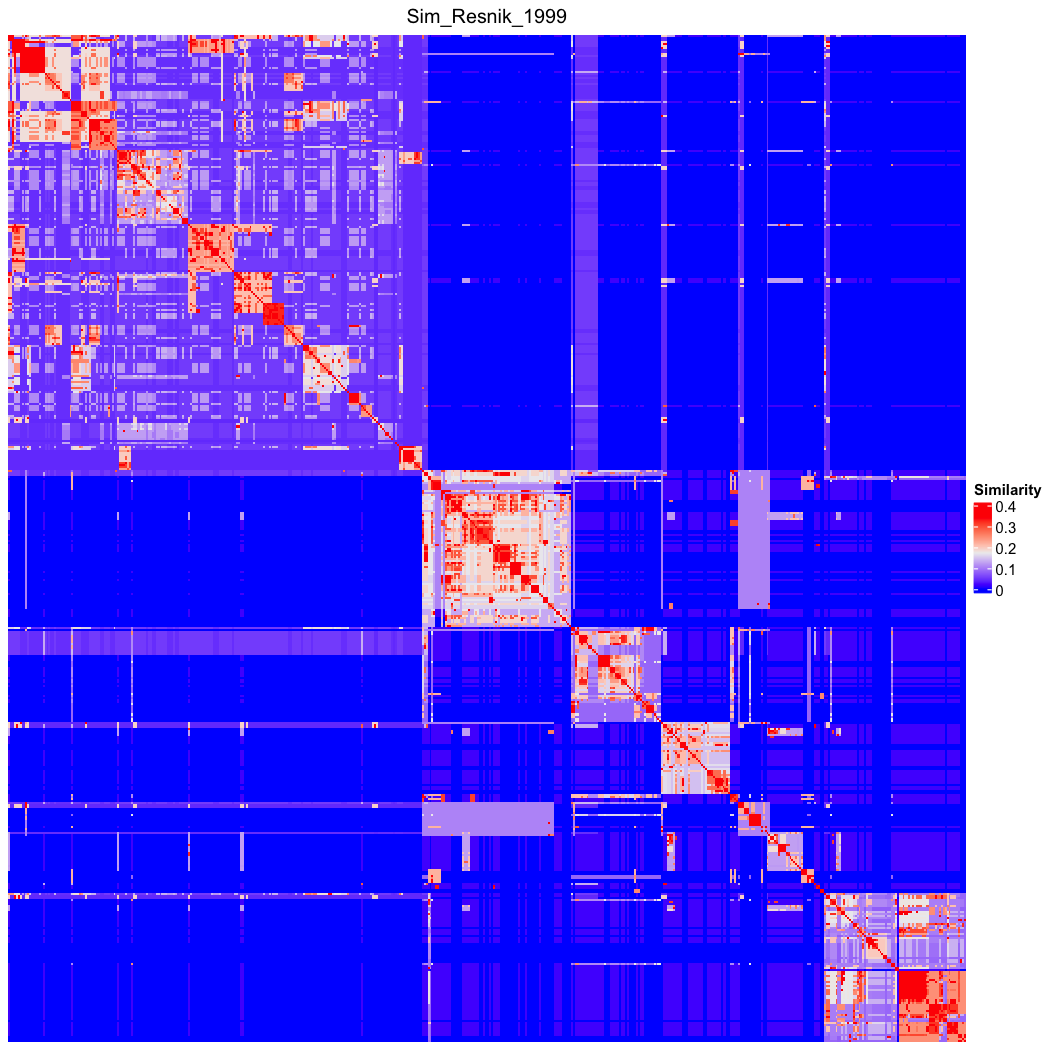

### go_bp_random_500_sim_Sim_Resnik_1999_Lin_order.png

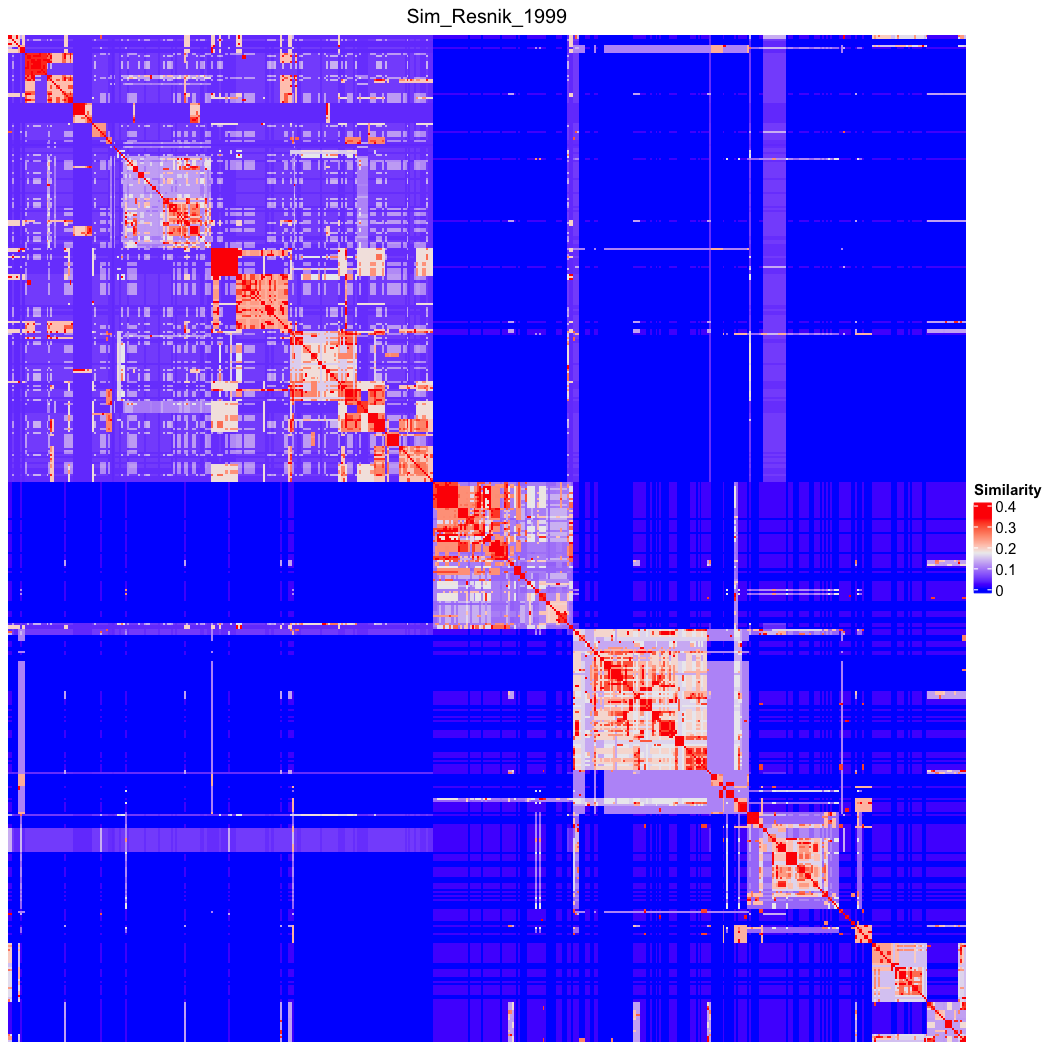

### go_bp_random_500_sim_Sim_Resnik_edge_2005.png

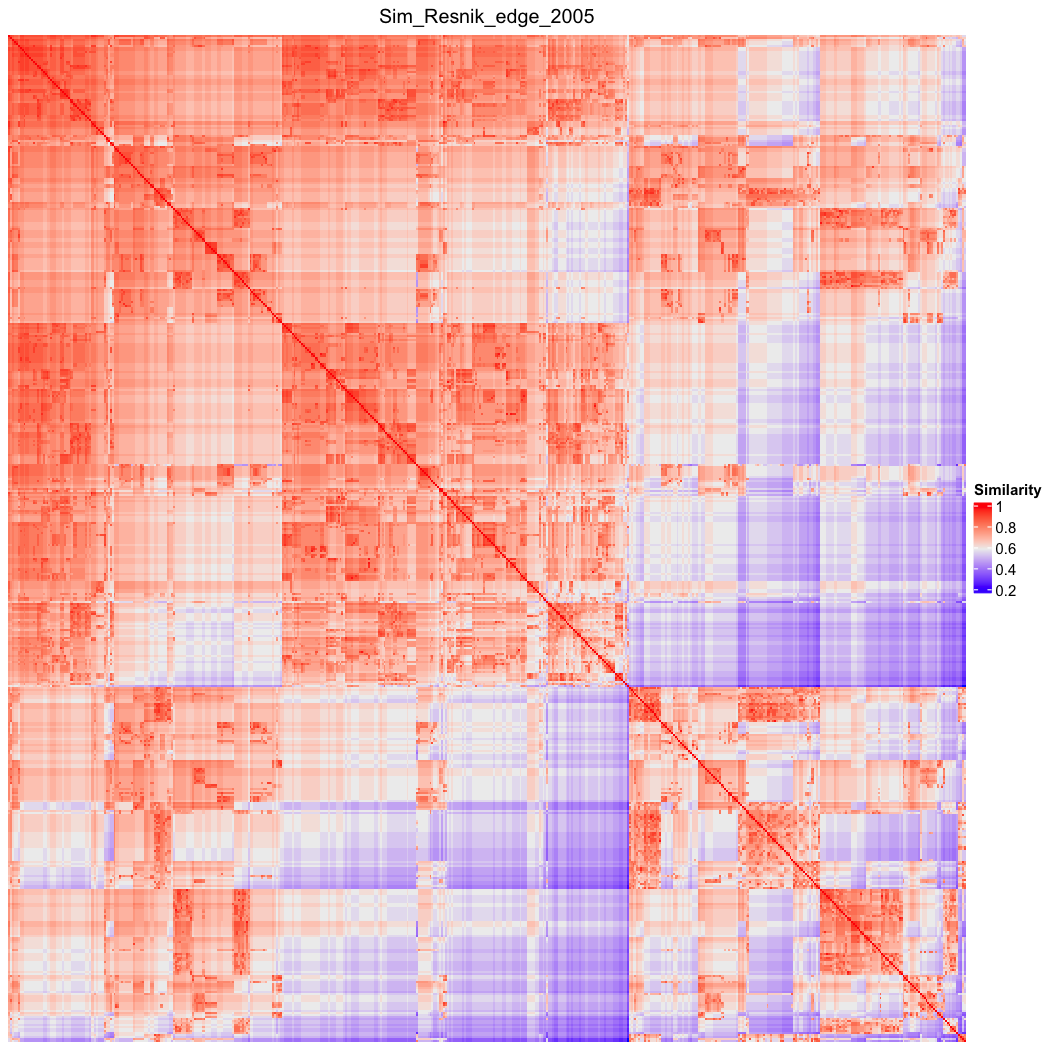

### go_bp_random_500_sim_Sim_Resnik_edge_2005_Lin_order.png

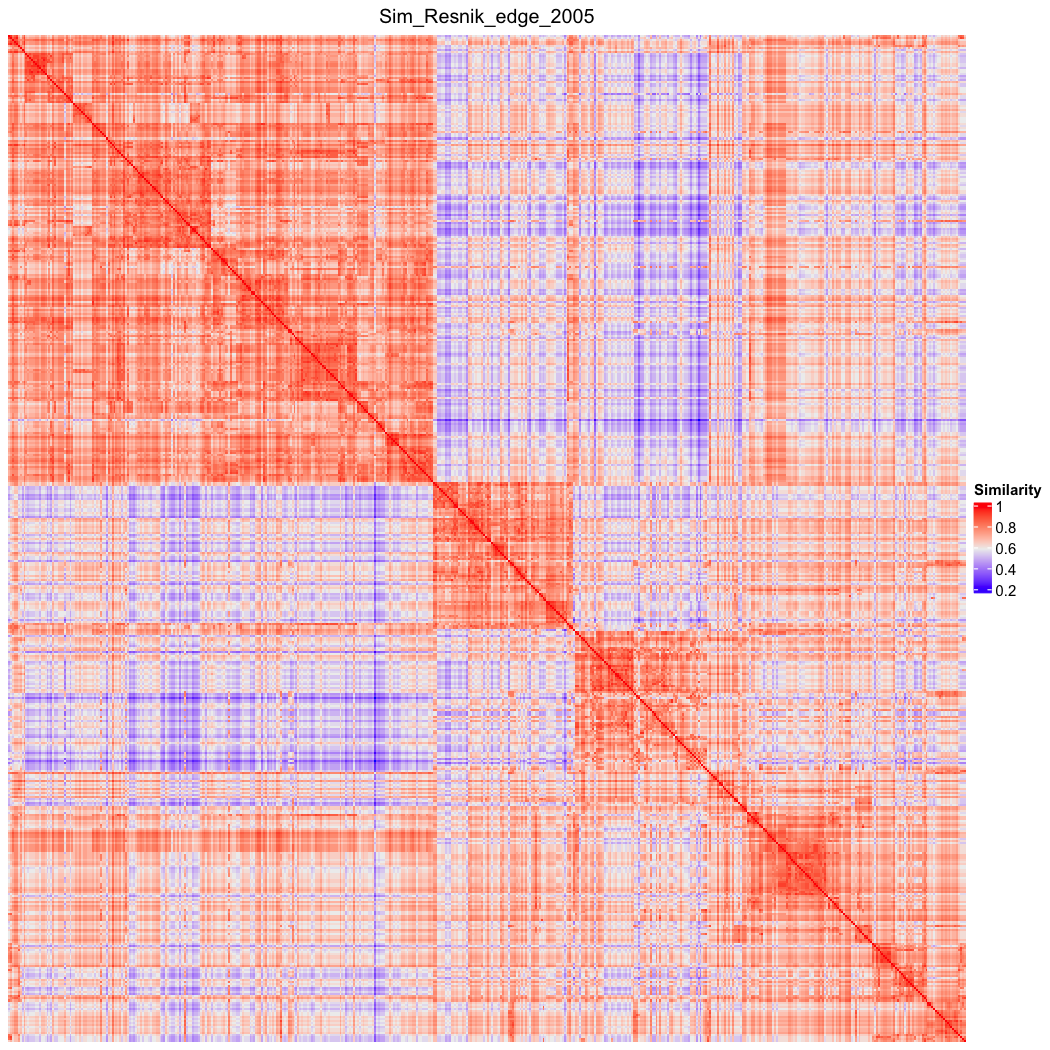

### go_bp_random_500_sim_Sim_RSS_2013.png

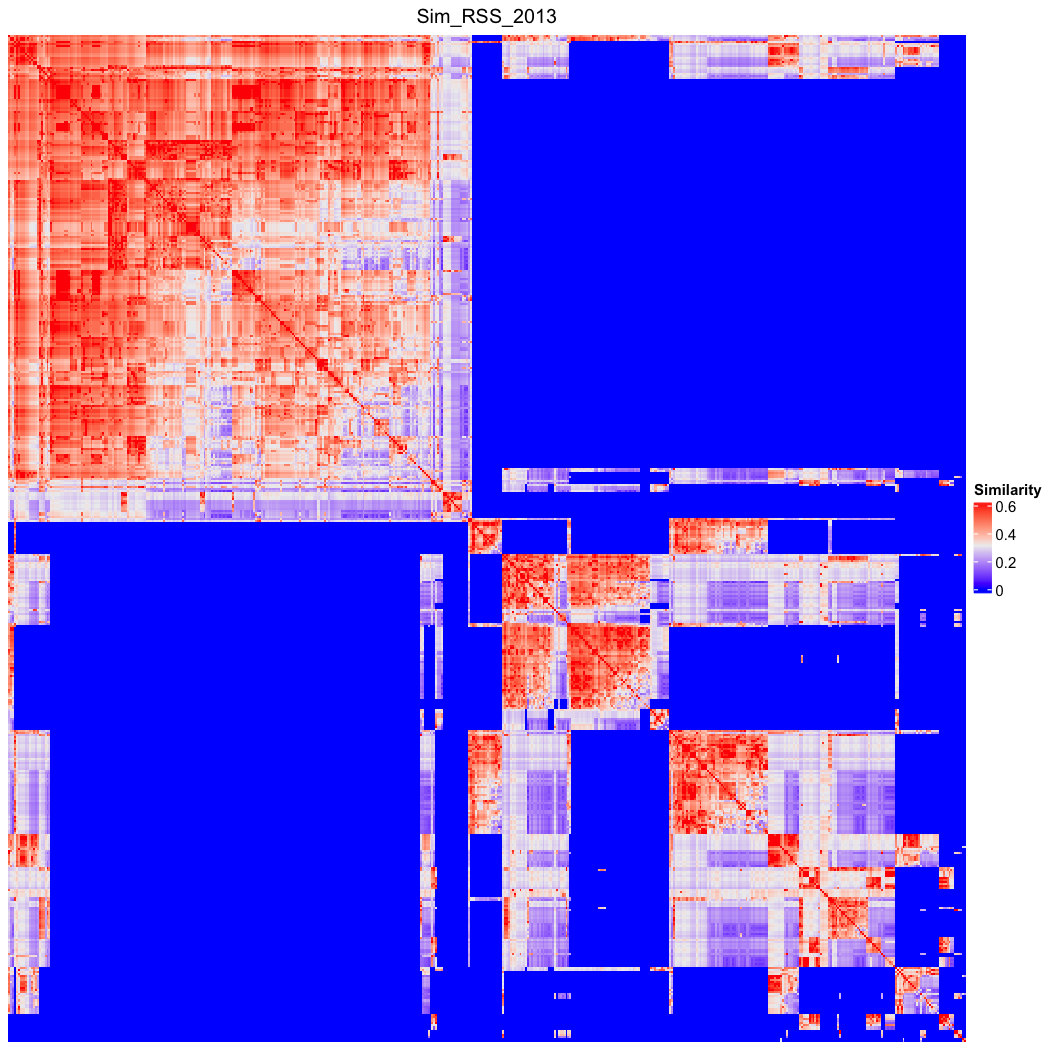

### go_bp_random_500_sim_Sim_RSS_2013_Lin_order.png

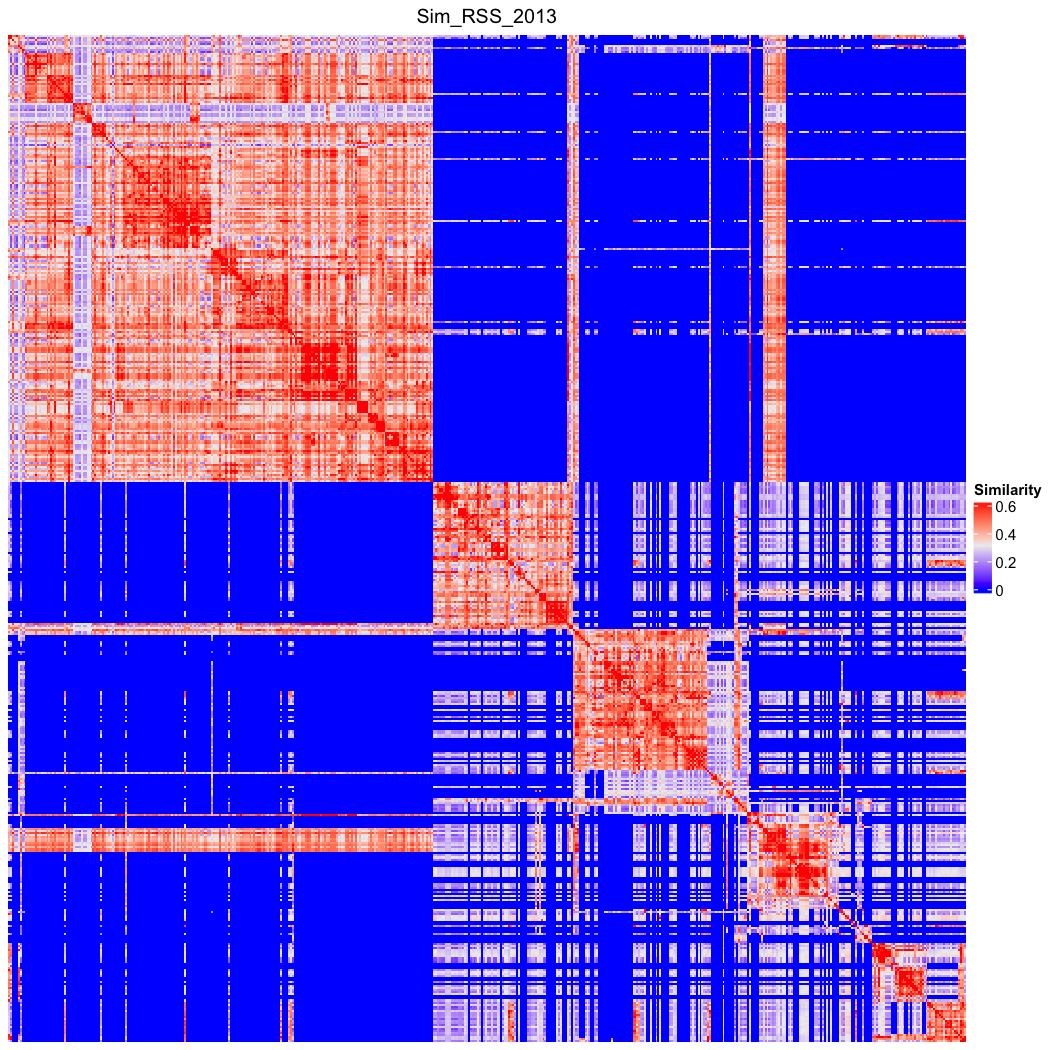

### go_bp_random_500_sim_Sim_Shen_2010.png

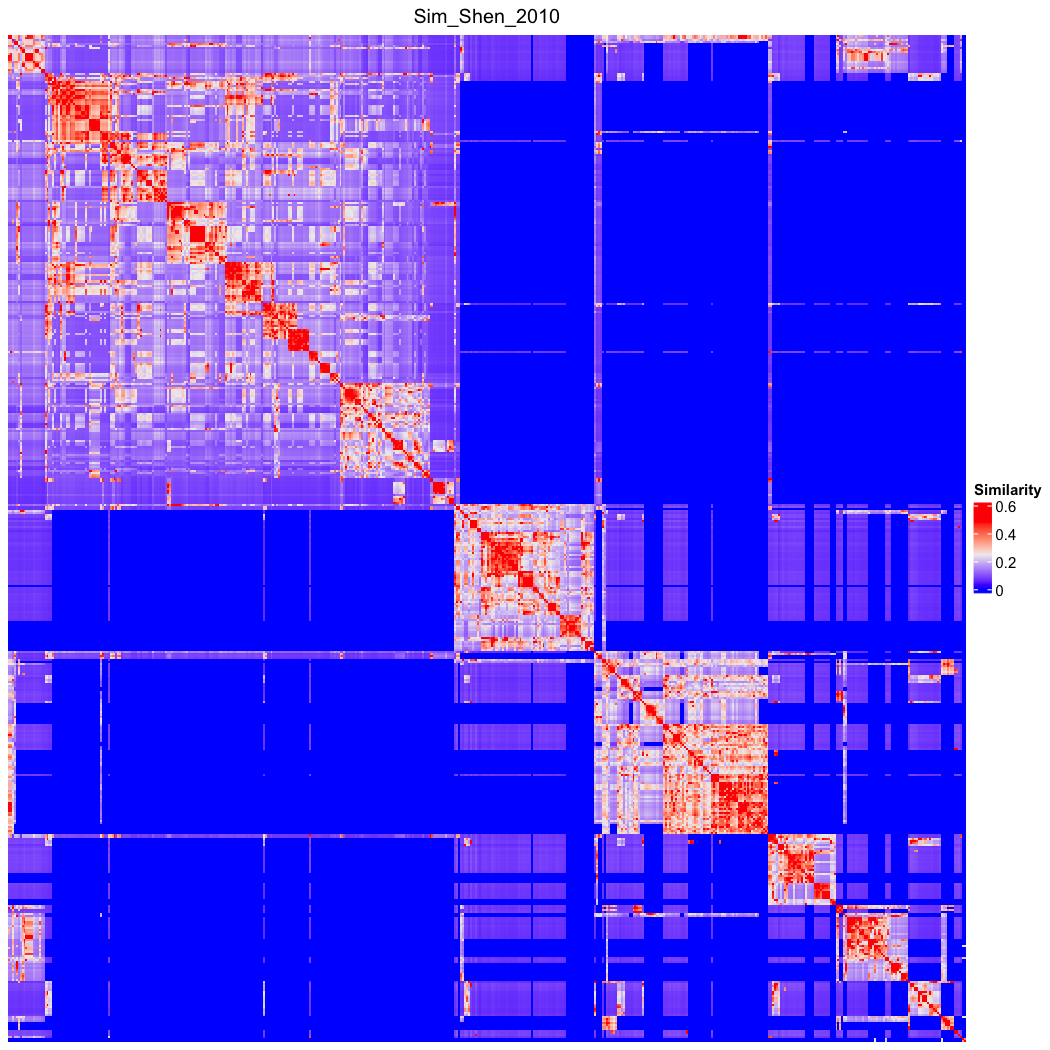

### go_bp_random_500_sim_Sim_Shen_2010_Lin_order.png

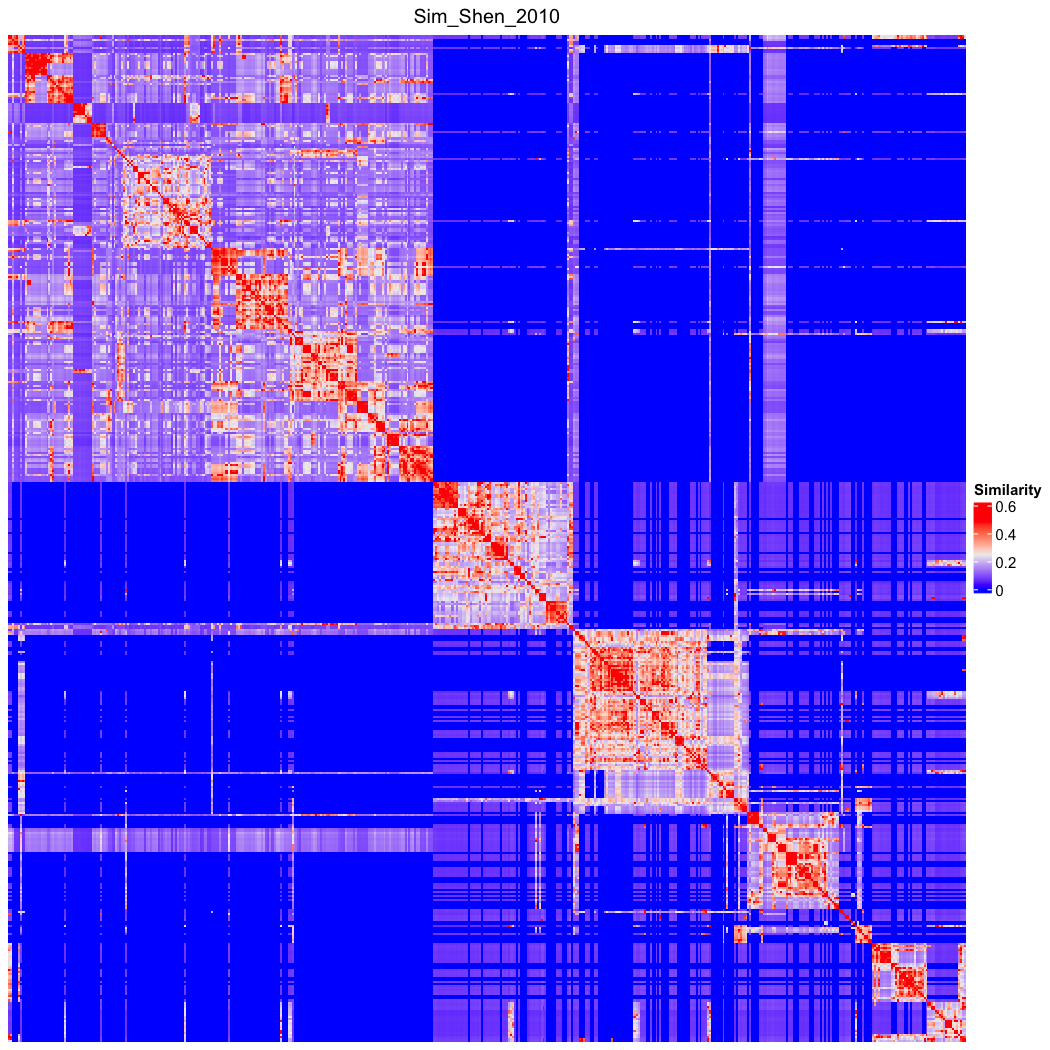

### go_bp_random_500_sim_Sim_Shenoy_2012.png

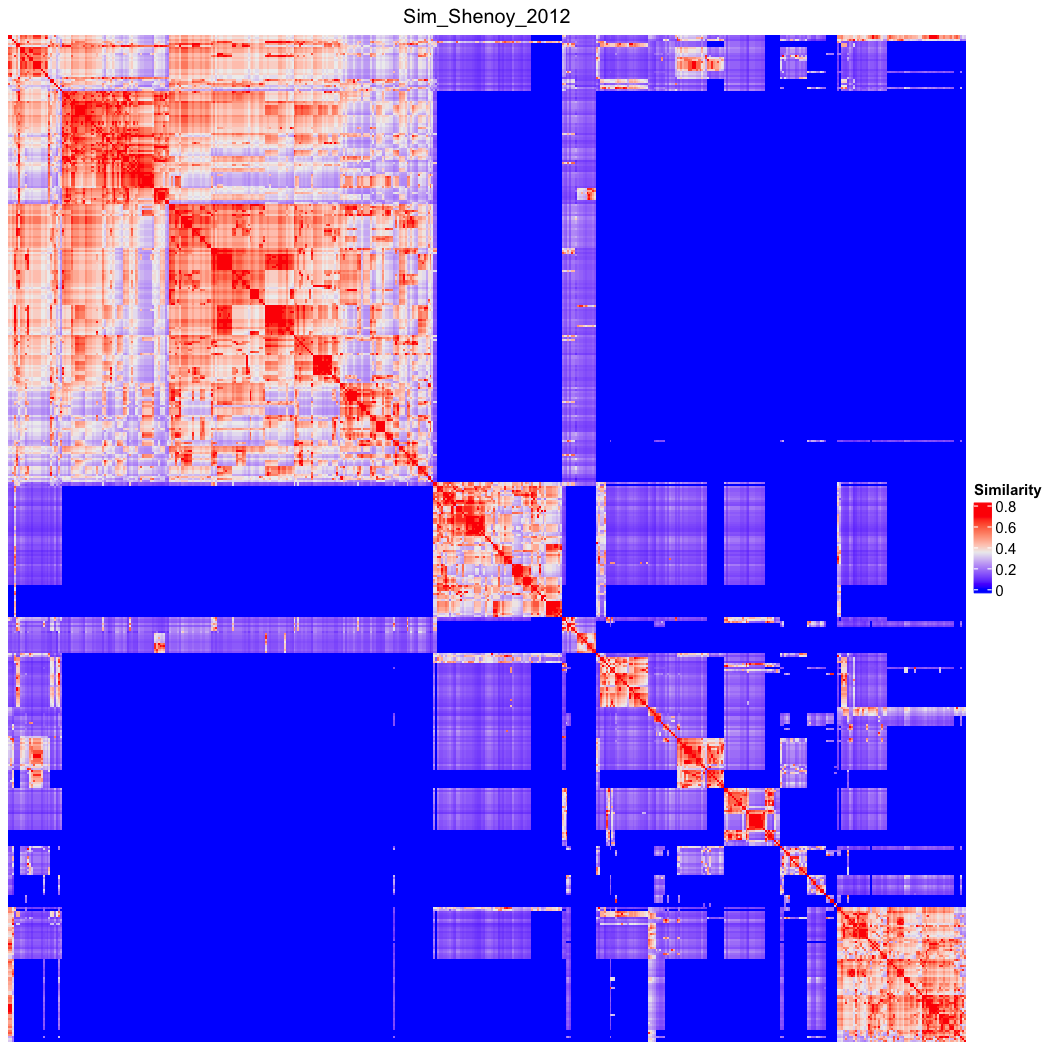

### go_bp_random_500_sim_Sim_Shenoy_2012_Lin_order.png

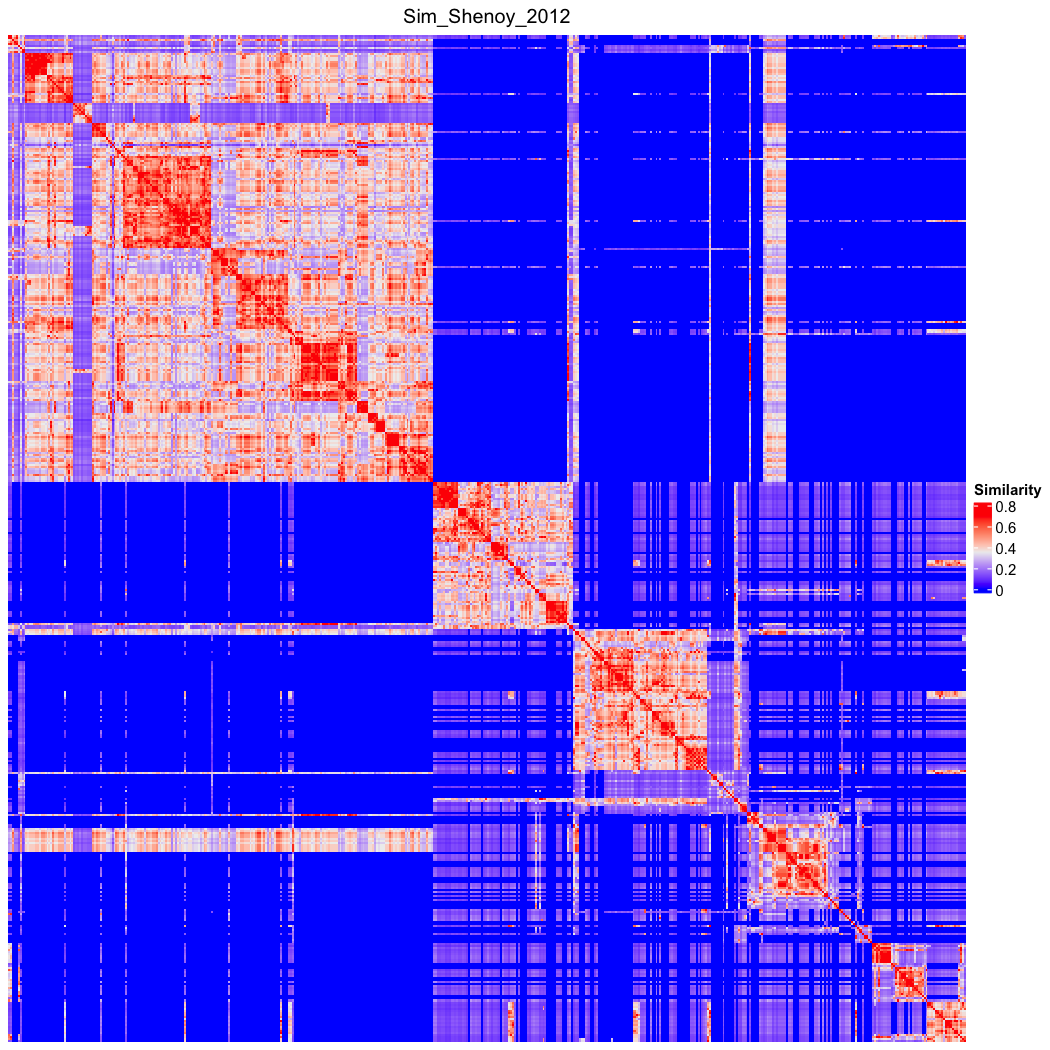

### go_bp_random_500_sim_Sim_SimIC_2010.png

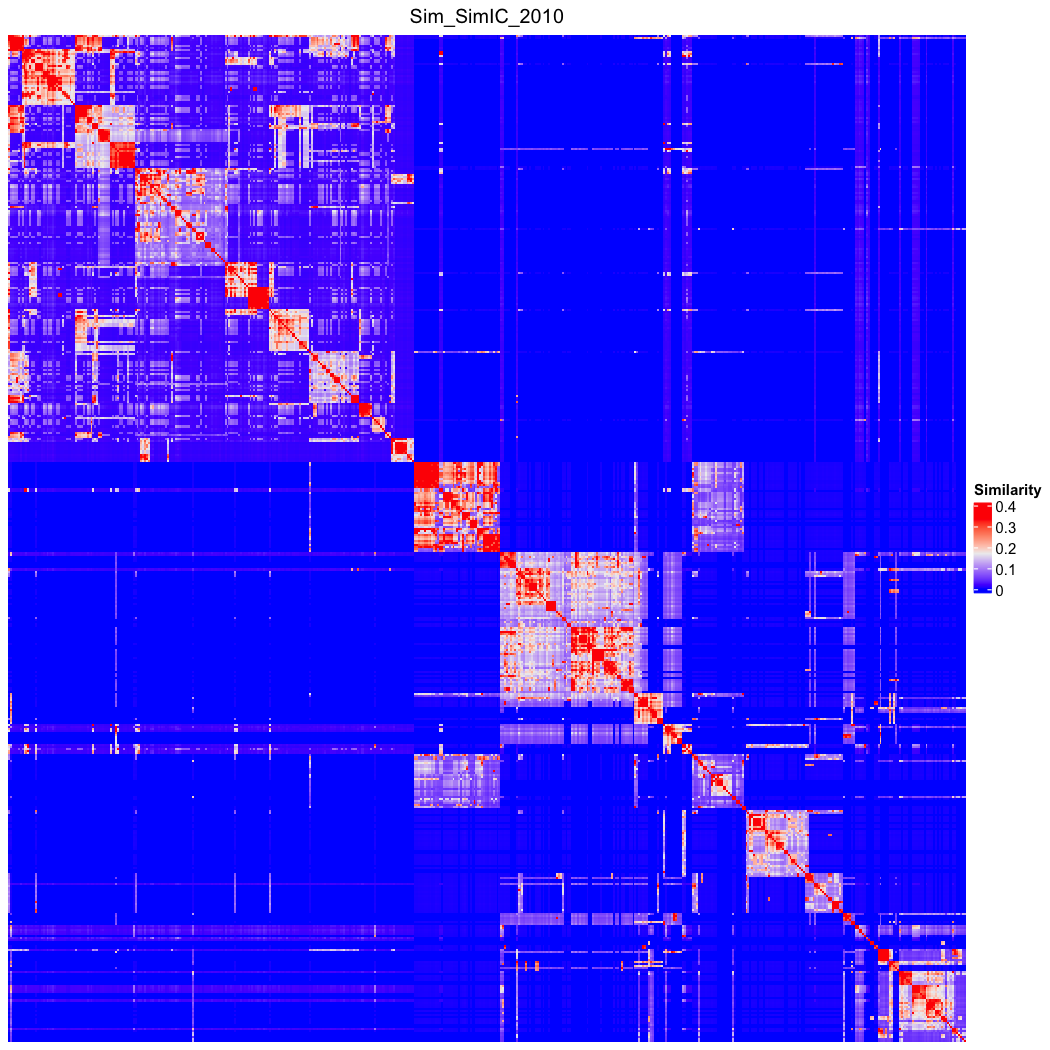

### go_bp_random_500_sim_Sim_SimIC_2010_Lin_order.png

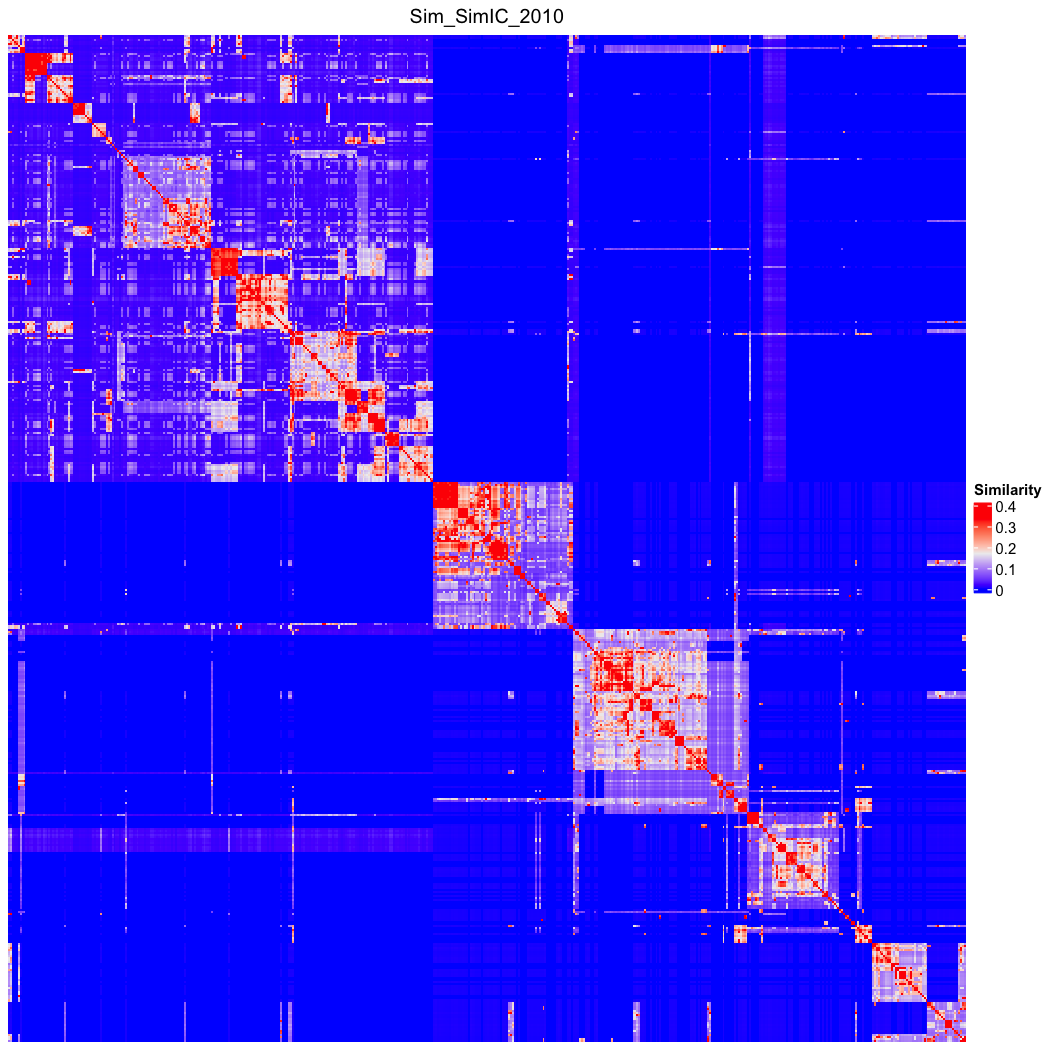

### go_bp_random_500_sim_Sim_Slimani_2006.png

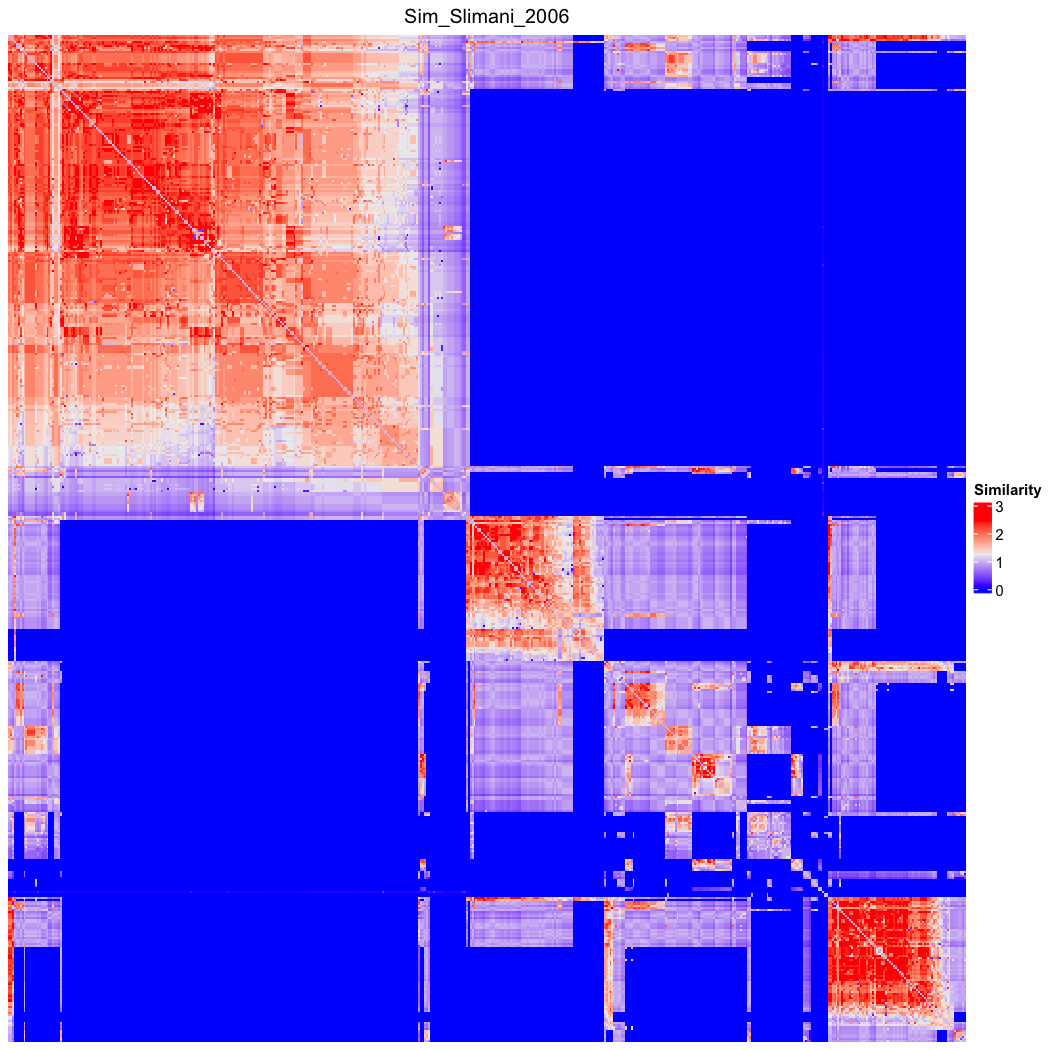

### go_bp_random_500_sim_Sim_Slimani_2006_Lin_order.png

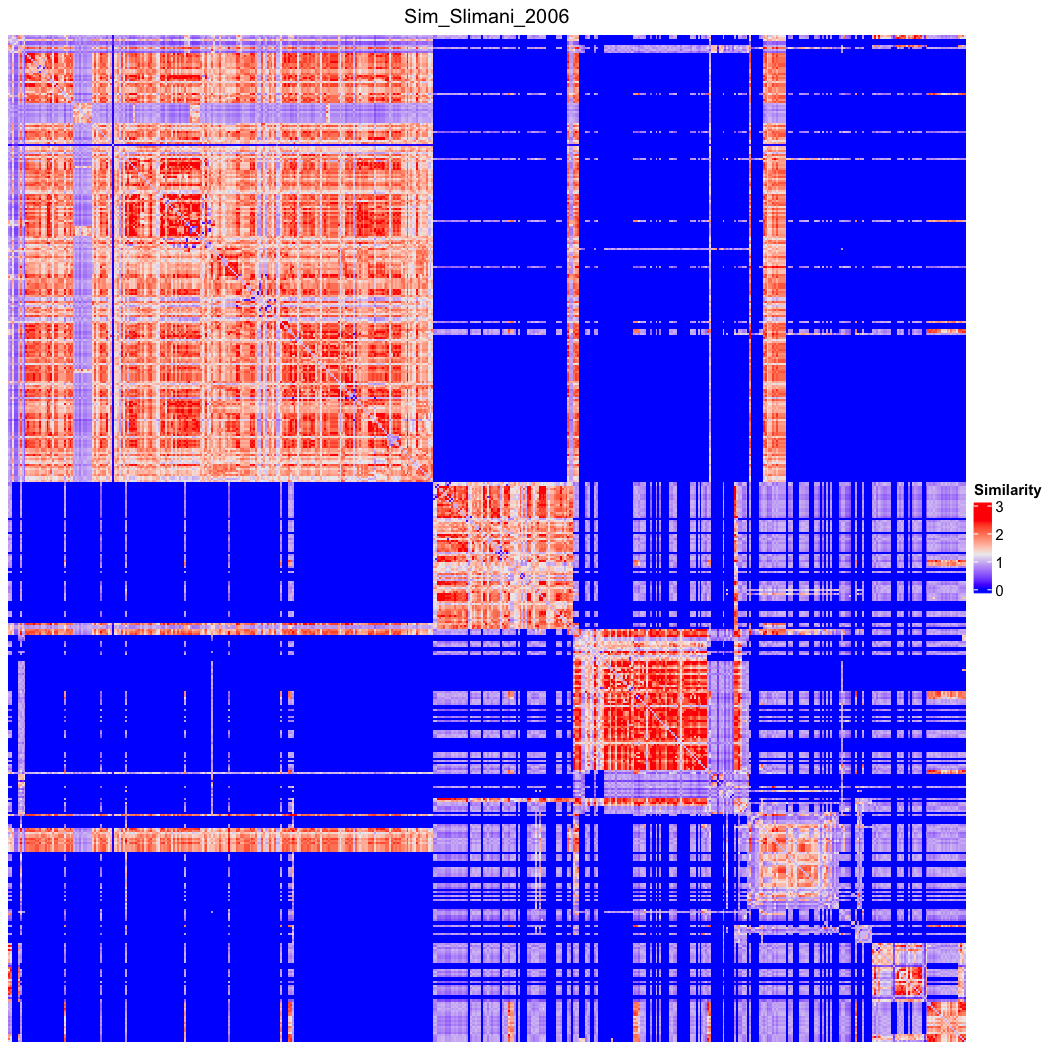

### go_bp_random_500_sim_Sim_SSDD_2013.png

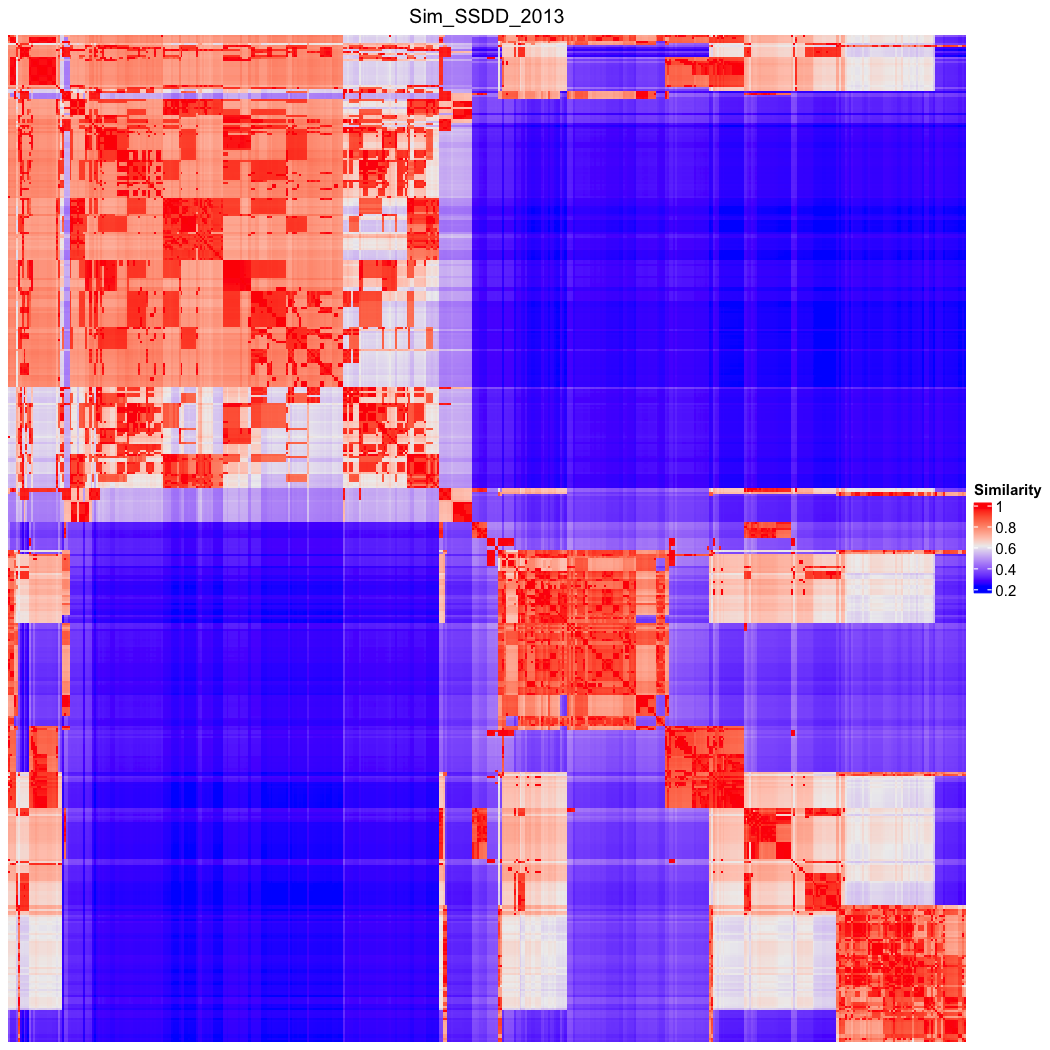

### go_bp_random_500_sim_Sim_SSDD_2013_Lin_order.png

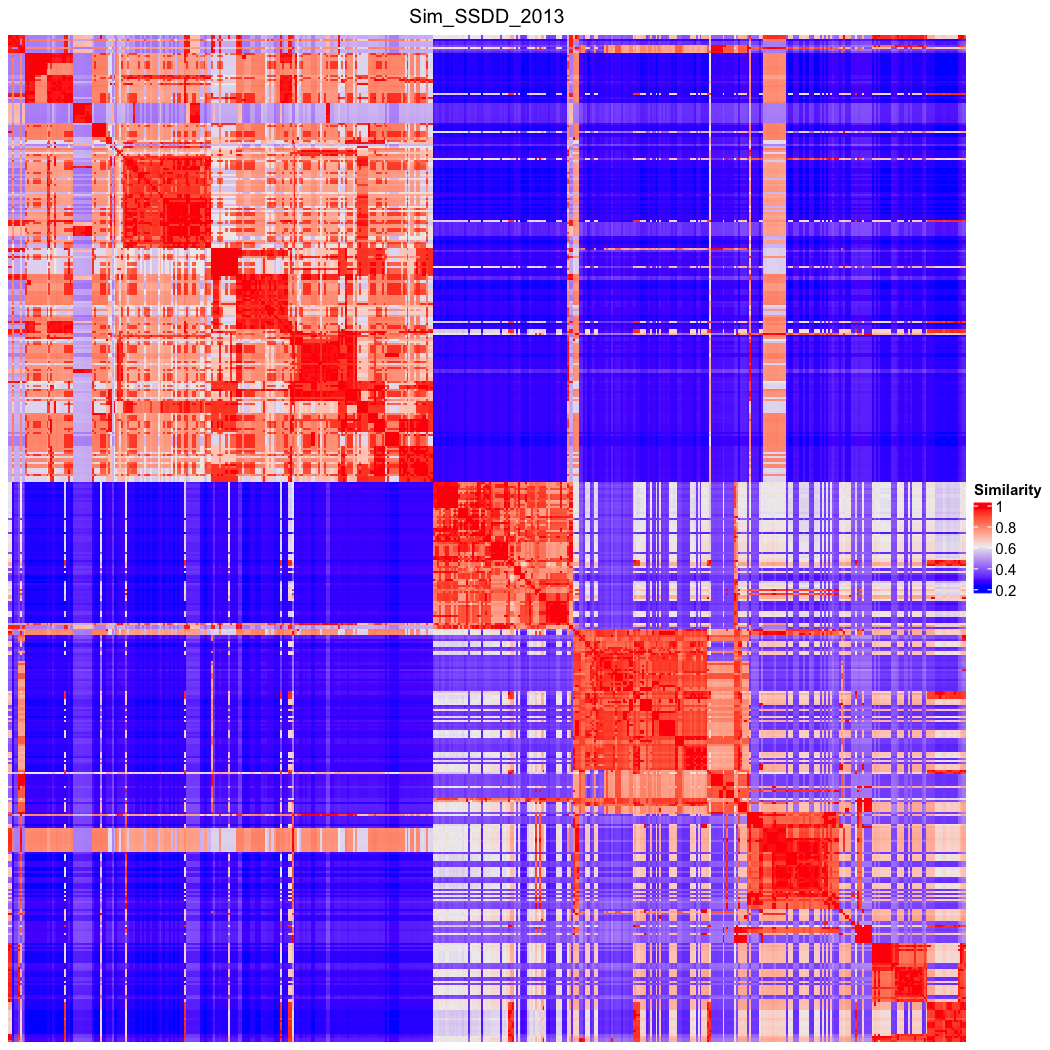

### go_bp_random_500_sim_Sim_Stojanovic_2001.png

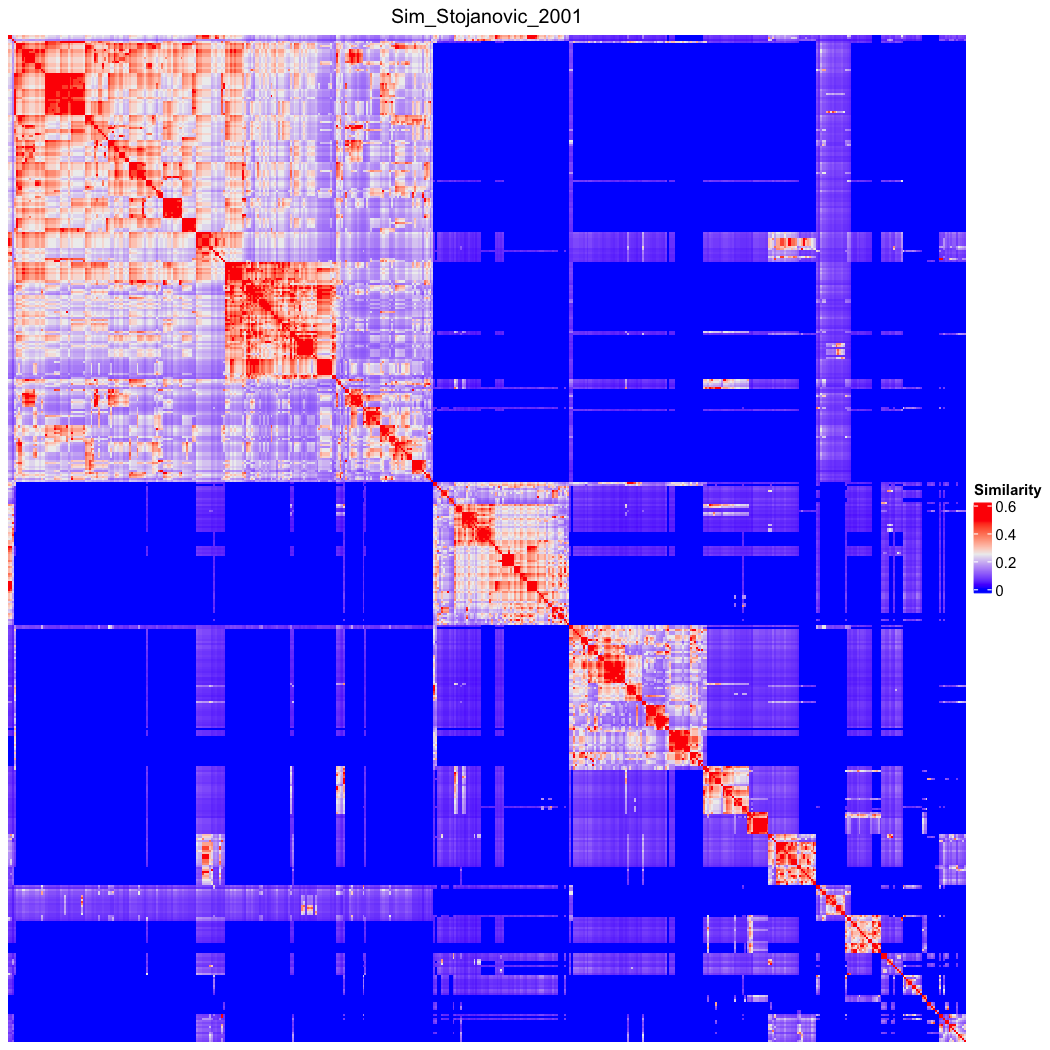

### go_bp_random_500_sim_Sim_Stojanovic_2001_Lin_order.png

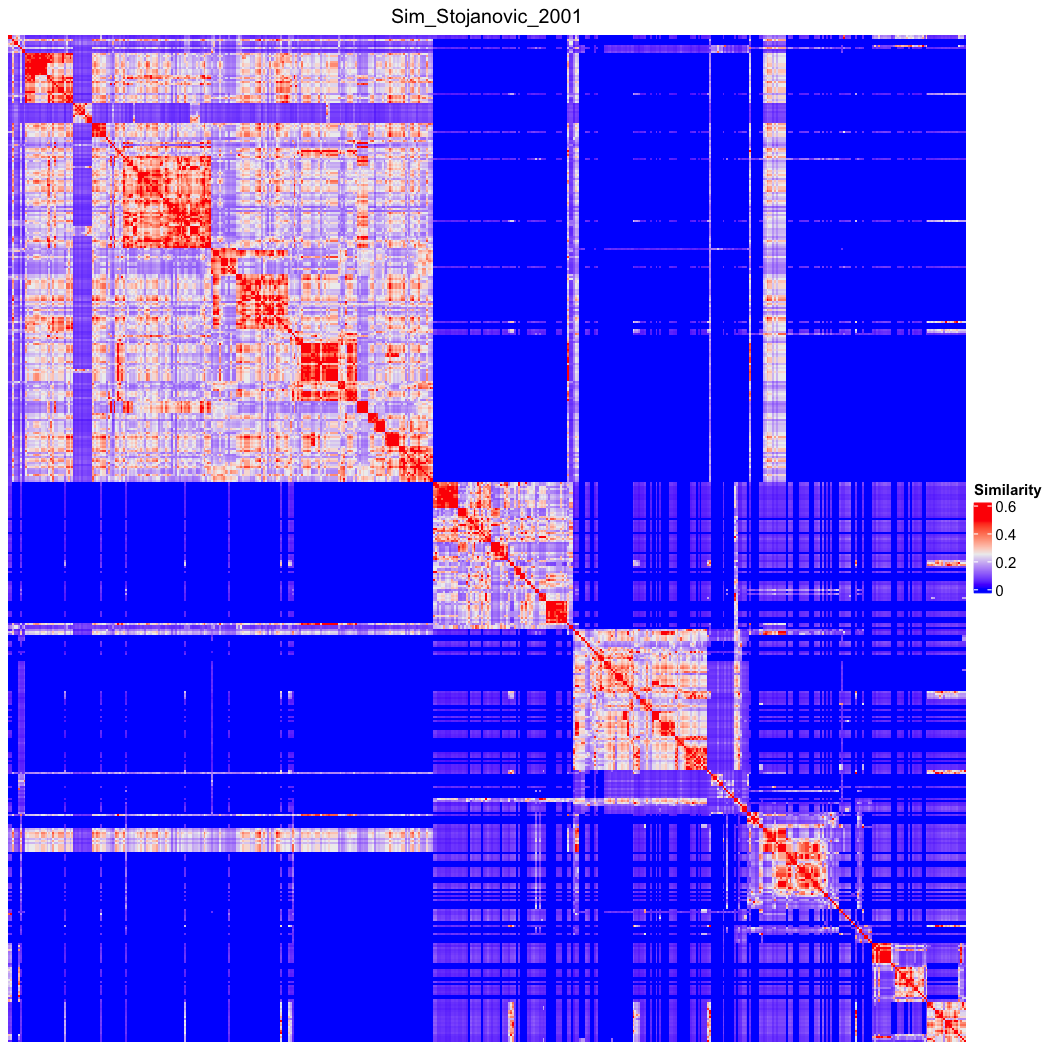

### go_bp_random_500_sim_Sim_universal.png

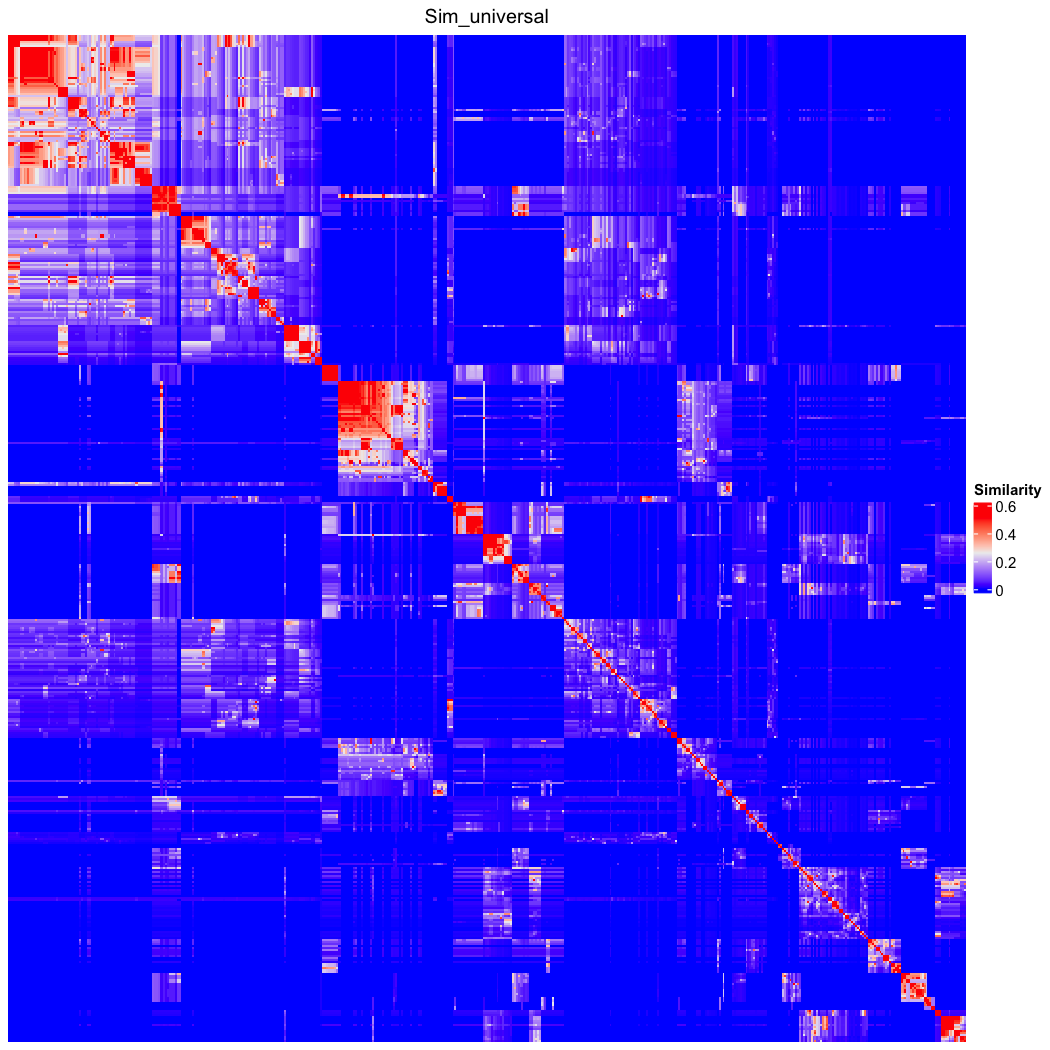

### go_bp_random_500_sim_Sim_universal_Lin_order.png

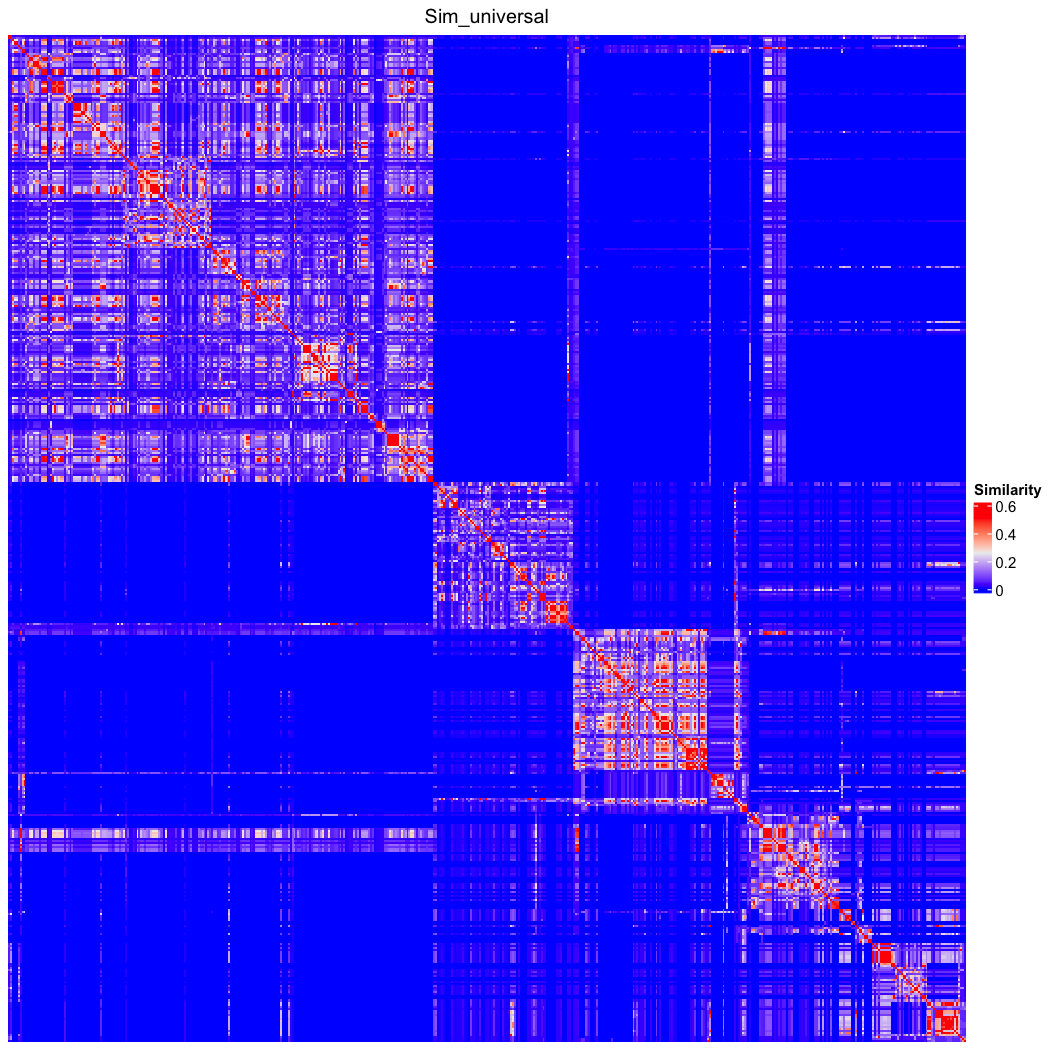

### go_bp_random_500_sim_Sim_Wang_2007.png

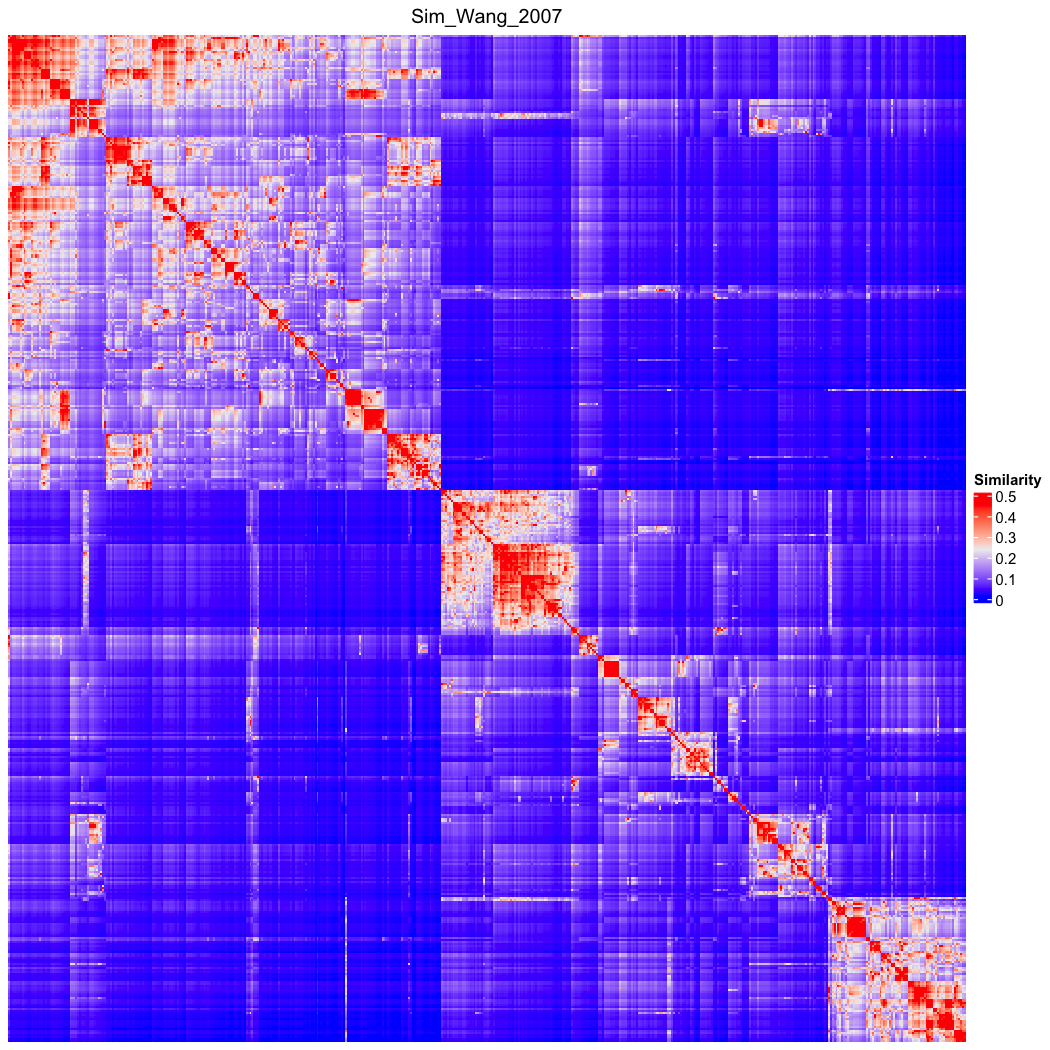

### go_bp_random_500_sim_Sim_Wang_2007_Lin_order.png

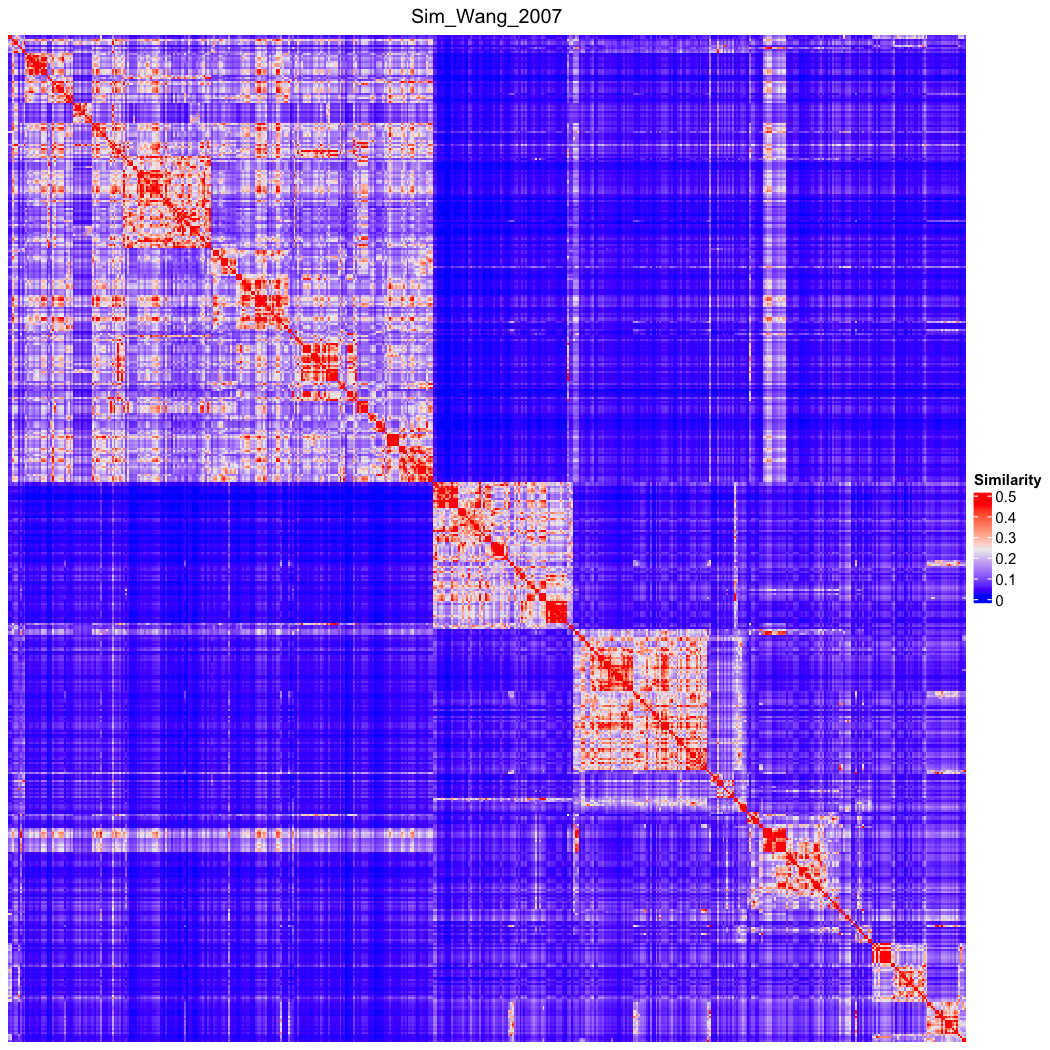

### go_bp_random_500_sim_Sim_Wang_edge_2012.png

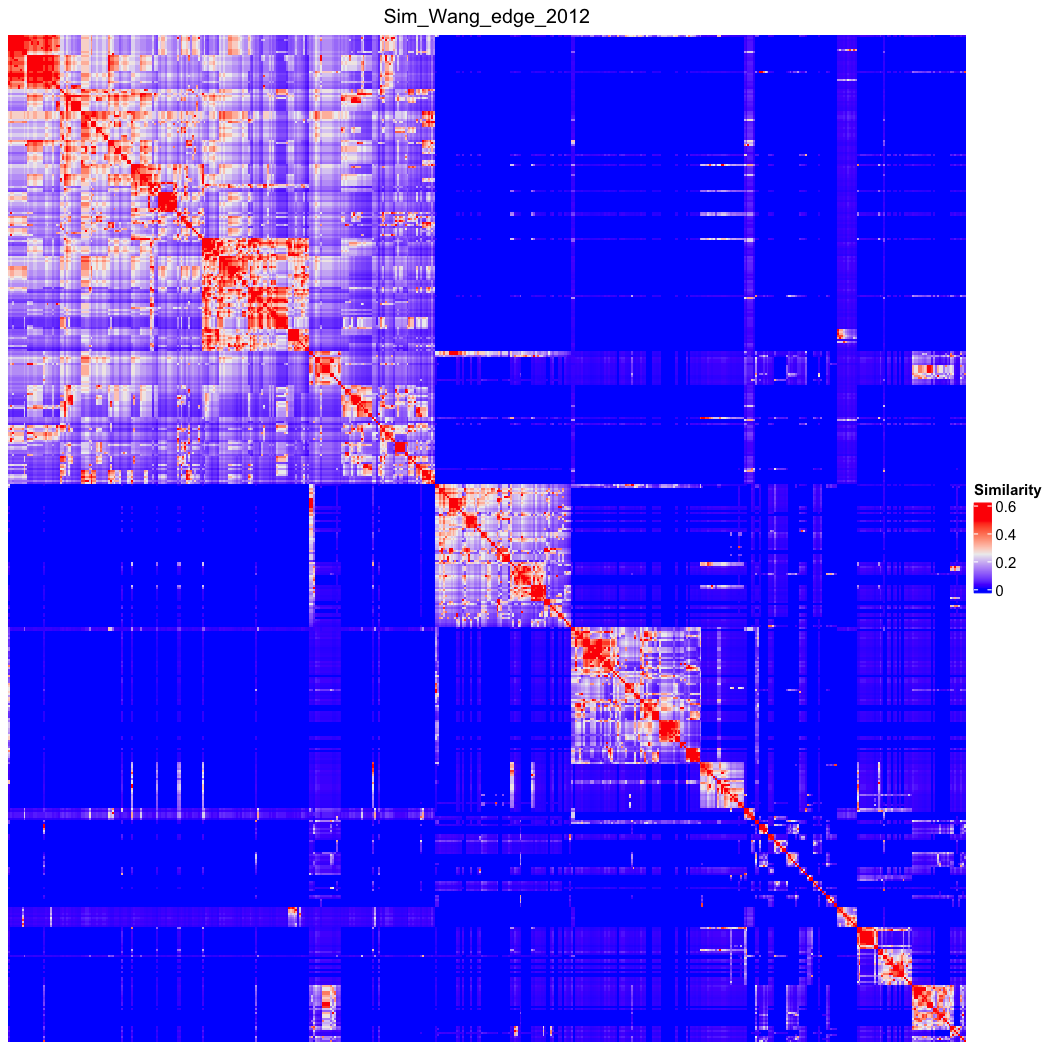

### go_bp_random_500_sim_Sim_Wang_edge_2012_Lin_order.png

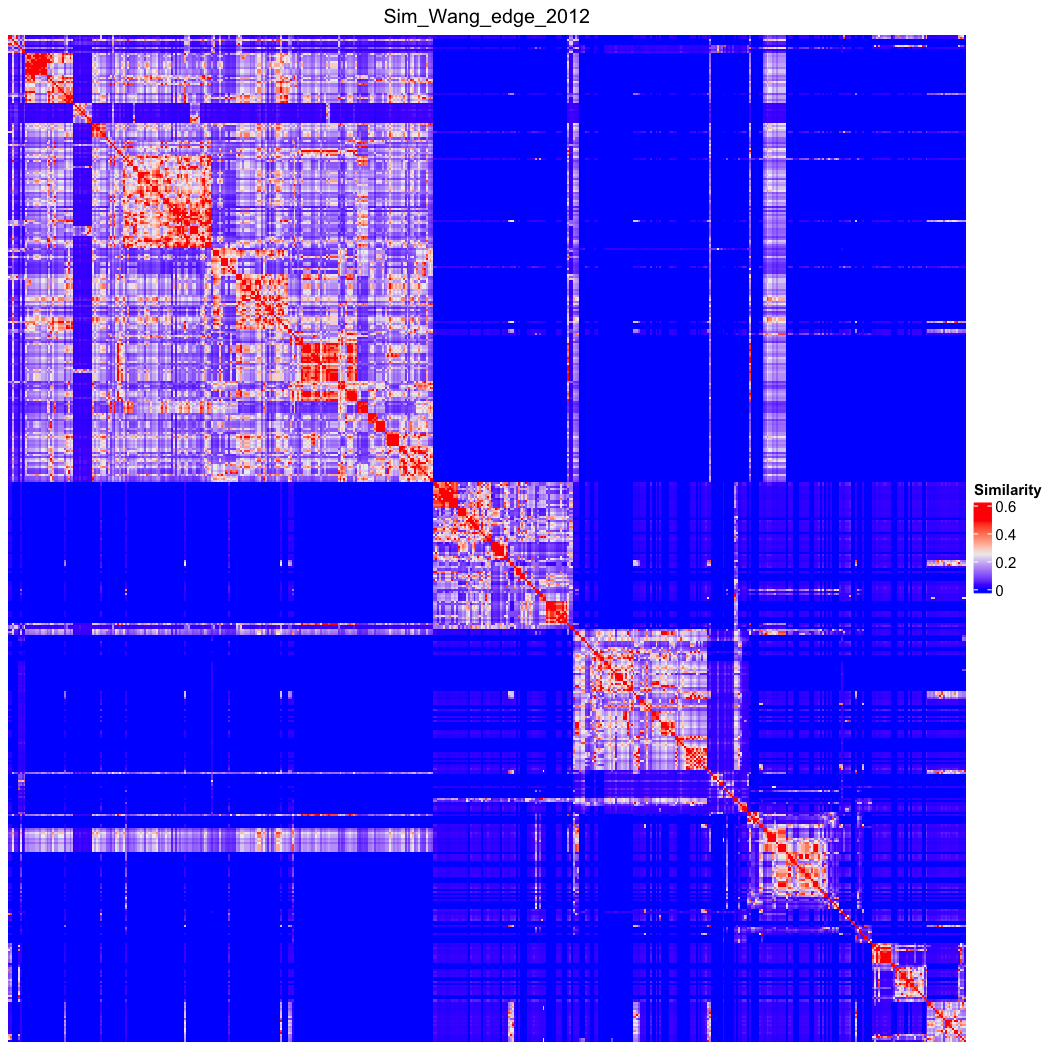

### go_bp_random_500_sim_Sim_WP_1994.png

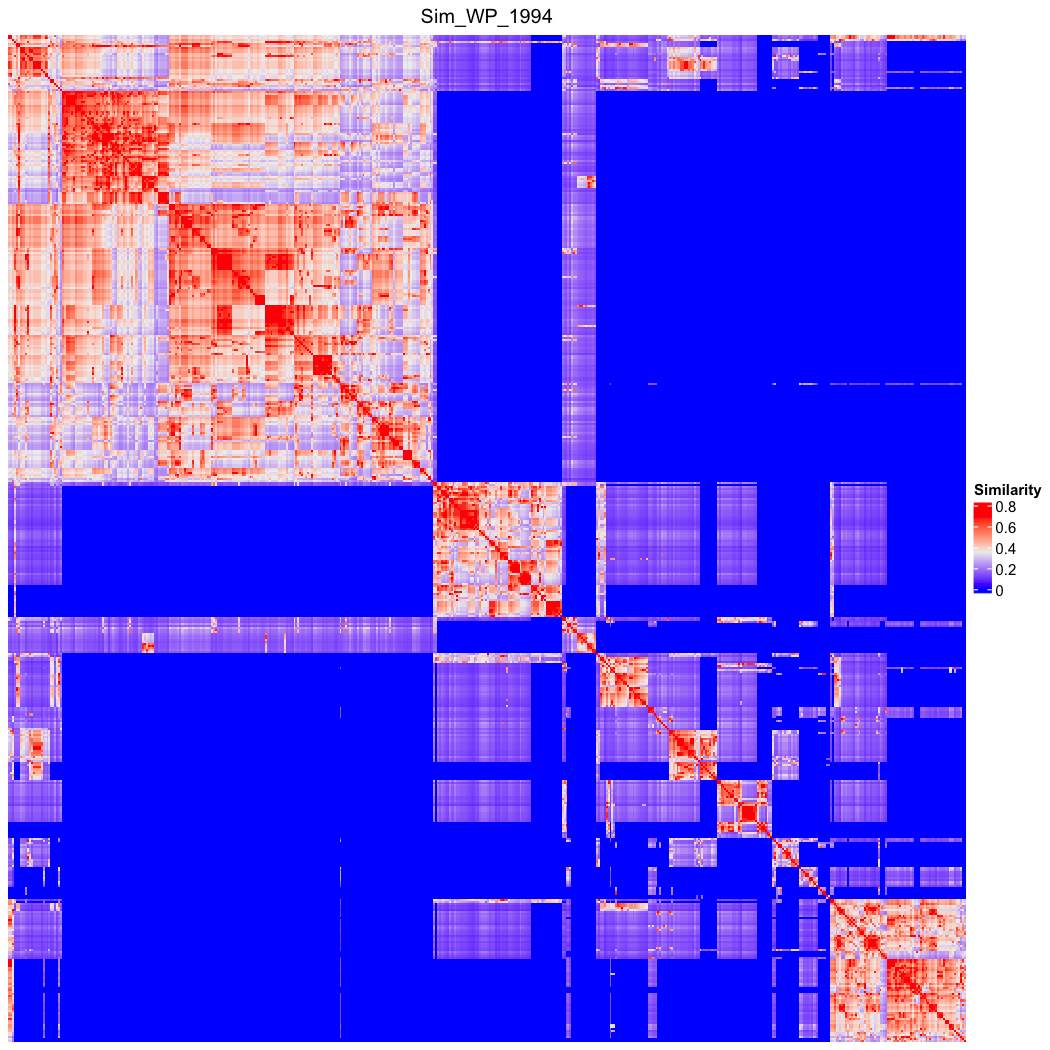

### go_bp_random_500_sim_Sim_WP_1994_Lin_order.png

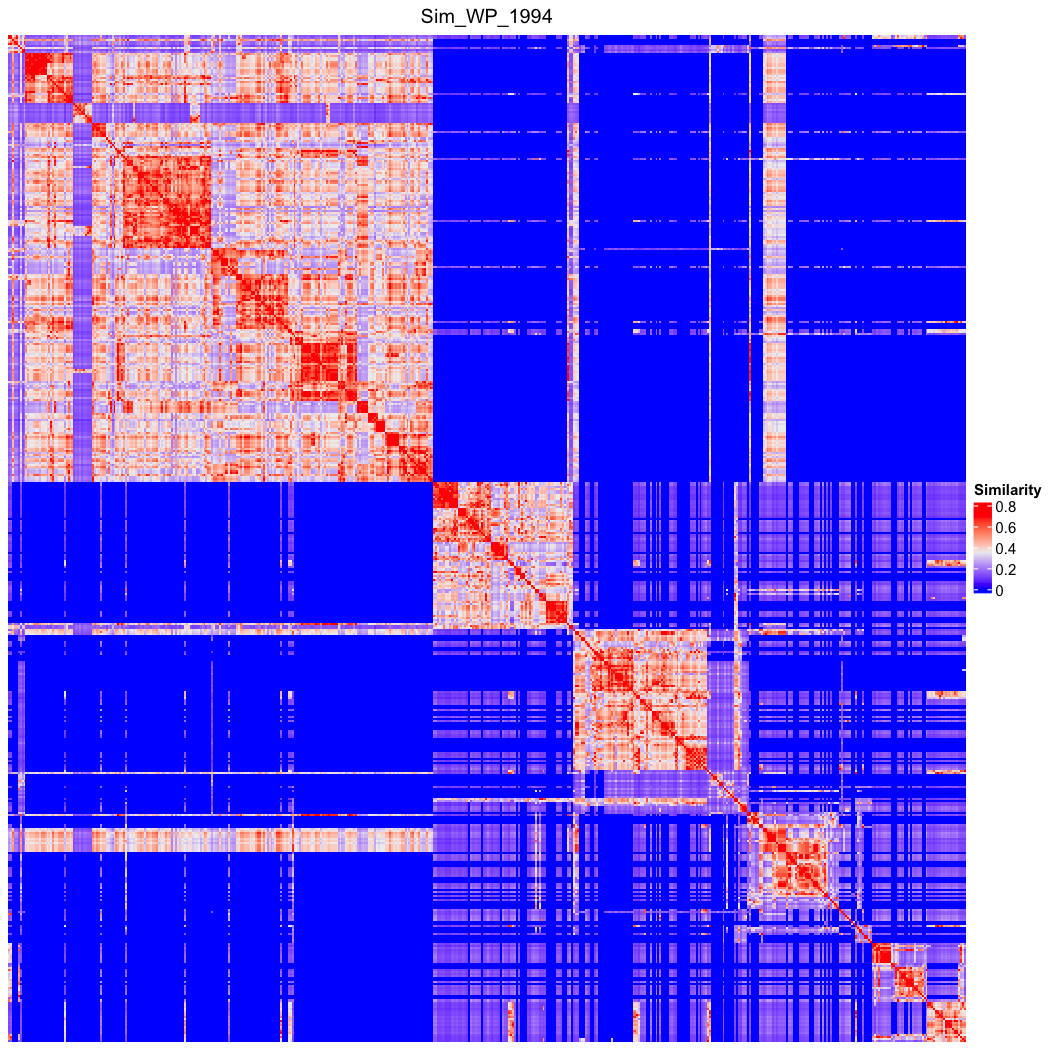

### go_bp_random_500_sim_Sim_XGraSM_2013.png

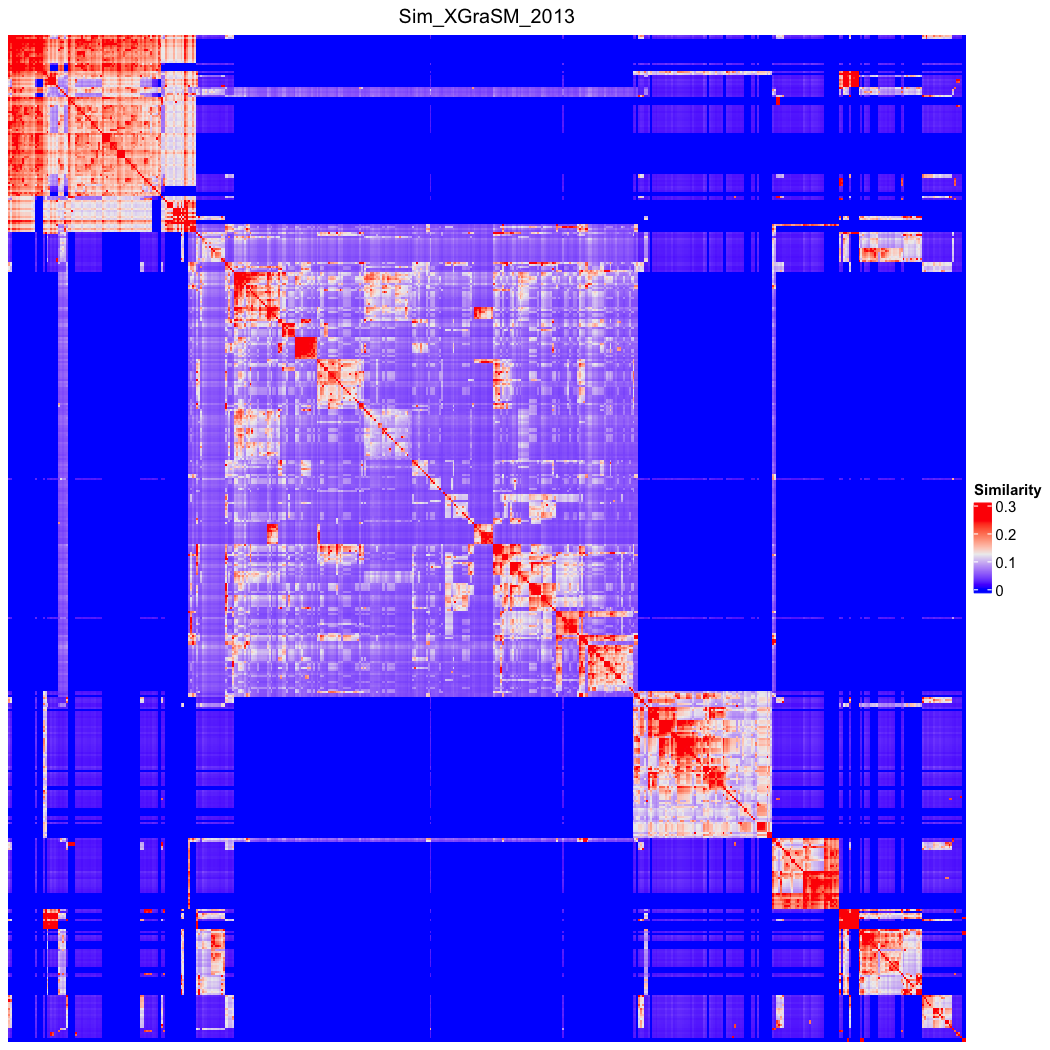

### go_bp_random_500_sim_Sim_XGraSM_2013_Lin_order.png

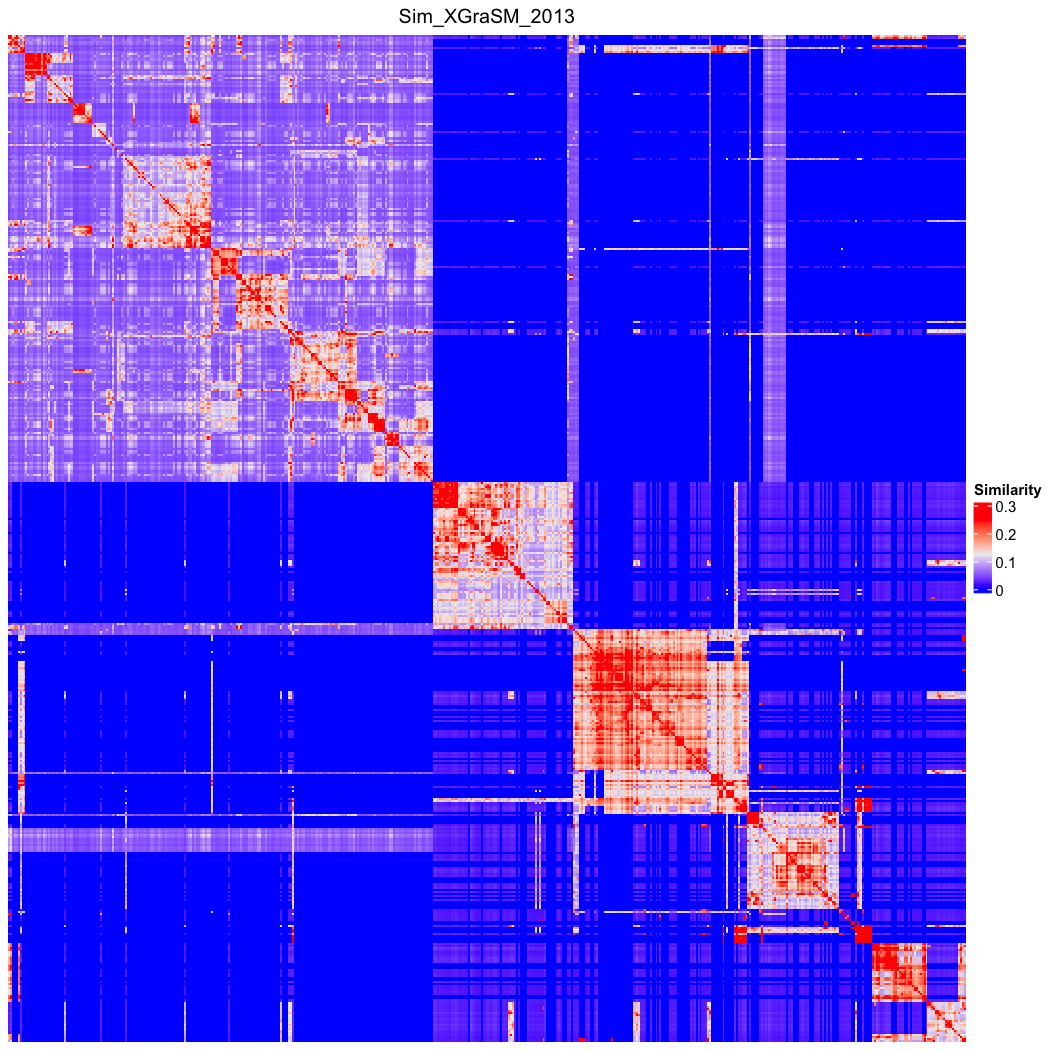

### go_bp_random_500_sim_Sim_Zhang_2006.png

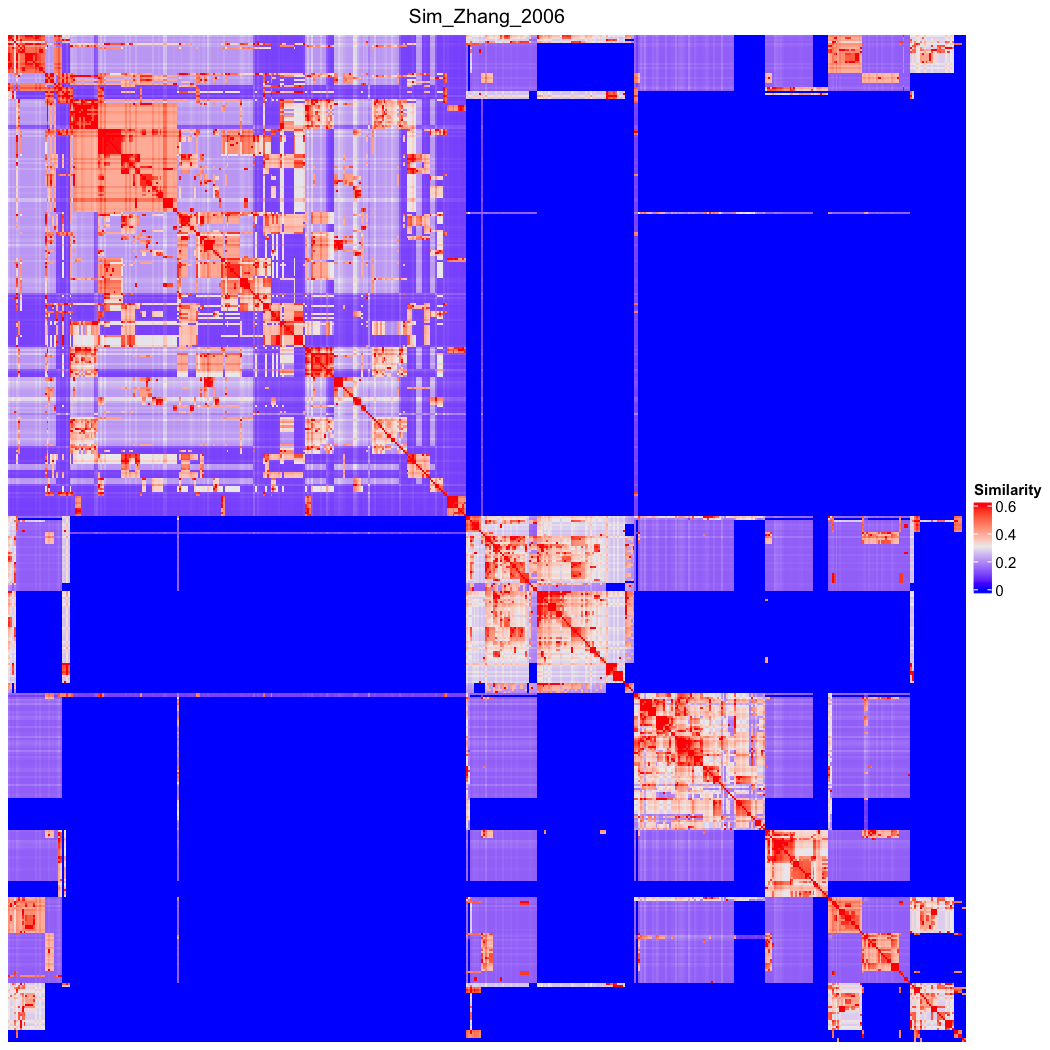

### go_bp_random_500_sim_Sim_Zhang_2006_Lin_order.png

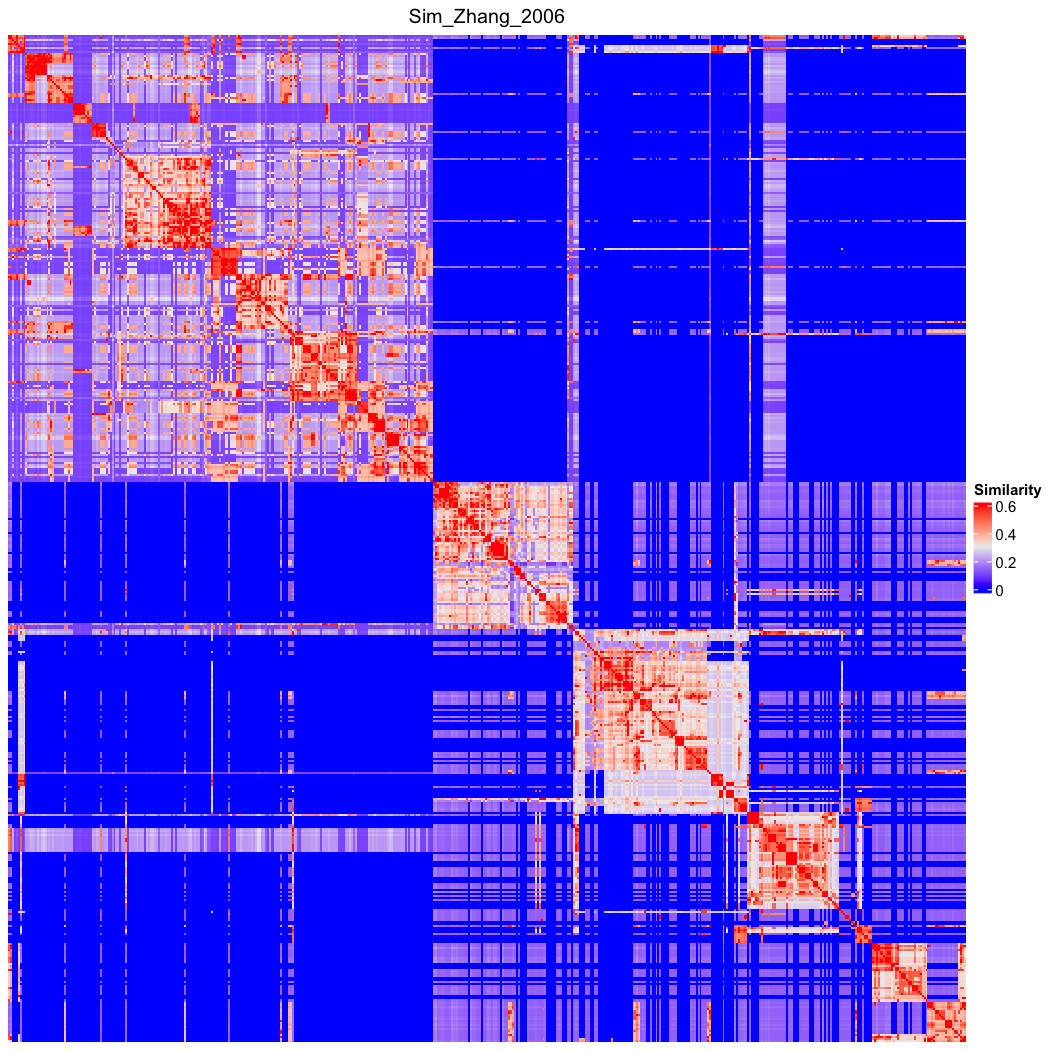

### go_bp_random_500_sim_Sim_Zhong_2002.png

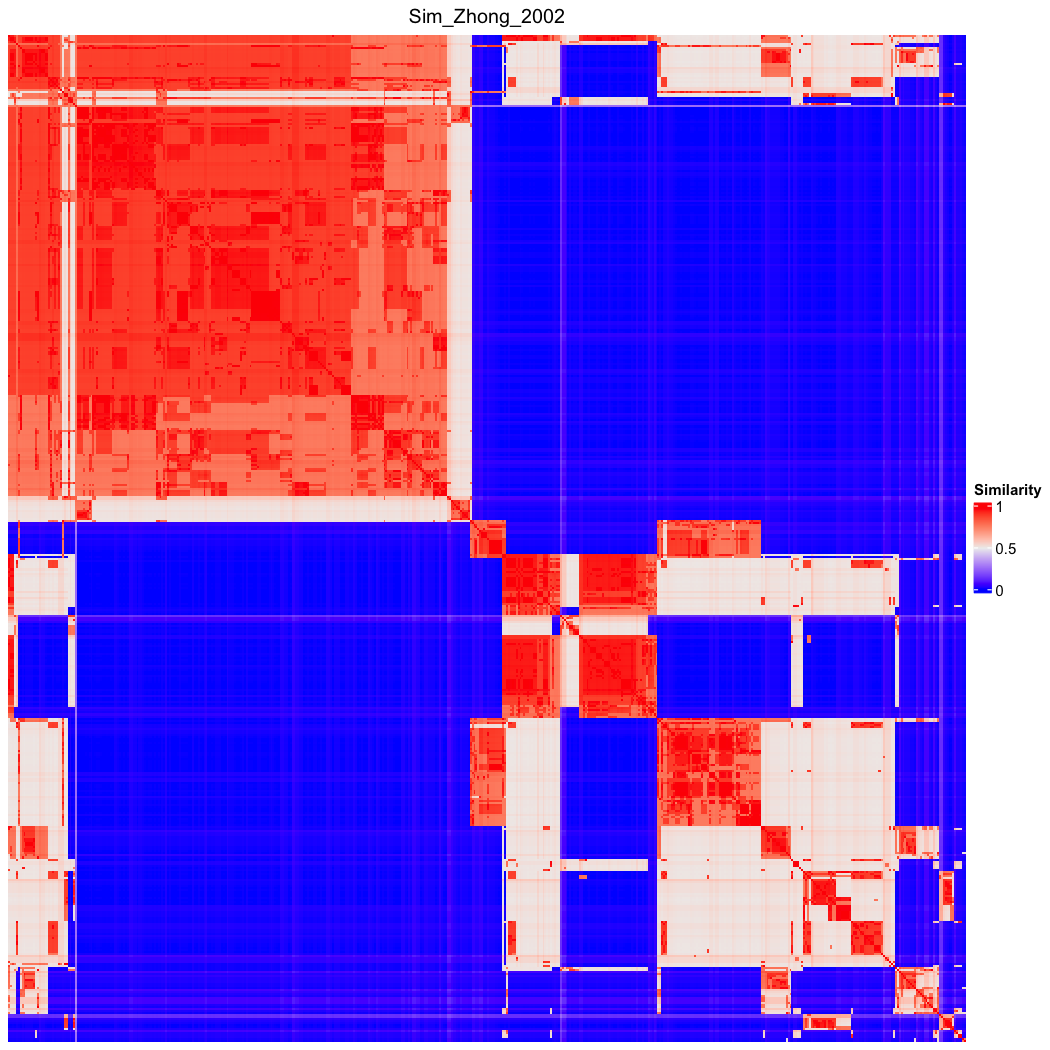

### go_bp_random_500_sim_Sim_Zhong_2002_Lin_order.png

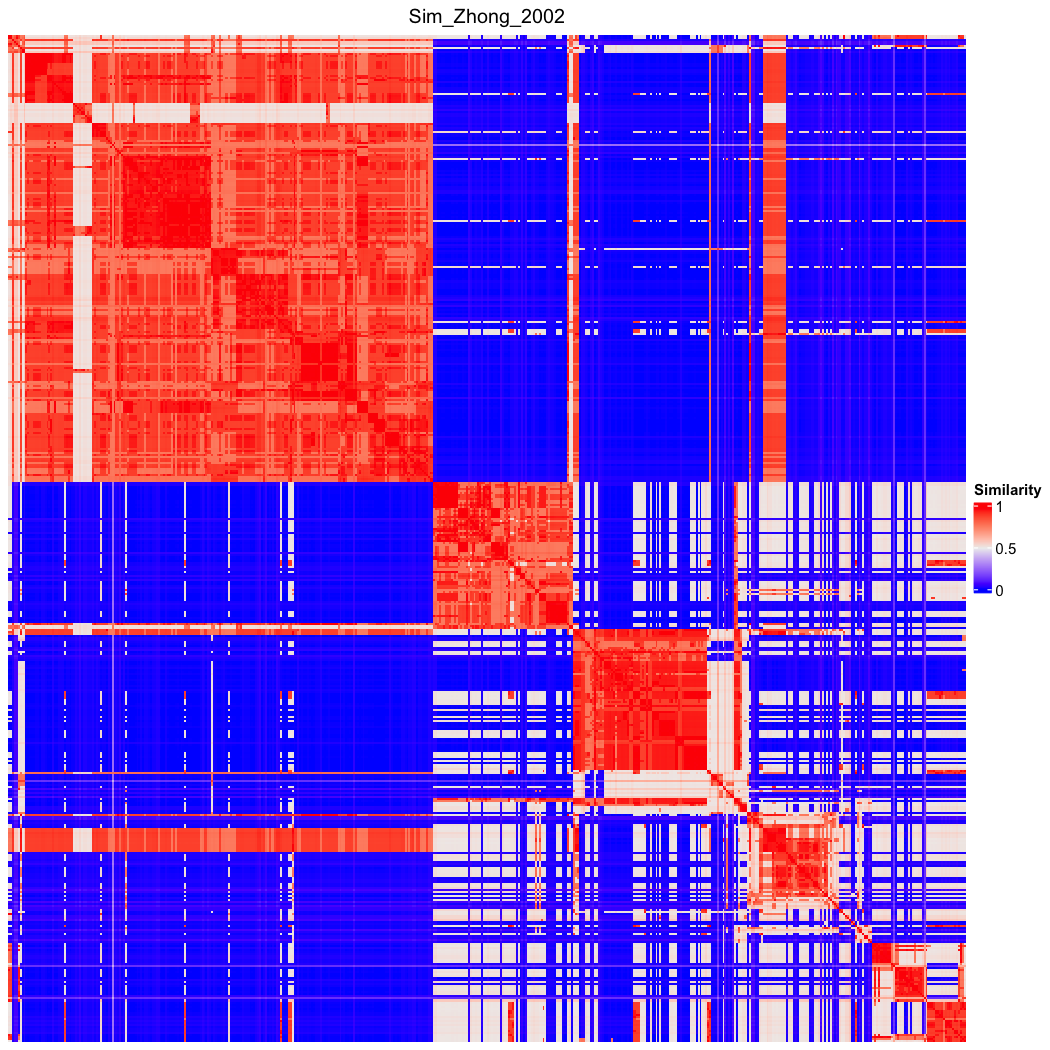

### OBOFoundry_bspo_heatmap.png

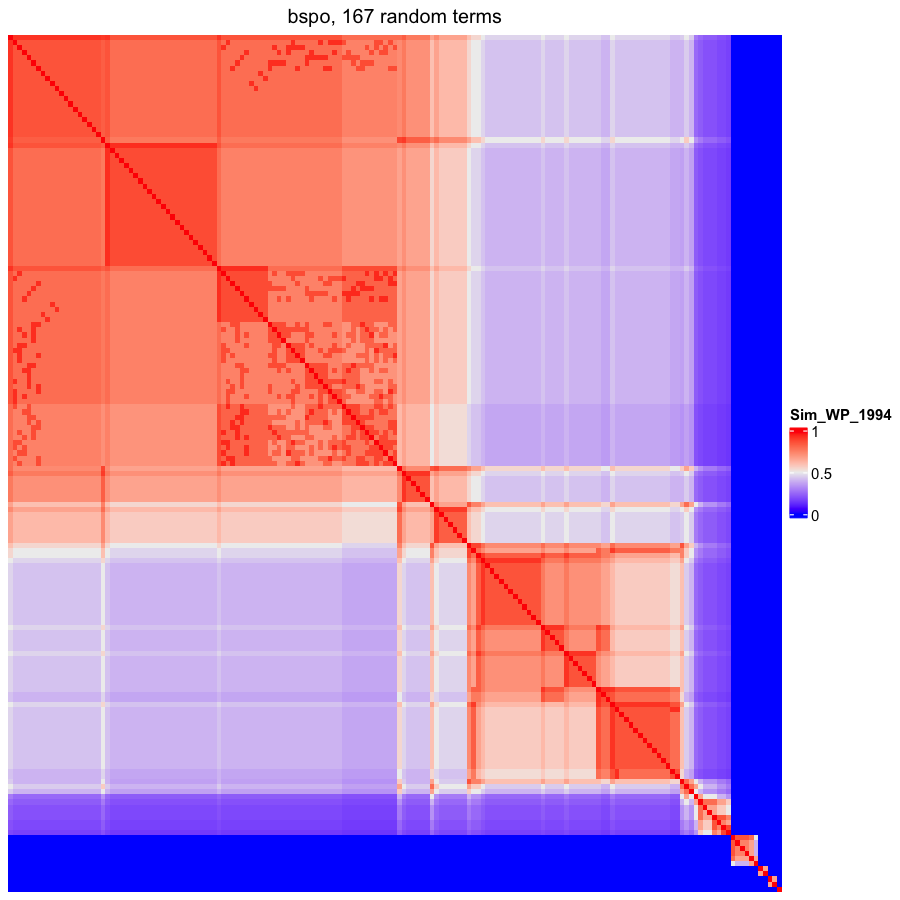

### OBOFoundry_ceph_runtime.png

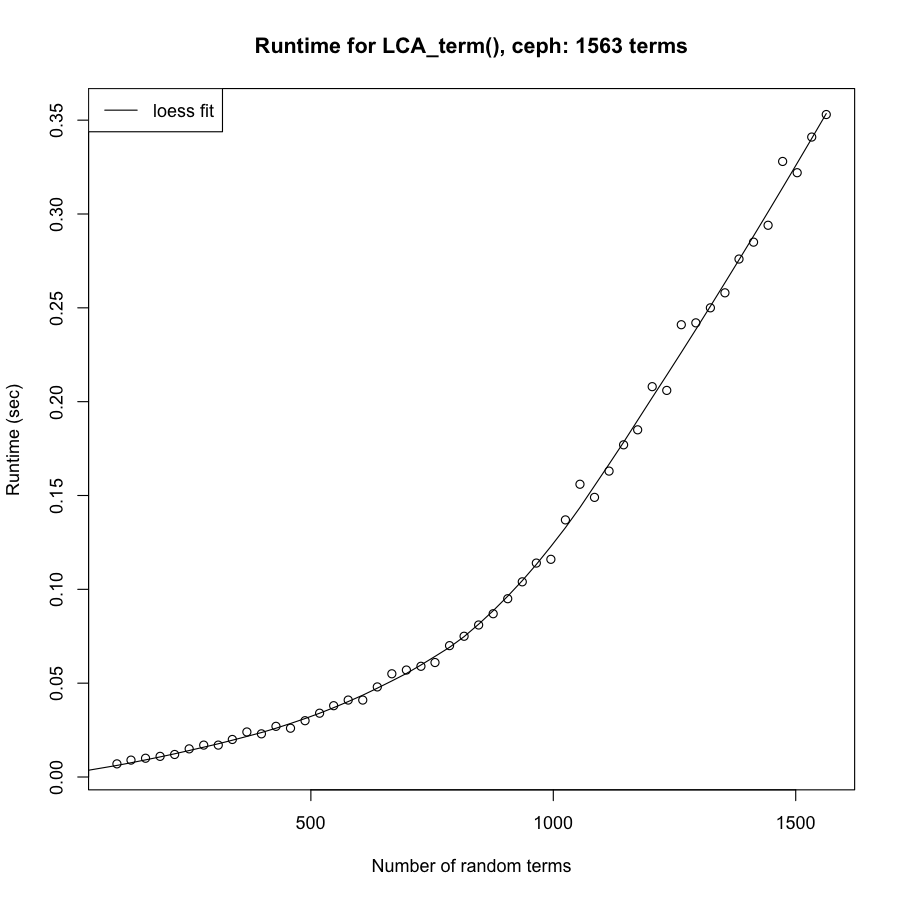

### OBOFoundry_cmo_heatmap.png

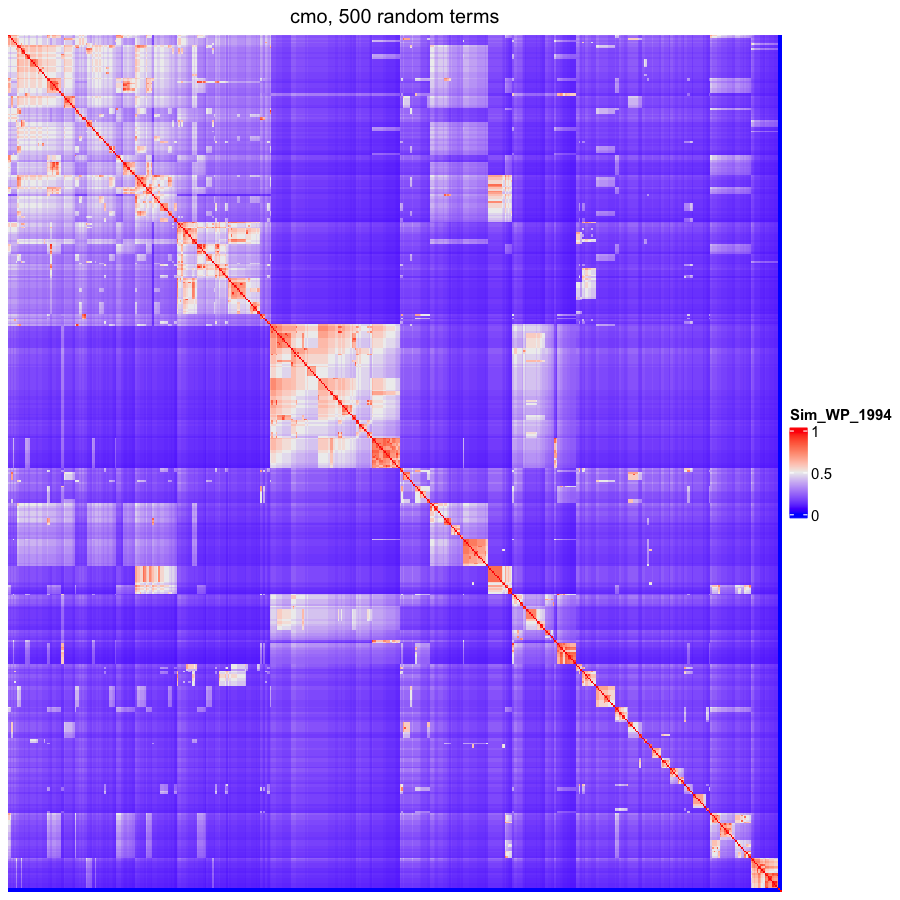

### OBOFoundry_emapa_runtime.png

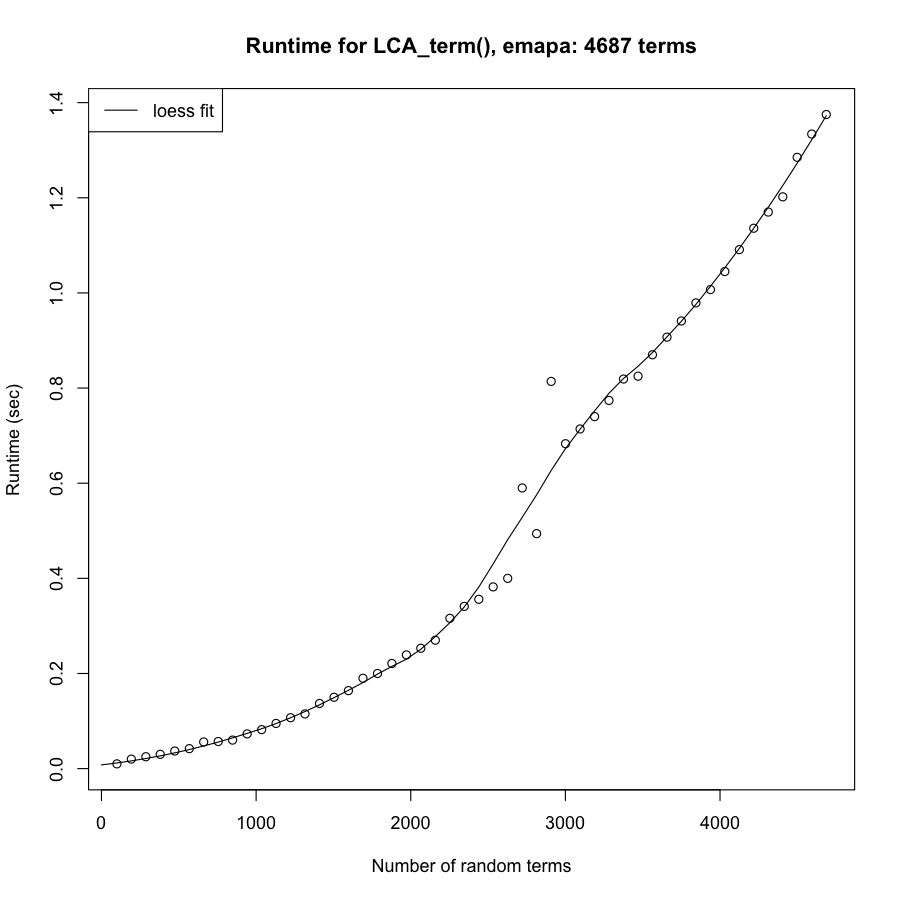

### OBOFoundry_flopo_runtime.png

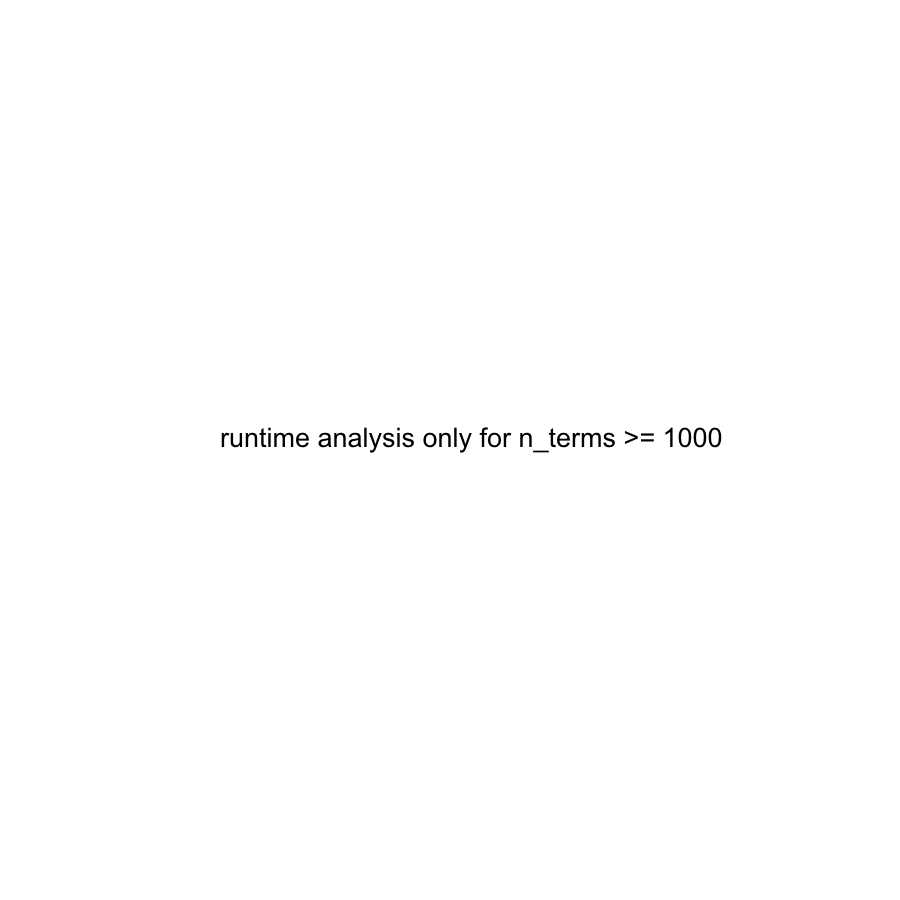

### OBOFoundry_fma_heatmap.png

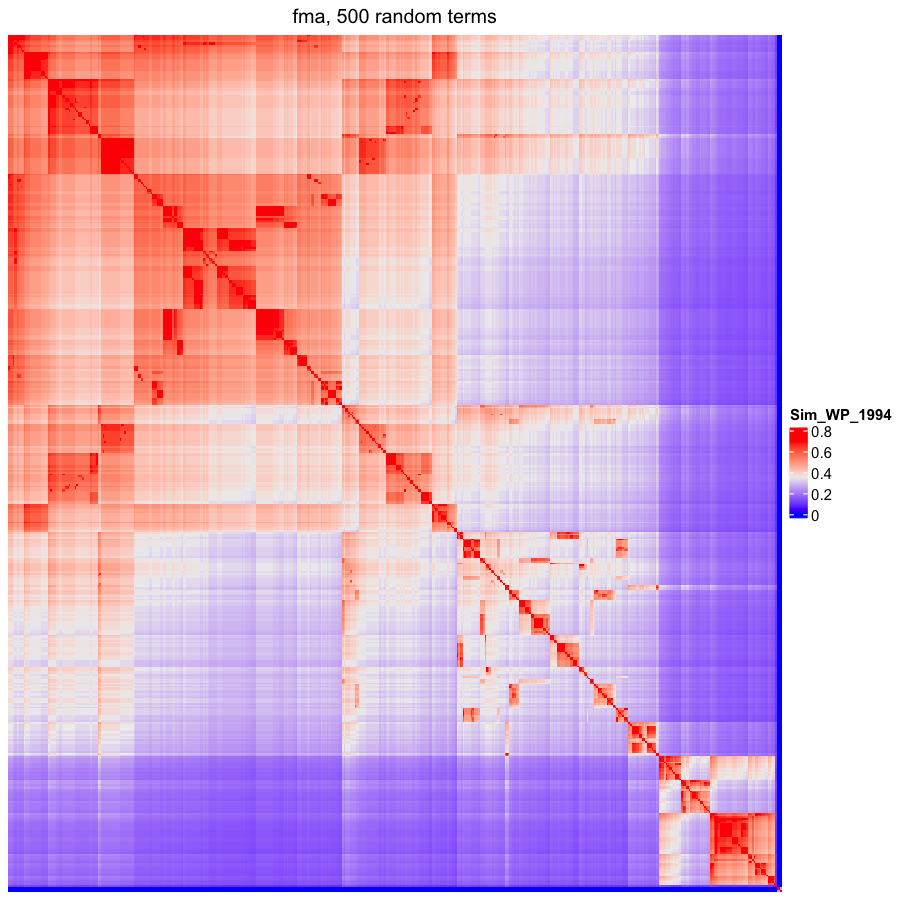

### OBOFoundry_mcro_heatmap.png

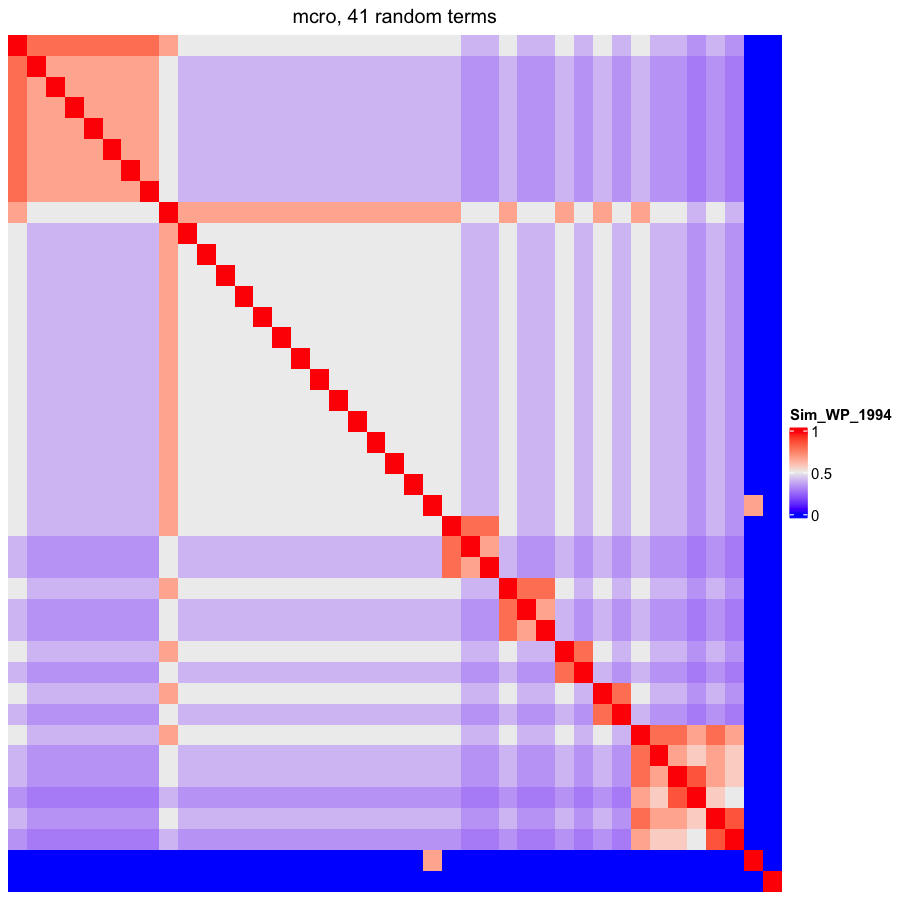

### OBOFoundry_mf_heatmap.png

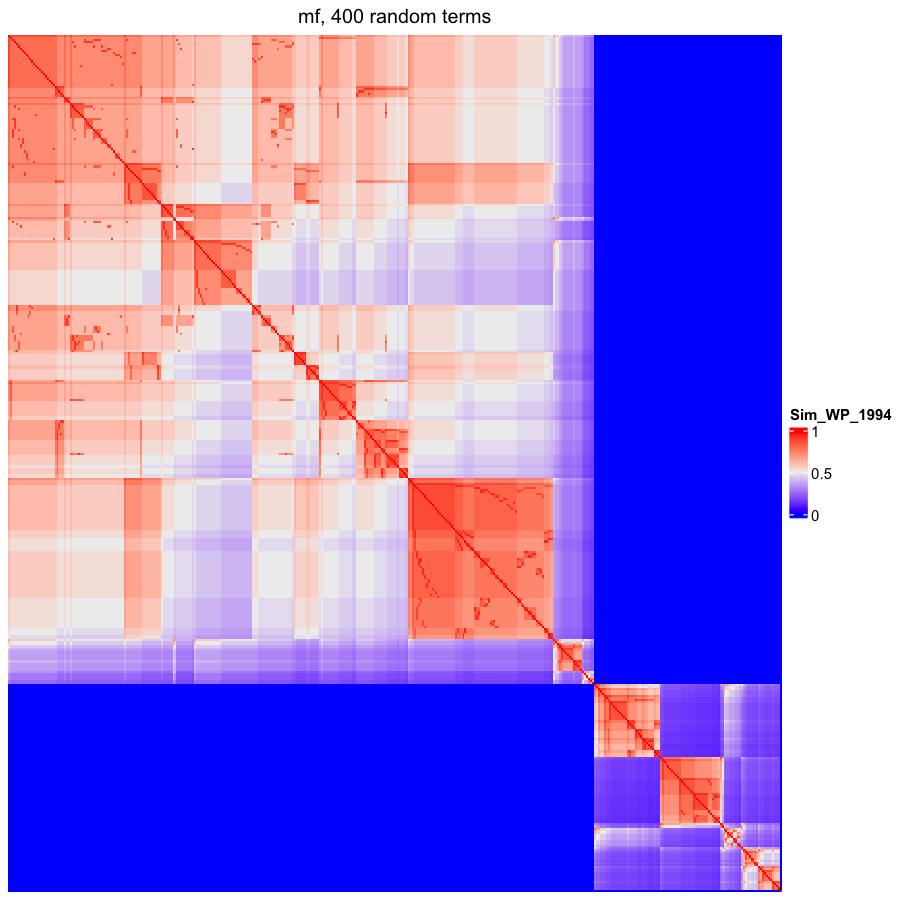

### OBOFoundry_mmo.png

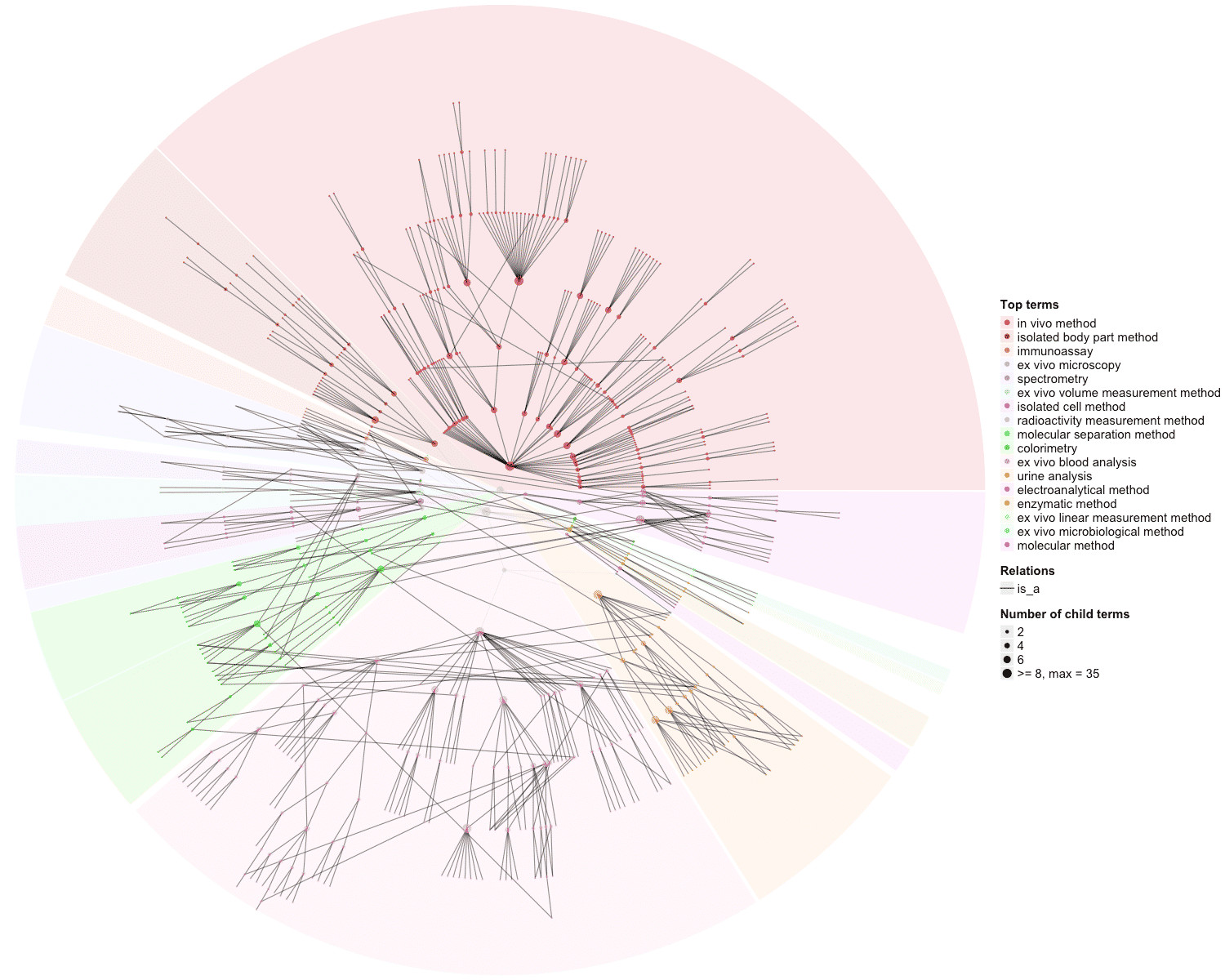

### OBOFoundry_mop.png

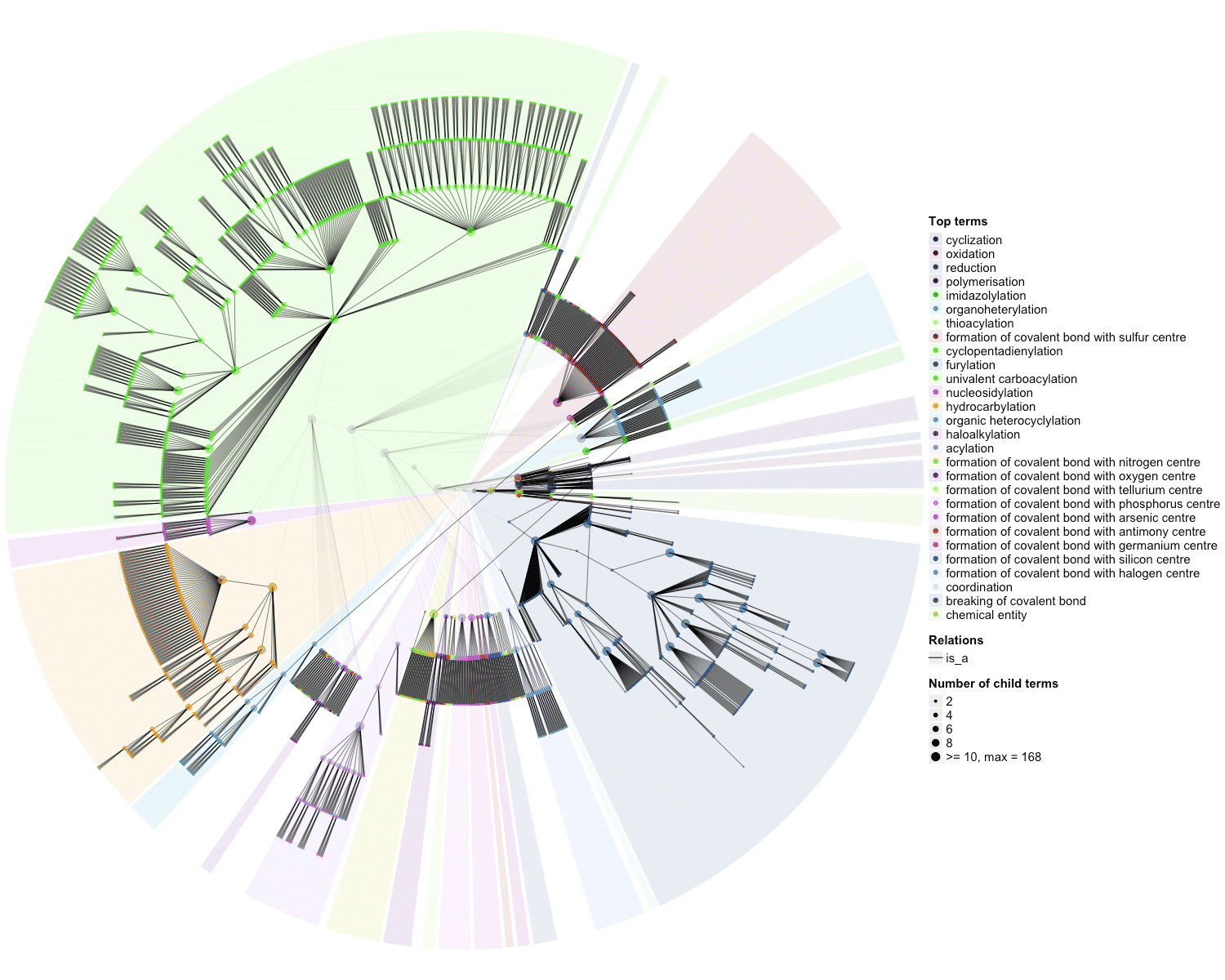

### OBOFoundry_obib.png

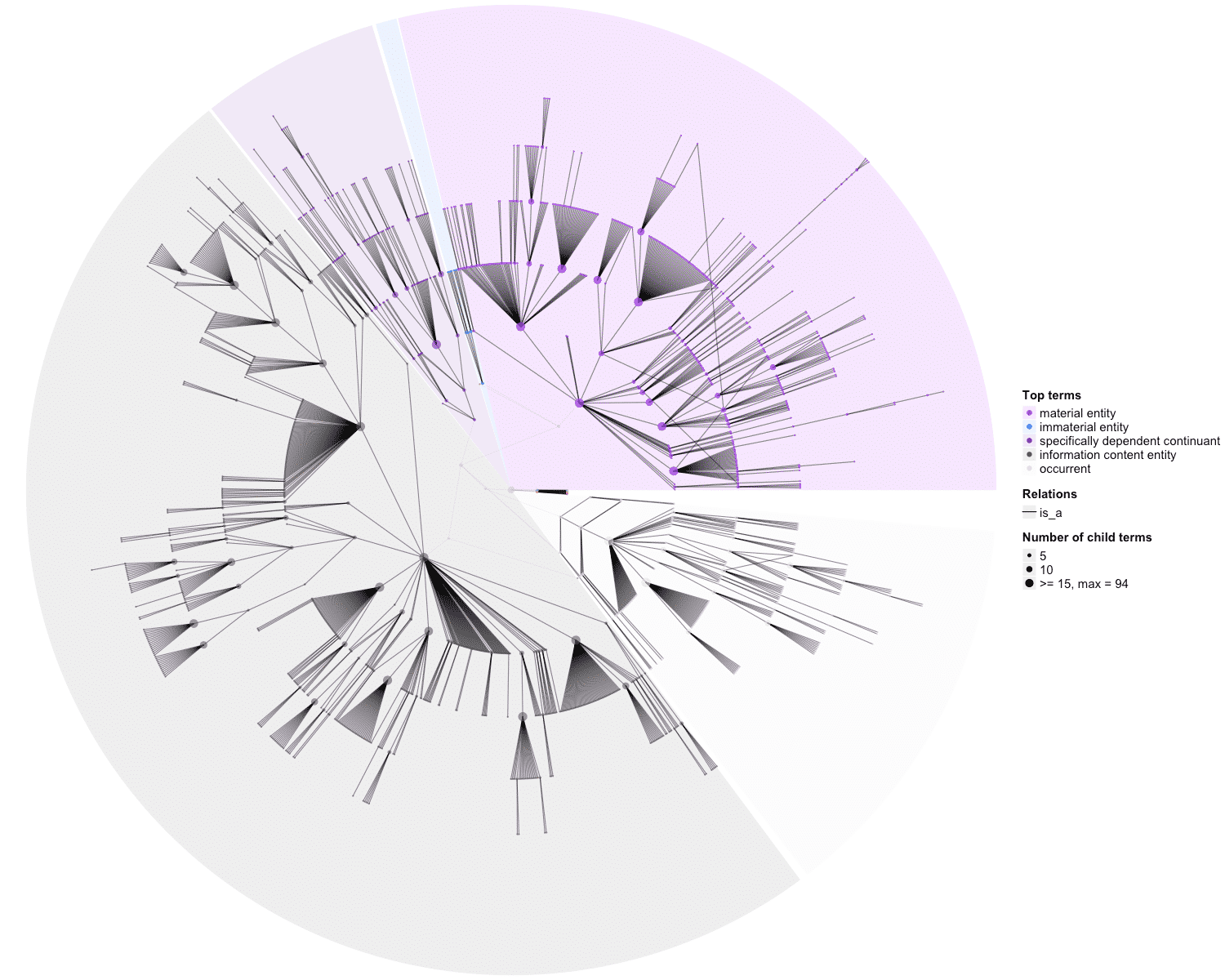

### OBOFoundry_one_heatmap.png

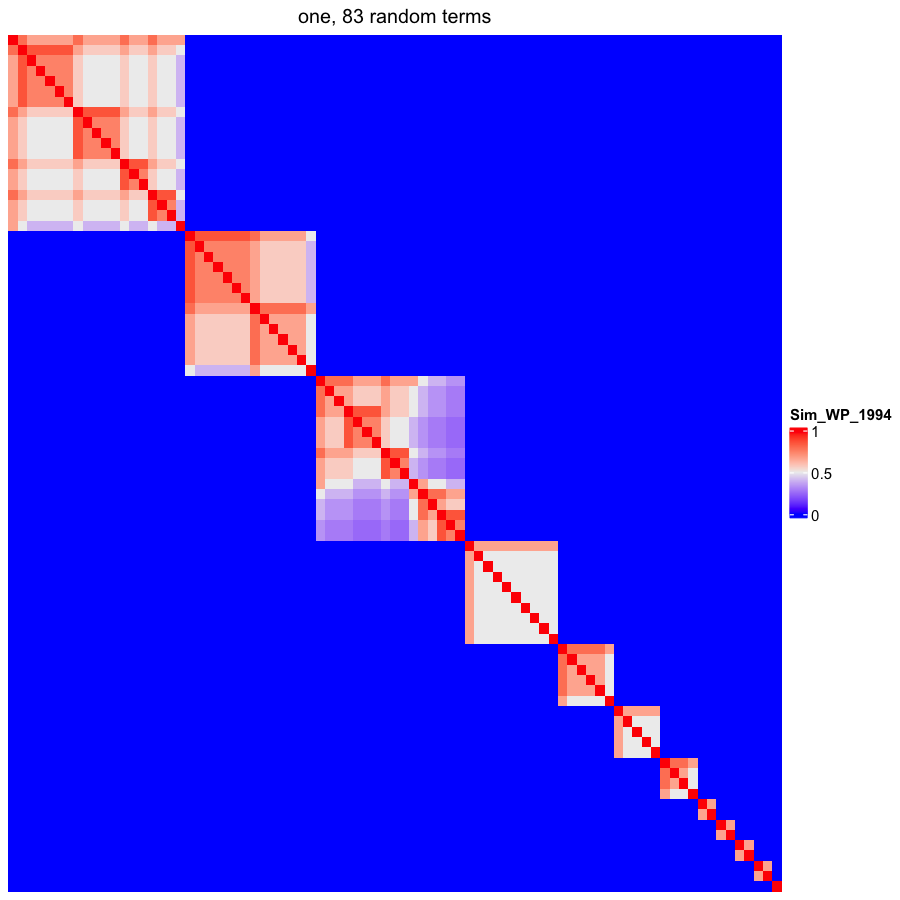

### OBOFoundry_oostt.png

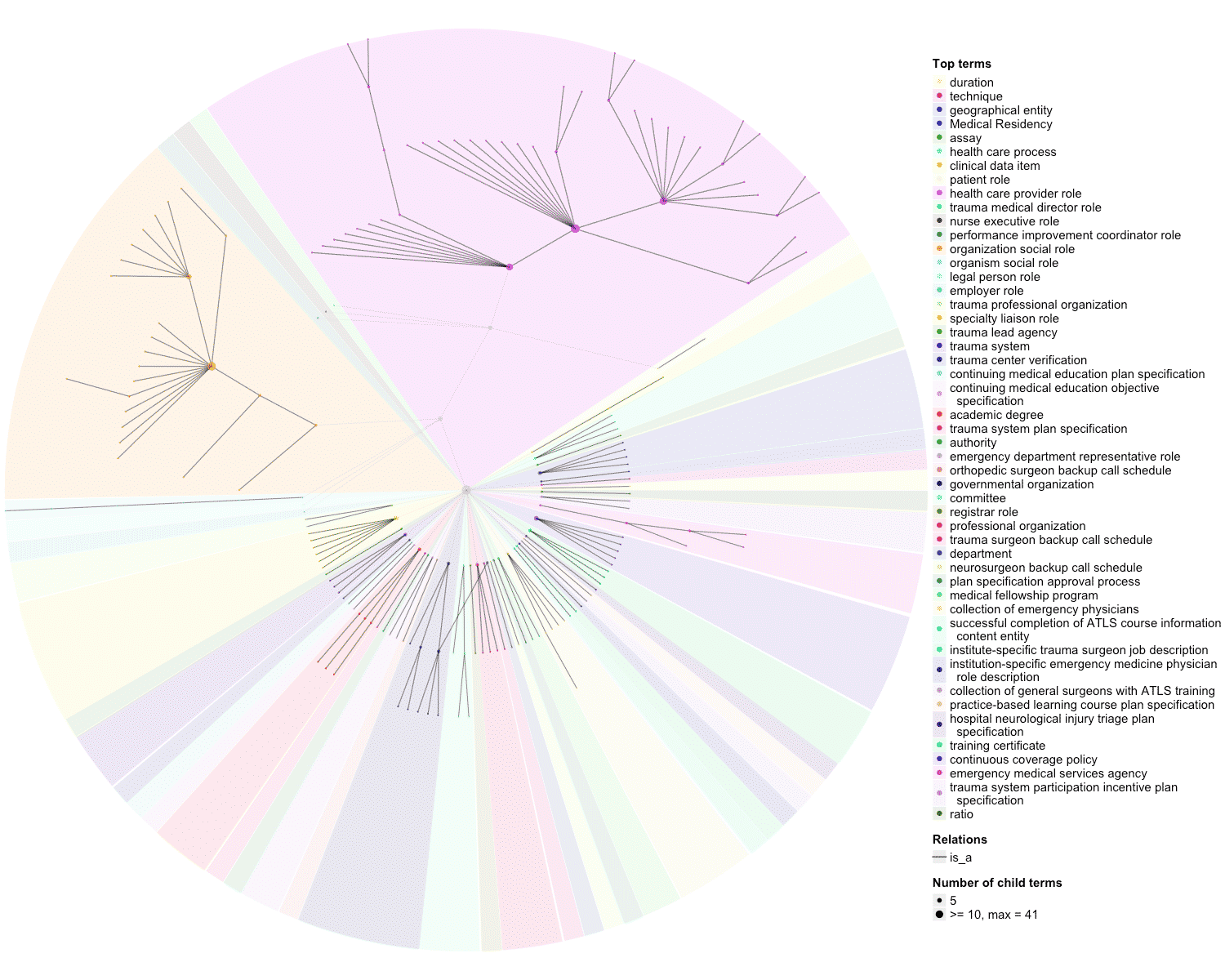

### OBOFoundry_opmi_runtime.png

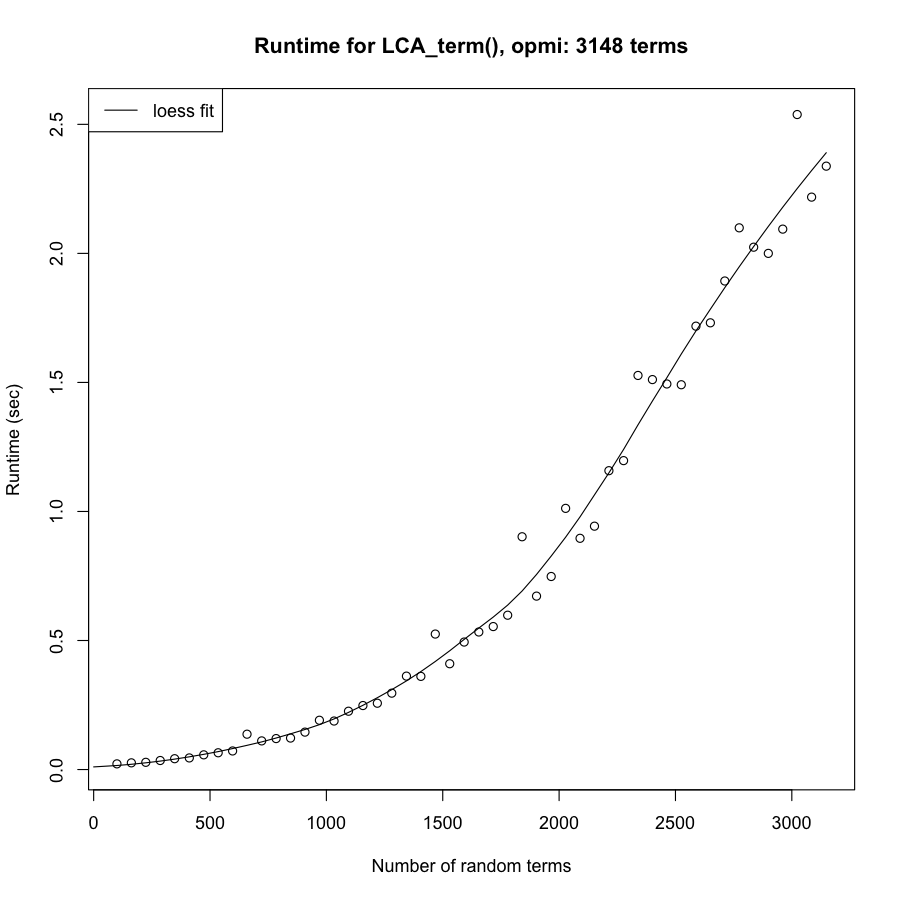

### OBOFoundry_peco.png

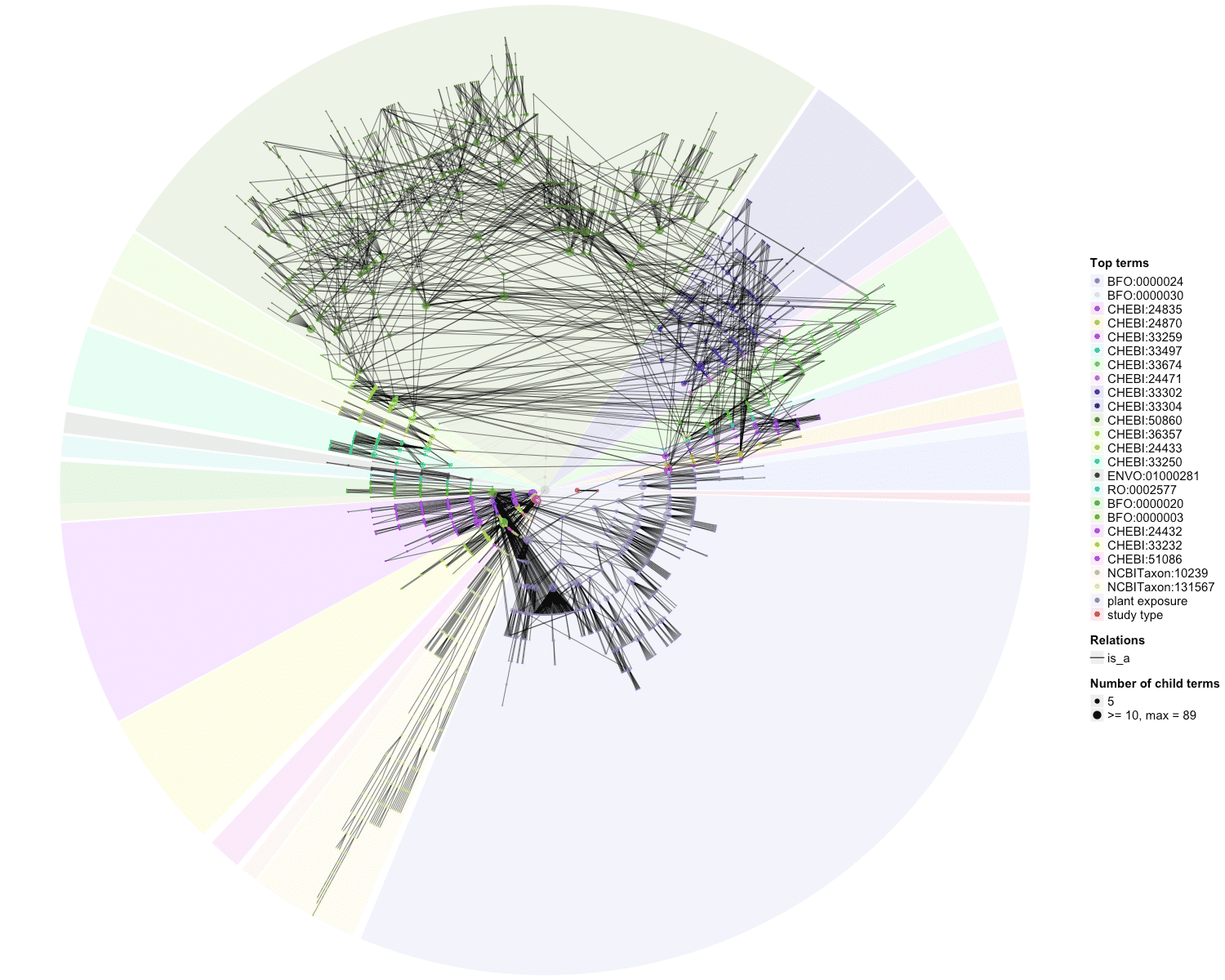

### OBOFoundry_planp_heatmap.png

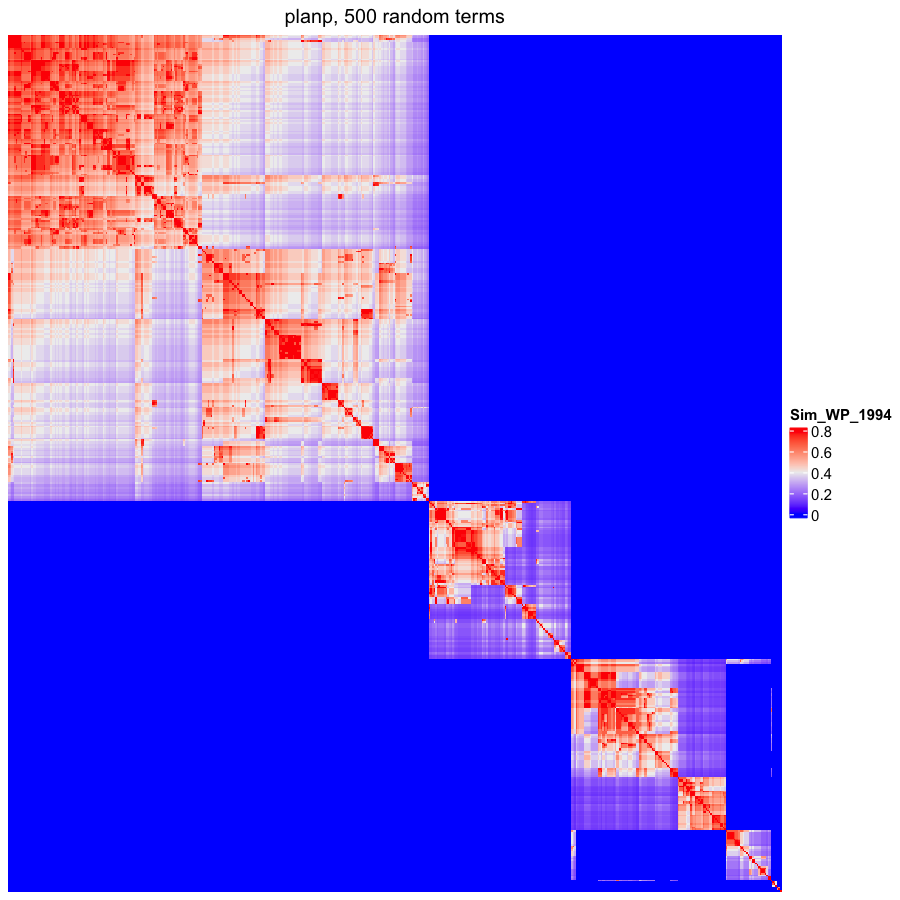

### OBOFoundry_po_runtime.png

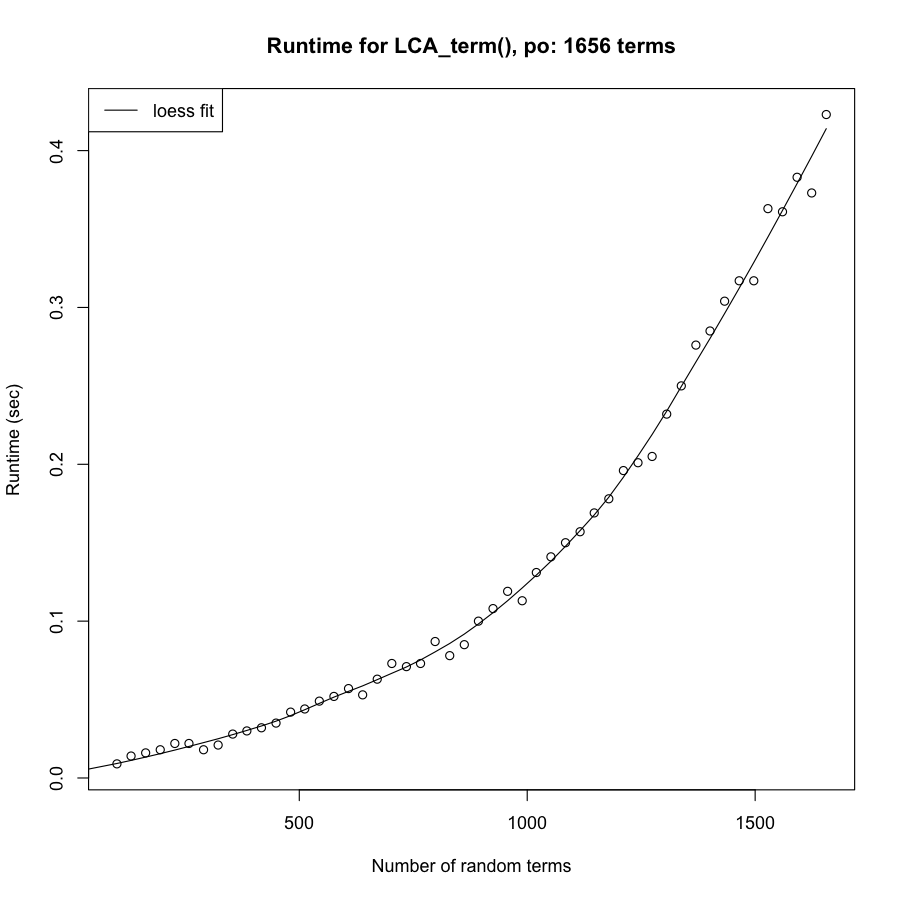

### OBOFoundry_ro_heatmap.png

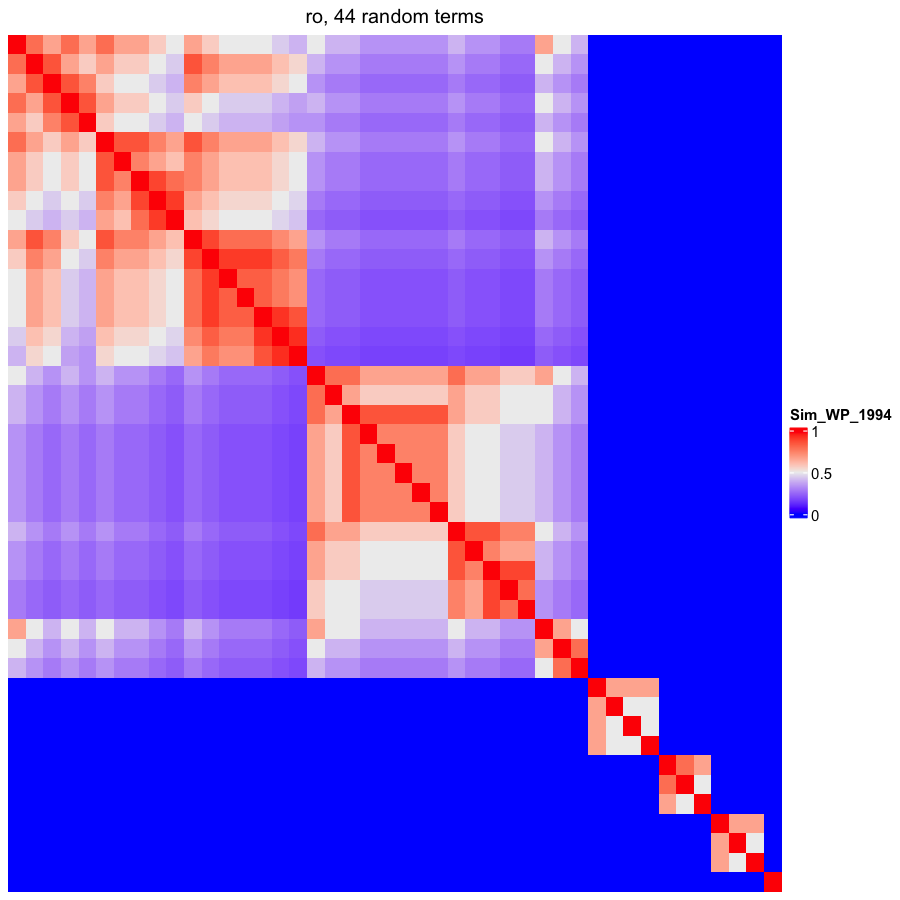

### OBOFoundry_rxno_heatmap.png

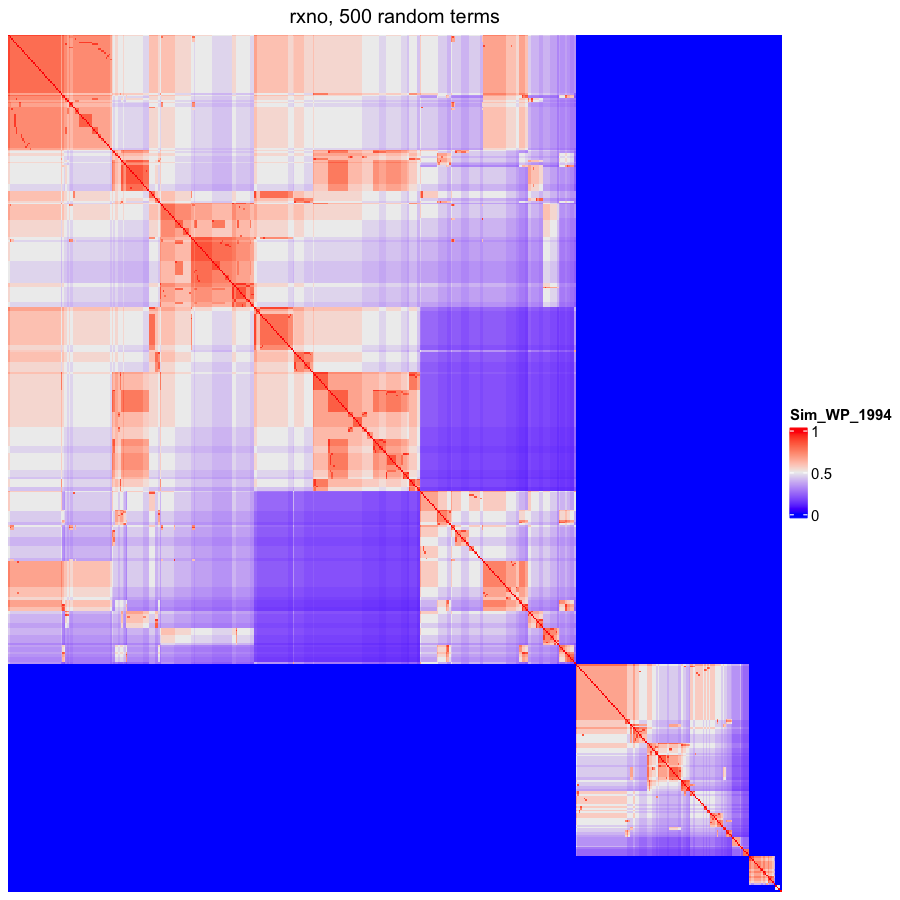

### OBOFoundry_swo_heatmap.png

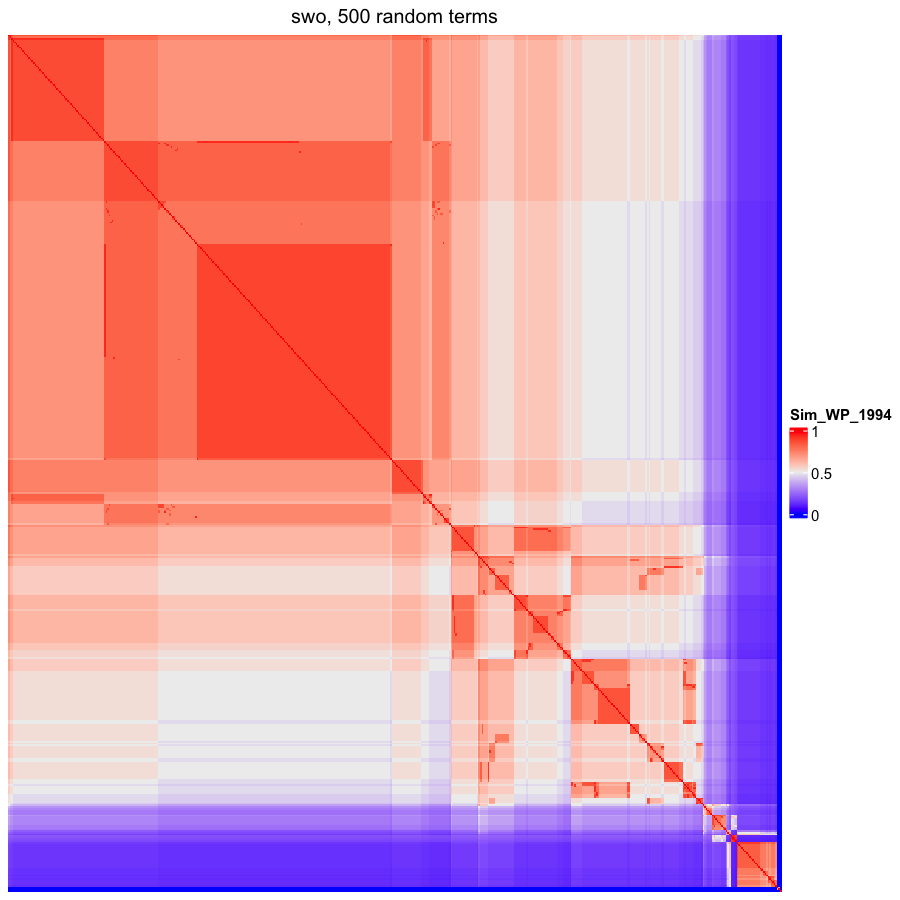

### OBOFoundry_tao.png

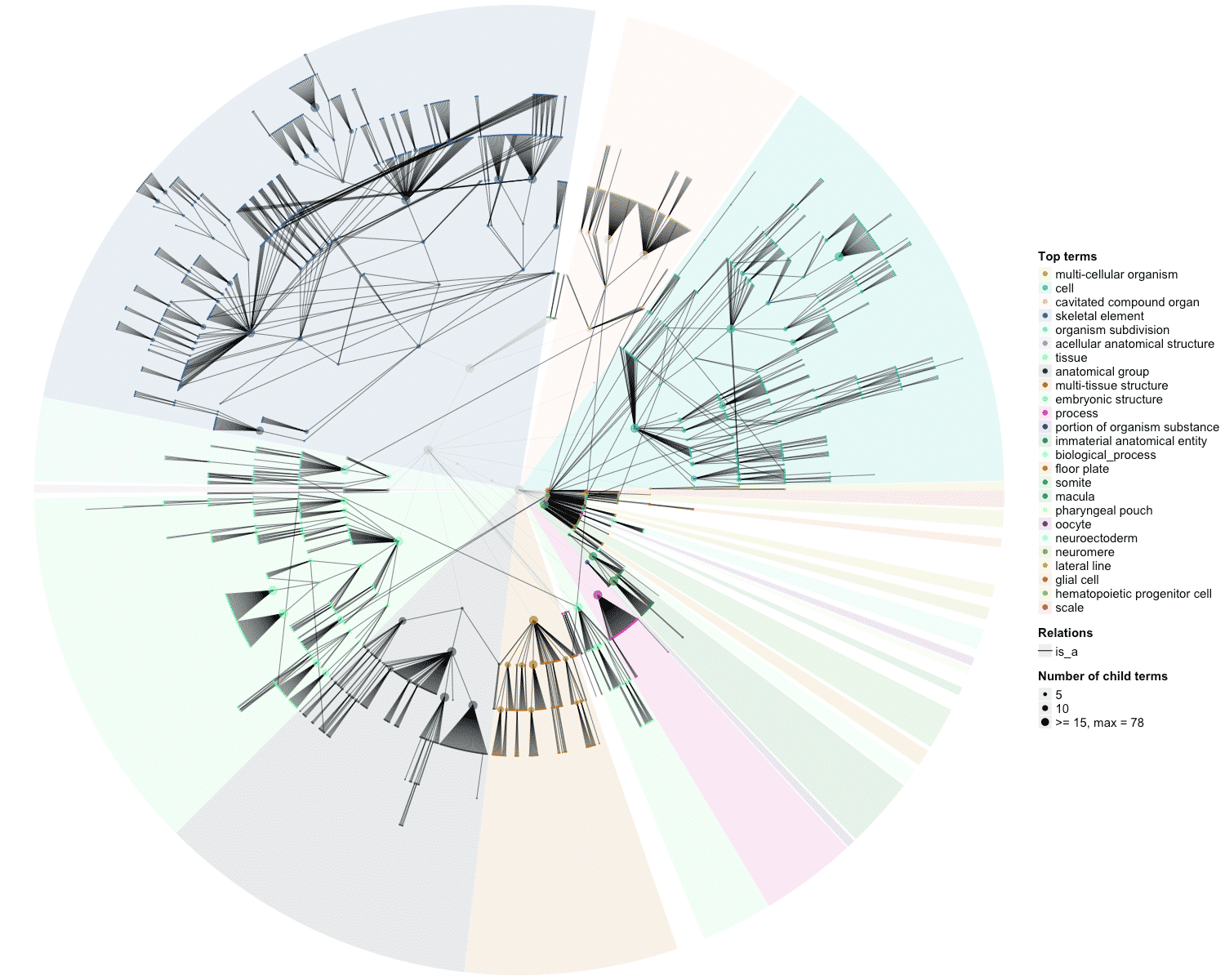

### OBOFoundry_tto.png

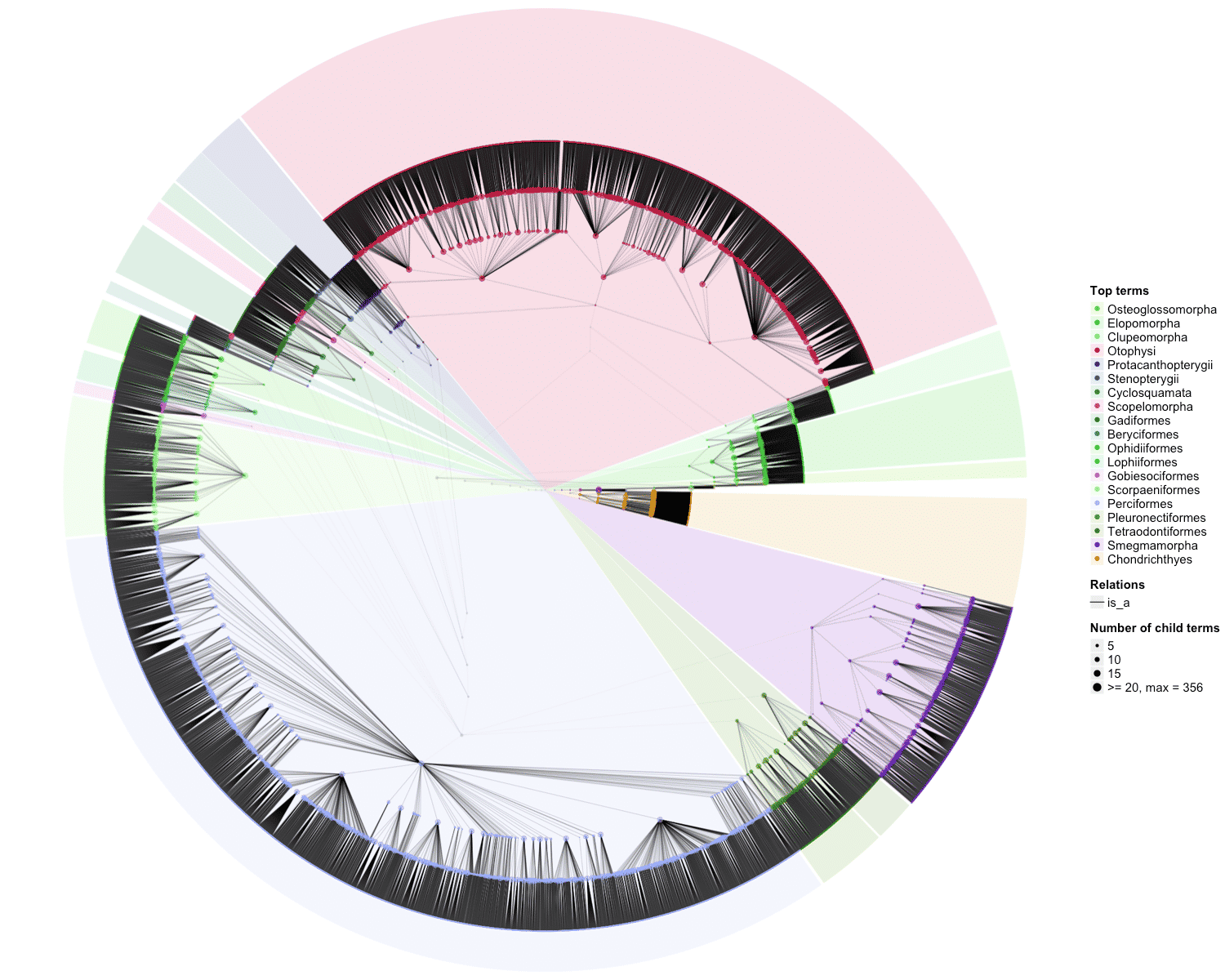

### OBOFoundry_uo_heatmap.png

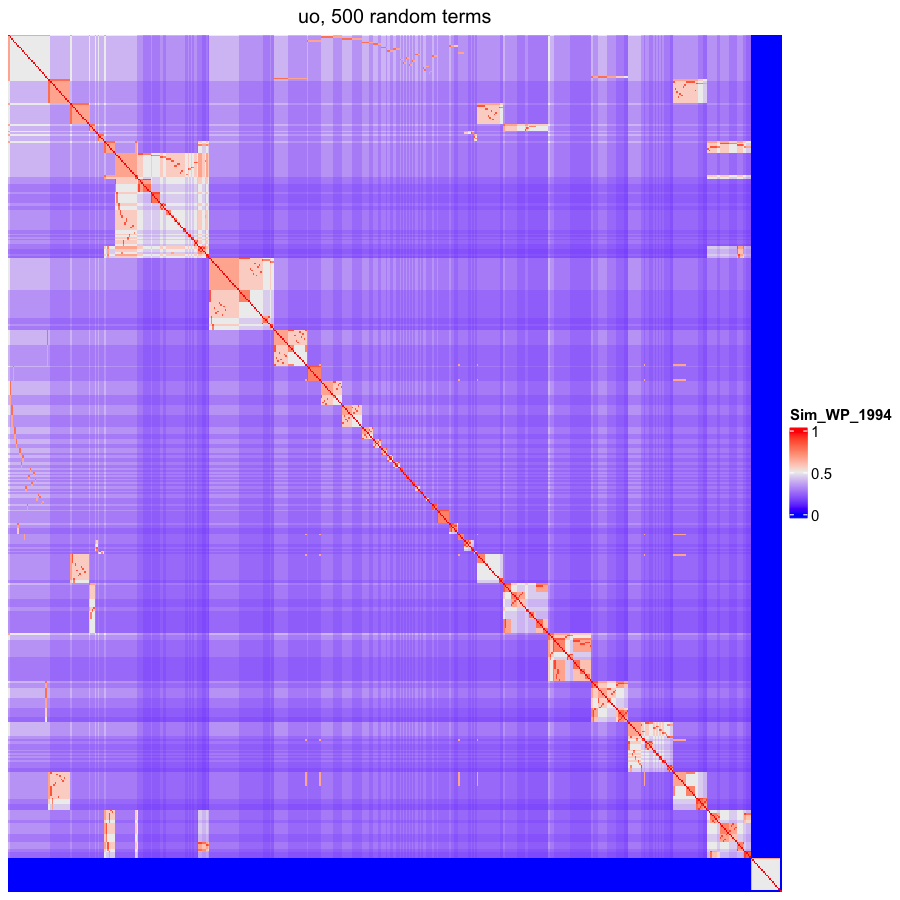

### OBOFoundry_wbphenotype.png

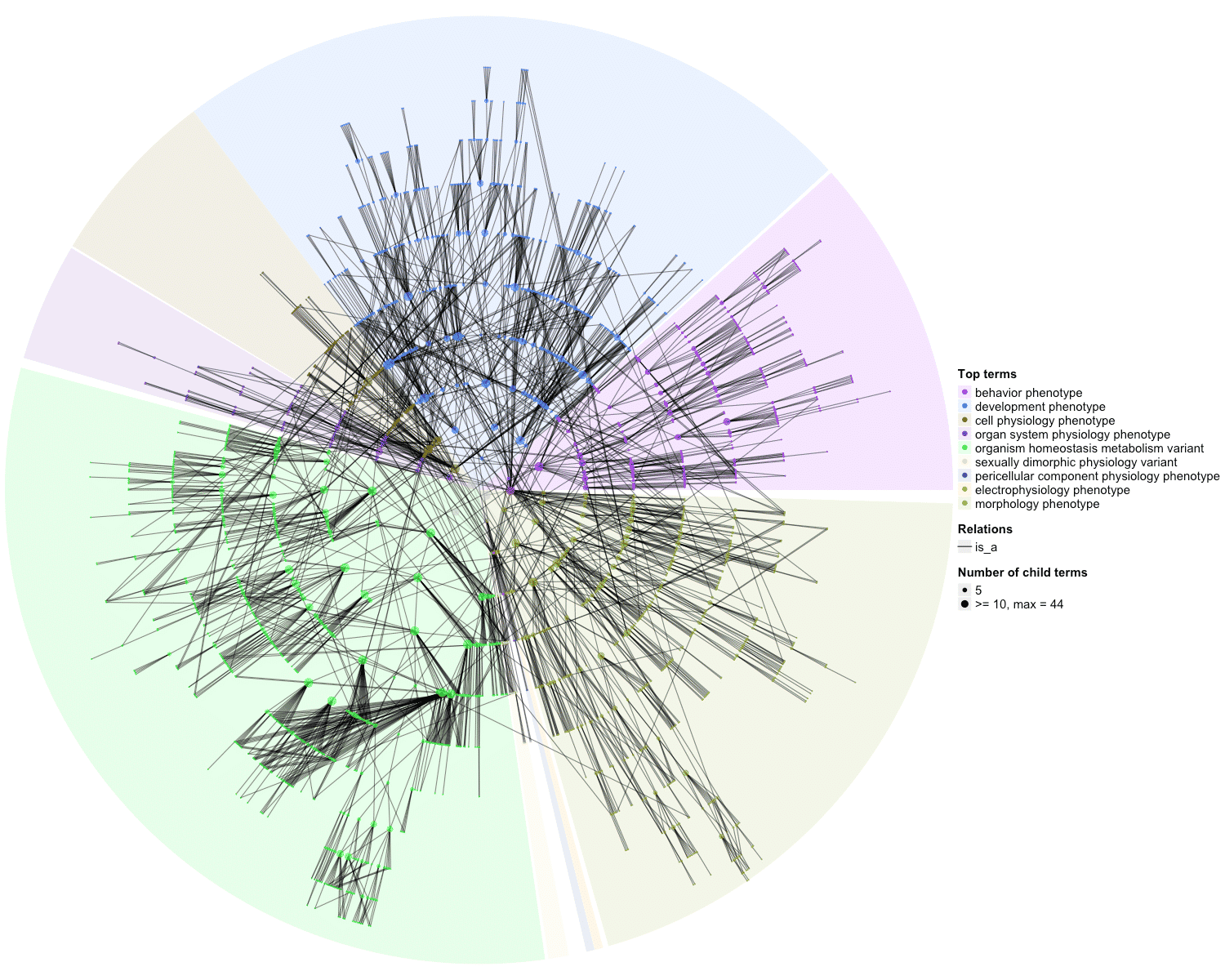

### OBOFoundry_zfa_heatmap.png

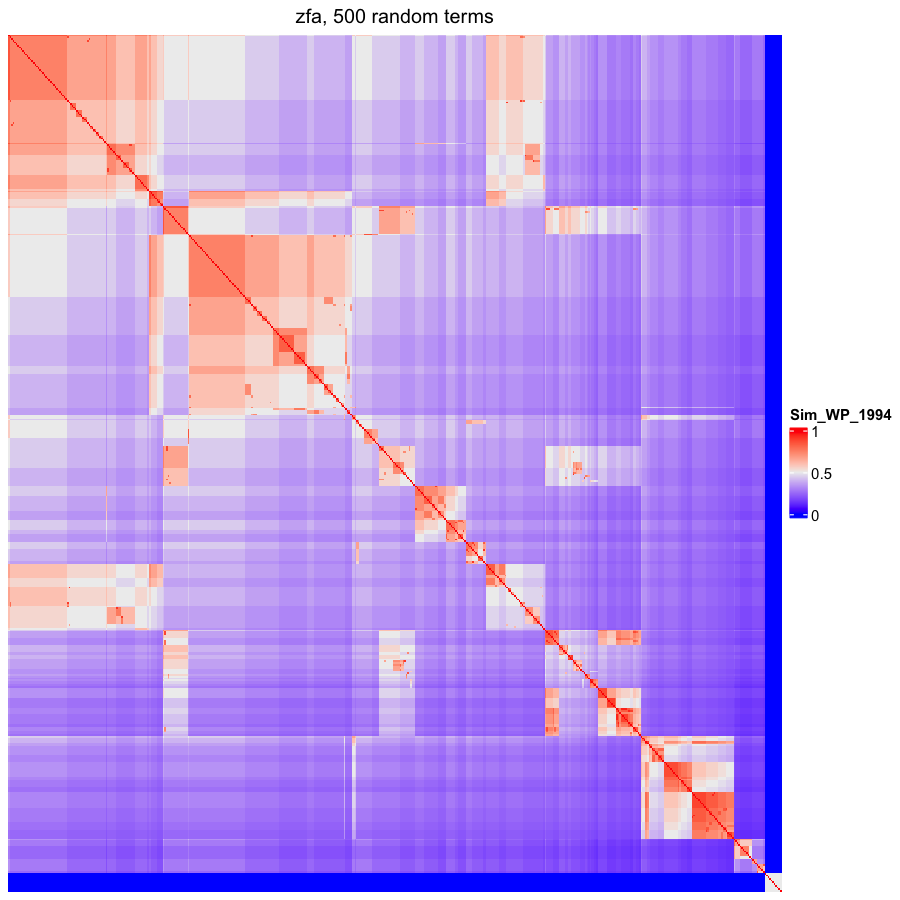

### scatterplot_group_1.png

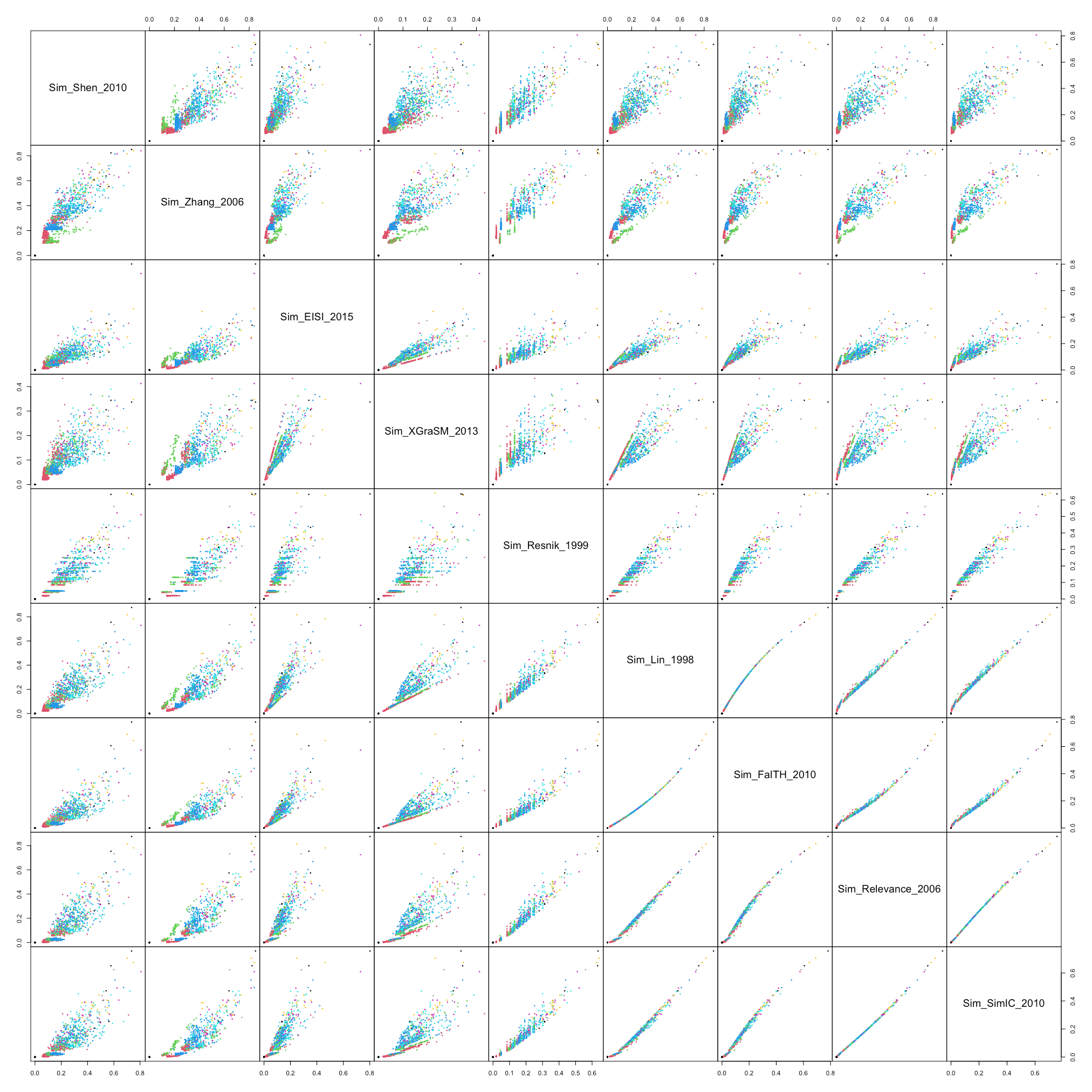

### scatterplot_group_2.png

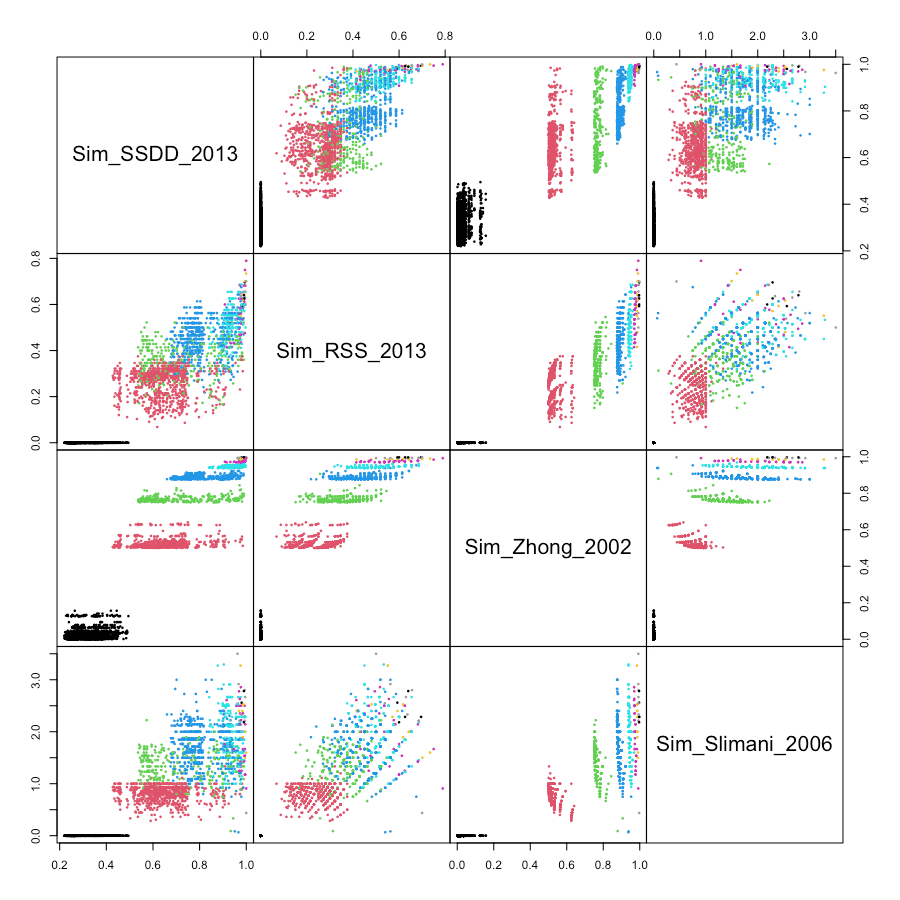

### scatterplot_group_3.png

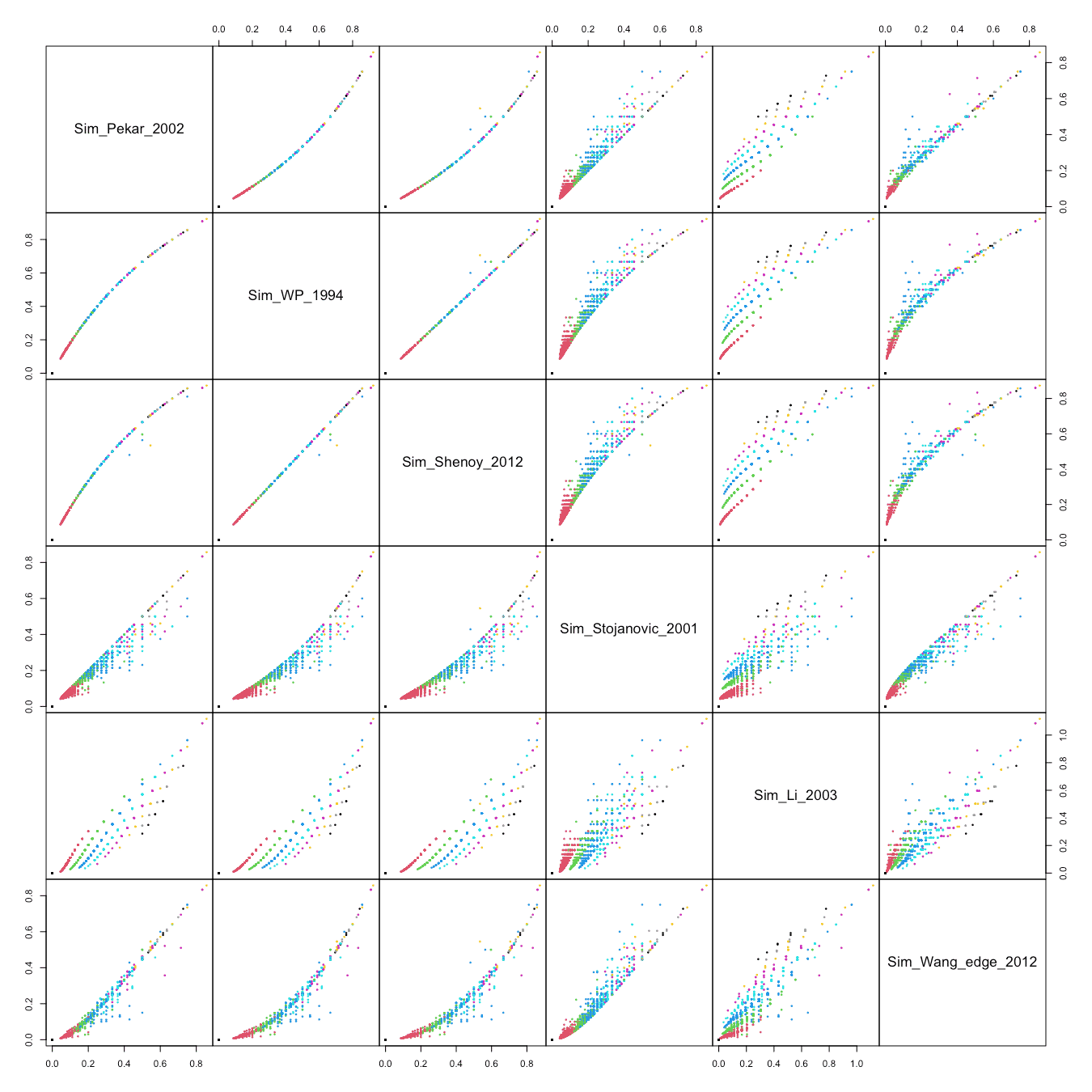

### scatterplot_group_4.png

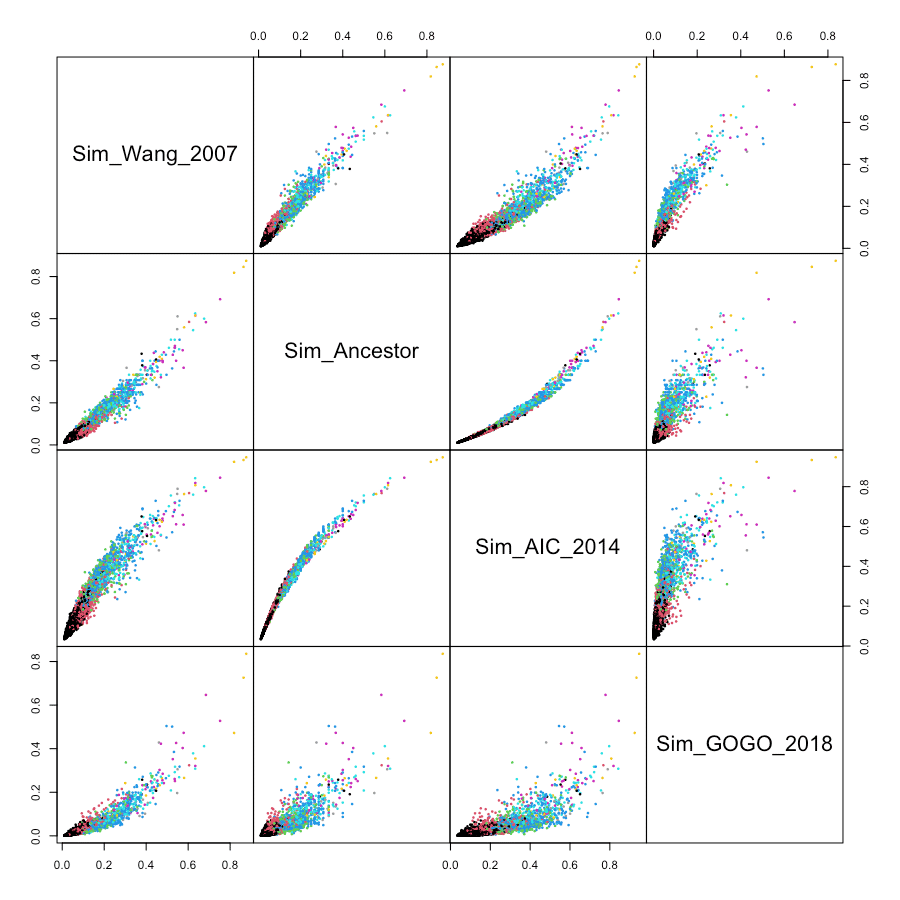

### scatterplot_group_5.png

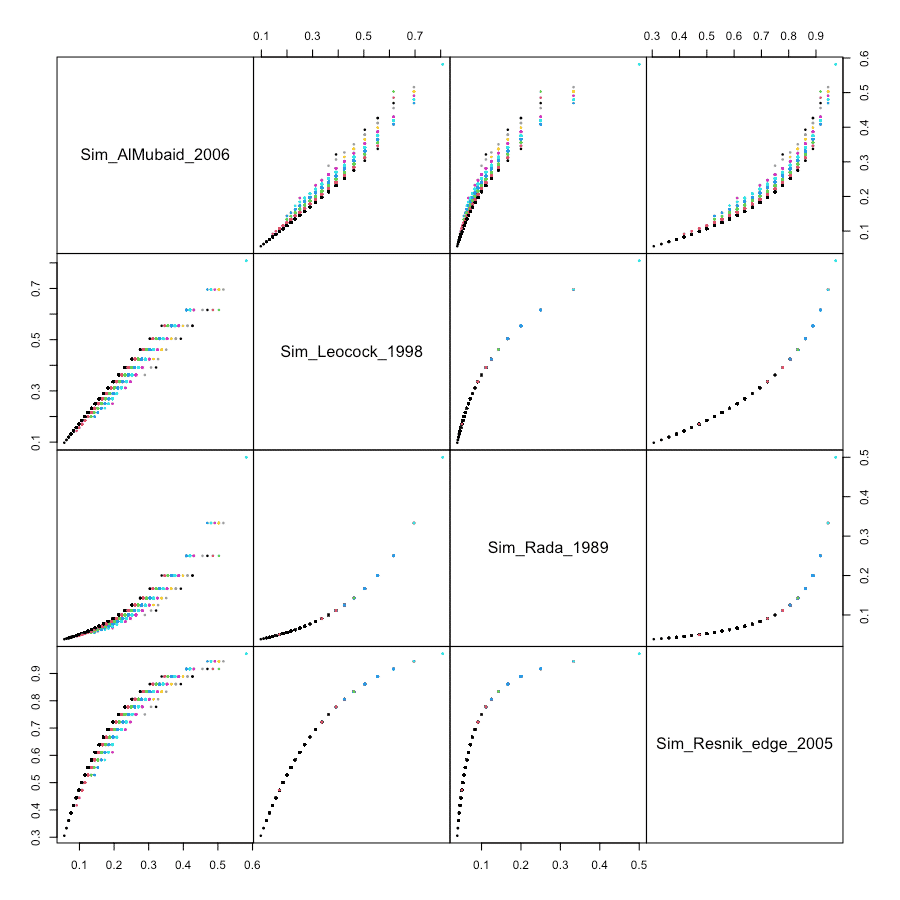
