## Supplementary File 3 for "*simona:* a comprehensive R package for semantic similarity analysis on bio-ontologies": compare_ic_methods.html


### Supplementary File 3. Compare IC methods

###### Zuguang Gu

#### 2024-08-27

**simona** supports many methods for calculating information contents (IC). In this document, we compare different IC methods using Gene Ontology as the test ontology.

By default, we use the Biological Process (BP) namespace in GO. We only take `"is_a"` and `"part_of"` as the relation types. The `org_db` argument is set to human for the `IC_annotation` method.

```
library(simona)
dag = create_ontology_DAG_from_GO_db(org_db = "org.Hs.eg.db")
dag
```

```
## An ontology_DAG object:
##   Source: GO BP / GO.db package 3.19.1 
##   27186 terms / 54178 relations
##   Root: GO:0008150 
##   Terms: GO:0000001, GO:0000002, GO:0000003, GO:0000011, ...
##   Max depth: 18 
##   Avg number of parents: 1.99
##   Avg number of children: 1.87
##   Aspect ratio: 356.46:1 (based on the longest distance from root)
##                 756.89:1 (based on the shortest distance from root)
##   Relations: is_a, part_of
##   Annotations: 18888 items
##                291, 1890, 4205, 4358, ...
## 
## With the following columns in the metadata data frame:
##   id, name, definition
```

All IC methods supported in **simona** are listed below. The full description of these IC methods can be found in the vignettes of **simona**.

```
all_term_IC_methods()
```

```
##  [1] "IC_offspring"     "IC_height"        "IC_annotation"    "IC_universal"    
##  [5] "IC_Zhang_2006"    "IC_Seco_2004"     "IC_Zhou_2008"     "IC_Seddiqui_2010"
##  [9] "IC_Sanchez_2011"  "IC_Meng_2012"     "IC_Wang_2007"
```

We calculate IC for all GO BP terms with different IC methods and save the results in a list.

```
lt = lapply(all_term_IC_methods(), function(method) {
    term_IC(dag, method)
})
names(lt) = all_term_IC_methods()
df = as.data.frame(lt)
```

We calculate the correlations between IC vectors from different methods and make the correlation heatmap. As 1 - correlation is also a valid dissimilarity measurement, we directly generate the hierarchical clustering based on `1 - cor`.

```
cor = cor(df, use = "pairwise.complete.obs")
hc = hclust(as.dist(1 - cor))
library(ComplexHeatmap)
Heatmap(cor, name = "correlation", cluster_rows = hc, cluster_columns = hc,
    row_dend_reorder = TRUE, column_dend_reorder = TRUE,
    column_title = "IC correlation, GO BP")
```

Figure S3.1. Correlation heatmap of ICs of GO BP terms by various IC methods.

The heatmap shows the IC methods can be put into at least two groups. The groups can be manually extracted by observing sub-trees from the dendrogram and the patterns on the heatmap.

We put `"IC_Wang_2007"` and `"IC_universal"` into a group called `"others"` because these two IC vectors show overall low similarities to other IC vectors.

```
group = c("IC_height"        = "1",
          "IC_Sanchez_2011"  = "1",
          "IC_Zhou_2008"     = "1",
          "IC_Meng_2012"     = "1",
          "IC_Seddiqui_2010" = "2",
          "IC_Seco_2004"     = "2",
          "IC_offspring"     = "2",
          "IC_Zhang_2006"    = "2",
          "IC_annotation"    = "2",
          "IC_Wang_2007"     = "others",
          "IC_universal"     = "others")
```

Next we perform MDS (multi-dimension scaling) analysis on the `1-cor` distance matrix and visualize the first two dimensions.

```
library(ggrepel)
library(ggplot2)
loc = cmdscale(as.dist(1 - cor))
loc = as.data.frame(loc)
colnames(loc) = c("x", "y")
loc$method = rownames(loc)

loc$group = group[rownames(loc)]

ggplot(loc, aes(x, y, label = method, col = factor(group))) + 
    geom_point() + 
    geom_text_repel(show.legend = FALSE, size = 3) +
    labs(x = "Dimension 1", y = "Dimension 2", col = "Group") +
    ggtitle("MDS based on the correlation between ICs from different IC methods")
```

Figure S3.2. MDS plot of similarities between various IC methods on GO BP terms.

Similar to the heatmap, the two methods of `"IC_Wang_2007"` and `"IC_universal"` are quite far from other points in the MDS plot.

We can directly compare different IC methods with pairwise scatterplots, colored by the depths of terms, which helps to see how term depths are weighted in different IC methods.

```
col = c("IC_height"        = 2,
        "IC_Sanchez_2011"  = 2,
        "IC_Zhou_2008"     = 2,
        "IC_Meng_2012"     = 2,
        "IC_Seddiqui_2010" = 3,
        "IC_Seco_2004"     = 3,
        "IC_offspring"     = 3,
        "IC_Zhang_2006"    = 3,
        "IC_annotation"    = 3,
        "IC_Wang_2007"     = 4,
        "IC_universal"     = 4)
pairs(df[, names(group)], pch = ".", col = dag_depth(dag), gap = 0, 
    text.panel=function(x, y, labels, cex, font, ...) {
        text(x, y, labels, col = col[labels])
    })
```

Figure 3.3. Pairwise comparison between various IC methods on GO BP terms.

```
sessionInfo()
```

```
## R version 4.4.1 (2024-06-14)
## Platform: x86_64-apple-darwin13.4.0
## Running under: macOS Big Sur ... 10.16
## 
## Matrix products: default
## BLAS/LAPACK: /Users/guz/opt/miniconda3/envs/R-4.4.1/lib/libopenblasp-r0.3.27.dylib;  LAPACK version 3.12.0
## 
## locale:
## [1] en_GB.UTF-8/en_GB.UTF-8/en_GB.UTF-8/C/en_GB.UTF-8/en_GB.UTF-8
## 
## time zone: Europe/Berlin
## tzcode source: system (macOS)
## 
## attached base packages:
## [1] grid      stats     graphics  grDevices utils     datasets  methods  
## [8] base     
## 
## other attached packages:
## [1] ggrepel_0.9.5         ggplot2_3.5.1         ComplexHeatmap_2.20.0
## [4] simona_1.3.12         knitr_1.48            rmarkdown_2.28       
## 
## loaded via a namespace (and not attached):
##  [1] tidyselect_1.2.1        farver_2.1.2            dplyr_1.1.4            
##  [4] blob_1.2.4              Biostrings_2.72.1       fastmap_1.2.0          
##  [7] promises_1.3.0          digest_0.6.37           mime_0.12              
## [10] lifecycle_1.0.4         cluster_2.1.6           KEGGREST_1.44.1        
## [13] RSQLite_2.3.7           magrittr_2.0.3          compiler_4.4.1         
## [16] rlang_1.1.4             sass_0.4.9              tools_4.4.1            
## [19] igraph_2.0.3            utf8_1.2.4              yaml_2.3.10            
## [22] labeling_0.4.3          bit_4.0.5               scatterplot3d_0.3-44   
## [25] xml2_1.3.6              RColorBrewer_1.1-3      withr_3.0.1            
## [28] BiocGenerics_0.50.0     stats4_4.4.1            fansi_1.0.6            
## [31] xtable_1.8-4            colorspace_2.1-1        GO.db_3.19.1           
## [34] scales_1.3.0            iterators_1.0.14        cli_3.6.3              
## [37] crayon_1.5.3            generics_0.1.3          httr_1.4.7             
## [40] rjson_0.2.22            DBI_1.2.3               cachem_1.1.0           
## [43] zlibbioc_1.50.0         parallel_4.4.1          AnnotationDbi_1.66.0   
## [46] XVector_0.44.0          matrixStats_1.3.0       vctrs_0.6.5            
## [49] jsonlite_1.8.8          IRanges_2.38.1          GetoptLong_1.0.5       
## [52] S4Vectors_0.42.1        bit64_4.0.5             clue_0.3-65            
## [55] foreach_1.5.2           jquerylib_0.1.4         glue_1.7.0             
## [58] codetools_0.2-20        Polychrome_1.5.1        shape_1.4.6.1          
## [61] gtable_0.3.5            later_1.3.2             GenomeInfoDb_1.40.1    
## [64] UCSC.utils_1.0.0        munsell_0.5.1           tibble_3.2.1           
## [67] pillar_1.9.0            htmltools_0.5.8.1       GenomeInfoDbData_1.2.12
## [70] circlize_0.4.16         R6_2.5.1                doParallel_1.0.17      
## [73] evaluate_0.24.0         shiny_1.9.1             Biobase_2.64.0         
## [76] highr_0.11              png_0.1-8               memoise_2.0.1          
## [79] httpuv_1.6.15           bslib_0.8.0             Rcpp_1.0.13            
## [82] org.Hs.eg.db_3.19.1     xfun_0.47               pkgconfig_2.0.3        
## [85] GlobalOptions_0.1.2
```
