## Supplementary File 4 for "*simona:* a comprehensive R package for semantic similarity analysis on bio-ontologies": compare_sim_methods.html

Supplementary File 4. Compare semantic similarity methods


### Supplementary File 4. Compare semantic similarity methods

###### Zuguang Gu

#### 2024-08-27

**simona** supports many methods for calculating semantic similarities. In this document, we compare different term similarity methods using Gene Ontology as the test ontology.

By default, we use the Biological Process (BP) namespace in GO. We only take `"is_a"` and `"part_of"` as the relation types. The `org_db` argument is also set to human for the annotation-based method.

```
library(simona)
dag = create_ontology_DAG_from_GO_db(org_db = "org.Hs.eg.db")
dag
```

```
## An ontology_DAG object:
##   Source: GO BP / GO.db package 3.19.1 
##   27186 terms / 54178 relations
##   Root: GO:0008150 
##   Terms: GO:0000001, GO:0000002, GO:0000003, GO:0000011, ...
##   Max depth: 18 
##   Avg number of parents: 1.99
##   Avg number of children: 1.87
##   Aspect ratio: 356.46:1 (based on the longest distance from root)
##                 756.89:1 (based on the shortest distance from root)
##   Relations: is_a, part_of
##   Annotations: 18888 items
##                291, 1890, 4205, 4358, ...
## 
## With the following columns in the metadata data frame:
##   id, name, definition
```

All term similarity methods supported in **simona** are listed as follows. The full description of these methods can be found in the vignettes of **simona**.

```
all_term_sim_methods()
```

```
##  [1] "Sim_Lin_1998"         "Sim_Resnik_1999"      "Sim_FaITH_2010"      
##  [4] "Sim_Relevance_2006"   "Sim_SimIC_2010"       "Sim_XGraSM_2013"     
##  [7] "Sim_EISI_2015"        "Sim_AIC_2014"         "Sim_Zhang_2006"      
## [10] "Sim_universal"        "Sim_Wang_2007"        "Sim_GOGO_2018"       
## [13] "Sim_Rada_1989"        "Sim_Resnik_edge_2005" "Sim_Leocock_1998"    
## [16] "Sim_WP_1994"          "Sim_Slimani_2006"     "Sim_Shenoy_2012"     
## [19] "Sim_Pekar_2002"       "Sim_Stojanovic_2001"  "Sim_Wang_edge_2012"  
## [22] "Sim_Zhong_2002"       "Sim_AlMubaid_2006"    "Sim_Li_2003"         
## [25] "Sim_RSS_2013"         "Sim_HRSS_2013"        "Sim_Shen_2010"       
## [28] "Sim_SSDD_2013"        "Sim_Jiang_1997"       "Sim_Kappa"           
## [31] "Sim_Jaccard"          "Sim_Dice"             "Sim_Overlap"         
## [34] "Sim_Ancestor"
```

We compare all supported term similarity methods, using 500 random GO terms. Since we also compare annotation-based methods, we randomly sample 500 GO terms from those having gene annotations.

```
set.seed(123)
ic = term_IC(dag, method = "IC_annotation")
ic = ic[!is.na(ic)]
go_id = sample(names(ic), 500)
```

We calculate similarities with different term similarity methods and save the results in a list.

```
lt = lapply(all_term_sim_methods(), function(method) {
    term_sim(dag, go_id, method)
})
names(lt) = all_term_sim_methods()
```

For comparison, we take the lower triangle matrix and merge all similarities values from different methods into a data frame.

```
df = as.data.frame(lapply(lt, function(x) x[lower.tri(x)]))
```

We calculate the correlations between similarity vectors from different methods and make the correlation heatmap. As 1 - correlation is also a valid dissimilarity measurement, we directly generate the hierarchical clustering based on `1 - cor`.

```
cor = cor(df, use = "pairwise.complete.obs")
hc = hclust(as.dist(1 - cor))
library(ComplexHeatmap)
Heatmap(cor, name = "correlation", cluster_rows = hc, cluster_columns = hc,
    row_dend_reorder = TRUE, column_dend_reorder = TRUE,
    column_title = "Semantic similarity correlation, GO BP")
```

Figure S4.1. Correlation heatmap of semantic similarities of 500 random GO BP terms by various similarity methods.

We next remove the following similarity methods which show very different patterns to other methods, and remake the heatmap.

```
ind = which(colnames(df) %in% c("Sim_Jiang_1997", "Sim_HRSS_2013", "Sim_universal",
    "Sim_Dice", "Sim_Kappa", "Sim_Jaccard", "Sim_Overlap"))
cor2 = cor[-ind, -ind]
df2 = df[, -ind]

hc = hclust(as.dist(1 - cor2))
Heatmap(cor2, name = "correlation", cluster_rows = hc, cluster_columns = hc,
    row_dend_reorder = TRUE, column_dend_reorder = TRUE,
    column_title = "Semantic similarity correlation, GO BP")
```

Figure S4.2. Correlation heatmap of semantic similarities after removing seven methods that generate very different patterns from other methods. Please note the color scale is differennt from Figure S4.1.

We can observe that all the similarity methods can be put into five groups. We manually add group labels to these methods:

```
group = c("Sim_Shen_2010"        = 1,
          "Sim_Zhang_2006"       = 1,
          "Sim_EISI_2015"        = 1,
          "Sim_XGraSM_2013"      = 1,
          "Sim_Resnik_1999"      = 1,
          "Sim_Lin_1998"         = 1,
          "Sim_FaITH_2010"       = 1,
          "Sim_Relevance_2006"   = 1,
          "Sim_SimIC_2010"       = 1,
          "Sim_SSDD_2013"        = 2,
          "Sim_RSS_2013"         = 2,
          "Sim_Zhong_2002"       = 2,
          "Sim_Slimani_2006"     = 2,
          "Sim_Pekar_2002"       = 3,
          "Sim_WP_1994"          = 3,
          "Sim_Shenoy_2012"      = 3,
          "Sim_Stojanovic_2001"  = 3,
          "Sim_Li_2003"          = 3,
          "Sim_Wang_edge_2012"   = 3,
          "Sim_Wang_2007"        = 4,
          "Sim_Ancestor"         = 4,
          "Sim_AIC_2014"         = 4,
          "Sim_GOGO_2018"        = 4,
          "Sim_AlMubaid_2006"    = 5,
          "Sim_Leocock_1998"     = 5,
          "Sim_Rada_1989"        = 5,
          "Sim_Resnik_edge_2005" = 5)
```

Next we perform MDS (multidimension scaling) analysis on the `1-cor` distance matrix and visualize the first two dimensions.

```
library(ggrepel)
library(ggplot2)
loc = cmdscale(as.dist(1-cor2))
loc = as.data.frame(loc)
colnames(loc) = c("x", "y")
loc$method = rownames(loc)

loc$group = group[rownames(loc)]

ggplot(loc, aes(x, y, label = method, col = factor(group))) + 
    geom_point() + 
    geom_text_repel(show.legend = FALSE, size = 3) +
    labs(x = "Dimension 1", y = "Dimension 2", col = "Group") +
    ggtitle("MDS on similarity methods on random 500 GO terms")
```

Figure S4.3. MDS plot of various similarity methods.

Following are the pairwise scatterplots of semantic similarities from two term similarity methods. The five groups are from the heatmap in Figure S4.2. Since each point correspond to a term-pair, the color of the point corresponds to the depth of their LCA term. Note that for those methods based on MICA terms, as shown in the document “Compare topology and annotation-based semantic similarity methods”, majority of LCA terms and MICA terms are the same. So in this comparison, we can take \(\mathrm{LCA} \approx \mathrm{MICA}\).

The methods in the five groups are summarized as follows:

- Group 1: methods that integrate information content. In the comparison we did here, it is IC\_annotation.
- Group 2: so-called “hybrid” methods.
- Group 3: methods that based on the depth of terms (e.g. LCA terms).
- Group 4: methods that aggregate from all common ancestor terms.
- Group 5: methods that based on the distance of two terms in the DAG.

- Group 1
- Group 2
- Group 3
- Group 4
- Group 5

Figure S4.4. Pairwise comparison between various similarity methods on 500 random GO BP terms. The five groups are defined in Figure S4.2.

---

Finally, you can select an individual heatmap of similarities calculated by:  Sim\_Lin\_1998 Sim\_Resnik\_1999 Sim\_FaITH\_2010 Sim\_Relevance\_2006 Sim\_SimIC\_2010 Sim\_XGraSM\_2013 Sim\_EISI\_2015 Sim\_AIC\_2014 Sim\_Zhang\_2006 Sim\_universal Sim\_Wang\_2007 Sim\_GOGO\_2018 Sim\_Rada\_1989 Sim\_Resnik\_edge\_2005 Sim\_Leocock\_1998 Sim\_WP\_1994 Sim\_Slimani\_2006 Sim\_Shenoy\_2012 Sim\_Pekar\_2002 Sim\_Stojanovic\_2001 Sim\_Wang\_edge\_2012 Sim\_Zhong\_2002 Sim\_AlMubaid\_2006 Sim\_Li\_2003 Sim\_RSS\_2013 Sim\_HRSS\_2013 Sim\_Shen\_2010 Sim\_SSDD\_2013 Sim\_Jiang\_1997 Sim\_Kappa Sim\_Jaccard Sim\_Dice Sim\_Overlap Sim\_Ancestor   Use order from Sim\_Lin\_1998

Prev method: Sim\_Ancestor Curr method: Sim\_Lin\_1998 Next method: Sim\_Resnik\_1999

Figure S4.5. Similarity heatmap of the 500 random GO BP terms under different similarity methods.

```
sessionInfo()
```

```
## R version 4.4.1 (2024-06-14)
## Platform: x86_64-apple-darwin13.4.0
## Running under: macOS Big Sur ... 10.16
## 
## Matrix products: default
## BLAS/LAPACK: /Users/guz/opt/miniconda3/envs/R-4.4.1/lib/libopenblasp-r0.3.27.dylib;  LAPACK version 3.12.0
## 
## locale:
## [1] en_GB.UTF-8/en_GB.UTF-8/en_GB.UTF-8/C/en_GB.UTF-8/en_GB.UTF-8
## 
## time zone: Europe/Berlin
## tzcode source: system (macOS)
## 
## attached base packages:
## [1] grid      stats     graphics  grDevices utils     datasets  methods  
## [8] base     
## 
## other attached packages:
## [1] GetoptLong_1.0.5      ggrepel_0.9.5         ggplot2_3.5.1        
## [4] ComplexHeatmap_2.20.0 simona_1.3.12         knitr_1.48           
## [7] rmarkdown_2.28       
## 
## loaded via a namespace (and not attached):
##  [1] tidyselect_1.2.1        farver_2.1.2            dplyr_1.1.4            
##  [4] blob_1.2.4              Biostrings_2.72.1       fastmap_1.2.0          
##  [7] promises_1.3.0          digest_0.6.37           mime_0.12              
## [10] lifecycle_1.0.4         cluster_2.1.6           KEGGREST_1.44.1        
## [13] RSQLite_2.3.7           magrittr_2.0.3          compiler_4.4.1         
## [16] rlang_1.1.4             sass_0.4.9              tools_4.4.1            
## [19] utf8_1.2.4              igraph_2.0.3            yaml_2.3.10            
## [22] labeling_0.4.3          bit_4.0.5               scatterplot3d_0.3-44   
## [25] xml2_1.3.6              RColorBrewer_1.1-3      withr_3.0.1            
## [28] BiocGenerics_0.50.0     stats4_4.4.1            fansi_1.0.6            
## [31] xtable_1.8-4            colorspace_2.1-1        GO.db_3.19.1           
## [34] scales_1.3.0            iterators_1.0.14        cli_3.6.3              
## [37] crayon_1.5.3            generics_0.1.3          ragg_1.3.2             
## [40] httr_1.4.7              rjson_0.2.22            DBI_1.2.3              
## [43] cachem_1.1.0            zlibbioc_1.50.0         parallel_4.4.1         
## [46] AnnotationDbi_1.66.0    XVector_0.44.0          proxyC_0.4.1           
## [49] matrixStats_1.3.0       vctrs_0.6.5             Matrix_1.7-0           
## [52] jsonlite_1.8.8          IRanges_2.38.1          S4Vectors_0.42.1       
## [55] bit64_4.0.5             clue_0.3-65             systemfonts_1.1.0      
## [58] foreach_1.5.2           jquerylib_0.1.4         glue_1.7.0             
## [61] codetools_0.2-20        Polychrome_1.5.1        shape_1.4.6.1          
## [64] gtable_0.3.5            later_1.3.2             GenomeInfoDb_1.40.1    
## [67] UCSC.utils_1.0.0        munsell_0.5.1           tibble_3.2.1           
## [70] pillar_1.9.0            htmltools_0.5.8.1       GenomeInfoDbData_1.2.12
## [73] circlize_0.4.16         R6_2.5.1                textshaping_0.4.0      
## [76] doParallel_1.0.17       evaluate_0.24.0         shiny_1.9.1            
## [79] Biobase_2.64.0          lattice_0.22-6          highr_0.11             
## [82] png_0.1-8               memoise_2.0.1           httpuv_1.6.15          
## [85] bslib_0.8.0             Rcpp_1.0.13             org.Hs.eg.db_3.19.1    
## [88] xfun_0.47               pkgconfig_2.0.3         GlobalOptions_0.1.2
```
