## Supplementary File 4 for "*simona:* a comprehensive R package for semantic similarity analysis on bio-ontologies": compare_sim_methods.Rmd

````
---
title: "Supplementary File 4. Compare semantic similarity methods"
author: "Zuguang Gu"
date: '`r Sys.Date()`'
output: 
  html_document:
    css: main.css
    toc: true
    fig_caption: true
---

```{r, echo = FALSE, message = FALSE}
library(knitr)
knitr::opts_chunk$set(
    error = FALSE,
    tidy  = FALSE,
    message = FALSE,
    warning = FALSE,
    fig.width = 6, fig.height = 6,
    fig.align = "center")

knitr::knit_hooks$set(pngquant = knitr::hook_pngquant)

knitr::opts_chunk$set(
  message = FALSE,
  dev = "ragg_png",
  fig.align = "center",
  pngquant = "--speed=10 --quality=30"
)
```

**simona** supports many methods for calculating semantic similarities. In
this document, we compare different term similarity methods using Gene Ontology as the test
ontology.

By default, we use the Biological Process (BP) namespace in GO. We only take
`"is_a"` and `"part_of"` as the relation types. The `org_db` argument is also
set to human for the annotation-based method.


```{r}
library(simona)
dag = create_ontology_DAG_from_GO_db(org_db = "org.Hs.eg.db")
dag
```


All term similarity methods supported in **simona** are listed as follows. The full
description of these methods can be found in the vignettes of **simona**.

```{r}
all_term_sim_methods()
```

We compare all supported term similarity methods, using 500 random GO terms.
Since we also compare annotation-based methods, we randomly sample 500 GO terms
from those having gene annotations.

```{r}
set.seed(123)
ic = term_IC(dag, method = "IC_annotation")
ic = ic[!is.na(ic)]
go_id = sample(names(ic), 500)
```

We calculate similarities with different term similarity methods and save the results in a list.

```{r}
lt = lapply(all_term_sim_methods(), function(method) {
    term_sim(dag, go_id, method)
})
names(lt) = all_term_sim_methods()
```

For comparison, we take the lower triangle matrix and merge all similarities values from
different methods into a data frame.

```{r}
df = as.data.frame(lapply(lt, function(x) x[lower.tri(x)]))
```

We calculate the correlations between similarity vectors from different methods and
make the correlation heatmap. As 1 - correlation is also a valid dissimilarity
measurement, we directly generate the hierarchical clustering based on `1 -
cor`.


```{r, fig.width = 9, fig.height = 8, fig.cap = "Figure S4.1. Correlation heatmap of semantic similarities of 500 random GO BP terms by various similarity methods."}
cor = cor(df, use = "pairwise.complete.obs")
hc = hclust(as.dist(1 - cor))
library(ComplexHeatmap)
Heatmap(cor, name = "correlation", cluster_rows = hc, cluster_columns = hc,
    row_dend_reorder = TRUE, column_dend_reorder = TRUE,
    column_title = "Semantic similarity correlation, GO BP")
```


We next remove the following similarity methods which show very different patterns
to other methods, and remake the heatmap.

```{r, fig.width = 8, fig.height = 8, fig.cap = "Figure S4.2. Correlation heatmap of semantic similarities after removing seven methods that generate very different patterns from other methods. Please note the color scale is differennt from Figure S4.1."}
ind = which(colnames(df) %in% c("Sim_Jiang_1997", "Sim_HRSS_2013", "Sim_universal",
    "Sim_Dice", "Sim_Kappa", "Sim_Jaccard", "Sim_Overlap"))
cor2 = cor[-ind, -ind]
df2 = df[, -ind]

hc = hclust(as.dist(1 - cor2))
Heatmap(cor2, name = "correlation", cluster_rows = hc, cluster_columns = hc,
    row_dend_reorder = TRUE, column_dend_reorder = TRUE,
    column_title = "Semantic similarity correlation, GO BP")
```

We can observe that all the similarity methods can be put into five groups. We manually add
group labels to these methods:

```{r}
group = c("Sim_Shen_2010"        = 1,
          "Sim_Zhang_2006"       = 1,
          "Sim_EISI_2015"        = 1,
          "Sim_XGraSM_2013"      = 1,
          "Sim_Resnik_1999"      = 1,
          "Sim_Lin_1998"         = 1,
          "Sim_FaITH_2010"       = 1,
          "Sim_Relevance_2006"   = 1,
          "Sim_SimIC_2010"       = 1,
          "Sim_SSDD_2013"        = 2,
          "Sim_RSS_2013"         = 2,
          "Sim_Zhong_2002"       = 2,
          "Sim_Slimani_2006"     = 2,
          "Sim_Pekar_2002"       = 3,
          "Sim_WP_1994"          = 3,
          "Sim_Shenoy_2012"      = 3,
          "Sim_Stojanovic_2001"  = 3,
          "Sim_Li_2003"          = 3,
          "Sim_Wang_edge_2012"   = 3,
          "Sim_Wang_2007"        = 4,
          "Sim_Ancestor"         = 4,
          "Sim_AIC_2014"         = 4,
          "Sim_GOGO_2018"        = 4,
          "Sim_AlMubaid_2006"    = 5,
          "Sim_Leocock_1998"     = 5,
          "Sim_Rada_1989"        = 5,
          "Sim_Resnik_edge_2005" = 5)
```

Next we perform MDS (multidimension scaling) analysis on the `1-cor` distance matrix and
visualize the first two dimensions. 


```{r, fig.width = 7, fig.height = 6, fig.cap = "Figure S4.3. MDS plot of various similarity methods."}
library(ggrepel)
library(ggplot2)
loc = cmdscale(as.dist(1-cor2))
loc = as.data.frame(loc)
colnames(loc) = c("x", "y")
loc$method = rownames(loc)

loc$group = group[rownames(loc)]

ggplot(loc, aes(x, y, label = method, col = factor(group))) + 
    geom_point() + 
    geom_text_repel(show.legend = FALSE, size = 3) +
    labs(x = "Dimension 1", y = "Dimension 2", col = "Group") +
    ggtitle("MDS on similarity methods on random 500 GO terms")
```


```{r, echo = FALSE, eval = !file.exists("image")}
dir.create("image", showWarnings = FALSE)
od = hclust(dist(lt[["Sim_Lin_1998"]]))$order
for(nm in names(lt)) {
    png(paste0("image/go_bp_random_500_sim_", nm, ".png"), width = 700*1.5, height = 700*1.5, res = 72*1.5)
    draw(Heatmap(lt[[nm]], name = "Similarity", column_title = nm,
        show_row_names = FALSE, show_column_names = FALSE,
        show_row_dend = FALSE, show_column_dend = FALSE))
    dev.off()

    png(paste0("image/go_bp_random_500_sim_", nm, "_Lin_order.png"), width = 700*1.5, height = 700*1.5, res = 72*1.5)
    draw(Heatmap(lt[[nm]], name = "Similarity", column_title = nm,
        row_order = od, column_order = od,
        show_row_names = FALSE, show_column_names = FALSE))
    dev.off()
}
```


Following are the pairwise scatterplots of semantic similarities from two term similarity methods. The five groups
are from the heatmap in Figure S4.2. Since each point correspond to a term-pair, the color of the point
corresponds to the depth of their LCA term. Note that for those methods based on MICA terms, as shown in the document ["Compare topology and annotation-based semantic similarity methods"](../compare_topo_anno/compare_topo_anno.html), majority of LCA terms and MICA terms are the same. So in this comparison, we can take $\mathrm{LCA} \approx \mathrm{MICA}$.

The methods in the five groups are summarized as follows:

- Group 1: methods that integrate information content. In the comparison we did here, it is IC_annotation.
- Group 2: so-called "hybrid" methods.
- Group 3: methods that based on the depth of terms (e.g. LCA terms).
- Group 4: methods that aggregate from all common ancestor terms.
- Group 5: methods that based on the distance of two terms in the DAG.

```{r, eval = !file.exists("image/scatterplot_group_1_foo.png"), echo = FALSE}
LCA_depth = LCA_depth(dag, go_id)
LCA_depth = LCA_depth[lower.tri(LCA_depth)]

library(GetoptLong)
for(i in 1:5) {
    l = group == i
    g = names(group)[l]
    ind = sample(nrow(df2), 5000)

    png(qq("image/scatterplot_group_@{i}_foo.png"), 
        width = 150*sum(l)*1.5, height = 150*sum(l)*1.5, res = 72*1.5)
    pairs(df2[ind, g], pch = 16, cex = 0.5, gap = 0, col = LCA_depth[ind]+1)
    dev.off()

    system(qq("pngquant --speed=10 --quality=60 image/scatterplot_group_@{i}_foo.png -o image/scatterplot_group_@{i}.png --force"))
    file.remove(qq("image/scatterplot_group_@{i}_foo.png"))

}
```


<style>
#scatterplot ul li {
    padding-right:20px;
    cursor: pointer;
    display: inline-block;
    color: #337ab7;
}
#scatterplot ul {
    padding-inline-start: 0px;
    margin-top: 20px;
}
</style>

<div id="scatterplot">
<ul>
<li><a id="scatterplot_group1" style="color:#337ab7" onclick="switch_scatterplot(1);">Group 1</a></li>
<li><a id="scatterplot_group2" style="color:grey" onclick="switch_scatterplot(2);">Group 2</a></li>
<li><a id="scatterplot_group3" style="color:grey" onclick="switch_scatterplot(3);">Group 3</a></li>
<li><a id="scatterplot_group4" style="color:grey" onclick="switch_scatterplot(4);">Group 4</a></li>
<li><a id="scatterplot_group5" style="color:grey" onclick="switch_scatterplot(5);">Group 5</a></li>
</ul>
<div class="figure" style="text-align:center">
<img id="scatterplot_image" src="image/scatterplot_group_1.png" />
<p class="caption">Figure S4.4. Pairwise comparison between various similarity methods on 500 random GO BP terms. The five groups are defined in Figure S4.2.</p>
</div>
</div>

<script>
scatterplot_size = [9, 4, 6, 4, 4];
$("#scatterplot_image").width(scatterplot_size[0]*150);
function switch_scatterplot(i) {
    $("#scatterplot_image").attr("src", "image/scatterplot_group_" + i + ".png");
    $("#scatterplot_image").width(scatterplot_size[i-1]*150);
    $("#scatterplot_group1").css("color", "grey");
    $("#scatterplot_group2").css("color", "grey");
    $("#scatterplot_group3").css("color", "grey");
    $("#scatterplot_group4").css("color", "grey");
    $("#scatterplot_group5").css("color", "grey");
    $("#scatterplot_group" + i).css("color", "#337ab7");
}
</script>

<hr>

<style>
#prev, #next {
    text-decoration: underline;
    color: #337ab7;
}
#prev:hover, #next:hover {
    cursor: pointer;
}
#prev, #curr {
    margin-right: 50px;
}
#image p {
    text-align: center;
}
</style>

Finally, you can select an individual heatmap of similarities calculated by: <select id="select">
  <option value='0'>Sim_Lin_1998</option>
  <option value='1'>Sim_Resnik_1999</option>
  <option value='2'>Sim_FaITH_2010</option>
  <option value='3'>Sim_Relevance_2006</option>
  <option value='4'>Sim_SimIC_2010</option>
  <option value='5'>Sim_XGraSM_2013</option>
  <option value='6'>Sim_EISI_2015</option>
  <option value='7'>Sim_AIC_2014</option>
  <option value='8'>Sim_Zhang_2006</option>
  <option value='9'>Sim_universal</option>
  <option value='10'>Sim_Wang_2007</option>
  <option value='11'>Sim_GOGO_2018</option>
  <option value='12'>Sim_Rada_1989</option>
  <option value='13'>Sim_Resnik_edge_2005</option>
  <option value='14'>Sim_Leocock_1998</option>
  <option value='15'>Sim_WP_1994</option>
  <option value='16'>Sim_Slimani_2006</option>
  <option value='17'>Sim_Shenoy_2012</option>
  <option value='18'>Sim_Pekar_2002</option>
  <option value='19'>Sim_Stojanovic_2001</option>
  <option value='20'>Sim_Wang_edge_2012</option>
  <option value='21'>Sim_Zhong_2002</option>
  <option value='22'>Sim_AlMubaid_2006</option>
  <option value='23'>Sim_Li_2003</option>
  <option value='24'>Sim_RSS_2013</option>
  <option value='25'>Sim_HRSS_2013</option>
  <option value='26'>Sim_Shen_2010</option>
  <option value='27'>Sim_SSDD_2013</option>
  <option value='28'>Sim_Jiang_1997</option>
  <option value='29'>Sim_Kappa</option>
  <option value='30'>Sim_Jaccard</option>
  <option value='31'>Sim_Dice</option>
  <option value='32'>Sim_Overlap</option>
  <option value='33'>Sim_Ancestor</option>
</select> <input id="use_lin_order" type="checkbox" checked /> Use order from Sim_Lin_1998

<span id="prev" value="33">Prev method: Sim_Ancestor</span> <span id="curr">Curr method: Sim_Lin_1998</span> <span id="next" value="1">Next method: Sim_Resnik_1999</span> 

<div id="image" class="figure">
<img src="image/go_bp_random_500_sim_Sim_Lin_1998_Lin_order.png" width="700" />
<p class="caption">Figure S4.5. Similarity heatmap of the 500 random GO BP terms under different similarity methods.</p>
</div>


```{r}
sessionInfo()
```

<script>
methods = ["Sim_Lin_1998", "Sim_Resnik_1999", "Sim_FaITH_2010", "Sim_Relevance_2006", "Sim_SimIC_2010", "Sim_XGraSM_2013", "Sim_EISI_2015", "Sim_AIC_2014", "Sim_Zhang_2006", "Sim_universal", "Sim_Wang_2007", "Sim_GOGO_2018", "Sim_Rada_1989", "Sim_Resnik_edge_2005", "Sim_Leocock_1998", "Sim_WP_1994", "Sim_Slimani_2006", "Sim_Shenoy_2012", "Sim_Pekar_2002", "Sim_Stojanovic_2001", "Sim_Wang_edge_2012", "Sim_Zhong_2002", "Sim_AlMubaid_2006", "Sim_Li_2003", "Sim_RSS_2013", "Sim_HRSS_2013", "Sim_Shen_2010", "Sim_SSDD_2013", "Sim_Jiang_1997", "Sim_Kappa", "Sim_Jaccard", "Sim_Dice", "Sim_Overlap", "Sim_Ancestor"];
$("#select").on("change", function() {
    i = parseInt($(this).val());
    suffix = ".png"
    if($("#use_lin_order").is(':checked')) {
        suffix = "_Lin_order.png"
    }
    $("#image img").attr("src", "image/go_bp_random_500_sim_" + methods[i] + suffix);
    $("#curr").text("Curr method: " + methods[i]);

    prev_i = i - 1;
    if(prev_i < 0) {
        prev_i = 33
    }
    $("#prev").attr("value", prev_i);
    $("#prev").text("Prev method: " + methods[prev_i]);

    next_i = i + 1;
    if(next_i > 33) {
        next_i = 0;
    }
    $("#next").attr("value", next_i);
    $("#next").text("Next method: " + methods[next_i]);
});
$("#prev").click(function() {
    i = parseInt($(this).attr("value"));
    suffix = ".png"
    if($("#use_lin_order").is(':checked')) {
        suffix = "_Lin_order.png"
    }
    $("#image img").attr("src", "image/go_bp_random_500_sim_" + methods[i] + suffix);
    $("#curr").text("Curr method: " + methods[i]);

    prev_i = i - 1;
    if(prev_i < 0) {
        prev_i = 33
    }
    $("#prev").attr("value", prev_i);
    $("#prev").text("Prev method: " + methods[prev_i]);

    next_i = i + 1;
    if(next_i > 33) {
        next_i = 0;
    }
    $("#next").attr("value", next_i);
    $("#next").text("Next method: " + methods[next_i]);

    $("#select").val(i);
})
$("#next").click(function() {
    i = parseInt($(this).attr("value"));
    suffix = ".png"
    if($("#use_lin_order").is(':checked')) {
        suffix = "_Lin_order.png"
    }
    $("#image img").attr("src", "image/go_bp_random_500_sim_" + methods[i] + suffix);
    $("#curr").text("Curr method: " + methods[i]);

    prev_i = i - 1;
    if(prev_i < 0) {
        prev_i = 33
    }
    $("#prev").attr("value", prev_i);
    $("#prev").text("Prev method: " + methods[prev_i]);

    next_i = i + 1;
    if(next_i > 33) {
        next_i = 0;
    }
    $("#next").attr("value", next_i);
    $("#next").text("Next method: " + methods[next_i]);

    $("#select").val(i);
})
$("#use_lin_order").change(function() {
    i = parseInt($("#select").val());
    if(this.checked) {
        $("#image img").attr("src", "image/go_bp_random_500_sim_" + methods[i] + "_Lin_order.png");
    } else {
        $("#image img").attr("src", "image/go_bp_random_500_sim_" + methods[i] + ".png");
    }
})

</script>
````
