## Supplementary File 5 for "*simona:* a comprehensive R package for semantic similarity analysis on bio-ontologies": compare_topo_anno.html

Supplementary File 5. Compare topology-based and annotation-based semantic similarity methods


### Supplementary File 5. Compare topology-based and annotation-based semantic similarity methods

###### Zuguang Gu

#### 2024-08-28

- Compare MICA and LCA
- Semantic similarity based on MICA and LCA
- Session info

Existing R packages for semantic similarity analysis are only based on gene annotations, thus mainly applicable on Gene Ontology. Besides annotation-based methods, **simona** also supports a large number of methods based on the topology of GO DAG, with no need of external annotation data. In this document, we compare the semantic similarity calculated based on gene annotations and based on topology of the GO DAG.

We use Gene Ontology (Biological Process namespace) as the test ontology and human as the organism.

```
library(simona)
dag = create_ontology_DAG_from_GO_db(org_db = "org.Hs.eg.db")
dag
```

```
## An ontology_DAG object:
##   Source: GO BP / GO.db package 3.19.1 
##   27186 terms / 54178 relations
##   Root: GO:0008150 
##   Terms: GO:0000001, GO:0000002, GO:0000003, GO:0000011, ...
##   Max depth: 18 
##   Avg number of parents: 1.99
##   Avg number of children: 1.87
##   Aspect ratio: 356.46:1 (based on the longest distance from root)
##                 756.89:1 (based on the shortest distance from root)
##   Relations: is_a, part_of
##   Annotations: 18888 items
##                291, 1890, 4205, 4358, ...
## 
## With the following columns in the metadata data frame:
##   id, name, definition
```

Since some GO terms have no gene annotated and the information contents (IC) cannot be calculated, we random sample 500 GO terms that have genes annotations (i.e. with IC values).

```
set.seed(123)
ic = term_IC(dag, method = "IC_annotation")
ic = ic[!is.na(ic)]
go_id = sample(names(ic), 500)
```

#### Compare MICA and LCA

For the IC-based method (which is here annotation-based. We name it as “IC\_annotation” for easier discussion), the similarity is determined by the IC of the *most informative commmon ancestor (MICA)* of two terms, while for the topology-based method, the similarity is determined by the *lowest common ancestor (LCA)*. We compare, for a given set of GO terms, how different their MICA and LCA are.

We have already set the gene annotation when calling `create_ontology_DAG_from_GO_db()`, thus calling `MICA_term()` directly uses IC\_annotation as the IC method. `MICA_term()` and `LCA_term()` returns a symmetric matrix, thus we only take the lower triangle of the square matrix.

```
term1 = MICA_term(dag, go_id)
term1 = term1[lower.tri(term1)]

term2 = LCA_term(dag, go_id)
term2 = term2[lower.tri(term2)]
```

In all `500*(500-1)/2 = 124750` pairs of terms, we check how many MICA and LCA are the same.

```
sum(term1 == term2)
```

```
## [1] 122774
```

```
sum(term1 == term2)/length(term1)
```

```
## [1] 0.9841603
```

It shows for more than 98% of the term pairs, their MICA and LCA are the same terms. For the remaining 1976 (2%) term pairs, we check how different their MICA and LCA are.

```
l = term1 != term2
term1 = term1[l]
term2 = term2[l]
```

Although the MICA and LCA of these 1976 term pairs are different, we ask are their MICA and LCA in ancestor/offspring relations? I.e. is the MICA of two terms an ancestor or offspring of their LCA? The function `shortest_distances_directed()` calculates *directed* distance in the DAG. If term \(a\) cannot reach term \(b\), taking directions into consideration, the distance value is assigned to -1.

```
term_unique = unique(c(term1, term2))
dist = shortest_distances_directed(dag, term_unique) # a non-symmetric matrix with terms as names
x = numeric(length(term1)) # dist from term1 to term2 or from term2 to term1
for(i in seq_along(term1)) {
    x[i] = ifelse(dist[term1[i], term2[i]] >= 0, dist[term1[i], term2[i]], dist[term2[i], term1[i]]) 
}
table(x)
```

```
## x
##   -1 
## 1976
```

It shows for the 1976 term pairs in this random dataset, none of their MICA and LCA are in ancestor/offspring relations.

To demonstrate it, we can take the MICA and LCA of the first term pair as an example. Here the syntax `dag[, c(term1[1], term2[1])]` returns an induced sub-DAG only taking `c(term1[1], term2[1])` as leaf terms. We color the MICA term in red, and the LCA term in blue. `"GO:0008150"` is the root of the DAG (biological\_process). The following plot shows the MICA term “`GO:0044281`” and the LCA term `"GO:0044238"` although are not in ancestor/offspring relations, are still close in the DAG.

```
dag2 = dag[, c(term1[1], term2[1])]
color = c("red", "blue")
names(color) = c(term1[1], term2[1])

dag_graphviz(dag2, node_param = list(color = color))
```

Figure S5.1. Sub-DAG of ancestors of the MICA term GO:0044281 and the LCA term GO:0044238.

Based on the example graph above, we guess it might be a general pattern that the MICA and LCA of the 1976 term pairs are in sibling relations, i.e. under the same parent term, or at least very close to each other. Next we calculate the distance between MICA and LCA taking DAG as *undirected*. The function `shortest_distances_via_NCA()` calculates the shortest distance of two terms passing through their nearest common ancestor (NCA).

```
term_unique = unique(c(term1, term2))
dist = shortest_distances_via_NCA(dag, term_unique) # a symmetric matrix with terms as names
x = numeric(length(term1))
for(i in seq_along(term1)) {
    x[i] = dist[term1[i], term2[i]]
}
barplot(table(x), 
    xlab = "Undirected distance between MICA and LCA of a term pair", 
    ylab = "Number of term pairs")
```

Figure S5.2. Numbers of term pairs whose MICA and LCA have different distances.

The barplot shows 73.0% of the 1976 term pairs, their MICA and LCA are siblings (i.e. sharing the same parent, with undirected distance of 2). Most of the MICA and LCA are also close in the DAG for other term pairs.

#### Semantic similarity based on MICA and LCA

In general, based on the results we have generated, we could conclude that the MICA method and the LCA method are very similar. In most of the time, they are the same set or very close in the DAG.

If thinking depth as a generalized measure of information content where deeper in the DAG, a larger value of of depth a term will have (note the depth of a parent is always smaller than the depth of its child terms because the depth is the longest distance from root), In this way, LCA is also a type of MICA. we compare the relations between the global IC\_annotation and depth.

```
ic = term_IC(dag, method = "IC_annotation")
depth = dag_depth(dag)

par(mfrow = c(1, 2))
# because depth is discrete, we use boxplot to visualize the relations
boxplot(ic ~ depth, xlab = "Depth", ylab = "IC_annotation")
barplot(table(depth), xlab = "Depth", ylab = "Number of terms")
```

Figure S5.3. Relations between IC\_annotation and depth. Left: boxplots of IC\_annotation on each depth; Right: Numbers of terms on each depth.

It shows depth and IC\_annotation have very strong positive relations. Together with the high compatible sets of MICA and LCA, it can be expected that the semantic similarities based on MICA and LCA should also be highly similar.

In the following example, we compare `"Sim_Lin_1998"` and `"Sim_WP_1994"` methods. `"Sim_Lin_1998"` is based on the IC\_annotation of MICA and `"Sim_WP_1994"` is based on the depth of LCA. The two methods have similar forms.

For `"Sim_Lin_1998"`, the similarity is defined as (let \(c\) be the MICA of \(a\) and \(b\)):

\[ \frac{2\*\mathrm{IC}(c)}{\mathrm{IC}(a) + \mathrm{IC}(b)} \]

For `"Sim_WP_1994"`, the similarity is defined as (let \(c\) be the LCA of \(a\) and \(b\)):

\[ \frac{2\*\mathrm{len}(r, c)}{\mathrm{len}\_c(r, a) + \mathrm{len}\_c(r, a)} \]

where \(\mathrm{len}(r, c)\) is the longest distance from root \(r\) to \(c\), i.e. the depth of \(c\), and \(\mathrm{len}\_c(r, a)\) is the longest distance from \(r\) to \(a\), passing through \(c\).

Note that the following two heatmaps share the same row orders and column orders.

- Use Sim\_Lin\_1998 order
- Use Sim\_WP\_1994 order

```
mat1 = term_sim(dag, go_id, method = "Sim_Lin_1998")
mat2 = term_sim(dag, go_id, method = "Sim_WP_1994")

od = hclust(dist(mat1))$order

library(ComplexHeatmap)

Heatmap(mat1, name = "similarity_lin", 
        show_row_names = FALSE, show_column_names = FALSE,
        row_order = od, column_order = od,
        column_title = "Sim_Lin_1998, based on MICA/IC_annotation") +
Heatmap(mat2, name = "similarity_wp", 
        show_row_names = FALSE, show_column_names = FALSE,
        row_order = od, column_order = od,
        column_title = "Sim_WP_1994, based on LCA/depth")
```

Figure S5.4. Heatmap of semantic similarities from the Lin 1998 method and WP 1994 method. The two heatmaps have the same row orders and column orders.

```
mat1 = term_sim(dag, go_id, method = "Sim_Lin_1998")
mat2 = term_sim(dag, go_id, method = "Sim_WP_1994")

od = hclust(dist(mat2))$order

library(ComplexHeatmap)

Heatmap(mat1, name = "similarity_lin", 
        show_row_names = FALSE, show_column_names = FALSE,
        row_order = od, column_order = od,
        column_title = "Sim_Lin_1998, based on MICA/IC_annotation") +
Heatmap(mat2, name = "similarity_wp", 
        show_row_names = FALSE, show_column_names = FALSE,
        row_order = od, column_order = od,
        column_title = "Sim_WP_1994, based on LCA/depth")
```

Figure S5.4. Heatmap of semantic similarities from the Lin 1998 method and WP 1994 method.

The two heatmaps show very similar patterns. Heatmap under `"Sim_WP_1994"` in general shows overall stronger similarity signals, but also higher level of “inter-block” signals.

And the scatter plot of the similarity values from the two methods:

```
plot(mat1[lower.tri(mat1)], mat2[lower.tri(mat2)], pch = 16, col = "#00000010",
    xlab = "Sim_Lin_1998 / IC_annotation / MICA", ylab = "Sim_WP_1994 / depth / LCA")
```

Figure S5.5. Scatterplot of semantic similarities calculated by the Lin 1998 method and WP 1994 method.

It is actually not strange that MICA/IC\_annotation method is very similar to LCA/depth method, because IC\_annotation is calculated based on the aggregation of gene annotations from offspring terms, which actually also takes account of the DAG topology.

Finally, let’s make a simplified example to show the relations between IC\_annotation and depth. Consider an ontology with a binary tree structure, i.e. each term has two children and each term has an unique gene annotated. All the leaf terms in the ontology have depth of \(d\_\mathrm{max}\). Then the total number of genes annotated to the root is \(2^{d\_\mathrm{max}+1}-1\). For a term with depth \(d\) where \(0 \leq d \leq d\_\mathrm{max}\), the number of genes annotated to it is \(2^{d\_\mathrm{max}-d+1}-1\), then IC\_annotation of this term would be:

\[
\begin{align\*}
\mathrm{IC}\_\mathrm{annotation} & = -\log\left(\frac{2^{d\_\mathrm{max}-d + 1} - 1}{2^{d\_\mathrm{max}+1} - 1} \right) \\
& \approx -\log\left(\frac{2^{d\_\mathrm{max}-d + 1}}{2^{d\_\mathrm{max}+1}} \right) \\
& = -\log\left(2^{-d} \right ) \\
& = d \* \log2
\end{align\*} \]

#### Session info

```
sessionInfo()
```

```
## R version 4.4.1 (2024-06-14)
## Platform: x86_64-apple-darwin20
## Running under: macOS Sonoma 14.5
## 
## Matrix products: default
## BLAS:   /Library/Frameworks/R.framework/Versions/4.4-x86_64/Resources/lib/libRblas.0.dylib 
## LAPACK: /Library/Frameworks/R.framework/Versions/4.4-x86_64/Resources/lib/libRlapack.dylib;  LAPACK version 3.12.0
## 
## locale:
## [1] en_GB.UTF-8/en_GB.UTF-8/en_GB.UTF-8/C/en_GB.UTF-8/en_GB.UTF-8
## 
## time zone: Europe/Berlin
## tzcode source: internal
## 
## attached base packages:
## [1] grid      stats     graphics  grDevices utils     datasets  methods  
## [8] base     
## 
## other attached packages:
## [1] ComplexHeatmap_2.20.0 simona_1.3.12         knitr_1.48           
## [4] rmarkdown_2.28       
## 
## loaded via a namespace (and not attached):
##  [1] KEGGREST_1.44.1         circlize_0.4.16         shape_1.4.6.1          
##  [4] rjson_0.2.22            xfun_0.47               bslib_0.8.0            
##  [7] visNetwork_2.1.2        htmlwidgets_1.6.4       GlobalOptions_0.1.2    
## [10] Biobase_2.64.0          Cairo_1.6-2             vctrs_0.6.5            
## [13] tools_4.4.1             stats4_4.4.1            parallel_4.4.1         
## [16] Polychrome_1.5.1        AnnotationDbi_1.66.0    RSQLite_2.3.7          
## [19] highr_0.11              cluster_2.1.6           blob_1.2.4             
## [22] pkgconfig_2.0.3         RColorBrewer_1.1-3      S4Vectors_0.42.1       
## [25] scatterplot3d_0.3-44    GenomeInfoDbData_1.2.12 lifecycle_1.0.4        
## [28] compiler_4.4.1          Biostrings_2.72.1       codetools_0.2-20       
## [31] clue_0.3-65             httpuv_1.6.15           GenomeInfoDb_1.40.1    
## [34] htmltools_0.5.8.1       sass_0.4.9              yaml_2.3.10            
## [37] later_1.3.2             crayon_1.5.3            jquerylib_0.1.4        
## [40] GO.db_3.19.1            cachem_1.1.0            org.Hs.eg.db_3.19.1    
## [43] iterators_1.0.14        foreach_1.5.2           mime_0.12              
## [46] digest_0.6.37           fastmap_1.2.0           colorspace_2.1-1       
## [49] cli_3.6.3               DiagrammeR_1.0.11       magrittr_2.0.3         
## [52] UCSC.utils_1.0.0        promises_1.3.0          bit64_4.0.5            
## [55] XVector_0.44.0          httr_1.4.7              matrixStats_1.3.0      
## [58] igraph_2.0.3            bit_4.0.5               png_0.1-8              
## [61] GetoptLong_1.0.5        memoise_2.0.1           shiny_1.9.1            
## [64] evaluate_0.24.0         IRanges_2.38.1          doParallel_1.0.17      
## [67] rlang_1.1.4             Rcpp_1.0.13             glue_1.7.0             
## [70] xtable_1.8-4            DBI_1.2.3               xml2_1.3.6             
## [73] BiocGenerics_0.50.0     rstudioapi_0.16.0       jsonlite_1.8.8         
## [76] R6_2.5.1                zlibbioc_1.50.0
```
