## Supplementary File 6 for "*simona:* a comprehensive R package for semantic similarity analysis on bio-ontologies": OBOFoundry.R

```
library(simona)
library(GetoptLong)
library(ComplexHeatmap)

setwd("~/manuscript/simona/suppl6_OBOFoundry_gallery")

# http://obofoundry.org/

# OBOFoundry_meta_table.rds is imported from http://obofoundry.org/registry/ontologies.jsonld
meta = readRDS("OBOFoundry_meta_table.rds")

# all the ontology files are downloaded by systme.file("scripts", "download.R", package = "simona")
files = scan(pipe("ls ~/workspace/ontology/OBOFoundry/*/*"), what = "character")
files = files[grep("(obo|owl|xml|json|rds|ttl)", files)]


df = data.frame(
	file = files,
	id = basename(dirname(files)),
	basename = basename(files),
	type = gsub("^.*?\\.", "", basename(files))
)

weight = c("obo" = 1, "owl" = 2, "rds" = 3, "ttl" = 4, "xml" = 5, "json" = 6)
i = tapply(seq_len(nrow(df)), df$id, function(ind) {
	ind[which.min(weight[df$type[ind]] + ifelse(grepl("base", df$basename[ind]), 1, 0))]
})

df = df[i, ]

id2title = structure(names = meta$id, meta$title)

df$title = id2title[df$id]

id2purl = structure(names = meta$id, meta$ontology_purl)

df$purl = id2purl[df$id]

df$n_terms = -1
df$n_relations = -1
df$avg_parents = -1
df$avg_children = -1
df$asp1 = -1
df$asp2 = -1
df$depth_max = -1
df$depth_q99 = -1
df$method = ""


for(i in seq_len(nrow(df))) {
	qqcat("\n===============@{i}, @{basename(df$file[i])}===========\n")
	oe = try({ 
		if(df$type[i] == "obo") {
			dag = import_obo(df$file[i], remove_cyclic_paths = TRUE, remove_rings = TRUE)
			df[i, "method"] = "import_obo"
		} else if(df$type[i] == "owl") {
			dag = import_owl(df$file[i], remove_cyclic_paths = TRUE, remove_rings = TRUE)
			df[i, "method"] = "import_owl"
		} else if(df$type[i] == "ttl") {
			dag = import_ttl(df$file[i], remove_cyclic_paths = TRUE, remove_rings = TRUE)
			df[i, "method"] = "import_ttl"
		} 
	})
	oe = try({
		if(inherits(oe, "try-error")) {
			dag = import_ontology(df$file[i], remove_cyclic_paths = TRUE, remove_rings = TRUE, robot_jar = "~/Downloads/robot.jar")
			df[i, "method"] = "import_ontology"
		} else if(! df$type[i] %in% c("obo", "owl", "ttl")) {
			dag = import_ontology(df$file[i], remove_cyclic_paths = TRUE, remove_rings = TRUE, robot_jar = "~/Downloads/robot.jar")
			df[i, "method"] = "import_ontology"
		}
		
	})

	if(inherits(oe, "try-error")) {
		next
	}

	df[i, "n_terms"] = dag_n_terms(dag)
	df[i, "n_relations"] = sum(sapply(dag@lt_children, length))
	df[i, "avg_parents"] = mean(n_parents(dag)[-dag_root(dag, in_labels = FALSE)])
	df[i, "avg_children"] = mean(n_children(dag)[-dag_leaves(dag, in_labels = FALSE)])
	df[i, "asp1"] = dag@aspect_ratio[1]
	df[i, "asp2"] = dag@aspect_ratio[2]
	df[i, "depth_max"] = max(dag_depth(dag))
	df[i, "depth_q99"] = quantile(dag_depth(dag), 0.99)

	png(qq("image/OBOFoundry_@{df[i, 'id']}_foo.png"), width = 1000*1.5, height = 800*1.5, res = 72*1.5)
	dag_circular_viz(dag, partition_by_size = round(dag_n_terms(dag)/5))
	dev.off()

	system(qq("pngquant --speed=10 --quality=60 image/OBOFoundry_@{df[i, 'id']}_foo.png -o image/OBOFoundry_@{df[i, 'id']}.png --force"))
	file.remove(qq("image/OBOFoundry_@{df[i, 'id']}_foo.png"))

	terms = random_terms(dag, min(500, dag_n_terms(dag)))
	sim = term_sim(dag, terms, method = "Sim_WP_1994")
	png(qq("image/OBOFoundry_@{df[i, 'id']}_heatmap.png"), width = 600*1.5, height = 600*1.5, res = 72*1.5)
	draw(Heatmap(sim, name = "Sim_WP_1994", 
		column_title = qq("@{df$id[i]}, @{length(terms)} random terms"),
		show_row_names = FALSE, show_column_names = FALSE,
		show_row_dend = FALSE, show_column_dend = FALSE))
	dev.off()

}

saveRDS(df, file = "OBOFoundry_summary.rds")
```
