## Supplementary File 6 for "*simona:* a comprehensive R package for semantic similarity analysis on bio-ontologies": OBOFoundry_viz.html

Supplementary File 6. OBO Foundry gallery


### Supplementary File 6. OBO Foundry gallery

###### Zuguang Gu

#### 2024-08-29

OBO Foundry is a database of public biological ontologies. In this document, we import all ontologies on OBO Foundry and generate circular visualization, similarity heatmap and runtime performance on them.

Columns in the table:

- `id`: ID of the ontology.
- `title`: Title of the ontology.
- `n_terms`: Number of terms.
- `n_relations`: Number of relations.
- `is_tree`: Whether the DAG is a tree. The DAG is a tree if n\_terms == n\_relations + 1.
- `pa_avg`: Average number of parent terms.
- `ch_avg`: Average number of child terms.
- `asp1`: `width/height`, where `width` is the largest number of terms on a specific distance to root (i.e. `max(table(largest_dist_to_root(dag)))`). `height` is the 99% percentile of heights of all terms.
- `asp2`: `width/height`, where `width` is the largest number of terms on a specific distance to root (i.e. `max(table(shortest_dist_to_root(dag)))`). `height` is the 99% percentile of heights of all terms.
- `d_max`: Maximal depth of terms.


---


---

[close]

##### Plots for ontology: ado <ado.owl>

- Circular visualization
- Similarity heatmap
- Runtime

```
# the circular visualization is generated by:
dag_circular_viz(dag, partition_by_size = round(dag_n_terms(dag)/5))
```

```
# the heatmap is generated by:
terms = random_terms(dag, min(500, dag_n_terms(dag)))
sim = term_sim(dag, terms, method = "Sim_WP_1994")
ComplexHeatmap::Heatmap(sim, name = "Sim_WP_1994",
    show_row_names = FALSE, show_column_names = FALSE,
    show_row_dend = FALSE, show_column_dend = FALSE)
```

The runtime is tested by different numbers of terms randomly picked from the ontology.

```
benchmark_runtime = function(dag) {
    invisible(dag_depth(dag))  # depth will be cached

    n_terms = dag_n_terms(dag)
    k = seq(100, min(20000, n_terms), length = 50)
    k = floor(k)
    t = rep(NA_real_, length(k))
    n_an = rep(0, length(k))
    for(i in seq_along(k)) {
        terms = sample(n_terms, k[i])
        t[i] = system.time(LCA_term(dag, terms))[3]
    }
    data.frame(k = k, t = t)
}
df = benchmark_runtime(dag)
plot(df$k, df$t, xlab = "Number of random terms", ylab = "Runtime (sec)", 
    main = "Runtime for LCA_term()")
x = c(0, df$k)
y = c(0, df$t)
fit = loess(y ~ x, span = 0.5)
lines(x, predict(fit))
legend("topleft", lty = 1, legend = "loess fit")
```
