## Supplementary File 6 for "*simona:* a comprehensive R package for semantic similarity analysis on bio-ontologies": OBOFoundry_viz.Rmd

````
---
title: "Supplementary File 6. OBO Foundry gallery"
author: "Zuguang Gu"
date: '`r Sys.Date()`'
output: 
  html_document:
    toc: true
---

```{r, echo = FALSE, message = FALSE}
library(knitr)
knitr::opts_chunk$set(
    error = FALSE,
    tidy  = FALSE,
    message = FALSE,
    warning = FALSE,
    fig.width = 6, fig.height = 6,
    fig.align = "center")
```

[OBO Foundry](http://obofoundry.org/) is a database of public biological
ontologies. In this document, we import all ontologies on OBO Foundry and
generate circular visualization, similarity heatmap and runtime performance on them.

<details>

<summary>Columns in the table:</summary>

- `id`: ID of the ontology.
- `title`: Title of the ontology.
- `n_terms`: Number of terms.
- `n_relations`: Number of relations.
- `is_tree`: Whether the DAG is a tree. The DAG is a tree if n_terms == n_relations + 1.
- `pa_avg`: Average number of parent terms.
- `ch_avg`: Average number of child terms.
- `asp1`: `width/height`, where `width` is the largest number of terms on a specific distance to root (i.e. `max(table(largest_dist_to_root(dag)))`). `height` is the 99% percentile of heights of all terms.
- `asp2`: `width/height`, where `width` is the largest number of terms on a specific distance to root (i.e. `max(table(shortest_dist_to_root(dag)))`). `height` is the 99% percentile of heights of all terms.
- `d_max`: Maximal depth of terms.

</details>


<hr>

```{r, echo = FALSE}
library(DT)
library(GetoptLong)
df = readRDS("OBOFoundry_summary.rds")
df = df[df$n_terms > 0, ]
id = df$id
df$id = qq("<a class='show_image' onclick='show_image(\"@{id}\", \"@{df$basename}\")'>@{df$id}</a>", collapse = FALSE)
df$avg_parents = round(df$avg_parents, 1)
df$avg_children = round(df$avg_children, 1)
df$is_tree = ifelse(df$n_terms == df$n_relations + 1, "tree", "")
df = df[, c("id", "title", "n_terms", "n_relations", "is_tree", "avg_parents", "avg_children", "asp1", "asp2", "depth_max")]
df = df[order(-df$n_terms), ]
datatable(df, escape = FALSE, rownames = FALSE,
    colnames = c("id", "title", "n_terms", "n_relations", "is_tree", "pa_avg", "ch_avg", "asp1", "asp2", 'd_max'))
```

<hr>

<style>
.show_image {
    text-decoration: underline;
    color: #337ab7;
}
.show_image:hover {
    cursor: pointer;
}
#image_wrapper {
    width:1200px;
    height:1300px;
    display:none;
    background-color: rgba(0, 0, 0, 0.5);
}
#image {
    width: 1100px;
    margin:20px auto;
    padding:20px;
    border: 1px solid grey;
    background-color: white;
}
#image ul li {
    font-size: 16px;
    padding-right:10px;
    cursor: pointer;
    display: inline-block;
    color: #337ab7;
}
#image ul {
    padding-inline-start: 0px;
    margin-top: 20px;
}
pre.code {
    background-color:#EFEFEF;
}
#close_div:hover{
    cursor: pointer;
}
</style>

<div id="image_wrapper">
<div id="image">
<p><a onclick='$("#image_wrapper").css("display", "none");' id="close_div">[close]</a><p>
<h3>Plots for ontology: <a href='http://obofoundry.org/ontology/ado.html' target="_blank">ado 
    <svg xmlns="http://www.w3.org/2000/svg" width="16" height="16" fill="currentColor" class="bi bi-box-arrow-up-right" viewBox="0 0 16 16">
  <path fill-rule="evenodd" d="M8.636 3.5a.5.5 0 0 0-.5-.5H1.5A1.5 1.5 0 0 0 0 4.5v10A1.5 1.5 0 0 0 1.5 16h10a1.5 1.5 0 0 0 1.5-1.5V7.864a.5.5 0 0 0-1 0V14.5a.5.5 0 0 1-.5.5h-10a.5.5 0 0 1-.5-.5v-10a.5.5 0 0 1 .5-.5h6.636a.5.5 0 0 0 .5-.5"/>
  <path fill-rule="evenodd" d="M16 .5a.5.5 0 0 0-.5-.5h-5a.5.5 0 0 0 0 1h3.793L6.146 9.146a.5.5 0 1 0 .708.708L15 1.707V5.5a.5.5 0 0 0 1 0z"/>
</svg>
</a> &lt;ado.owl&gt;</h3>
<ul>
<li><a id="circular_click">Circular visualization</a></li>
<li><a id="heatmap_click" style="color:grey">Similarity heatmap</a></li>
<li><a id="runtime_click" style="color:grey">Runtime</a></li>
</ul>
<div id="circular">
<pre class="code"># the circular visualization is generated by:
dag_circular_viz(dag, partition_by_size = round(dag_n_terms(dag)/5))
</pre>
<img id="image1" src="image/OBOFoundry_ado.png" /></div>
<div id="heatmap" style="display:none;">
<pre class="code"># the heatmap is generated by:
terms = random_terms(dag, min(500, dag_n_terms(dag)))
sim = term_sim(dag, terms, method = "Sim_WP_1994")
ComplexHeatmap::Heatmap(sim, name = "Sim_WP_1994",
    show_row_names = FALSE, show_column_names = FALSE,
    show_row_dend = FALSE, show_column_dend = FALSE)
</pre>
<img id="image2" src="image/OBOFoundry_ado_heatmap.png" width=600 />
</div>
<div id="runtime" style="display:none;">
<p>The runtime is tested by different numbers of terms randomly picked from the ontology.</p>
<pre class="code">benchmark_runtime = function(dag) {
    invisible(dag_depth(dag))  # depth will be cached

    n_terms = dag_n_terms(dag)
    k = seq(100, min(20000, n_terms), length = 50)
    k = floor(k)
    t = rep(NA_real_, length(k))
    n_an = rep(0, length(k))
    for(i in seq_along(k)) {
        terms = sample(n_terms, k[i])
        t[i] = system.time(LCA_term(dag, terms))[3]
    }
    data.frame(k = k, t = t)
}
df = benchmark_runtime(dag)
plot(df$k, df$t, xlab = "Number of random terms", ylab = "Runtime (sec)", 
    main = "Runtime for LCA_term()")
x = c(0, df$k)
y = c(0, df$t)
fit = loess(y ~ x, span = 0.5)
lines(x, predict(fit))
legend("topleft", lty = 1, legend = "loess fit")
</pre>
<img id="image3" src="image/OBOFoundry_ado_runtime.png" width=600 />
</div>
</div>
</div>


<script>
function show_image(id, basename) {

    $("#image1").attr("src", "image/OBOFoundry_" + id + ".png");
    $("#image2").attr("src", "image/OBOFoundry_" + id + "_heatmap.png");
    $("#image3").attr("src", "image/OBOFoundry_" + id + "_runtime.png");
    $("#image h3").html("Plots for ontology: <a href='http://obofoundry.org/ontology/" + id + ".html' target='_blank'>" + id + ' <svg xmlns="http://www.w3.org/2000/svg" width="16" height="16" fill="currentColor" class="bi bi-box-arrow-up-right" viewBox="0 0 16 16"><path fill-rule="evenodd" d="M8.636 3.5a.5.5 0 0 0-.5-.5H1.5A1.5 1.5 0 0 0 0 4.5v10A1.5 1.5 0 0 0 1.5 16h10a1.5 1.5 0 0 0 1.5-1.5V7.864a.5.5 0 0 0-1 0V14.5a.5.5 0 0 1-.5.5h-10a.5.5 0 0 1-.5-.5v-10a.5.5 0 0 1 .5-.5h6.636a.5.5 0 0 0 .5-.5"/> <path fill-rule="evenodd" d="M16 .5a.5.5 0 0 0-.5-.5h-5a.5.5 0 0 0 0 1h3.793L6.146 9.146a.5.5 0 1 0 .708.708L15 1.707V5.5a.5.5 0 0 0 1 0z"/></svg>' + "</a> &lt;" + basename + "&gt;");

    $("#image_wrapper").css("display", "block");
    $("#image_wrapper").css("position", "absolute");
    $("#image_wrapper").css("top", "0px");

    document.getElementById("image_wrapper").scrollIntoView();
}

$( function() {

    $( "#circular_click" ).click(function() {
        $("#circular").show();
        $("#heatmap").hide();
        $("#runtime").hide();
        $("#circular_click").css("color", "#337ab7");
        $("#heatmap_click").css("color", "grey");
        $("#runtime_click").css("color", "grey");
    });
    $( "#heatmap_click" ).click(function() {
        $("#heatmap").show();
        $("#circular").hide();
        $("#runtime").hide();
        $("#circular_click").css("color", "grey");
        $("#heatmap_click").css("color", "#337ab7");
        $("#runtime_click").css("color", "grey");
    })
    $( "#runtime_click" ).click(function() {
        $("#runtime").show();
        $("#heatmap").hide();
        $("#circular").hide();
        $("#runtime_click").css("color", "#337ab7");
        $("#heatmap_click").css("color", "grey");
        $("#circular_click").css("color", "grey");
    });

    $(".main-container").css("max-width", "1200px");
} );
</script>
````
