## Supplementary File 6 for "*simona:* a comprehensive R package for semantic similarity analysis on bio-ontologies": run_time_OBOFoundry.R

```
benchmark_runtime = function(dag) {
    invisible(dag_depth(dag))  # depth will be cached

    n_terms = dag_n_terms(dag)
    k = seq(100, min(20000, n_terms), length = 50)
    k = floor(k)
    t = rep(NA_real_, length(k))
    n_an = rep(0, length(k))
    for(i in seq_along(k)) {
        message(k[i], "/", max(k), "...")
        terms = sample(n_terms, k[i]) # numeric indicies are also allowed

        t[i] = system.time(LCA_term(dag, terms))[3]
    }
    data.frame(k = k, t = t)
}


library(simona)
library(GetoptLong)

# all the ontology files are downloaded by systme.file("scripts", "download.R", package = "simona")
files = scan(pipe("ls ~/workspace/ontology/OBOFoundry/*/*"), what = "character")
files = files[grep("(obo|owl|xml|json|rds|ttl)", files)]

df = data.frame(
    file = files,
    id = basename(dirname(files)),
    basename = basename(files),
    type = gsub("^.*?\\.", "", basename(files))
)

weight = c("obo" = 1, "owl" = 2, "rds" = 3, "ttl" = 4, "xml" = 5, "json" = 6)
i = tapply(seq_len(nrow(df)), df$id, function(ind) {
    ind[which.min(weight[df$type[ind]] + ifelse(grepl("base", df$basename[ind]), 1, 0))]
})

df = df[i, ]


lt = list()
for(i in seq_len(nrow(df))) {
    qqcat("\n===============@{i}, @{basename(df$file[i])}===========\n")
    oe = try({ 
        if(df$type[i] == "obo") {
            dag = import_obo(df$file[i], remove_cyclic_paths = TRUE, remove_rings = TRUE)
            df[i, "method"] = "import_obo"
        } else if(df$type[i] == "owl") {
            dag = import_owl(df$file[i], remove_cyclic_paths = TRUE, remove_rings = TRUE)
            df[i, "method"] = "import_owl"
        } else if(df$type[i] == "ttl") {
            dag = import_ttl(df$file[i], remove_cyclic_paths = TRUE, remove_rings = TRUE)
            df[i, "method"] = "import_ttl"
        } 
    })
    oe = try({
        if(inherits(oe, "try-error")) {
            dag = import_ontology(df$file[i], remove_cyclic_paths = TRUE, remove_rings = TRUE, robot_jar = "~/Downloads/robot.jar")
            df[i, "method"] = "import_ontology"
        } else if(! df$type[i] %in% c("obo", "owl", "ttl")) {
            dag = import_ontology(df$file[i], remove_cyclic_paths = TRUE, remove_rings = TRUE, robot_jar = "~/Downloads/robot.jar")
            df[i, "method"] = "import_ontology"
        }
        
    })

    if(inherits(oe, "try-error")) {
        next
    }

    df[i, "n_terms"] = dag_n_terms(dag)
    df[i, "n_relations"] = sum(sapply(dag@lt_children, length))
    df[i, "avg_parents"] = mean(n_parents(dag)[-dag_root(dag, in_labels = FALSE)])
    df[i, "avg_children"] = mean(n_children(dag)[-dag_leaves(dag, in_labels = FALSE)])
    df[i, "asp1"] = dag@aspect_ratio[1]
    df[i, "asp2"] = dag@aspect_ratio[2]
    df[i, "depth_max"] = max(dag_depth(dag))
    df[i, "depth_q99"] = quantile(dag_depth(dag), 0.99)

    nm = df$id[i]

    if(dag_n_terms(dag) < 1000) {
        png(qq("image/OBOFoundry_@{nm}_runtime.png"), width = 600*1.5, height = 600*1.5, res= 72*1.5)
        plot(NULL, xlim = c(0, 1), ylim = c(0, 1), axes = FALSE, ann = FALSE)
        text(0.5, 0.5, "runtime analysis only for n_terms >= 1000", cex = 1.5)
        dev.off()
    } else {
        lt[[nm]] = benchmark_runtime(dag)

        png(qq("image/OBOFoundry_@{nm}_runtime.png"), width = 600*1.5, height = 600*1.5, res= 72*1.5)
        plot(lt[[nm]]$k, lt[[nm]]$t, xlab = "Number of random terms", ylab = "Runtime (sec)", 
            main = paste0("Runtime for LCA_term(), ", nm, ": ", df[i, "n_terms"], " terms"))
        x = c(0, lt[[nm]]$k)
        y = c(0, lt[[nm]]$t)
        fit = loess(y ~ x, span = 0.5)
        lines(x, predict(fit))

        legend("topleft", lty = 1, legend = "loess fit")
        dev.off()
    }
}


rownames(df) = df$id
df = df[names(lt), ]

save(lt, df, file = "runtime_OBOFoundry_all.RData")
```
