## Supplementary figures and images for "*simona:* a comprehensive R package for semantic similarity analysis on bio-ontologies"

### OBOFoundry_ado.png

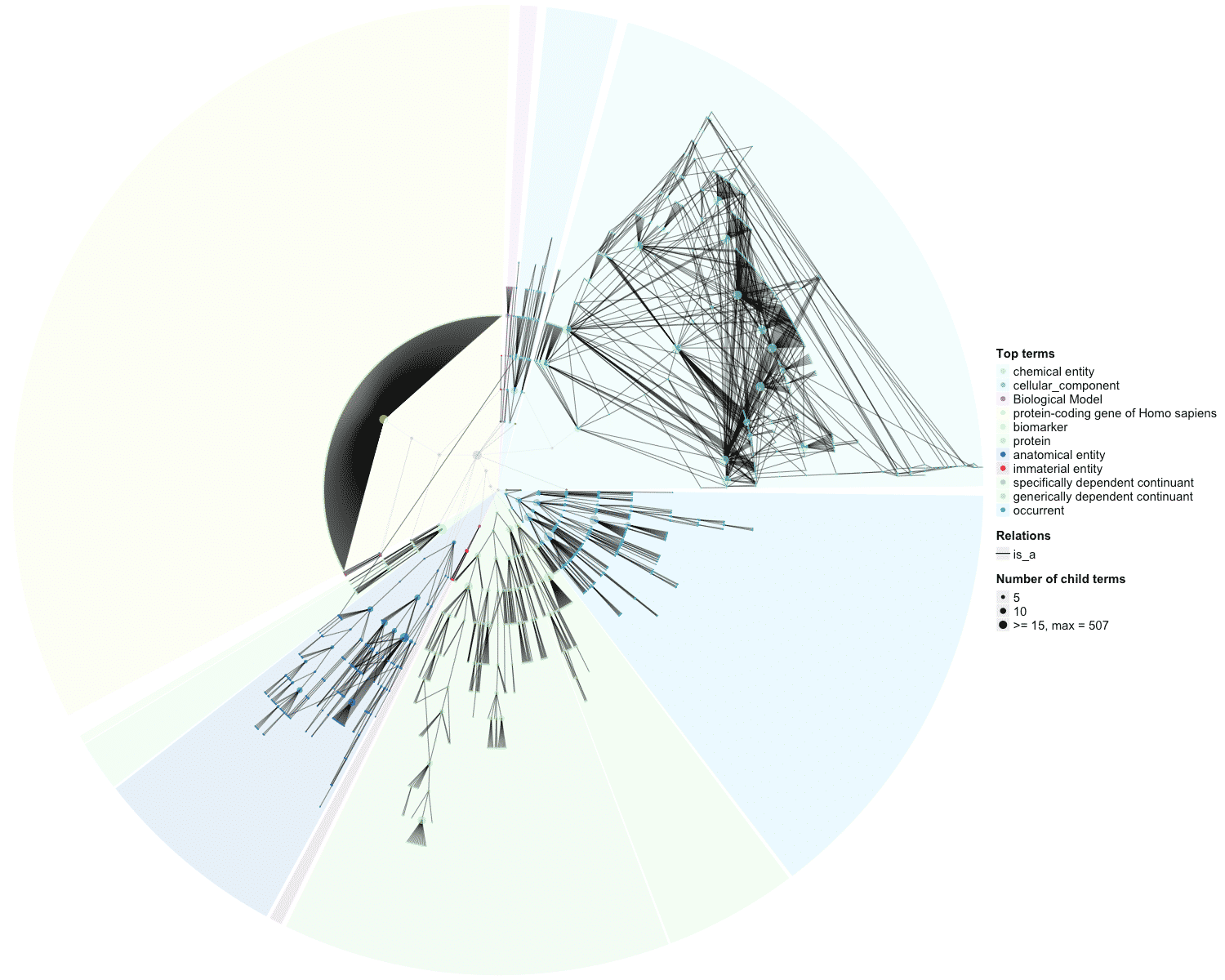

### OBOFoundry_aeo_heatmap.png

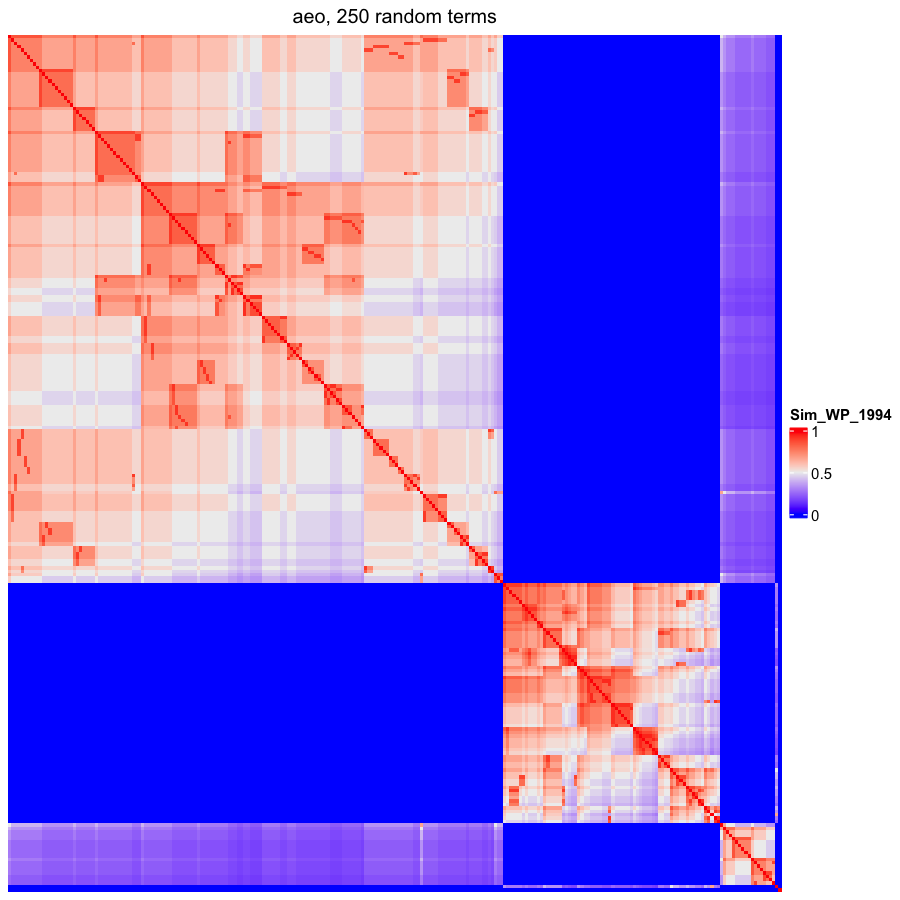

### OBOFoundry_cdao.png

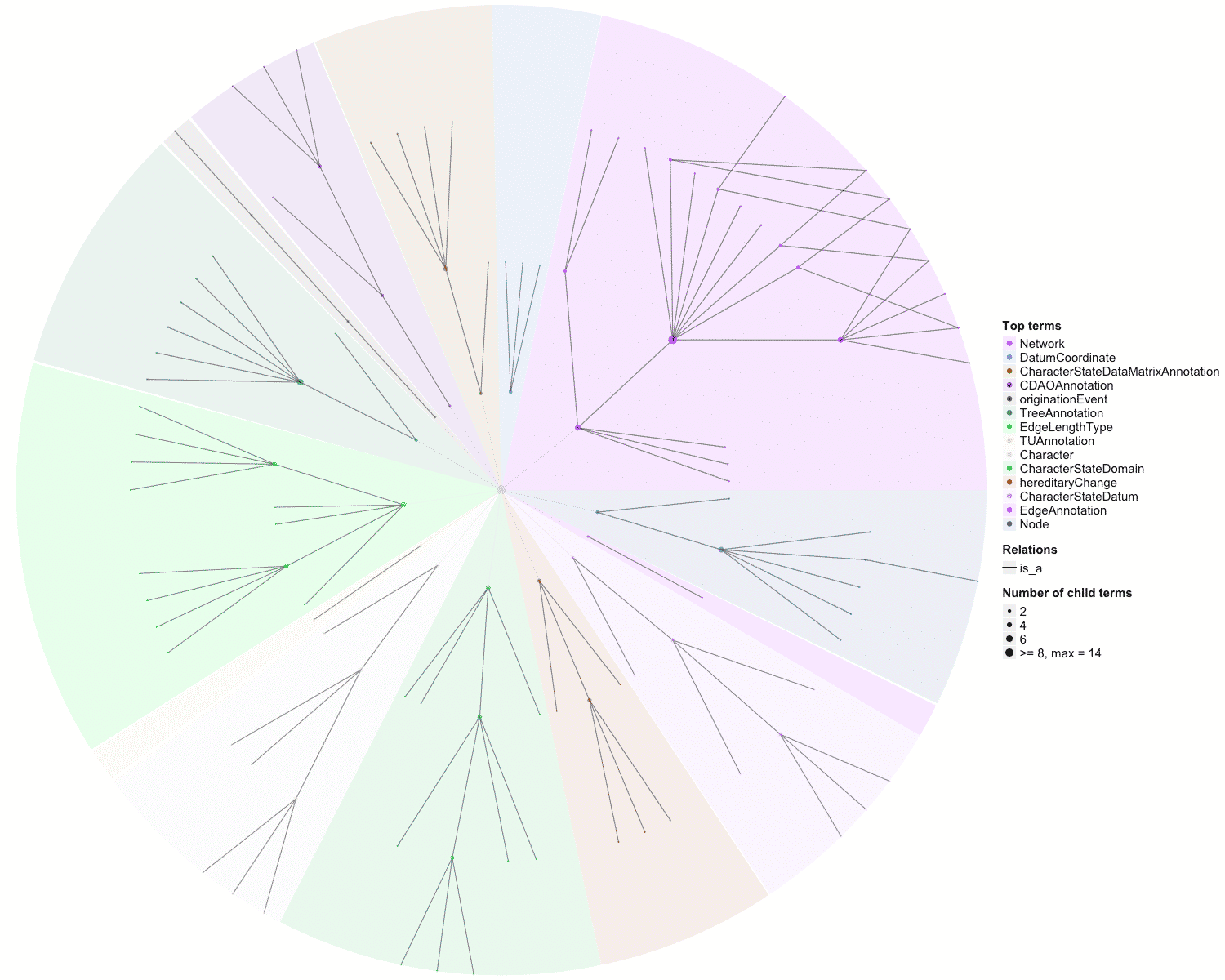

### OBOFoundry_chmo_runtime.png

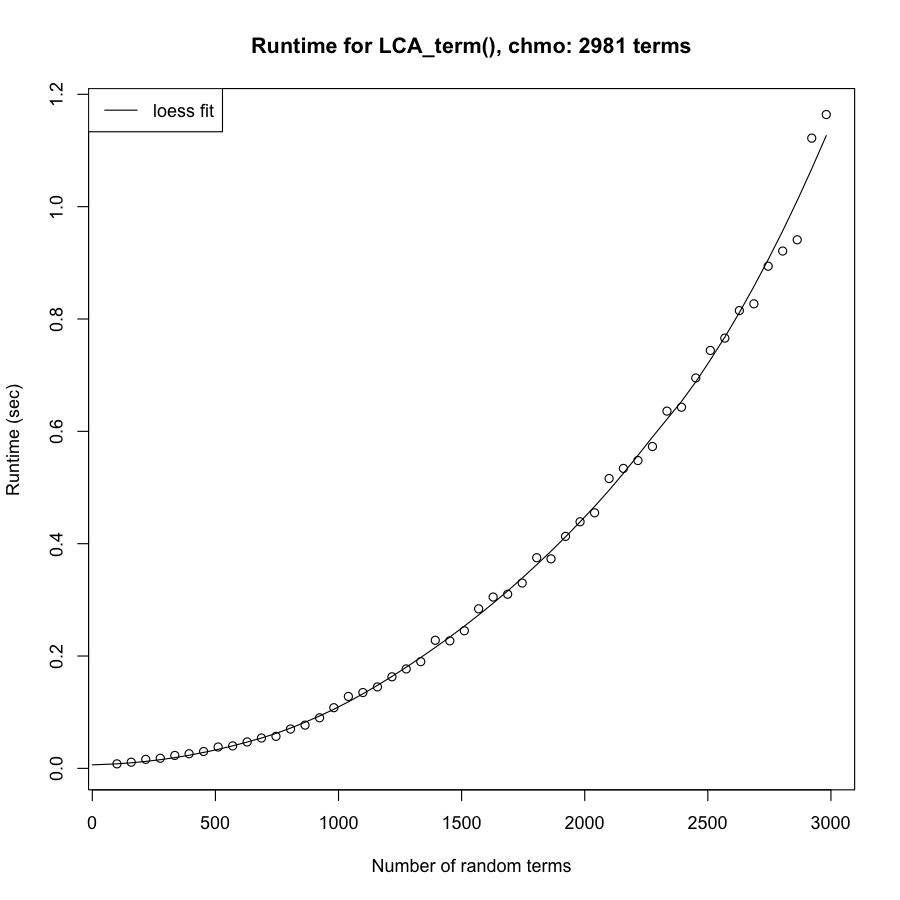

### OBOFoundry_doid.png

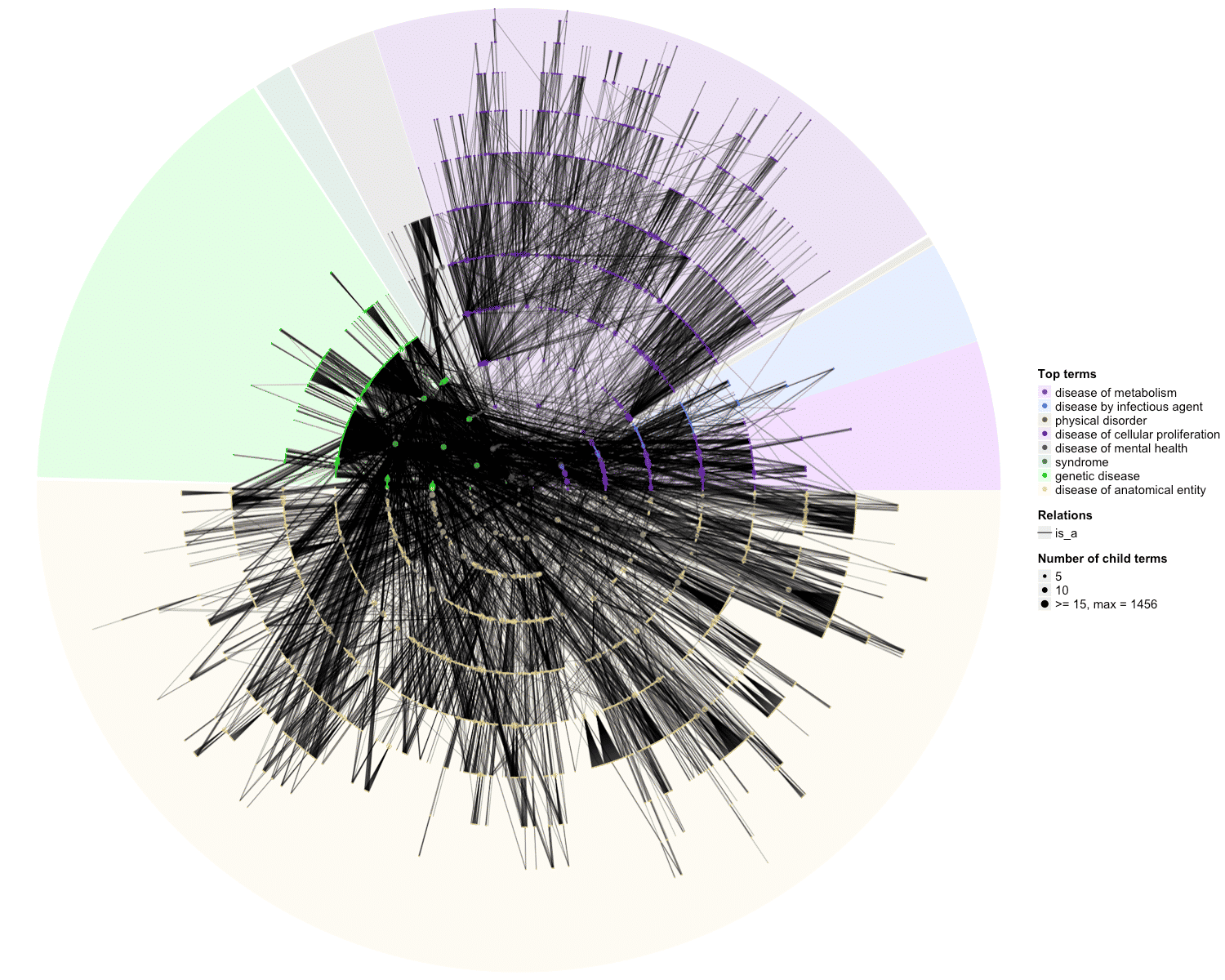

### OBOFoundry_ecao_heatmap.png

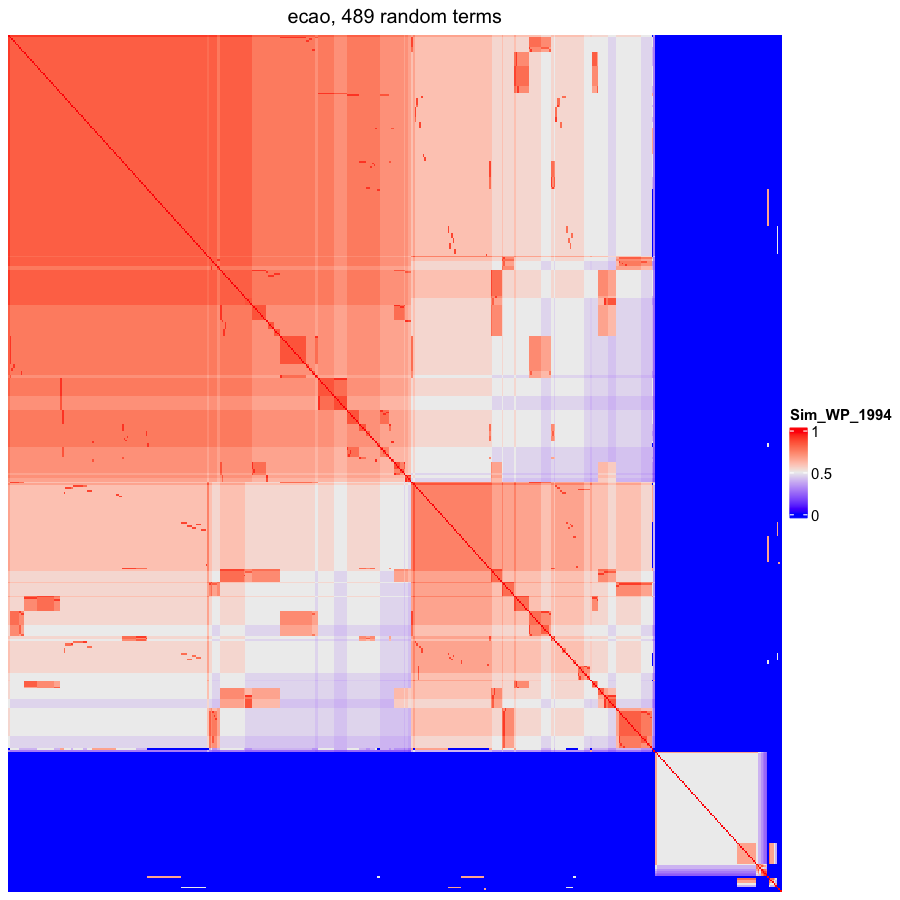

### OBOFoundry_ehdaa2_runtime.png

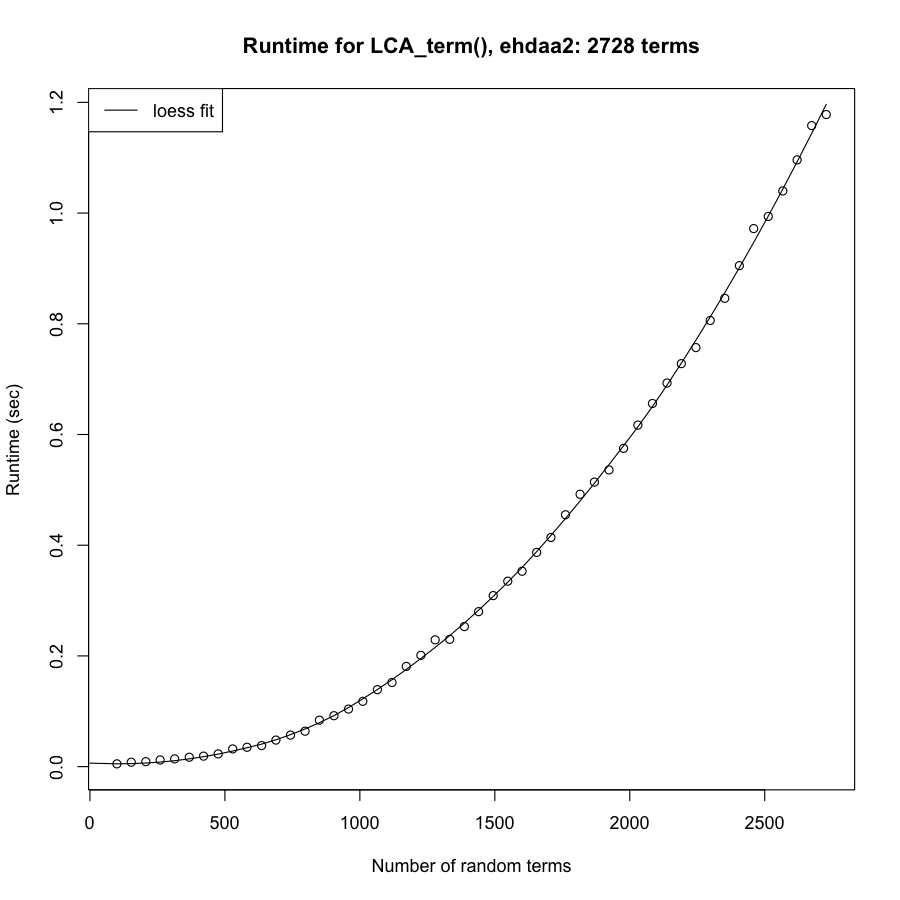

### OBOFoundry_fbbi.png

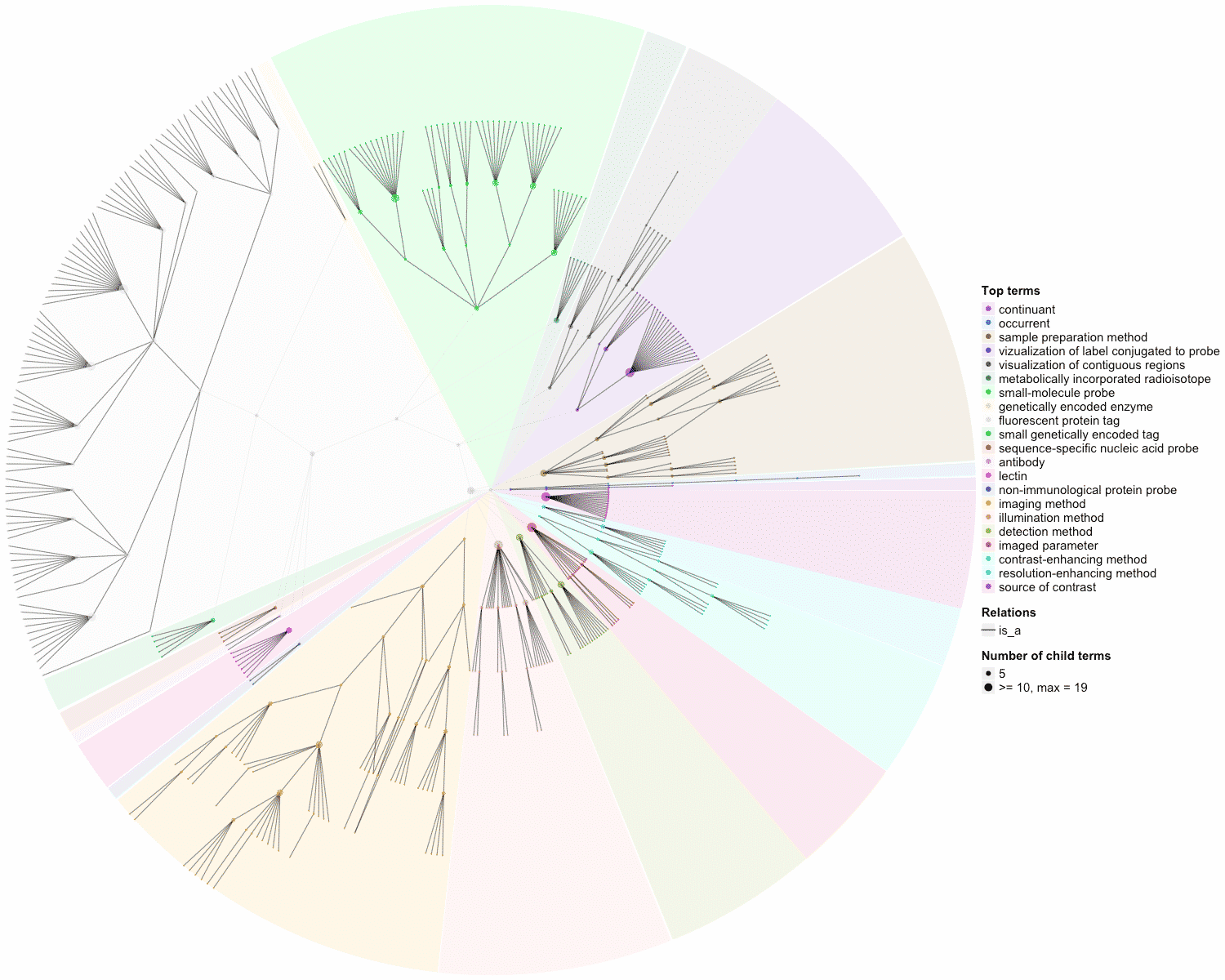

### OBOFoundry_fbbt_runtime.png

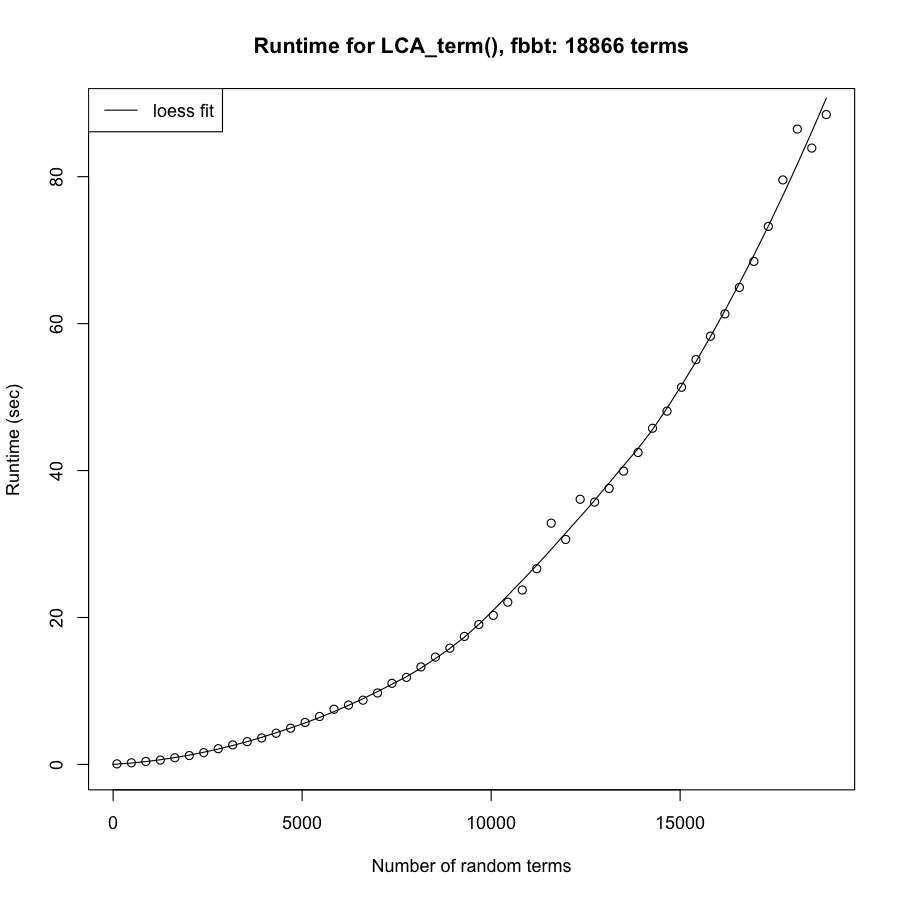

### OBOFoundry_fbcv_heatmap.png

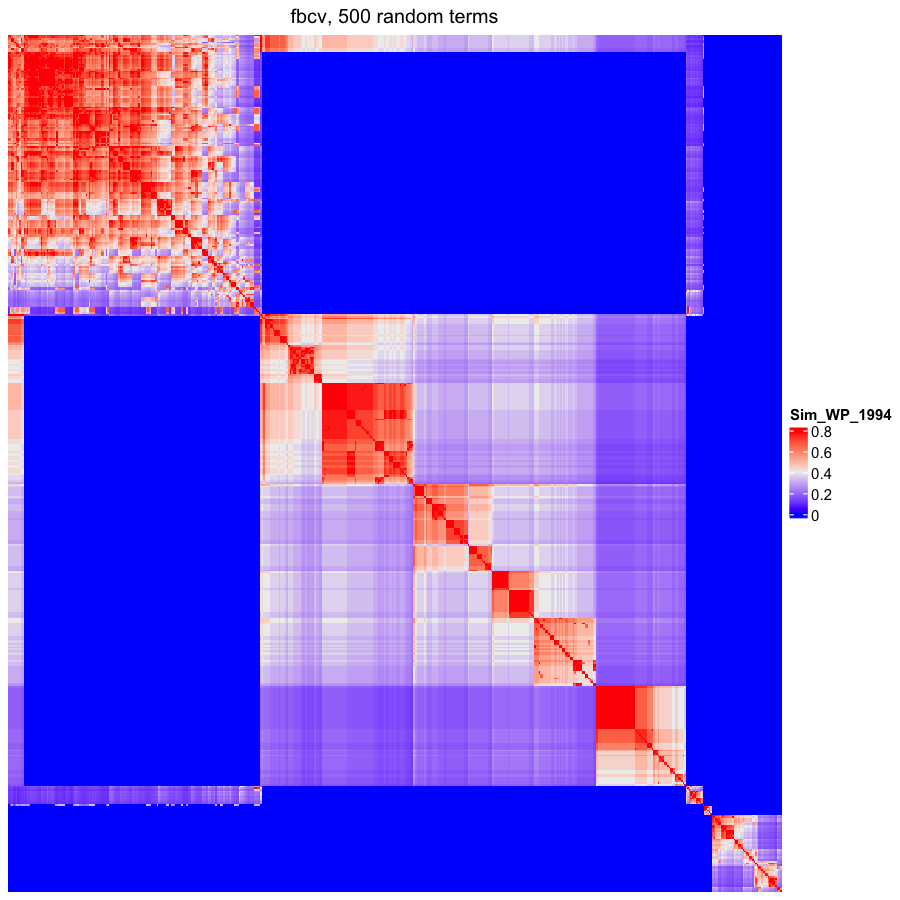

### OBOFoundry_fbdv_heatmap.png

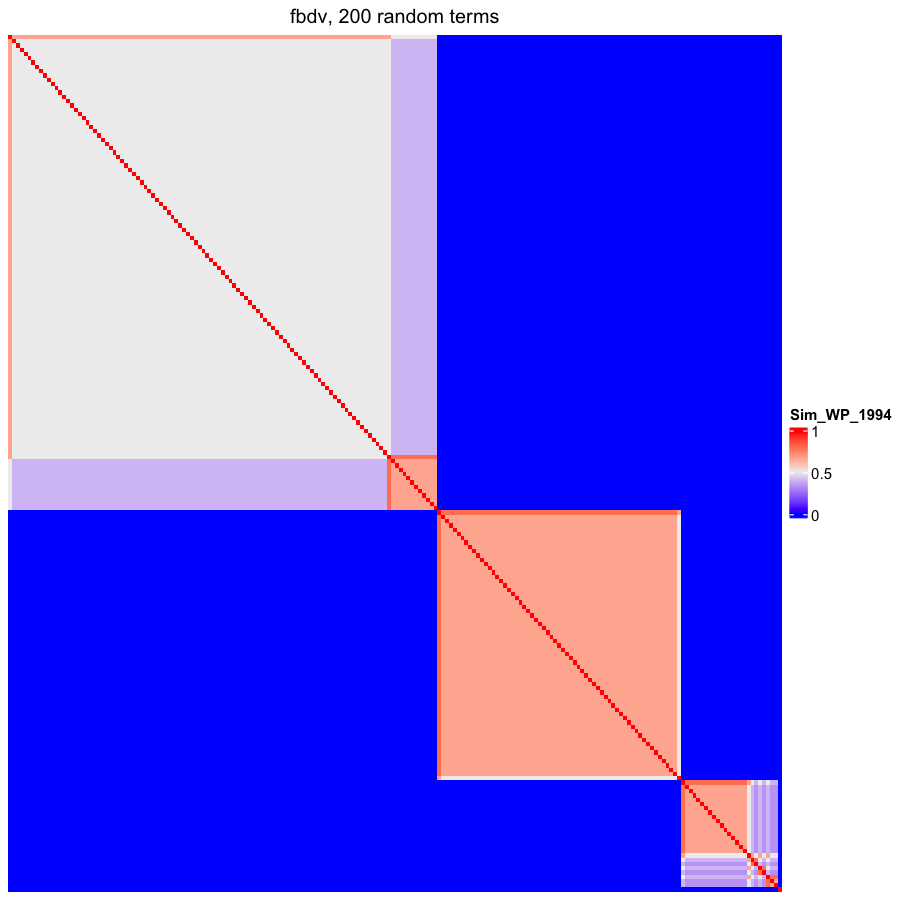

### OBOFoundry_flu.png

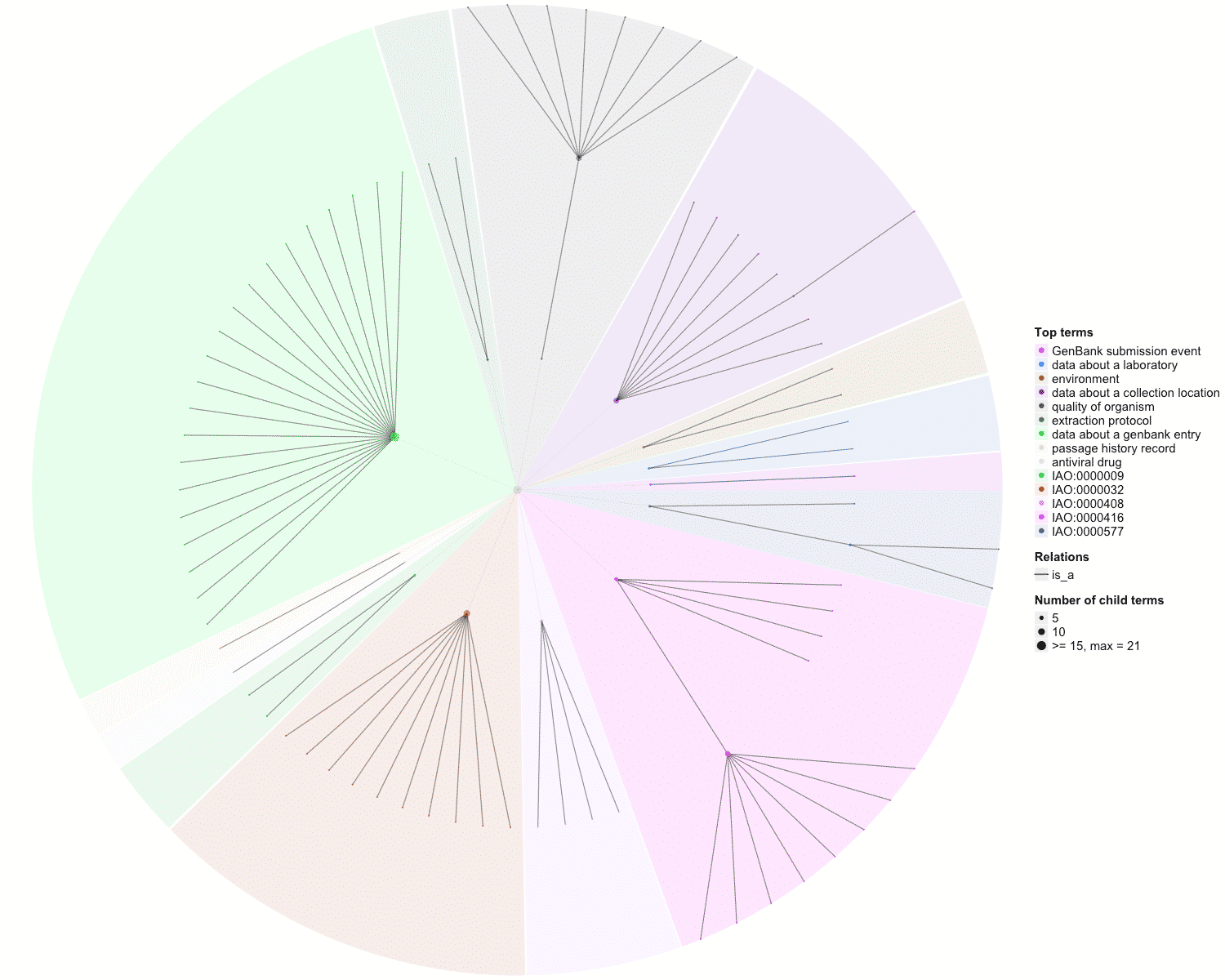

### OBOFoundry_foodon_runtime.png

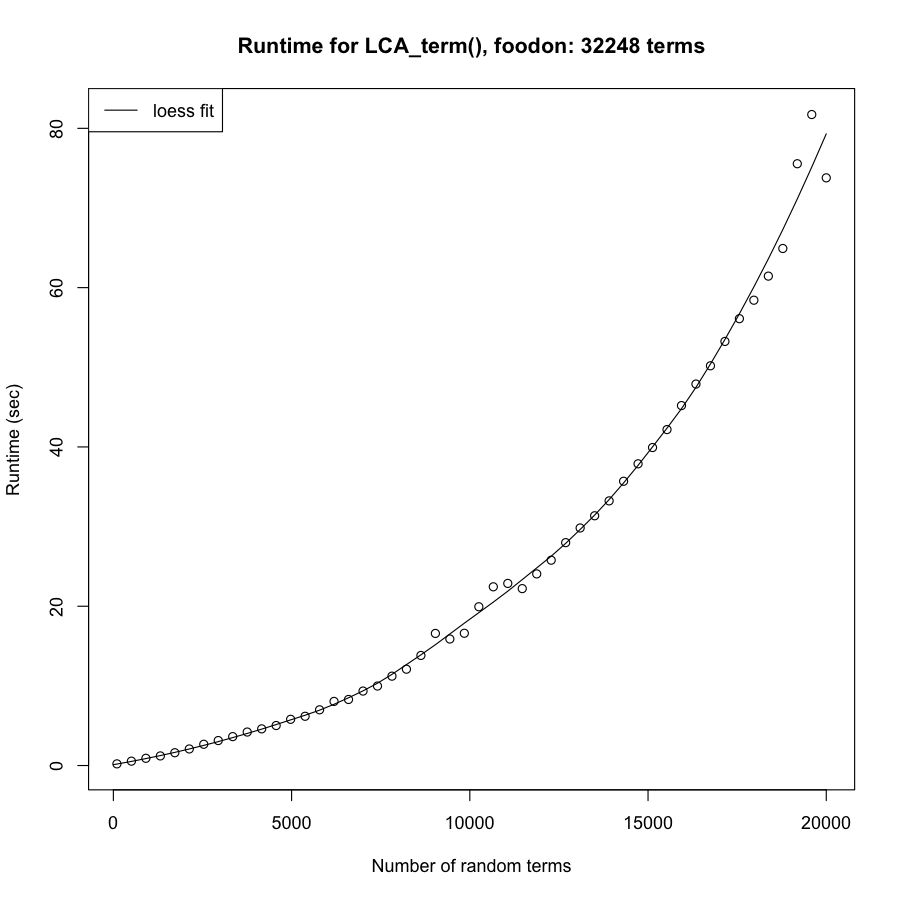

### OBOFoundry_fovt_runtime.png

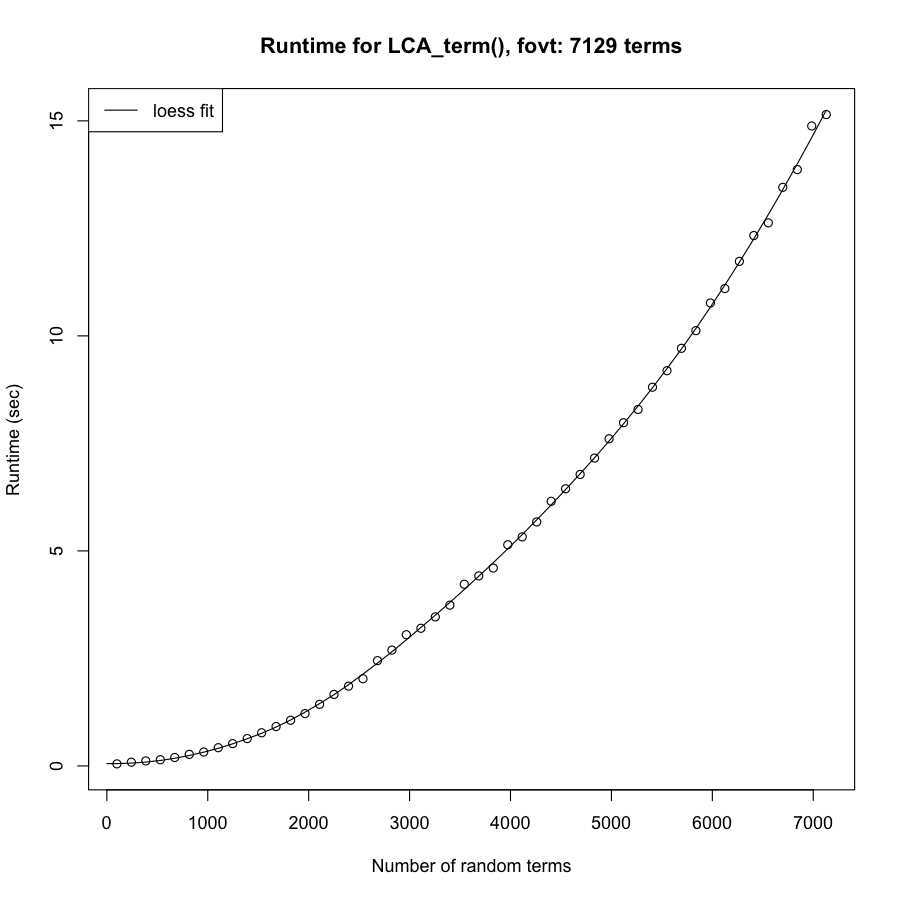

### OBOFoundry_gecko.png

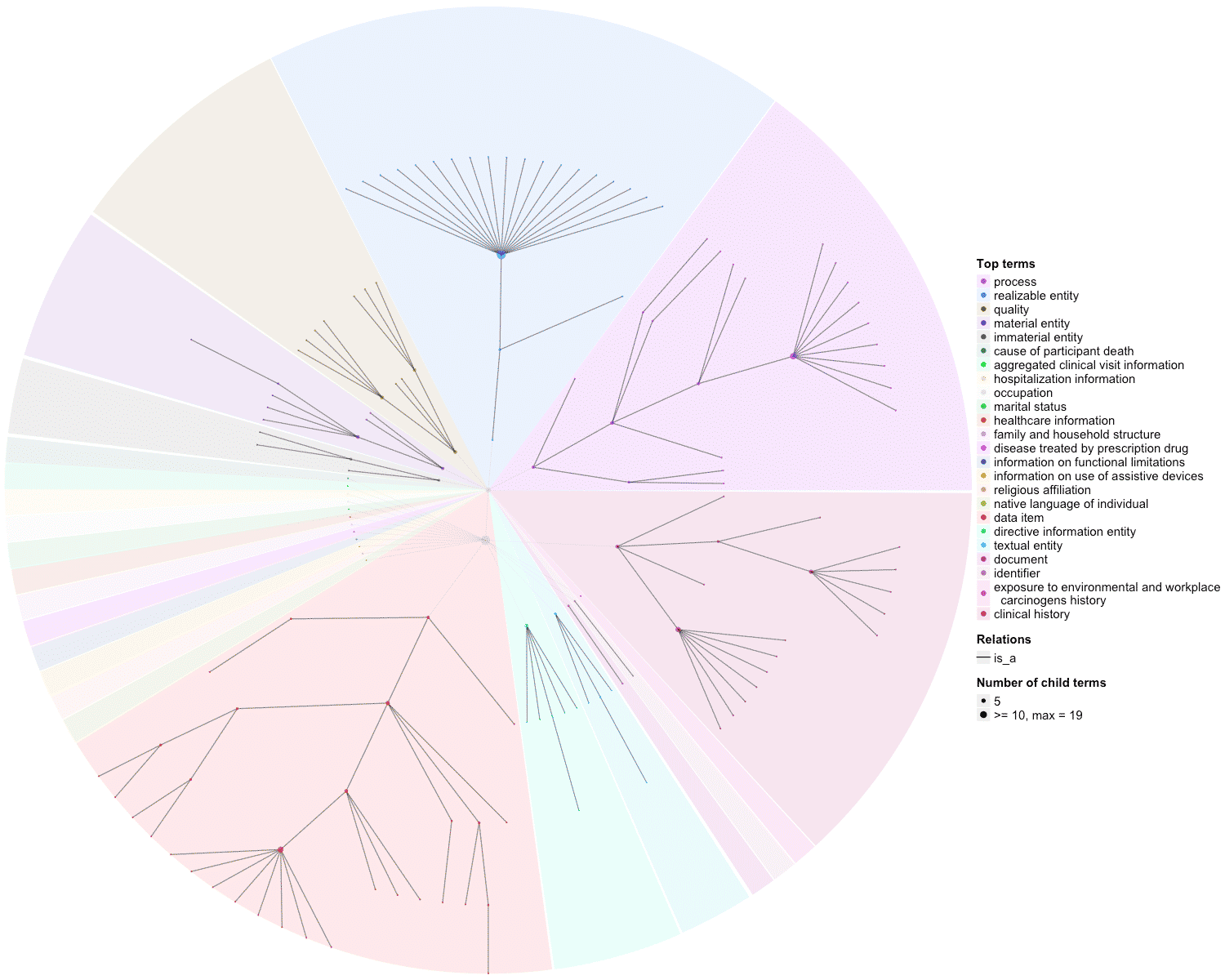

### OBOFoundry_genepio.png

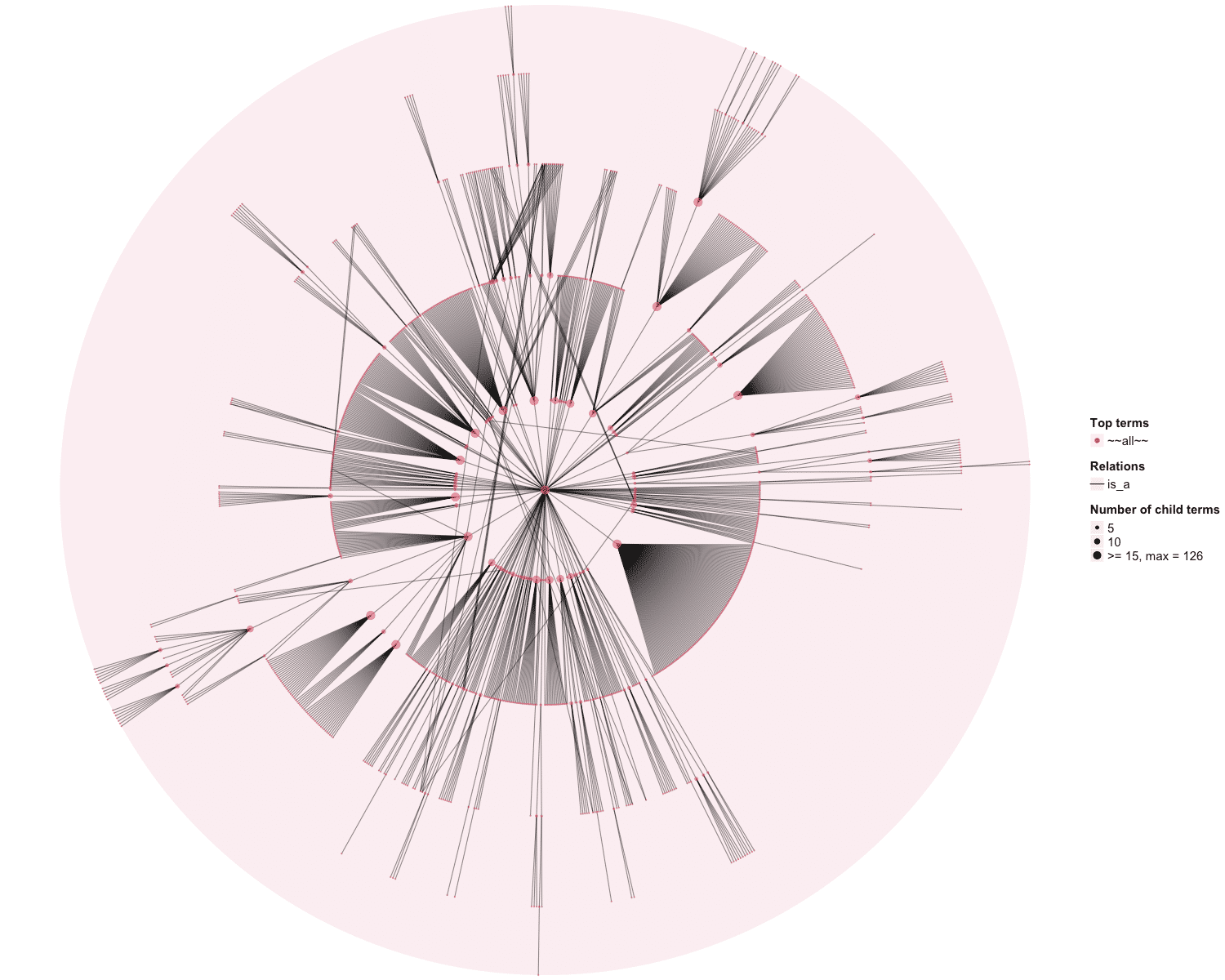

### OBOFoundry_geo.png

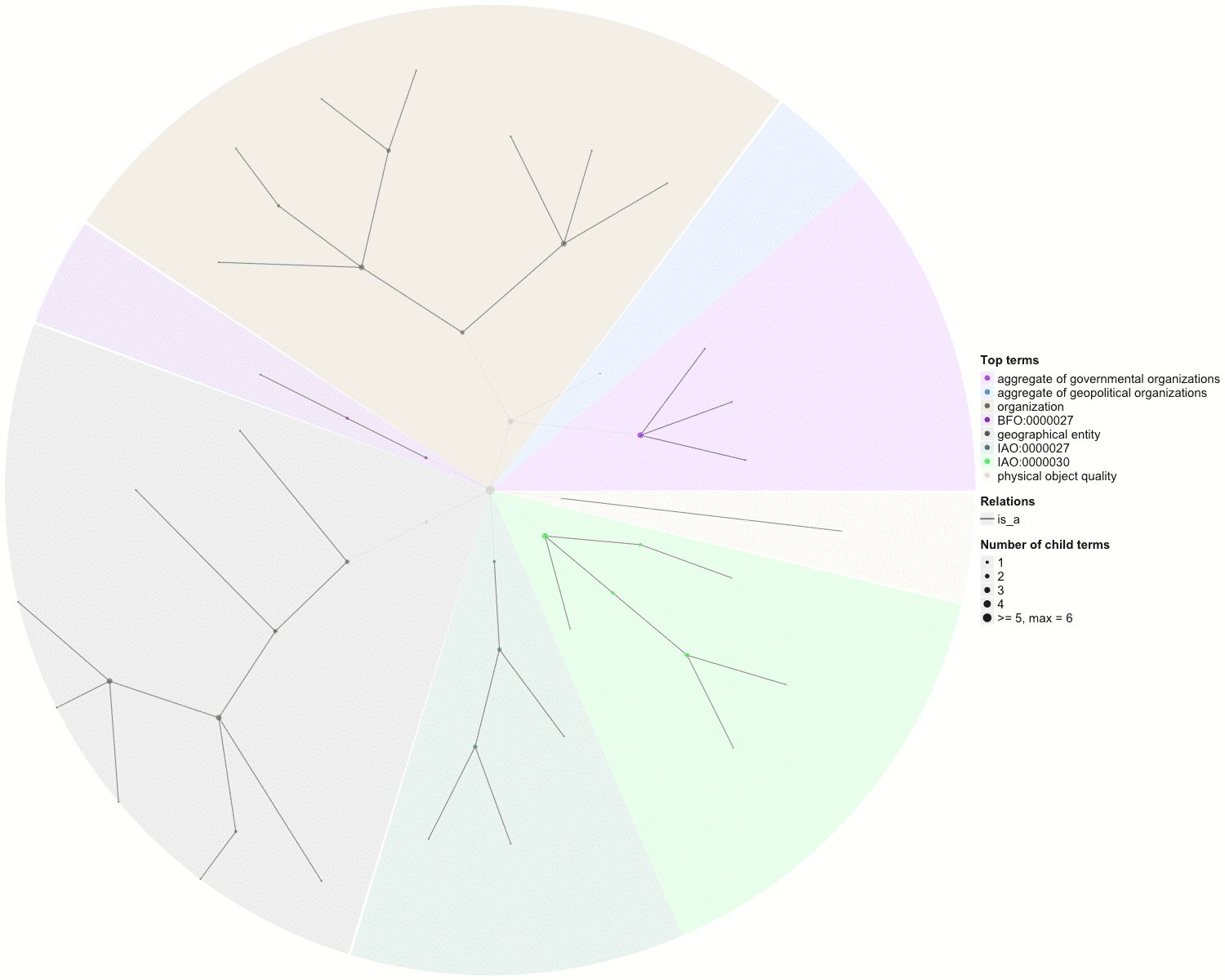

### OBOFoundry_gsso_runtime.png

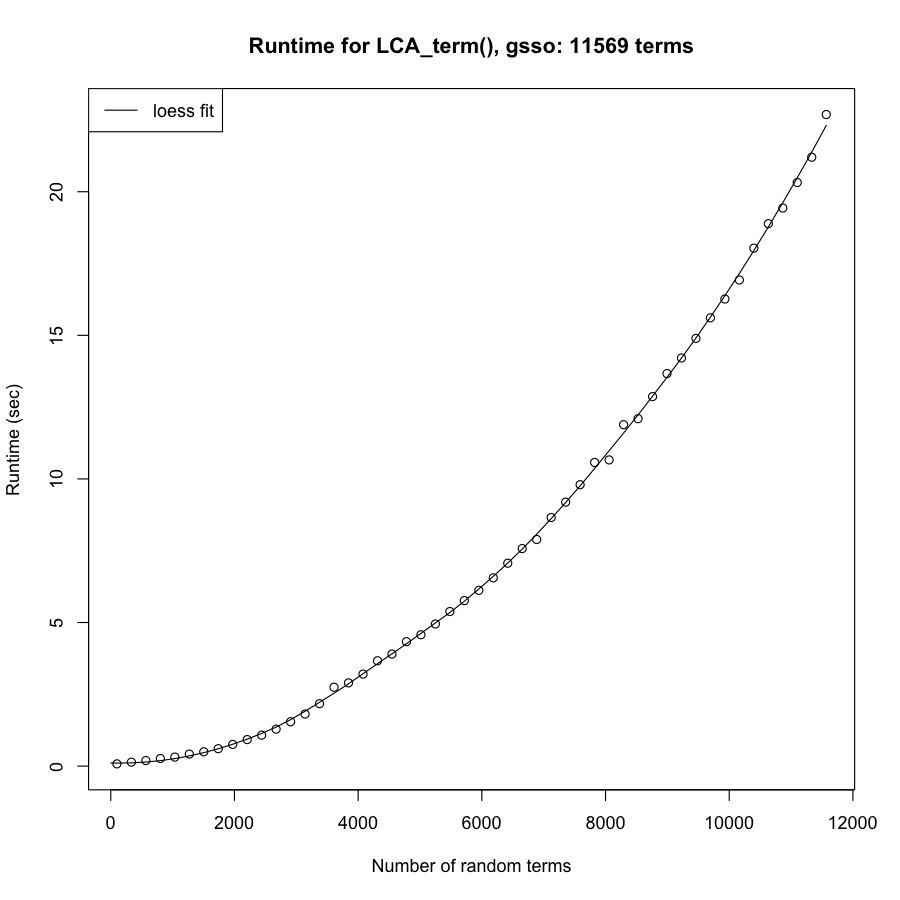

### OBOFoundry_idomal_heatmap.png

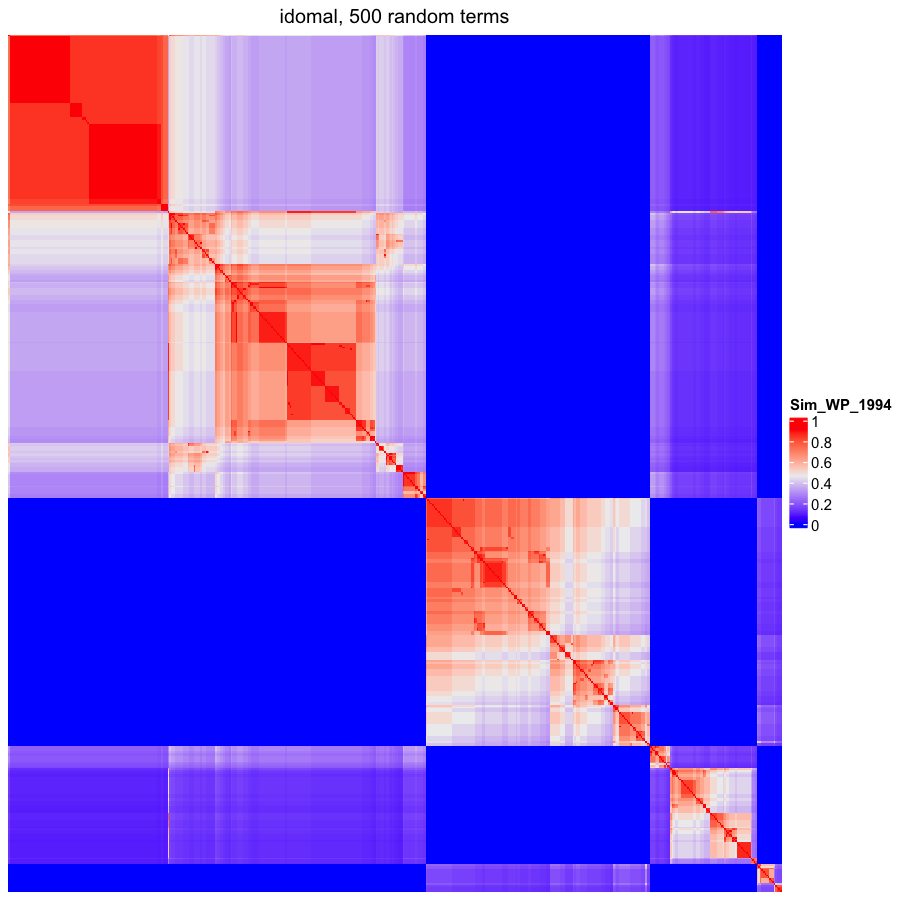

### OBOFoundry_maxo_runtime.png

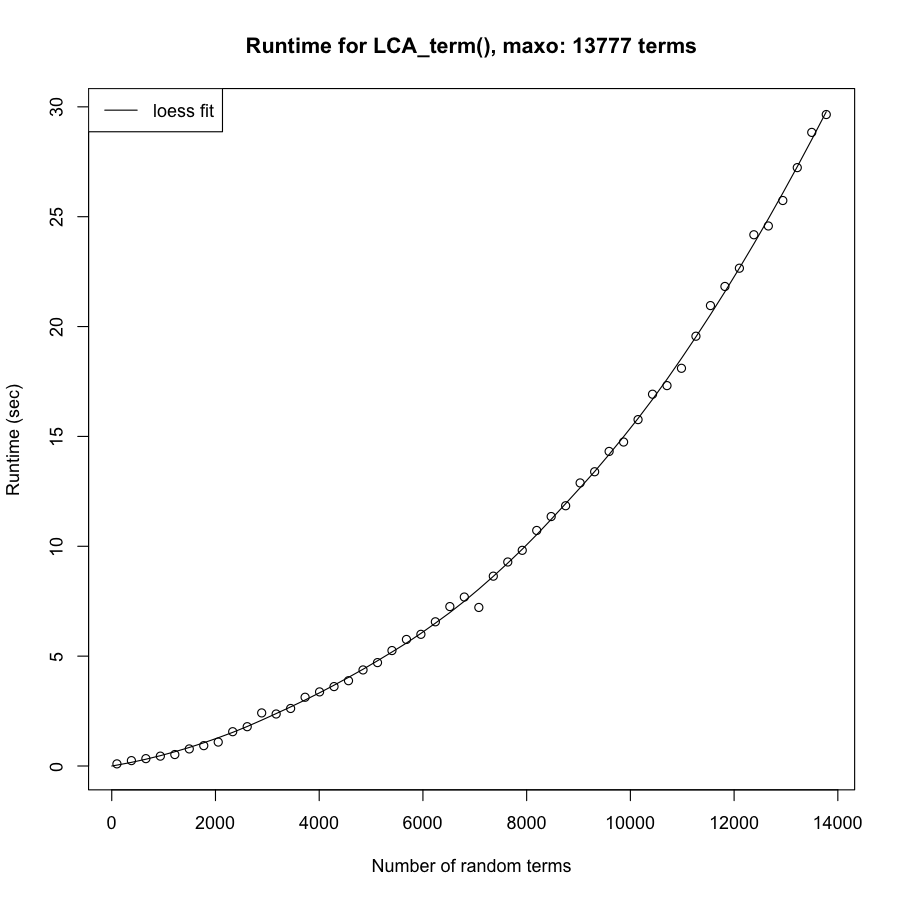

### OBOFoundry_mcro.png

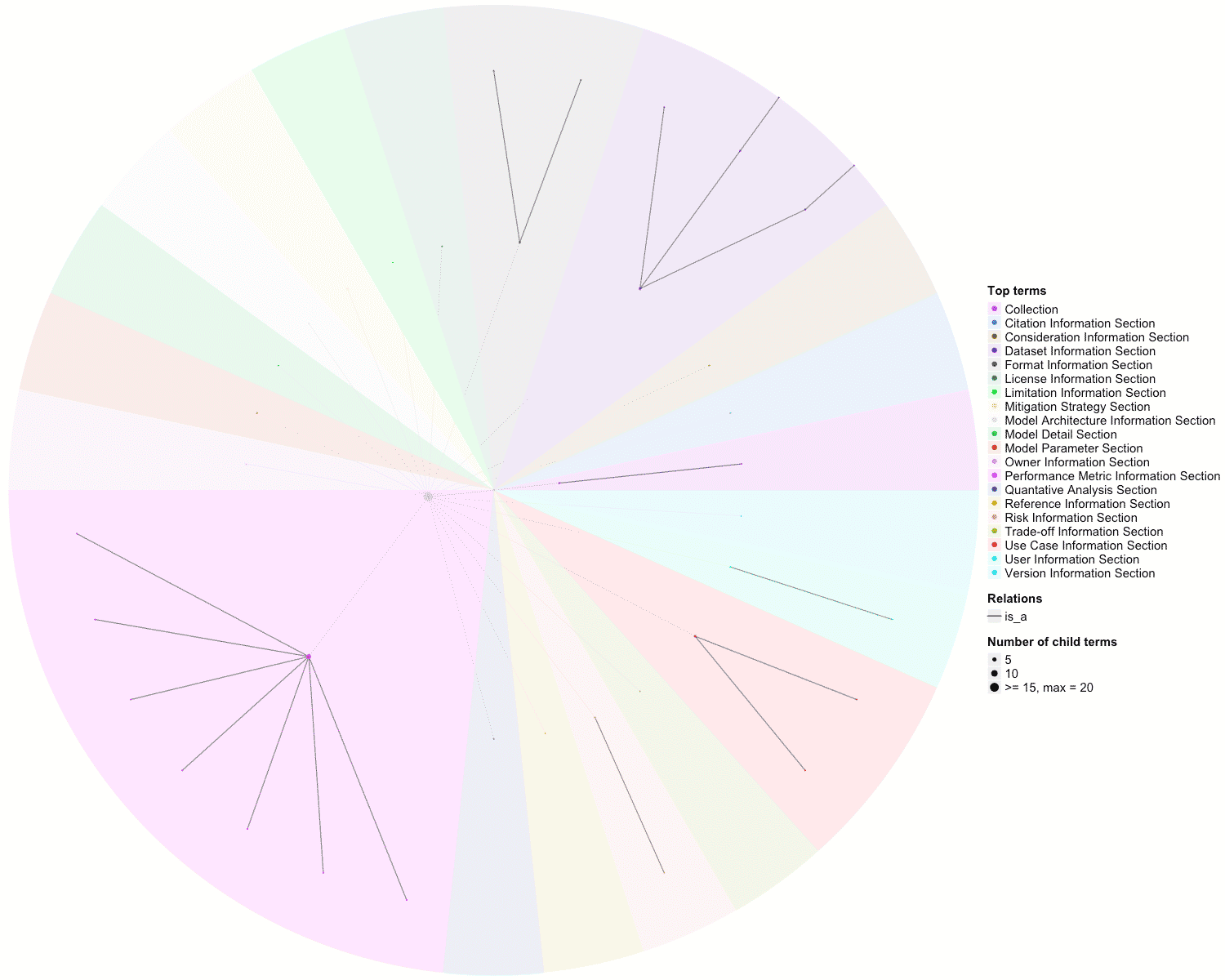

### OBOFoundry_micro_heatmap.png

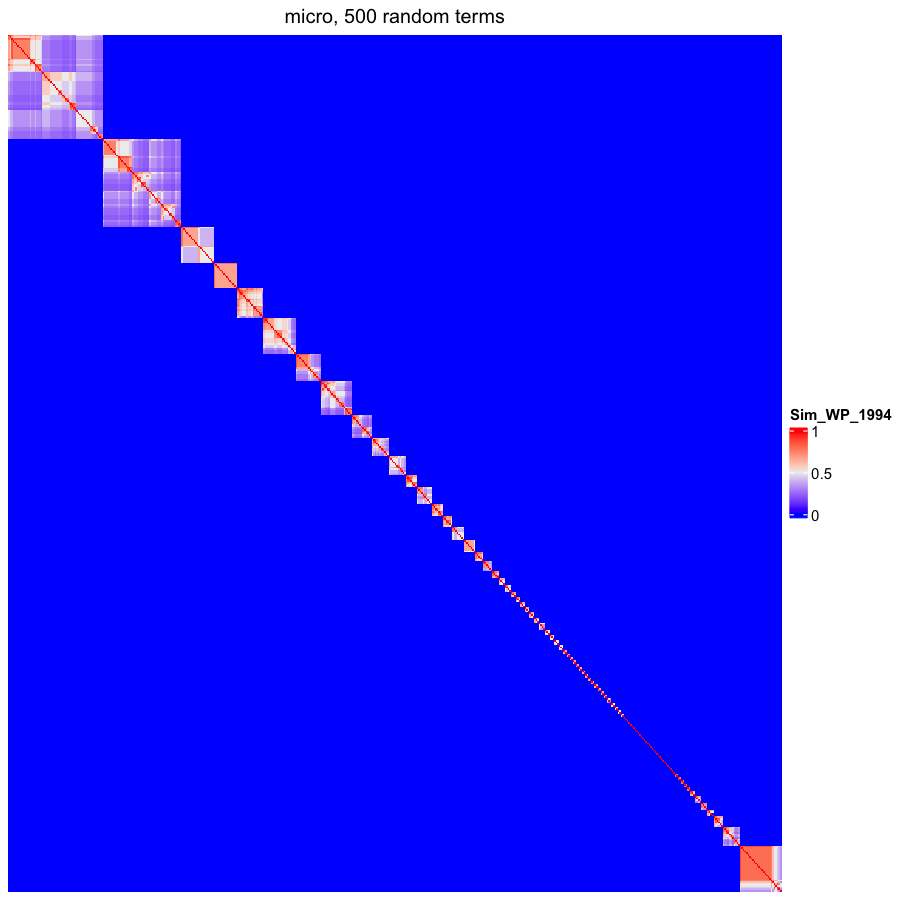

### OBOFoundry_mod_heatmap.png

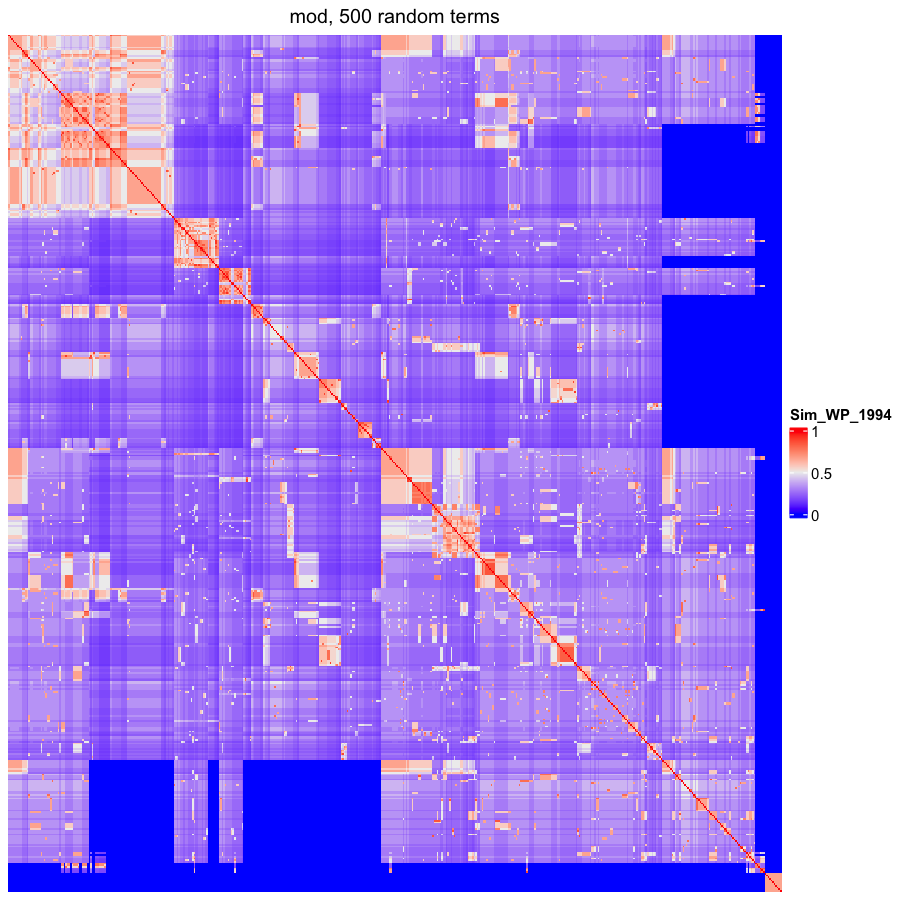

### OBOFoundry_mro.png

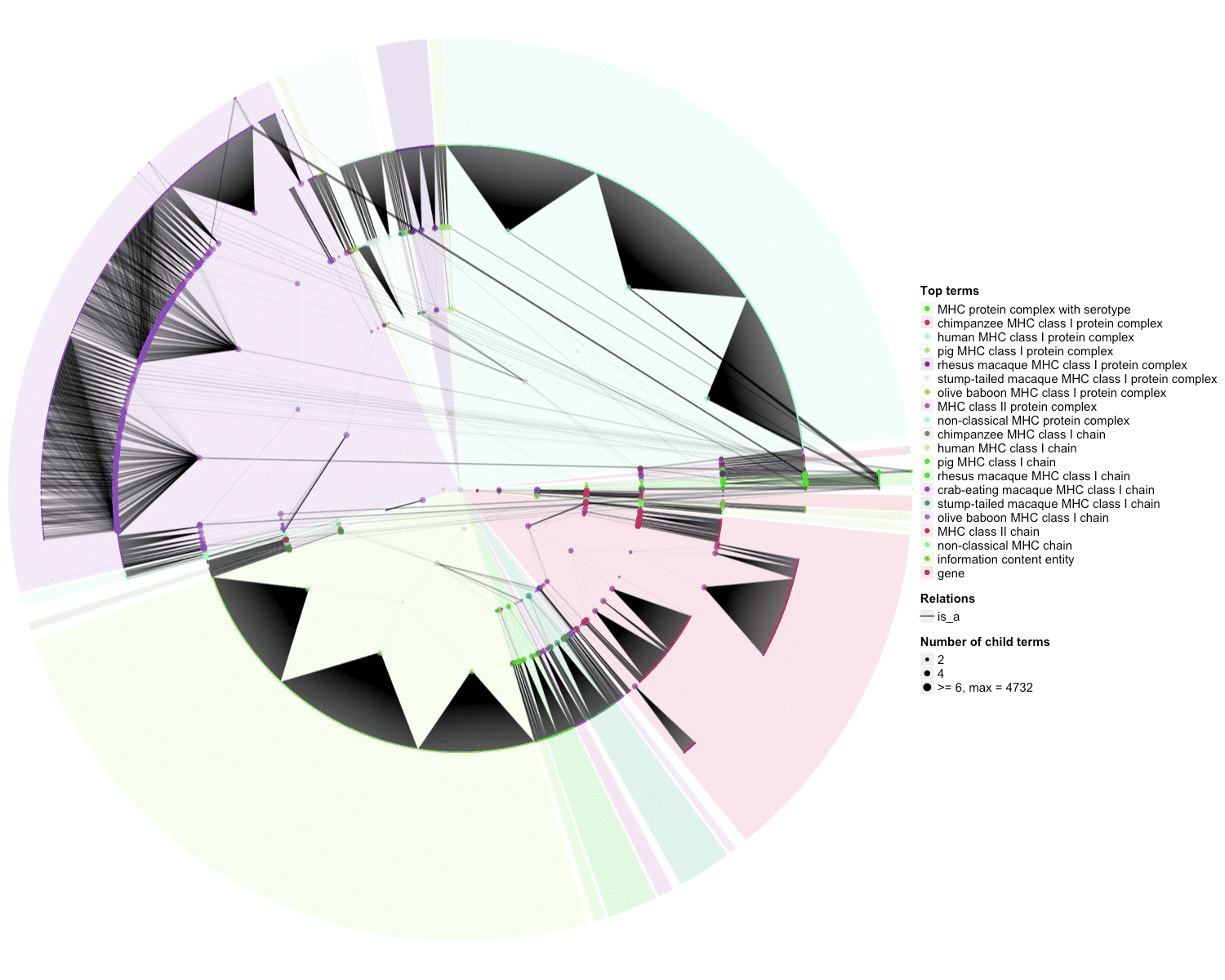

### OBOFoundry_ncro_heatmap.png

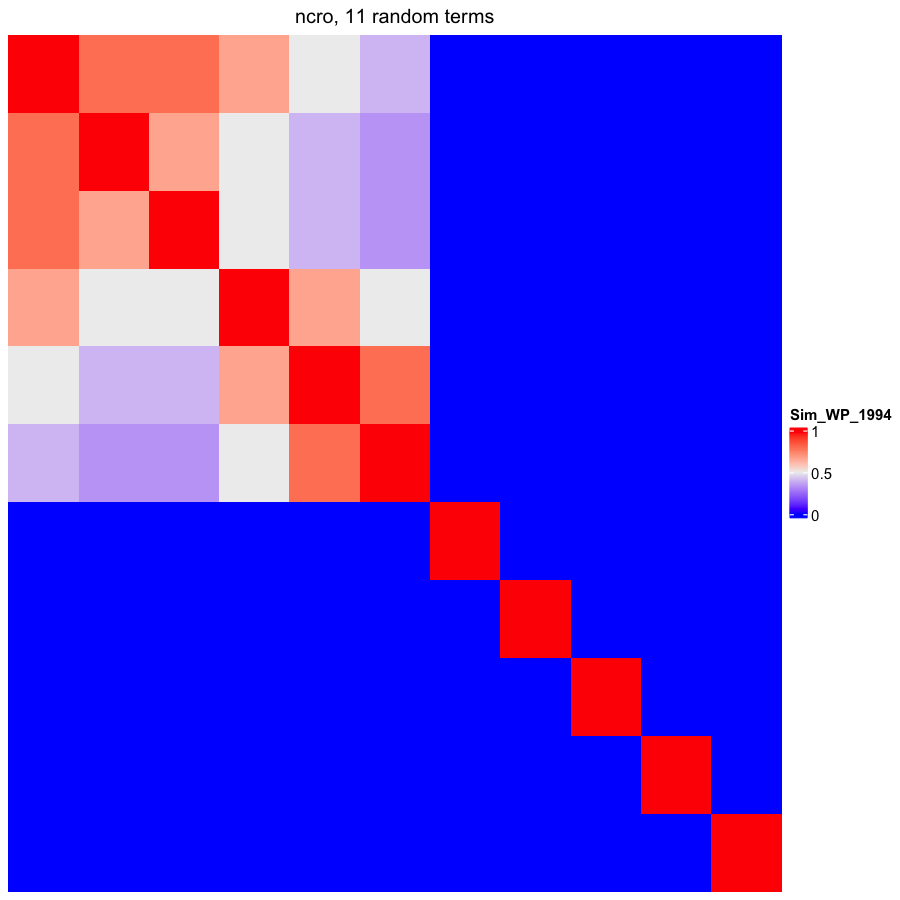

### OBOFoundry_oarcs_heatmap.png

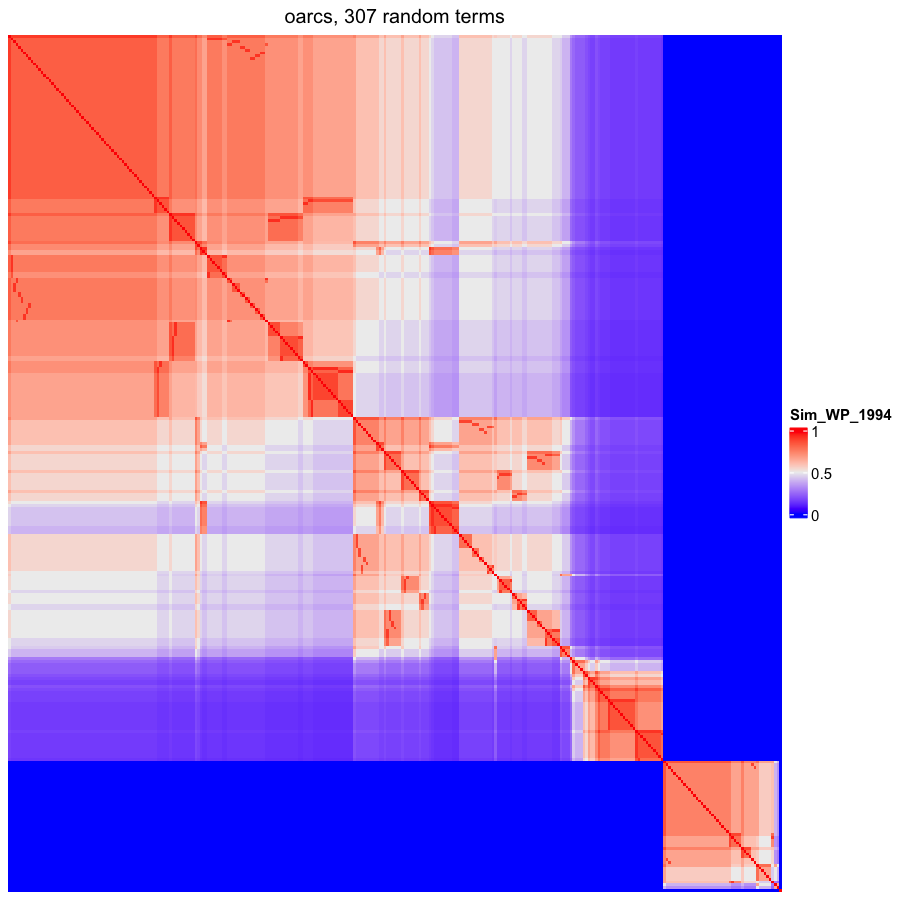

### OBOFoundry_obcs_heatmap.png

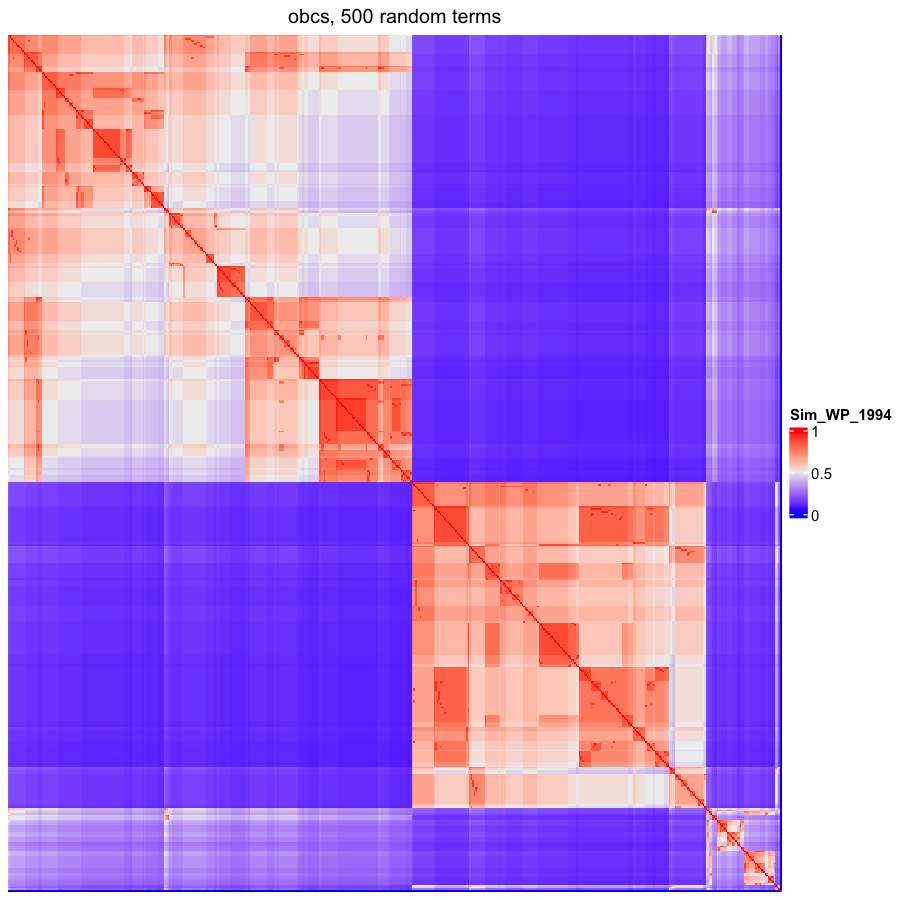

### OBOFoundry_ogsf_heatmap.png

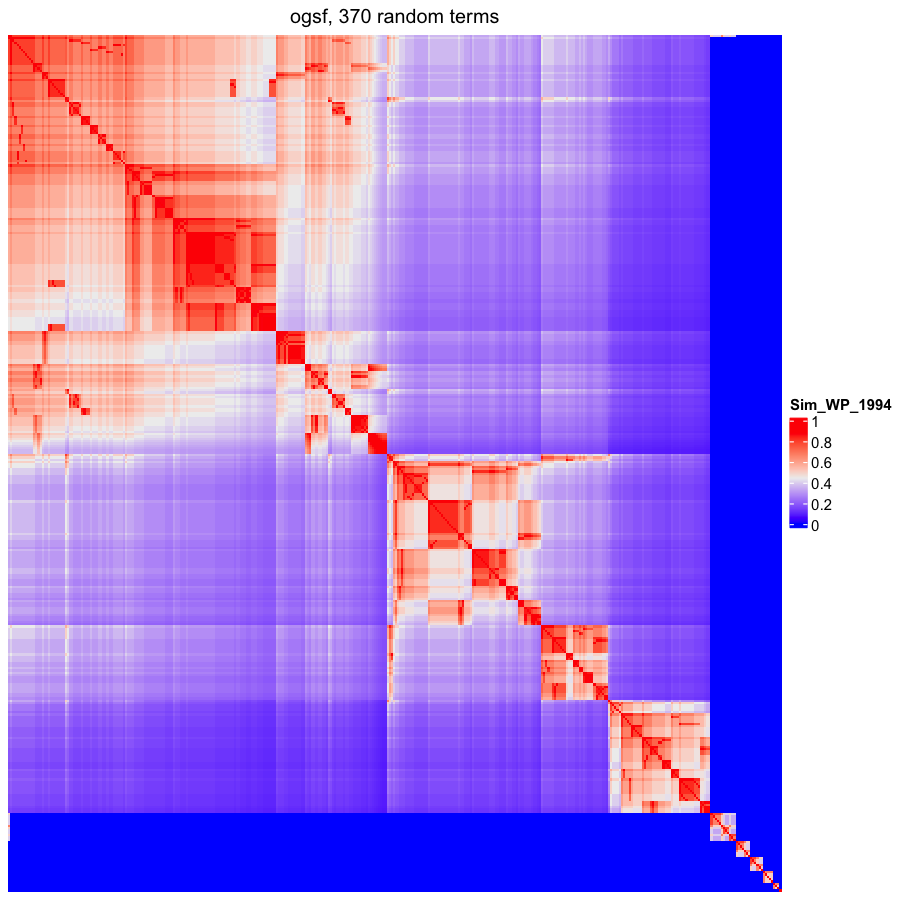

### OBOFoundry_ohpi_heatmap.png

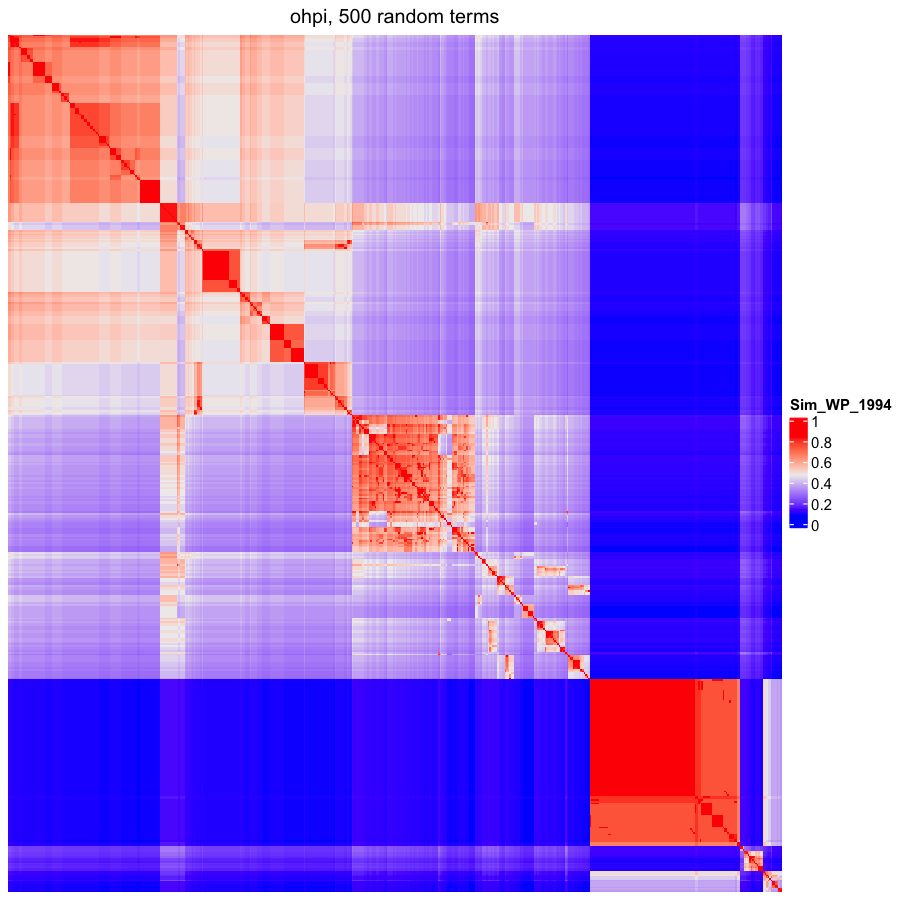

### OBOFoundry_omiabis_heatmap.png

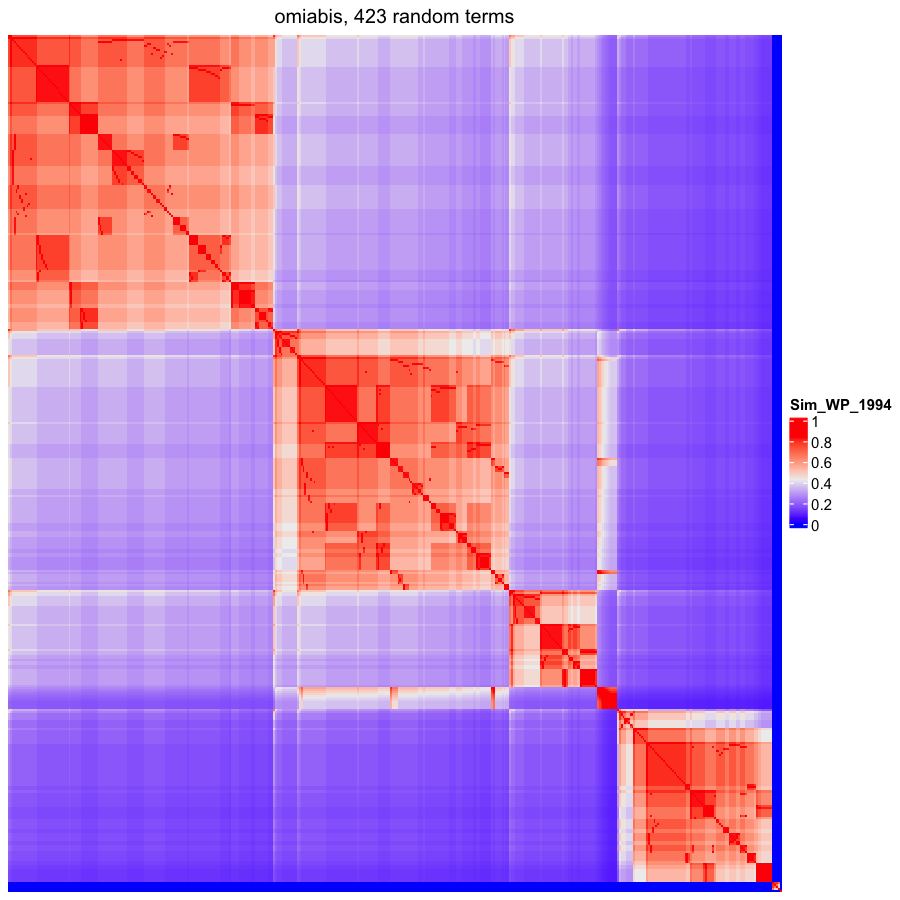

### OBOFoundry_omp_runtime.png

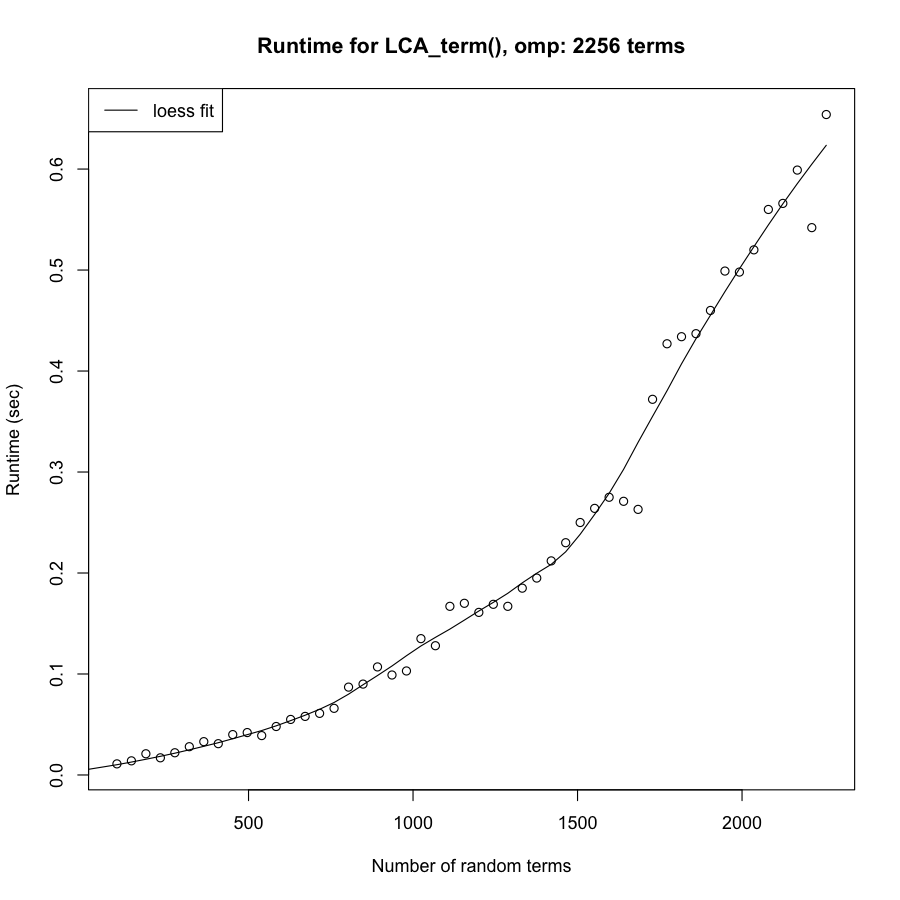

### OBOFoundry_ontoavida_heatmap.png

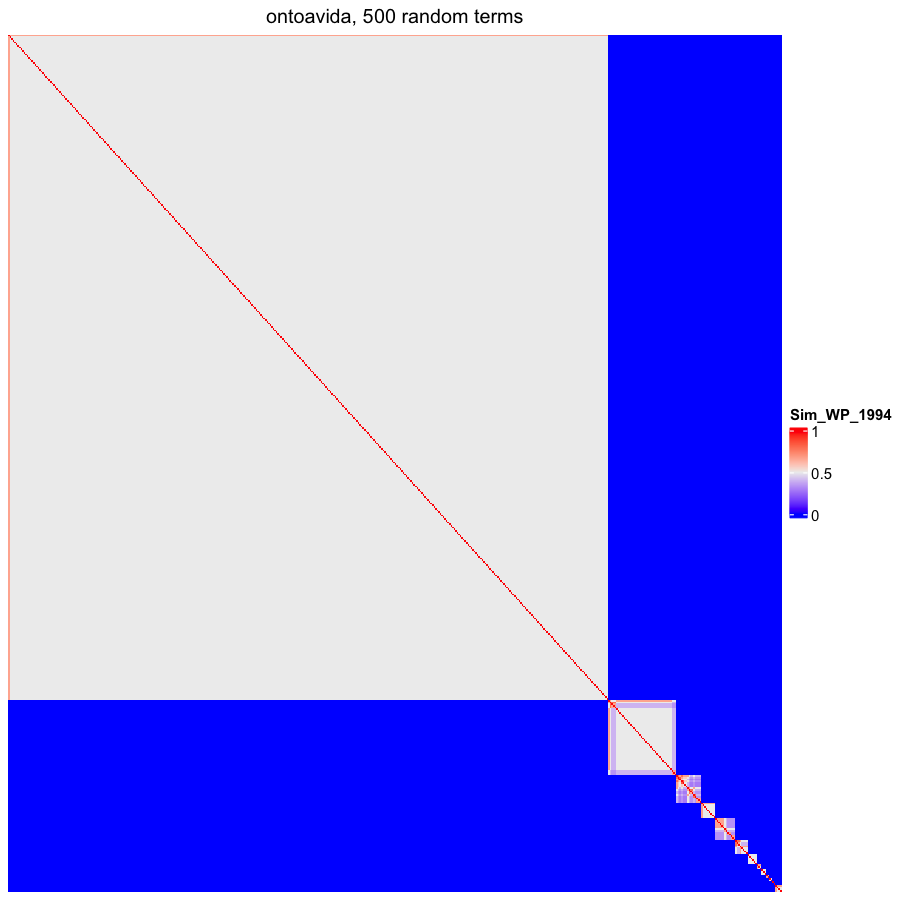

### OBOFoundry_ovae_heatmap.png

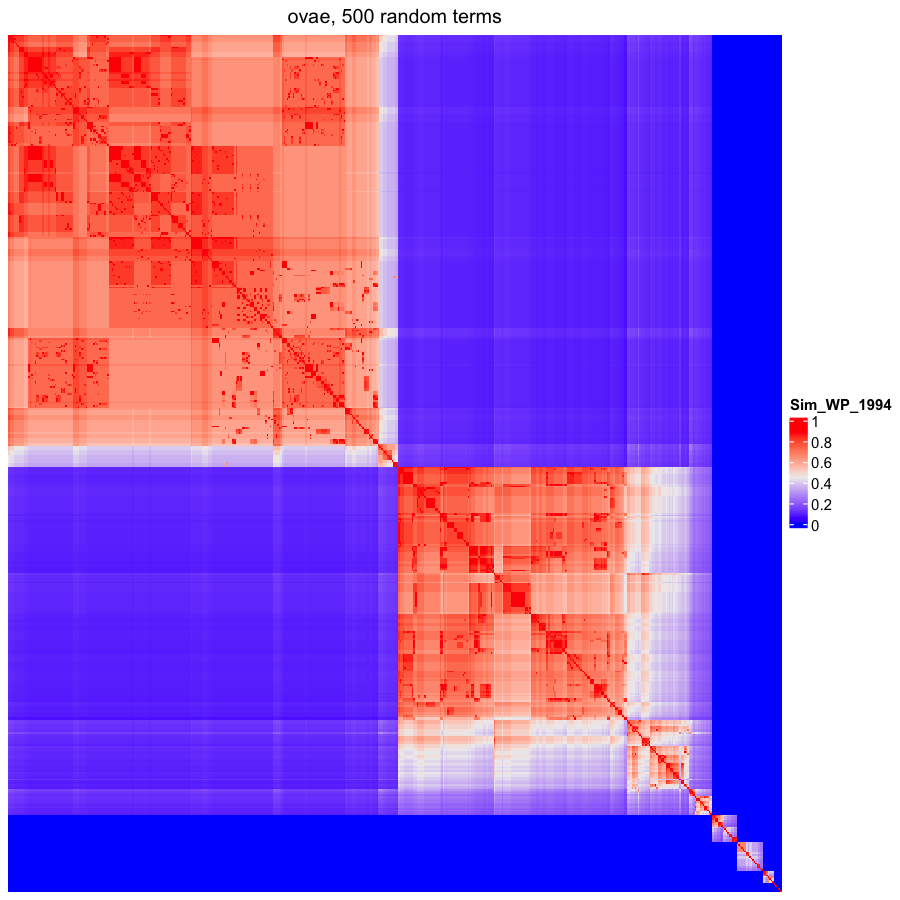

### OBOFoundry_poro.png

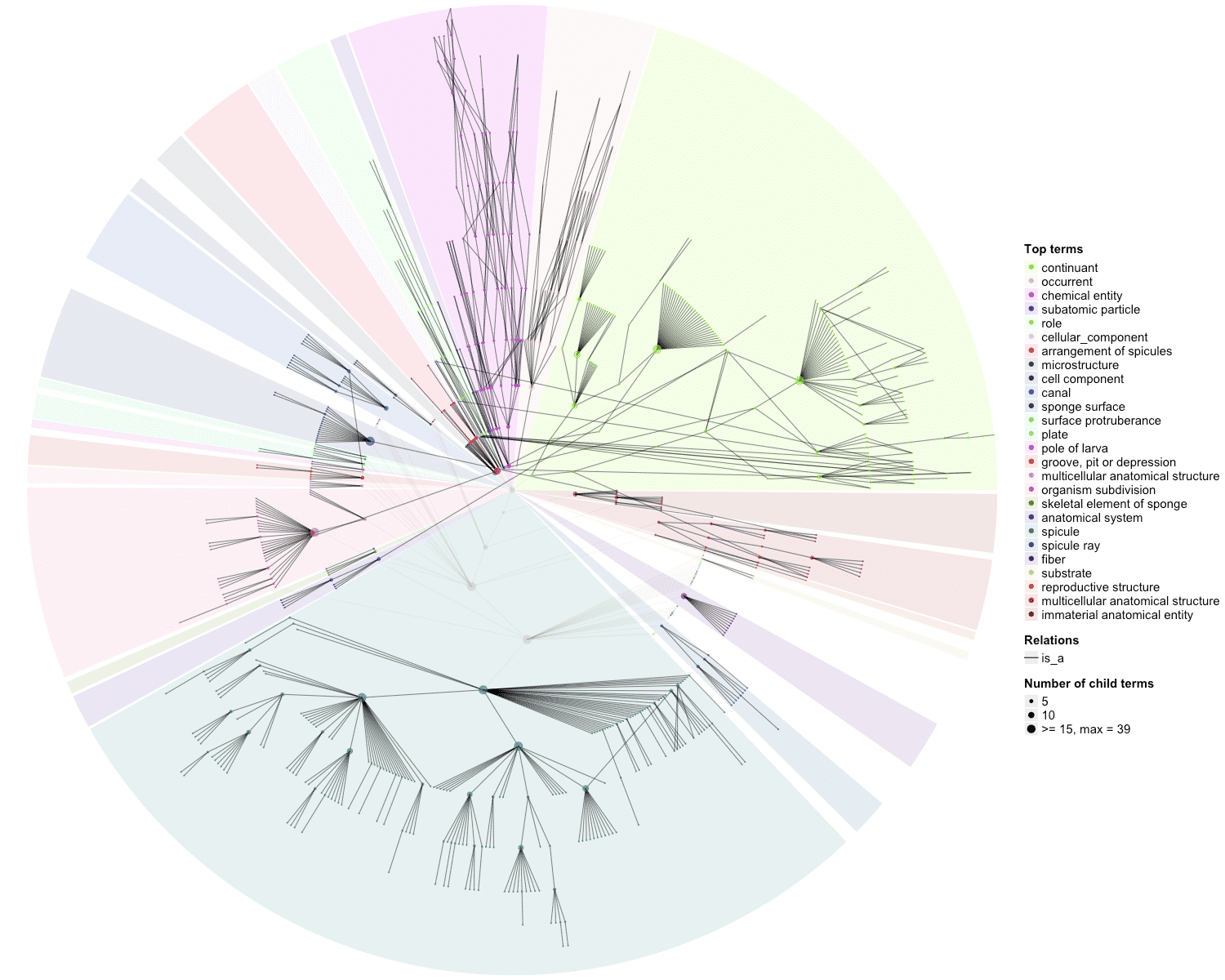

### OBOFoundry_pso.png

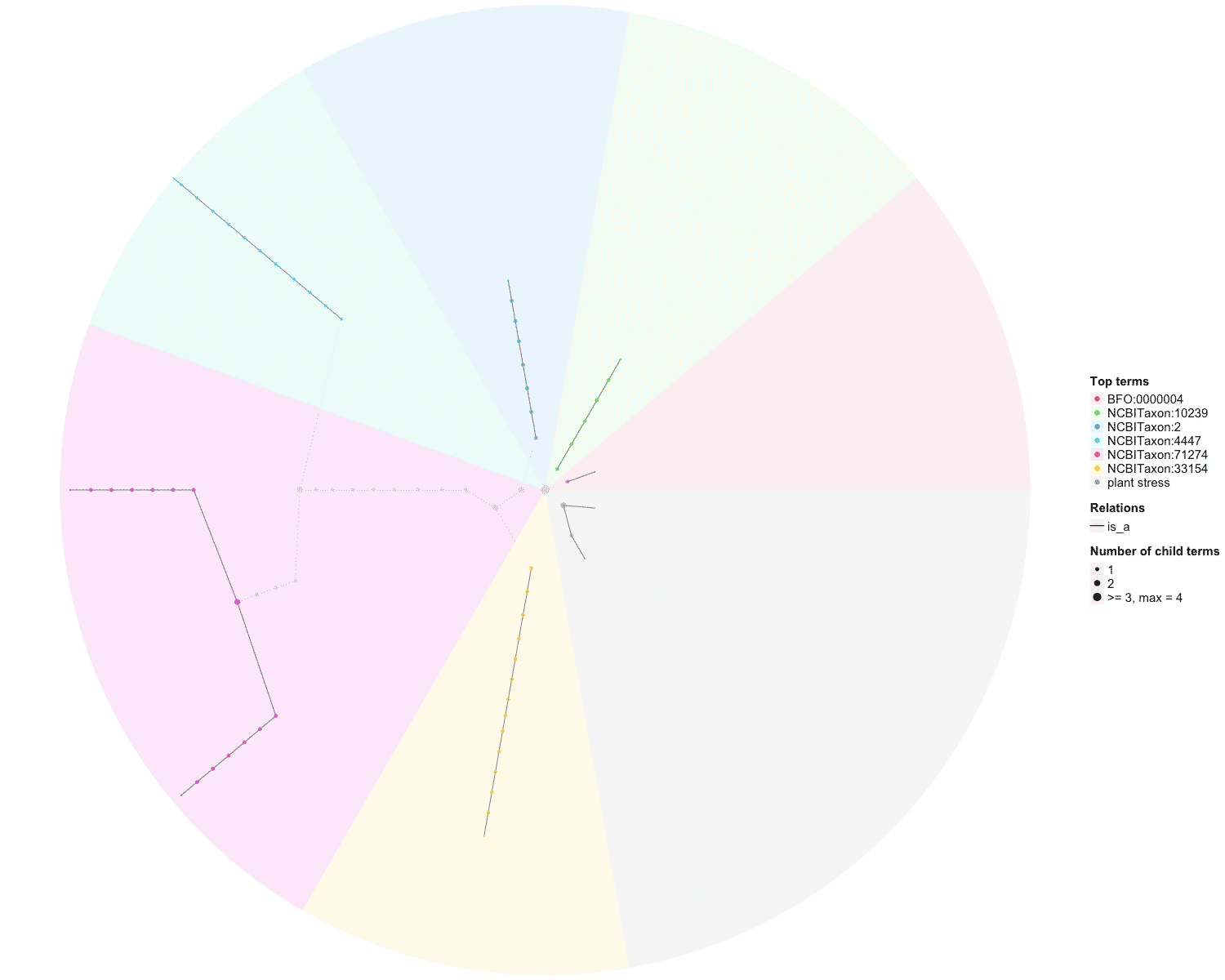

### OBOFoundry_rxno.png

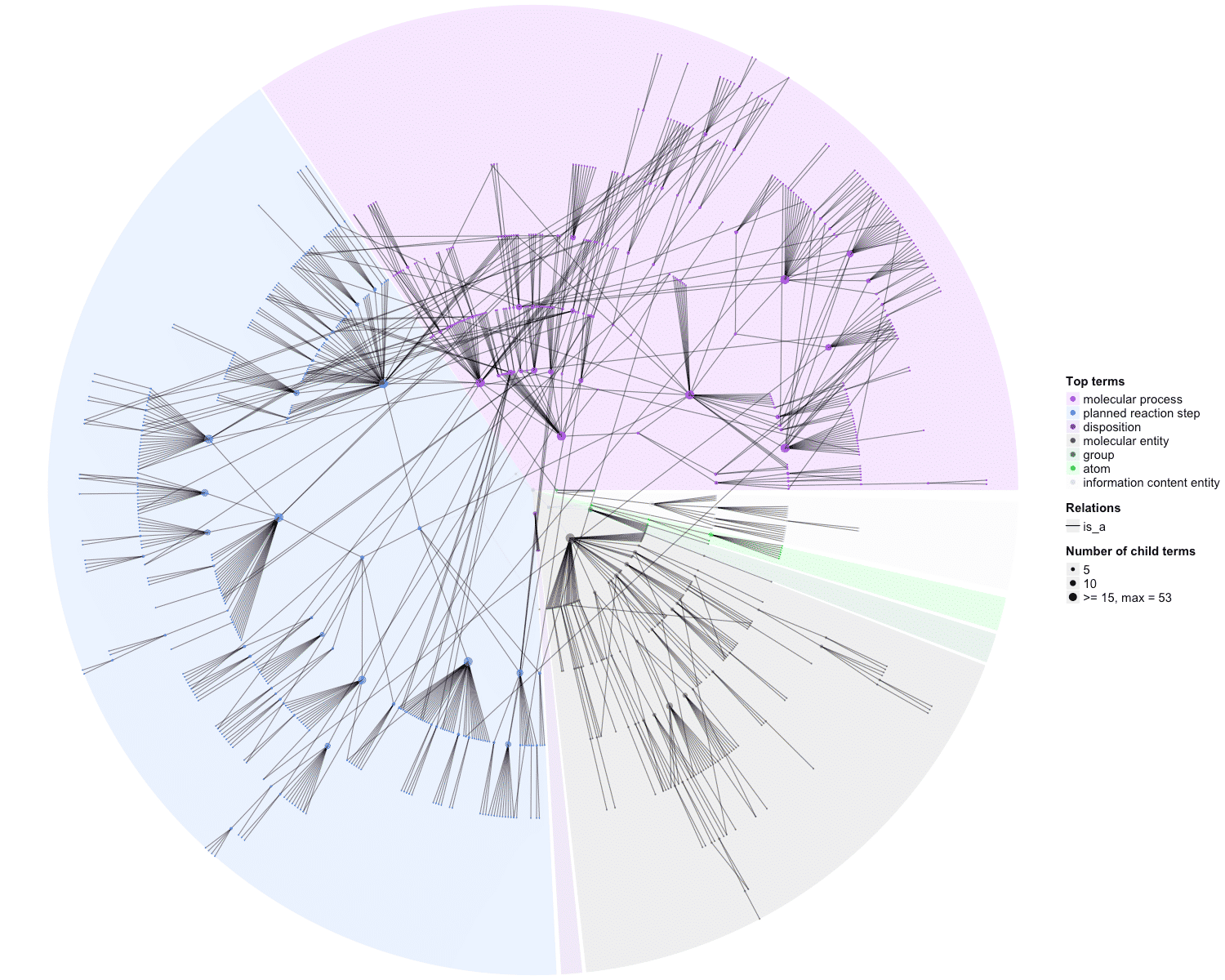

### OBOFoundry_tgma.png

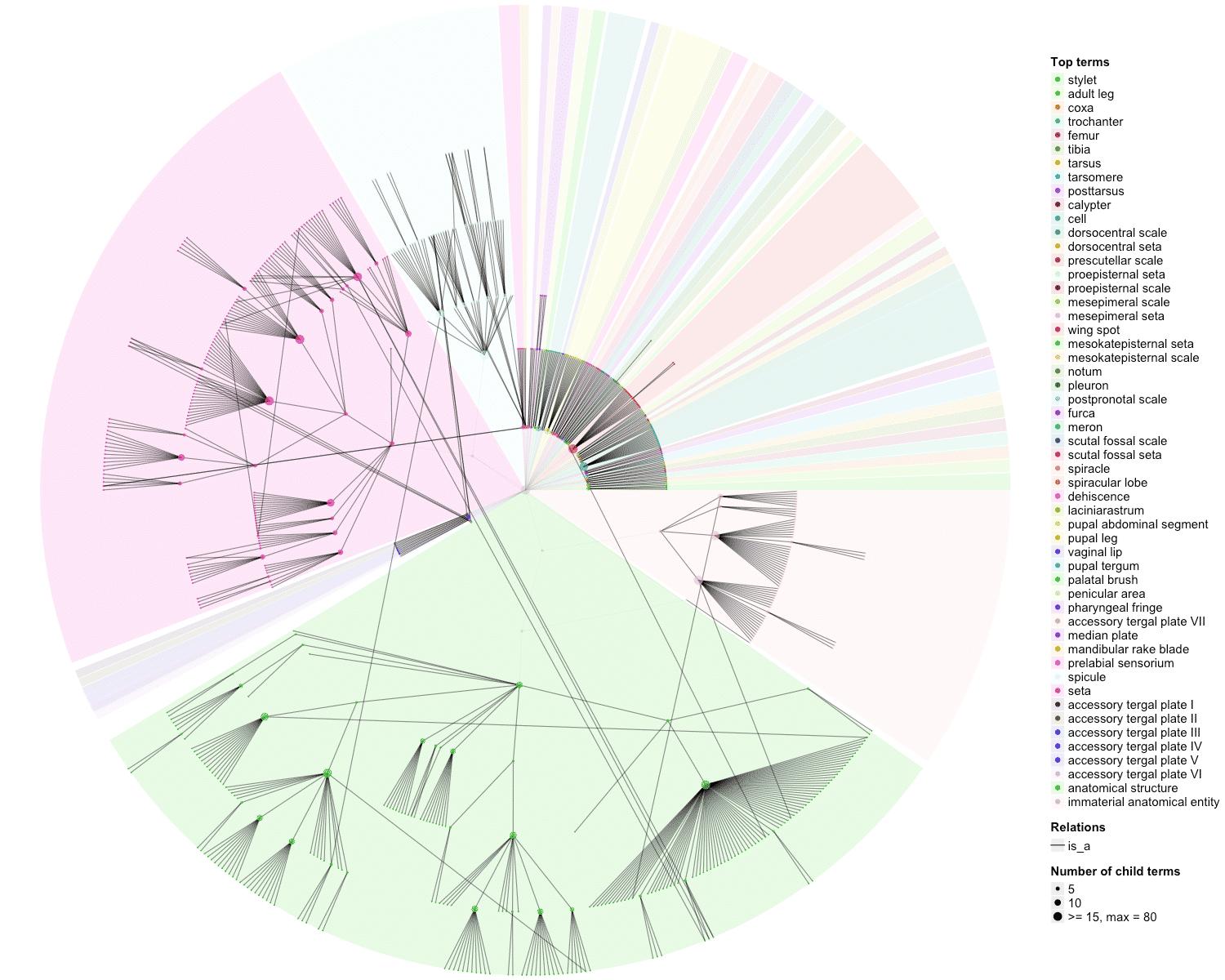

### OBOFoundry_uberon_runtime.png

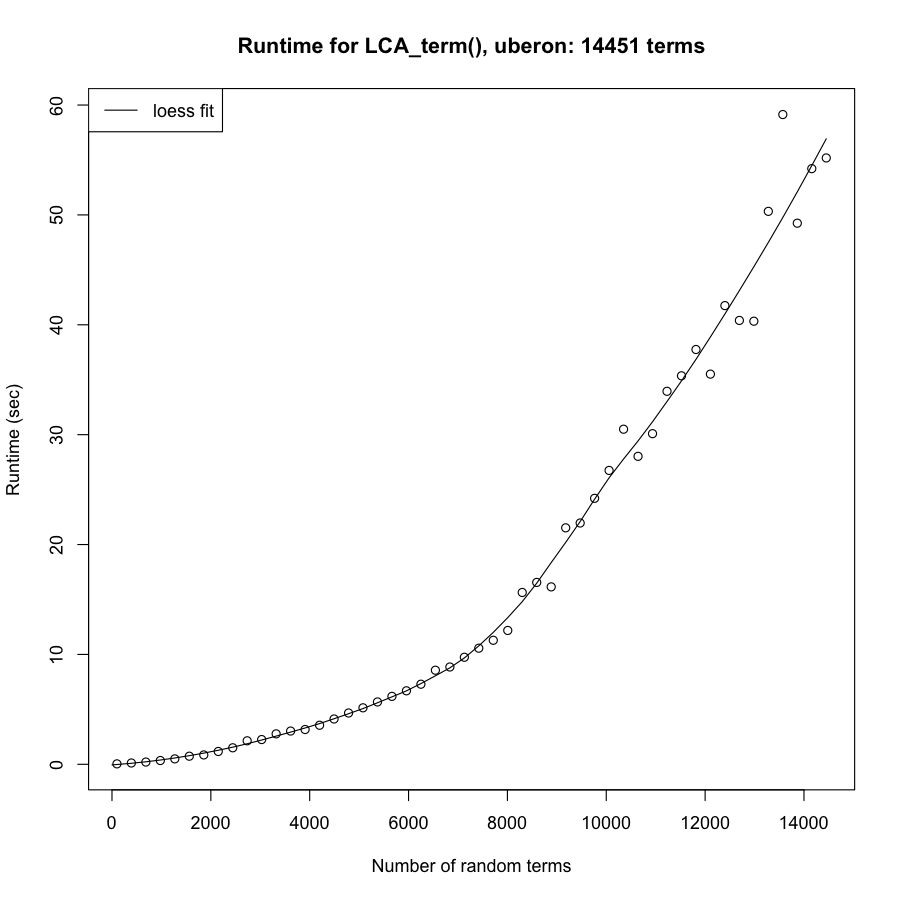

### OBOFoundry_vsao_heatmap.png

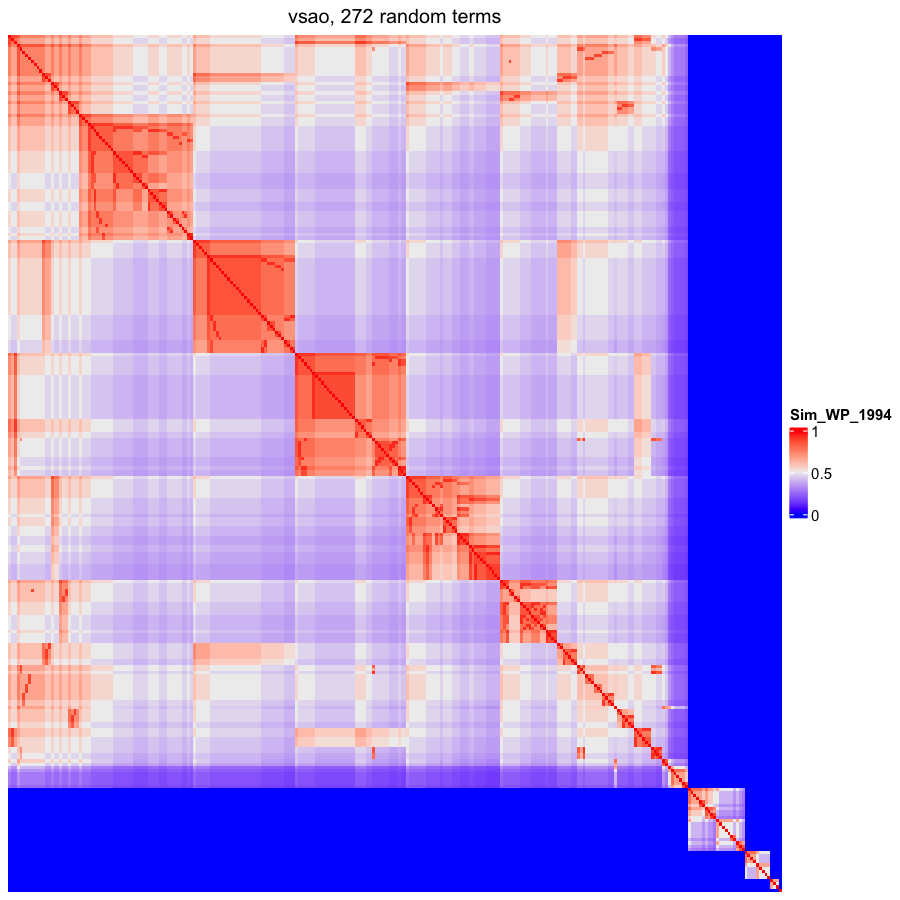

### OBOFoundry_vt_heatmap.png

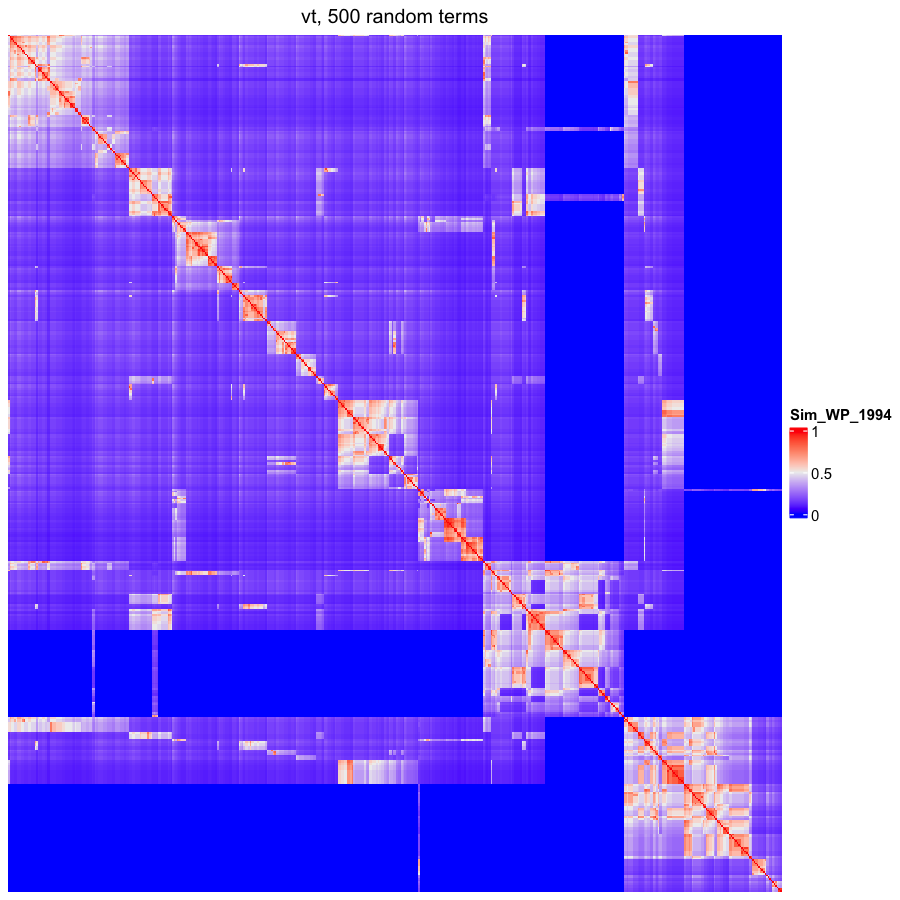
