## Supplementary figures and images for "*simona:* a comprehensive R package for semantic similarity analysis on bio-ontologies"

### OBOFoundry_aism_runtime.png

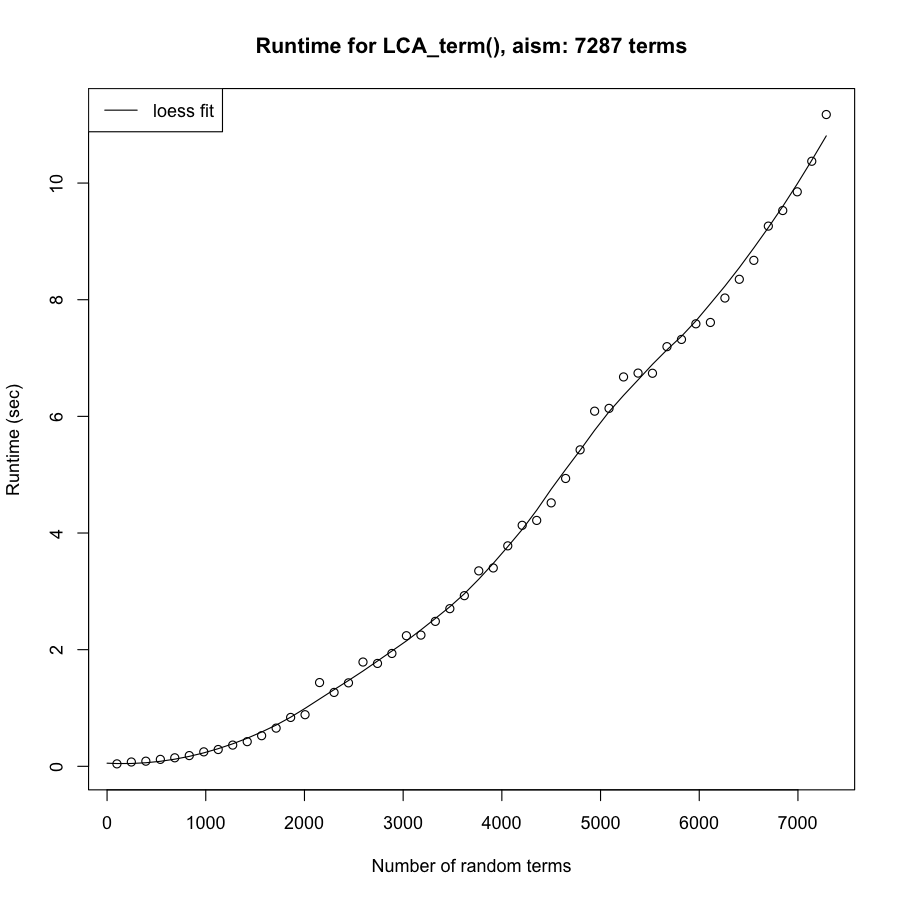

### OBOFoundry_amphx_heatmap.png

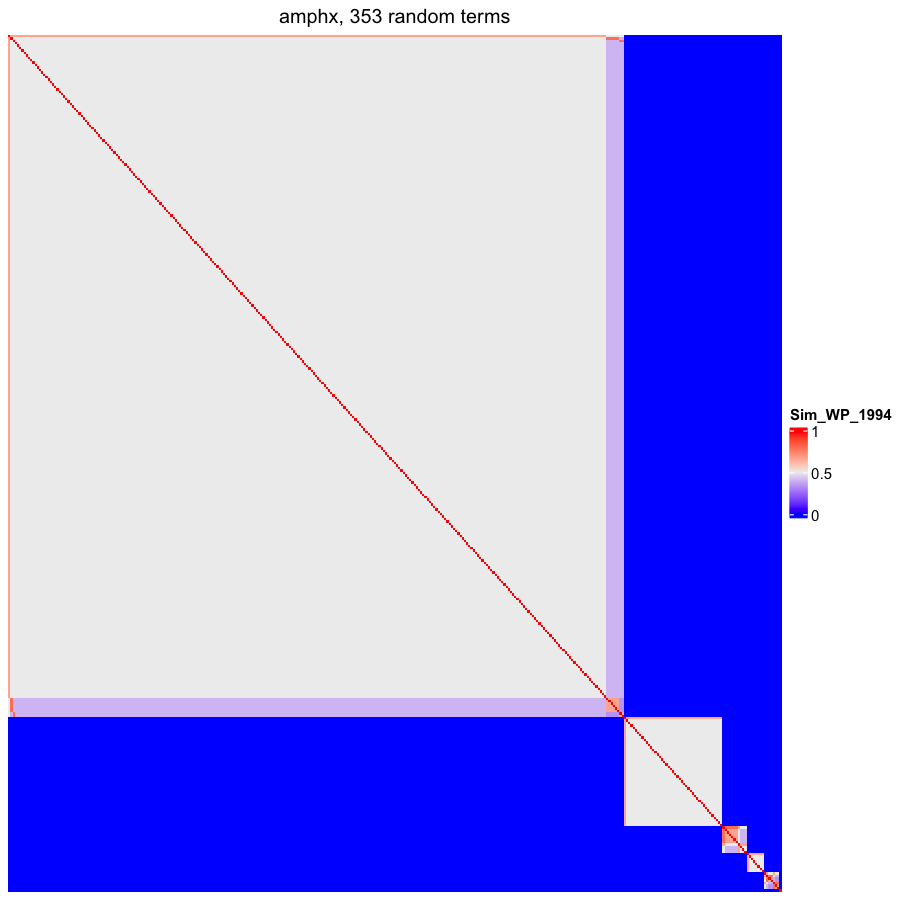

### OBOFoundry_apo_heatmap.png

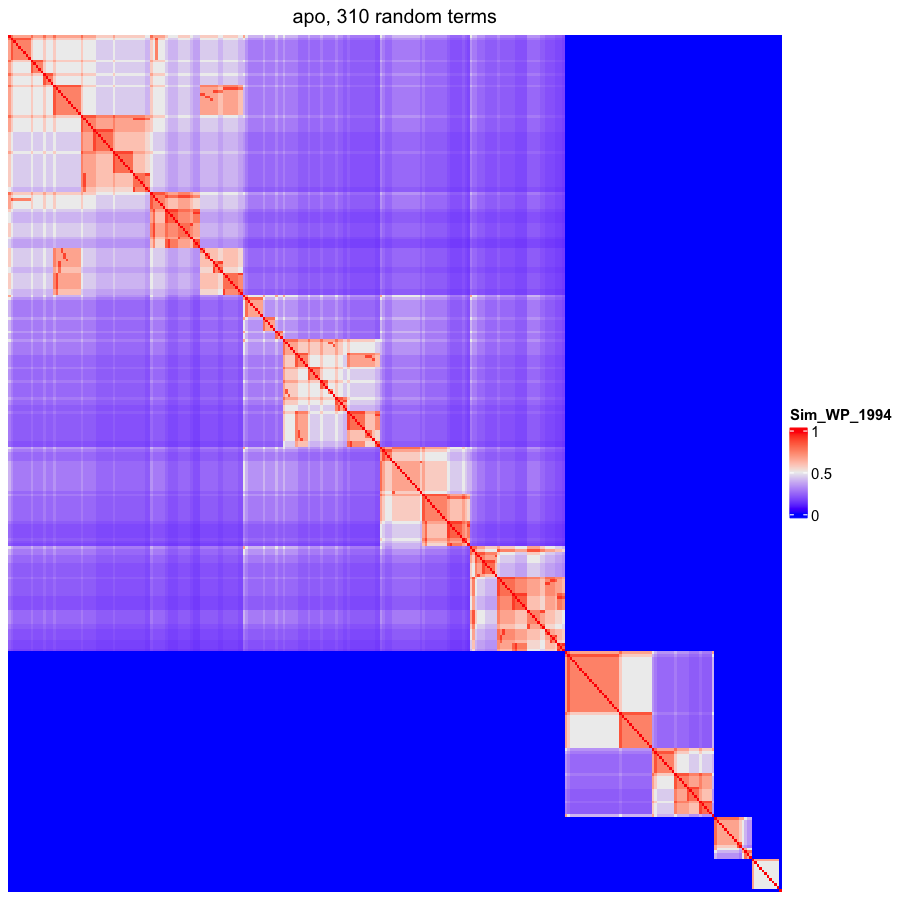

### OBOFoundry_apollo_sv_runtime.png

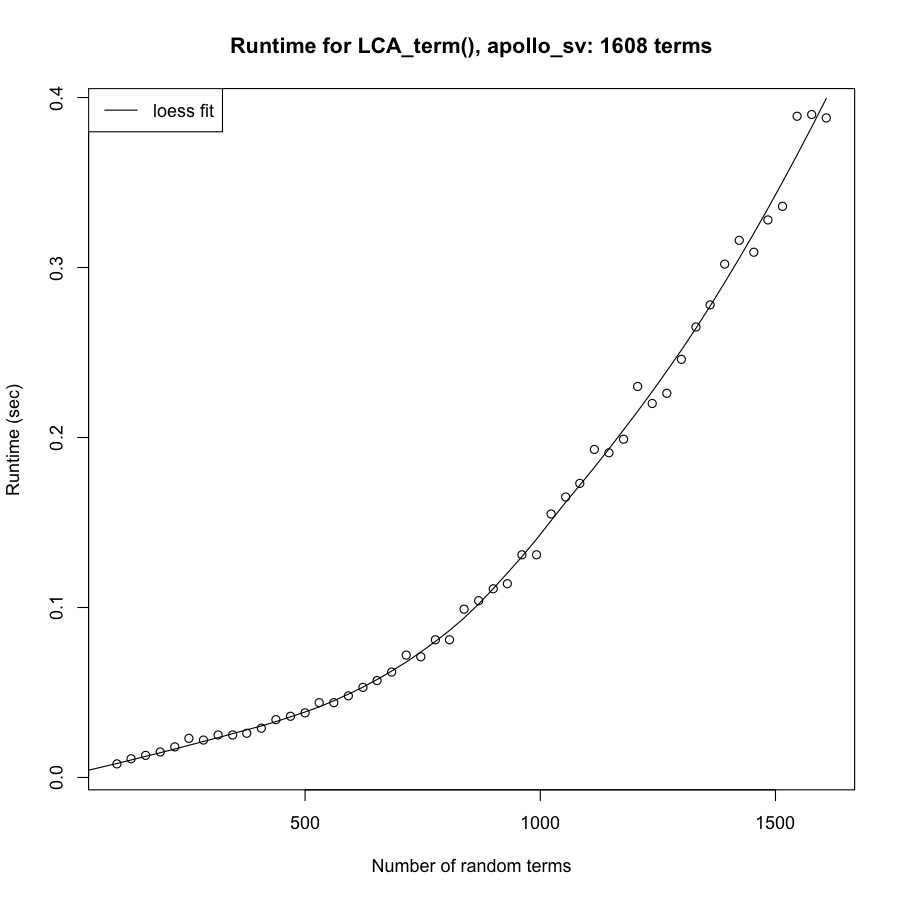

### OBOFoundry_aro_runtime.png

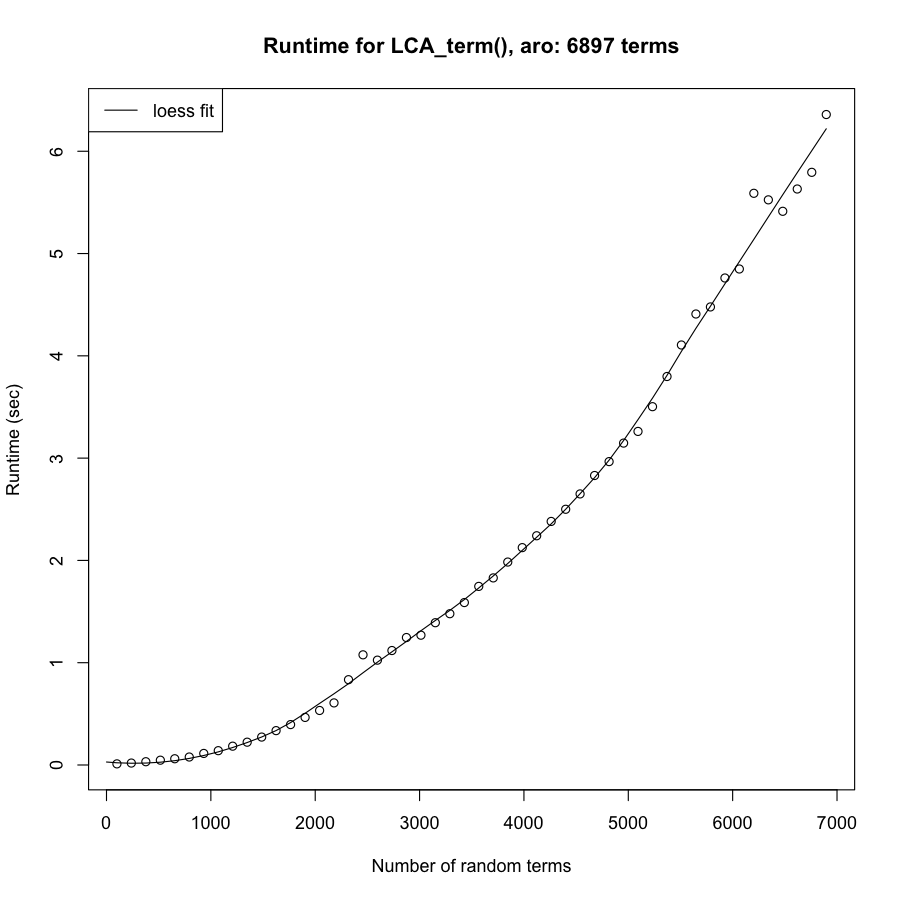

### OBOFoundry_bto_runtime.png

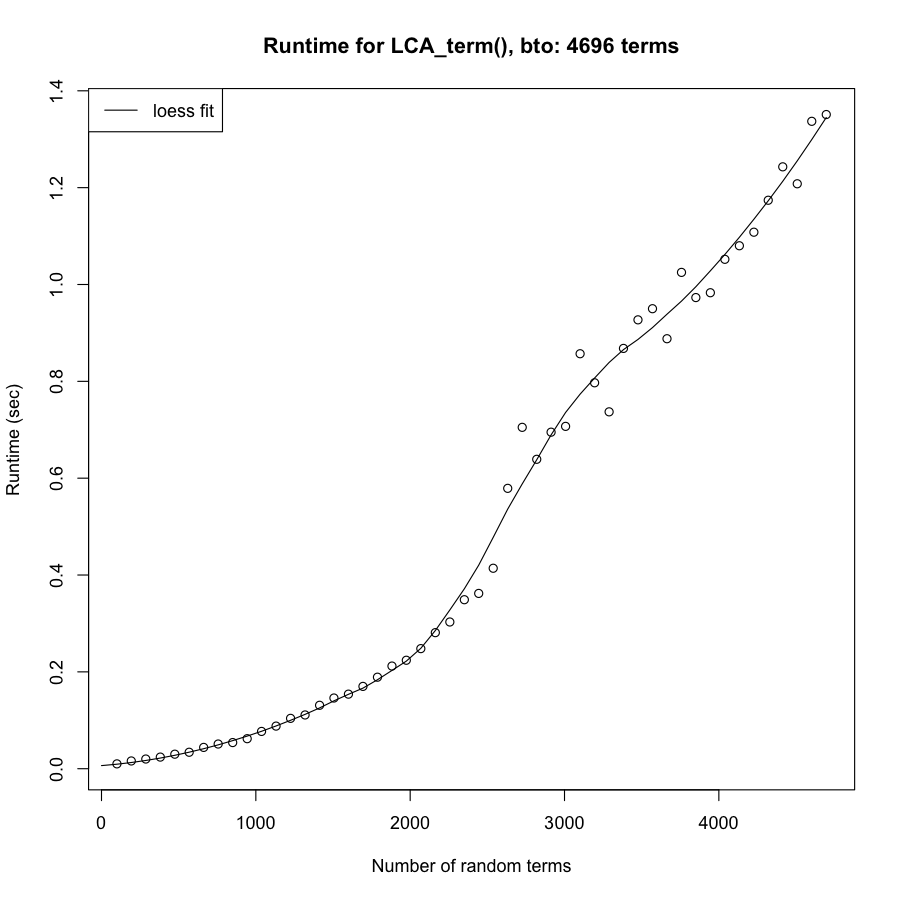

### OBOFoundry_cdno_runtime.png

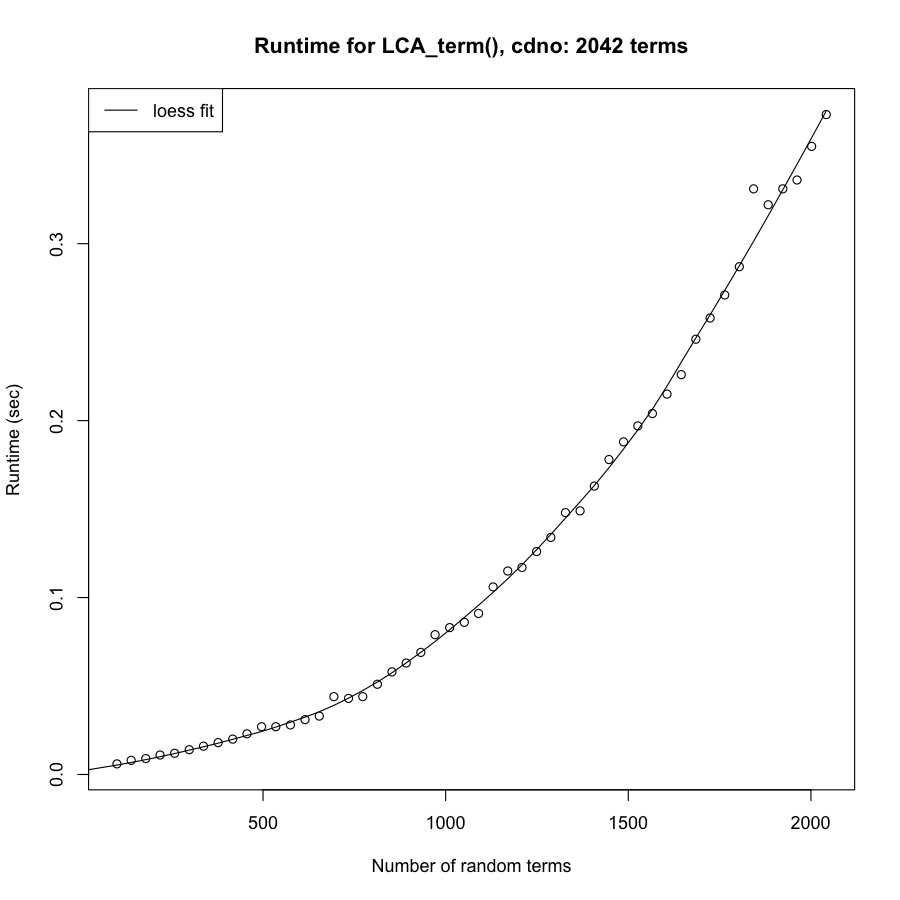

### OBOFoundry_chebi_heatmap.png

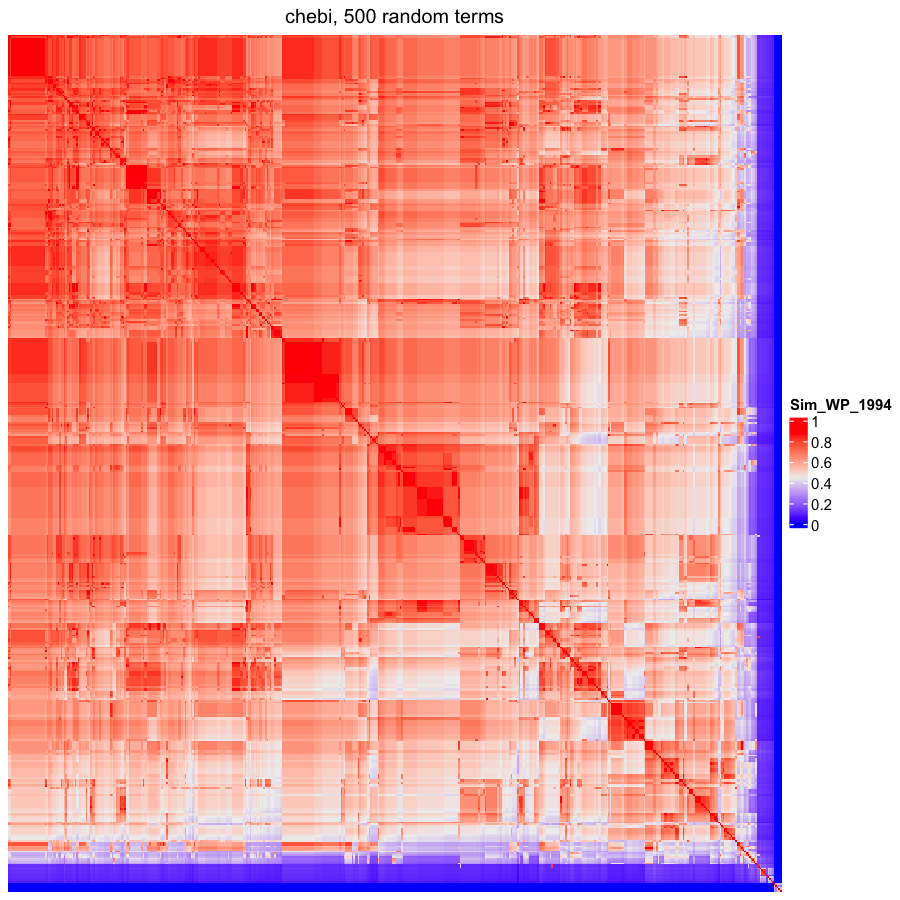

### OBOFoundry_cheminf.png

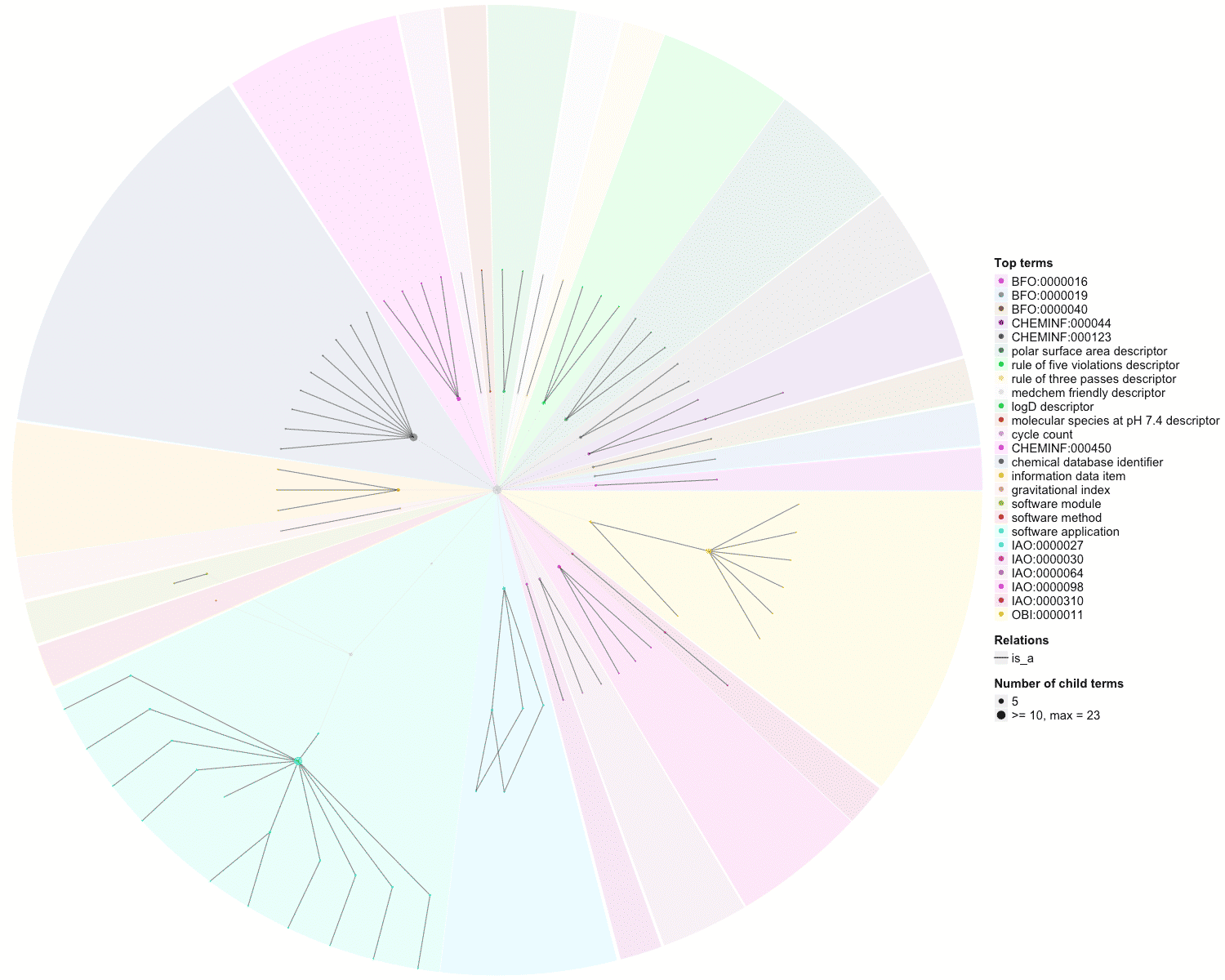

### OBOFoundry_cido_runtime.png

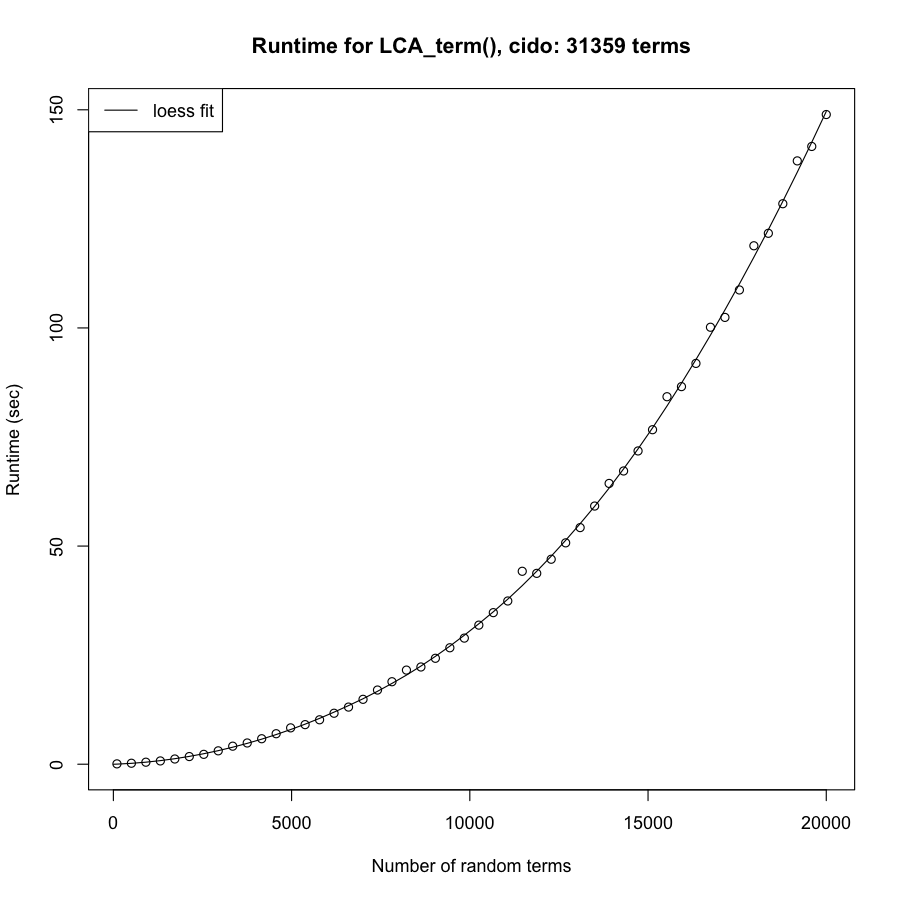

### OBOFoundry_cl_runtime.png

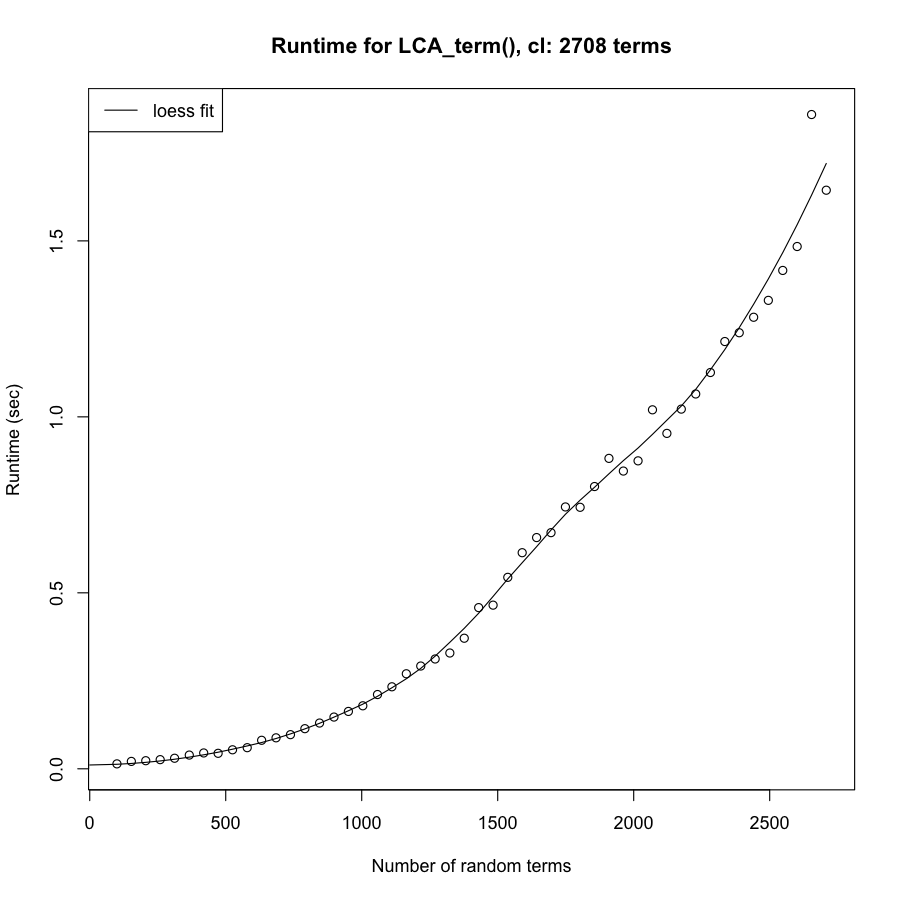

### OBOFoundry_clao_runtime.png

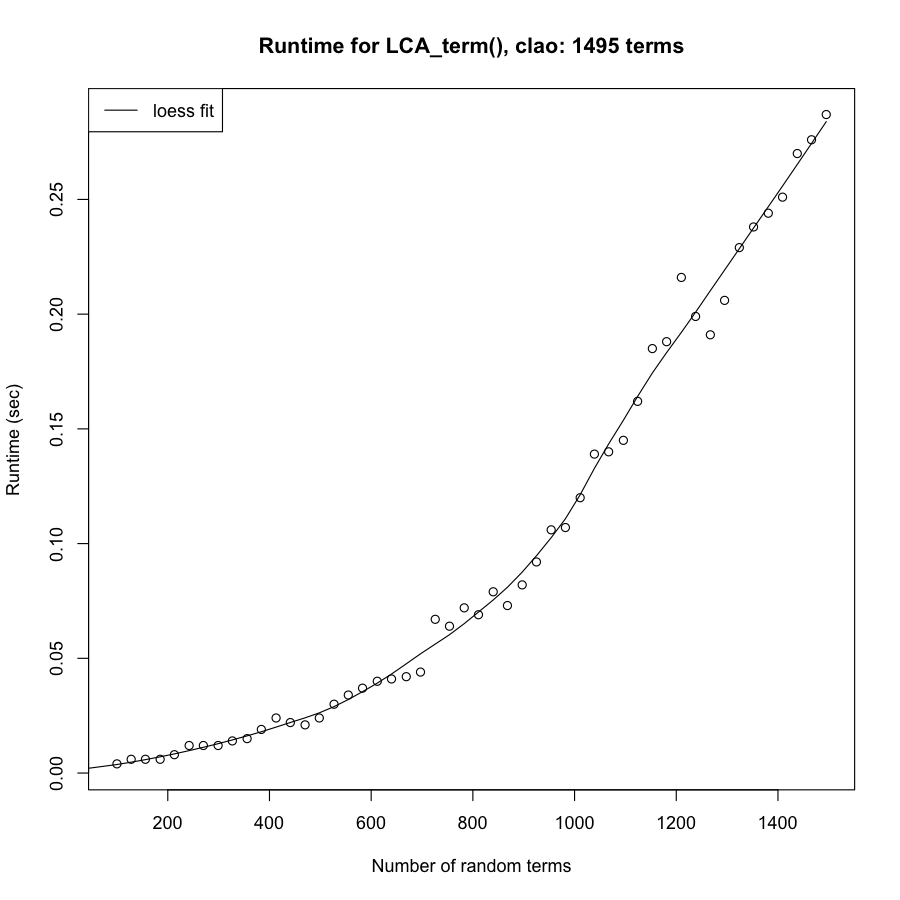

### OBOFoundry_clo_heatmap.png

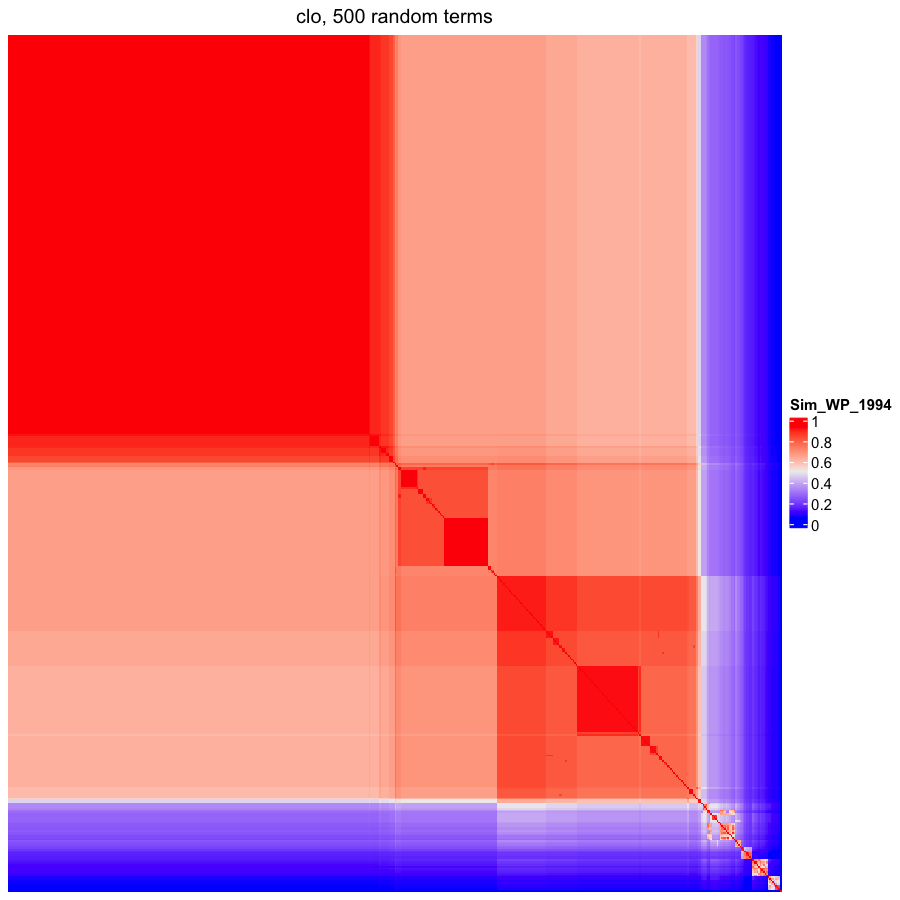

### OBOFoundry_clyh_heatmap.png

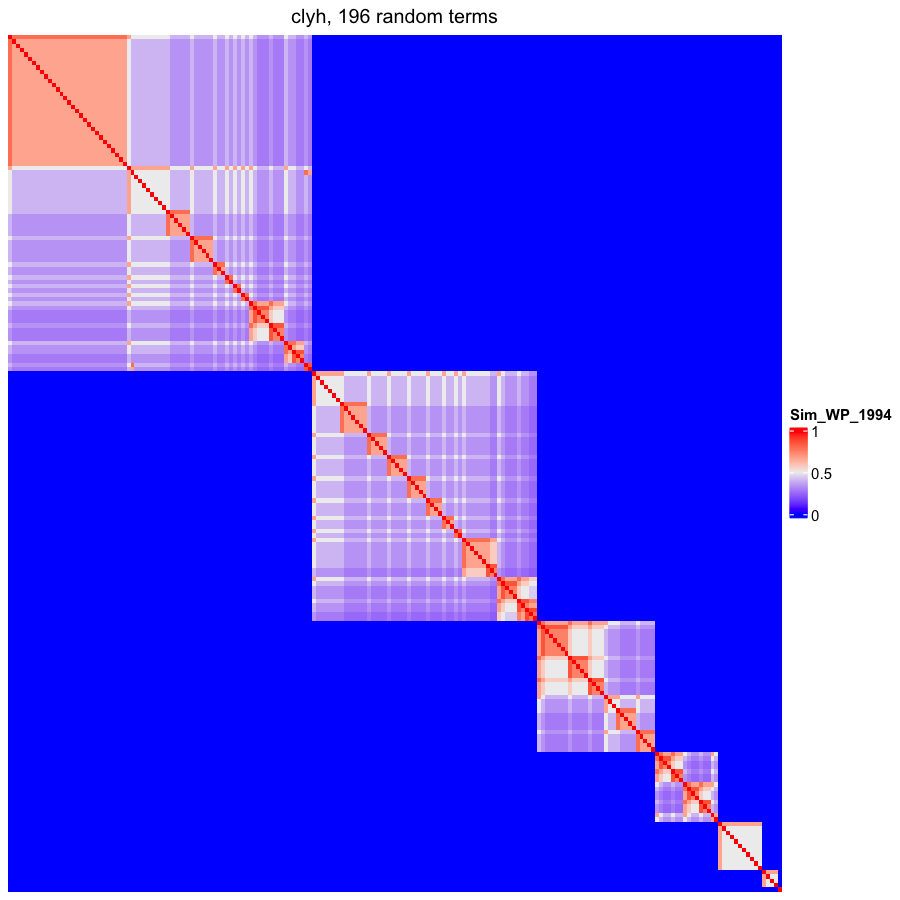

### OBOFoundry_colao.png

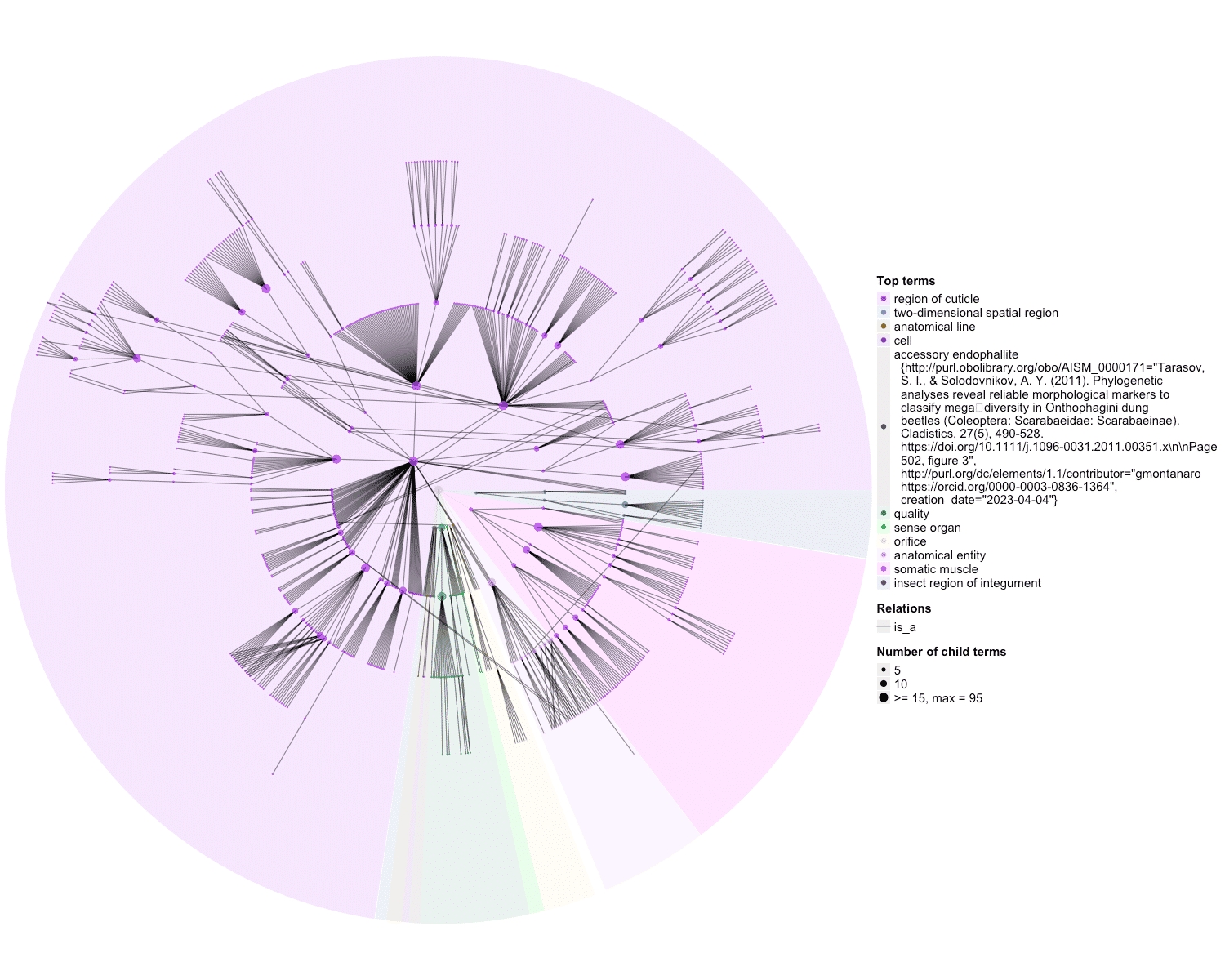

### OBOFoundry_cro_heatmap.png

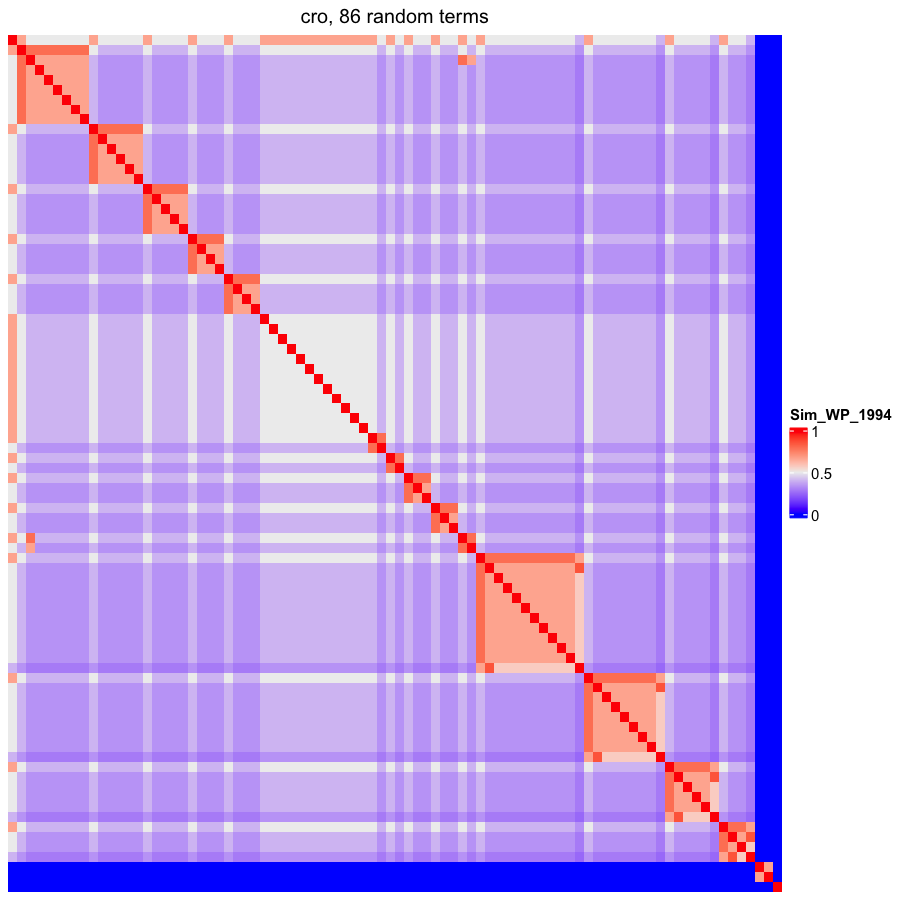

### OBOFoundry_cto_heatmap.png

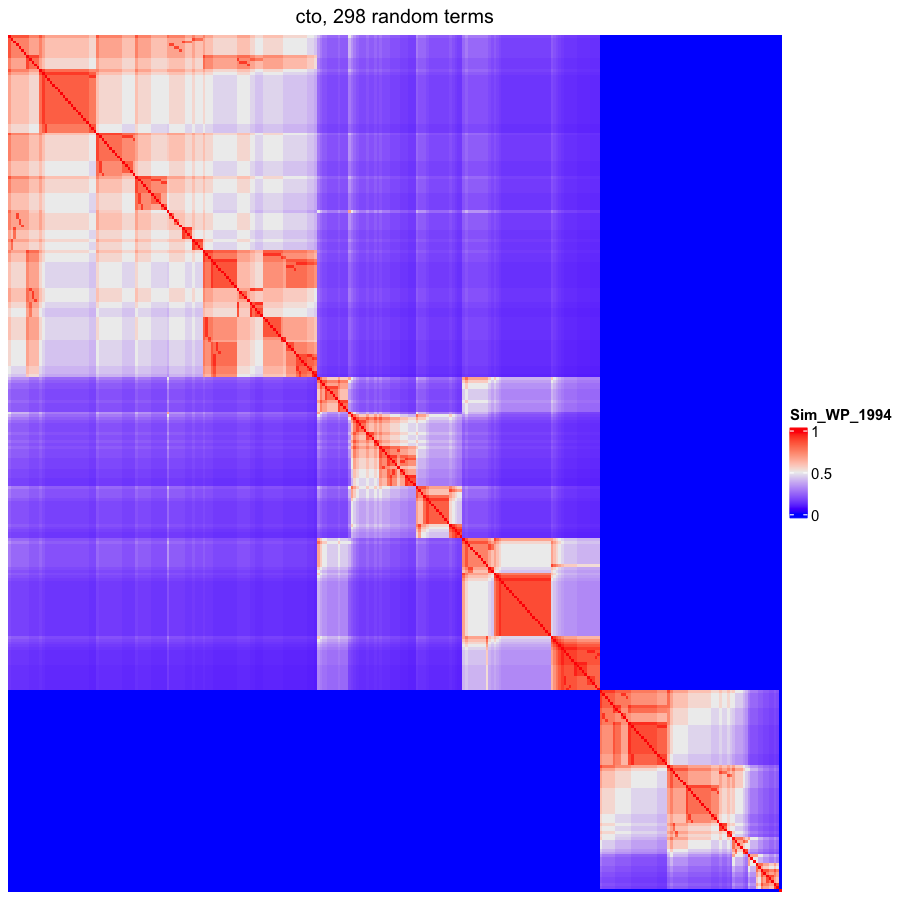

### OBOFoundry_ddanat_heatmap.png

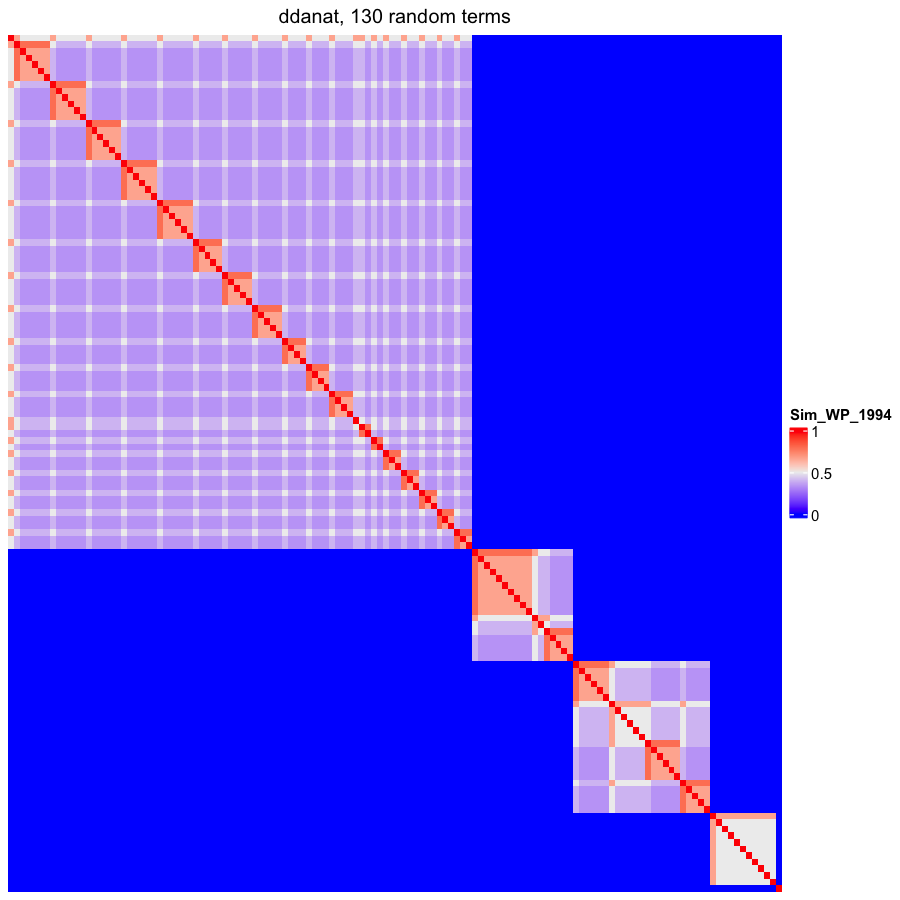

### OBOFoundry_dpo.png

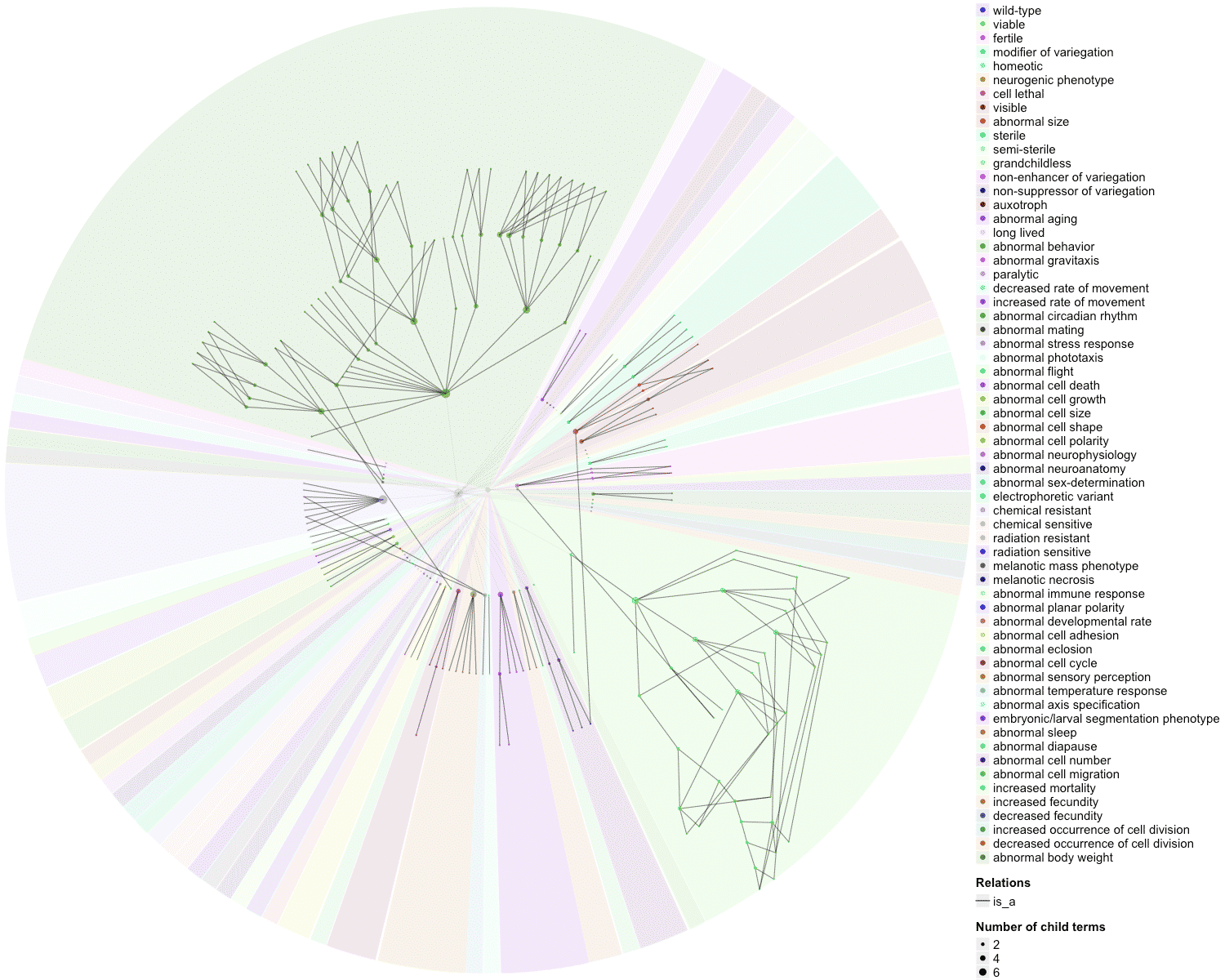

### OBOFoundry_duo_heatmap.png

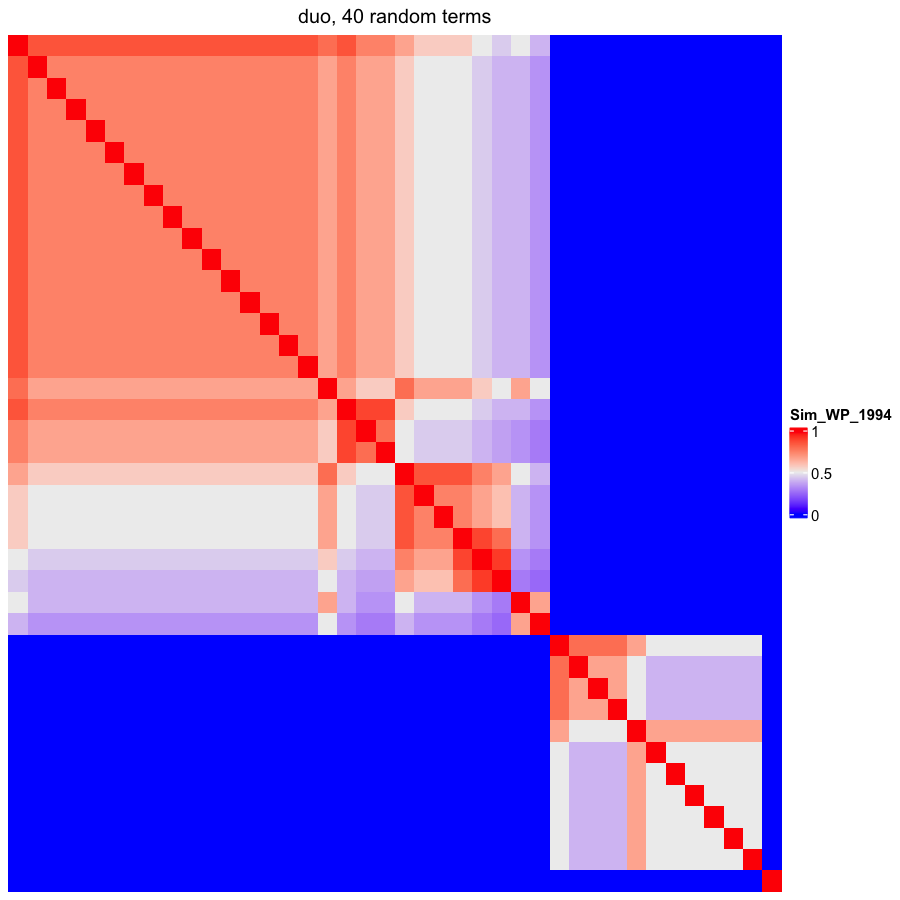

### OBOFoundry_eco.png

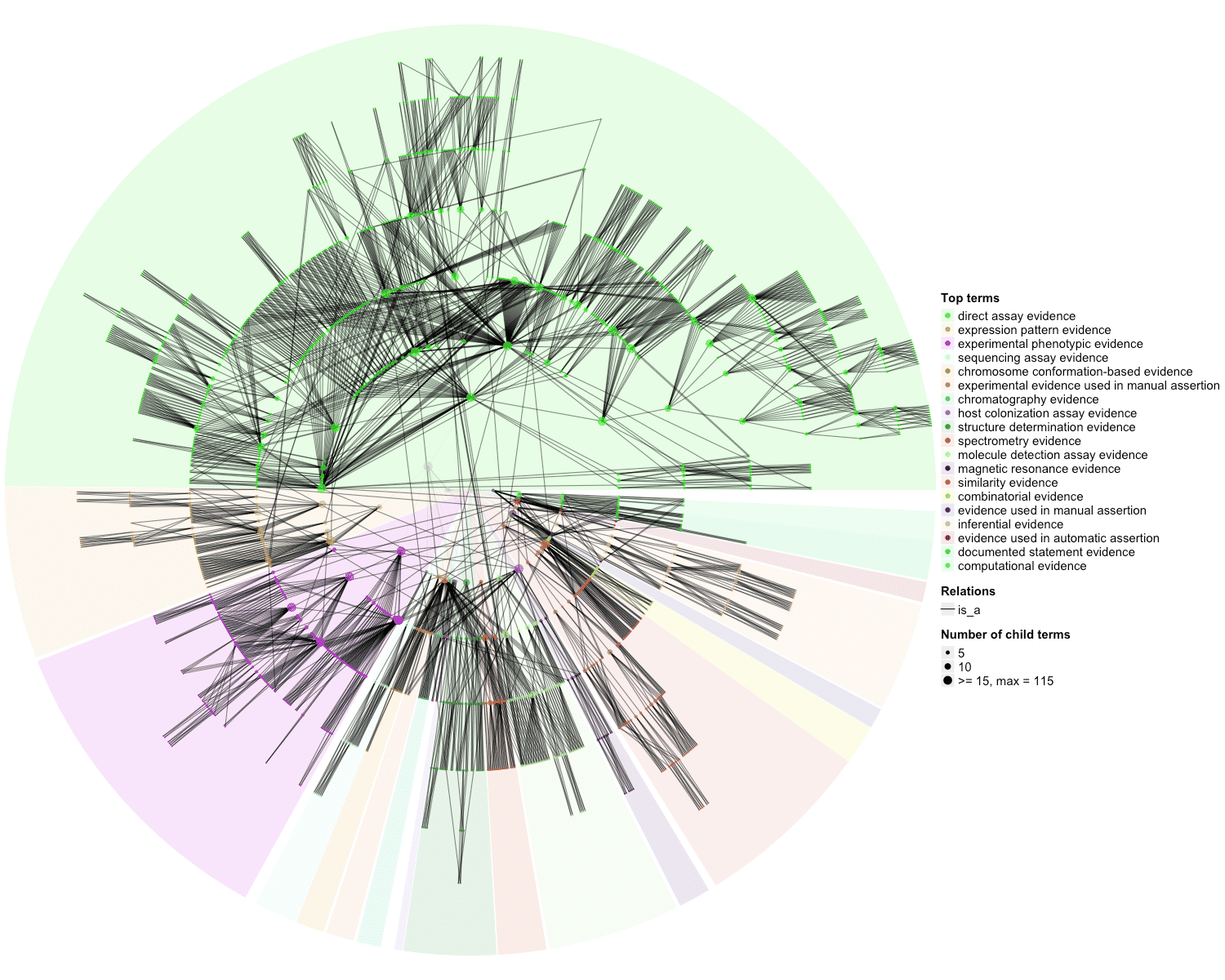

### OBOFoundry_ecocore_heatmap.png

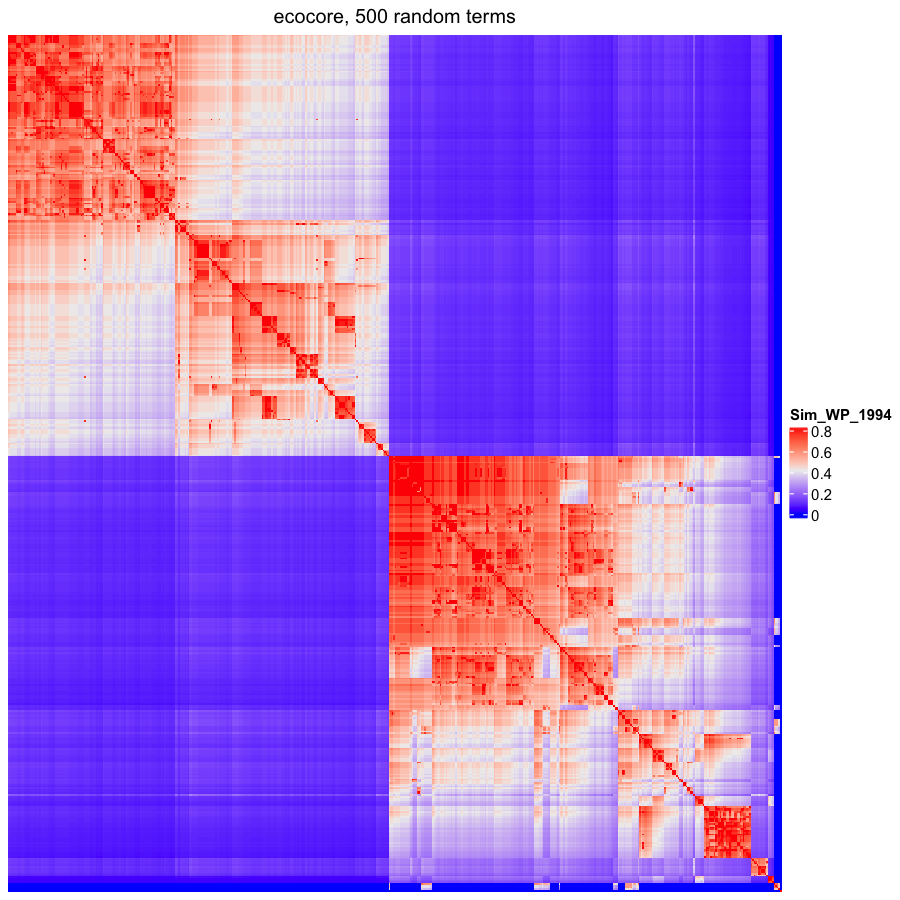

### OBOFoundry_ecto_heatmap.png

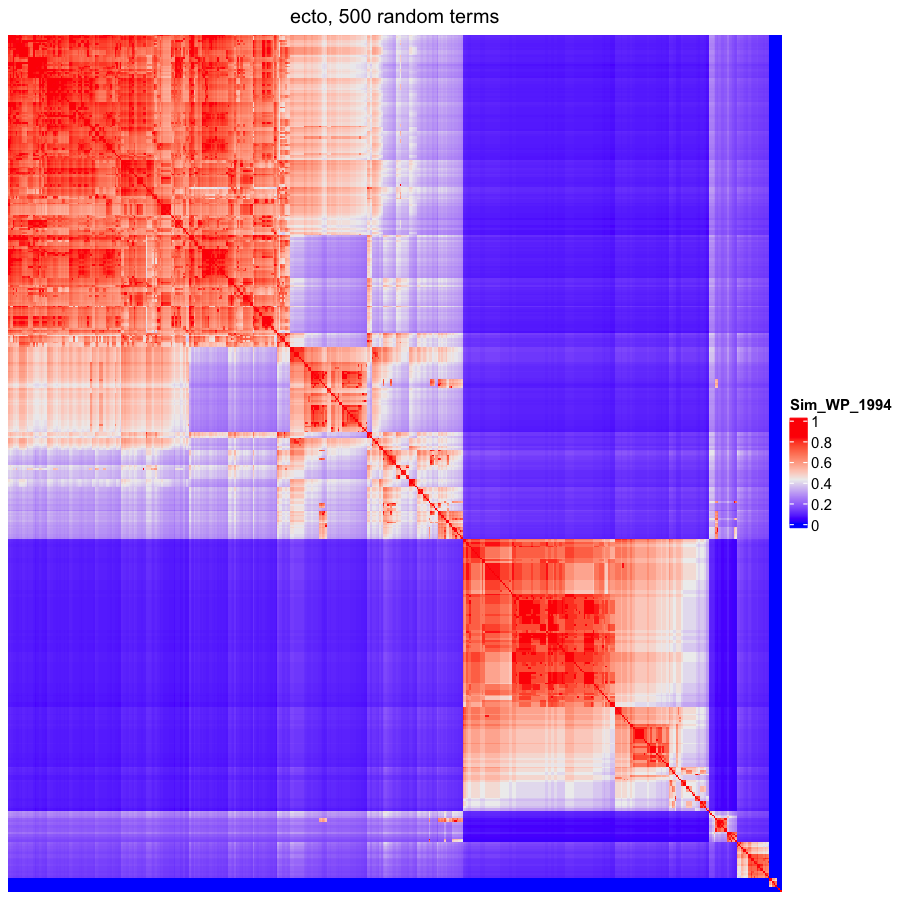

### OBOFoundry_ehdaa2.png

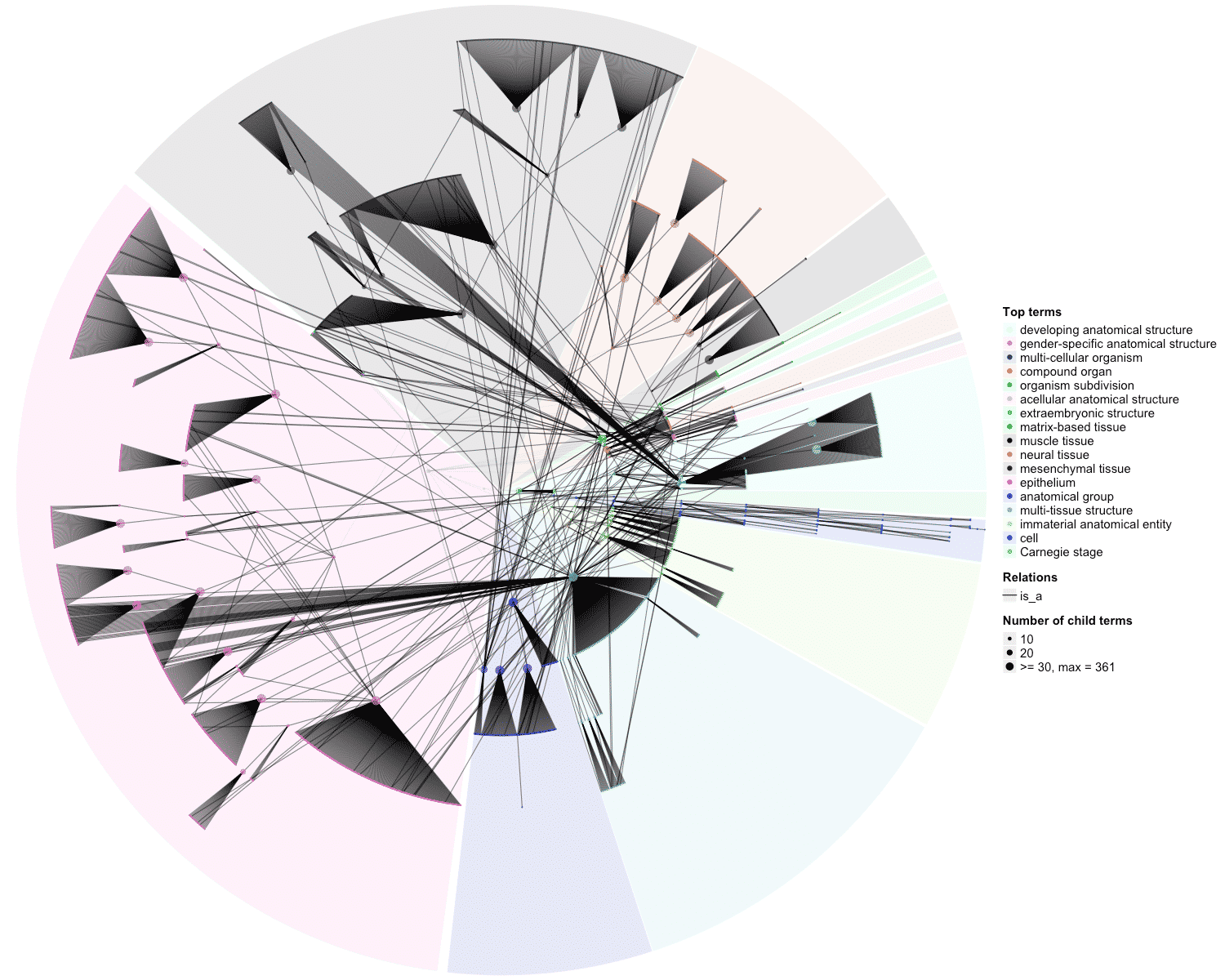

### OBOFoundry_emap_heatmap.png

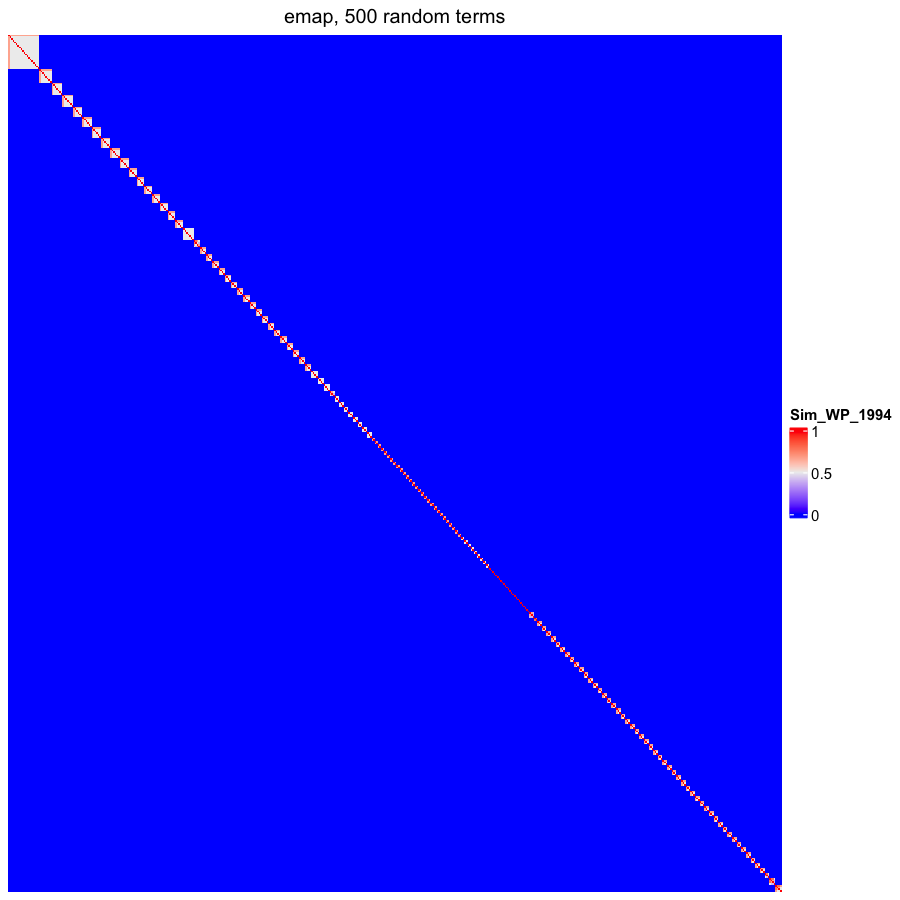

### OBOFoundry_envo.png

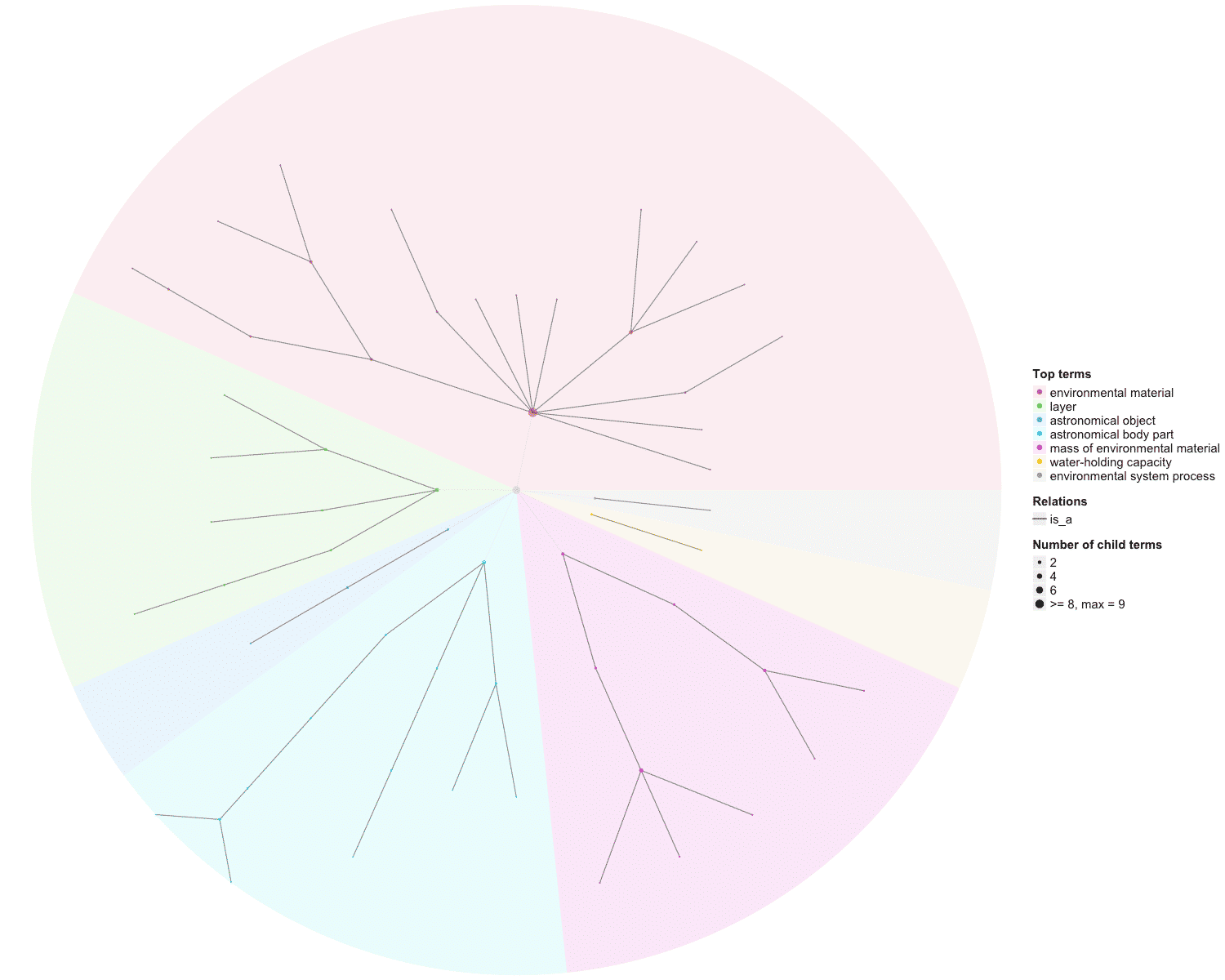

### OBOFoundry_eupath_runtime.png

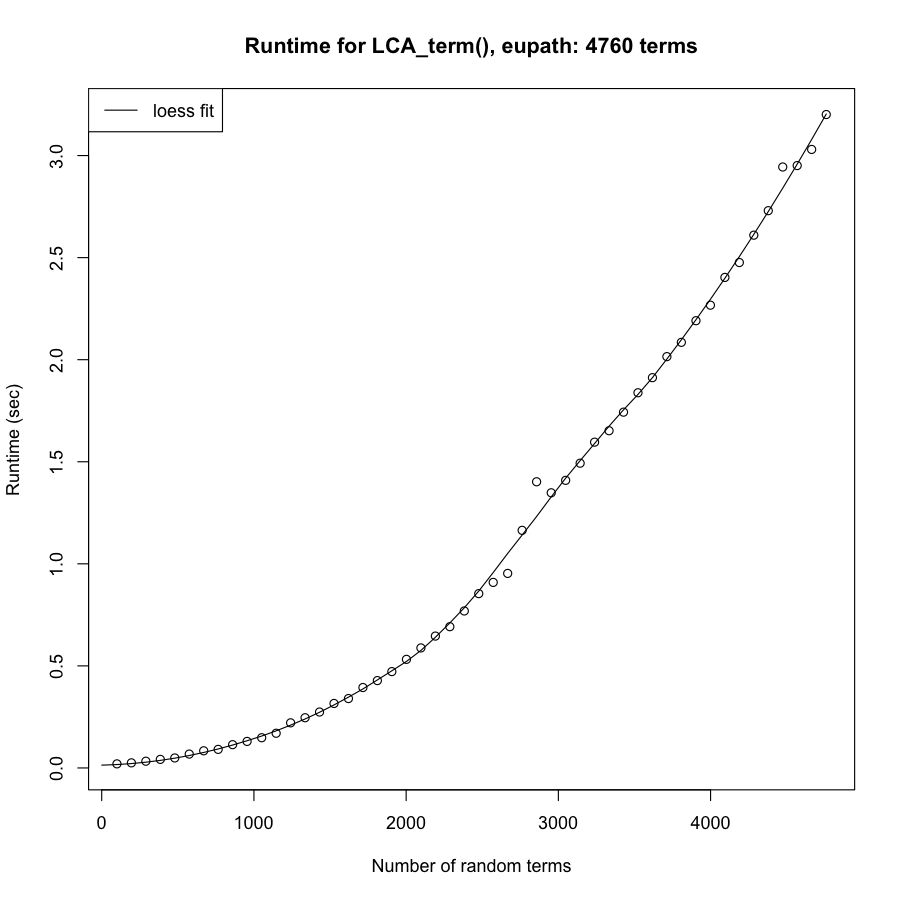

### OBOFoundry_fbsp.png

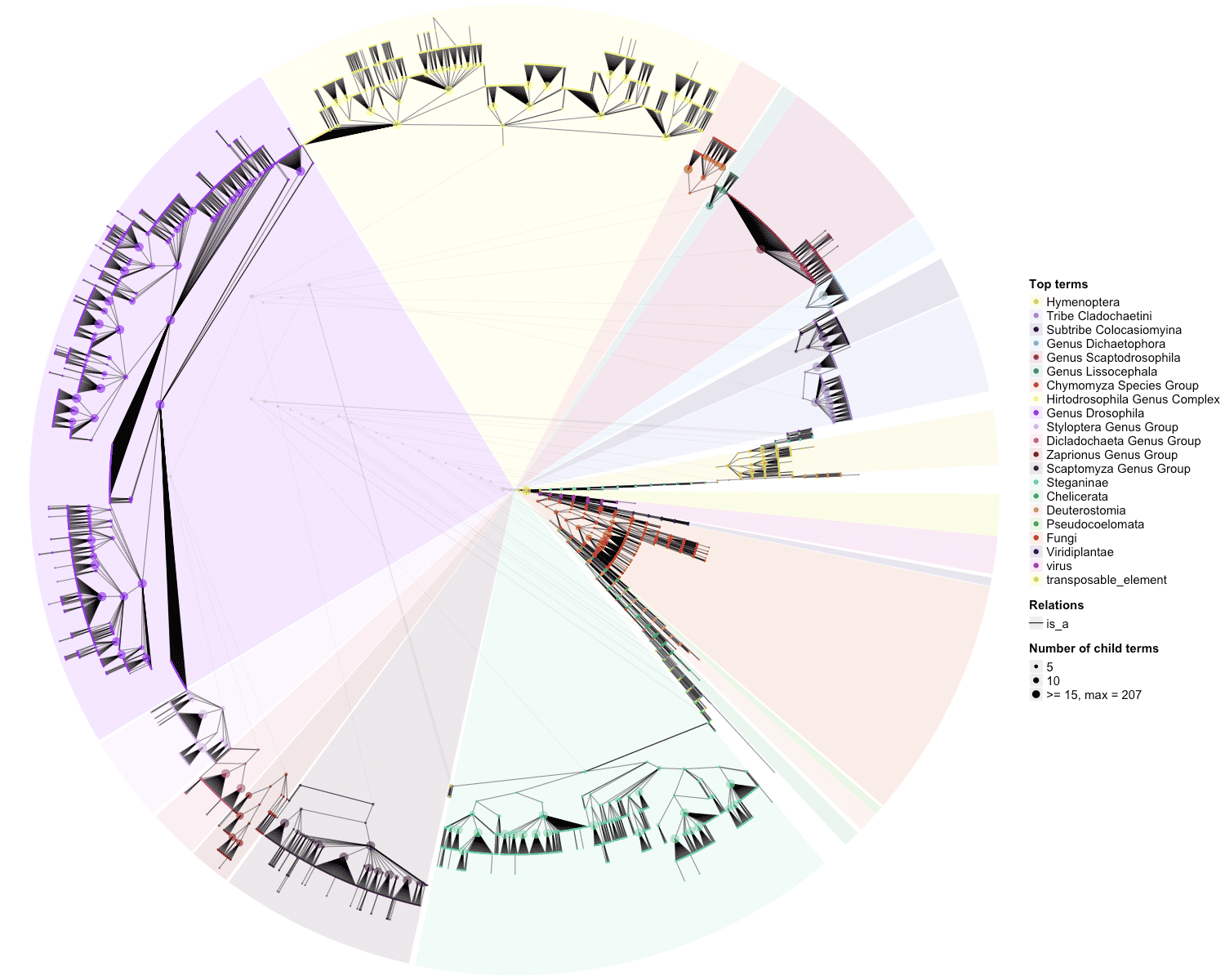

### OBOFoundry_fix.png

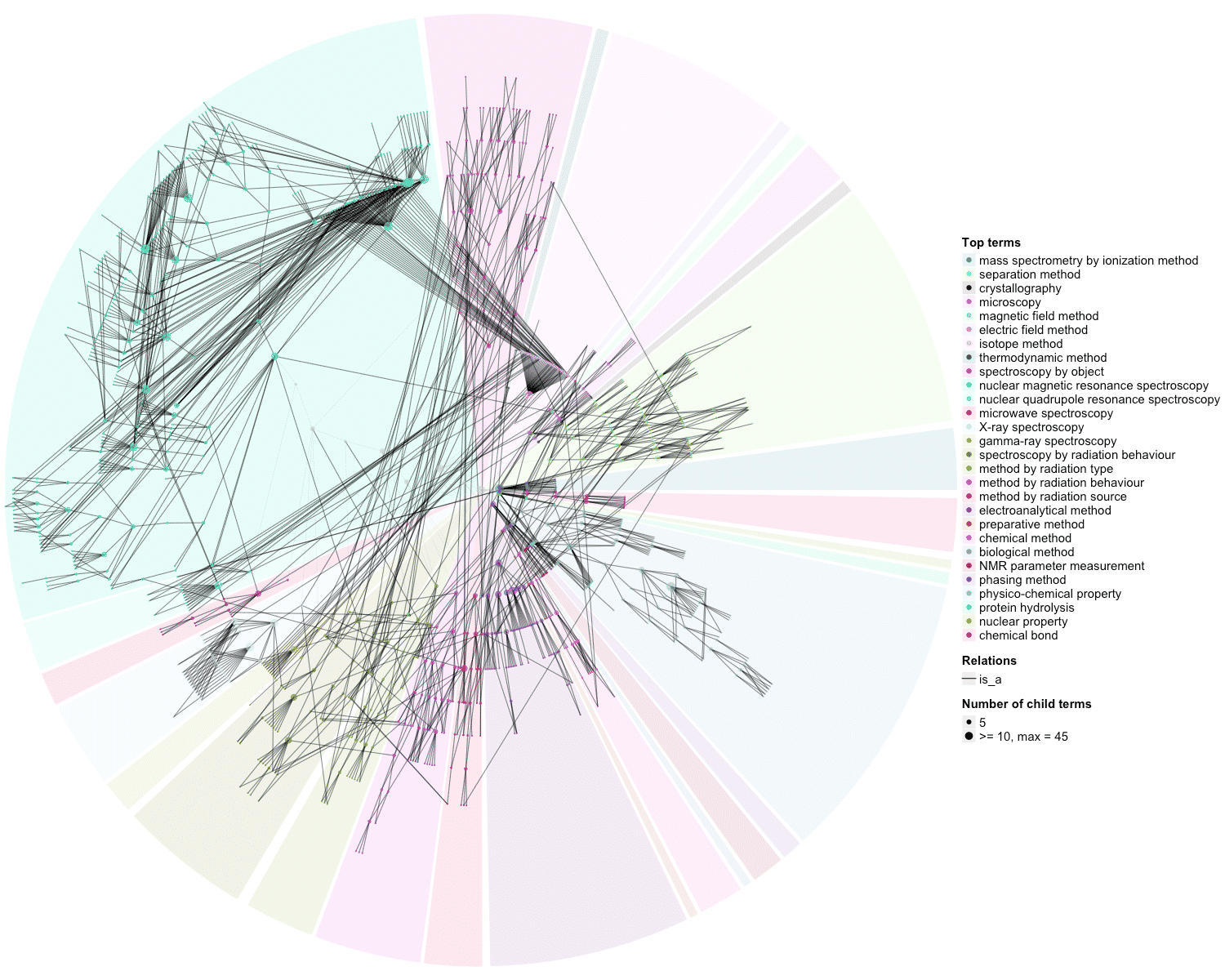

### OBOFoundry_fobi_heatmap.png

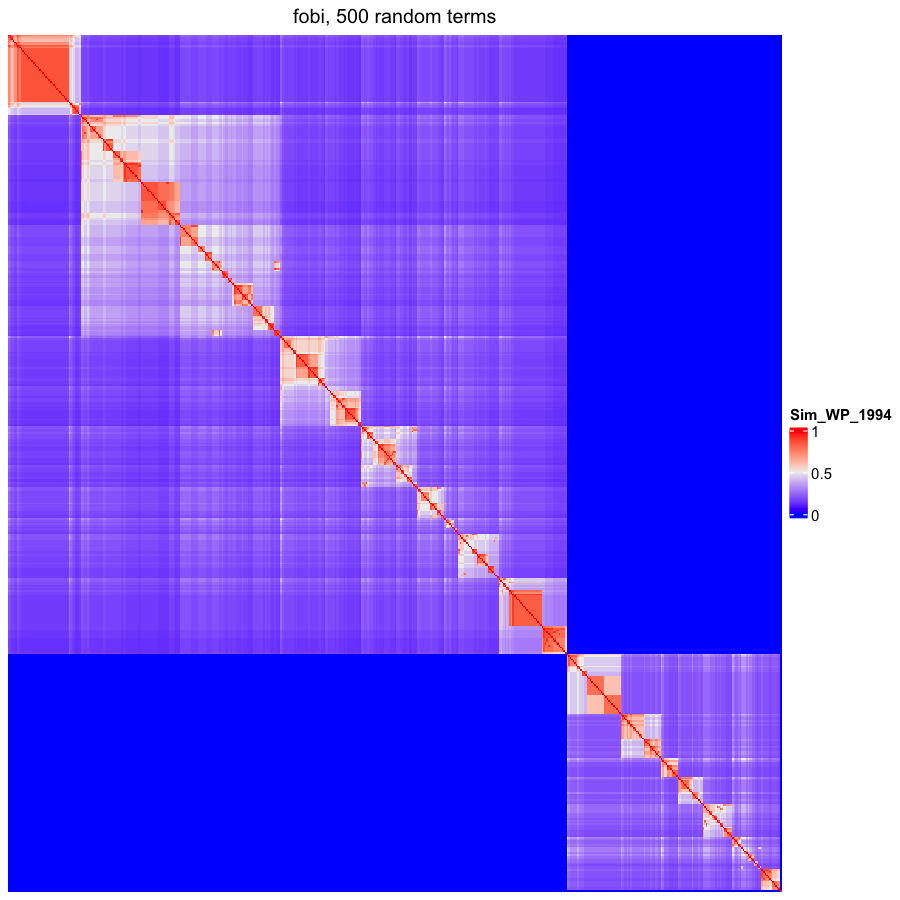

### OBOFoundry_fovt.png

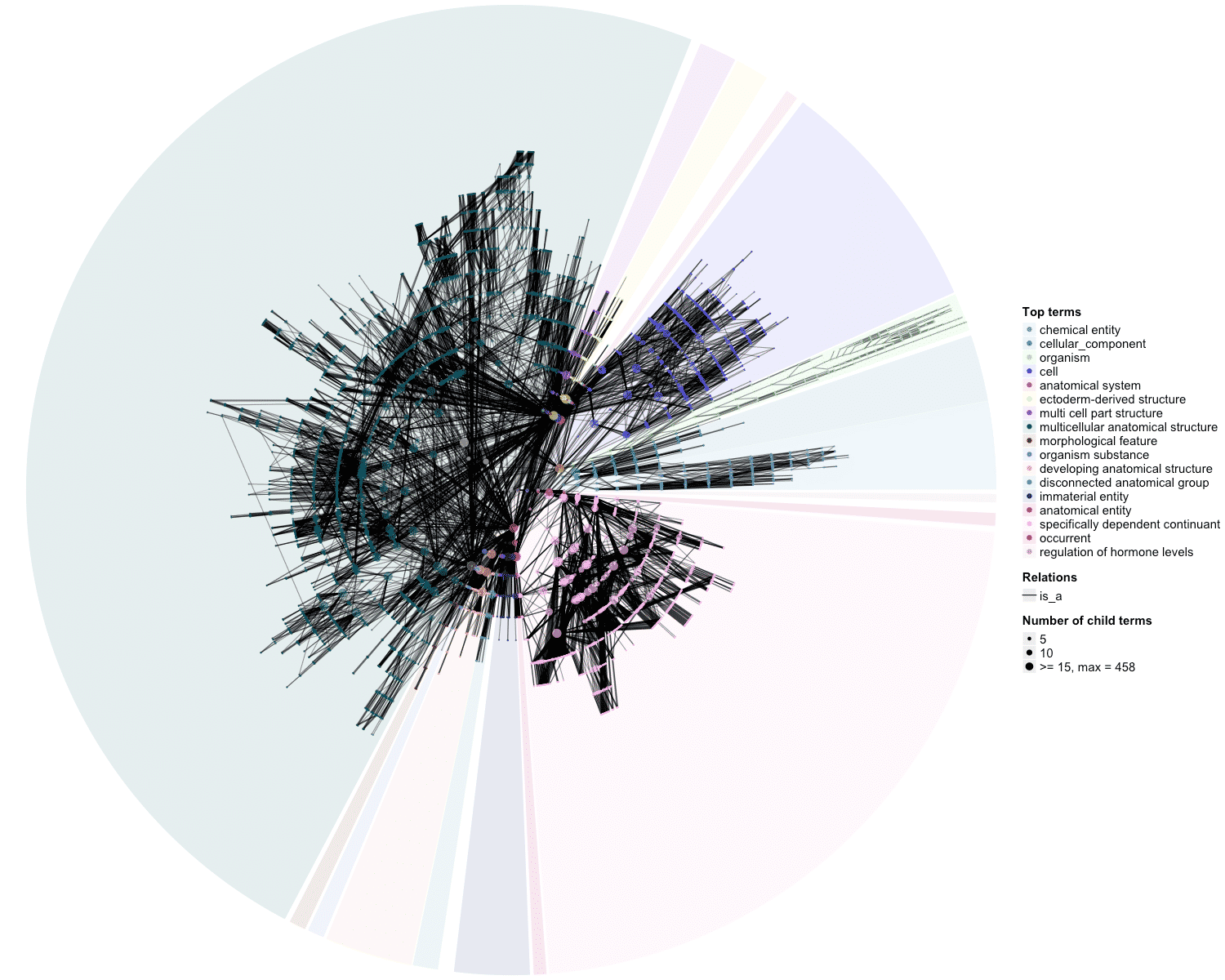

### OBOFoundry_fypo.png

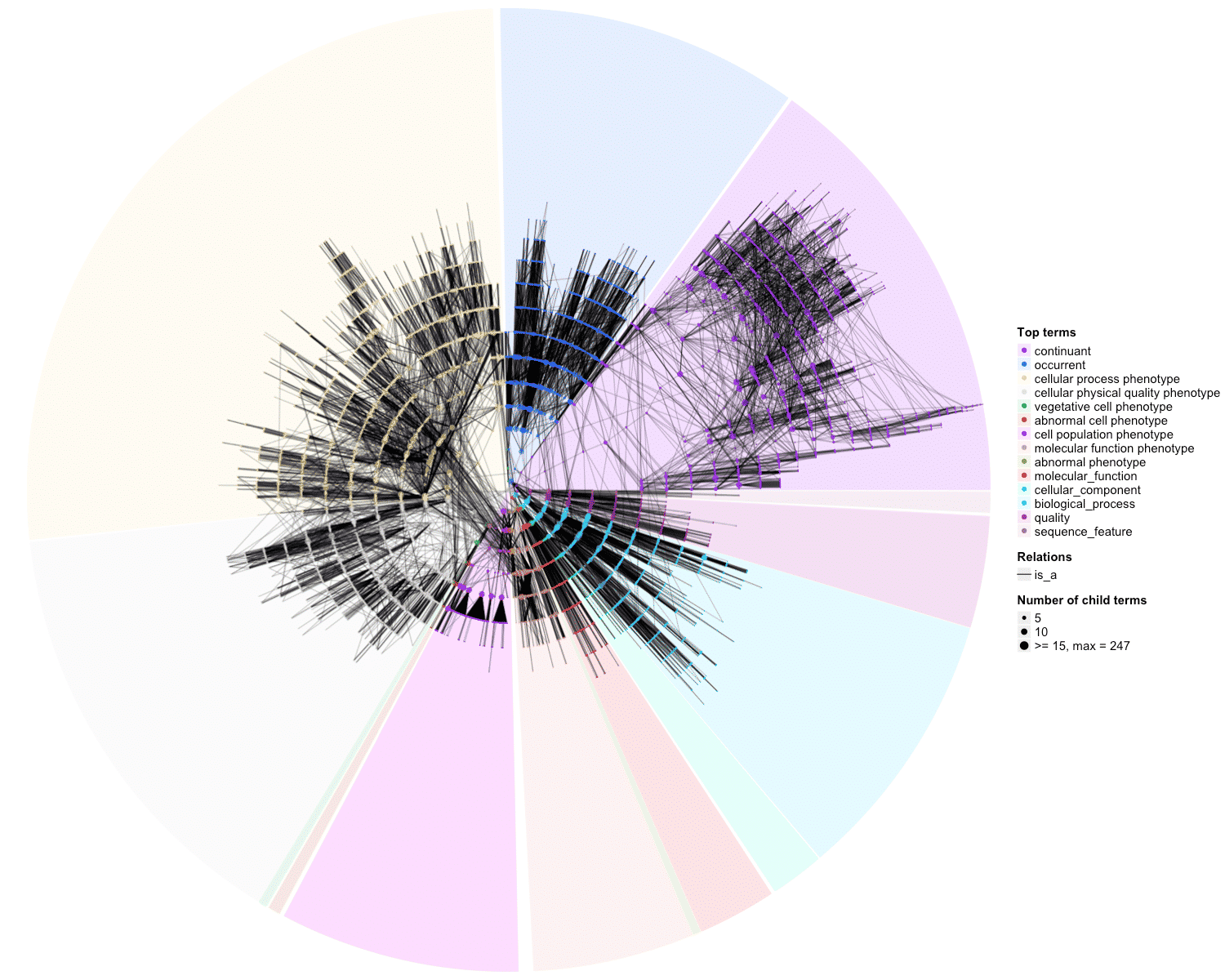

### OBOFoundry_fypo_heatmap.png

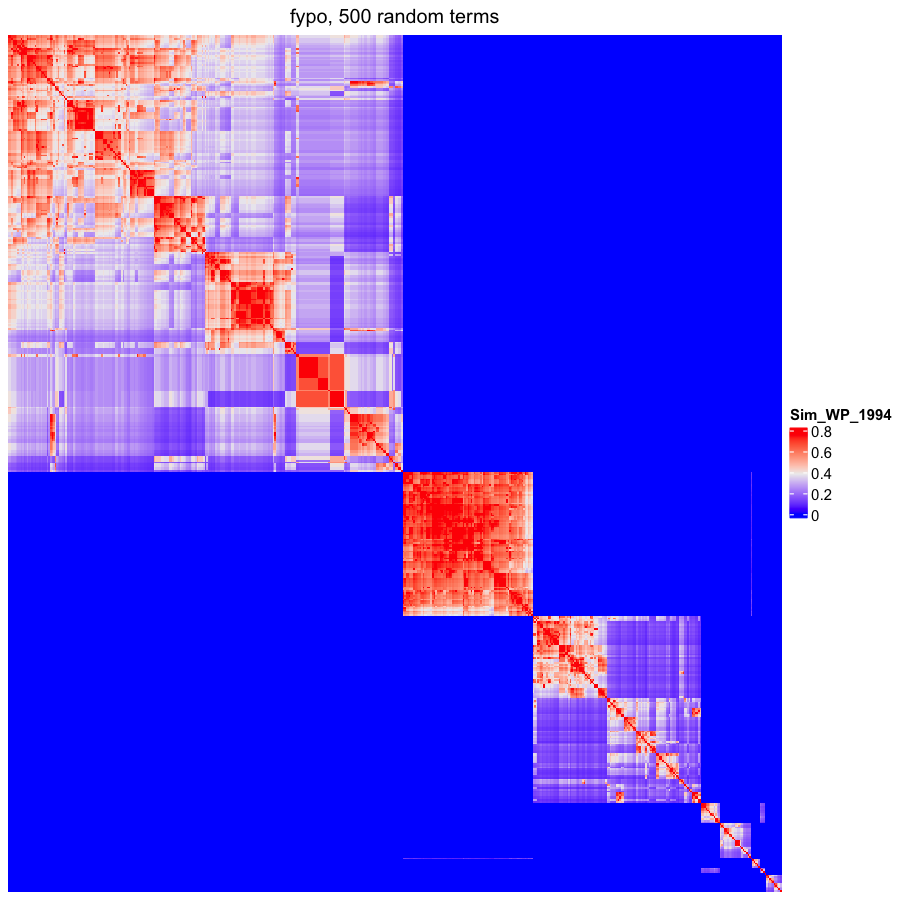

### OBOFoundry_gaz_heatmap.png

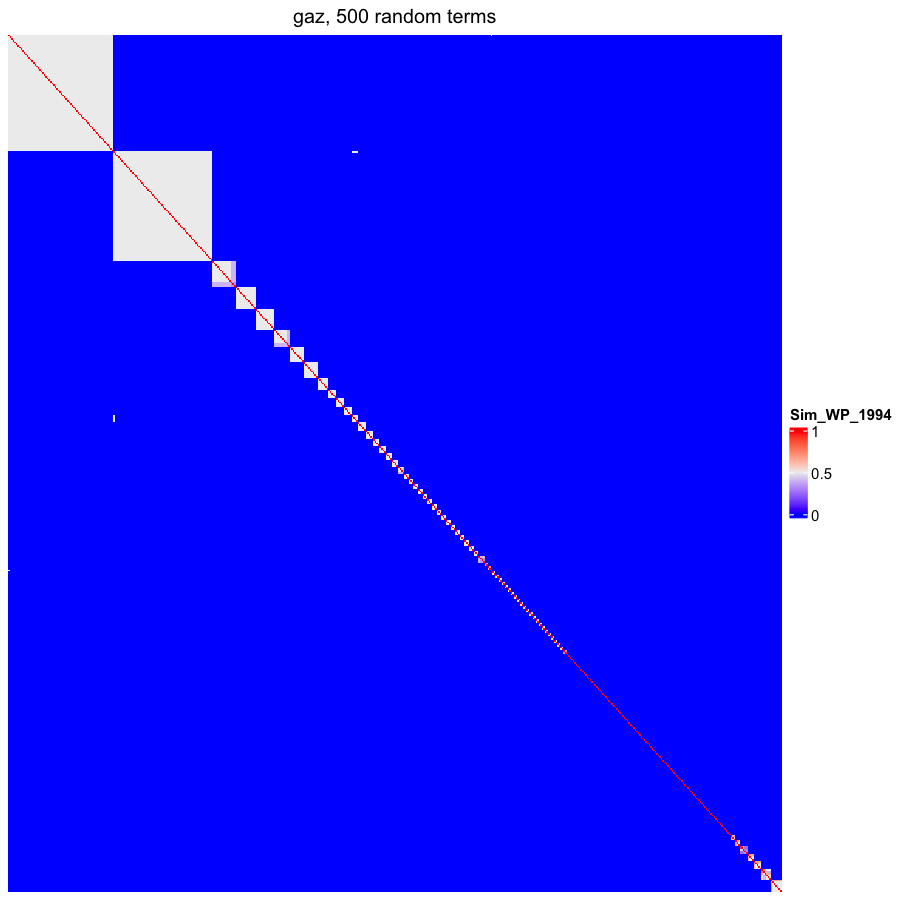

### OBOFoundry_gno.png

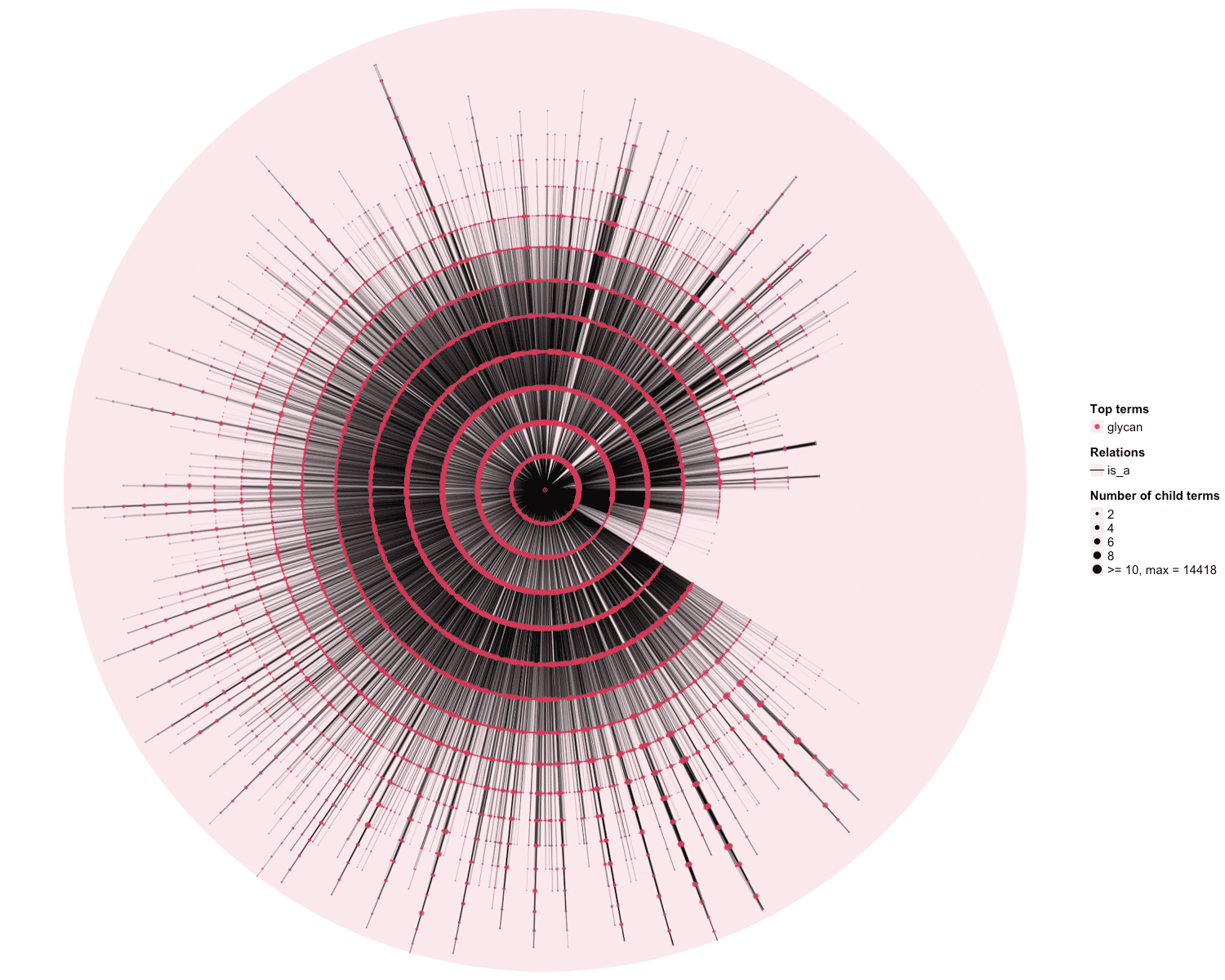

### OBOFoundry_gno_runtime.png

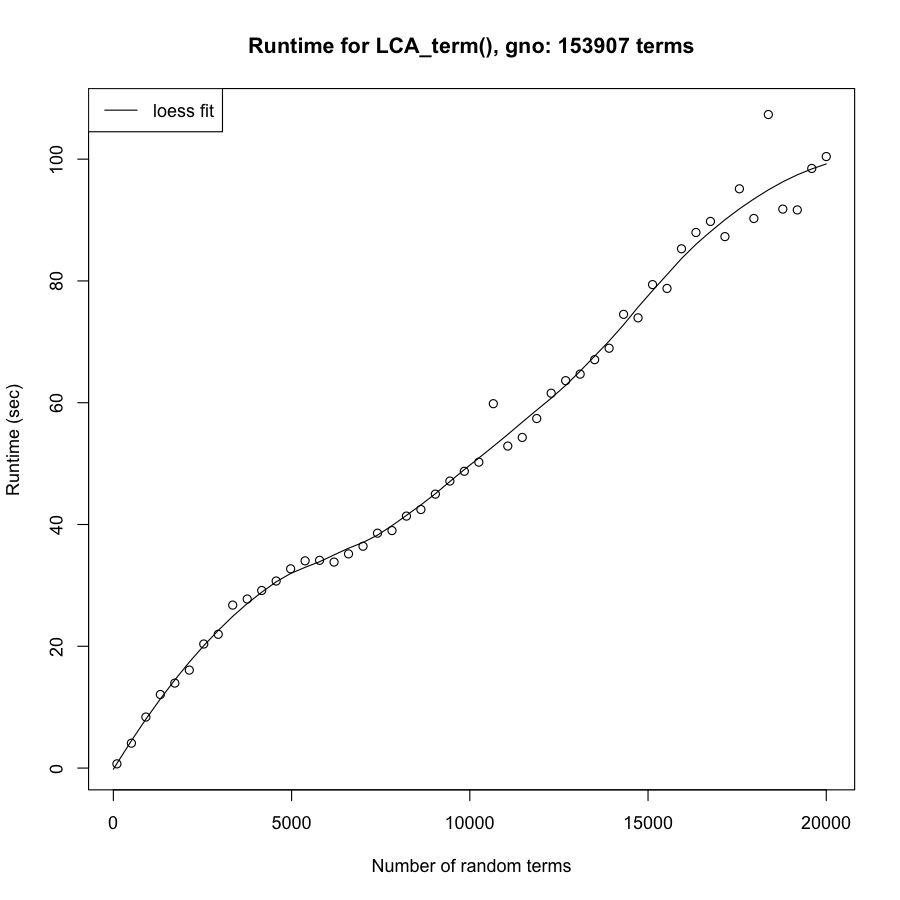

### OBOFoundry_hao_runtime.png

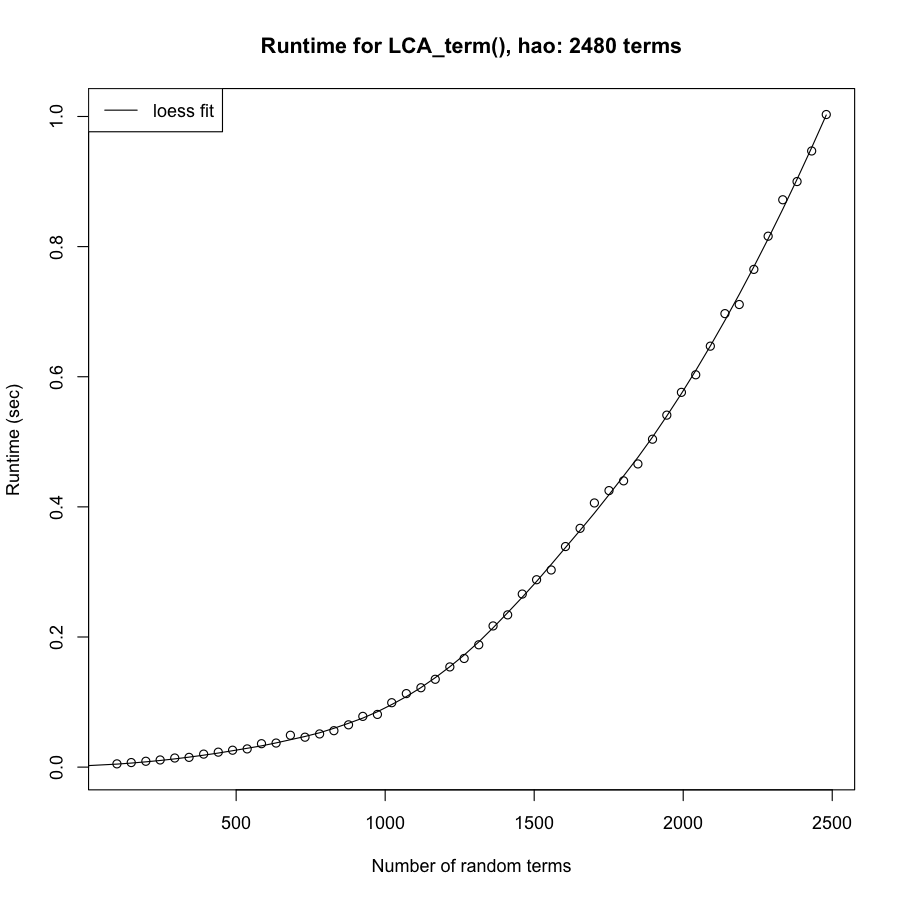

### OBOFoundry_hp_runtime.png

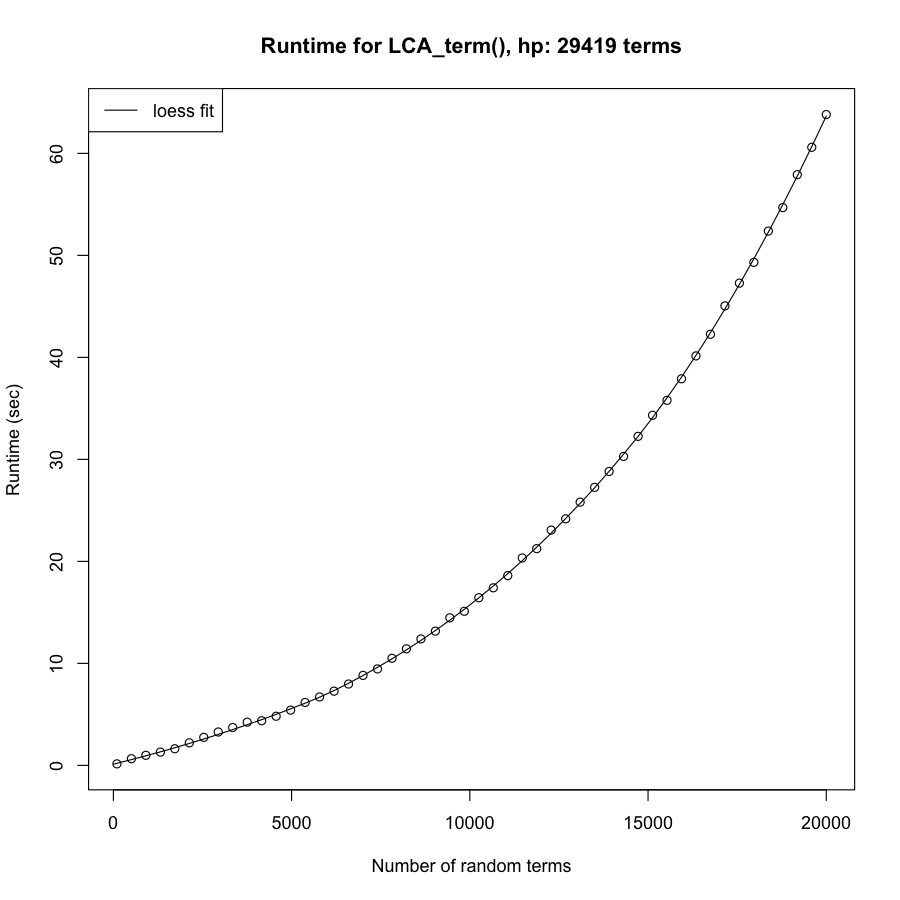

### OBOFoundry_hso.png

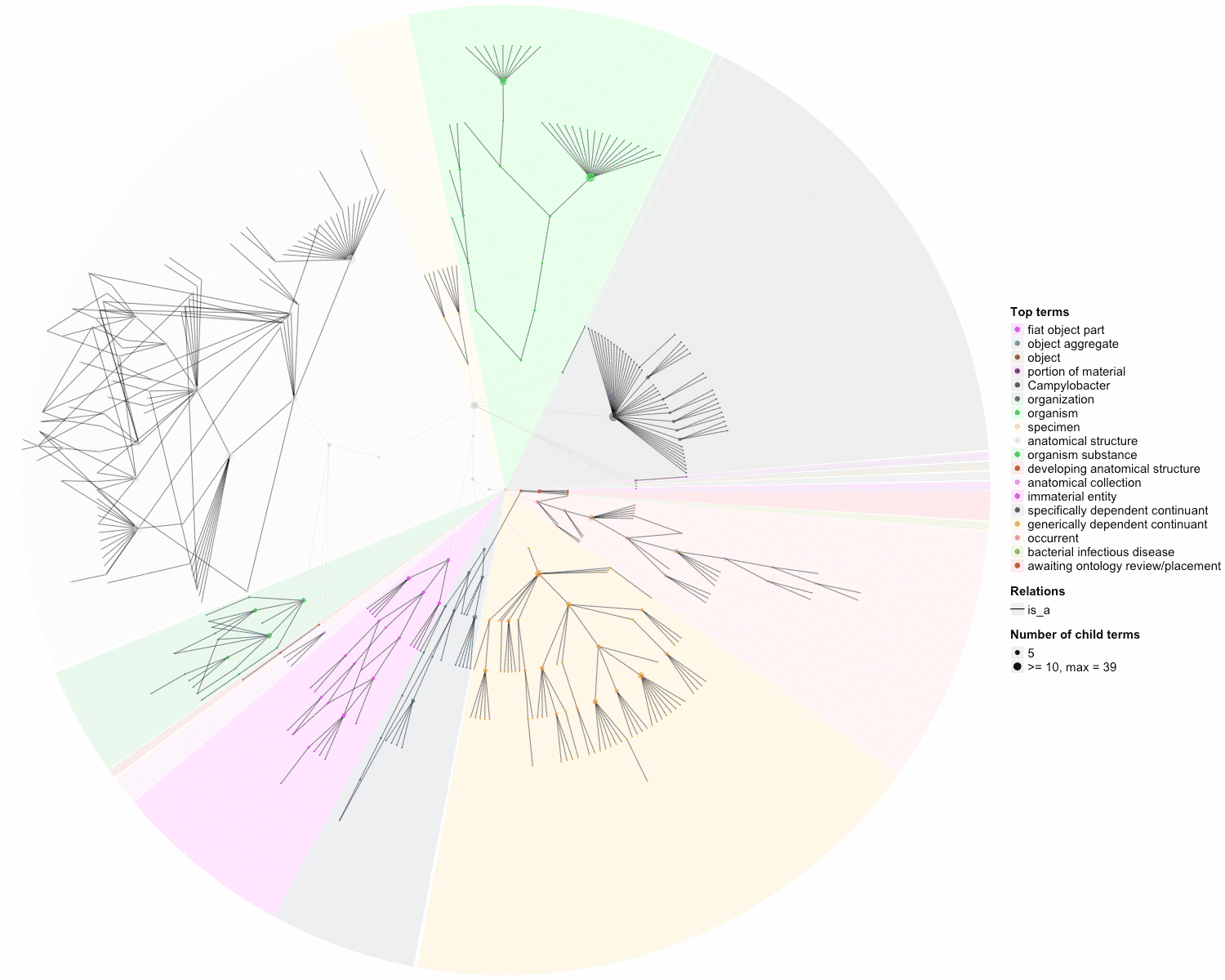

### OBOFoundry_htn_heatmap.png

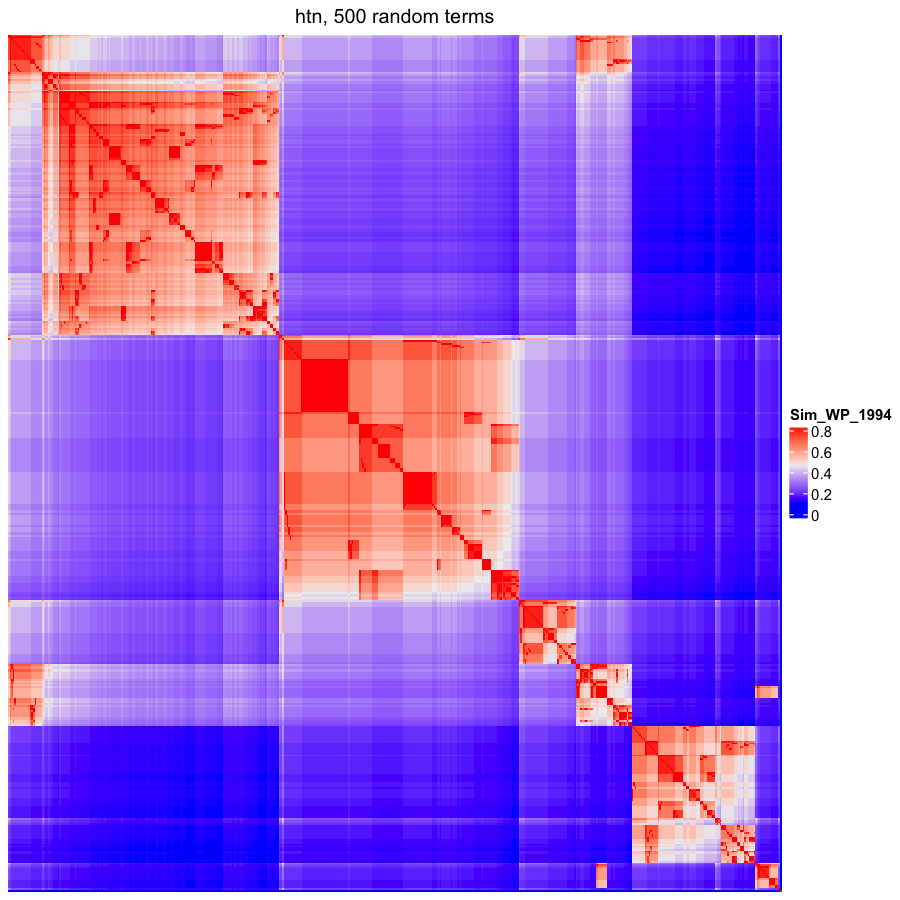

### OBOFoundry_iao.png

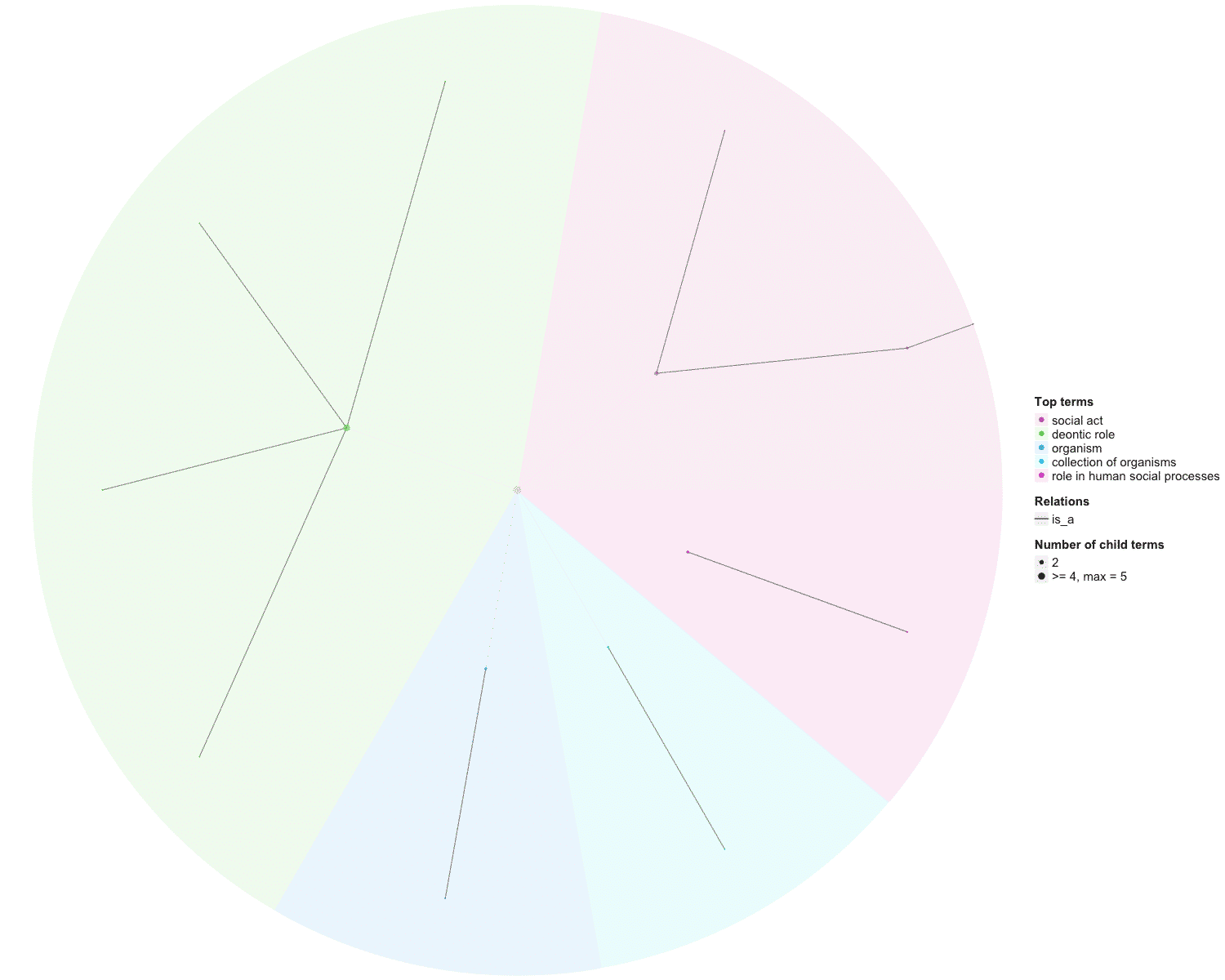

### OBOFoundry_iao_heatmap.png

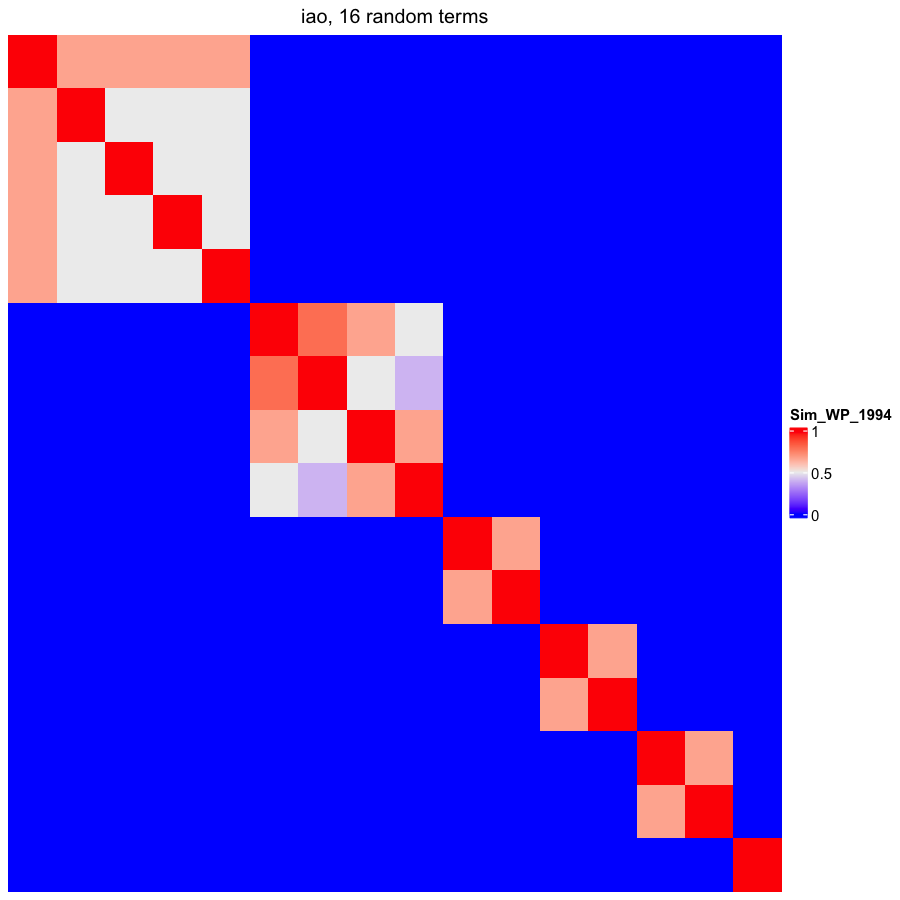

### OBOFoundry_iceo_runtime.png

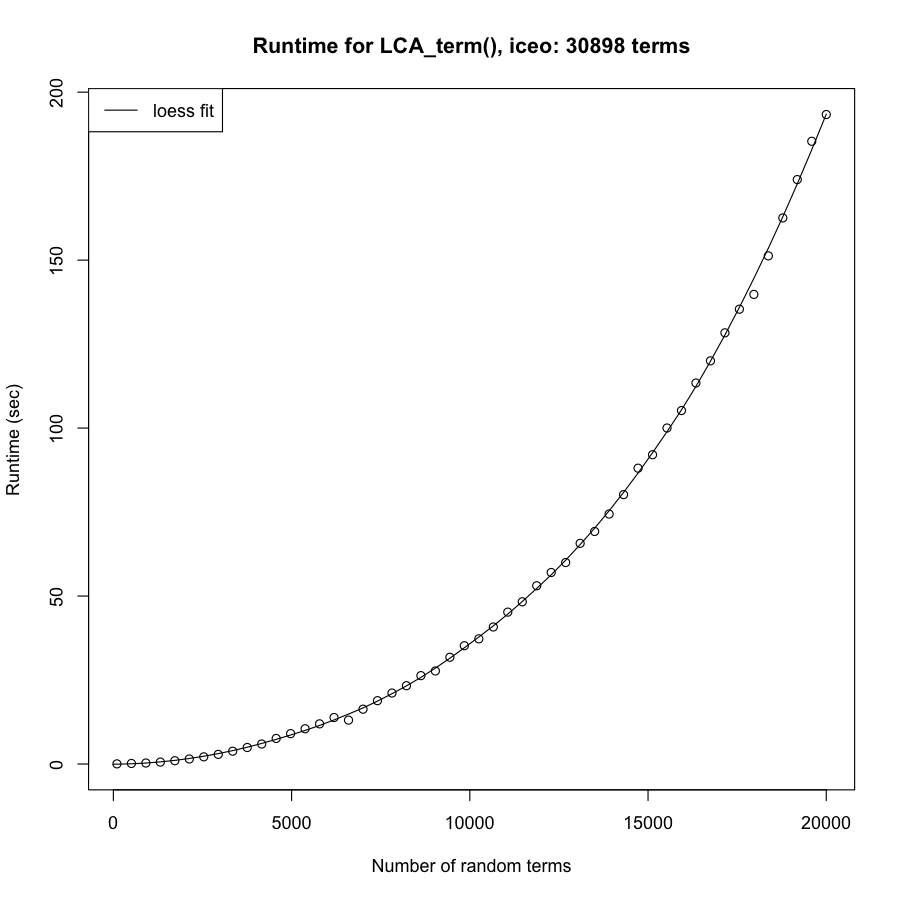

### OBOFoundry_ma_runtime.png

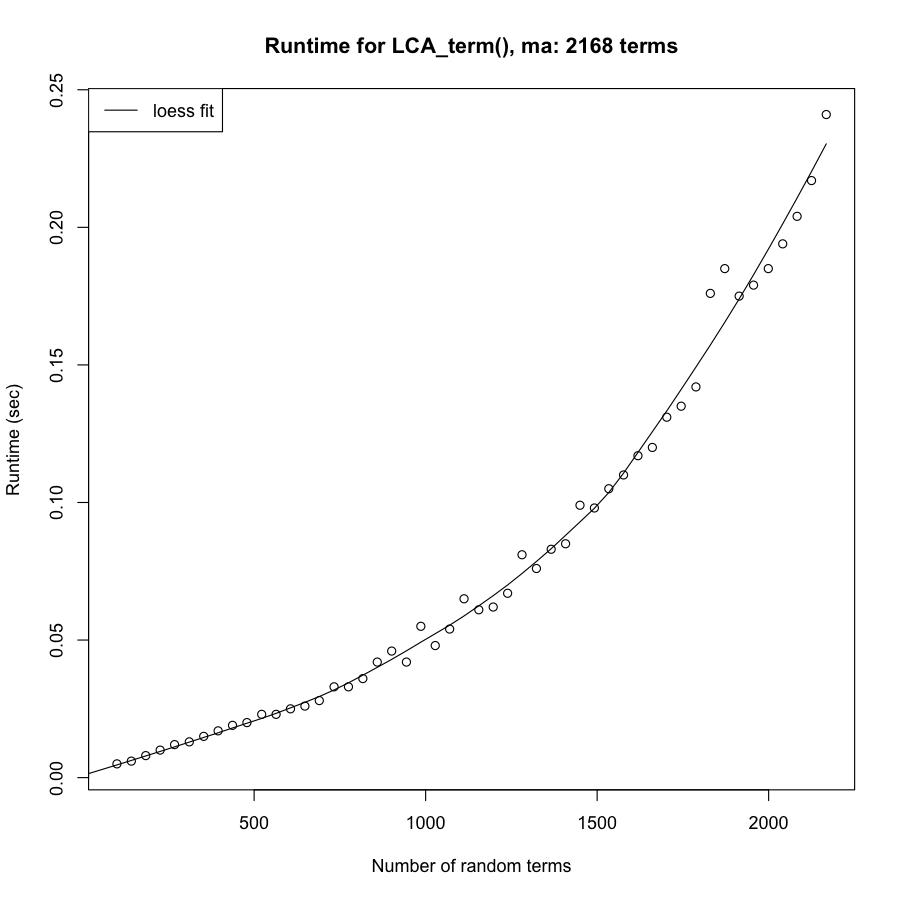

### OBOFoundry_mamo.png

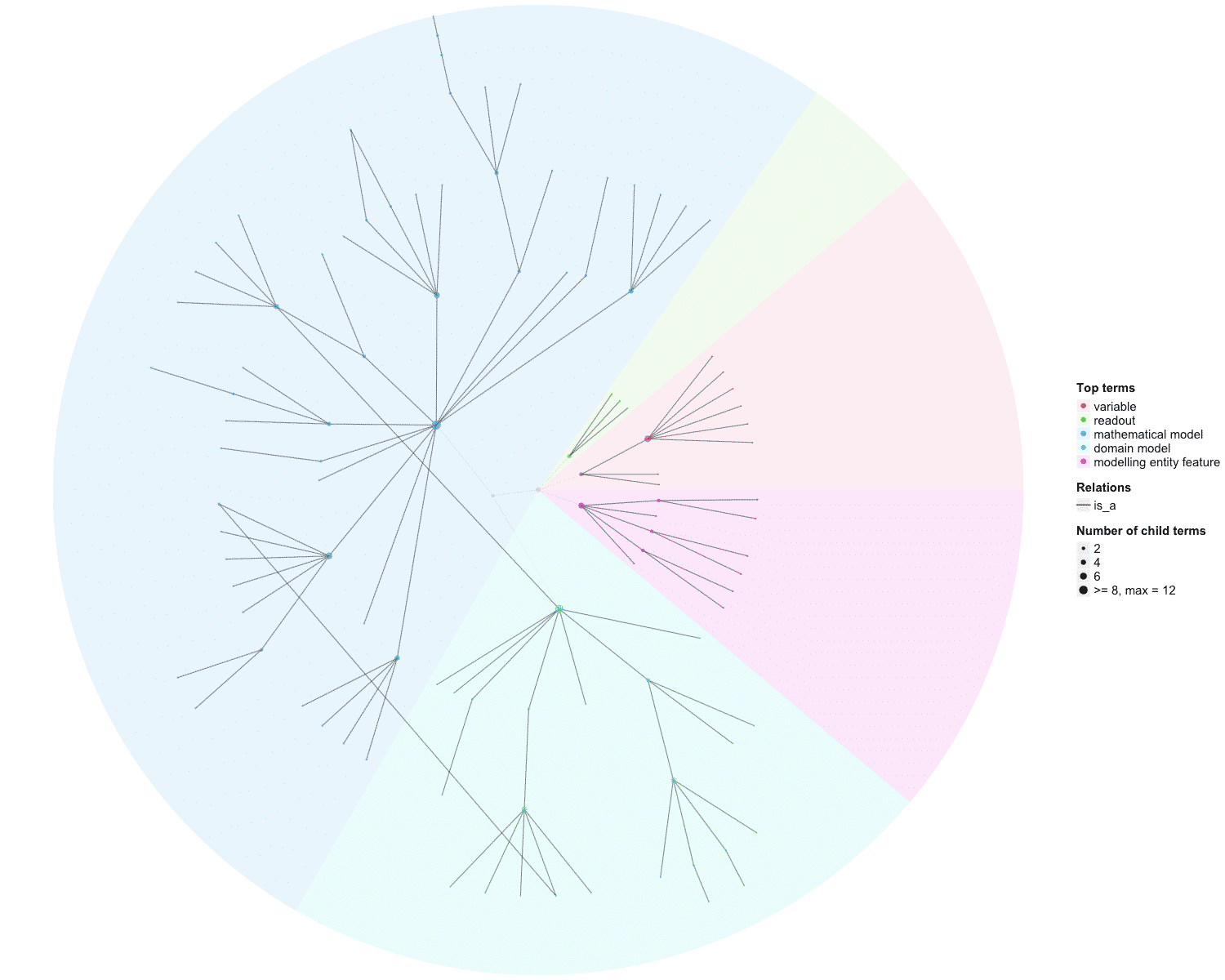

### OBOFoundry_maxo.png

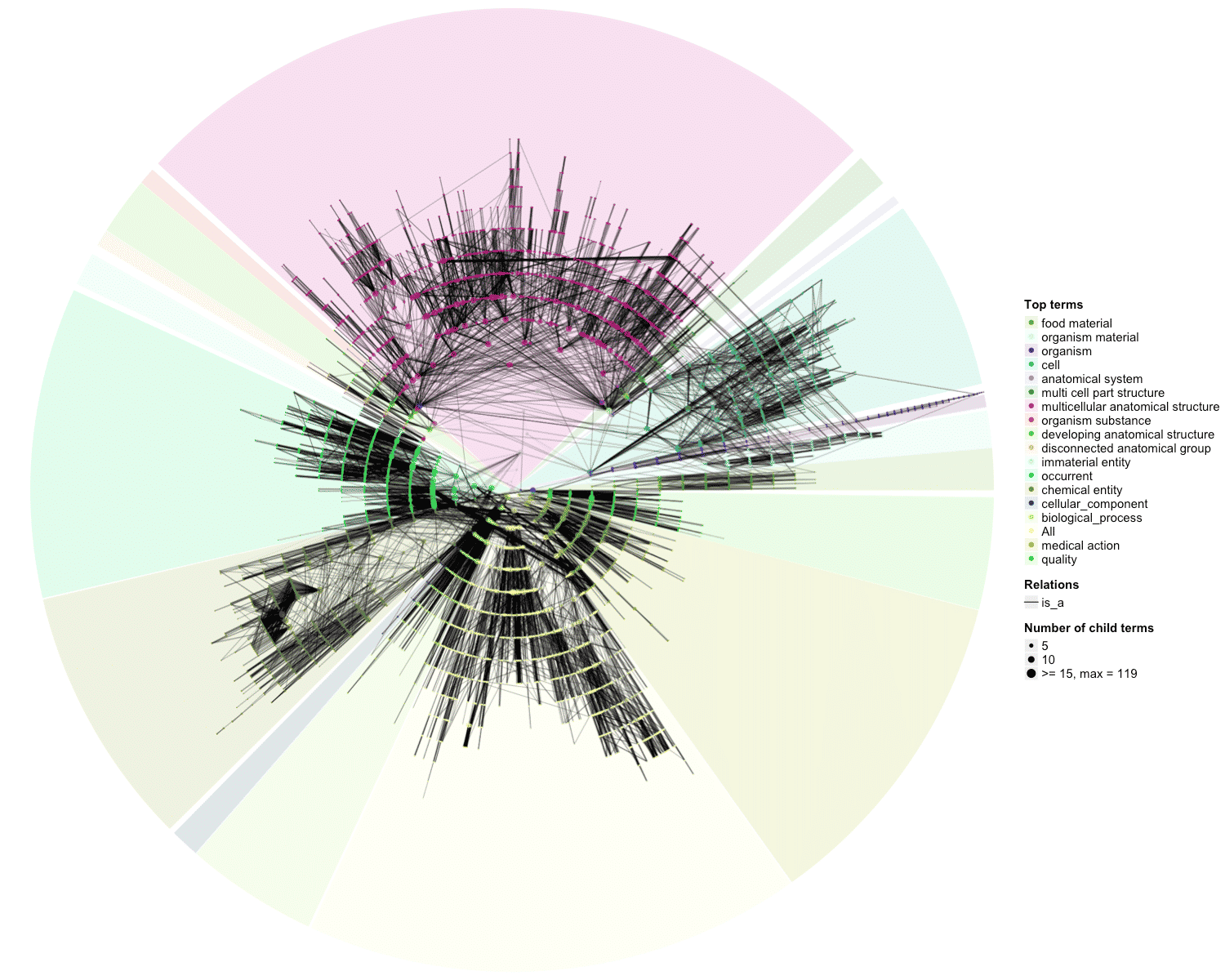

### OBOFoundry_mco_runtime.png

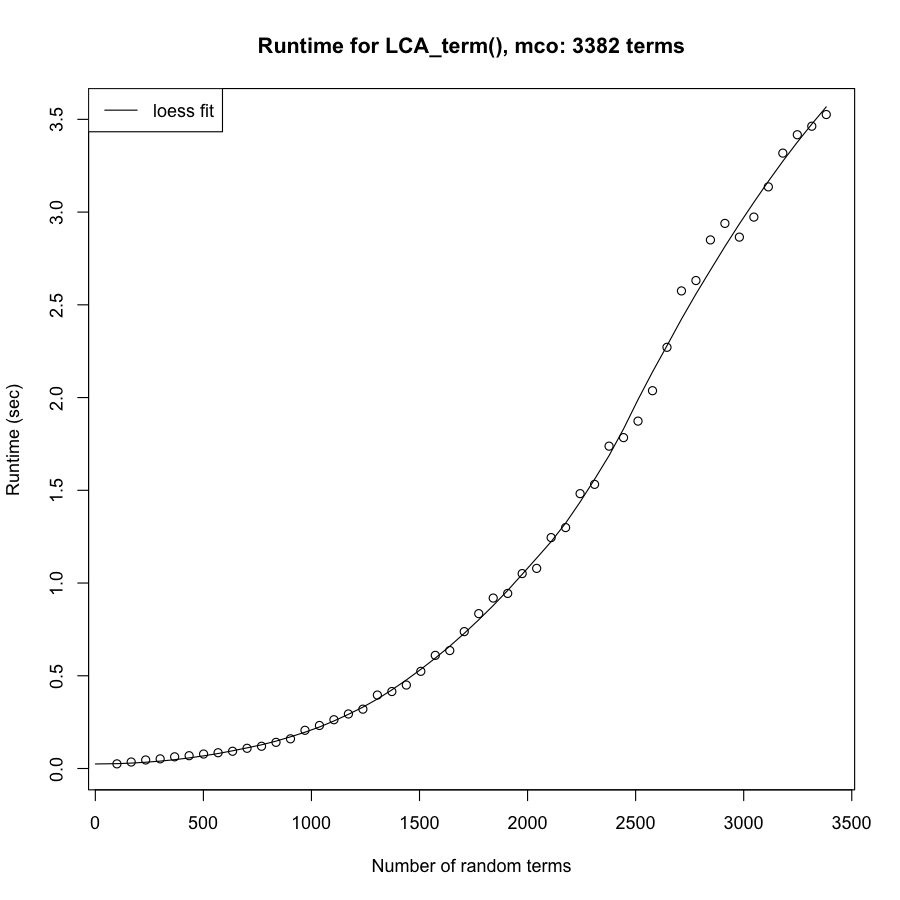

### OBOFoundry_miapa.png

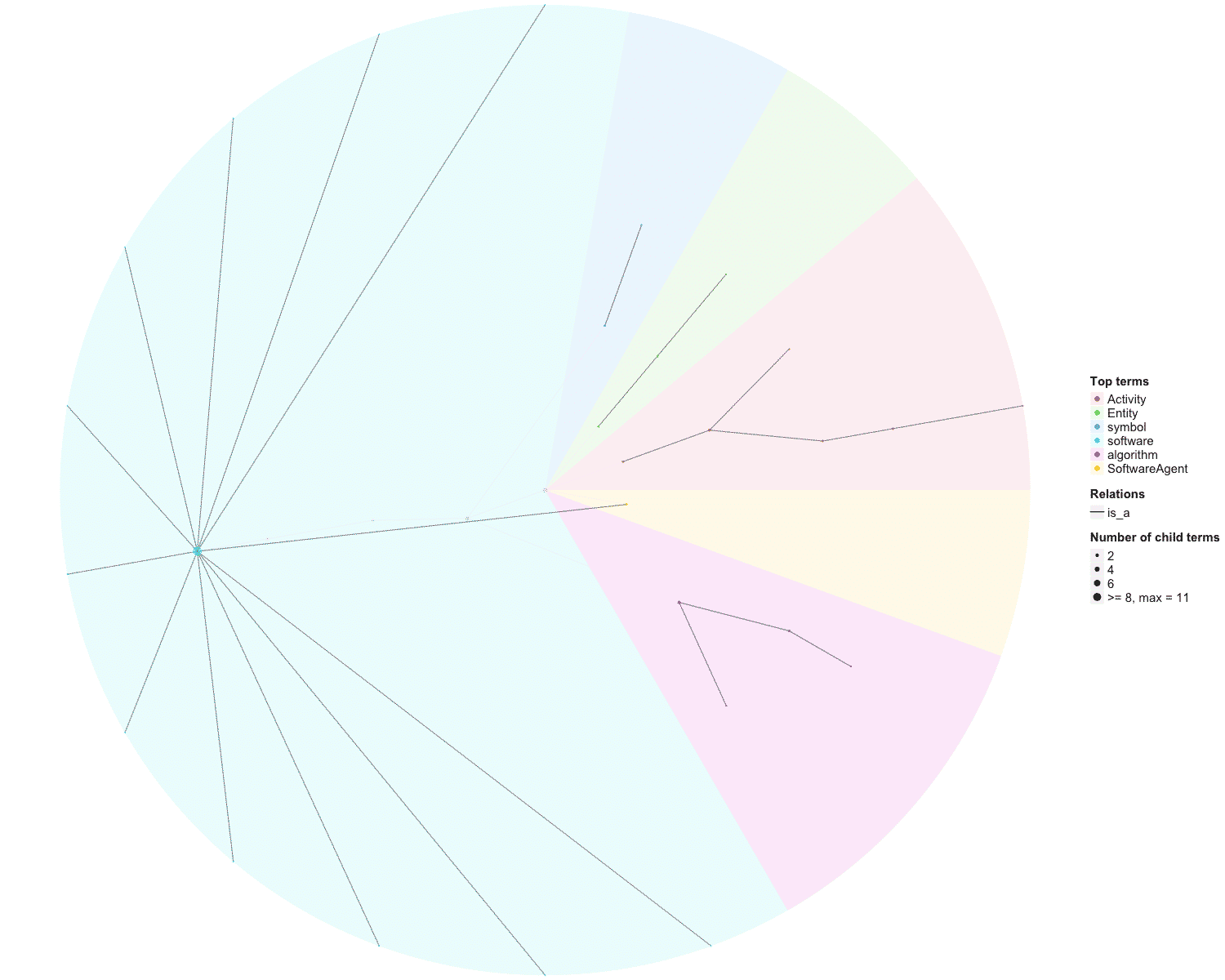

### OBOFoundry_micro.png

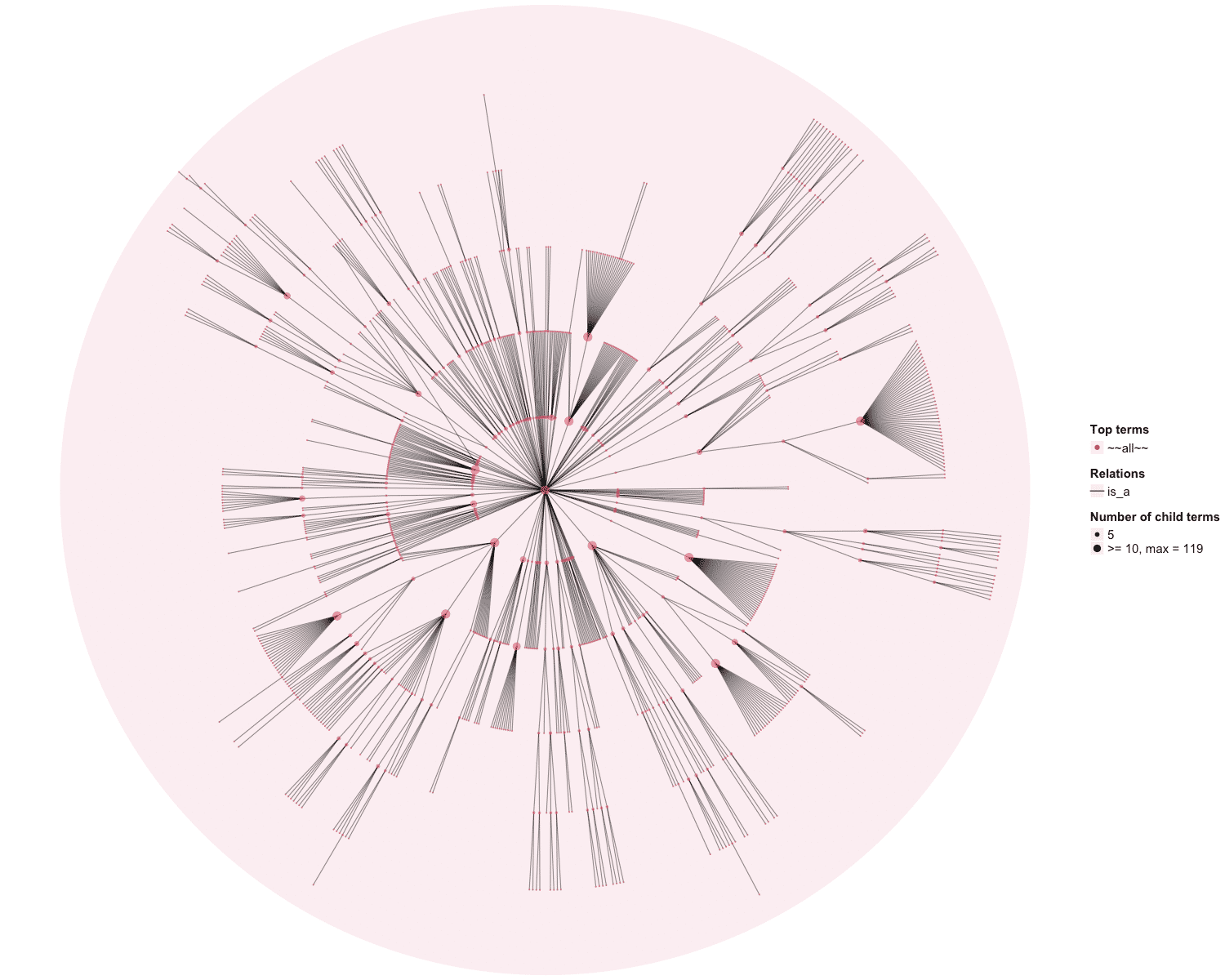

### OBOFoundry_mod.png

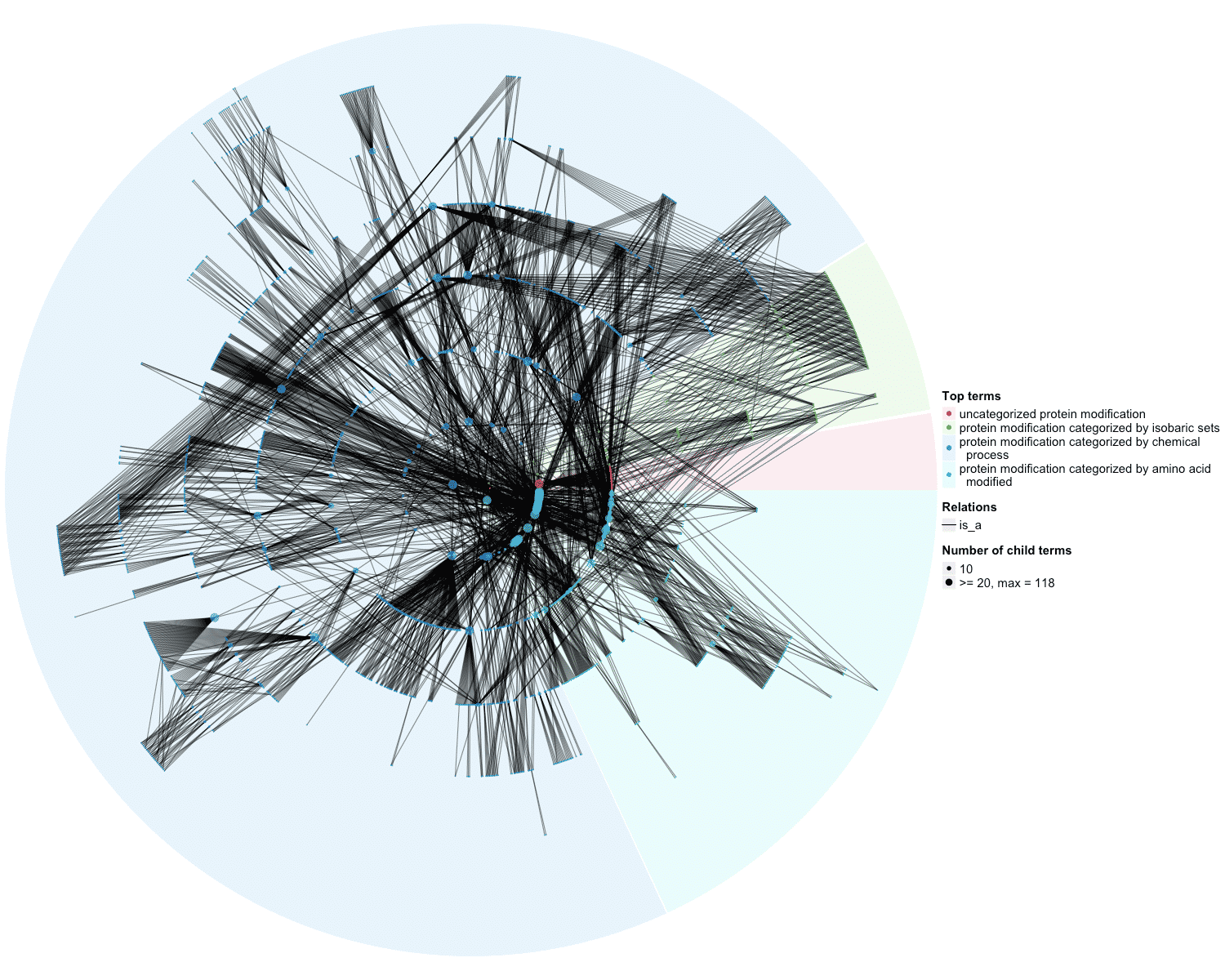

### OBOFoundry_mondo.png

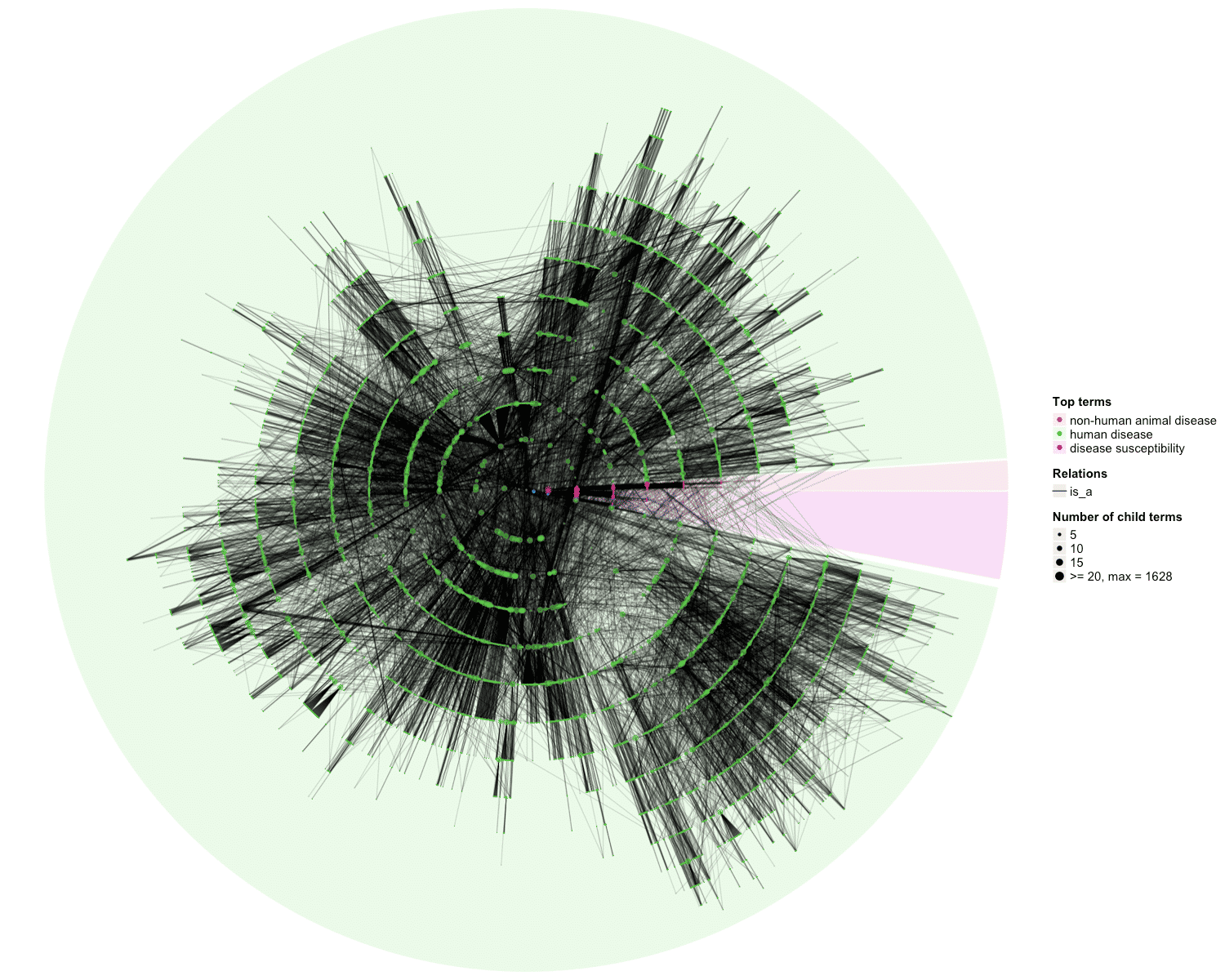

### OBOFoundry_mp_heatmap.png

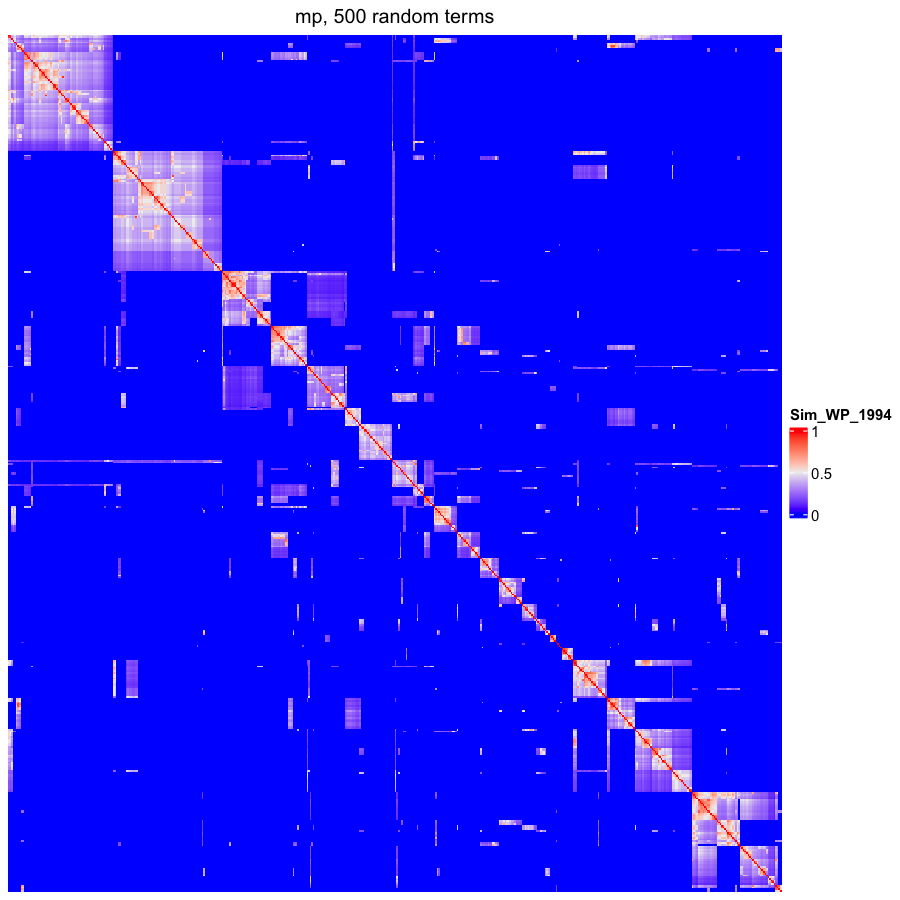

### OBOFoundry_mpio_heatmap.png

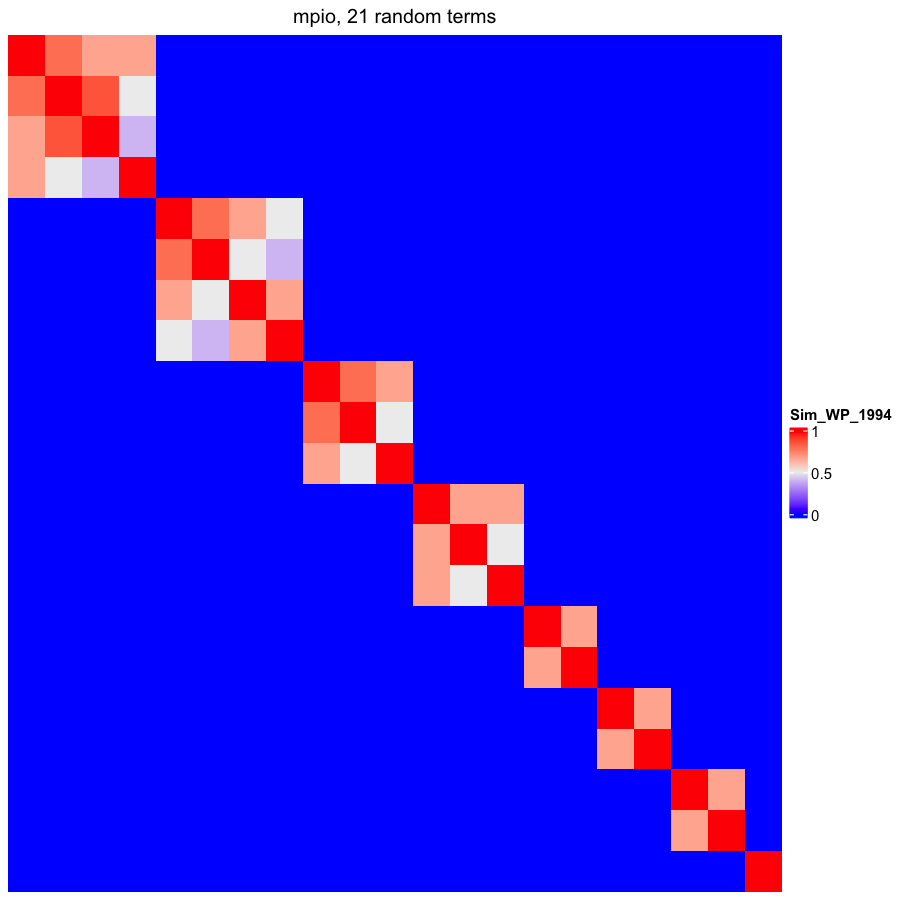

### OBOFoundry_nbo_runtime.png

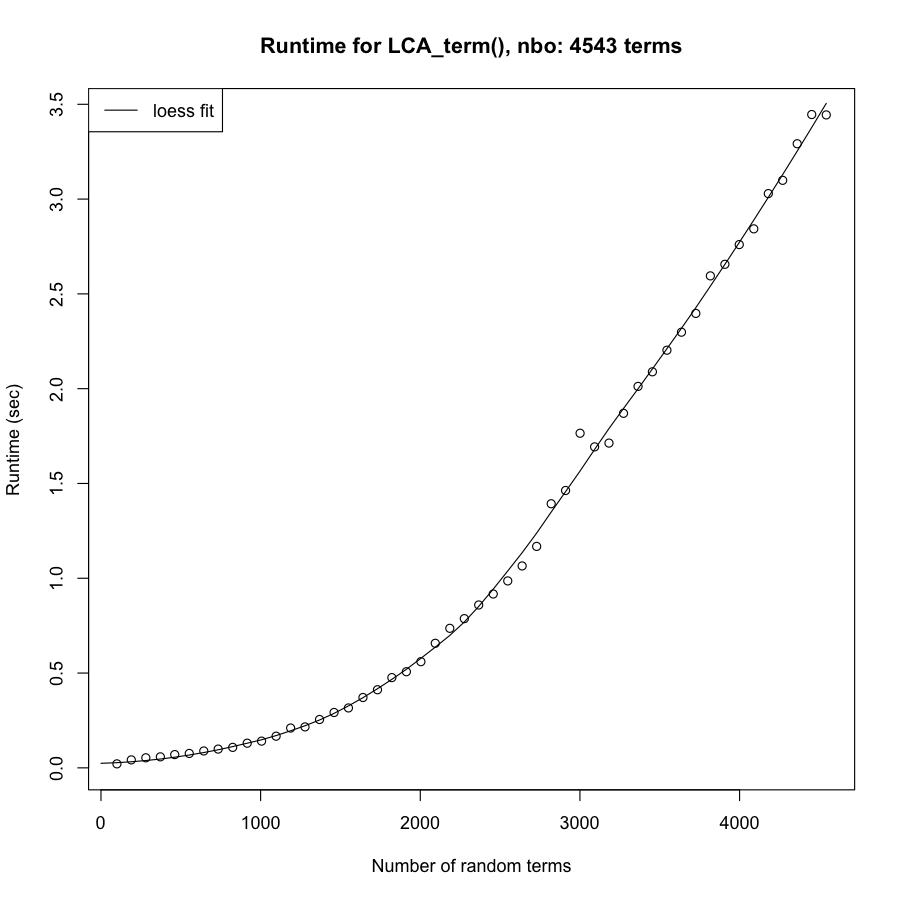

### OBOFoundry_ncbitaxon.png

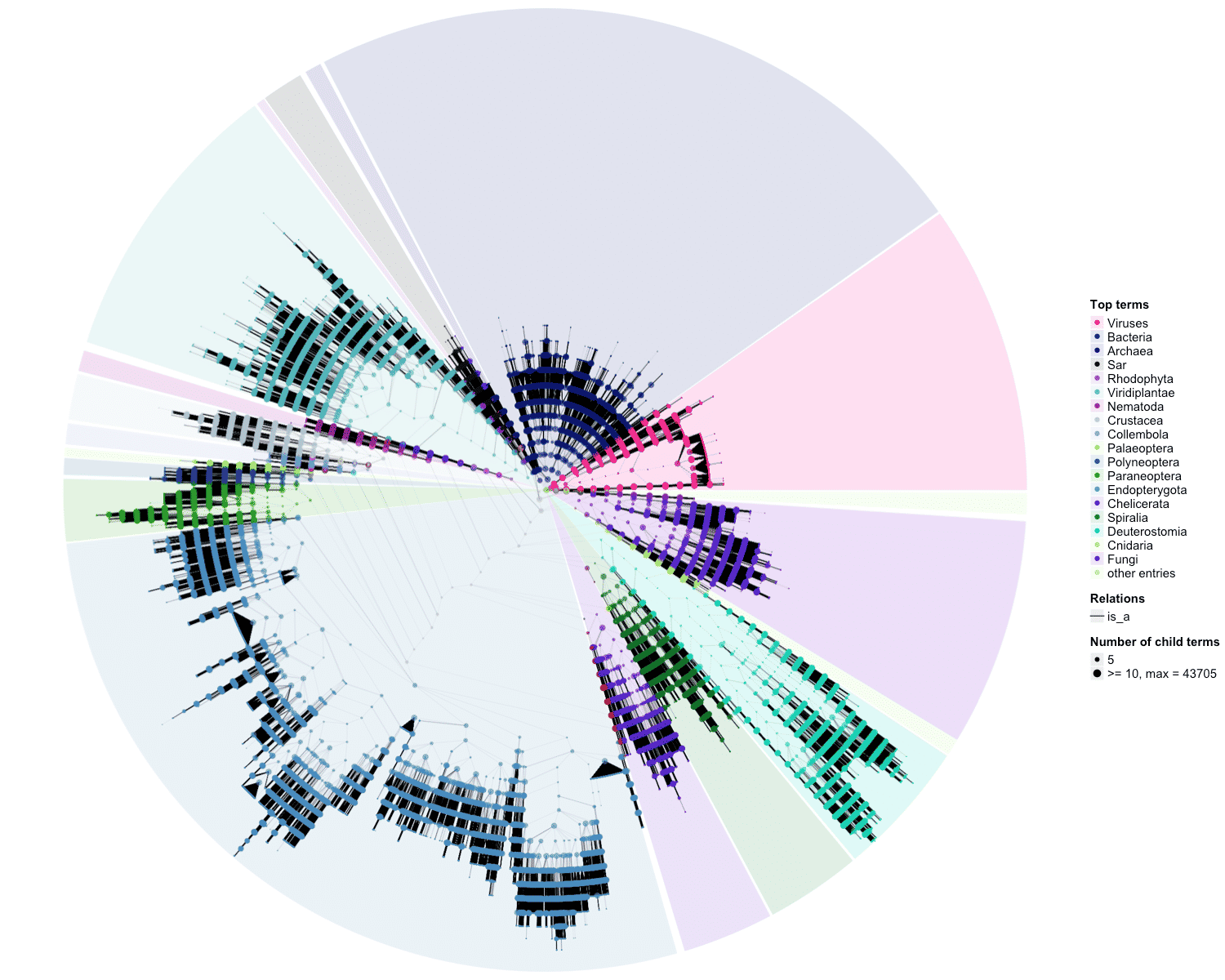

### OBOFoundry_ncit_heatmap.png

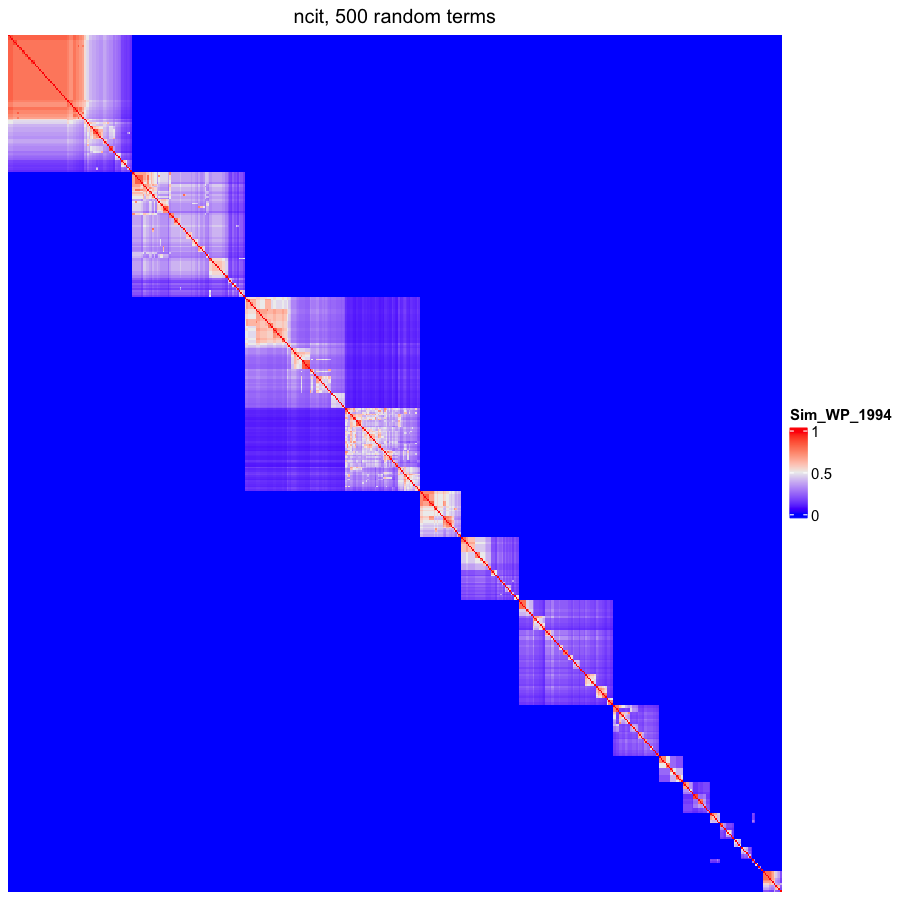

### OBOFoundry_ngbo_runtime.png

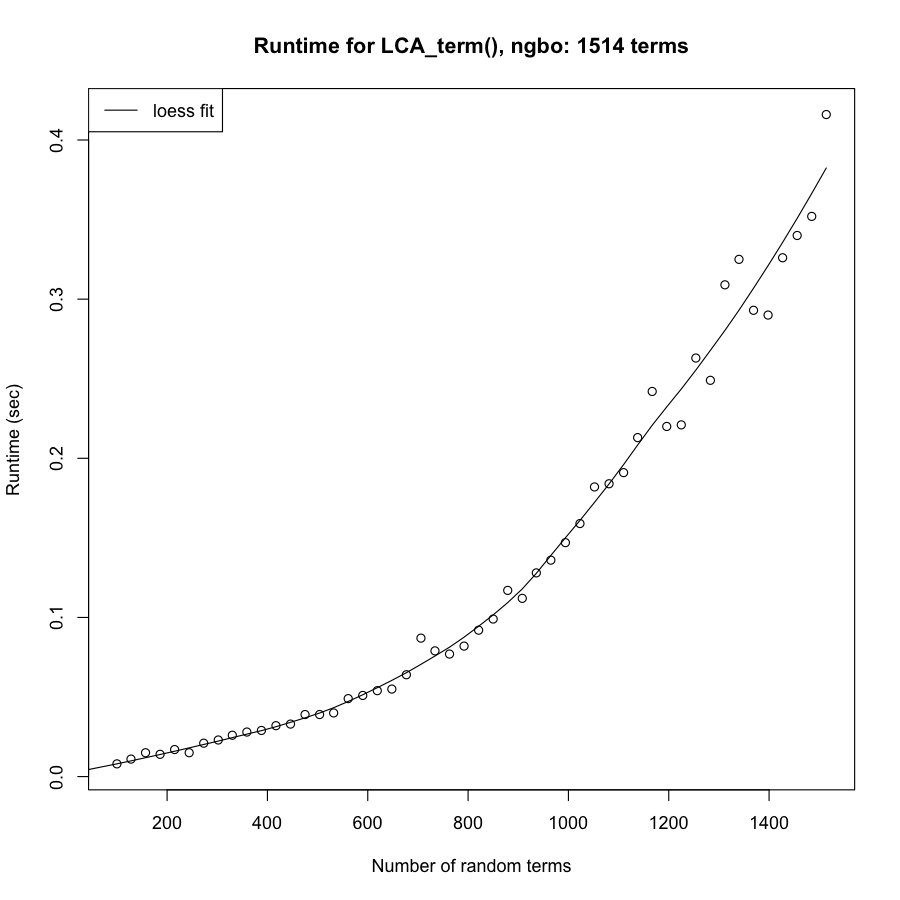

### OBOFoundry_nomen.png

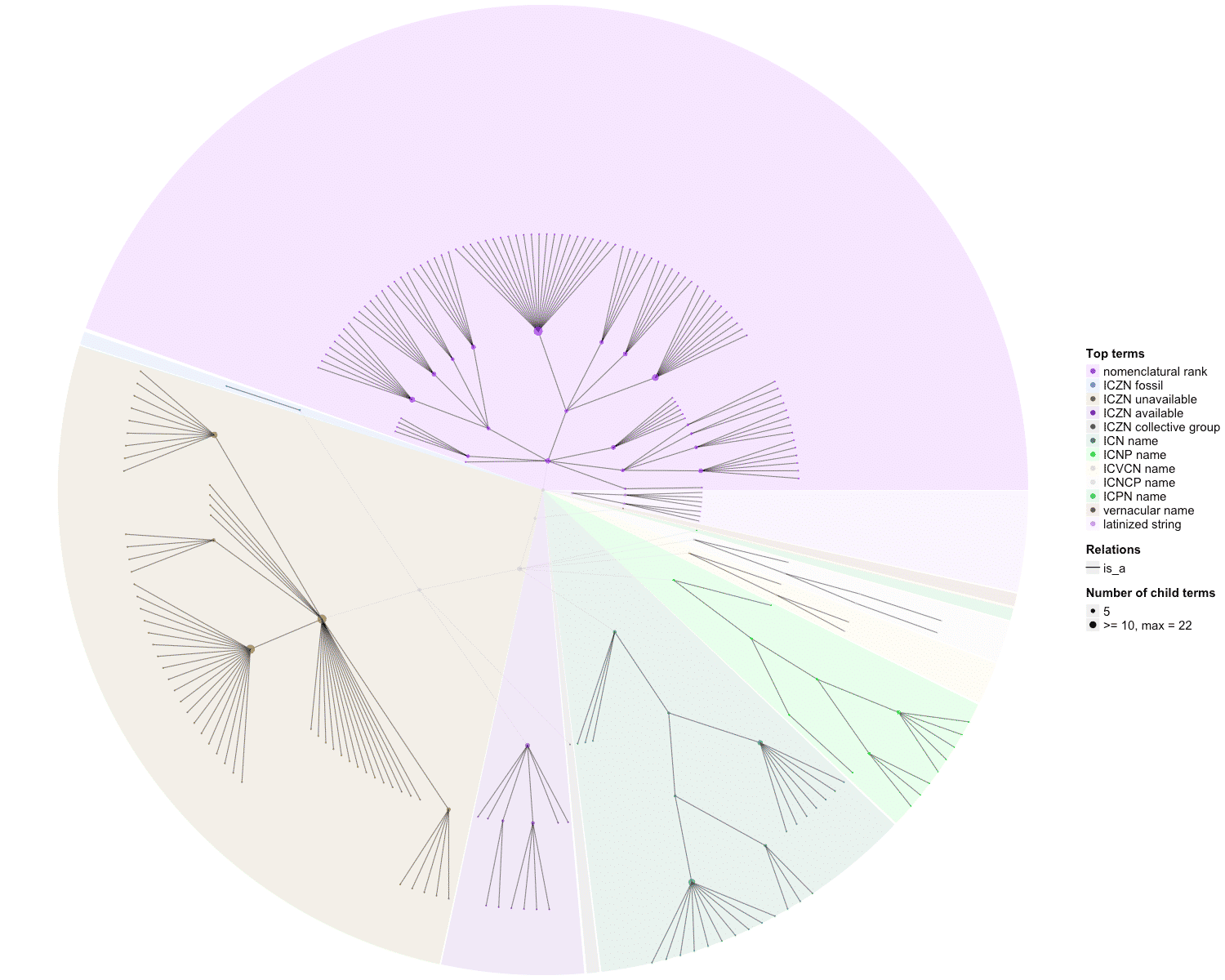

### OBOFoundry_oba.png

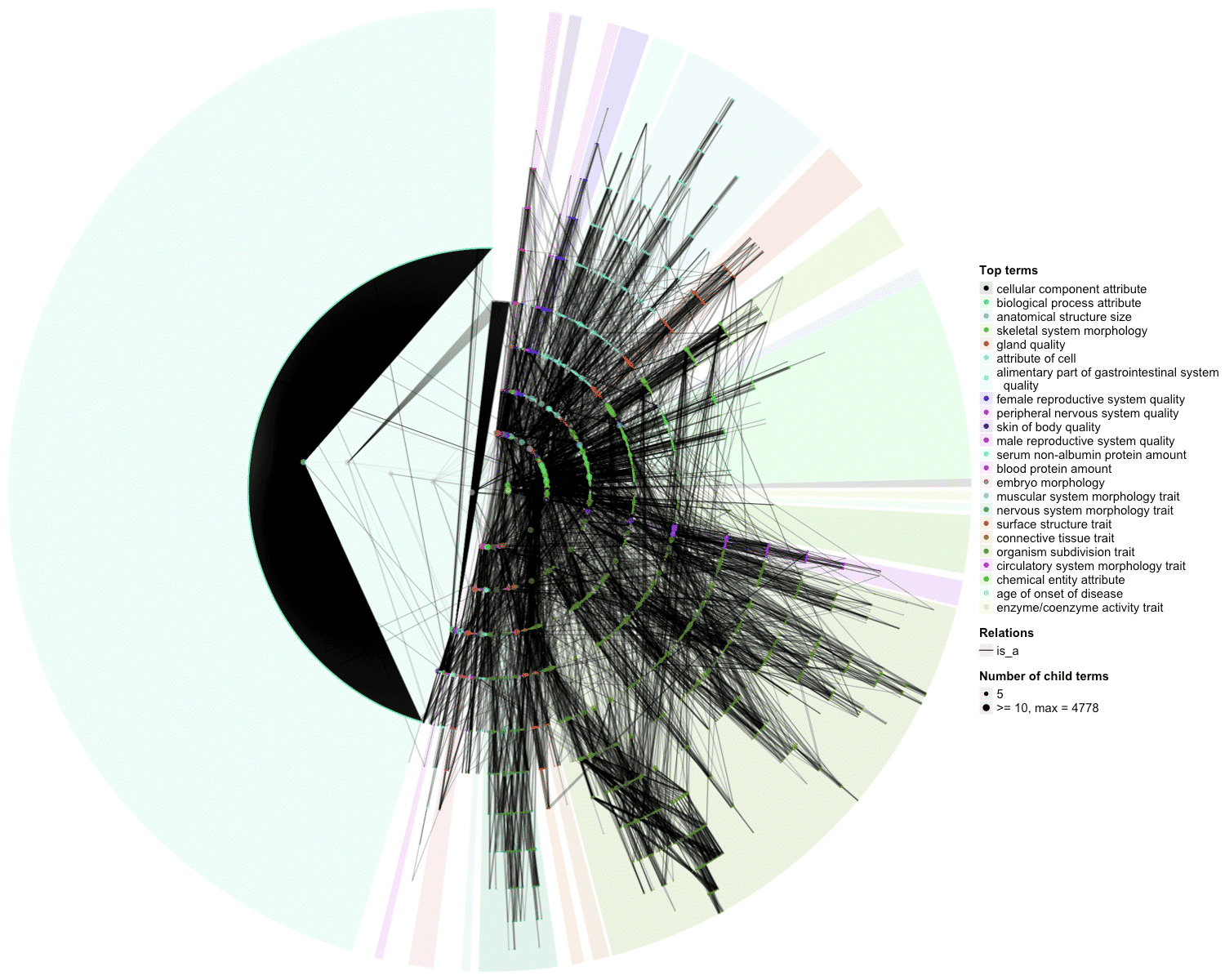

### OBOFoundry_oba_runtime.png

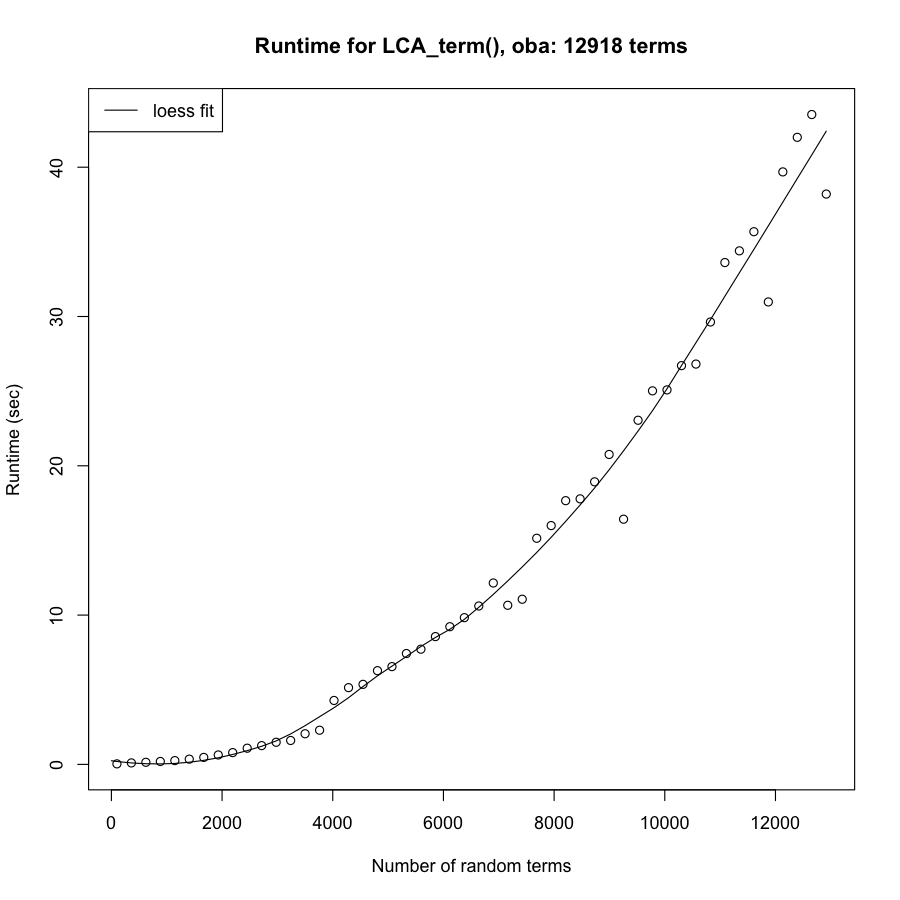

### OBOFoundry_obib_runtime.png

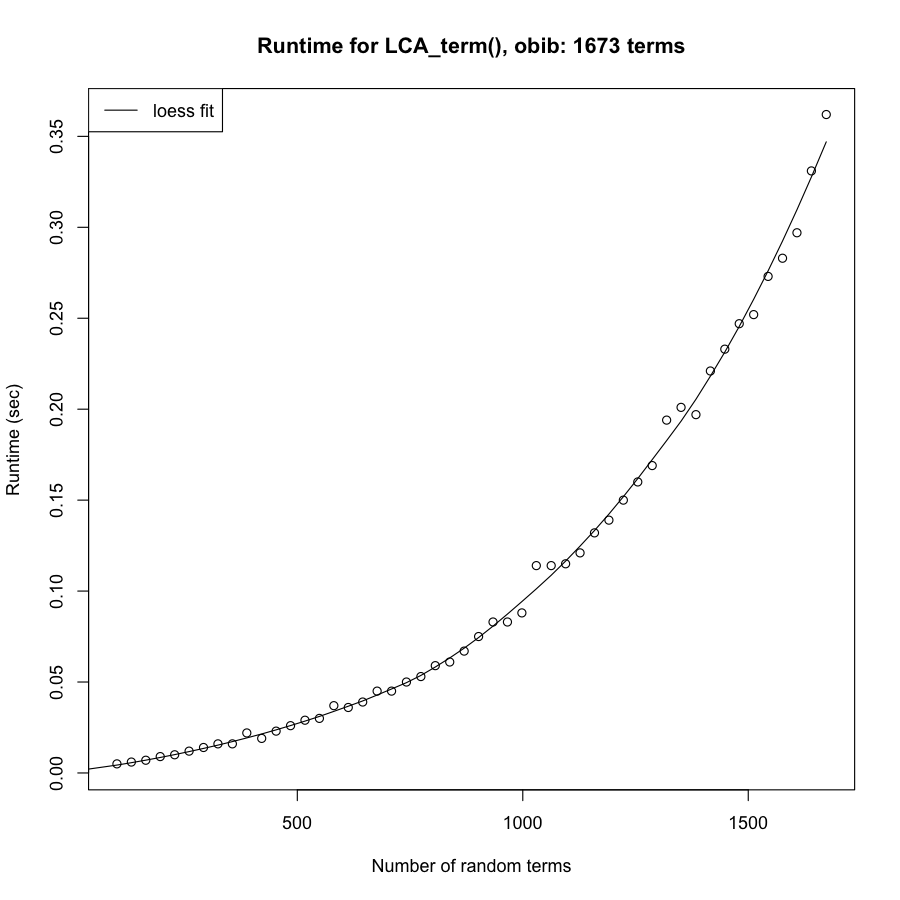

### OBOFoundry_ogms_heatmap.png

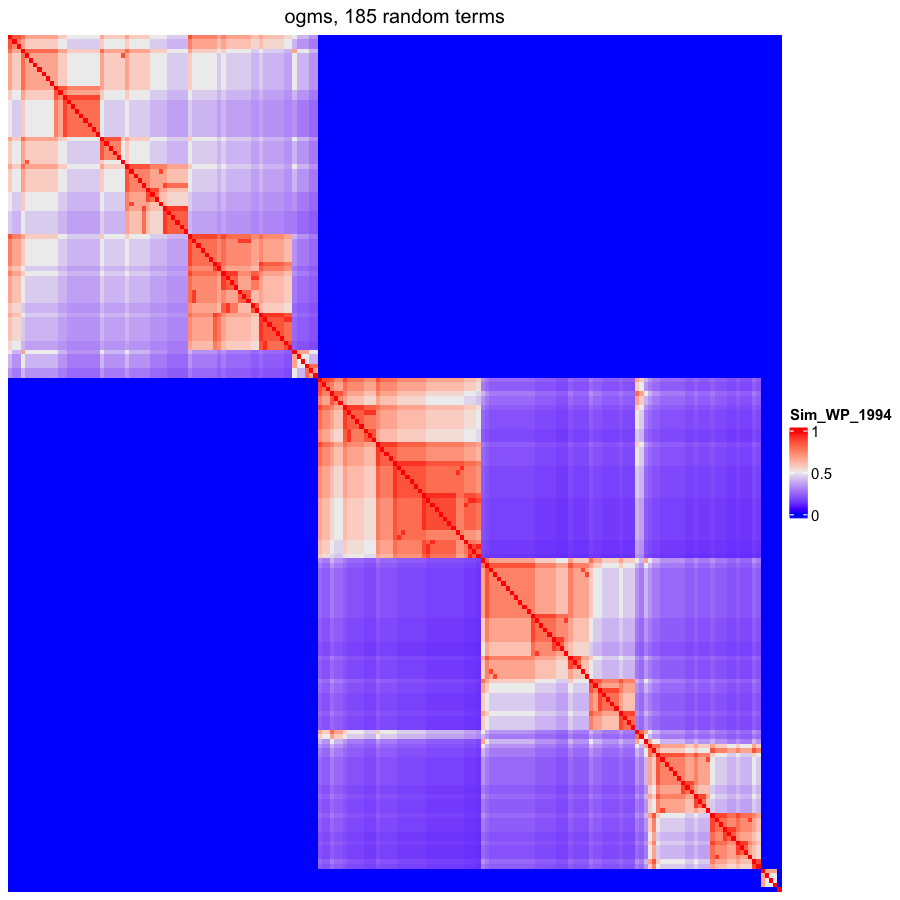

### OBOFoundry_ohpi.png

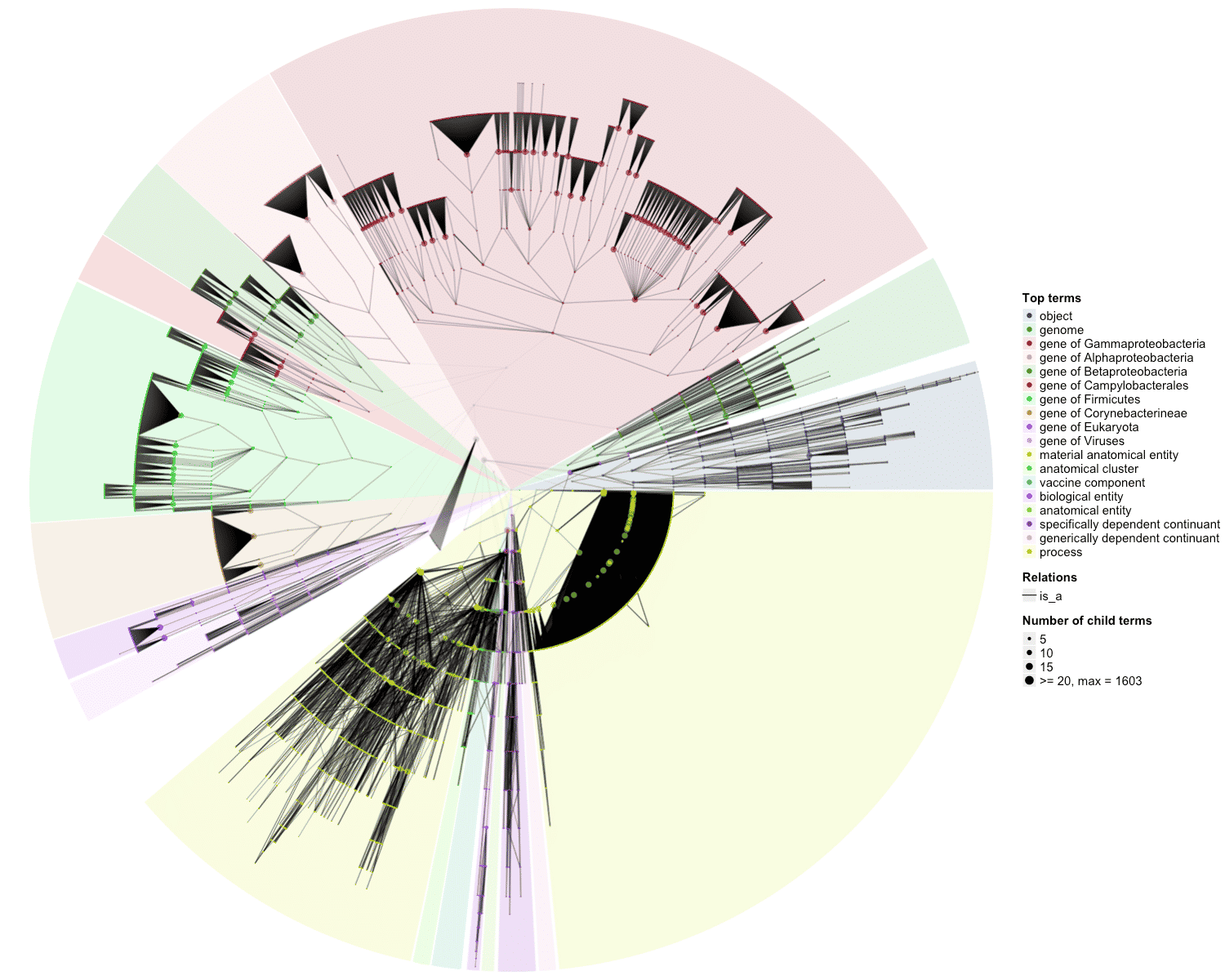

### OBOFoundry_omo.png

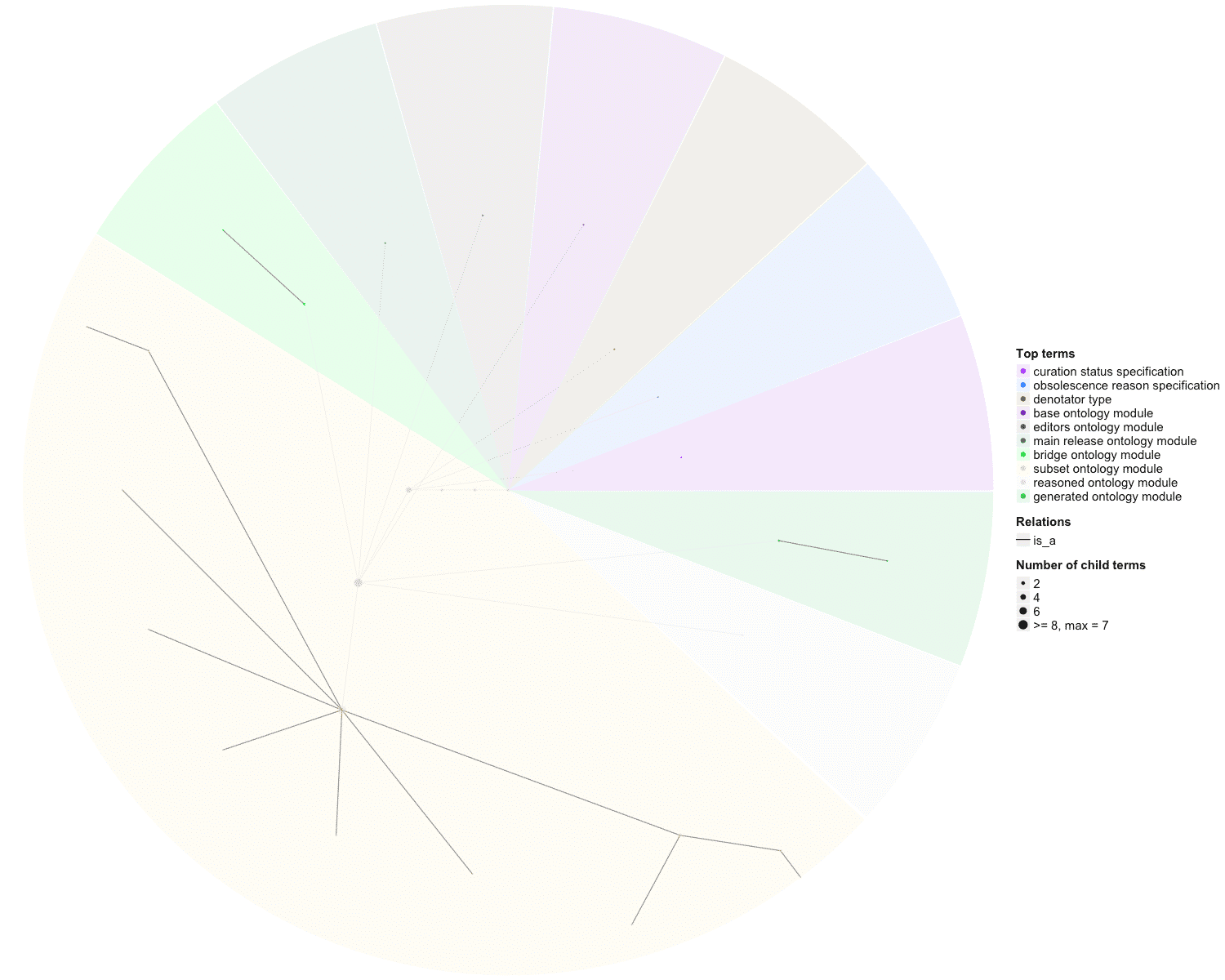

### OBOFoundry_omrse.png

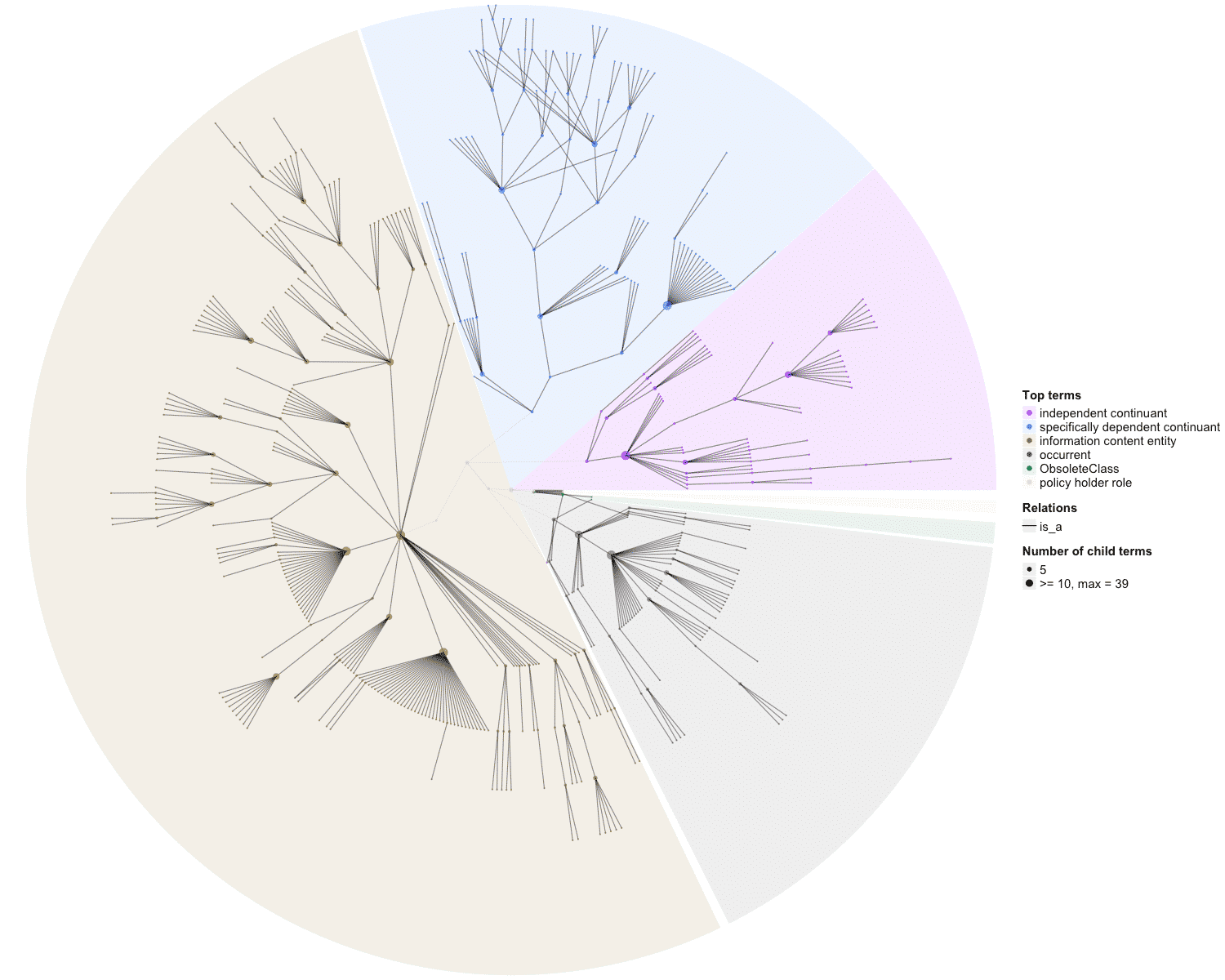

### OBOFoundry_one.png

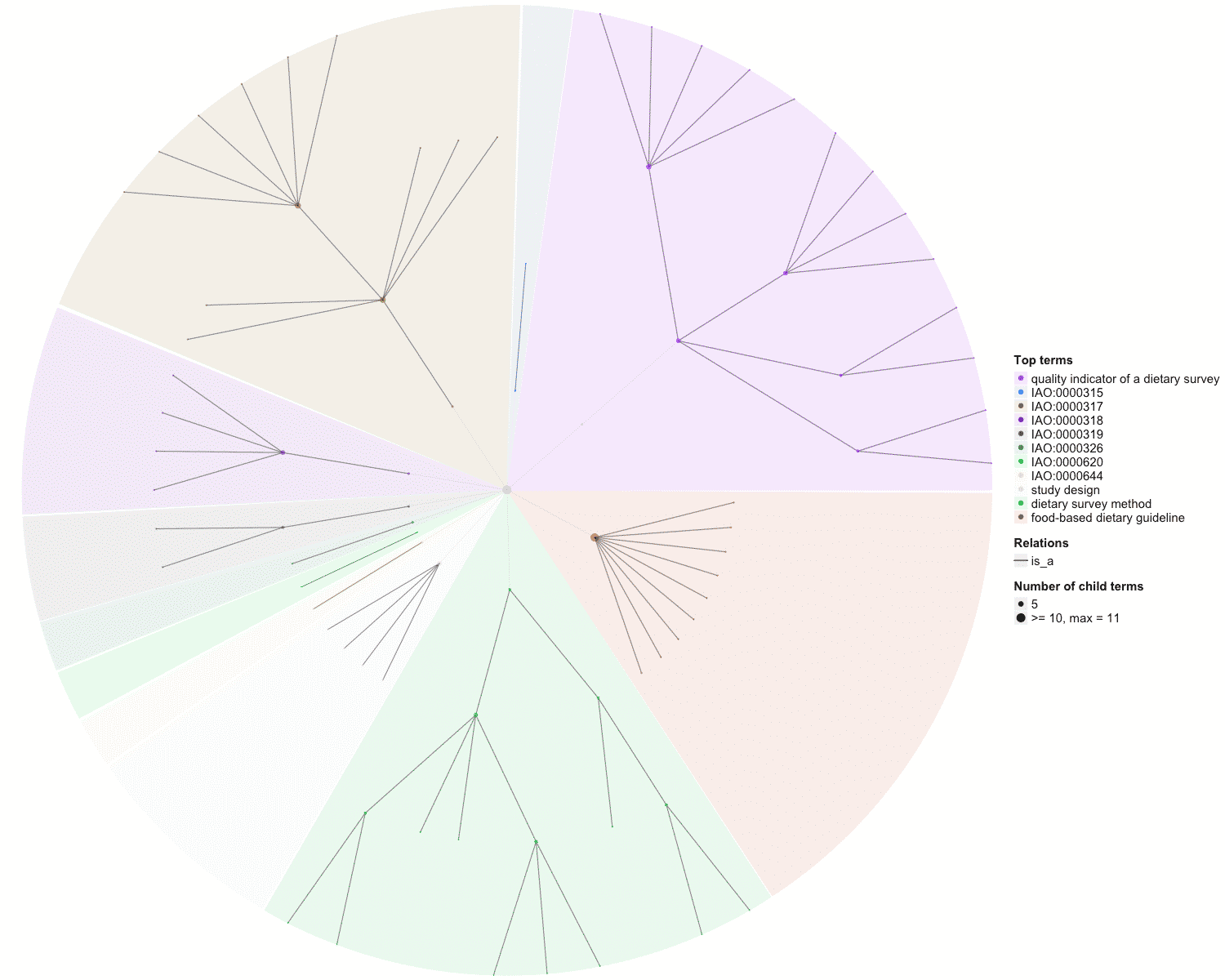

### OBOFoundry_ons.png

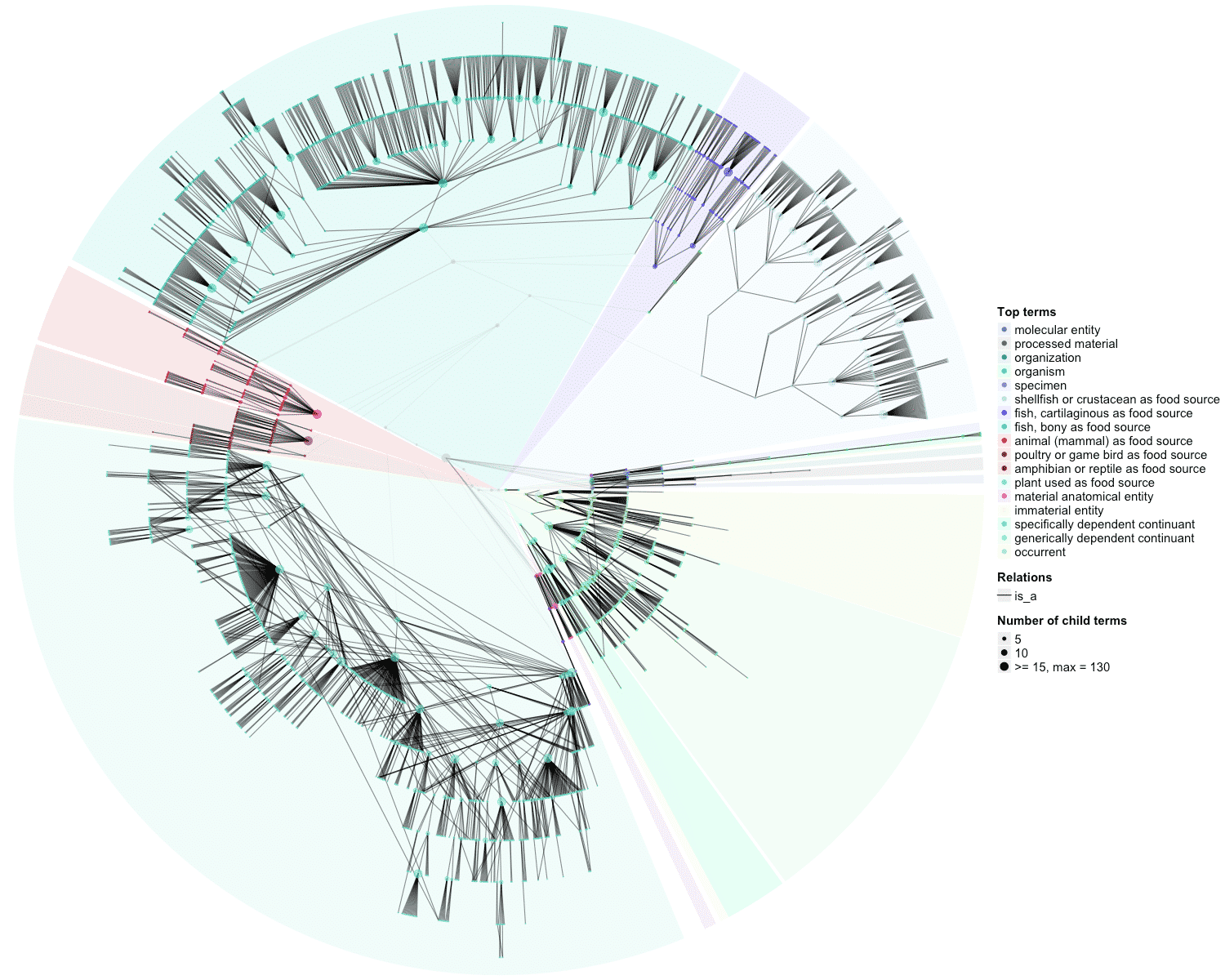

### OBOFoundry_ons_heatmap.png

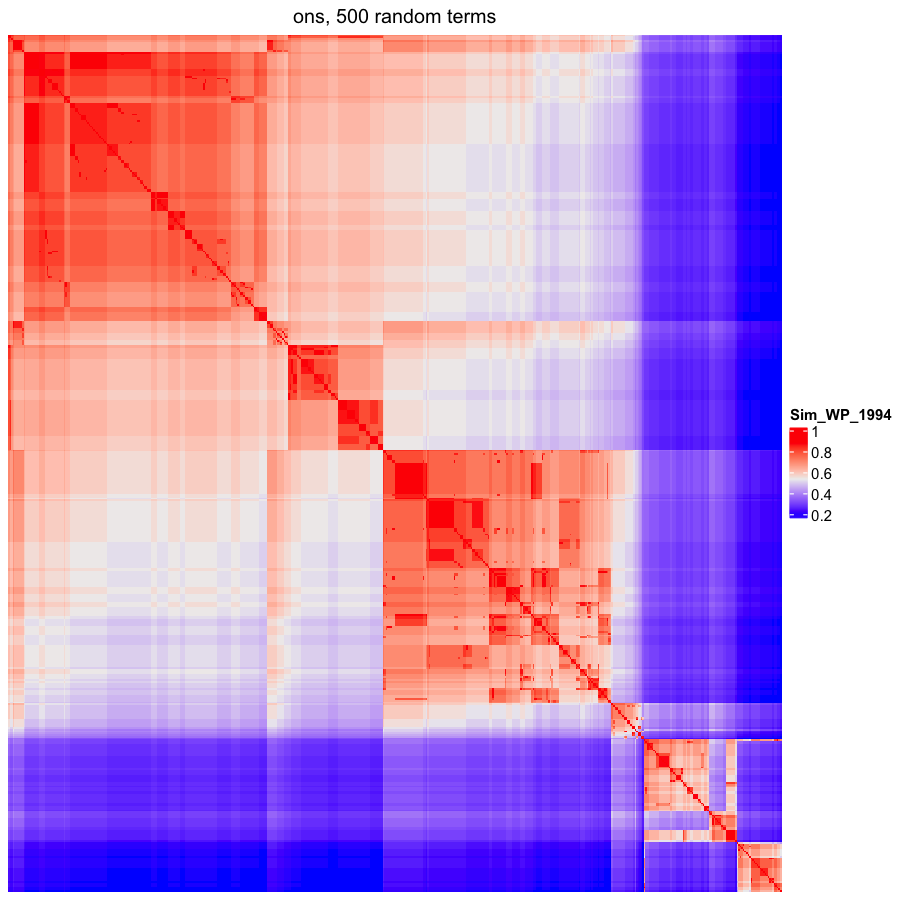

### OBOFoundry_ontoneo.png

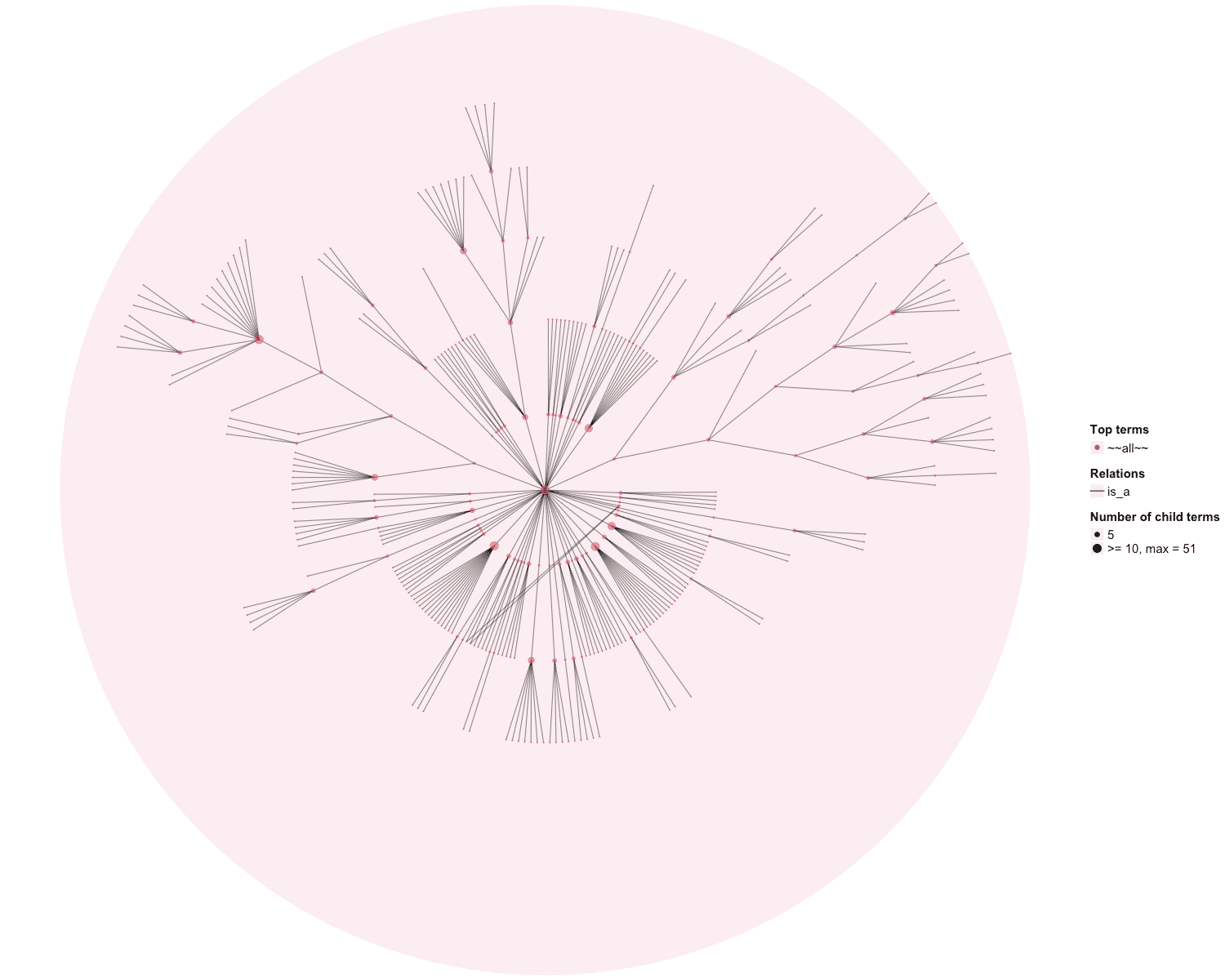

### OBOFoundry_ontoneo_heatmap.png

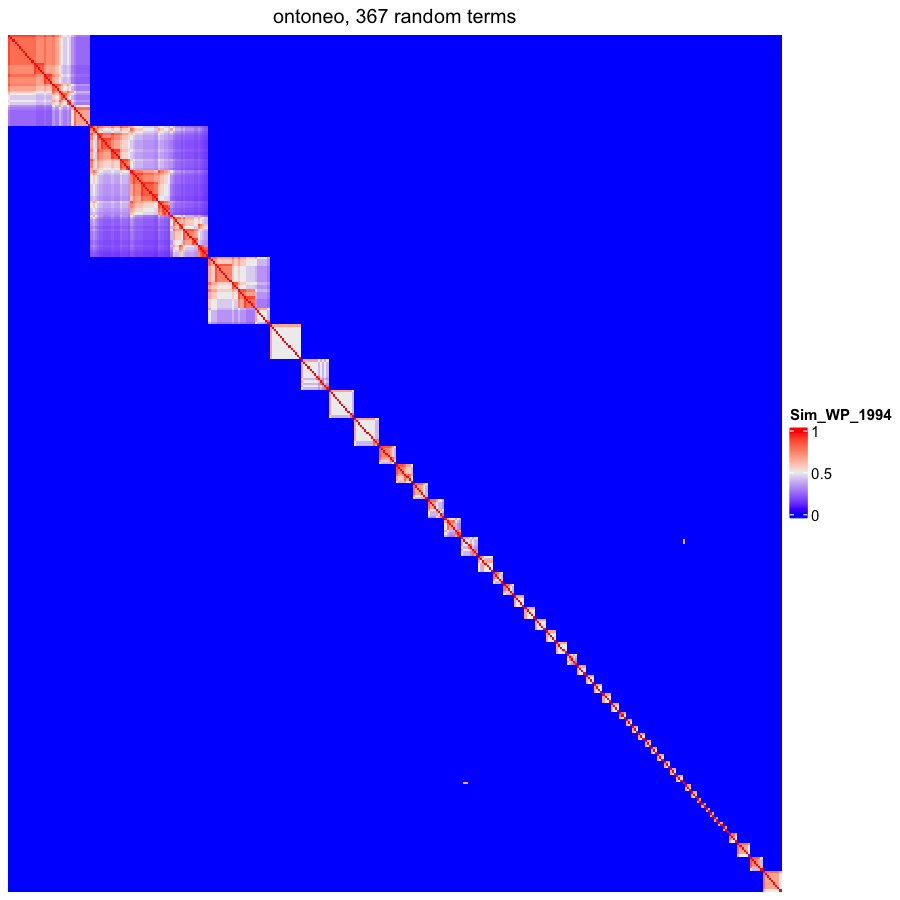

### OBOFoundry_opl_heatmap.png

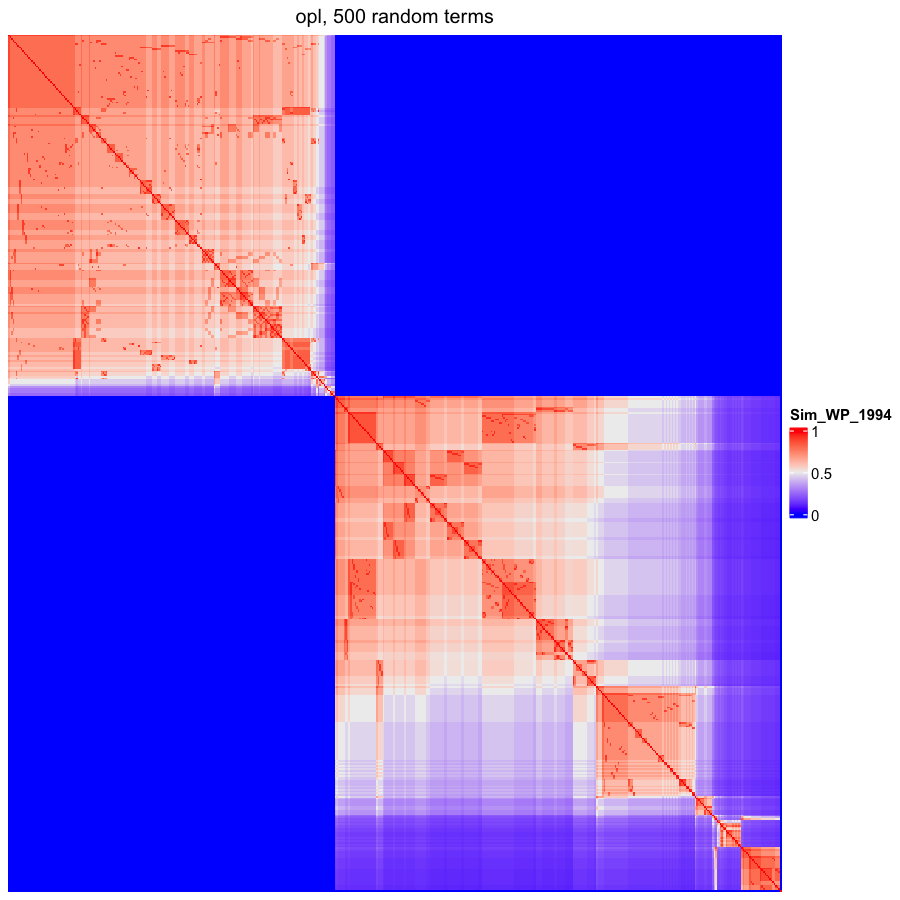

### OBOFoundry_ovae.png

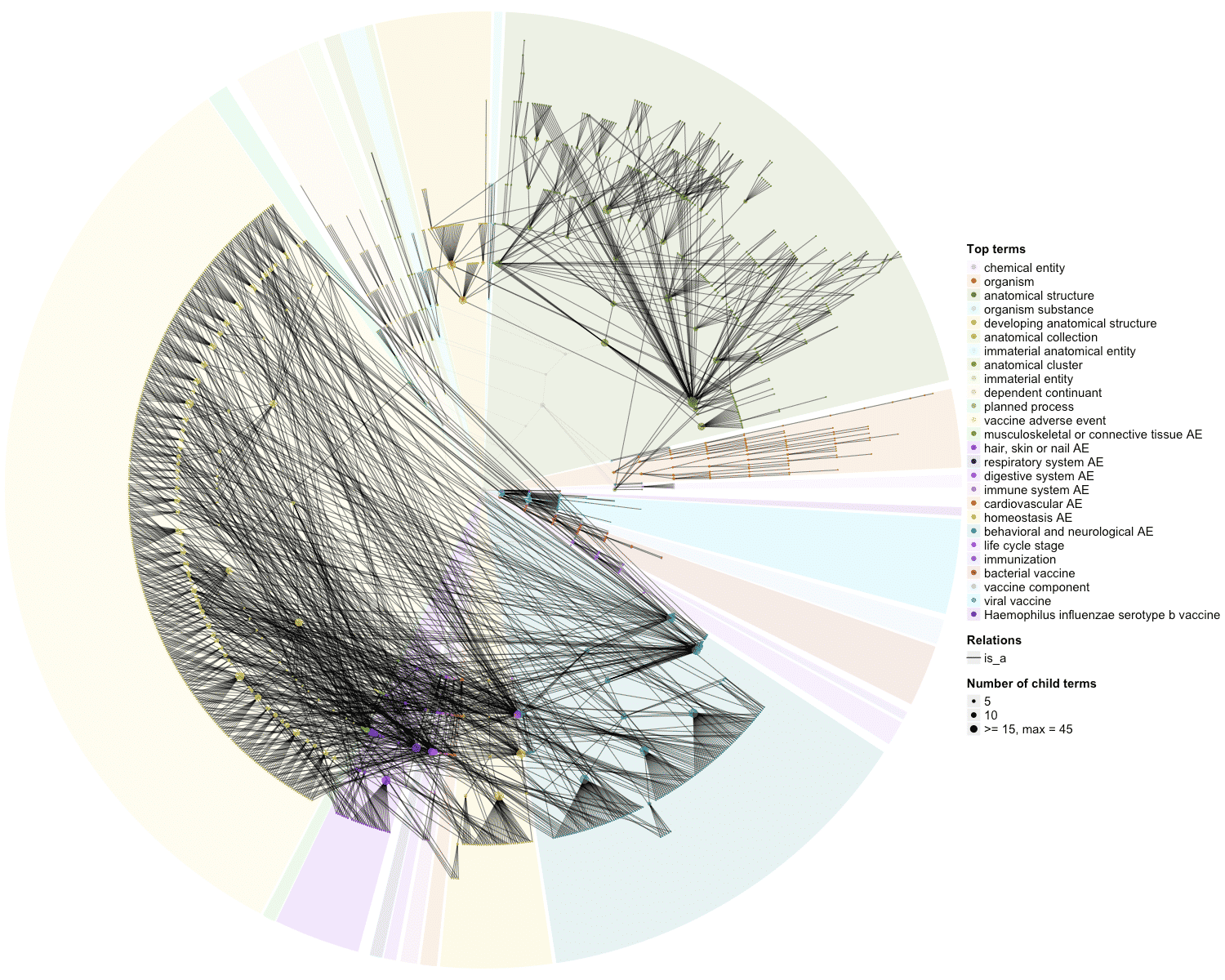

### OBOFoundry_pato_heatmap.png

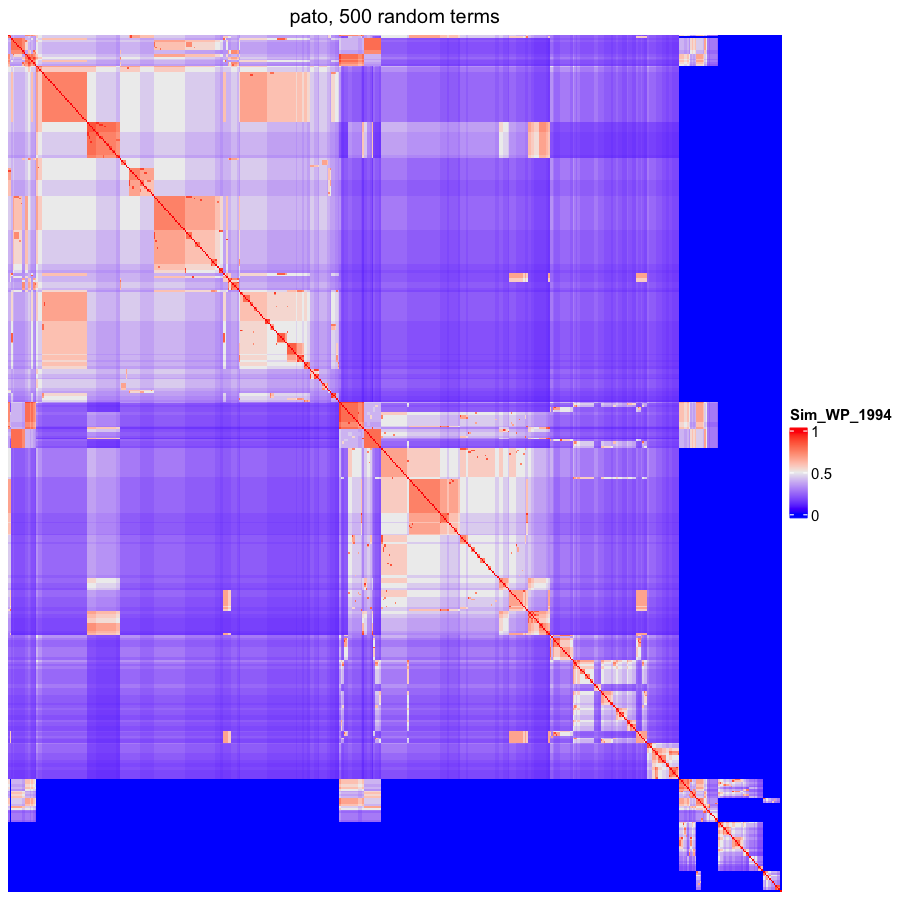

### OBOFoundry_phipo_runtime.png

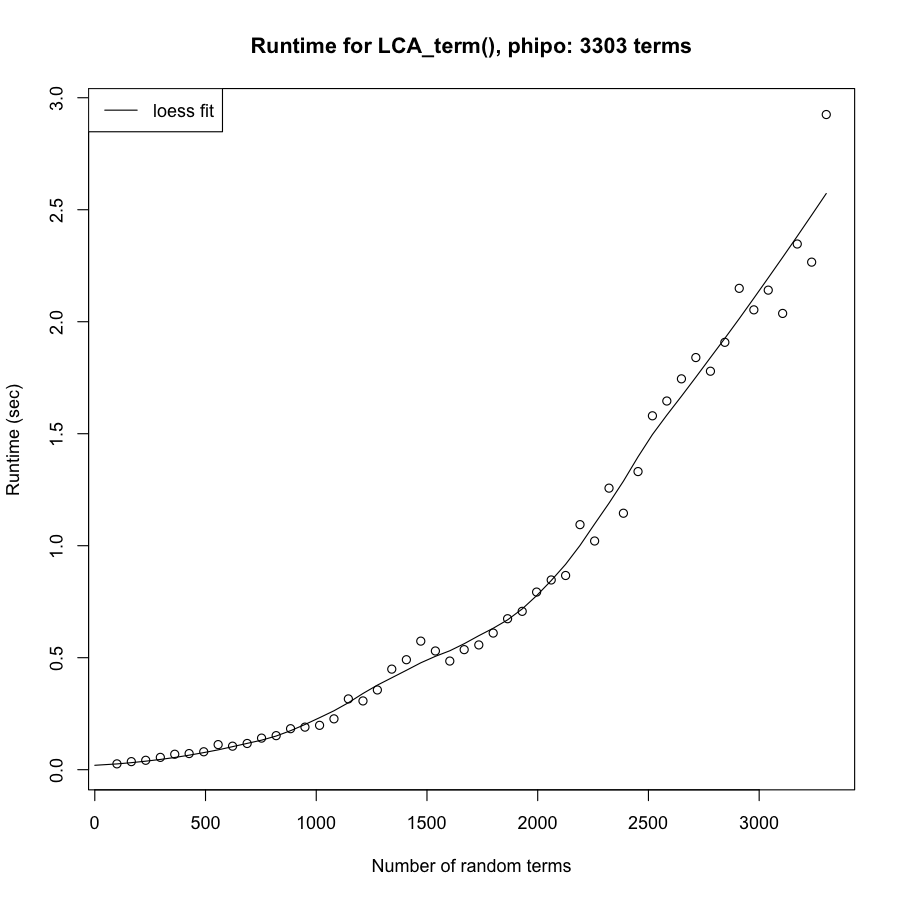

### OBOFoundry_po.png

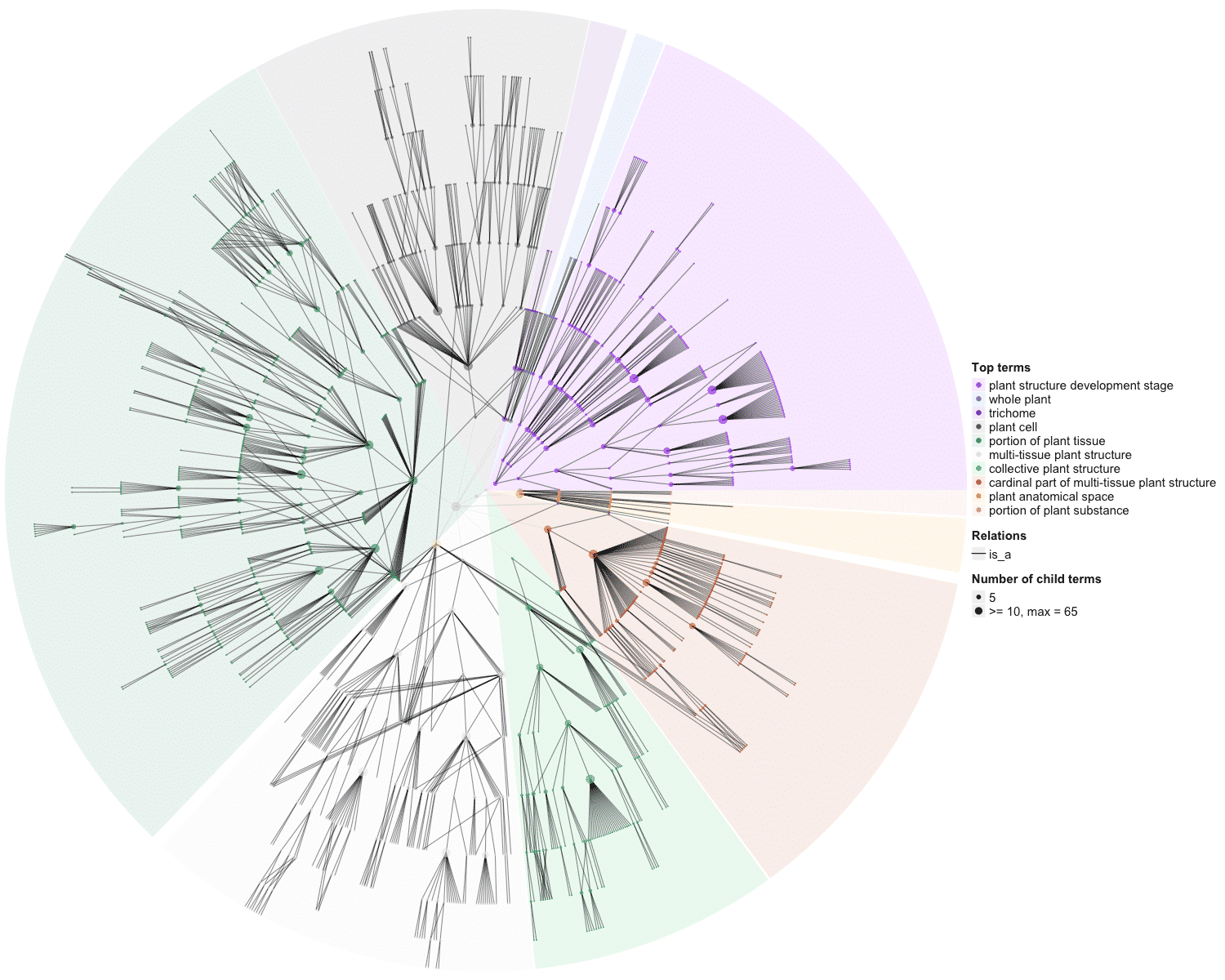

### OBOFoundry_ppo_heatmap.png

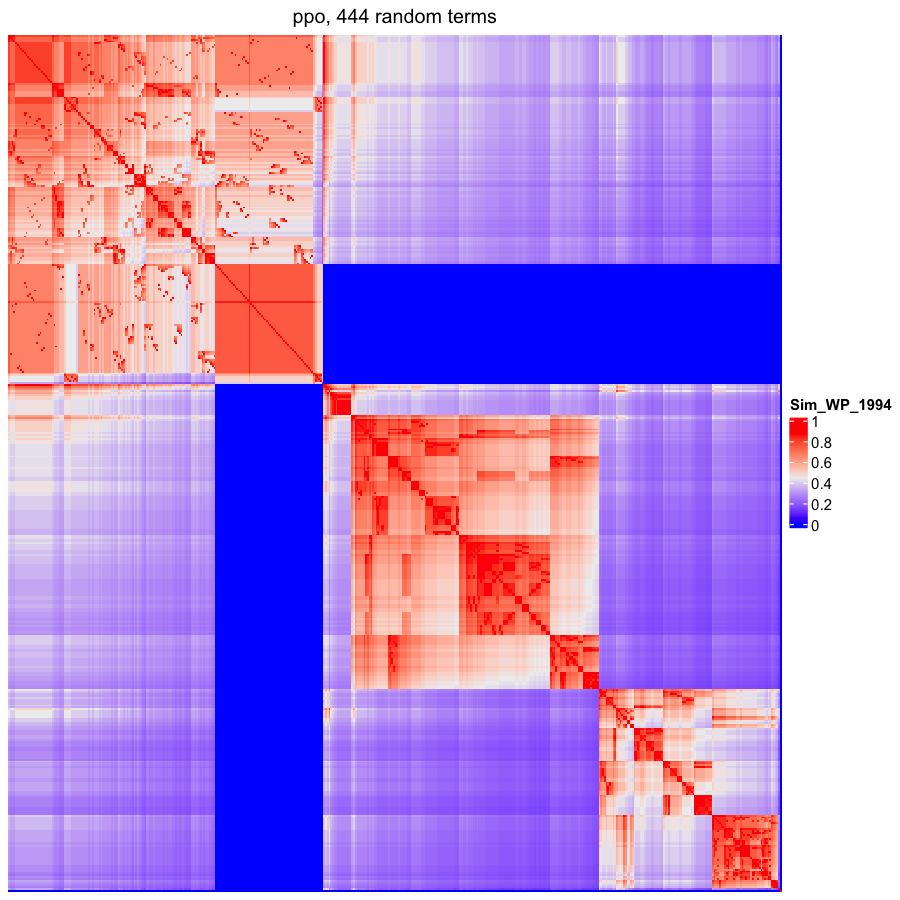

### OBOFoundry_pr_heatmap.png

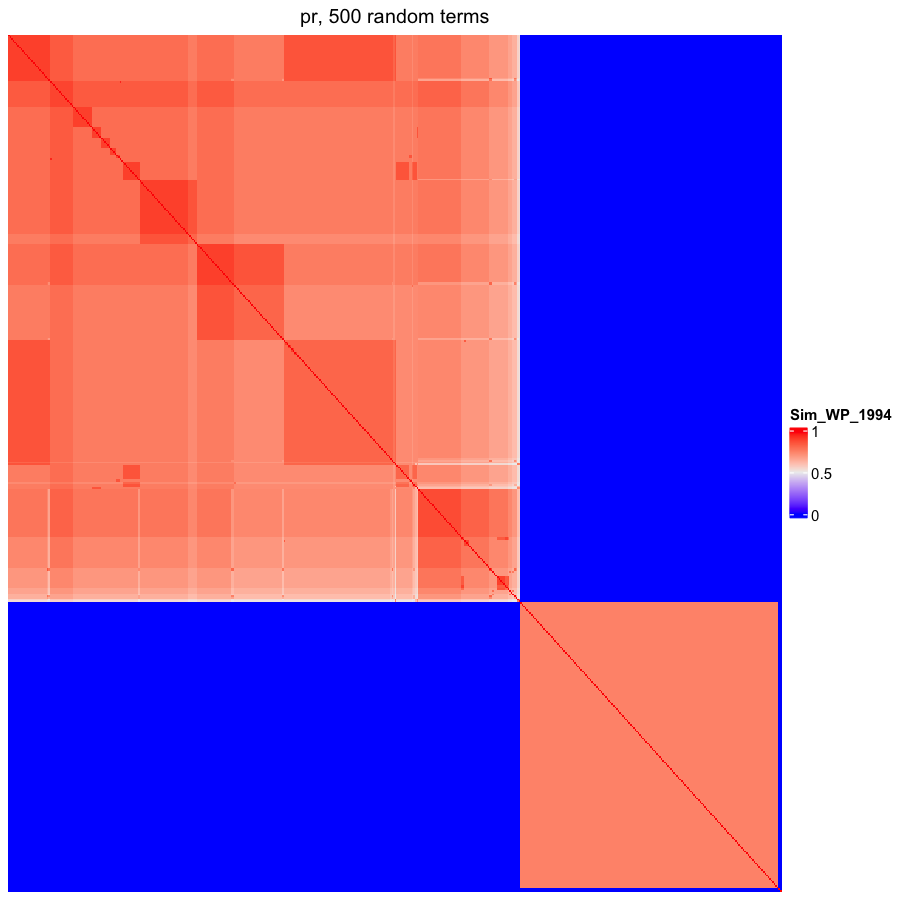

### OBOFoundry_pw_heatmap.png

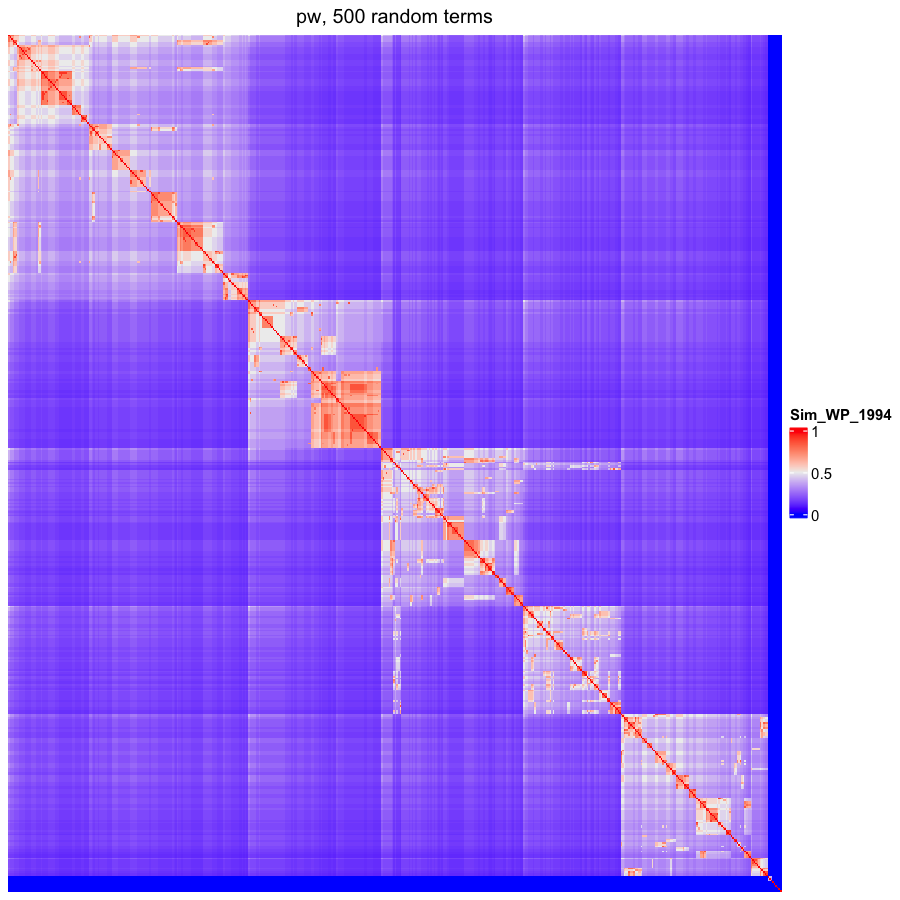

### OBOFoundry_rex_heatmap.png

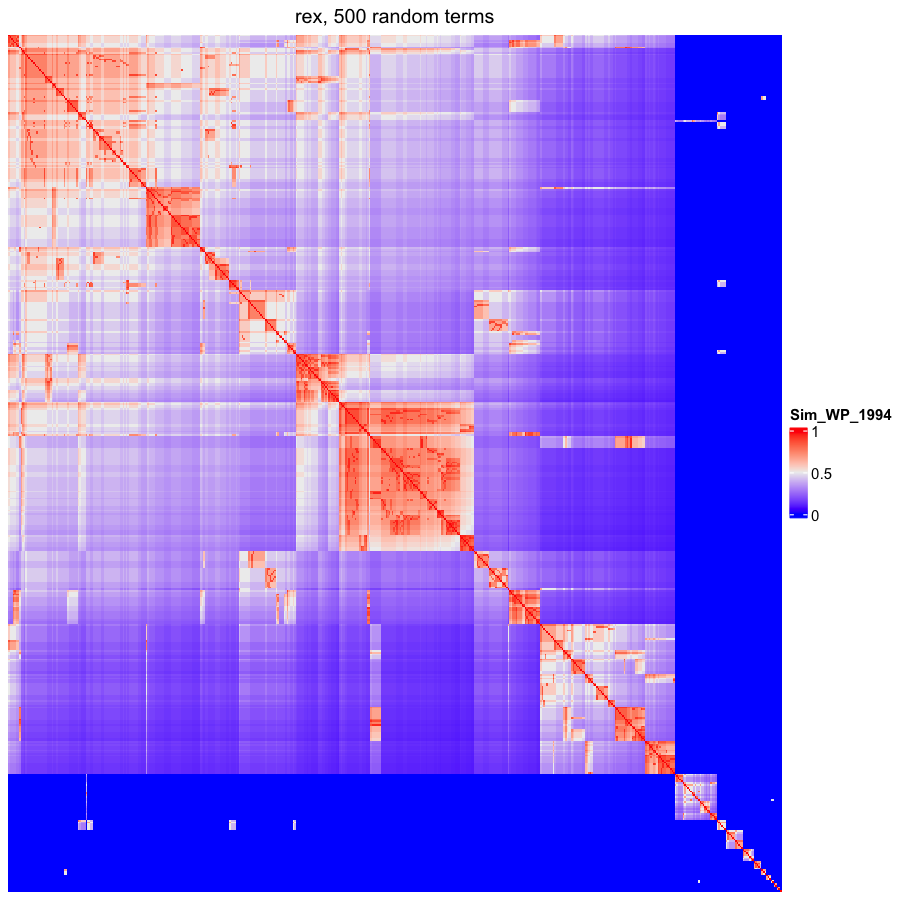

### OBOFoundry_rs.png

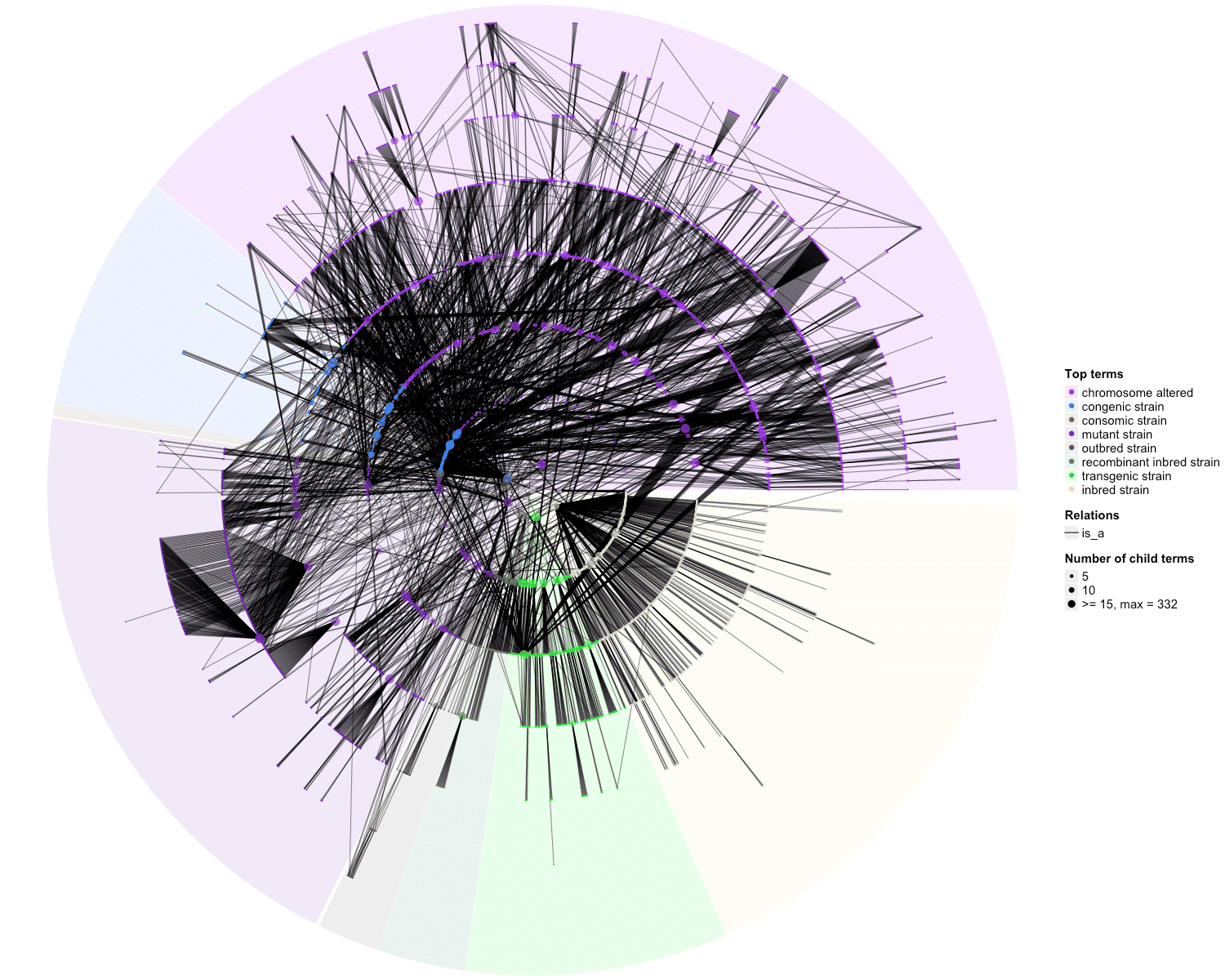

### OBOFoundry_scdo_heatmap.png

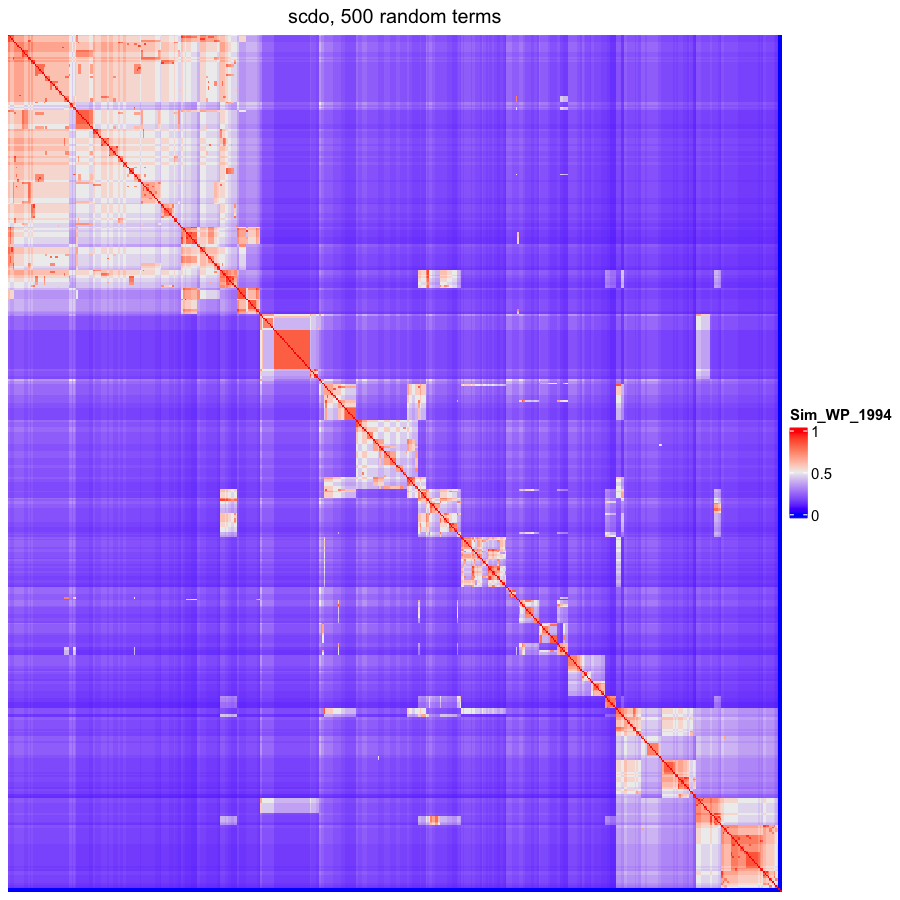

### OBOFoundry_so_heatmap.png

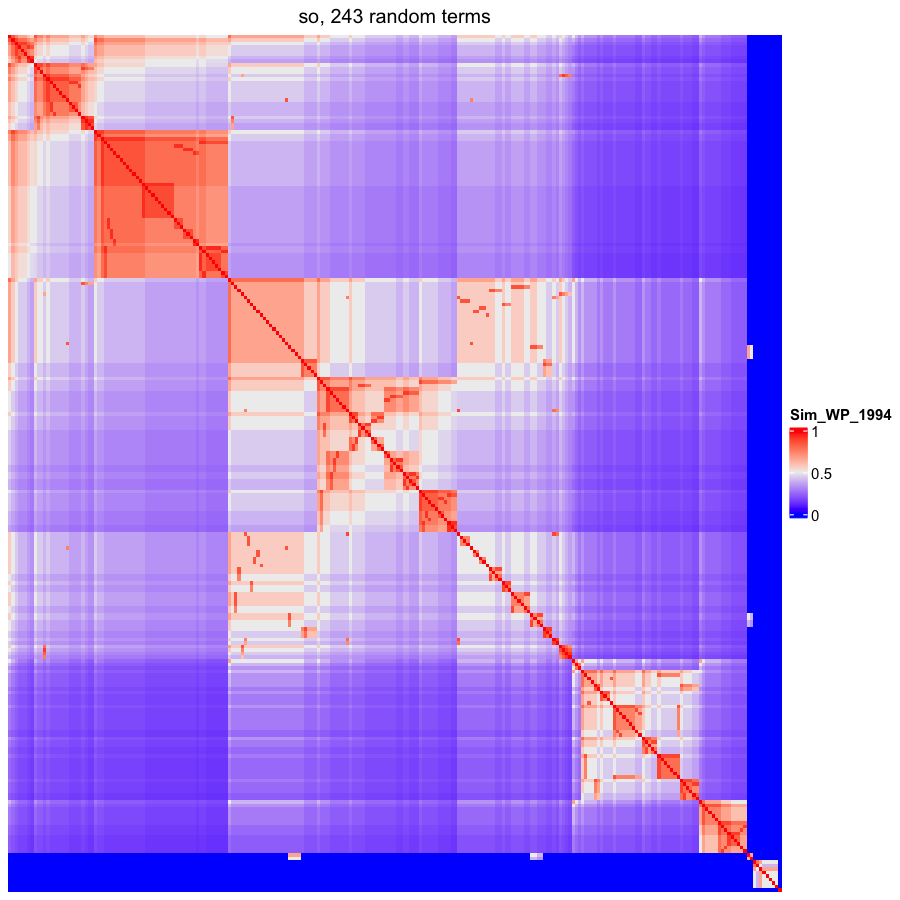

### OBOFoundry_spd.png

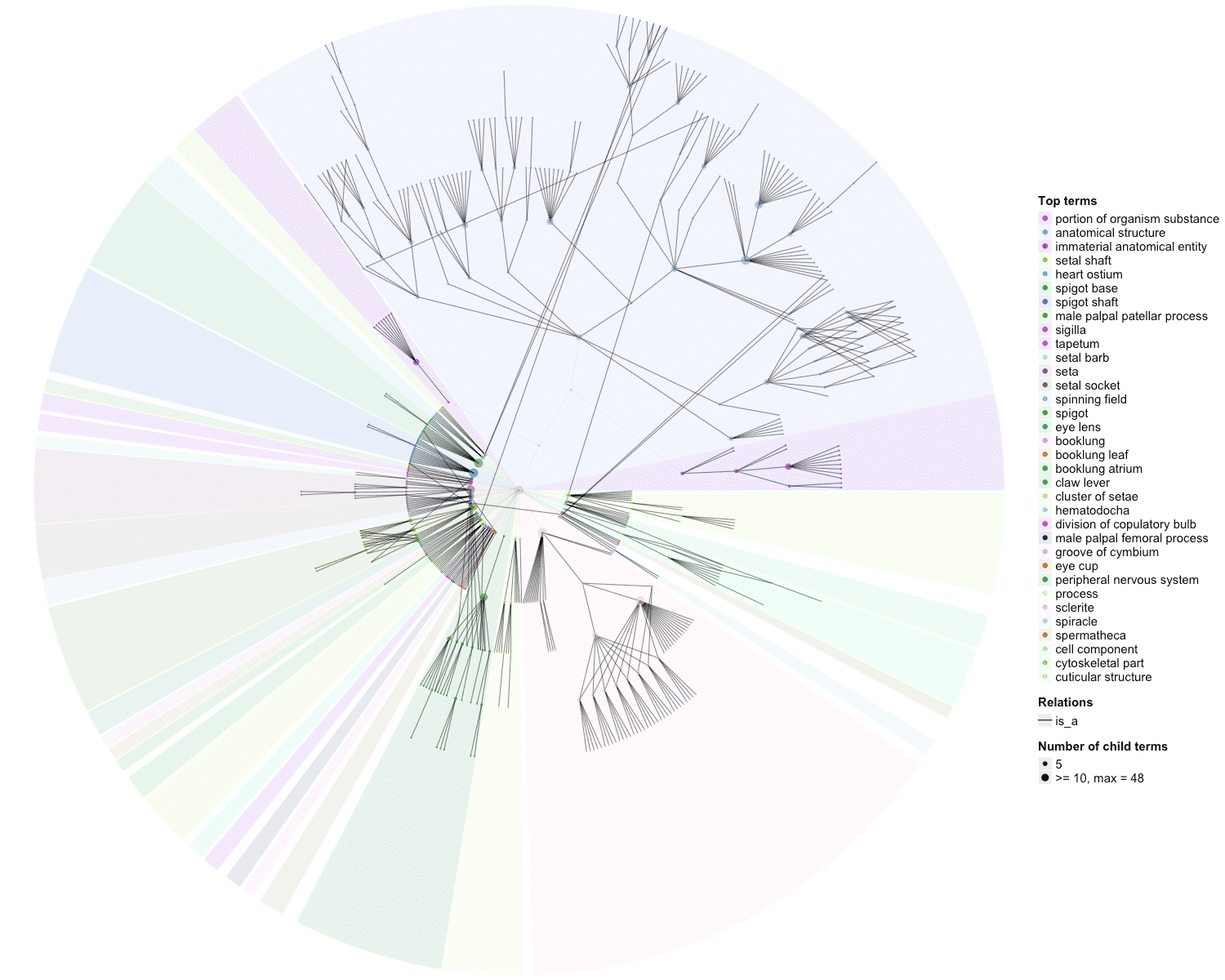

### OBOFoundry_symp_heatmap.png

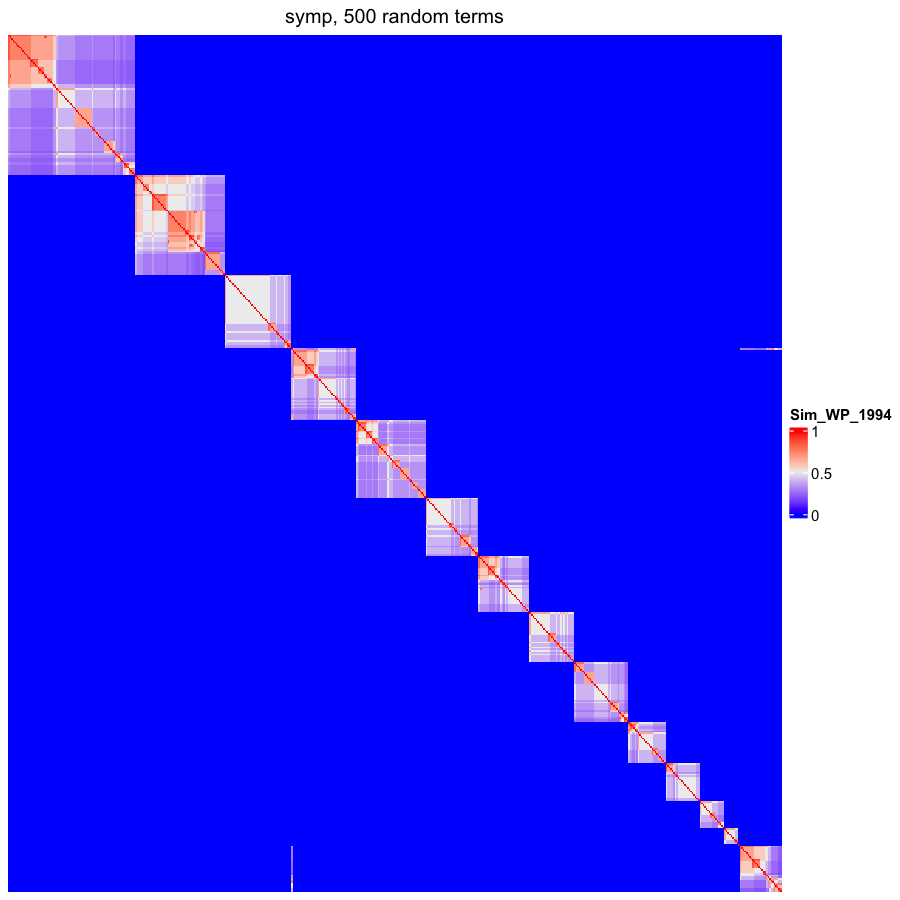

### OBOFoundry_taxrank_heatmap.png

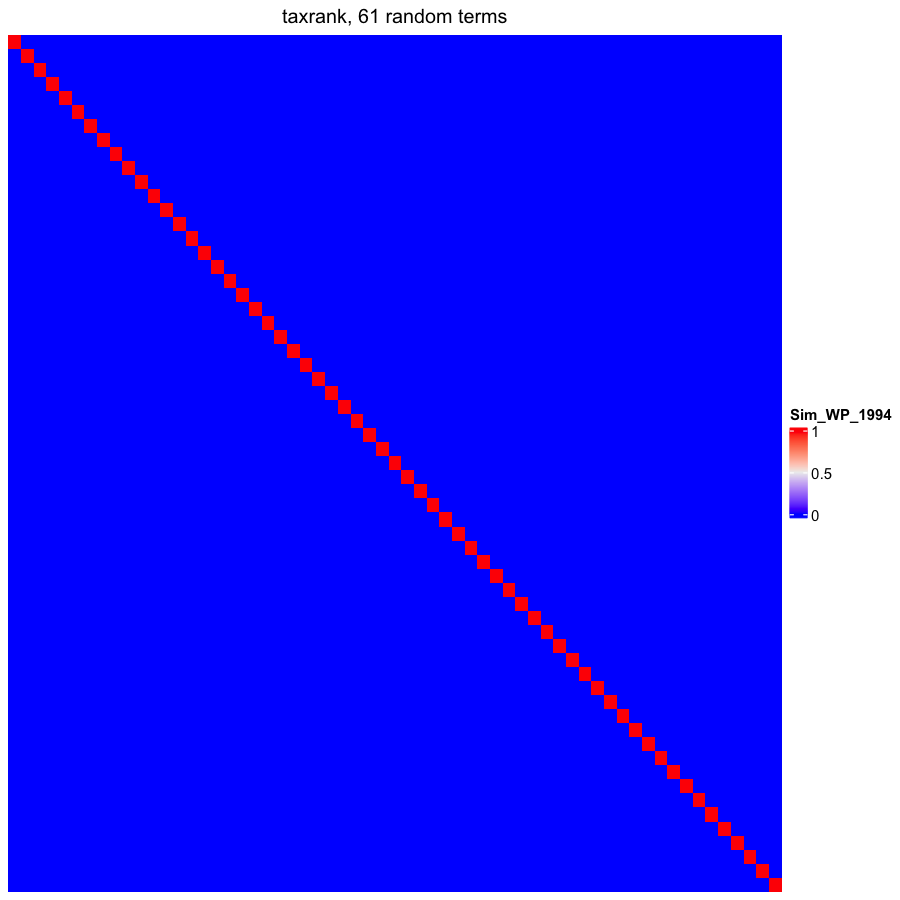

### OBOFoundry_to_heatmap.png

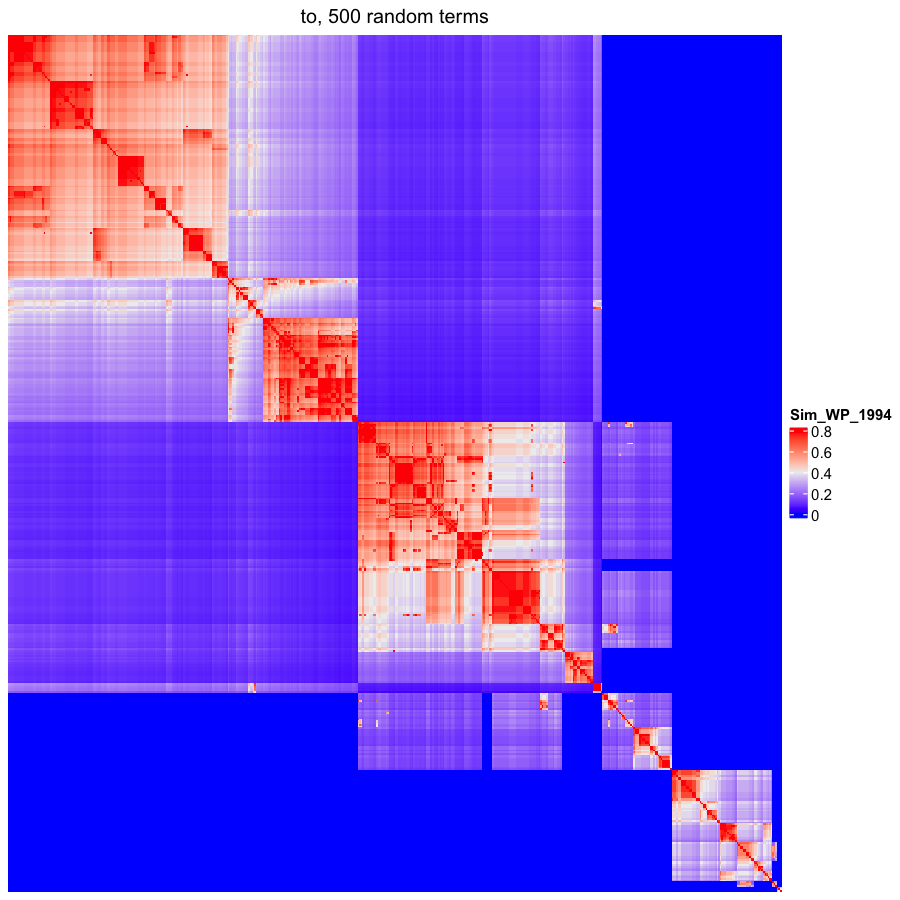

### OBOFoundry_trans.png

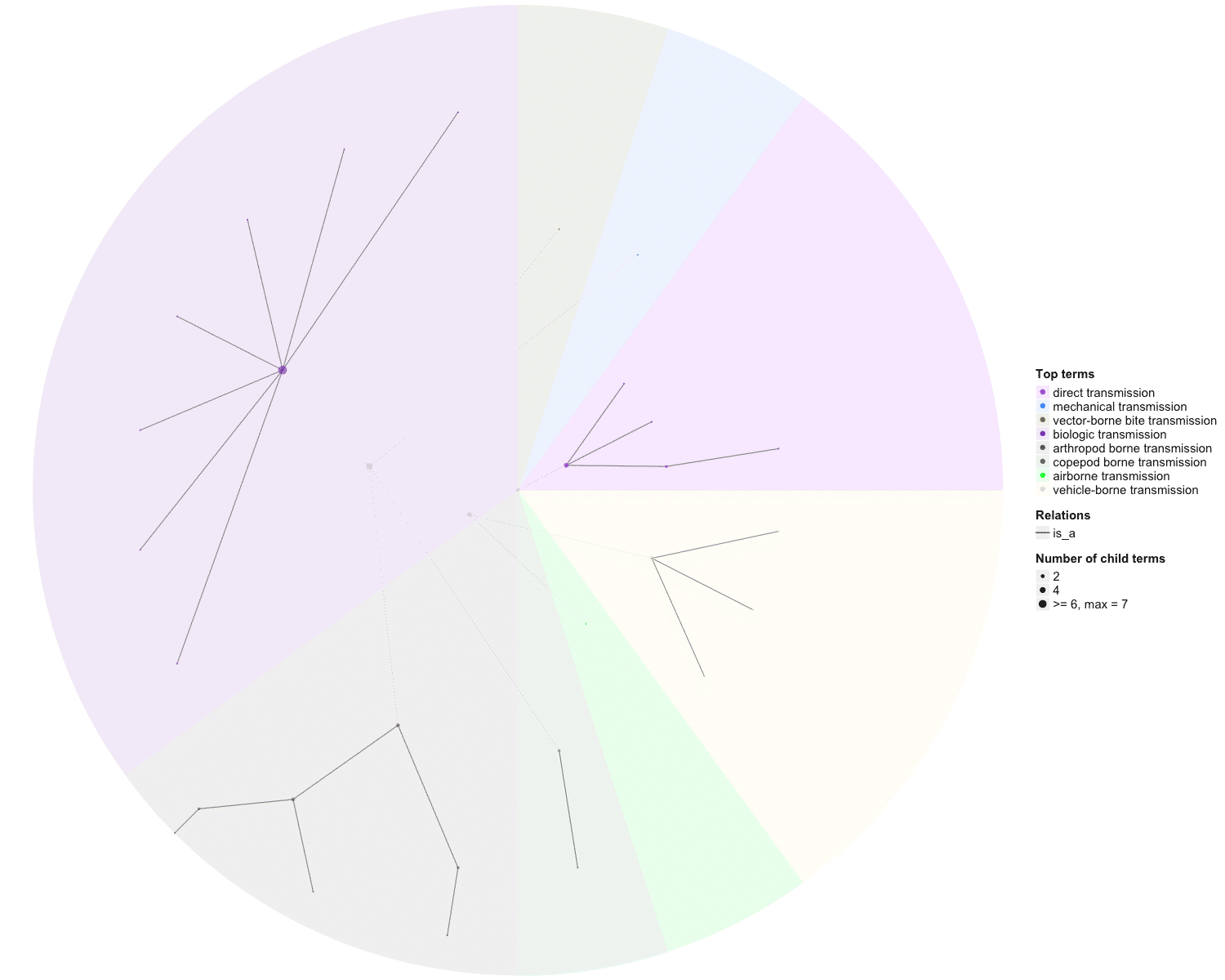

### OBOFoundry_tto_runtime.png

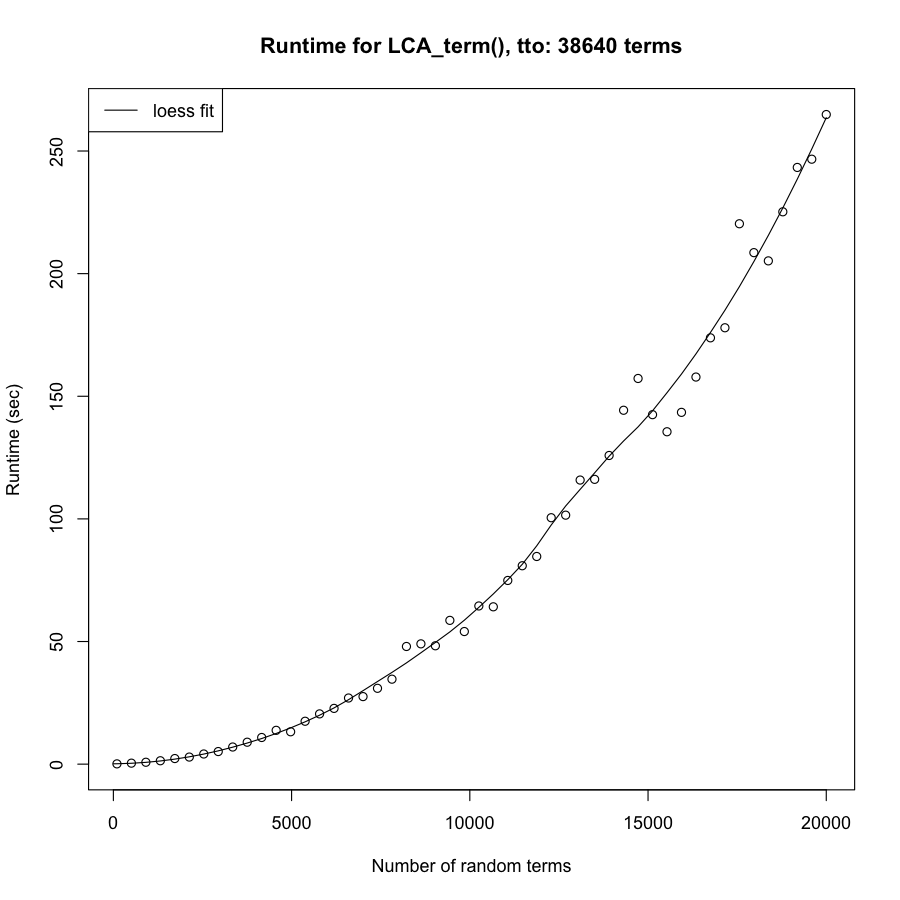

### OBOFoundry_txpo.png

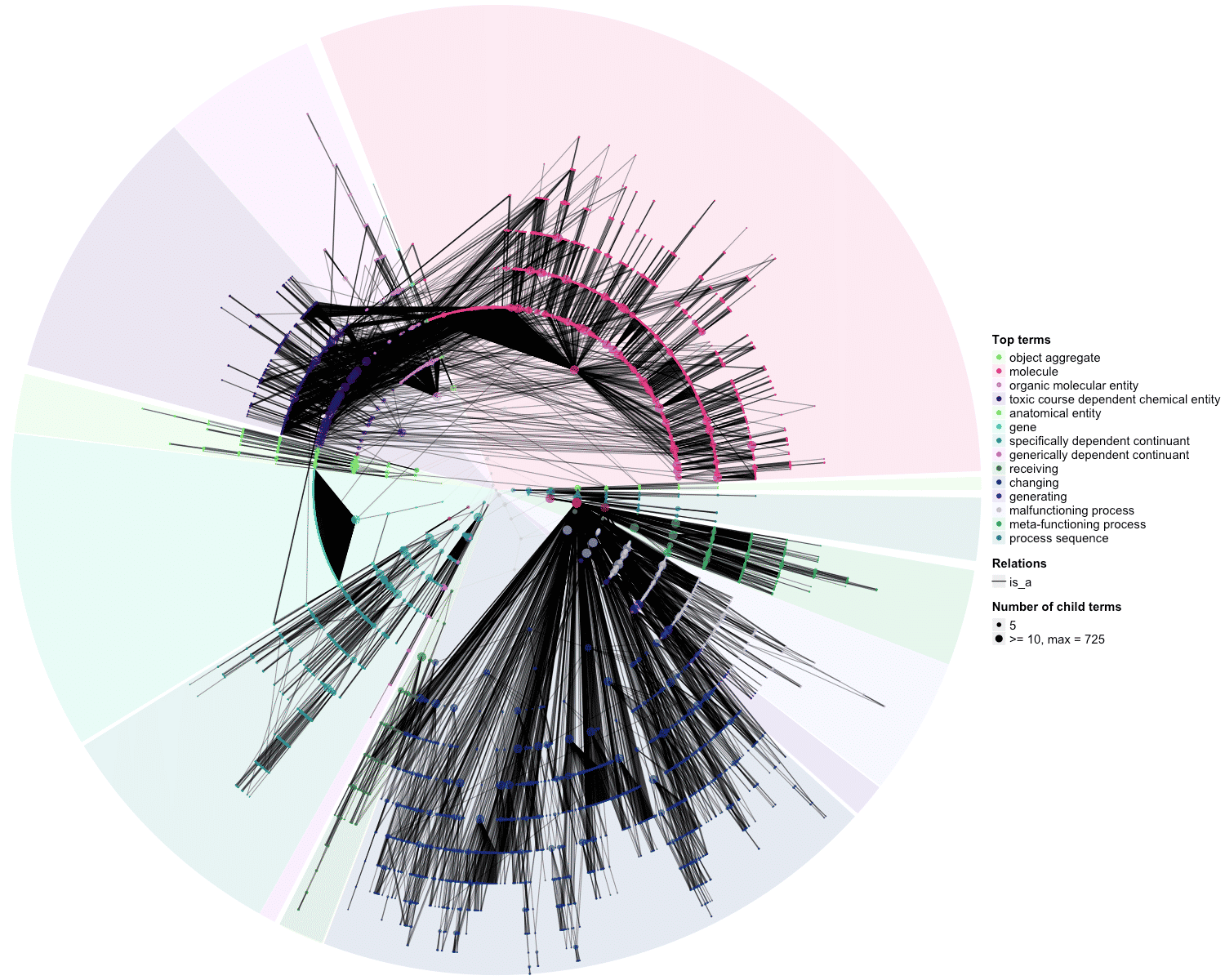

### OBOFoundry_txpo_heatmap.png

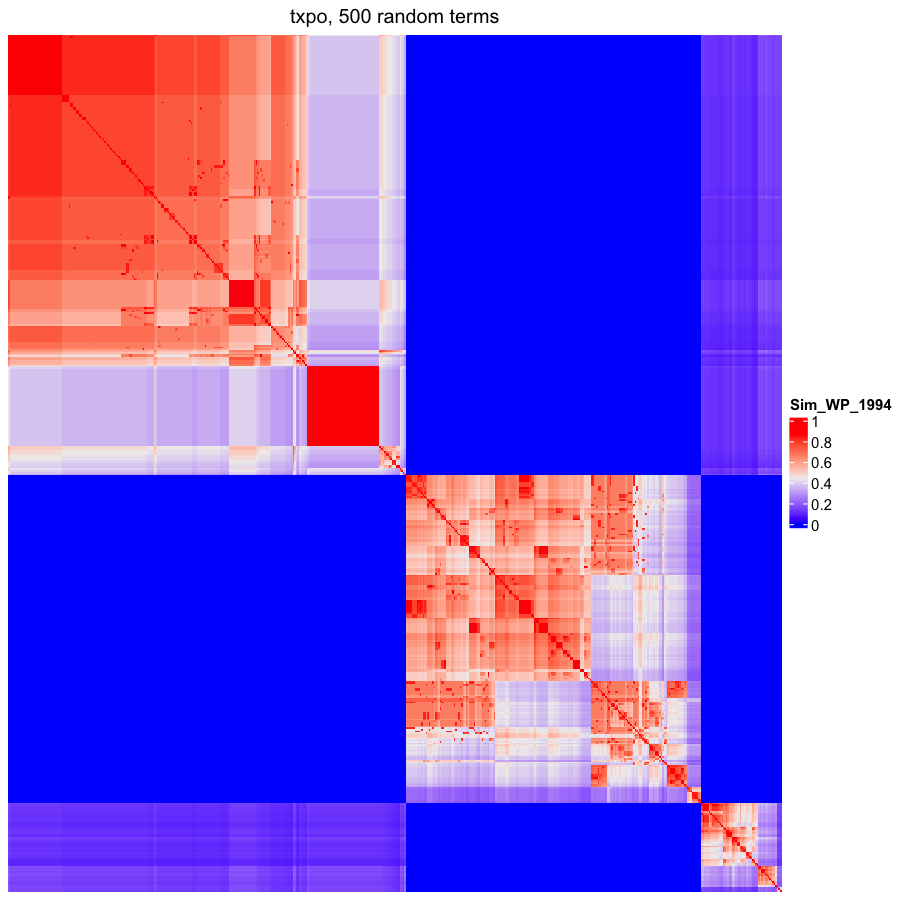

### OBOFoundry_upa.png

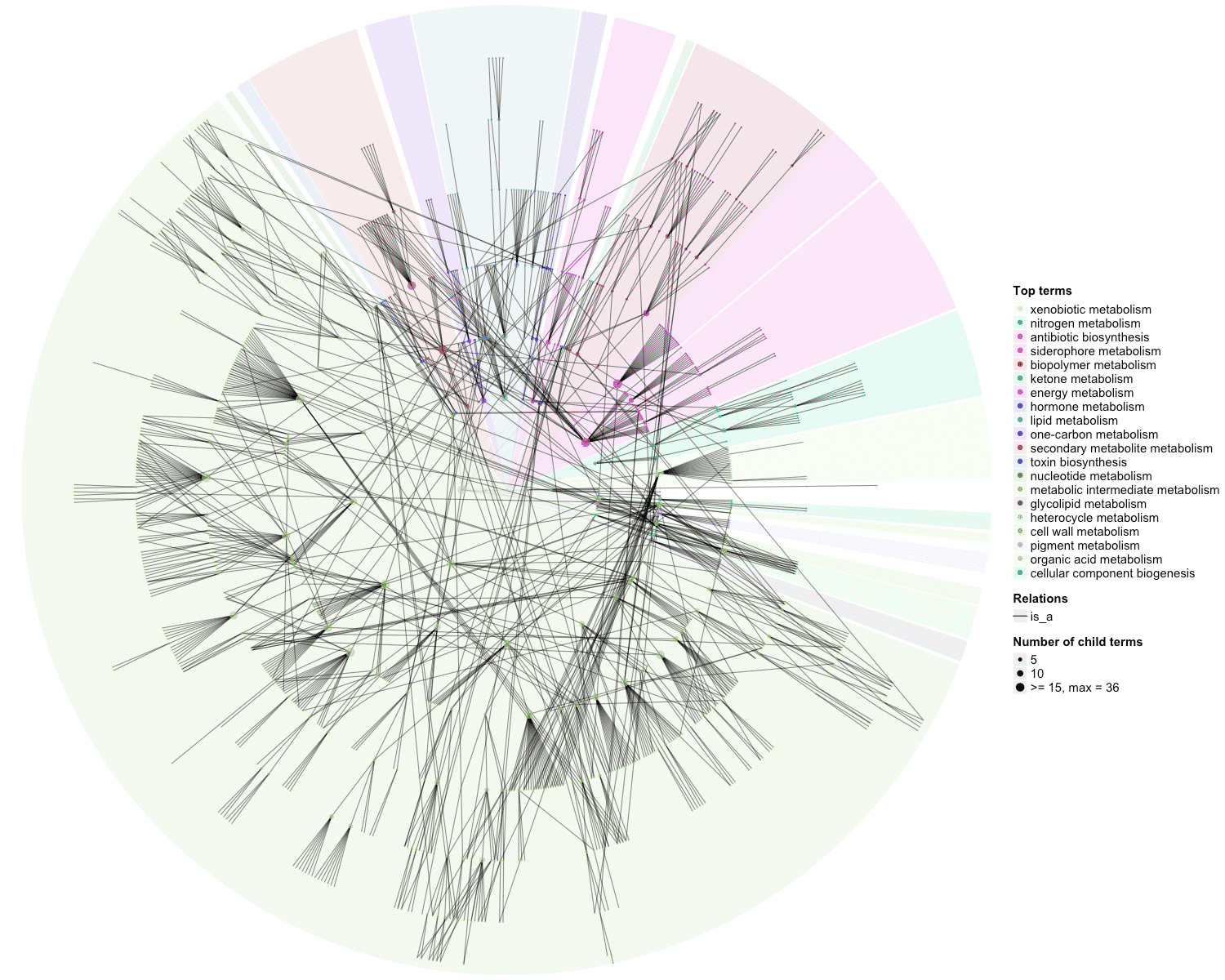

### OBOFoundry_upa_heatmap.png

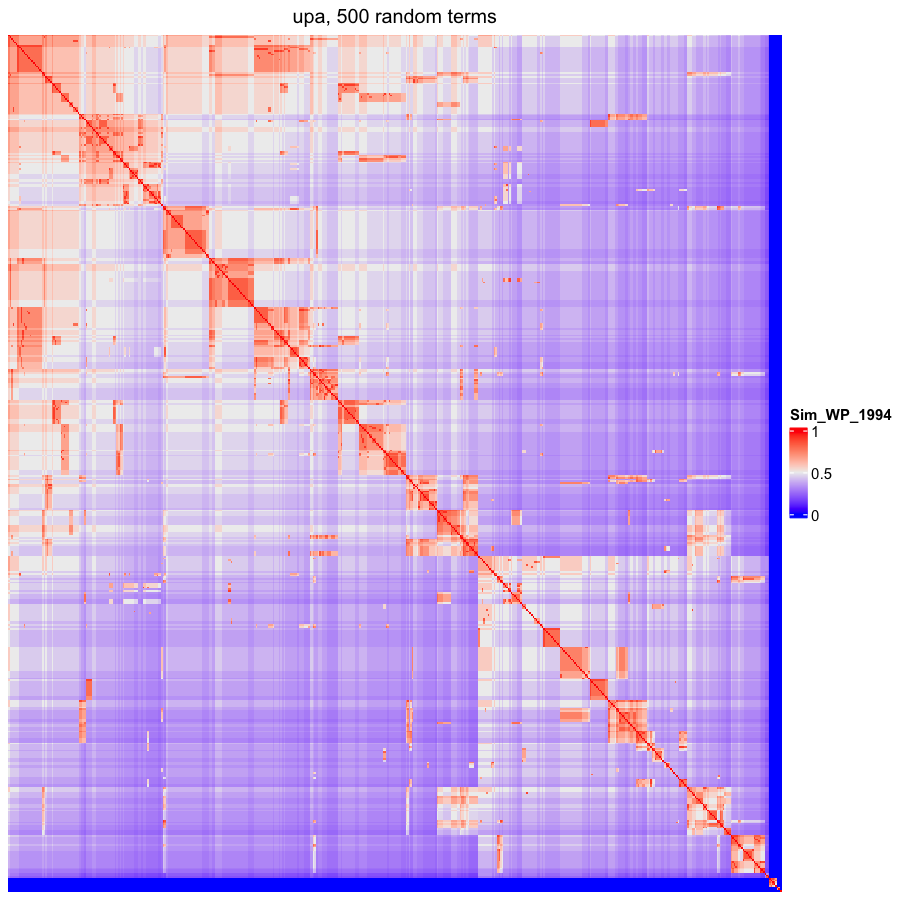

### OBOFoundry_upheno.png

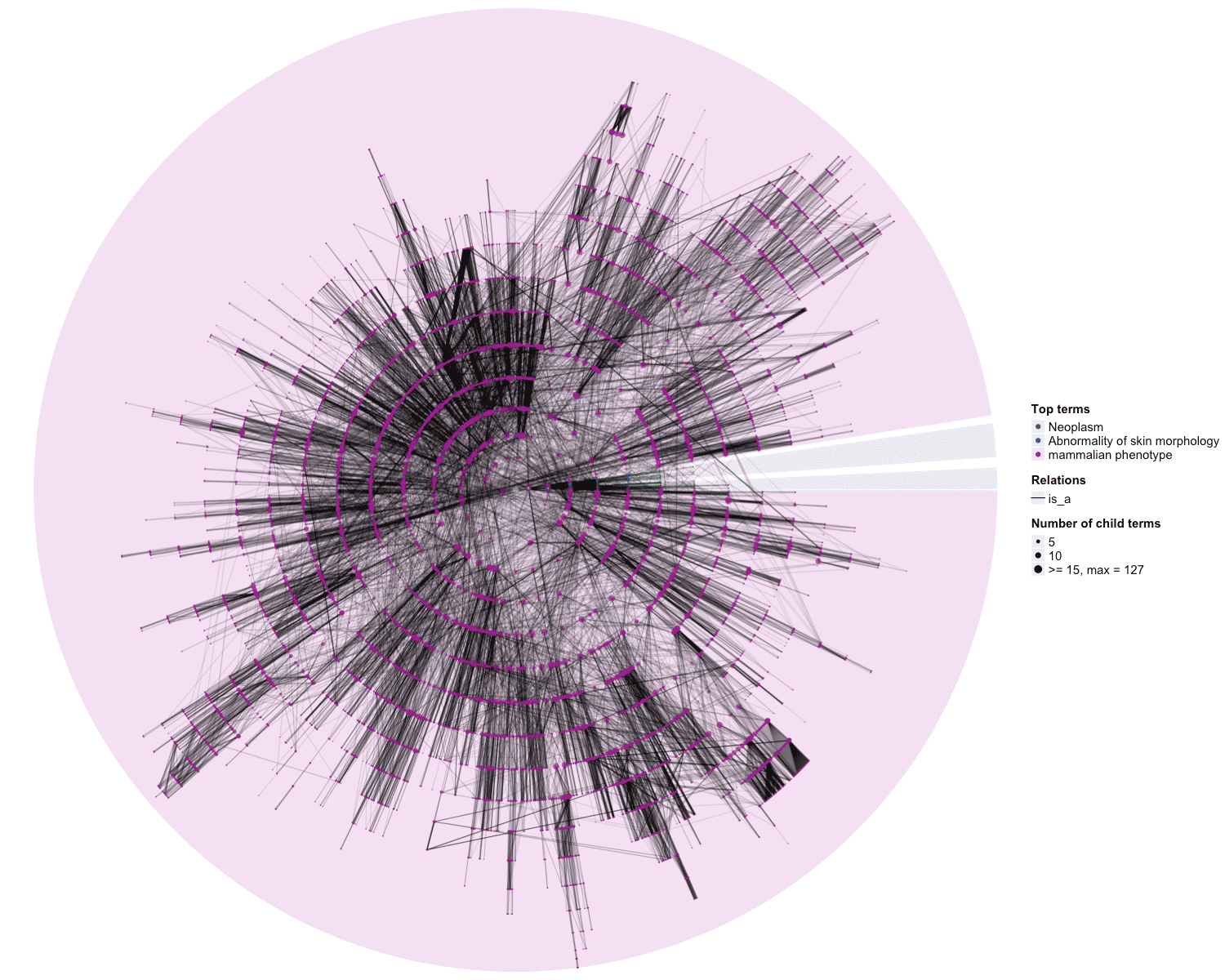

### OBOFoundry_vo_runtime.png

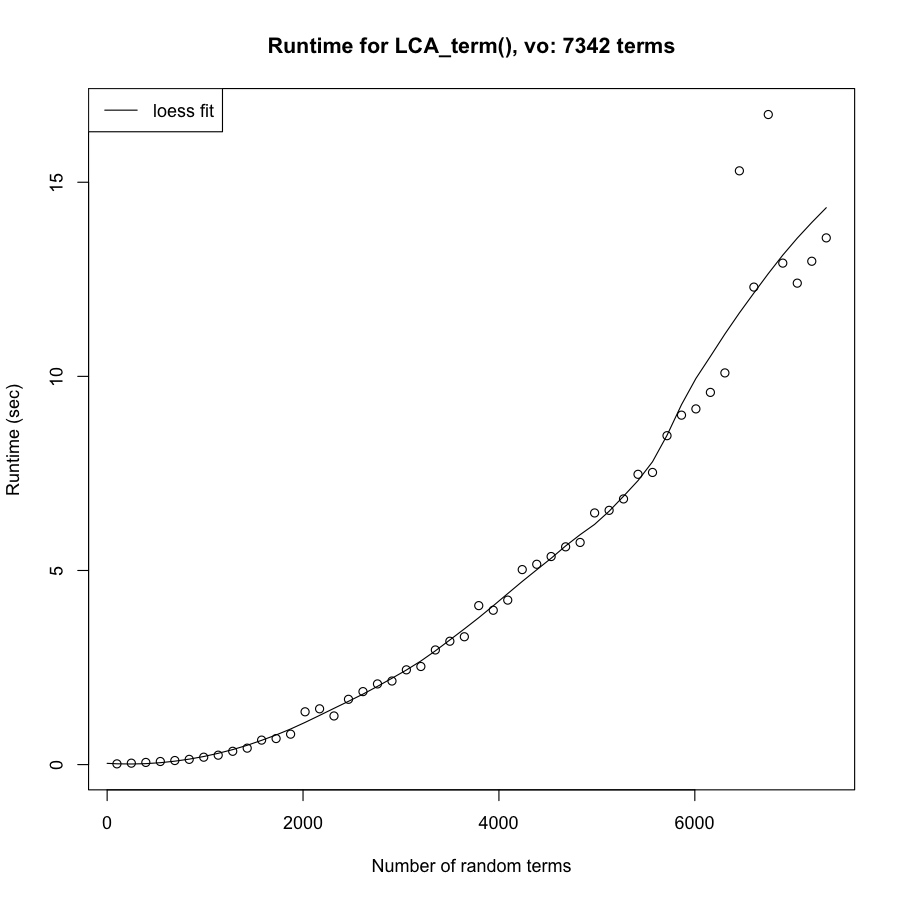

### OBOFoundry_vsao.png

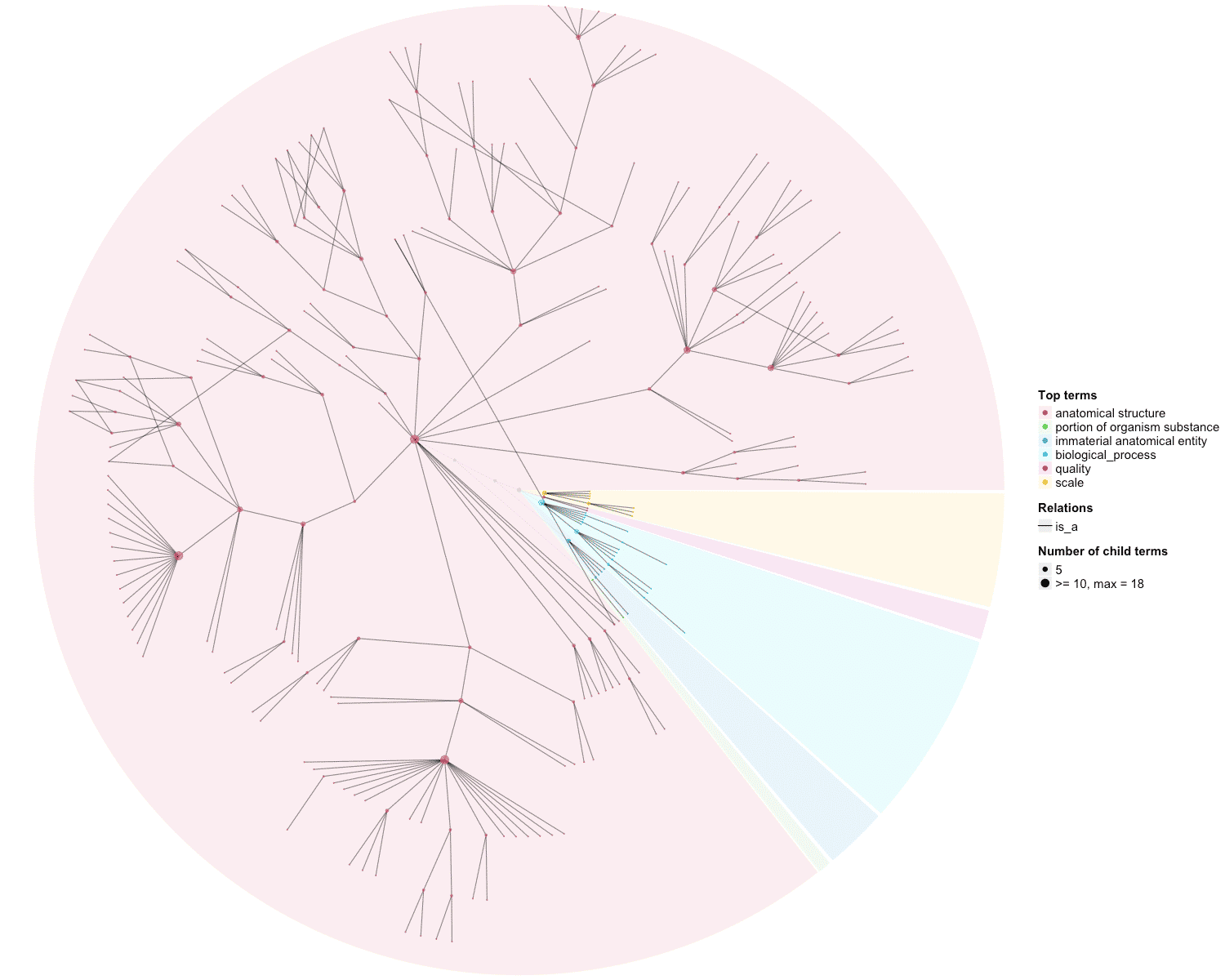

### OBOFoundry_vto.png

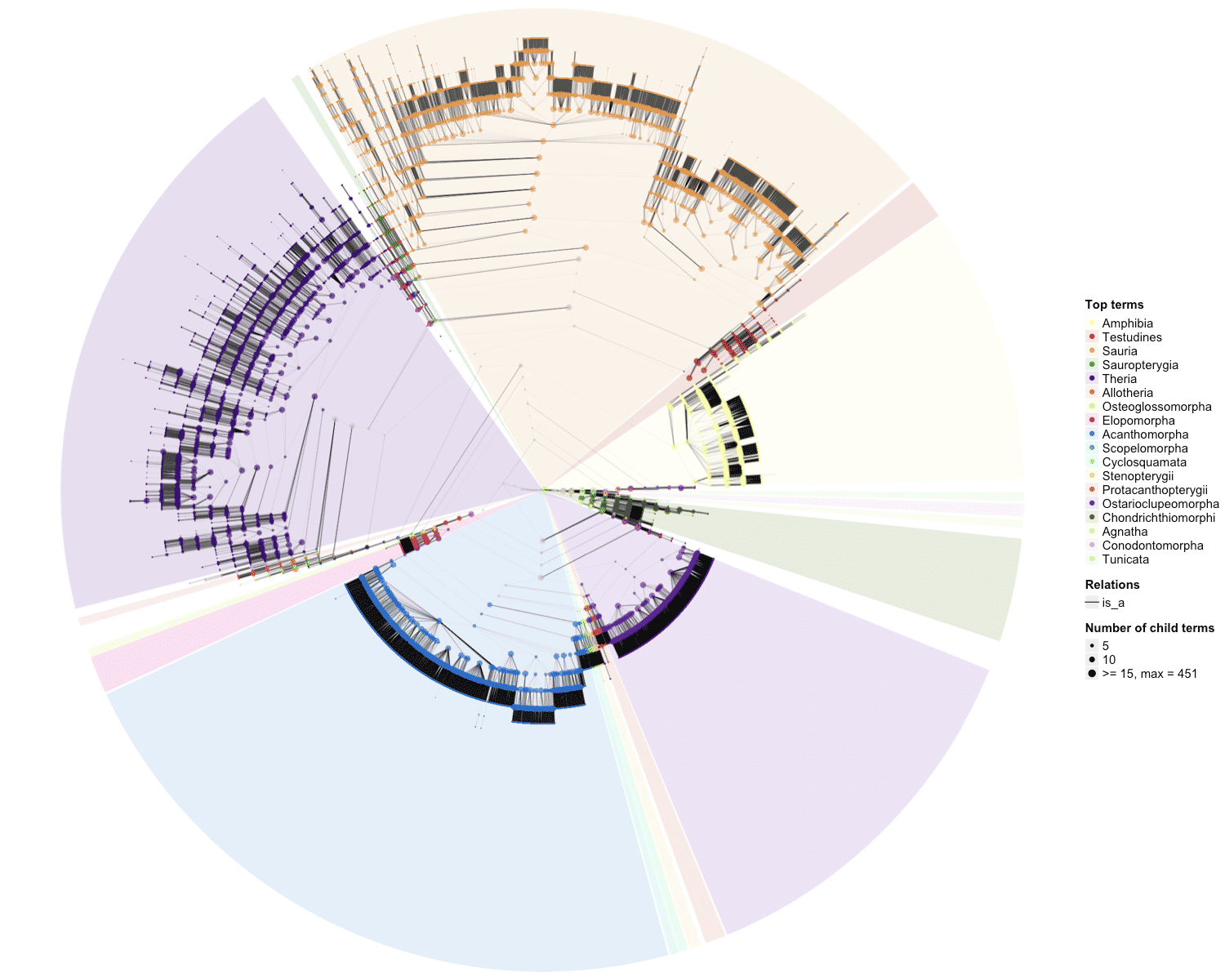

### OBOFoundry_vto_heatmap.png

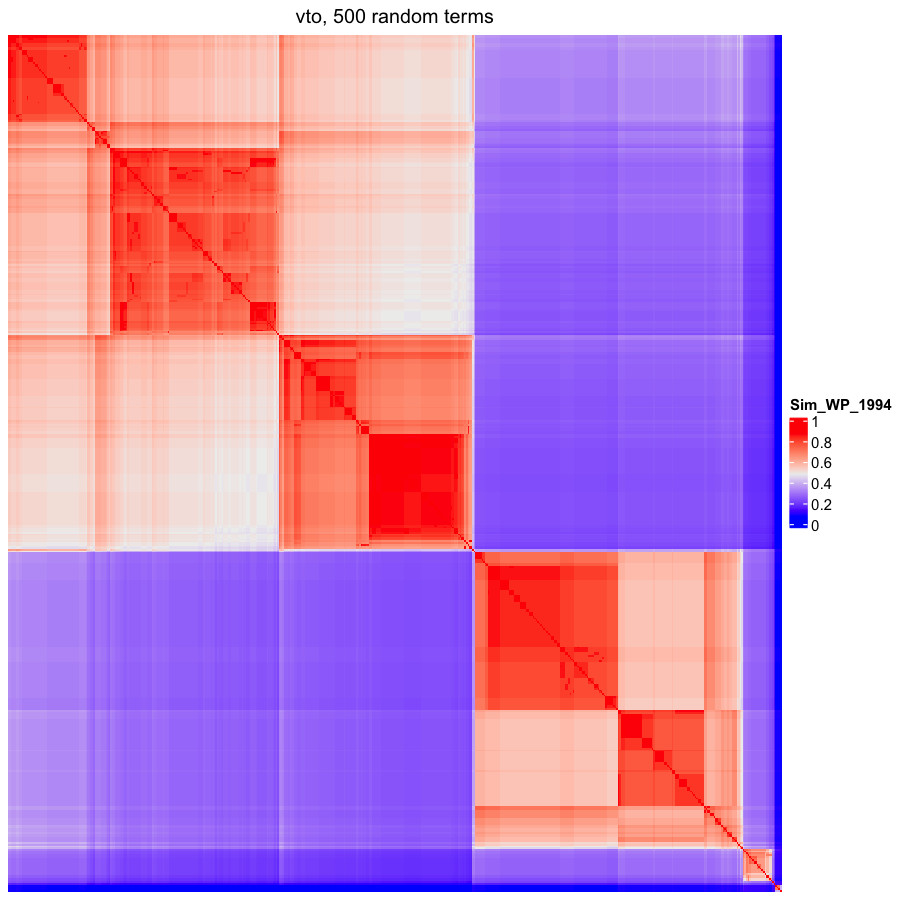

### OBOFoundry_wbbt_heatmap.png

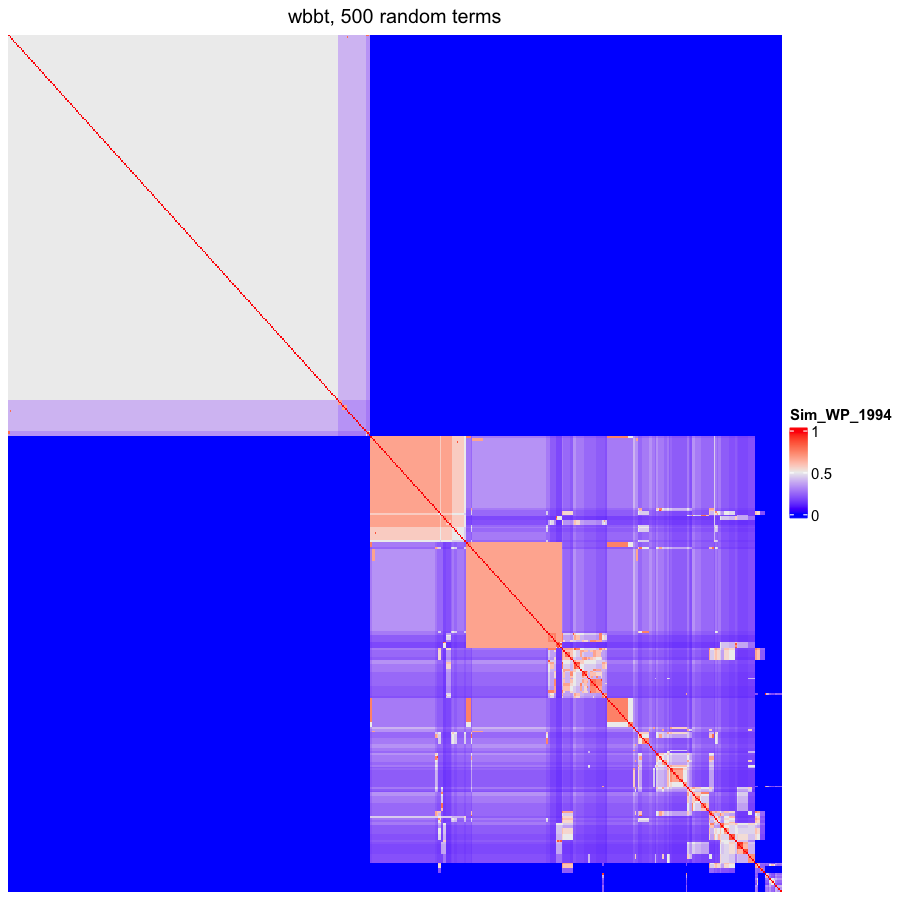

### OBOFoundry_xao_heatmap.png

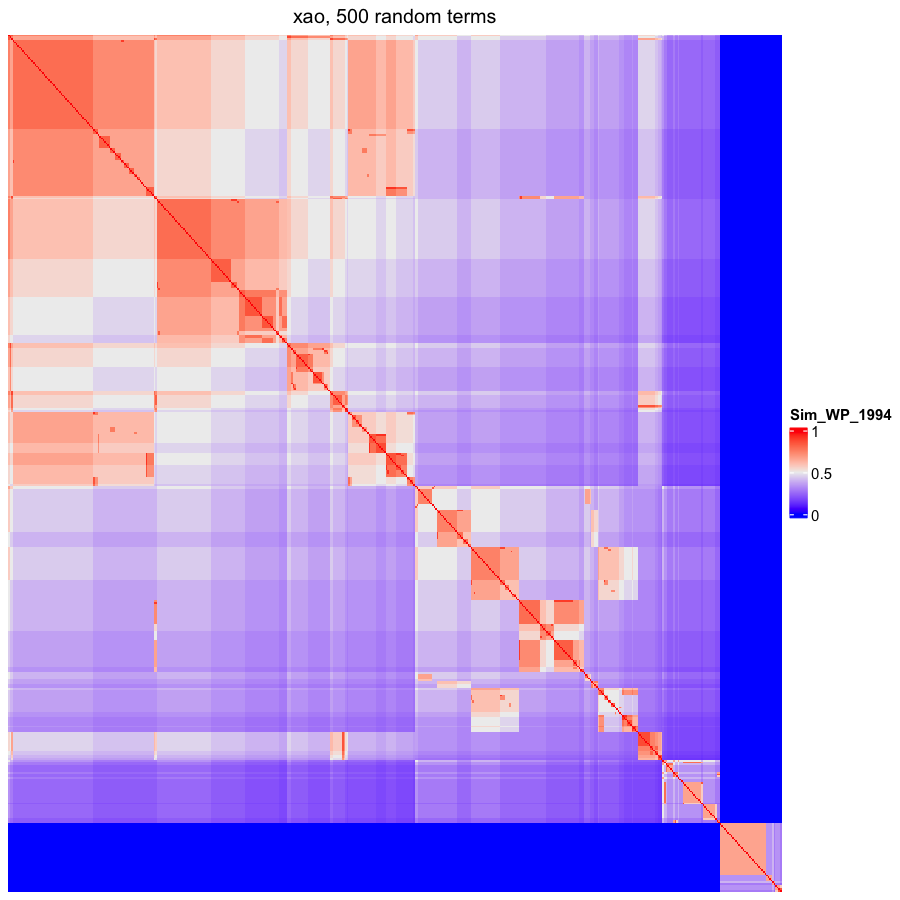

### OBOFoundry_xco.png

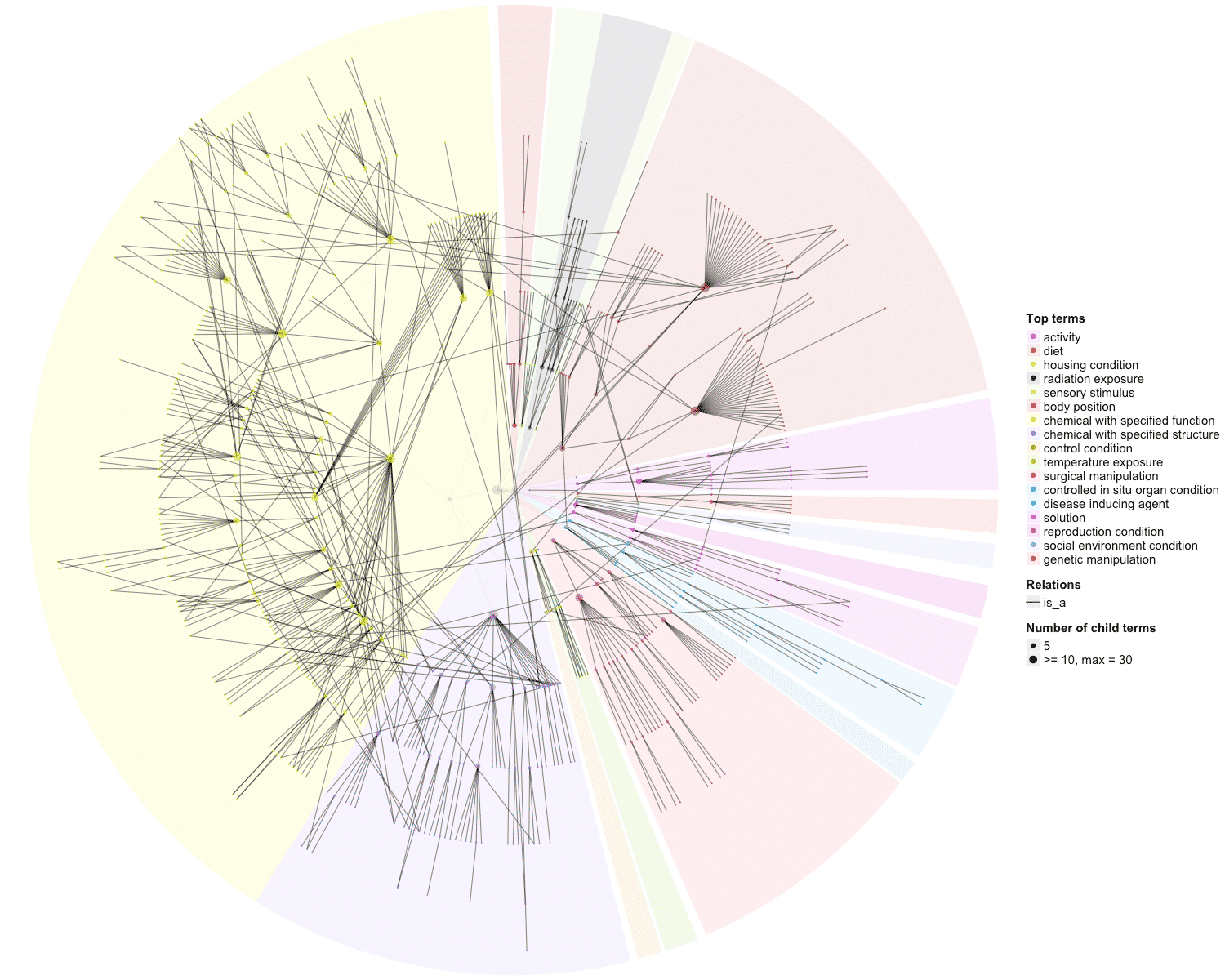
